# Discovering the N-terminal Methylome by Repurposing of Proteomic Datasets

**DOI:** 10.1101/2021.04.14.439552

**Authors:** Panyue Chen, Tiago Jose Paschoal Sobreira, Mark C. Hall, Tony R. Hazbun

## Abstract

Protein α-N-methylation is an underexplored post-translational modification involving the covalent addition of methyl groups to the free α-amino group at protein N-termini. To systematically explore the extent of α-N-terminal methylation in yeast and humans, we reanalyzed publicly accessible proteomic datasets to identify N-terminal peptides contributing to the α-N-terminal methylome. This repurposing approach found evidence of α-N-methylation of established and novel protein substrates with canonical N-terminal motifs of established α-N-terminal methyltransferases, including human NTMT1/2 and yeast Tae1. NTMT1/2 are implicated in cancer and aging processes but have unclear and context-dependent roles. Moreover, α-N-methylation of non-canonical sequences was surprisingly prevalent, suggesting unappreciated and cryptic methylation events. Analysis of the amino acid frequencies of α-N-methylated peptides revealed a [S]_1_-[S/A/Q]_2_ pattern in yeast and [A/N/G]_1_-[A/S/V]_2_-[A/G]_3_ in humans, which differs from the canonical motif. We delineated the distribution of the two types of prevalent N-terminal modifications, acetylation, and methylation, on amino acids at the 1^st^ position. We tested three potentially methylated proteins and confirmed the α-N-terminal methylation of Hsp31 by additional proteomic analysis and immunoblotting. The other two proteins, Vma1 and Ssa3, were found to be predominantly acetylated, indicating proteomic searching for α-N-terminal methylation requires careful consideration of mass spectra. This study demonstrates the feasibility of reprocessing proteomic data for global α-N-terminal methylome investigations.

The raw MS data that supports the findings of this study were deposited with PRIDE identifier: PXD022833.

Graphical Abstract (For TOC only).

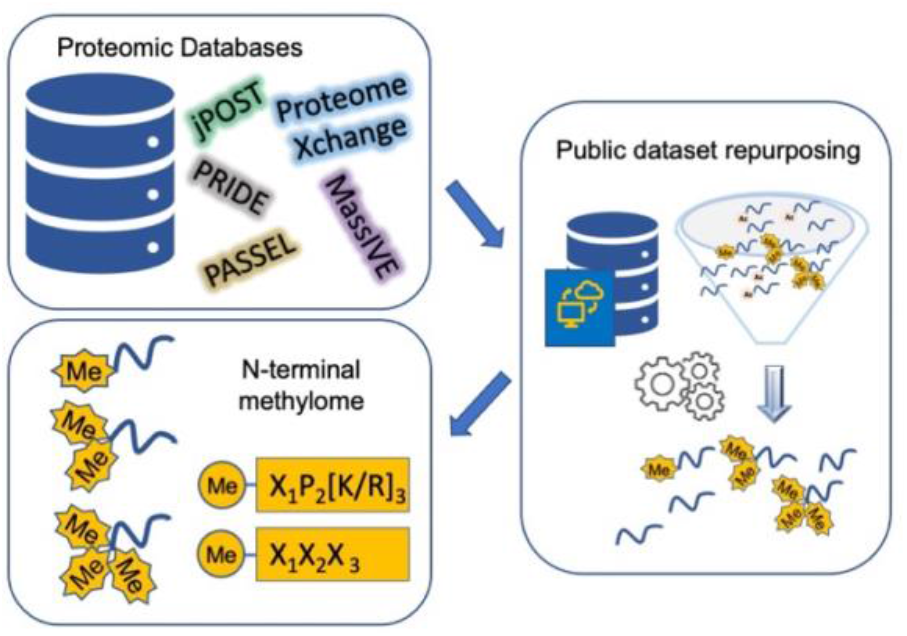

## Introduction

Protein N-terminal methylation is a novel post-translational modification (PTM) that was first reported decades ago in prokaryotes, whereas the eukaryotic methyltransferases were identified more recently. This PTM involves adding up to three methyl groups on the free α-amino group of the protein N-terminus. The fully methylated state is trimethylation except for proline dimethylation and can result in a pH insensitive positive charge on the protein N-termini.^1^ The Tae1/NTMT1/NTMT2 α-N-terminal methyltransferases are highly conserved between yeast and humans, and the enzymes recognize substrates with similar N-terminal sequence motifs, X_1_-P_2_-[K/R]_3_ (X position could be A, S, G or P), hereinafter referred to as the canonical N-terminal motif.^2,3^ In the motif, the initiating methionine (iMet) is commonly removed during protein maturation, leaving the α-amino group on the X amino acid exposed, which is subsequently targeted for methylation.

Most of the canonical N-terminal motif-containing proteins have not been validated as methylated in yeast and humans. The major N-terminal methyltransferase in yeast that recognizes the X_1_-P_2_-[K/R]_3_ motifs, Tae1 (<u>t</u>ranslational <u>a</u>ssociated <u>e</u>lement 1), was previously shown to modify two ribosomal subunits, Rpl25a/b and Rps12ab, and one proteasome subunit Rpt1.^4,5^ These three substrates are involved in protein synthesis and degradation, but the modification’s exact role is unclear. The prevailing view is that N-terminal methylation affects polysome assembly and thus contributes to protein synthesis efficiency and fidelity.^6^ Notably, a total of 45 proteins in the yeast proteome contain the canonical motif, and Tae1 is implicated in multiple biological pathways (Table S1). In humans, there are two primary enzymes responsible for N-terminal methylation, NTMT1 and NTMT2, that target a broad range of substrates associated with diverse biological pathways (Table 1) and are implicated in cancer and aging pathology.^4,5,7–17^ N-terminal methylation has been demonstrated to regulate protein-protein interactions and protein-DNA interactions.^2^ Trimethylation on CENP-A is crucial for constitutive centromere complex formation by recruiting CENP-T and CENP-I, which contributes to cell cycle progress and cell survival.^9,10^ Loss of trimethylation on CENP-B prevents binding to its centromeric DNA motif.^11^ Other research also suggests that N-terminal methylation might regulate protein-DNA interactions by creating a positive charge on the substrate protein, thus granting an ion-ion interaction with the negatively charged DNA backbone.^18^ In addition, N-terminal methylation may cooperate with N-terminal acetylation to regulate protein localization and differential interactions. N-terminal methylation increases MYL9 binding to DNA in the nucleus, while N-terminal acetylation contributes to its interaction with cytoskeletal proteins.^14^ The other enzyme with known α-N-terminal methylation activity besides Tae1 in yeast is Nnt1/Efm7, which is also capable of lysine methylation. Nnt1 appears to target a single substrate, the translational factor Tef1/eEF1A, and recognizes the Tef1 N-terminal G_1_-K_2_-E_3_-K_4_ sequence.^7^ In humans, METTL13 methylates eEF1α and is the functional homolog to Nnt1 despite the lack of sequence similarity between the enzymes (Table 1).^16,19^ To our knowledge, N-terminal methylation of proteins without the X_1_-P_2_-[K/R]_3_ motif or G_1_-K_2_-E_3_-K_4_ sequence has not been previously reported in eukaryotes and α-N-terminal methylation of non-canonical sequences is unclear.

**Table 1.** N-terminal methyltransferases and verified α-N-terminal methylated substrates in yeast and humans. * indicates the corresponding methyltransferase is not confirmed experimentally.

| N-terminal methyltransferase | Substrate | N-terminal sequence | Reference |
| --- | --- | --- | --- |
| yTae1 | Rpl12ab | PPKF | Webb, KJ. (2010) |
|  | Rps25a/b | PPKQ | Webb, KJ. (2010) |
|  | Rpt1 * | PPKE | Kimura, Y. (2013) |
| yNnt1/yEfm7 | Tef1(eEF1A) | GKEK | Hamey, JJ. (2016) |
| hNTMT1/2 | Rb | PPKT | Tooley, CS. (2010) |
|  | CENP-A | GPKR | Bailey, AO. (2013) |
|  |  |  | Sathyan, KM. (2017) |
|  | CENP-B | GPKR | Dai, X. (2013) |
|  | RCC1 | SPKR | Tooley, CS. (2010) |
|  | DDB2 | APKK | Cai, Q. (2014) |
|  | PARP3 | APKP | Dai, X. (2015) |
| | SETα | APKR | Tooley, CS. (2010) |
|  | KLHL31 | APKK | Tooley, CS. (2010) |
|  | MYL2 | APKK | Tooley, CS. (2010) |
|  | MYL3 | APKK | Tooley, CS. (2010) |
|  | MYL9 | SSKR | Neivtt, C. (2018) |
|  | Rpl23A | APKA | Tooley, CS. (2010) |
|  | Rpl12 | PPKF | Webb, KJ. (2010) |
|  | Rps25 | PPKD | Webb, KJ. (2010) |
|  | OLA1 | PPKK | Jia, K. (2019) |
|  | MRG15 | APKQ | Bade, D. (2021) |
| hMETTL13 | eEF1α | GKEK | Jakobsson, ME. (2018) |

Several methods have been used to confirm α-N-terminal methylation, including immunodetection and tandem mass spectrometry. Although antibodies have been developed for specific N-terminally methylated proteins, they have limited utility and are not suited for proteomic enrichment. Mass spectrometry is widely used for protein identification, detecting PTMs at a single protein level or proteomic level and studying protein N-terminomics. Numerous strategies have been developed for specific enrichment or quantitation of PTMs on protein N-termini, such as α-N-terminal acetylation. These techniques include quantitative isotopic labeling methods such as TAILS and SILAC,^20^ and negative enrichment methods such as COFRADIC^21^ and ChaFRADIC^22^ analysis. Recently, there has been an effort to encourage the archiving of proteomic datasets on various proteomic consortium databases with public accessibility, such as <u>PR</u>oteomics <u>IDE</u>ntification Database (PRIDE)^23^, ProteomeXchange^24^ and iProx^25^. Proteomic datasets deposited in these databases were generated for varied purposes, but they contain information about N-terminal methylation that was not previously searched or exploited. Thus, by crafting the search parameters for N-terminal methylation in sequence-based searching engines, reanalysis of datasets or repurposing would be an unbiased approach to investigating the N-terminal methylome landscape. Besides, many platforms and tools are established to facilitate datasets reanalyses such as the trans proteomics pipeline/PTMProphet^26,27^ and SearchGUI/PeptideShaker.^28^

This report demonstrates the utility of searching proteomic datasets generated by several techniques for methylated protein N-terminal peptides. We verified the presence of known α-N-methylated substrates and detected a collection of potentially α-N-terminal methylated proteins. We observed a close association between α-N-terminal methylation and the amino acid type at the first position of the protein sequence by analyzing the sequence pattern. By investigating the distribution of α-N-terminal methylation and α-N-terminal acetylation on various amino acids at the first position, we could characterize the sequence patterns of modifications across the proteome and between the two species. Surprisingly, the majority of N-terminal peptides identified by MS search did not have the canonical motif. We endeavored to validate three of the potential protein hits with purified overexpressed protein from yeast. We found that detecting methylation from this dataset repurposing approach can be elusive because experimental analysis of two non-canonical candidate hits, Ssa3 and Vma1 revealed full acetylation rather than the methylation initially predicted. However, we confirmed that the canonical motif-containing protein, Hsp31, is methylated. This study reveals intriguing global patterns of α-N-terminal methylation and serves as a foundation for further proteomic investigations of the N-terminal methylome.

## Experimental Procedures

### Script for Mascot automatic searching

A Perl script was created to automate searching and avoid overloading of the server. The Perl script allows downloading of raw files from the ftp server such as PRIDE or iProx by wget and subsequently convert them into .mgf files by msconverter tool, Proteowizard. MINE files with parameters described below were generated for corresponding datasets and used for Mascot searching by the command “nph-mascot.exe”. Mascot searching results were exported into .csv file by the command “export_dal_2.pl”. Finally, the methylated peptides were parsed and retrieved into .csv files. The script will be made available upon request.

### Yeast culturing and protein overexpression

Candidate proteins were purified using the movable open reading frame (MORF) collection from Dharmacon, where yeast genes were Gateway cloned into pBG1805 in frame with a C-terminal triple affinity tag, 6xHis-HA-3C protease site-Protein A. Protein expression is regulated by the *GAL* promotor.^29^ Yeast storage, culture and protein purification protocol were slightly modified from the Dharmacon technical manual.^29^ Yeast strains were streaked and stored on a synthetic complete agar plate with uracil dropout (SC-URA). On day 1, a single colony was inoculated into 2 mL SC-URA plus 2% glucose liquid media for overnight culture. On day 2, the culture was diluted into 20 mL SC-URA plus 2% raffinose at OD_600_=0.1 and incubated overnight. The culture was diluted into 800 mL of SC-URA plus 2% raffinose at day 3. When OD_600_ reached 0.8-1.2, 400 mL of 3X YP (3% yeast extract, 6% peptone and 6% galactose) media was added to induce the protein expression for 6 h. The yeast pellet was harvested by centrifuging at 4200 rpm for 10 min at 4 °C. The pellet was washed with 10 mL sterilized deionized water once and stored at −80 °C.

### Protein purification

Frozen yeast pellets were thawed from −80 °C and resuspended in CE buffer (50 mM pH 7.5 Tris-Cl, 1 mM EDTA, 4 mM MgCl_2_, 5 mM DTT, 10% glycerol, 0.75 M NaCl). PMSF (Promega) was added to 1 mM. The cell suspension was distributed into microcentrifuge tubes with an equal volume of 0.5 mm glass beads (Biospec). Yeast cells were lysed by bead beater (Biospec) for 7 cycles of 25 s interspersed with cooling on ice for 30 s. The lysate was centrifuged at 13000 rpm for 15 min. Clarified lysate supernatant was diluted into IPP0 buffer (10 mM pH 8 Tris-Cl, 0.1% NP40) and incubated with triple-washed IgG Sepharose 6 Fast Flow bead (Cytiva) for 2 h. IgG bead was separated, washed twice with IPP150 buffer (10 mM pH 8 Tris-Cl, 0.1% NP40, 150 mM NaCl) and subsequently wash twice with 3C cleavage buffer (10 mM pH8 Tris, 150 mM NaCl, 0.5 mM EDTA, 1 mM DTT, and 0.1% NP40). Bead was resuspended in 3C cleavage buffer and protein of interest was cleaved from bead using 3C protease (Acro Biosystems) with overnight digestion. The 3C protease was removed by glutathione agarose (Scientific). The eluent was passed through a 0.22 μM Spin-X centrifuge tube filter (Corning). Protein concentration was determined by BCA assay (Thermo Scientific).

### In solution digestion for Mass Spectrometry

The sample preparation protocol was adjusted from the literature.^30^ Protein purified and eluted from protein A agarose bead was used for sample preparation. Cold acetone (−20 °C) was added to the protein solution and stored at −20 °C overnight. Precipitated protein was collected by centrifugation at 15,000 x g for 10 min at 4 °C. The protein pellet was washed once with cold acetone and resuspended in 10 μl of 10 mM dithiothreitol (DTT, Sigma) in 25 mM ammonium bicarbonate, followed by incubation at 37 °C for 1 h. 10 μl of alkylation reagent mixture (97.5% acetonitrile, 0.5% triethylphosphine and 2% iodoethanol) was added to each sample and incubated at 37 °C for 1 h. The sample was dried in a vacuum centrifuge and digested using a barocycler (PBI). Hsp31, Ssa3 and Vma1 were digested by AspN (Sigma), GluC (Promega) and Trypsin (Sigma-Aldrich), respectively. Digested protein was cleaned with an UltraMicroSpin C18 column (The Nest Group) and dried into a pellet. The peptide pellet was reconstituted in 97% deionized water, 3% acetonitrile and 0.1% formic acid (v/v). An aliquot of 2-3 μl was run into the Q Exactive HF hybrid Quadrupole Orbitrap instrument (Thermo Scientific).^30^ The α-N-acetylation on Ssa3 was also confirmed by Orbitrap Fusion Lumos Tribrid mass spectrometer (ThermoFisher). Detailed protocols are included in Supporting data.^31–34^

### N-terminal methylation searching parameters in Mascot Daemon

Datasets were searched by Mascot Daemon with customized parameters to search for N-terminal methylated peptides in four different datasets as outlined in Table 2 and Table 3. For the ChaFRADIC dataset, iProx SILAC labeling dataset and MISL dataset, manual searches with Mascot Daemon (2.5.1 or 2.7.0.1) were conducted and the N-terminal peptide hit lists were matched with the results from a Perl-driven automatic searching process. A fourth dataset which we termed the Chemical Labeling dataset was searched using the automated script to identify the methylated N-terminal peptides. The exact search parameters used for each dataset are outlined in Table 2 including the variable and fixed modifications, tolerance level, peptide charge state and missed cleavages. Decoy databases were searched in all four cases and only matched N-terminal peptides with scores greater than 20 were used for hits selection and subsequent bioinformatics analysis. The false discovery rate (FDR) of each Mascot search for the ChaFRADIC dataset, iProx dataset and MISL dataset were manually checked and collected into an excel file (Supporting information Table S2).

**Table 2.** Searching parameters for ChaFRADIC, iProx and MISL datasets. The parameters used for repurposing each dataset is same as used in the original study except that the variable modifications are crafted for α-N-methylation.

|  | ChaFRADIC dataset<br>(PXD000292) | iProx SILAC labeling<br>(IPX0001550000) | MISL dataset<br>(PXD000606) |
| --- | --- | --- | --- |
| Species<br>(Repurposed<br>in this paper) | *S. cerevisiae* (WT and *icp55Δ*) | *S. cerevisiae* (*met6Δ*) and<br>Hela | S-adenosylmethionine (SAM) auxotroph<br>*YDR502C S. cerevisiae*<br>(*sam1Δ, sam2Δ*) |
| Database | Uniprot_S288C | Uniprot_S288C;<br>Uniprot_human | Uniprot_S288C |
| Variable<br>modification | Trimethyl (protein N-term)<br>Dimethyl (protein-N-terminal proline)<br>Dimethyl (K)<br>Dimethyl: 2H(4) (K) | Methyl:<br>2H(3)13C(1)(protein N-term)<br>Dimethyl: 2H(6)13C(2)<br>(protein N-term)<br>Trimethyl: 2H(9)13C(3)<br>(protein N-term) | For light isotope SAM labeled MS files:<br>Acetyl (protein N-term)<br>Methyl (protein N-term)<br>Dimethyl (protein N-term)<br>Trimethyl (protein N-term)<br>For heavy isotope SAM labeled MS files:<br>Acetyl (Protein N-term)<br>Methyl: 2H(3) (protein N-term)<br>Dimethyl: 2H(6) (protein N-term)<br>Trimethyl: 2H(9) (protein N-term) |
| Fixed<br>modification | Carbamidomethyl (C) | Carbamidomethyl (C)<br>Label: 13C(1)2H(3)(M) | Carbamidomethyl (C) |
| MS<br>tolerance | 10 ppm | 10 ppm | 50 ppm |
| MS/MS<br>tolerance | 0.02 Da | 0.5 Da | 0.8 Da |
| Enzyme | semi ArgC | trypsin/P | trypsin/P |
| Instrument | Q-Exactive mass<br>spectrometer (Thermo<br>Scientific) | Orbitrap Fusion mass<br>spectrometer (Thermo<br>Scientific) | LTQ-Orbitrap XL mass spectrometer<br>(Thermo) |
| Peptide<br>charge state | Charge +1 or<br>Charge +2 or<br>Charge +3 or<br>Charge +4 or<br>Charge +5 or<br>Based on which raw file is<br>searched | +2, +3 and +4 | +2, +3 and +4 |
| # missed<br>cleavages | 2 | 2 | 2 |

**Table 3.** Parameter settings for Chemical Labeling dataset (PXD0055831). Four subsets are generated with different combinations of enzyme and blocking reagent. Each of them is searched with same parameter settings as in original study but the variable modification is crafted for α-N-methylation.

|  | Group 1 | Group 2 | Group 3 | Group 4 |
| --- | --- | --- | --- | --- |
| Cell line | HEK293T | HEK293T | HEK293T | HEK293T |
| Variable modification | 1. Methyl (protein N-term)<br>2. Dimethylation (protein N-terminal)<br>3. Trimethyl (protein N-term)<br>4. D6-acetylation: 2H(3) (protein N-terminal) |  | 1. Methyl (protein N-term)<br>2. Dimethylation (protein N-terminal)<br>3. Trimethyl (protein N-term)<br>4. Propionylation (protein N-terminal) |  |
| Fixed modification | 1. Carbamidomethyl (C)<br>2. D6-acetylation: 2H(3) (K) |  | 1. Carbamidomethyl (C)<br>2. Propionylation (K) |  |
| MS tolerance | 15 ppm |  |  |  |
| MS/MS tolerance | 0.5 Da |  |  |  |
| Enzyme | Trypsin/P | GluC | Trypsin/P | GluC |
| Instrument | Q-Exactive mass spectrometer (Thermo Scientific) |  |  |  |
| Peptide charge state | +1, +2, +3 |  |  |  |
| # missed cleavage | 1 |  |  |  |

### Bioinformatic analysis of methylated peptides and proteins

Sequence conservation was graphically visualized using two tools, WebLogo^35^ and iceLogo^36^. Four types of sequence logos were generated for each dataset or dataset combinations with the redundant peptide list including iMet, redundant peptide list omitting iMet, nonredundant peptide list including iMet or nonredundant peptide list without iMet. The logo range was limited from the first amino acid position to the third of identified peptides. To generate the redundant lists of peptide sequences, methylated peptides were directly extracted from Mascot search export .csv file for each dataset and trimmed to the first three N-terminal amino acids.

Duplicated peptide sequences were removed to generate nonredundant lists. For logo analysis on iMet cleaved proteins, peptides containing iMet were excluded from each peptide list. In the WebLogo application, the small sample correction option was utilized to generate the frequency and content logo plots. In the iceLogo web application, the percentage scoring system was used to generate both logos and heat maps (p-value 0.05). The proteome background reference was utilized based on Swiss-Prot database composition of yeast or human proteins. The α-N-terminal methylation sequence patterns of yeast and humans were extracted from heat maps of yeast and humans combined peptide lists (nonredundant and iMet omitted).

Venn diagrams were generated with an online tool, Venny 2.1^37^, to analyze the reproducibility across datasets of each species. A list of nonredundant gene names of corresponding protein hits was extracted from the Mascot search export .csv file for each dataset. Three yeast gene lists and two human gene lists were used to generate the corresponding Venn diagram of yeast and humans.

### Western blot analysis for α-N-terminal methylation on Hsp31

The pBG1805 plasmid with Hsp31 ORF from Dharmacon was transformed into BY4741 WT yeast, *tae1Δ* BY4741 and *tae1Δ /Δ* BY4743 yeast. Protein expression and purification were conducted as described in the previous method section. Purified Hsp31 protein was analyzed via SDS-PAGE and western blot. A primary antibody capable of detecting mono-/di-N-terminal methylation (anti-N-me2Ser 2 antibody, a gift from Dr. Schaner-Tooley) was used at 1:1000 dilution.^38^ Chemiluminescent anti-rabbit (1:10,000) was used as secondary anybody. The western blot image was developed with ECL reagents, followed by X-ray film exposure.

## Results

### Discovery of N-terminal methylation of proteins with canonical motif and non-canonical motifs from four proteomic datasets

The following properties are important when considering repurposing a dataset to discover α-N-methylation: the N-terminal peptide enrichment methods applied, the chemical or isotopic labeling methods employed, the level of resolution of the instrument, and the use of an appropriate protease. Enrichment for protein N-terminal peptides increases the representation of α-N-methylation in the sample. Though employing positive enrichment is hindered by the lack of appropriate N-terminal methylation antibodies, many negative enrichment processes have been developed that deplete internal peptides and have successfully been applied to investigate α-N-acetylation. Hence, we focused on MS datasets using negative selection methods that eliminate internal peptides and increase the protein α-N-methylome representation. Isobaric modifications, such as α-N-terminal trimethylation (42.047 Da) and α-N-terminal acetylation (42.011 Da), require extra efforts to differentiate with fidelity. MS datasets generated by high-resolution instruments, isotopic labeling of either methyl group or acetyl group, and efficient fragmentation methods are thus favored. Trypsin is the most used protease in proteomics studies, but it is not optimal for identifying substrates containing the X_1_-P_2_-[K/R]_3_ canonical sequences. It cleaves at the carboxyl side of [K/R]_3_ and yields N-terminal peptides consisting of three amino acids that are poorly detectable.^39^ Hence, MS datasets generated by alternative proteases are more optimal for detecting N-terminal peptides.

Based on the above criteria, we focused on three types of datasets generated by the following methods: (1) Negative selective N-terminal enrichment method based on charge shift, such as ChaFRADIC method (e.g., ChaFRADIC dataset repurposed in this study); (2) Negative selection N-terminal enrichment method that depletes internal peptides, such as COFRADIC and chemical labeling methods (e.g., Chemical labeling dataset in this work); (3) Quantitative isotopic labeling methods such as ^13^C labeling (e.g., iProx dataset and MISL datasets in this work). Based on the above factors, we selected four datasets to demonstrate the prevalence of α-N-terminal methylation in yeast and humans. We performed manual searching for three experimental datasets in Mascot Daemon (herein referred to as the ChaFRADIC dataset^22^, iProx dataset^40^ and MISL dataset^41^). The parameters used for repurposing each dataset were identical to the original study except that variable modification was crafted individually to detect α-N-methylation (Table 2). A more complex Chemical Labeling dataset^42^ was searched using a Perl script to automate the searching process with the Mascot server (Table 3). All the datasets in ChaFRADIC had FDRs in the 2-8% range under specified significance level. Most Mascot searches of the iProx dataset had an FDR of 1-4% and most MISL searches had an FDR in the range of 4-9%. The frequency of α-N-terminal methylated peptides identified from decoy databases was monitored for ChaFradic, iProx and MISL datasets and was summarized in Table S2.

The MS/MS spectra of the ChaFRADIC, iProx and MISL datasets were manually examined to verify the site of methylation on protein N-termini, and the spectra of yeast hits are available in Figure S1. The number and identity of b/y ions, the maximum number of consecutive b/y ions and peak intensity were checked and recorded in Supporting information (Table S3). All the MS2 spectra have at least two b/y ion matches, although some spectra have weak peak signals. The MS2 spectra for the Chemical Labeling set were not inspected because it was not feasible to check all the spectra associated with this dataset manually. For most proteins identified, the MS2 spectra support the site of modification on protein N-termini. We refer to the identified proteins as hits in this study due to the limited number of peptides and relatively weak signal of ion fragmentation. Many software programs have functions available to measure PTM localization probability when more than one modification site might be present in one peptide, such as Mascot site analysis and MaxQuant PTM scoring^43^, and they all utilize MS/MS data. However, it is not possible to calculate the probability of methylation on protein N-termini without introducing other potential modification sites. Considering that lysine and arginine are the two sites most preferred to be methylated, we included mono-/di-/tri-methylation on lysine and mono-/di-methylation on arginine into the Mascot searches and monitored the site analysis probability score of Mascot. We recapitulated part of our initial hits with a very high probability of protein N-terminal localization (Table S4). These site localization analysis results are consistent with the manual interpretation based on MS/MS spectra and confirm the effectiveness of our manual examination on MS2 spectra and b/y ion series.

In the ChaFRADIC dataset, N-termini of yeast proteins are negatively enriched from *S. cerevisiae* spheroplasts based on charge reduction and two consecutive SCX separations. Briefly, free α- and ɛ-amino groups on lysine side chains or protein N-termini in the samples were blocked with a dimethyl group by formaldehyde and cyanoborohydride. N-terminal proline is the only exception and can only be mono-methylated during the blocking process.^44^ Trypsin was used for sample digestion with ArgC specificity in this case because trypsin recognition and subsequent cleavage are blocked by dimethylated lysine. Digested peptides were fractionated based on their charge state by the 1^st^ SCX (Strong Cation Exchange) separation. Internal peptides in each fraction with free α-amino group released by trypsin digestion were deuteron-acetylated and had their positive charge reduced by one. The change in the charge states of internal peptides leads to a peak shift in the 2^nd^ SCX separation and are depleted. Only the fractions remaining unchanged in the two SCX separations were used for further MS/MS identification by Q-Exactive mass spectrometry, which contains increased representation of protein N-termini, either protected by extraneous dimethylation or innate modifications.

The ChaFRADIC method selectively enriches modified N-terminal peptides and excludes the internal peptides. However, the blocking method obscures the origin of monomethylation and dimethylation (except dimethylated proline) on protein α-N-amino groups; hence, we focused on identifying native trimethylation at protein α-N-termini and dimethylation on protein N-terminal proline. We identified 42 N-terminal methylated peptides corresponding to 12 yeast proteins from 5 SCX fraction MS files, including Ssa3, Por1, Gpm1, Rpl15a, Rpl16b, Hom2, Rpl12a, Rpl24a, Rpl24b, Rpl26a, Rpl27a and Tef1. It should be noted that Ssa3 and Tef1 do not have a C-terminal tryptic end but were included as possible hits. The frequency of identified peptides carrying a non-tryptic end in the complete dataset is ~16%-20%, possibly due to non-specific digestion, solution degradation or impurity of the endonuclease.^45^ All of them were putatively identified as protein N-terminal trimethylated, except that Gpm1 and Rpl12a are dimethylated on P1. The α-N-trimethylation for all proteins was on the first amino acid after iMet is excised, except for Rpl24a/b (Figure S1). The site localization of each hit was confirmed by MS2 spectra and the site analysis score of Mascot (Table S4). Two proteins, Tef1 and Rpl12ab, have previously been reported to be α-N-methylated. Tef1 is the yeast translation elongation factor 1a (eEF1A) known to be α-N-trimethylated by the dual lysine and protein α-N-terminal methyltransferase, Nnt1 (Table 4 and Table S5).^46^ Rpl12ab is encoded by two separate genes, *RPL12A* and *RPL12B*, resulting in identical polypeptide sequences. Five methyl groups were assigned to the first three amino acid positions of Rpl12ab in our Mascot searching, which is equivalent to the full methylation state on both the protein N-termini and the K_3_ side chain. This methylation state agrees with a previous report for Rpl12ab.^4,47^ To our knowledge, this is the first report of α-N-terminal trimethylation for the other five hits and surprisingly, they do not contain the canonical motif X_1_-P_2_-[K/R]_3_ motifs recognized by the predominant α-N-terminal methyltransferases in yeast, Tae1.

**Table 4.**
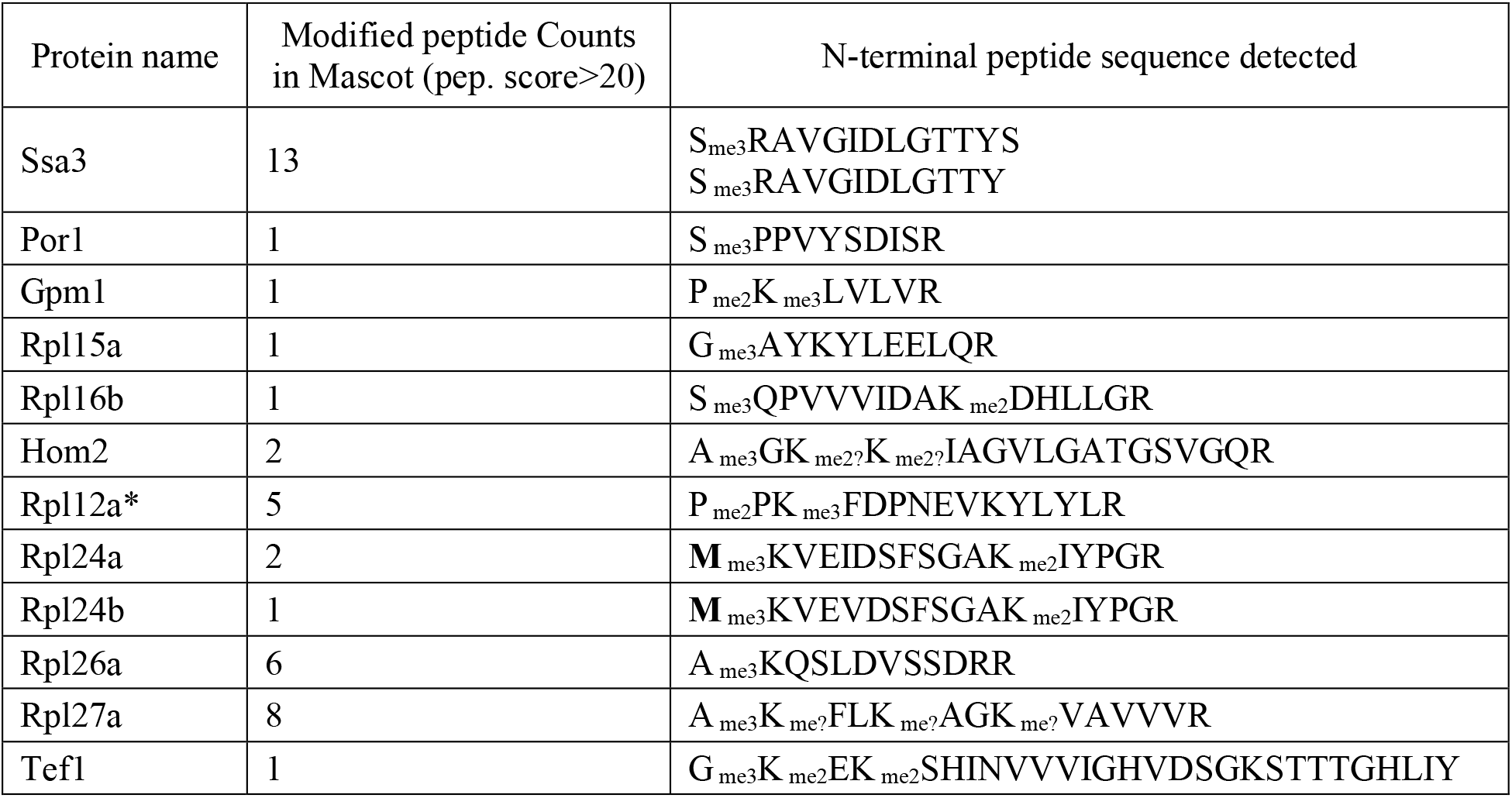
Methylated hits from the ChaFRADIC dataset. 12 proteins are tri-methylated while none has the canonical motif recognized by Tae1. The modification localizations were determined by Mascot site analysis and MS2 spectra. *Rpl12a peptide is detected to have 5 methyl groups which is consistent with reported proline_1_ dimethylation and lysine_3_ trimethylation. Ambiguous localization of lysine methylation is labeled with “?”.

The iProx isotopic labeling dataset was designed initially for identifying lysine methylation using ^13^CD_3_-methionine in *E. coli*, yeast and Hela cell line.^40^ We examined ^13^CD_3_ labeled N-terminal methylation in the datasets for both eukaryotes (yeast and Hela cell line), which differentiates the α-N-acetylation (42.011 Da, UNIMOD) from α-N-terminal trimethylation (54.113 Da, UNIMOD). This iProx dataset and the MISL dataset described below are the two proteomic datasets in our report that used heavy isotope label methyl donors. Here, we searched for all heavy isotope α-N-terminal methylation species (mono-/di-/tri-methylation), whereas only α-N-monomethylation was searched in the original publication.^40^ We discovered 11 unique N-terminal methylated peptides (iMet cleaved) from nine unique protein methylation events in the HeLa dataset, including seven α-N-monomethylation, three α-N-dimethylation and one α-N-trimethylation. Only the ATG3 (Q_1_-N_2_-V_3_) protein was found to have two α-N-methylation species, monomethylation and dimethylation (Table S5). Identification of multiple methylation states increases the confidence that these proteins are biologically methylated. For the yeast dataset, 11 unique α-N-methylated peptides (iMet cleaved) were detected from two unique proteins, including three α-N-monomethylated peptides and eight α-N-dimethylated peptides. One protein, the gene product of *ARB1* (P_1_-P_2_-V_3_), had evidence of multiple N-terminal peptide methylation states (Table S5). Arb1 was the only protein previously reported to be α-N-monomethylated in the associated publication from this dataset and the detection of methylation of Bdh2 (R_1_-A_2_-L_3_) is novel.^40^ Notably, none of the peptides contained the canonical recognition motif. Arb1 has an N-terminal sequence of P_1_-P_2_-V_3,_ which has similarity to the P_1_-P_2_-K_3_ canonical motif but is missing the crucial lysine residue at the third position.

The MISL dataset was generated using a strategy called <u>M</u>ethylation by <u>I</u>sotope <u>L</u>abeled <u>S</u>AM (MISL). Briefly, *S. cerevisiae* (*sam1Δ* and *sam2Δ* BY4741 background) SAM deficient cells were metabolically labeled with either the heavy form of CD_3_-SAM or the light form of CH_3_-SAM. The subsequent lysates were mixed at 1:1 ratio and digested with trypsin before MS/MS analysis. The original study only searched for methylation on amino acid side chains and did not investigate α-N-terminal methylation. This data was repurposed by searching for the light isotope form of α-N-methylation species and the heavy form of α-N-methylation species with the same search parameters as in the original study. We focused on heavy labeled methylation because it easily distinguishes trimethylation from acetylation and demonstrates *in vivo* methylation. 49 unique heavy-N-terminal methylated peptides (iMet cleaved) were found for 48 unique proteins, including 23 α-N-monomethylated peptides, 13 α-N-dimethylated peptides and 13 α-N-trimethylated peptides. Several proteins were found as multiple methylation species or in multiple datasets. Ahp1, with the S_1_-D_2_-L_3_ amino acid sequence, was found to be α-N-monomethylated and α-N-trimethylated with heavy isotope label and α-N-trimethylated with the light isotope. Rpn13 (S_1_-M_2_-S_3_) was α-N-trimethylated with heavy isotope label and α-N-dimethylated with the light label. Asc1 (A_1_-S_2_-N_3_) was found to be α-N-trimethylated in both light and heavy isotope labels. Only one protein, Rps25a/b (P_1_-P_2_-K_3_), contained the canonical motif and was shown to be heavy isotope α-N-monomethylated. Interestingly, four more proteins containing the canonical motif were shown to be heavy isotope α-N-methylated with low peptide scores, including Rpl12ab, Rpt1, Ola1 and Hsp31. Rpl12ab and Rpt1 were previously reported to be α-N-dimethylated.^4,5^ The use of trypsin in this study likely leads to suboptimal detection of canonical motif-containing peptides, as discussed earlier.

Another large proteomic data set was investigated that was generated using a negative N-terminal enrichment method. Four different individual sample preparations depending on the combination of endoprotease and chemical blocking reagent. Briefly, the free α- and ɛ-amino groups in HEK293T cell lysate were blocked (by propionylation or D3-acetylation), including protein α-amino group and amino groups on lysine side chains. The proteomic sample was then subjected to enzymatic digestion by GluC or Trypsin.

Internal peptides with the free amino group were removed by NHS activated agarose and thus protected protein N-termini were negatively selected. The results of this search are listed in Table S5. The majority of the hits identified did not contain the canonical motif while five unique protein hits contained the canonical X-P-K motif (Table S6). Two of these hits have previously been demonstrated to be methylated: SET (A_1_-P_2_-K_3_) was reported to be α-N-terminally trimethylated and RPL23A (A_1_-P_2_-K_3_) has previously been demonstrated to be α-N-terminally dimethylated and trimethylated.^8^ In addition, four other protein hits containing the canonical motif were identified that had corresponding N-terminal methylated peptides with scores near the cutoff.

Overall, we screened for potential N-terminal methylation events by searching and parsing three yeast datasets and two human datasets generated by various techniques. We identified three yeast proteins and five human proteins containing known α-N-terminal recognition motifs (X_1_-P_2_-[K/R]_3_ or G_1_-K_2_-E_3_-K_4_), in which all the yeast proteins and two human proteins were verified in earlier studies. This result indicates the efficacy of the repurposing method in identifying potential α-N-methylation hits with canonical motifs and yields a large set of proteins without the canonical motif. These results suggest a greater prevalence of α-N-terminal methylation in the yeast and human proteomes than previously appreciated. However, careful verification is needed to confirm potential hits due to possible false localization of a methyl group on lysine/arginine side chains instead of the N-terminus or because of ambiguity between isobaric modifications.

### Sequence analysis of methylated peptides shows semi-specific conservation at the first position of the α-N-terminal methylome

The identification of numerous peptides in multiple datasets allowed the investigation of the global conserved pattern of α-N-terminal methylation events. We applied WebLogo^35^ and iceLogo^48^ to analyze the sequence conserveness of α-N-methylated proteins in yeast and humans (Figure 1, Figure 2 and Figure S2).

**Figure 1.**
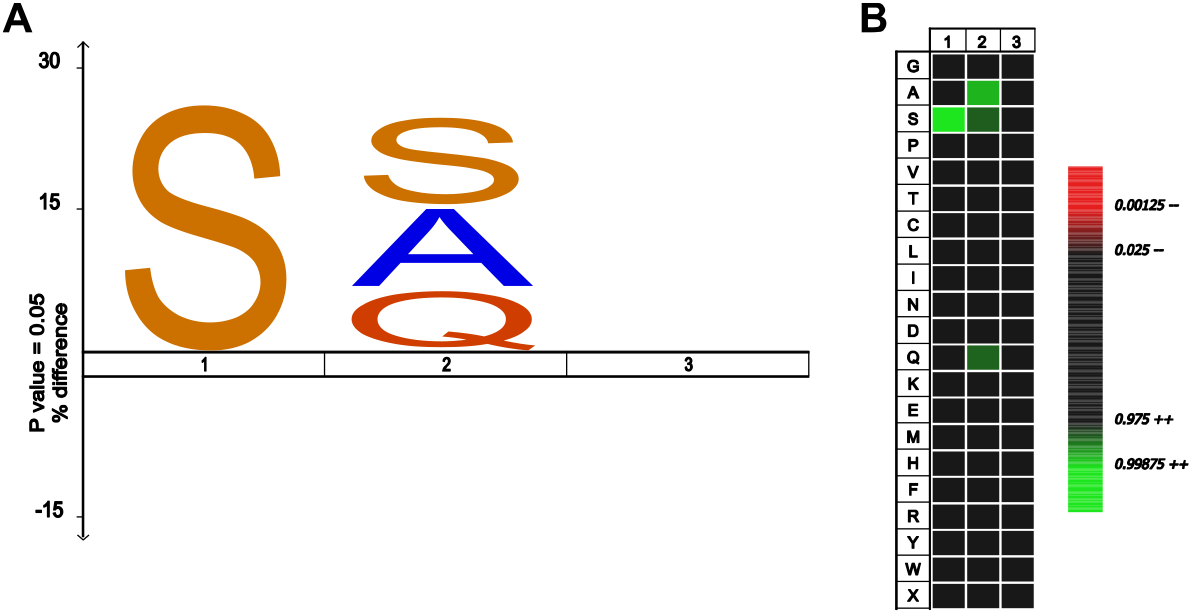
N-terminal amino acid conservation for methylated peptides in the yeast datasets. A. iceLogo was generated by iceLogo web application. Over-presented amino acids are show above center line and under-represented amino acids are below the center line. A S_1_-[S/A/Q]_2_ pattern is clearly enriched in yeast dataset. B. Heatmap demonstrates over-represented amino acids and under-represented amino acids colored green and red, respectively.

**Figure 2.**
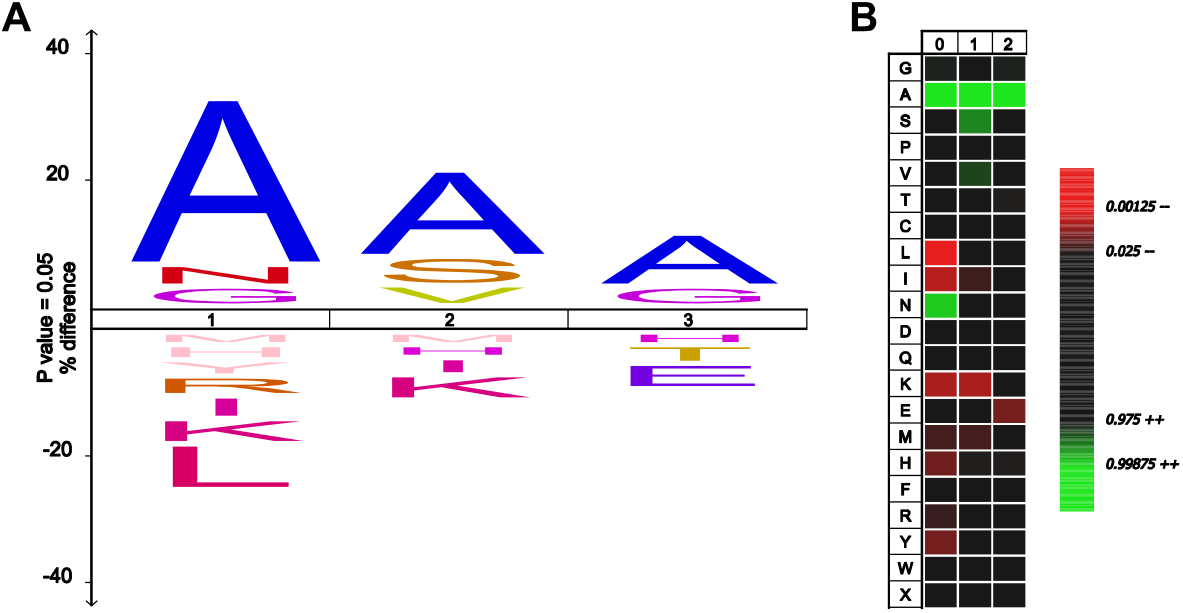
N-terminal amino acid conservation for methylated peptides in the human datasets. A. iceLogo generated by iceLogo web application as in Figure 1. Over-represented amino acids are show above center line and under-represented amino acids are below center line. A [A/N/G]_1_-[A/S/V]_2_-[A/G]_3_ pattern is clearly enriched in the human dataset. B. Heatmap demonstrates over-represented amino acids and under-represented amino acids colored green and red, respectively.

To avoid over-interpretation from the sequence analysis and the uncertainty introduced by isobaric modifications (especially the prevalent α-N-terminal acetylation), unlabeled trimethylated protein hits are excluded from the logo analysis. For yeast datasets, we only analyzed the heavy labeled hits from the iProx and MISL datasets to determine sequence pattern in yeast. Hits from ChaFRADIC were excluded from logo analysis. We used the heavy isotope labeled methylation hits from the iProx Hela subset and unlabeled mono-/di-methylated hits from the Chemical Labeling datasets for the human proteome logo analysis (Table S7). Our subsequent investigations focused on the lists of nonredundant peptides with iMet removed and other logo visualizations can be found in supporting information (Figure S2).

We combined a total of 47 nonredundant peptides (iMet cleaved) identified from the heavy isotope labeled MISL yeast dataset and iProx yeast datasets for yeast logo analyses (Table S7). iceLogo visualization suggests that the first two amino acid positions might be more determinant in α-N-terminal methylation events. Serine is significantly enriched at the 1^st^ position, while the second position appears to have a more diverse preference for serine, alanine and glutamine (Figure 1A and 1B). The 3^rd^, 4^th^, and 5^th^ positions are not conserved in the nonredundant and likely do not factor in influencing non-canonical N-terminal methylation (data not shown after the 3^rd^ position). No amino acid is under-represented. This result reveals a pattern of [S]_1_-[S/A/Q]_2_ for N-terminal methylation events in the yeast proteome.

In the human proteome methylation data, we analyzed a list of 222 nonredundant peptides (iMet cleaved) from the heavy isotope labeled iProx human dataset and the Chemical Labeling dataset. Similar to the yeast iceLogo, the first three amino acids are more relevant to the α-N-terminal methylation events in humans. iceLogo analysis demonstrates that alanine is significantly enriched at the 1^st^ position along with glycine and asparagine. In addition, several amino acids are significantly under-represented at the first three positions, which indicates that they are less favored for the α-N-terminal methylation. A pattern of [A/N/G]_1_-[A/S/V]_2_-[A/G]_3_ was indicated for α-N-terminal methylation events in the human proteome (Figure 2A and 2B).

Evidence for the NTMT1/2 canonical sequence, which prefers proline at the second position and lysine at the third position in human global N-terminal methylome was not detected. Eight protein hits containing the canonical motif were identified with peptide scores greater than 20, and an additional eight proteins with canonical motifs were identified with less confidence (Supporting information Table S6). However, these were not sufficient to influence the overall methylation pattern uncovered in our approach. Methylated Rpl12ab peptides were detected in the ChaFRADIC and MISL datasets with high and low scores, respectively. In total, there are 15 nonredundant protein hits containing canonical motifs from all four datasets. We found that nine out of the 15 were identified from the HEK cell line, and six were from the yeast datasets. Six protein hits have been verified in earlier literature, including Rpl12ab, Rps25a/b, eEF1 and Rpt1 in yeast and Rpl23a, SET in humans^4^. Overall, our motif analysis suggests the presence of an additional methylation process that is not restricted to the X_1_-P_2_-[K/R]_3_ motifs and is dependent mainly on the first amino acid for semi-specificity. The NTMT1/2 enzymes are likely not responsible for this non-canonical methylation, because extensive peptide specificity studies show strong specificity for the X_1_-P_2_-[K/R]_3_ motif or related motif.^4,8^

### Comparison of N-terminal methylation with N-terminal acetylated proteins

Two prevalent modifications on the protein α-N-termini are α-N-terminal acetylation and α-N-terminal methylation. There is evidence that the competition between these two modifications on the same protein regulates its dual localization, such as MYL9.^14^ Here, we investigate the overlap of α-N-terminal methylation events and α-N-terminal acetylation events. By integrating our α-N-terminal methylation repurposing results with α-N-terminal acetylation reported from the original papers, we classified the identified protein hits into four categories based on their N-termini modification states: only α-N-terminal acetylated, only α-N-terminal methylated, dual modification or not modified by either of the two. Each category was counted, and the frequency of each category was plotted in a pie chart (Figure 3A). The proportion for each class indicates the distribution of modifications for the proteome. To further explore the extent of both modifications in the global proteome, we plotted the frequency of each category over the 1^st^ amino acid type of the protein hits. The distribution plots reveal that subsets of proteins with some types of amino acids might be under dual control or prefer either of the two modifications. (Figure 3B and 3C).

**Figure 3.**
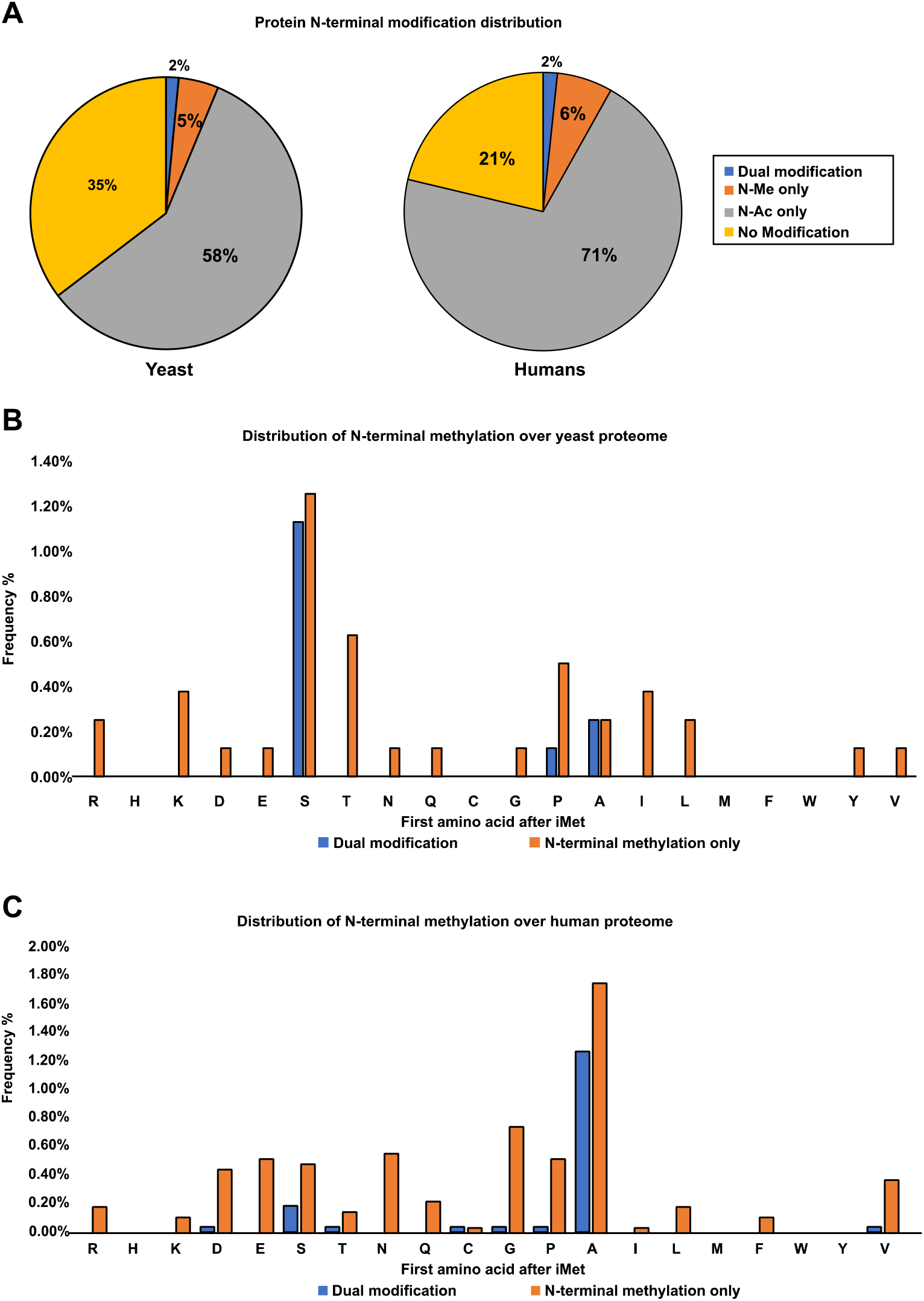
The frequency of proteins found to have only α-N-terminal methylation or both modifications are plotted according to the type of amino acid. Only iMet cleaved proteins are used for this graph. (A) Pie charts shows the proportion of yeast or human proteome undergo α-N-terminal methylation or α-N-terminal acetylation. (B) The frequency plot for yeast datasets, combining hits from MISL and iProx datasets. (C) The frequency plot for human datasets, combining hits from Chemical Labeling and iProx datasets. The frequency of N-terminal acetylation only was not depicted in the frequency plots because high levels of acetylation for several amino acids results in a scale that obscures the visualization of lower frequency methylation levels. The frequency plots including iMet retained proteins are in Figure S3.

We utilized the nonredundant proteins from our pattern analysis for analyzing the distribution of modifications. Analysis that included protein hits modified on iMet (which consist of more than 30% of total identified protein hits) showed that methionine did not change the overall distribution of α-N-terminal modifications in the proteome. However, iMet is predominantly modified by both methylation and acetylation (Figure S3). Both yeast and humans datasets had 60%-70% of protein hits that were α-N-terminal acetylated, consistent with other estimates of N-terminal acetylated proteins.^49,50^ The α-N-terminal methylated protein hits consist of 7-8% of the total proteome in both yeast and humans. Only 2% out of the 7-8% methylated protein hits could be under the dual control of α-N-terminal methylation and α-N-terminal acetylation. In addition, around 21% human proteome and 35% yeast proteome did not have evidence of either modification. The proportion of α-N-terminal acetylation is approximately 10 times as much as α-N-terminal methylation (Figure 3A). The high prevalence of α-N-terminal acetylation could be related to the close association of acetyltransferases with ribosomes.^49^ There is no current evidence that α-N-terminal methyltransferases are in close spatial proximity with ribosomes.

We examined each modification’s distribution and preference on the 1^st^ amino acid for both yeast and humans. The identified proteins with various α-N-terminal modification states were clustered by the 1^st^ amino acid, and the fraction of each α-N-terminal modification states (only α-N-terminal methylated, only α-N-terminal acetylated, dual modification or not modification) was determined by dividing the modified population of each cluster against the total identified protein population.

Yeast α-N-methylation was detected on all amino acids with less frequency for H, C, M, F and W. As expected, methylation was detected on the amino acids A, S and P as in the Tae1/NTMT canonical motif and amino acids with methylatable side chains such as R and K. Unexpectedly, methylation was also detected on D, E, N, and Q which are rarely reported to be N-methylated. The dual modification was observed predominantly on A and S (Figure 3B). In the human proteome, all amino acids are methylated except H, M, W and Y. The A and G constitute the majority of the N-terminal methylation detected. The dual-modification types were observed predominantly on A and S. Interestingly, although N-terminal proline is thought to block acetylation,^51^ we note that proline acetylation was identified in the original paper and an additional recent report has demonstrated proline acetylation (Figure 3C).^52^

To summarize, in both human and yeast datasets, around 7-8% of proteins are subject to α-N-methylation, which is one-tenth of the proportion of N-terminal acetylation. α-N-methylation is more prevalent in both species and not strictly conserved on the initial amino acid type. Both charged amino acids (D, N, Q, E, K and R) or hydrophobic amino acids (L, V, A and G) could be α-N-methylated. A and S at protein N-termini are the most frequent for α-N-methylation, while the combination of α-N-methylation and α-N-acetylation is also prevalent. Yeast and human datasets showed similar patterns in the distribution of α-N-methylation and α-N-acetylation.

### Ssa3 and Vma1 are predominantly α-N-terminal acetylated instead of methylated

PTM events identified from a proteomic dataset maybe not accurate and difficult to locate because of peptide misassignment and limited fragmentation detection. Thus, verification is necessary for ensuring α-N-terminal methylation is correctly identified from the repurposed datasets because of the closely matched mass of trimethylation and acetylation modifications. We demonstrate that the identification of peptide trimethylation is easily confused with acetylation. Importantly, we validated novel protein methylation of Hsp31. We attempted to verify three hits detected using our repurposed datasets; two yeast protein hits without the canonical motif (Ssa3 and Vma1) and one protein with a canonical motif were purified (Hsp31). To verify the results from proteomic dataset searching and semi-quantify the ratio between different N-terminal PTMs, we purified the proteins using the Dharmacon yeast ORF collection. The C-terminally tagged proteins of interest were overexpressed using the *GAL* promoter. Subsequently, the overexpressed protein was affinity-purified with Protein A agarose beads. All proteins were purified to greater than 95% purity as assessed by SDS-PAGE (Figure 4A) and examined by western blot (Figure S4). Ssa3 is a member of stress-inducible member of the heat shock protein 70 family. The Ssa3 band matches the predicted molecular weight and the Vma1 band matches the molecular weight predicted after intein splicing (Figure 4; Figure S5 and S6).^53^ The Vma1 intein splicing was verified based on the protein coverage map (Figure S7). Ssa3 was digested by GluC, while Vma1 was digested by trypsin in solution followed by intact protein analysis and tandem MS.

**Figure 4.**
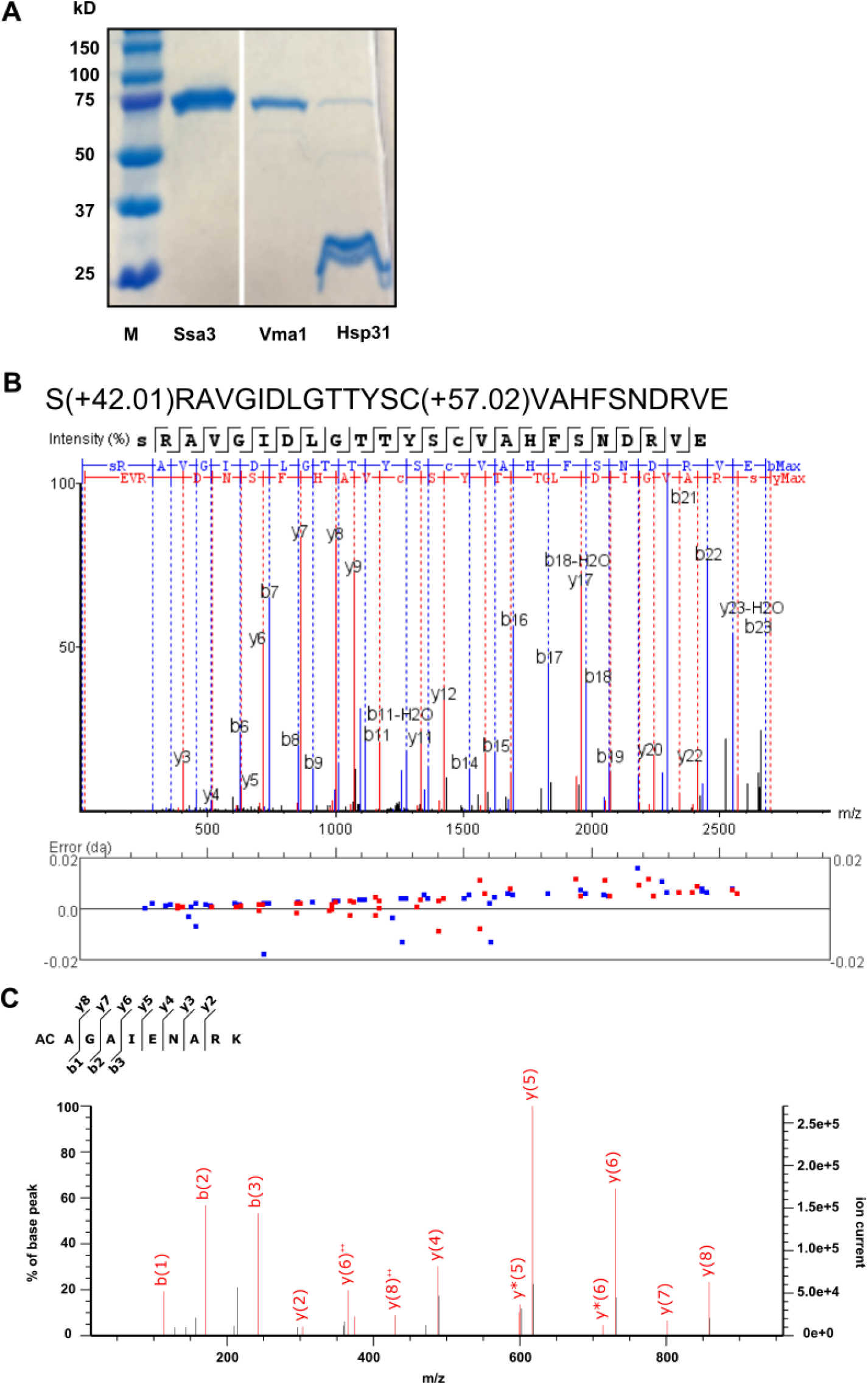
Verification of N-terminal modifications potentially methylated substrates. (A) Vma1-MORF, Ssa3-MORF and Hsp31-MORF were purified by protein A affinity purification and released from the beads using protease 3C. Purified proteins were analyzed by Coomassie blue staining. (B) Fragmentation of Ssa3 confirmed N-terminal acetylation on Ssa3 Serine_1_. Error is within 0.02 Da range. (C) MS2 spectrum indicates that Vma1 is acetylated on Alanine_1_.

Analysis of the purified Ssa3 by intact mass spectrometry suggested a major mass shift (+40.75 Da) corresponding to trimethylation (+42.047 Da) or acetylation (+42.0311 Da) (Figure S7). Furthermore, the tandem MS analysis using an Orbitrap Fusion Lumos Tribrid mass spectrometer (ThermoFisher) confirmed that the N-terminal peptides were fully acetylated instead of trimethylated as predicted with a protein sequence coverage of 99% (Figure 4B & Figure S7). Analysis of the b ions and y ions collected demonstrated the acetylation on the N-terminal serine. We performed both database searches and de novo searches with Mascot Daemon (MATRIX SCIENCE), PEAKS X plus (Bioinformatics Solutions Inc.) and BioPharma Finder software (ThermoFisher). In PEAKS search with 10ppm tolerance on the precursor and 0.02 Da fragment error, 309 peptides were matched to Ssa3, covering 99% of the protein sequence. 95 peptide-spectrum matches (PSM) of Ssa3 were found to be exclusively acetylated at α-N-terminus, consistent with the significant peak detected using intact mass spectrometry (Figure S7). All the PSM were +3 charge. The finding that Ssa3 is acetylated rather than trimethylated using Mascot is probably a result of utilizing the more sensitive Lumos instrument at a higher resolution (60K) than that used in the ChaFRADIC study. Interestingly, Ssa1 was also one of the top hits indicating it is a copurifying protein, and the peptides of Ssa1 also were identified to be α-N-terminal acetylated. Ssa1 is predicted to be acetylated, and mutations preventing N-terminal acetylation appear to decrease the ability of binding to prions.^54^ Although Ssa1 is assumed to be acetylated in the literature, the acetylation evidence is indirect, whereas our MS data is the first direct evidence confirming that heat shock protein 70s are N-terminal acetylated.^54,55^ The purified Vma1 has 55% protein coverage and 21 N-terminal peptides detected. All N-terminal peptides were α-N-terminal acetylated and had PSMs with +2 charge. The NatA dependent acetylation of Vma1 is consistent with earlier reports and annotations in PTM databases.^56^ Although the repurposed MS dataset suggested that Vma1 is α-N-monomethylated, our MS analysis of Vma1 did not detect methylation but predominantly α-N-acetylation. Methylation could occur at a low frequency because Vma1 monomethylation was detected in two separate repurposed datasets. The failure of detecting α-N-monomethylation is possibly due to the overexpression strategy employed to purify the protein or the low representation of Vma1 methylation under physiological conditions. These results demonstrate the importance of instrument sensitivity and approach in the verification of these modifications. Further comprehensive studies are needed to examine and verify methylation frequencies but are beyond this study’s scope.

### Identification of methylation of the canonical motif-containing protein, Hsp31

We then investigated another potential substrate consistent with the canonical α-N-terminal motif. Hsp31 was suggested to be α-N-methylated in the MISL dataset, although the peptide score was below the score cutoff of 20. We predicted that Tae1 is likely responsible for the methylation of Hsp31. Here, we determined the N-terminal modification on Hsp31 by tandem mass spectrometry and immunodetection. Hsp31 was purified from the yeast MORF collection as described earlier (Figure S4). For tandem mass spectrometry, purified Hsp31 was digested with AspN for MS/MS analysis. Protein coverage was 75%, and MS fragmentation was consistent with α-monomethylation (Figure 5A and Figure S8). 16 PSM corresponded to the Hsp31 N-terminus in total. Two PSM were consistent with monomethylated Hsp31 N-termini. The y12 ion in 1 PSM indicates the presence of monomethylation on Ala_1_ site. Five PSM were consistent with either N-terminal dimethylation or formylation, but MS2 spectra matched formylation with lower ppm error (~3.6 ppm for formylation and ~21.6 for dimethylation. All the N-terminal PSM had +2 charge or +3 charge, which could be explained due to the presence of multiple lysines. Hsp31 purified from wild type BY4741 yeast was shown to be methylated by a polyclonal antibody designed to recognize dimethylated SPK N-terminal sequences (a gift from Dr. Schaner-Tooley)^38^. This antibody also has cross-reactivity for mono-methylated N-termini and similar sequences such as A_1_-P_2_-K_3_ based on synthetic peptide dot blot assays (data not shown). The loss of methylation was observed in Hsp31 purified from haploid *tae1Δ* and diploid *tae1Δ/tae1Δ* yeast (Figure 5B). This data further supports monomethylation on the Hsp31 N-terminus, albeit at a low level under these culturing and expression conditions. The lack of signal in the *TAE1*-deficient yeast strains indicates that methylation is contributed mainly by Tae1. In conclusion, we were able to verify novel N-terminal methylation of a protein with a canonical motif demonstrating the repurposing approach’s validity.

**Figure 5.**
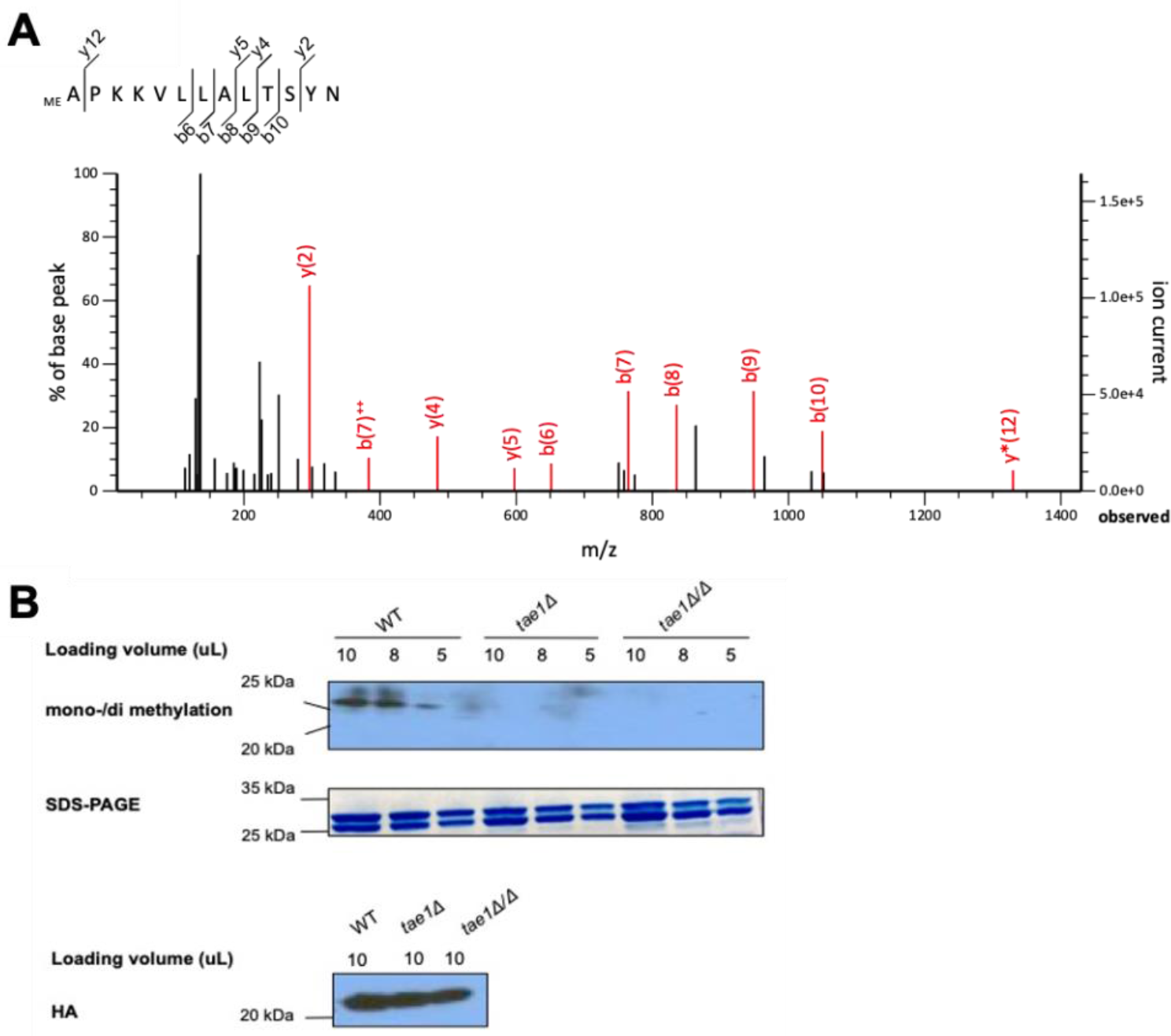
Hsp31 is N-terminally methylated. A) MS2 spectrum consistent with monomethylation. Mascot score for this PSM is 44. An additional monomethylation MS2 spectra is depicted in Figure S8. (B) A polyclonal antibody recognizing mono and dimethylated N-termini was used to probe Hsp31 purified from WT, haploid *tae1Δ* and diploid *tae1Δ*/*tae1Δ* yeast strains. The same samples were stained with coomassie blue after separate SDS-PAGE to show purity – an extra band is evident due to proteolysis during purification. Samples were also probed with anti-HA antibody which is a tag retained on the C-terminus after purification.

## Discussion

Protein N-terminal methylation is a novel protein modification type that has garnered increasing attention recently. Two classes of α-N-methyltransferases found in yeast and humans recognize distinct protein N-terminal sequence patterns. Yeast Tae1 and the human orthologs, NTMT1/NTMT2, recognize a canonical X_1_-P_2_-[K/R]_3_ motif on substrate proteins and contribute to most of the N-terminal methylation events reported to date. Another yeast N-terminal methyltransferase, Nnt1, is a more specific enzyme that appears only to have a sole target, Tef1.^46^ Tef1 contains a G_1_-K_2_-E_3_-K_4_ N-terminal sequence and does not share similarity with the X_1_-P_2_-[K/R]_3_ motifs. The functional, but not the sequence ortholog, of Nnt1 in humans appears to be METTL13 which also recognizes a similar sequence and methylates the orthologous substrate, eEF1a.^16,19^ Besides these two classes of N-terminal methyltransferases, the detection of N-terminal methylation events usually relies on assuming that the canonical motif sequence has the potential to be methylated, and proteins without the motif are unmethylated. Hence the potential of identifying non-canonical methylation events is largely excluded. An *in vitro* ^3^H-methylation assay with 9-mer synthetic N-terminal peptides suggests the X_1_-P_2_-[K/R]_3_ is the most favorable substrate for yeast and human N-terminal methyltransferases but also demonstrates varying amino acid preferences.^4^ There is an urgent need to develop an unbiased method to screen potential protein N-terminal methylation events regardless of the N-terminal sequence. Repurposing current datasets can verify previously characterized protein modifications at a proteomic level and enable the discovery of potential N-terminal methylation events in an unbiased way.

In this study, we expanded our knowledge of the extent of the N-terminal methylome in yeast cells and human cell lines. Only a small subset of the potential hits contains the canonical methylation motif. In the ChaFRADIC dataset, we identified two known α-N-methylated substrates, Rpl12ab (P_1_-P_2_-K_3_) and Tef1 (G_1_-K_2_-E_3_-K_4_). In the MISL dataset, we identified one known substrate Rps25ab (P_1_-P_2_-K_3_) with a peptide score >20 and two known substrates with lesser scores, Rpl12ab (P_1_-P_2_-K_3_) and Rpt1 (P_1_-P_2_-K_3_).^4,5^ We also identified two low score hits with the canonical motif but have not been demonstrated to be methylated, which are Hsp31 (A_1_-P_2_-K_3_) and Ola1 (P_1_-P_2_-K_3_). We confirmed α-N-monomethylation of Hsp31 by mass spectrometry and immunoblotting. From the Chemical Labeling dataset, we identified two known substrates, Rpl23a (A_1_-P_2_-K_3_) and SET (A_1_-P_2_-K_3_) with peptide scores >20. Three more protein hits with the canonical motif were identified that had not been previously reported, which are CMTM2 (A_1_-P_2_-K_3_), XPNPEP1 (P_1_-P_2_-K_3_) and ZC3H15 (P_1_-P_2_-K_3_). In addition, we found 4 other protein hits with lower scores, RGS11 (P_1_-P_2_-R_3_), PCSK6 (P_1_-P_2_-R_3_), MORF4L1 (A_1_-P_2_-K_3_) and TOMM34 (A_1_-P_2_-K_3_) (Table S6). There are 45 yeast proteins with the canonical motif in the yeast proteome, and many were not identified by our repurposing approach demonstrating that we are under-sampling the N-terminome (Table S1). The remaining hits identified by our study appeared to have moderate conservation in the first position but minimal conservation on adjacent amino acid positions. Only Arb1 contains a related sequence, P_1_-P_2_-V_3_, while the majority contain unrelated sequences. This indicates N-terminal methylation might be more widely occurring and is not restricted to the X_1_-P_2_-[K/R]_3_ motif-containing proteins. We found serine in yeast and alanine/serine/asparagine in humans are over-represented compared to proteome background at the first position, indicating that proteins with these amino acids are the most methylated in humans and yeast, respectively.^49^ We also note that some amino acids are under-represented, such as Met/His/Trp/Arg/Ile/Lys/Leu at the first position after iMet. These N-terminal peptides might be less favored, or they are not optimal for MS detection because of peptide length or low representation.

It should be noted that protein hits from the repurposing result should be carefully investigated and further confirmed. Our attempt to validate two proteins with non-canonical sequences indicated that they were predominantly acetylated. In the case of Ssa3, the repurposing approach initially indicated trimethylation of the protein, but our experiment with a high-resolution Orbitrap Fusion Lumos Tribrid mass spectrometer demonstrates that the protein is α-N-acetylated. In addition, Vma1 was detected as monomethylated in two different isotopic labeling datasets, but our experiment indicates that it was predominantly acetylated. We posit that Vma1 may be monomethylated at a nominal level relative to acetylation, and hence it is difficult to identify these low frequency methylated peptides. The lack of prevalent α-N-terminal methylation in these two proteins with non-canonical motifs implies that α-N-terminal methylation is a relatively rare modification for these proteins. Due to possible ambiguity of identifying α-N-terminal methylation from α-N-terminal acetylation as shown in Ssa3, we excluded any nonlabelled trimethylation hits from our downstream bioinformatics analysis for the sake of accuracy. Future proteomic assessment of α-N-terminal methylation representation and function would be enhanced by combining N-terminal enrichment techniques with isotopic labeling methods or depleting the α-N-terminal acetylated proteins.

The source of the non-canonical methylation events is unclear. N-terminal methyltransferases that prefer the canonical motifs may be responsible for a rare level of background methylation of proteins with non-canonical motifs. Another possibility is that an additional methyltransferase enzyme is performing this activity. In prokaryotes, the function of N-terminal methylation is found to be relatively promiscuous and does not follow canonical motif recognition.^2^ PrmA is the nonessential N-terminal methyltransferase in both *E.coli* and *T. thermophilus* and is reported to modify three amino acids on Rpl11, including N-terminal alanine, Lys3 and Lys39 while Rpl11.^57–59^ Notably, there is minimal sequence similarity between PrmA and the eukaryotic N-terminal transferases, NTMT1/2 and Tae1.^4^ Our studies raise the intriguing possibility that a functionally corresponding eukaryotic methyltransferase contributes to these general methylation events detected by our study. The source of these methylation events could be revealed by proteomic studies of genetically modified cell lines or yeast strains (such as methyltransferase gene deletions or functionally compromised strains). We expect non-canonical methylation is enzymatically driven because various isotopic labels resulted in detectable methylation. Another possibility is that nonenzymatic methylation of proteins occurs by reaction with SAM.^60^ We also note that bioinformatic analysis of non-canonical motif methylated proteins, including GO term analysis did not show enrichment in any category (data not shown). An additional question is if these methylation events have direct biological roles, or are possibly present at a very nominal and persistent level, or are sporadic events. Although the source of non-canonical methylation is unclear, our studies have shown it to be persistent and widespread across the proteome, indicating that this type of methylation warrants further investigation.

The reproducibility of hits across different samples is low. No protein hits are shared by the three yeast datasets and two proteins are reproduced in two human datasets (Figure 6). The low reproducibility in both humans and yeast is likely because of the limited number of protein hits, exclusion of the majority of trimethylated hits from bioinformatics analysis and insufficiency in the detection of the α-N-terminome. Besides, up to 2-3 peptide occurrences were identified for the same PTM for most protein hits (Table S5). Also of note is that we observed different peptide cleavage forms encompassing the same modification, which increases the confidence in a correct assignment. The low reproducibility across datasets demonstrates a need for the optimized design of experimental approaches and more comprehensive dataset searching strategies to improve α-methylation detection. Within the Chemical Labeling dataset, where four sample preparation methods (two proteases and two chemical labeling methods) were individually used to analyze the α-N-terminal methylome in the same sample, 33 proteins are found in all four subsets and 289 proteins were shared by two or more subsets (Figure 6). Moving forward, we demonstrate the utility of building an automated searching platform to repurposing appropriate datasets and provide the path to generating a compendium of potentially α-N-terminal methylated proteins.

**Figure 6.**
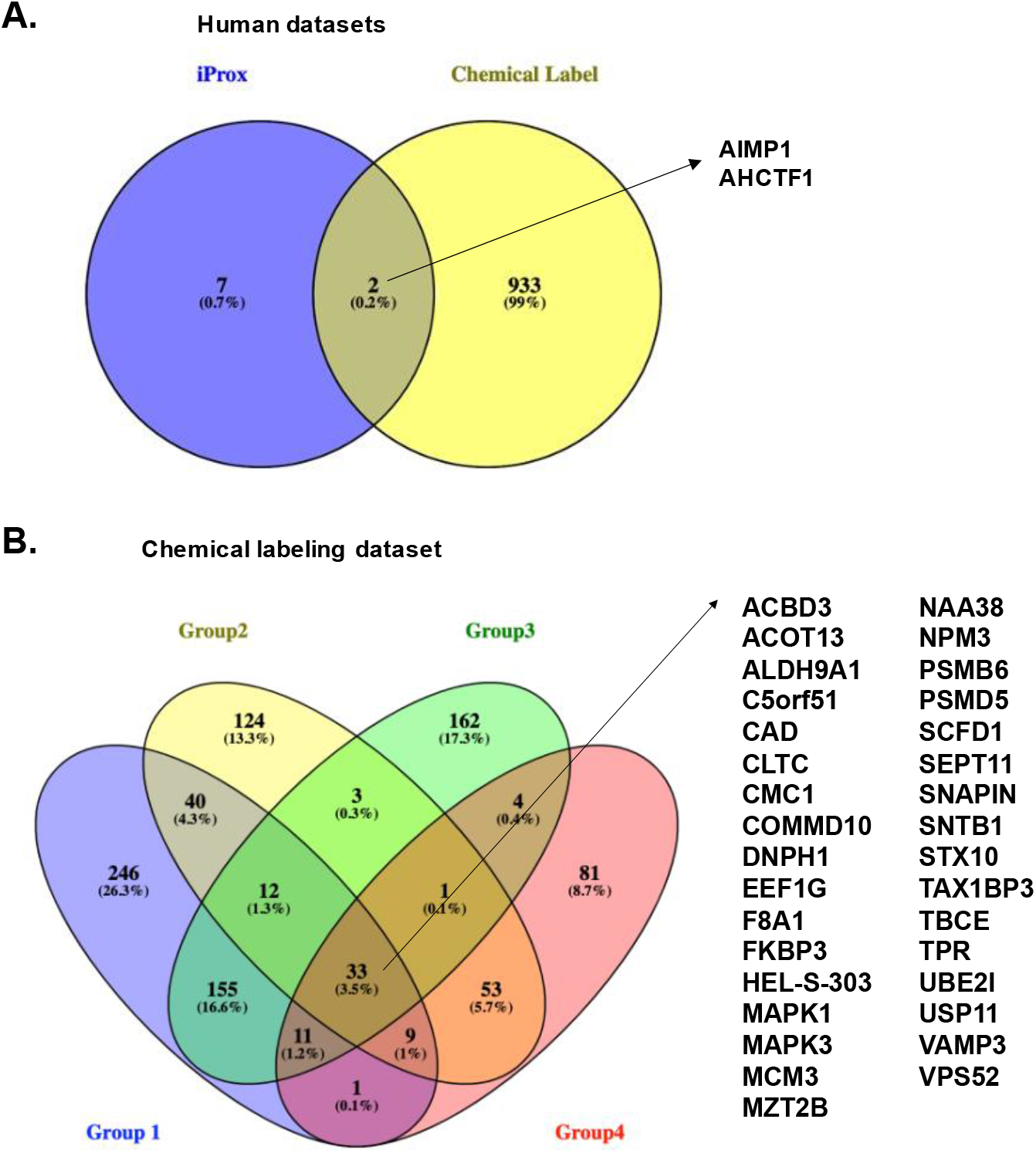
Venn diagram demonstrating the number of potential methylated proteins and the overlap from human datasets. (A) The human datasets yielded 2 overlapping proteins. (B) The Chemical Labeling subsets yielded 33 overlapping proteins present in all four subsets and 289 proteins that appeared in two or more subsets. Yeast datasets don’t have overlapping hits.

To avoid false positive hits and overinterpretation, extra attention should be paid to sample preparation techniques, resolution of mass spectrometry instruments, and the ambiguity caused by α-N-terminal acetylation. In some N-terminal enrichment techniques, dimethylation reagent is used to block the free α-N-terminus so that the free α-N-terminus of the internal peptides can be selectively removed. Only α-N-trimethylation would be meaningful in these datasets, while the α-N-monomethylation and α-N-dimethylation could be chemically formed during sample preparation. The sensitivity of an instrument is also another crucial factor. It is essential that high-resolution mass spectrometry instruments are used that are capable of differentiating a small mass difference, such as the Q-Exactive or higher resolution LTQ instruments. In addition, instruments may not be run at a high resolution for scanning speed reasons, and thus they cannot accurately differentiate acetylation from trimethylation. Another concern of generally searching for any specific protein modifications at the proteome level is the noise signal ratio. Less protein representation, protein contamination, insufficient MS/MS fractionation, and the vast number of peptides analyzed might lead to false positive results. To further optimize global N-terminal methylome investigations, two methodologies might yield higher quality and quantity: (i) Combination of SILAC and N-terminal enrichment methods and (ii) Negative selection method removing internal peptides and N-terminal acetylated peptides. Supplementary methods may be necessary to verify potentially modified proteins and accurate modification site assignment, such as probing the protein of interest under either physiological levels or overexpression followed by mass spectrometry.

The vast number of datasets generated for various purposes such as protein identification, modification detection and investigating proteome perturbances are openly accessible on proteome consortiums sites such as ProteomeXchange (http://www.proteomexchange.org)^24^ and databases such as iProx (https://www.iprox.org/)^25^ and PRIDE (https://www.ebi.ac.uk/pride/)^23^. Reanalysis of public datasets has been reported previously to discover novel protein-encoding genes^61^, finding missing human proteins^62^, identify novel PTMs^63^ and reveal PTM sites ^64^. One of the significant barriers for reusing proteomic datasets is the complexity of MS data. Both the sample preparation procedure and the mass spectrometry technique used in the datasets need to be closely examined to find suitable ones for repurposing. Efforts to mitigate these factors are underway including integrating experimental metadata and reanalysis tools in a platform such as the Reanalysis of Data User (ReDU) interface, which focuses on chemicals and metabolite proteomics data.^65^ There are abundant MS datasets that were designed for probing modified protein N-termini and repurposing those datasets for N-terminal methylation events will shorten the research cycle and reduce the expense of proteomic studies. By repurposing the current datasets, potentially α-N-terminal methylated proteins are generated with confidence depending on the type of instrument, the complexity of the sample, protein abundance and the accessibility of N-termini. Notably, no previous reports have examined the N-terminal methylome by employing the reanalysis of public datasets. Of course, it would be necessary to verify α-N-terminal methylation events with purified proteins or orthologous methods to investigate specific proteins further. The establishment of the prevalence of this α-N-terminal modification is a crucial step to build a compendium of methylated proteins and discover the functions of this cryptic modification.

## Supporting Information

Table S1. 45 canonical N-terminal motif-containing proteins in yeast.

Table S2. FDR information of each dataset.

Table S3. b/y ions of MS2 spectrum of α-N-methylated peptides.

Table S4. Mascot search results with alternative K/R methylation sites.

Table S5. Protein hits from each repurposed dataset.

Table S6. Protein hits containing canonical motifs.

Table S7. List of sequences used for WebLogo and iceLogo analysis.

Figure S1. MS/MS spectra for all hits from three repurposed yeast datasets.

Figure S2. WebLogos and iceLogos for repurposed datasets.

Figure S3. N-Ac and N-Me distribution plots (iMet retained hits included).

Figure S4. Western blot of purified proteins.

Figure S5. Intact mass spectrometry of Vma1.

Figure S6. Protein coverage map of Vma1.

Figure S7. Methods description and MS analysis of Ssa3.

Figure S8. MS2 spectra of monomethylated and formylated Hsp31.

Table S6, Figure S2-8 are integrated into a single Word file. Table S1-5, Table S7, Figure S1 are individual Excel files. Figure S1 is attached as a separate individual PDF file.

Mascot output files for the four repurposed datasets are compressed into the “Supporting information raw search output.zip” file.

## Acknowledgment

Analysis of Ssa3, Vma1 and Hsp31 was performed at the Indiana University School of Medicine (IUSM) proteomics core, Purdue Proteomics Facility and BGI Americas Corp. We gratefully acknowledge the Purdue University Center for Cancer Research (NIH P30CA023168) for supporting the Purdue Proteomics Facility and Dr. Uma Aryal for assistance with the MS analysis. Work in the IUSM Proteomics Core was supported, in part, with support from the Indiana Clinical and Translational Sciences Institute which is funded by Award Number UL1TR002529 from the National Institutes of Health, National Center for Advancing Translational Sciences, Clinical and Translational Sciences Award. We thank Dr. Amber Mosley and Dr. Emma Doud at the IUSM Proteomics Core for assistance with the mass spectrometry analysis. Acquisition of the IUSM Proteomics core instrumentation used for this project, the Orbitrap Fusion Lumos, was provided by the Indiana University Precision Health Initiative.

All data were deposited to the PRIDE Archive (http://www.ebi.ac.uk/pride/archive/) via the PRIDE partner repository and are publicly accessible with the data set identifier PXD022833.

## Notes

The authors declare no competing financial interest.

