## Supporting information for "Discovering the N-terminal Methylome by Repurposing of Proteomic Datasets"

### Supporting information list.


| | |
|---|---|
| Table S1. 45 canonical N-terminal motif-containing proteins in yeast. | Excel file |
| Table S2. FDR information of each dataset. | Excel file |
| Table S3. b/y ions of MS2 spectra of α-N-methylated peptides. | Excel file |
| Table S4. Mascot search results with alternative K/R methylation sites. | Excel file |
| Table S5. Protein hits from each repurposed dataset. | Excel file |
| Table S6. Protein hits containing canonical motifs. | Page S2-S3 |
| Table S7. List of sequences used for WebLogo and iceLogo analysis. | Excel file |
| Figure S1. MS/MS spectra for all hits from three repurposed yeast datasets. | Separate PDF |
| Figure S2. WebLogos and iceLogo for repurposed datasets. | Page S4-S5 |
| Figure S3. N-Ac and N-Me distribution plots (iMet retained hits included). | Page S6 |
| Figure S4. Western blot of purified proteins. | Page S7-S8 |
| Figure S5. Intact mass spectrometry of Vma1. | Page S9 |
| Figure S6. Protein coverage map of Vma1. | Page S10 |
| Figure S7. Methods description and MS analysis of Ssa3. | Page S11-S12 |
| Figure S8. MS2 spectra of monomethylated and formylated Hsp31. | Page S13-S16 |


Table S6, Figure S2-S8 are integrated into this single Word file.

Table S1-S5 and Table S7 are in a single Excel file.

Figure S1 is attached as a separate individual PDF file.

Mascot output files for the four repurposed datasets are compressed into the "Supporting information raw search output.zip" file.

**Table S6: Canonical motif-containing hits. (Yellow labeled rows have previously been reported in the literature).**

Dataset 1. 2 Hits in the CharFRADIC dataset containing X-P-K/R motif (Baker's yeast) with peptide score >20.

| Name | N-terminal sequence | Description |
|---|---|---|
| **Rpl12ab** | PPKFDPNEVKYLYLR (dimethyl)* | 60S ribosomal protein L12ab |
| **Tef1** | GKEKSHINVVVIGHVDSGKSTTT (Trimethyl) | Elongation factor 1-alpha |

*Rpl12ab was detected with 5 methyl groups within the first 5 amino acid positions in our hand with limited MS2 fragmentation. The methyl groups localization is most likely to be dimethylation on proline$_1$ and trimethylation on lysine$_3$.

Dataset 2. Hits in the iProx dataset containing X-P-K/R motif (Baker's yeast and Hela cell). None found.

Dataset 3. 5 hits in the chemical labeling dataset containing X-P-K/R motif (HEK293T cell) with peptide score>20.

| Group number and conditions | Name | N-terminal sequence | Description |
|---|---|---|---|
| 1.D6-acetylation +trypsin digestion | **XPNPEP1** | PPKVTSELLI (methyl) | Xaa-Pro aminopeptidase |
| 2.D6-acetylation +GluC digestion | **CMTM2** | APKAAKGAKPE (trimethyl & dimethyl) | CKLF-like MARVEL transmembrane domain-containing protein |
| 2.D6-acetylation +GluC digestion | **RPL23A** | APKAKKEAPAPPKAE (trimethyl & dimethyl) | 60S ribosomal protein L23a |
| 2.D6-acetylation +GluC digestion | **SET** | APKRQSPLPPQKKKPRPPPALGPEE (trimethyl) | SET |
| 2.D6-acetylation +GluC digestion | **ZC3H15** | PPKKQAQAGGSKKAE (dimethyl) | Zinc finger CCCH domain-containing protein 15 |

There are four protein hits from the chemical labeling dataset containing canonical N-terminal motif X-P-K/R with scores close to the cut-off value:

| Group number and conditions | Name | N-terminal sequence and peptide scores | Description |
|---|---|---|---|
| 3. Propionlyation +Trypsin | **RGS11** | PPRWLPPR (trimethyl, peptide score 19.22) | Alternative protein RGS11 |
| 3. Propionlyation +Trypsin | **PCSK6** | PPRAPPAPGPR (trimethyl, peptide score 19.91) | Proprotein convertase subtilisin/kexin type 6 |
| 2. D6-acetylation +GluC digestion | **MORF4L1** | APKQDPKPKFQE (dimethyl and trimethyl, peptide scores are 11, 18.27 and 16.27) | Mortality factor 4-like protein 1 |
| 2. D6-acetylation +GluC digestion | **TOMM34** | APKFPDSVE (monomethyl and trimethyl, peptide scores are 10.04 and 12.61, respectively) | Mitochondrial import receptor subunit TOM34 |

Dataset 4. One hit in the MISL dataset containing the X-P-K/R motif (Baker's yeast) with peptide score>20.

| Name | N-terminal sequence | Description |
|---|---|---|
| **Rps25a/b** | PPKQQLSKAAK (monomethyl) | 40S ribosomal protein S25-a/b |

There are four protein hits with peptide score <20 as follows:

| Name | N-terminal sequence | Description |
|---|---|---|
| **Rpl12ab** | PPKFDPNEVK (monomethyl & dimethyl) | 60S ribosomal protein L12ab |
| **Rpt1** | PPKEDWEKYK (dimethyl) | 26S proteasome regulatory subunit 7 homolog |
| **Ola1** | PPKKQVEEK (monomethyl) | Obg-like ATPase 1 |
| **Hsp31** | APKKVLLALTSYNDVFYSDGAK (dimethyl & trimethyl) | Glutathione-independent glyoxalase HSP31 |

In total, 8 protein hits identified from 4 datasets contain canonical N-terminal motifs X-P-K/R with a score >20. Five of them were identified from the HEK cell line sample and two out of the five were reported previously. One protein was identified from yeast, and it was previously reported in the literature. There are 8 protein hits identified from 4 datasets with less confidence (peptide score<20).

**Figure S2: WebLogos and iceLogo.**

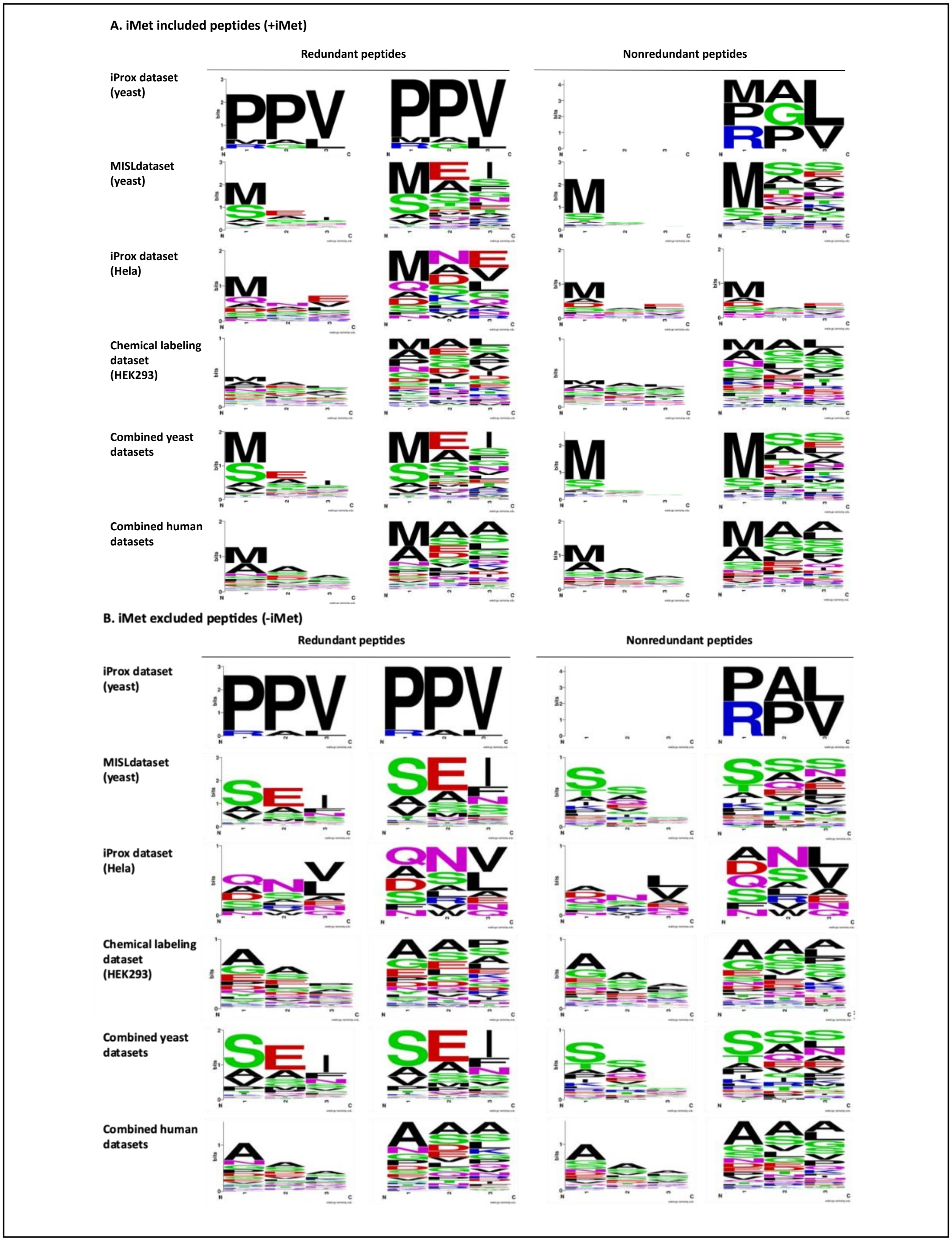

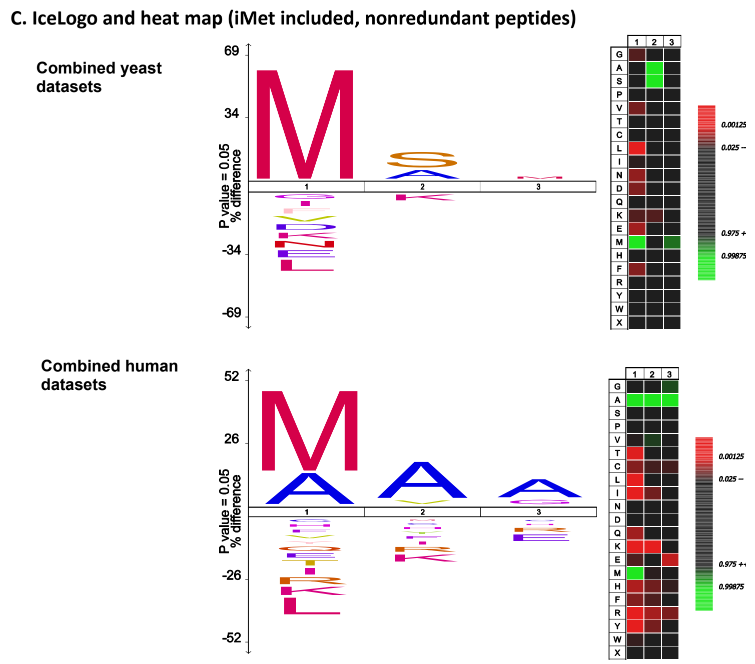


Figure S2. (A) and (B). Weblogo for all repurposed datasets. Four Web logos are generated for each individual datasets or combined datasets depending on whether the iMet is cleaved or peptide list is redundant. iProx yeast dataset lacks sequence logos for nonredundant peptides since too few nonredundant N-terminal peptides were identified to generate a sequence logo by WebLogo. (C) iceLogo and heatmap for nonredundant peptide lists of yeast and human datasets including iMet retained peptides.

Several factors might affect the sequence logo pattern, including iMet retention and peptide redundancy within the identified peptide lists. iMet is encoded by the starting codon of the mRNA but is usually removed during protein maturation by the methionine aminopeptidase.[1] More than 30% of the N-terminal peptides retain the iMet in each repurposed dataset except the ChaFRADIC dataset and iProx yeast subset. Specifically, the iMet retention rate was 47% for the MISL dataset, 47% for the iProx human dataset, and 34% for the Chemical Labeling dataset. Three iMet retained peptides were detected in the ChaFRADIC dataset and only one iMet retained peptide was detected in the iProx yeast subset.

We applied WebLogo and iceLogo to analyze the sequence conserveness of a-N-methylated proteins in yeast and humans. Both logo programs reflect the sequence pattern of a list of peptides based on different algorithms and are informative in identifying sequence conservation. WebLogo utilizes Shannon's theory and graphically displays the sequence pattern based on the frequency of amino acids from the aligned peptide sequences.[2] iceLogo builds on probability theory and finds significantly over-represented/under-represented amino acids by comparing the experimental set with a reference background set.[3] We used the Swiss-Prot yeast proteome and human proteome as reference sets for iceLogo analysis.

Individual WebLogos for each yeast and humans were generated for peptides. WebLogos for iMet cleaved peptide list or iMet retained peptide list of each dataset were correspondingly analyzed (Figure S2A and B). Redundancy and overrepresentation of identified peptides in the list could alter the conservation patterns, so we also investigated redundant and nonredundant lists for each dataset (Figure S2A and B). We found that retaining redundant peptides had a limited effect on the site frequency. The WebLogo for iMet retained peptides had methionine as the most frequent amino acid at the first position. The conserved pattern of the list of nonredundant iMet included peptides were predicted by iceLogo and heatmap (Figure S2C). It shows iMet is also significantly enriched for N-terminal methylation. iceLogos of nonredundant iMet excluded peptides were shown in Figure 1 and Figure 2.

**Figure S3. N-Ac and N-Me distribution plots (iMet retained hits included).**

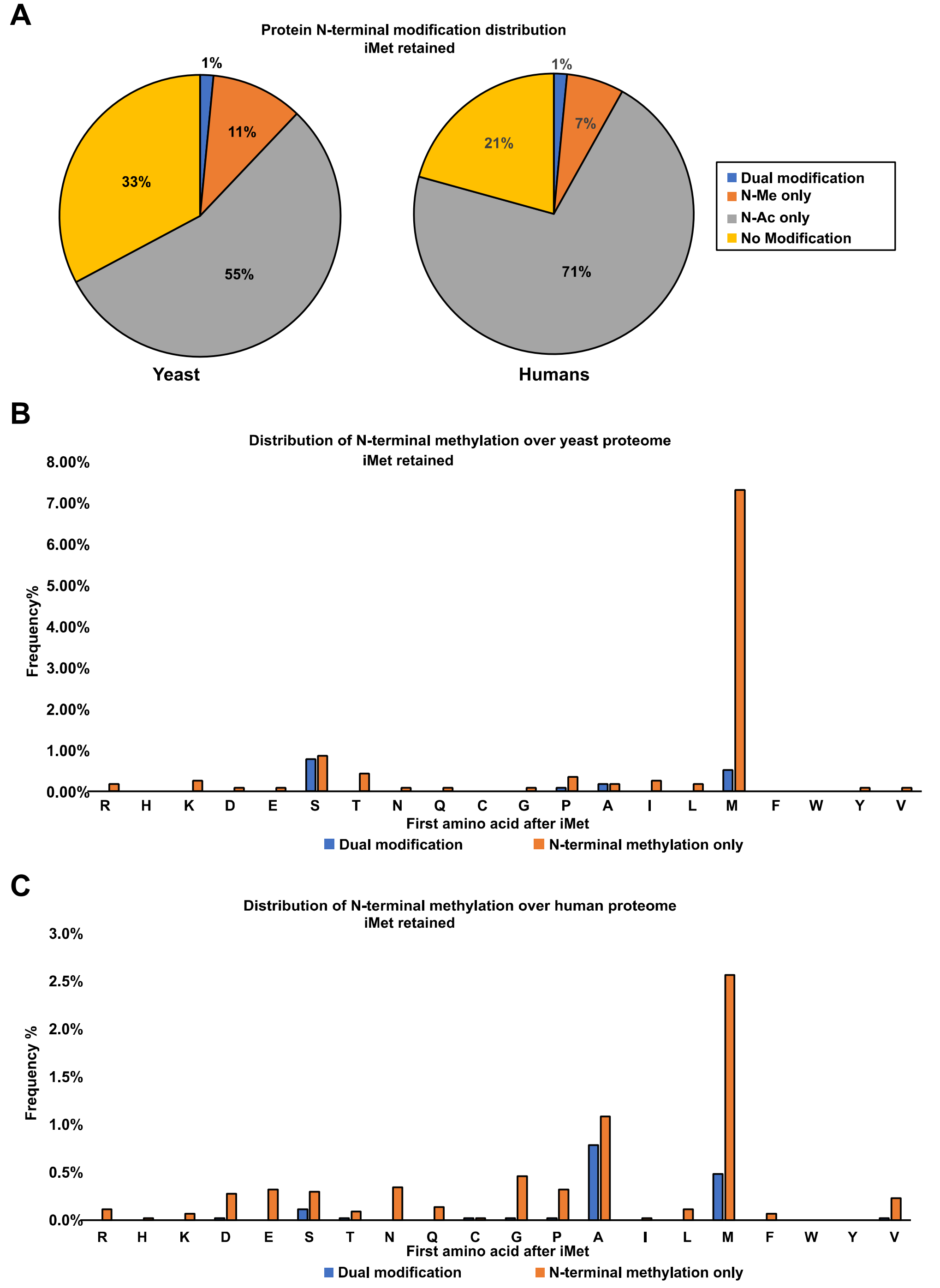


Figure S3. The frequency of proteins found to have only α-N-terminal methylation or both modifications are plotted according to the type of amino acid. Both iMet retained proteins and iMet cleaved proteins are used for this graph. (A) Pie charts show the proportion of yeast or human proteome found to be α-N-terminal methylated or α-N-terminal acetylated. (B) The frequency plot for yeast datasets, including MISL and iProx datasets. (C) The frequency plot for yeast datasets, including Chemical Labeling and iProx datasets. The frequency of acetylation was not depicted in the bar graphs because high levels of acetylation for several amino acids results in a scale that obscures the visualization of lower frequency methylation levels.

1165 unique proteins are identified from iProx yeast datasets and MISL datasets, in which iMet retained proteins are included. 141 (12%) proteins are found to be heavy isotope methylated. 18 (1%) out of 141 are under dual modification of α-N-terminal methylation and α-N-terminal acetylation, which the rest 123 (11%) proteins are only found to be α-N-terminal methylated. Around 642 (55%) proteins are found to be α-N-terminal acetylated and 382 (33%) proteins were not modified by either modification. (Figure S3A)

4320 unique proteins were identified from iProx Hela datasets and chemical labeling datasets, in which iMet retained proteins are included. Heavy isotope methylated proteins from iProx Hela datasets and mono/dimethylated proteins from Chemical labeling datasets constitute 351 (8%) unique proteins. 66 (1%) out of 351 are under dual modification of α-N-terminal methylation and α-N-terminal acetylation which the rest 285 (7%) proteins are only found to be α-N-terminal methylated. Around 3074 (71%) proteins are found to be α-N-terminal acetylated and 895 (21%) proteins did not carry either modification. (Figure S3A)

N-Ac and N-Me distribution on 1st amino acid of proteins in yeast and human is delineated in Figure S2B and Figure S2C, respectively. iMet retained proteins are included in this figure compared to Figure 3 used in the manuscript, and the proportions and frequencies of modifications did not change appreciably except for methionine (A).

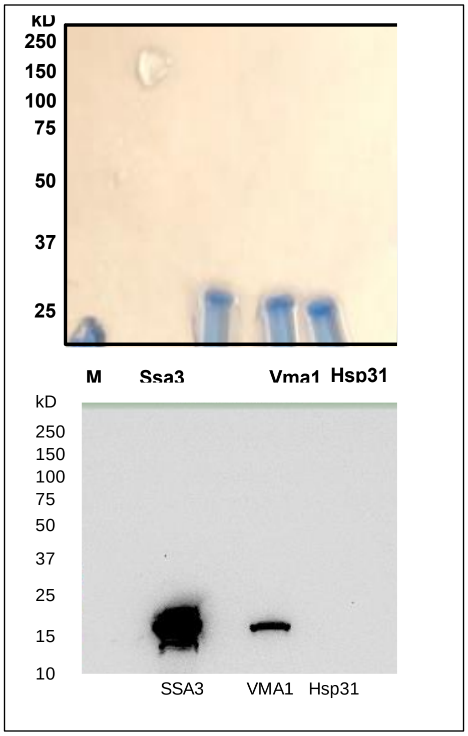


**Figure S4: Western blot analysis of purified proteins.** C-terminal MORF tagged Ssa3, Vma1 and Hsp31 were purified from yeast using IgG agarose bead and eluted with HRV-3C protease (Acro Biosystems). The released protein contained a C-terminal HA-6His tandem tag. ~10 µg protein was used for SDS-PAGE and subsequent chemiluminescent western blot analysis. The membrane was blocked with 5% milk and incubated with an anti-HA antibody (1:1000, BioVision) for 1 h. The stain was washed with PBS three times and followed by an anti-mouse antibody (1:10,000) incubation. The chemiluminescence image was acquired by a ChemiDoc imaging system (Bio-Rad).

**Figure S5. Intact mass spectrometry of Vma1.**

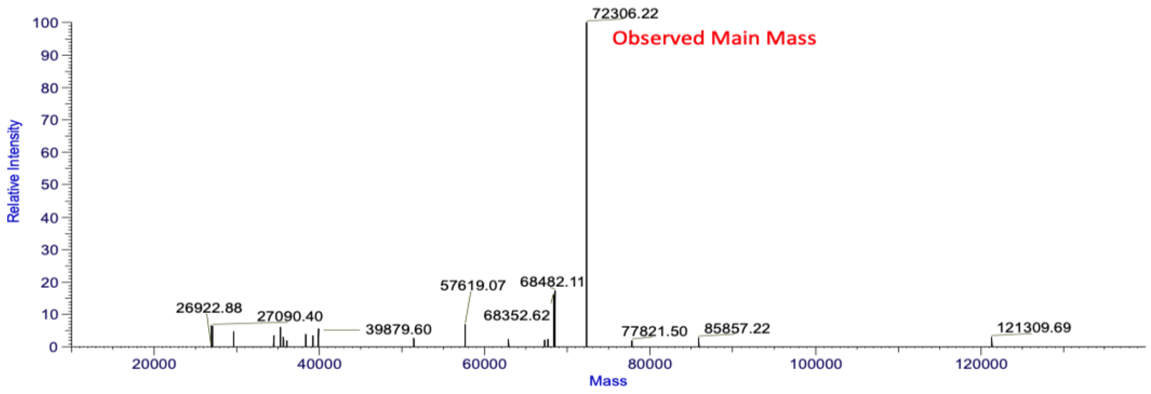


Figure S5. The intact protein spectrometry of Vma1 has a major peak at 72306.22 Da. to the calculated mass of the protein if the intein domain is autoprocessed and excised from the full-length protein.

**Figure S6. Protein coverage map of Vma1.**

Each identified peptide was mapped to the protein sequence of Vma1 (Gold – hatch boxes). No peptides were identified that corresponded to an area (AA 284-737 – black hatched boxes) that contained the Vma1 intein sequence, suggesting that the intein is indeed excised from the protein. Color boxes under the Vma1 sequence depict the percent of peptide recovery. Note that the Vma1 sequence also includes the remaining MORF affinity tag at the C-terminus, which has limited peptide coverage.

| Protein | # of MS Peaks | MS Peak Area | Sequence Coverage |
|---|---|---|---|
| Vma1 | 487 | 49.4% | 53.7% |

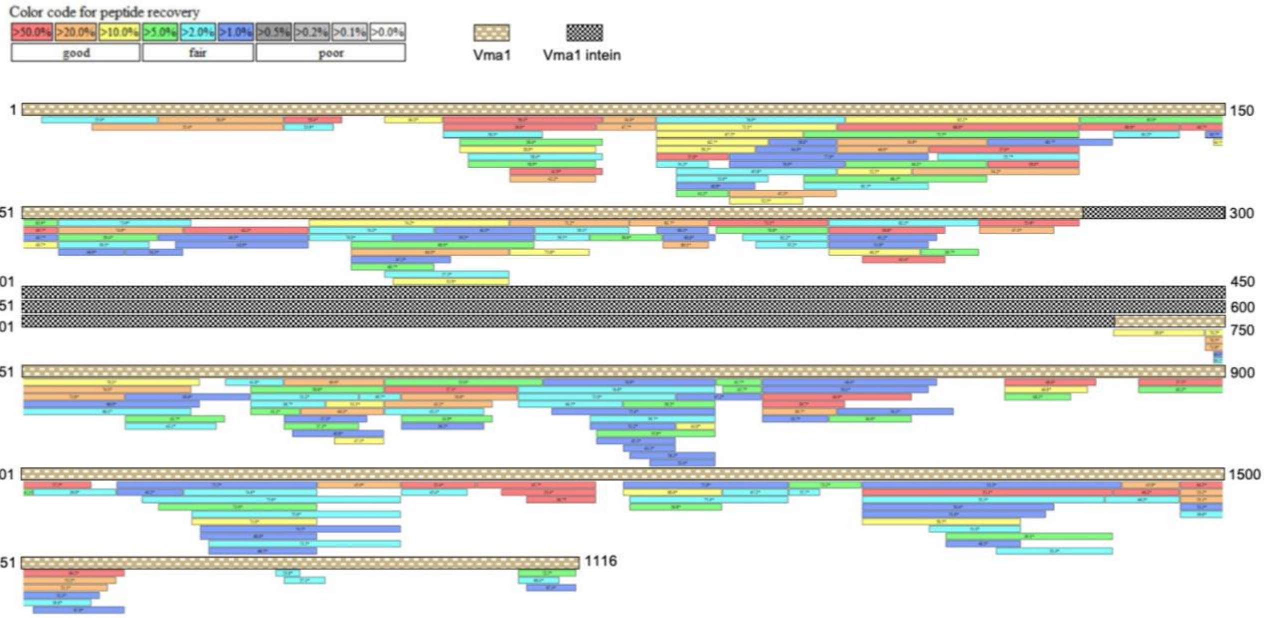

**Figure S7. Method description and MS analysis of Ssa3.**

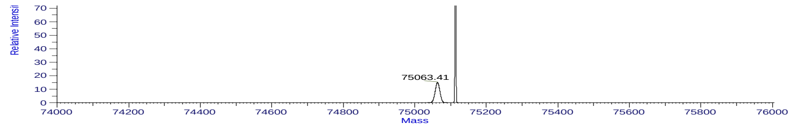


B

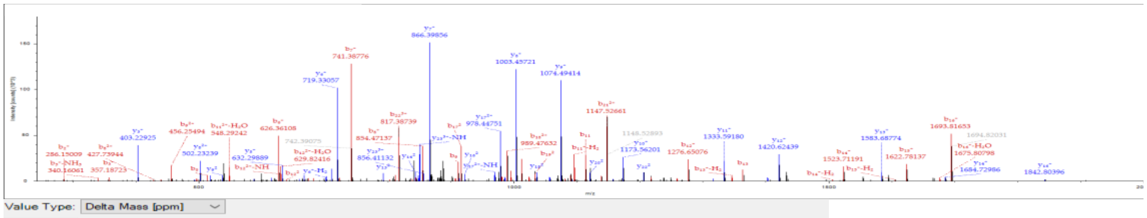

Value Type: Delta Mass [ppm]

Ion Series | Neutral Losses | Precursor Ions | Internal Fragments

| #1 | b+ | b2+ | b3+ | Seq. | y+ | y2+ | y3+ | #2 |
|---|---|---|---|---|---|---|---|---|
| 1 | | | | S-Acetyl | | | | 24 |
| 2 | +3.11 | | | R | | | +4.40 | 23 |
| 3 | +2.42 | | | A | | +2.49 | | 22 |
| 4 | | | | V | | +2.26 | | 21 |
| 5 | +1.70 | | | G | | +1.35 | | 20 |
| 6 | +1.53 | | | I | | +0.39 | | 19 |
| 7 | +1.64 | | | D | | +2.14 | | 18 |
| 8 | +1.96 | +1.68 | | L | | +1.97 | | 17 |
| 9 | +1.81 | -8.87 | | G | +1.94 | +1.06 | | 16 |
| 10 | +2.42 | -0.71 | | T | | | | 15 |
| 11 | +2.43 | | | T | +5.07 | +1.77 | | 14 |
| 12 | +1.91 | +1.38 | | Y | +1.88 | +1.54 | | 13 |
| 13 | +2.18 | -2.16 | | S | +2.11 | +1.65 | | 12 |
| 14 | +2.60 | | | C-Carbami... | +2.67 | +1.72 | | 11 |
| 15 | +1.79 | +6.19 | | V | +2.30 | +2.99 | | 10 |
| 16 | +2.87 | | | A | +2.01 | +1.05 | | 9 |
| 17 | | +1.80 | | H | +1.97 | +1.67 | | 8 |
| 18 | | +1.70 | | F | +1.98 | | | 7 |
| 19 | | +1.17 | | S | +1.79 | | | 6 |
| 20 | | +1.77 | | N | +1.49 | | | 5 |
| 21 | | +2.04 | | D | +0.72 | | | 4 |
| 22 | | | +1.68 | R | +1.76 | | | 3 |
| 23 | | | +1.51 | V | | | | 2 |
| 24 | | | | E | | | | 1 |

Figure S7 (A) Intact protein analysis of Ssa3. The major peak in the intact protein spectrum is 75113.59Da. (B) 95 PSMs with N term Ac, C14 Carbamidomethyl were detected for Ssa3. A typical matched peptide of Ssa3 and identified MS/MS fragmentation is shown.

### Methods for Ssa3 MS analysis

### Intact protein analysis

The intact protein analysis was performed in Q-Exactive HFX and resolved by BioPharma Finder. Ssa3 was cleaved from IgG agarose bead and had a residual tag sequence at the protein C-terminus resulting in a calculated protein mass of 75072.84 Da. The prominent peak in the intact protein spectrum is 75113.59 Da, which is 40.75 Da more than calculated. This mass shift is close to trimethylation or acetylation.

### LC/MS Materials and methods

Sample preparation, mass spectrometry analysis, bioinformatics and data evaluation were performed in collaboration with the Proteomics Core Facility at the Indiana University School of Medicine (IUSM). Methods described below, in brief, were adaptations from literature reports and vendor-provided protocols. 8 M urea was added to approximately 4 ug purified protein, reduced with 5 mM TCEP at room temperature for 30 min, and then alkylated with 10 mM chloroacetamide for 30 min in the dark at room temperature. Samples were diluted to 0.5 M Urea with 50 mM ammonium carbonate, pH 7.8, and digestions were performed using GluC (Sigma Aldrich #11420399001) at a 1:2 protease to substrate ratio, overnight at 25 °C. The reaction was quenched with 0.5 % TFA, and samples were desalted on 50 mg SepPak columns (Waters). Samples were dried down by a speed vacuum.

**LC/MS method**

Samples were analyzed using a 25 cm EasySpray (Thermo Fisher Scientific cat. num. 802A rev 2) column on an Easy nano1200 LC and Lumos Fusion orbitrap mass spectrometer (both Thermo Fisher Scientific). Solvent B was increased from 5%-35% over 55 min, to 85% over 3 min, and back to 5% over 2 min (Solvent A: water, 0.1% formic acid; Solvent B: 80% acetonitrile, 0.1% formic acid). A data-dependent top speed (3 sec cycle time) acquisition method was used with a MS scan range of 400-1500 m/z, orbitrap resolution of 120,000, standard AGC target, and auto maximum injection time (IT). MS2 filters of 2.5e4 intensity threshold, charge states of 2-7, and dynamic exclusion of 60 sec were applied. MS2 settings were quadrupole isolation window of 1 m/z, fixed normalized CID collision energy of 35%, standard AGC target, dynamic maximum IT, and Orbitrap resolution of 60,000.

**Data Analysis**

Data analysis, including de novo and database searches were performed using PEAKS X plus software (Bioinformatics Solutions)[4], with Orbi-Orbi parameters (10 ppm precursor and 0.02 Da fragment tolerance), including the following variable modifications: carbamidomethylation (C), phosphorylation (S, T, Y), Protein N-terminal Acetylation, Protein N-terminal trimethylation, Protein N-terminal demethylation, Protein N-terminal methylation, trimethylation (K), dimethylation (K, R), methylation (K, R). The search database included Uniprot *Saccharomyces cerevisiae* proteins (downloaded 021917), common contaminants, and including the sequence of Ssa3 without the N-terminal methionine and with C-terminal tags. Results were filtered for peptide FDR of ≥ 1%, A-Score≥ 12, mutation ion intensity ≥ 1%, and de novo confidence ≥ 80%.[5–7]

### Figure S8. MS2 spectra of monomethylated and formylated Hsp31.

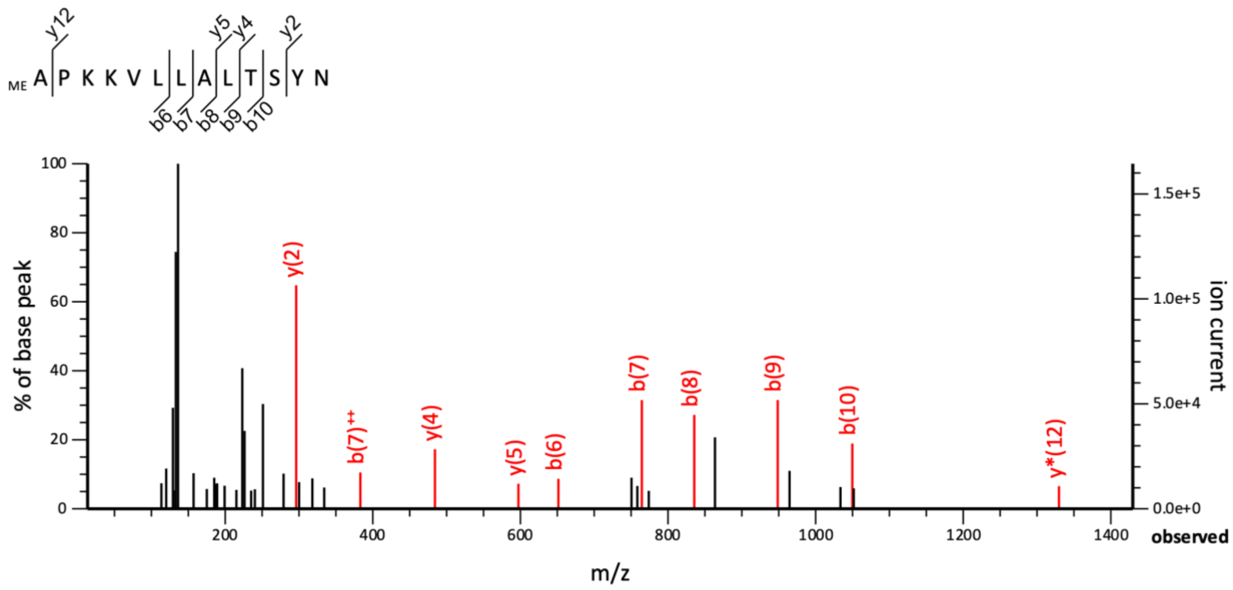


```
Monoisotopic mass of neutral peptide Mr(calc): 1430.8497
Fixed modifications: Carbamidomethyl (C) (apply to specified residues or termini only)
Variable modifications:
  N-term : Methyl (Protein N-term)
Ions Score: 44  Expect: 3.7e-05
Matches : 10/136 fragment ions using 11 most intense peaks   (help)
```

| # | a | a++ | a* | a*++ | b | b++ | b* | b*++ | Seq. | y | y++ | y* | y*++ | # |
|---|---|---|---|---|---|---|---|---|---|---|---|---|---|---|
| 1 | 58.0651 | 29.5362 | | | 86.0600 | 43.5337 | | | A | | | | | 13 |
| 2 | 155.1179 | 78.0626 | | | 183.1128 | 92.0600 | | | P | 1346.8042 | 673.9057 | 1329.7777 | 665.3925 | 12 |
| 3 | 283.2129 | 142.1101 | 266.1863 | 133.5968 | 311.2078 | 156.1075 | 294.1812 | 147.5942 | K | 1249.7514 | 625.3794 | 1232.7249 | 616.8661 | 11 |
| 4 | 411.3078 | 206.1575 | 394.2813 | 197.6443 | 439.3027 | 220.1550 | 422.2762 | 211.6417 | K | 1121.6565 | 561.3319 | 1104.6299 | 552.8186 | 10 |
| 5 | 510.3762 | 255.6918 | 493.3497 | 247.1785 | 538.3711 | 269.6892 | 521.3446 | 261.1759 | V | 993.5615 | 497.2844 | 976.5350 | 488.7711 | 9 |
| 6 | 623.4603 | 312.2338 | 606.4337 | 303.7205 | ***651.4552*** | 326.2312 | 634.4287 | 317.7180 | L | 894.4931 | 447.7502 | 877.4666 | 439.2369 | 8 |
| 7 | 736.5444 | 368.7758 | 719.5178 | 360.2625 | ***764.5393*** | 382.7733 | 747.5127 | 374.2600 | L | 781.4090 | 391.2082 | 764.3825 | 382.6949 | 7 |
| 8 | 807.5815 | 404.2944 | 790.5549 | 395.7811 | ***835.5764*** | 418.2918 | 818.5498 | 409.7786 | A | 668.3250 | 334.6661 | 651.2984 | 326.1529 | 6 |
| 9 | 920.6655 | 460.8364 | 903.6390 | 452.3231 | ***948.6605*** | 474.8339 | 931.6339 | 466.3206 | L | ***597.2879*** | 299.1476 | 580.2613 | 290.6343 | 5 |
| 10 | 1021.7132 | 511.3602 | 1004.6867 | 502.8470 | ***1049.7081*** | 525.3577 | 1032.6816 | 516.8444 | T | ***484.2038*** | 242.6055 | 467.1773 | 234.0923 | 4 |
| 11 | 1108.7452 | 554.8763 | 1091.7187 | 546.3630 | 1136.7402 | 568.8737 | 1119.7136 | 560.3604 | S | ***383.1561*** | 192.0817 | 366.1296 | 183.5684 | 3 |
| 12 | 1271.8086 | 636.4079 | 1254.7820 | 627.8946 | 1299.8035 | 650.4054 | 1282.7769 | 641.8921 | Y | ***296.1241*** | 148.5657 | 279.0975 | 140.0524 | 2 |
| 13 | | | | | | | | | N | 133.0608 | 67.0340 | 116.0342 | 58.5207 | 1 |

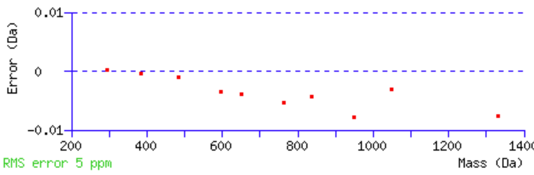


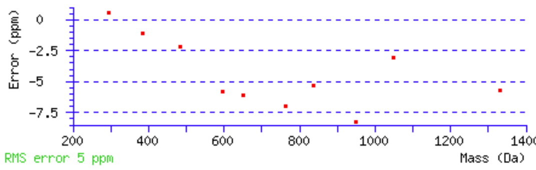


NCBI **BLAST** search of APKKVLLALTSYN
(Parameters: blastp, nr protein database, expect=20000, no filter, PAM30)
Other BLAST web gateways

**All matches to this query**

| Score | Mr(calc) | Delta | Sequence |
|---|---|---|---|
| 44.3 | 1430.8497 | 0.0002 | APKKVLLALTSYN |

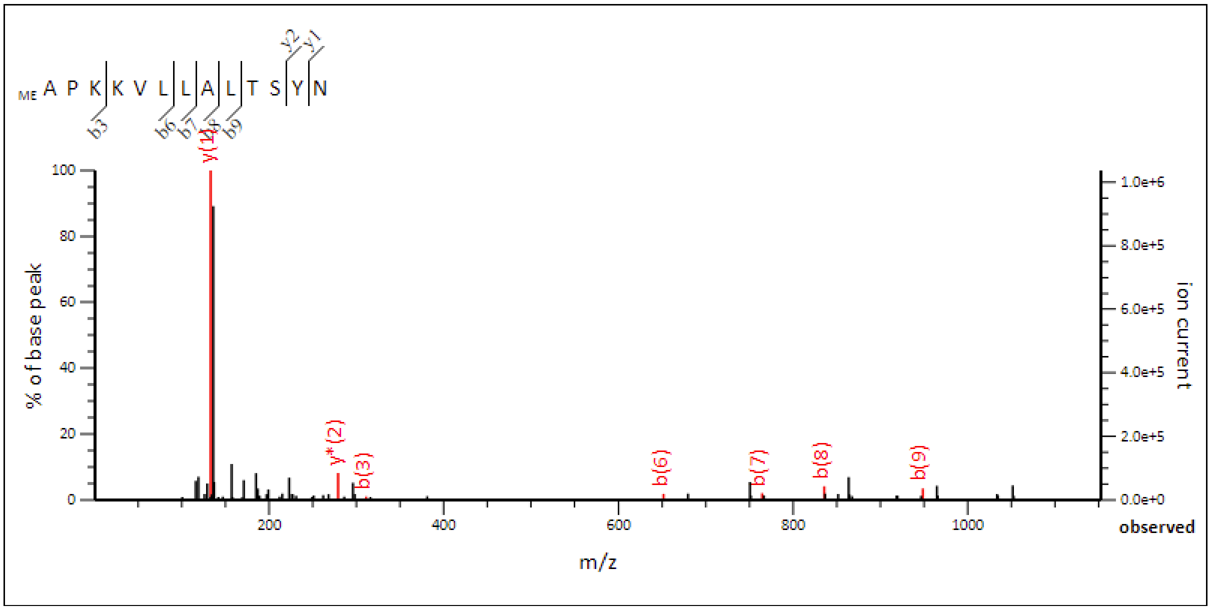


**Monoisotopic mass of neutral peptide Mr(calc):** 1430.8497
**Fixed modifications:** Carbamidomethyl (C) (apply to specified residues or termini only)
**Variable modifications:**
N-term : Methyl (Protein N-term)
**Ions Score:** 23 **Expect:** 0.0053
**Matches :** 7/136 fragment ions using 16 most intense peaks (help)

| # | a | $a^{++}$ | $a^{*}$ | $a^{*++}$ | b | $b^{++}$ | $b^{*}$ | $b^{*++}$ | Seq. | y | $y^{++}$ | $y^{*}$ | $y^{*++}$ | # |
|---|---|---|---|---|---|---|---|---|---|---|---|---|---|---|
| 1 | 58.0651 | 29.5362 | | | 86.0600 | 43.5337 | | | A | | | | | 13 |
| 2 | 155.1179 | 78.0626 | | | 183.1128 | 92.0600 | | | P | 1346.8042 | 673.9057 | 1329.7777 | 665.3925 | 12 |
| 3 | 283.2129 | 142.1101 | 266.1863 | 133.5968 | ***311.2078*** | 156.1075 | 294.1812 | 147.5942 | K | 1249.7514 | 625.3794 | 1232.7249 | 616.8661 | 11 |
| 4 | 411.3078 | 206.1575 | 394.2813 | 197.6443 | 439.3027 | 220.1550 | 422.2762 | 211.6417 | K | 1121.6565 | 561.3319 | 1104.6299 | 552.8186 | 10 |
| 5 | 510.3762 | 255.6918 | 493.3497 | 247.1785 | 538.3711 | 269.6892 | 521.3446 | 261.1759 | V | 993.5615 | 497.2844 | 976.5350 | 488.7711 | 9 |
| 6 | 623.4603 | 312.2338 | 606.4337 | 303.7205 | ***651.4552*** | 326.2312 | 634.4287 | 317.7180 | L | 894.4931 | 447.7502 | 877.4666 | 439.2369 | 8 |
| 7 | 736.5444 | 368.7758 | 719.5178 | 360.2625 | ***764.5393*** | 382.7733 | 747.5127 | 374.2600 | L | 781.4090 | 391.2082 | 764.3825 | 382.6949 | 7 |
| 8 | 807.5815 | 404.2944 | 790.5549 | 395.7811 | ***835.5764*** | 418.2918 | 818.5498 | 409.7786 | A | 668.3250 | 334.6661 | 651.2984 | 326.1529 | 6 |
| 9 | 920.6655 | 460.8364 | 903.6390 | 452.3231 | ***948.6605*** | 474.8339 | 931.6339 | 466.3206 | L | 597.2879 | 299.1476 | 580.2613 | 290.6343 | 5 |
| 10 | 1021.7132 | 511.3602 | 1004.6867 | 502.8470 | 1049.7081 | 525.3577 | 1032.6816 | 516.8444 | T | 484.2038 | 242.6055 | 467.1773 | 234.0923 | 4 |
| 11 | 1108.7452 | 554.8763 | 1091.7187 | 546.3630 | 1136.7402 | 568.8737 | 1119.7136 | 560.3604 | S | 383.1561 | 192.0817 | 366.1296 | 183.5684 | 3 |
| 12 | 1271.8086 | 636.4079 | 1254.7820 | 627.8946 | 1299.8035 | 650.4054 | 1282.7769 | 641.8921 | Y | 296.1241 | 148.5657 | 279.0975 | 140.0524 | 2 |
| 13 | | | | | | | | | N | **133.0608** | 67.0340 | 116.0342 | 58.5207 | 1 |

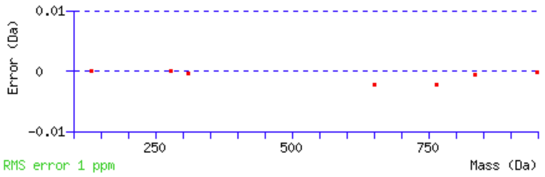


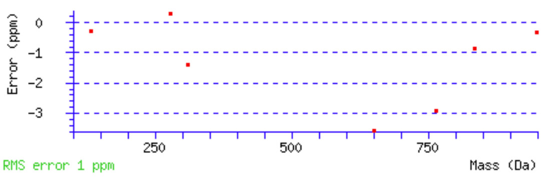


NCBI **BLAST** search of APKKVLLALTSYN
(Parameters: blastp, nr protein database, expect=20000, no filter, PAM30)
Other BLAST web gateways

**All matches to this query**

| Score | Mr(calc) | Delta | Sequence |
|---|---|---|---|
| 22.8 | 1430.8497 | -0.0014 | APKKVLLALTSYN |

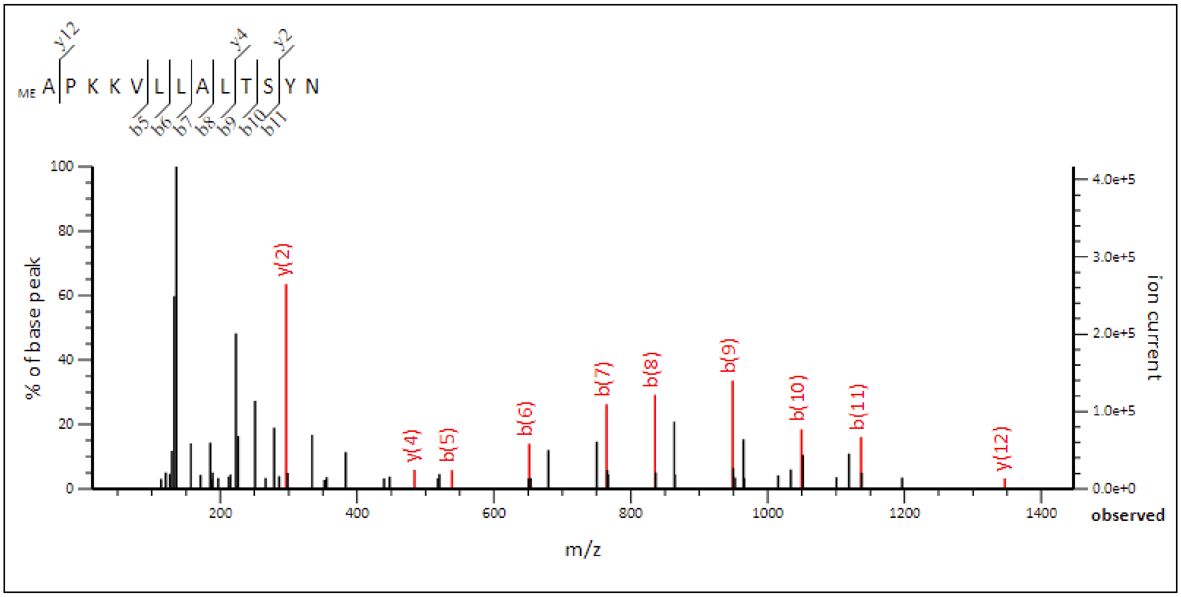


**Monoisotopic mass of neutral peptide Mr(calc):** 1430.8497
**Fixed modifications:** Carbamidomethyl (C) (apply to specified residues or termini only)
**Variable modifications:**
N-term : Methyl (Protein N-term)
**Ions Score:** 60 **Expect:** 1e-06
**Matches :** 10/136 fragment ions using 12 most intense peaks (help)

| # | a | a++ | a* | a*++ | b | b++ | b* | b*++ | Seq. | y | y++ | y* | y*++ | # |
|---|---|---|---|---|---|---|---|---|---|---|---|---|---|---|
| 1 | 58.0651 | 29.5362 | | | 86.0600 | 43.5337 | | | A | | | | | 13 |
| 2 | 155.1179 | 78.0626 | | | 183.1128 | 92.0600 | | | P | **1346.8042** | 673.9057 | 1329.7777 | 665.3925 | 12 |
| 3 | 283.2129 | 142.1101 | 266.1863 | 133.5968 | 311.2078 | 156.1075 | 294.1812 | 147.5942 | K | 1249.7514 | 625.3794 | 1232.7249 | 616.8661 | 11 |
| 4 | 411.3078 | 206.1575 | 394.2813 | 197.6443 | 439.3027 | 220.1550 | 422.2762 | 211.6417 | K | 1121.6565 | 561.3319 | 1104.6299 | 552.8186 | 10 |
| 5 | 510.3762 | 255.6918 | 493.3497 | 247.1785 | ***538.3711*** | 269.6892 | 521.3446 | 261.1759 | V | 993.5615 | 497.2844 | 976.5350 | 488.7711 | 9 |
| 6 | 623.4603 | 312.2338 | 606.4337 | 303.7205 | ***651.4552*** | 326.2312 | 634.4287 | 317.7180 | L | 894.4931 | 447.7502 | 877.4666 | 439.2369 | 8 |
| 7 | 736.5444 | 368.7758 | 719.5178 | 360.2625 | ***764.5393*** | 382.7733 | 747.5127 | 374.2600 | L | 781.4090 | 391.2082 | 764.3825 | 382.6949 | 7 |
| 8 | 807.5815 | 404.2944 | 790.5549 | 395.7811 | ***835.5764*** | 418.2918 | 818.5498 | 409.7786 | A | 668.3250 | 334.6661 | 651.2984 | 326.1529 | 6 |
| 9 | 920.6655 | 460.8364 | 903.6390 | 452.3231 | ***948.6605*** | 474.8339 | 931.6339 | 466.3206 | L | 597.2879 | 299.1476 | 580.2613 | 290.6343 | 5 |
| 10 | 1021.7132 | 511.3602 | 1004.6867 | 502.8470 | ***1049.7081*** | 525.3577 | 1032.6816 | 516.8444 | T | **484.2038** | 242.6055 | 467.1773 | 234.0923 | 4 |
| 11 | 1108.7452 | 554.8763 | 1091.7187 | 546.3630 | ***1136.7402*** | 568.8737 | 1119.7136 | 560.3604 | S | 383.1561 | 192.0817 | 366.1296 | 183.5684 | 3 |
| 12 | 1271.8086 | 636.4079 | 1254.7820 | 627.8946 | 1299.8035 | 650.4054 | 1282.7769 | 641.8921 | Y | **296.1241** | 148.5657 | 279.0975 | 140.0524 | 2 |
| 13 | | | | | | | | | N | 133.0608 | 67.0340 | 116.0342 | 58.5207 | 1 |

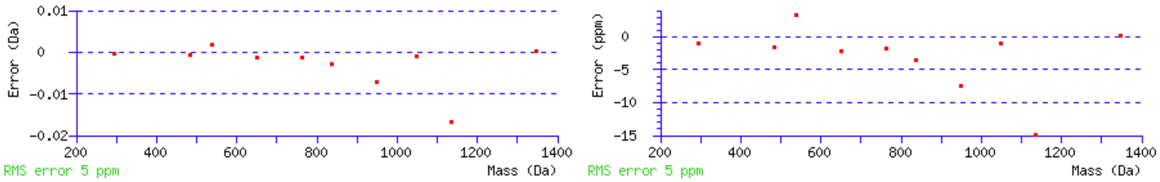


NCBI **BLAST** search of APKKVLLALTSYN
(Parameters: blastp, nr protein database, expect=20000, no filter, PAM30)
Other BLAST web gateways

**All matches to this query**

| Score | Mr(calc) | Delta | Sequence |
|---|---|---|---|
| 59.9 | 1430.8497 | -0.0010 | APKKVLLALTSYN |

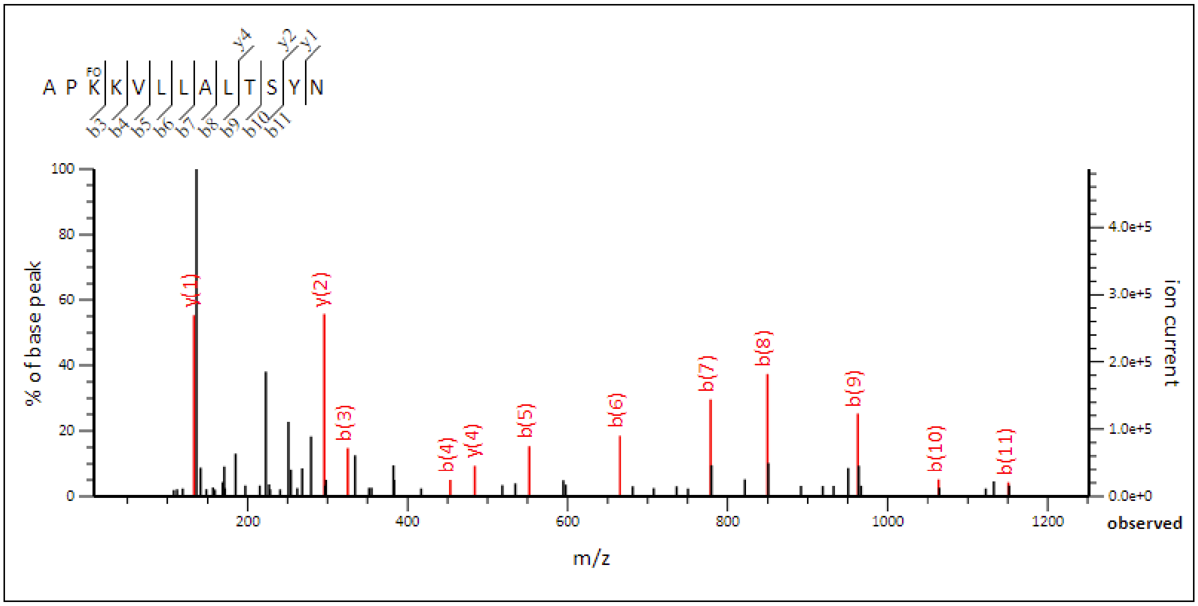


```
Monoisotopic mass of neutral peptide Mr(calc): 1444.8289
Fixed modifications: Carbamidomethyl (C) (apply to specified residues or termini only)
Variable modifications:
  K3 : Formyl (K)
Ions Score: 62  Expect: 9.2e-07
Matches : 12/136 fragment ions using 22 most intense peaks   (help)
```

| # | a | $a^{++}$ | a* | $a^{*++}$ | b | $b^{++}$ | b* | $b^{*++}$ | Seq. | y | $y^{++}$ | y* | $y^{*++}$ | # |
|---|---|---|---|---|---|---|---|---|---|---|---|---|---|---|
| 1 | 44.0495 | 22.5284 | | | 72.0444 | 36.5258 | | | A | | | | | 13 |
| 2 | 141.1022 | 71.0548 | | | 169.0972 | 85.0522 | | | P | 1374.7991 | 687.9032 | 1357.7726 | 679.3899 | 12 |
| 3 | 297.1921 | 149.0997 | 280.1656 | 140.5864 | ***325.1870*** | 163.0972 | 308.1605 | 154.5839 | K | 1277.7464 | 639.3768 | 1260.7198 | 630.8635 | 11 |
| 4 | 425.2871 | 213.1472 | 408.2605 | 204.6339 | ***453.2820*** | 227.1446 | 436.2554 | 218.6314 | K | 1121.6565 | 561.3319 | 1104.6299 | 552.8186 | 10 |
| 5 | 524.3555 | 262.6814 | 507.3289 | 254.1681 | ***552.3504*** | 276.6788 | 535.3239 | 268.1656 | V | 993.5615 | 497.2844 | 976.5350 | 488.7711 | 9 |
| 6 | 637.4396 | 319.2234 | 620.4130 | 310.7101 | ***665.4345*** | 333.2209 | 648.4079 | 324.7076 | L | 894.4931 | 447.7502 | 877.4666 | 439.2369 | 8 |
| 7 | 750.5236 | 375.7654 | 733.4971 | 367.2522 | ***778.5185*** | 389.7629 | 761.4920 | 381.2496 | L | 781.4090 | 391.2082 | 764.3825 | 382.6949 | 7 |
| 8 | 821.5607 | 411.2840 | 804.5342 | 402.7707 | ***849.5557*** | 425.2815 | 832.5291 | 416.7682 | A | 668.3250 | 334.6661 | 651.2984 | 326.1529 | 6 |
| 9 | 934.6448 | 467.8260 | 917.6183 | 459.3128 | ***962.6397*** | 481.8235 | 945.6132 | 473.3102 | L | 597.2879 | 299.1476 | 580.2613 | 290.6343 | 5 |
| 10 | 1035.6925 | 518.3499 | 1018.6659 | 509.8366 | ***1063.6874*** | 532.3473 | 1046.6608 | 523.8341 | T | **484.2038** | 242.6055 | 467.1773 | 234.0923 | 4 |
| 11 | 1122.7245 | 561.8659 | 1105.6980 | 553.3526 | ***1150.7194*** | 575.8633 | 1133.6929 | 567.3501 | S | 383.1561 | 192.0817 | 366.1296 | 183.5684 | 3 |
| 12 | 1285.7878 | 643.3976 | 1268.7613 | 634.8843 | 1313.7828 | 657.3950 | 1296.7562 | 648.8817 | Y | **296.1241** | 148.5657 | 279.0975 | 140.0524 | 2 |
| 13 | | | | | | | | | N | **133.0608** | 67.0340 | 116.0342 | 58.5207 | 1 |

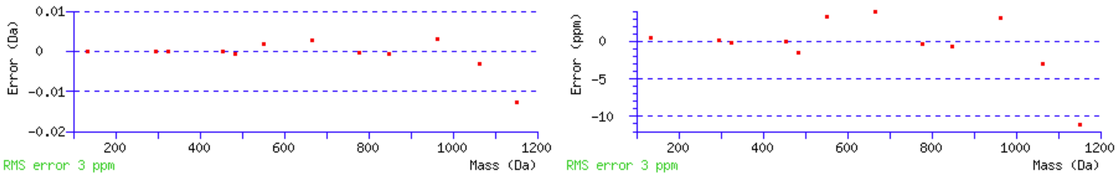


NCBI **BLAST** search of APKKVLLALTSYN
(Parameters: blastp, nr protein database, expect=20000, no filter, PAM30)
Other BLAST web gateways

**All matches to this query**

| Score | Mr(calc) | Delta | Sequence | Site Analysis |
|---|---|---|---|---|
| 62.2 | 1444.8289 | 0.0012 | APKKVLLALTSYN | Formyl K3 48.90% |
| 62.2 | 1444.8653 | -0.0351 | APKKVLLALTSYN | |
| 62.2 | 1444.8289 | 0.0012 | APKKVLLALTSYN | Formyl N-term 48.90% |
| 48.8 | 1444.8289 | 0.0012 | APKKVLLALTSYN | Formyl K4 2.21% |
