## Supplementary material for "Discovering the N-terminal Methylome by Repurposing of Proteomic Datasets": Figure S1

**Figure S1. MS/MS spectra for all hits from three repurposed yeast datasets**

Contents | Pages

**MS/MS spectra for hits in ChaFRADIC dataset** Page 1-43

**MS/MS spectra for hits from iProx dataset** Page 44-54

**MS/MS spectra for MISL hits (heavy labeled, iMet excluded)** Page 55-251

**MS/MS2 spectra for hits in ChaFRADIC dataset:**

*Only spectra of peptide score >20 will be listed here. Peptides have initial M cleaved.

1. F001801. SSA3 (13 peptides spectra)

MS/MS Fragmentation of **SRAVGIDLGTTY**
Found in **HSP73_YEAST** in **Uniprot_S288C_yeast**, sp|P09435|HSP73_YEAST Heat shock protein SSA3 OS=Saccharomyces cerevisiae (strain ATCC 204508 / S288c) OX=559292 GN=SSA3 PE=1 SV=3

Match to Query 16753: 1293.681804 from(647.848178,2+) intensity(1034732.4375) rtinseconds(4270.52364) index(6842)
Title: qExactive02198_+1.14602.14602.2
Data file D:\Panyue\enriched_fr.+1.mzML

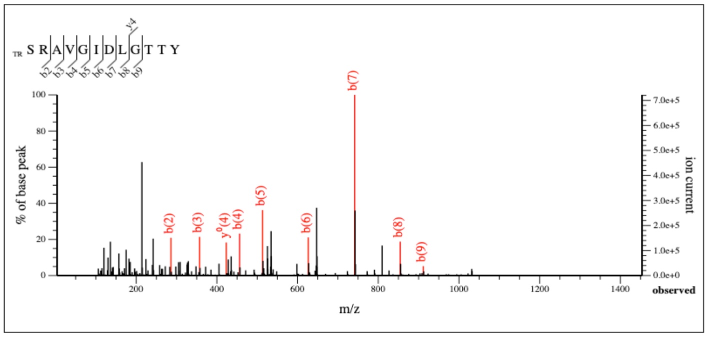


**Monoisotopic mass of neutral peptide Mr(calc):** 1293.6929
**Fixed modifications:** Carbamidomethyl (C) (apply to specified residues or termini only)
**Variable modifications:**
N-term : Trimethyl (Protein N-term)
**Ions Score:** 47 **Expect:** 0.00082
**Matches :** 10/108 fragment ions using 20 most intense peaks (help)

| # | b | b++ | b* | b*++ | b0 | b0++ | Seq. | y | y++ | y* | y*++ | y0 | y0++ | # |
|---|---|---|---|---|---|---|---|---|---|---|---|---|---|---|
| 1 | 130.0863 | 65.5468 | | | 112.0757 | 56.5415 | S | | | | | | | 12 |
| 2 | 286.1874 | 143.5973 | 269.1608 | 135.0840 | 268.1768 | 134.5920 | R | 1165.6212 | 583.3142 | 1148.5946 | 574.8009 | 1147.6106 | 574.3089 | 11 |
| 3 | 357.2245 | 179.1159 | 340.1979 | 170.6026 | 339.2139 | 170.1106 | A | 1009.5201 | 505.2637 | | | 991.5095 | 496.2584 | 10 |
| 4 | 456.2929 | 228.6501 | 439.2663 | 220.1368 | 438.2823 | 219.6448 | V | 938.4829 | 469.7451 | | | 920.4724 | 460.7398 | 9 |
| 5 | 513.3144 | 257.1608 | 496.2878 | 248.6475 | 495.3038 | 248.1555 | G | 839.4145 | 420.2109 | | | 821.4040 | 411.2056 | 8 |
| 6 | 626.3984 | 313.7028 | 609.3719 | 305.1896 | 608.3879 | 304.6976 | I | 782.3931 | 391.7002 | | | 764.3825 | 382.6949 | 7 |
| 7 | 741.4254 | 371.2163 | 724.3988 | 362.7030 | 723.4148 | 362.2110 | D | 669.3090 | 335.1581 | | | 651.2984 | 326.1529 | 6 |
| 8 | 854.5094 | 427.7584 | 837.4829 | 419.2451 | 836.4989 | 418.7531 | L | 554.2821 | 277.6447 | | | 536.2715 | 268.6394 | 5 |
| 9 | 911.5309 | 456.2691 | 894.5043 | 447.7558 | 893.5203 | 447.2638 | G | 441.1980 | 221.1026 | | | 423.1874 | 212.0974 | 4 |
| 10 | 1012.5786 | 506.7929 | 995.5520 | 498.2796 | 994.5680 | 497.7876 | T | 384.1765 | 192.5919 | | | 366.1660 | 183.5866 | 3 |
| 11 | 1113.6263 | 557.3168 | 1096.5997 | 548.8035 | 1095.6157 | 548.3115 | T | 283.1288 | 142.0681 | | | 265.1183 | 133.0628 | 2 |
| 12 | | | | | | | Y | 182.0812 | 91.5442 | | | | | 1 |

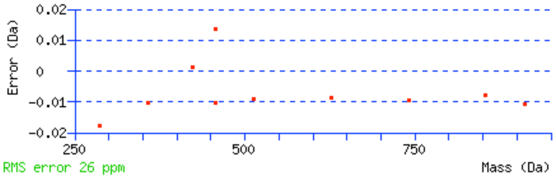


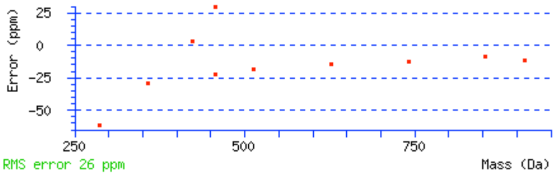

MS/MS Fragmentation of **SRAVGIDLGTTY**
Found in **HSP73_YEAST** in **Uniprot_S288C_yeast**, sp|P09435|HSP73_YEAST Heat shock protein SSA3 OS=Saccharomyces cerevisiae (strain ATCC 204508 / S288c) OX=559292 GN=SSA3 PE=1 SV=3

Match to Query 16754: 1293.682002 from(647.848277,2+) intensity(2806538.7500) rtinseconds(5450.31894) index(10596)
Title: qExactive02198_+1.18830.18830.2
Data file D:\Panyue\enriched_fr.+1.mzML

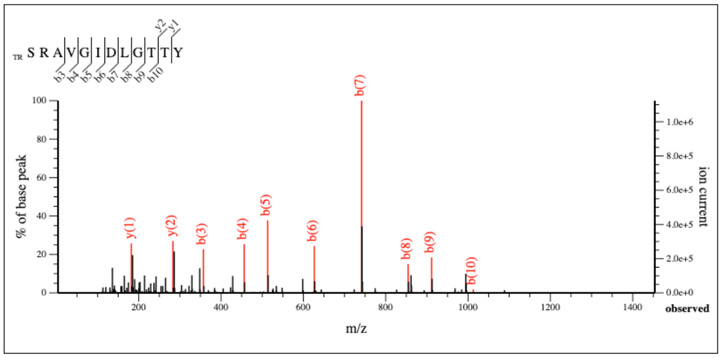


**Monoisotopic mass of neutral peptide Mr(calc):** 1293.6929
**Fixed modifications:** Carbamidomethyl (C) (apply to specified residues or termini only)
**Variable modifications:**
N-term : Trimethyl (Protein N-term)
**Ions Score:** 88 **Expect:** 6.4e-08
**Matches :** 11/108 fragment ions using 10 most intense peaks (help)

| # | b | b++ | b* | b*++ | b0 | b0++ | Seq. | y | y++ | y* | y*++ | y0 | y0++ | # |
|---|---|---|---|---|---|---|---|---|---|---|---|---|---|---|
| 1 | 130.0863 | 65.5468 | | | 112.0757 | 56.5415 | S | | | | | | | 12 |
| 2 | 286.1874 | 143.5973 | 269.1608 | 135.0840 | 268.1768 | 134.5920 | R | 1165.6212 | 583.3142 | 1148.5946 | 574.8009 | 1147.6106 | 574.3089 | 11 |
| 3 | 357.2245 | 179.1159 | 340.1979 | 170.6026 | 339.2139 | 170.1106 | A | 1009.5201 | 505.2637 | | | 991.5095 | 496.2584 | 10 |
| 4 | 456.2929 | 228.6501 | 439.2663 | 220.1368 | 438.2823 | 219.6448 | V | 938.4829 | 469.7451 | | | 920.4724 | 460.7398 | 9 |
| 5 | 513.3144 | 257.1608 | 496.2878 | 248.6475 | 495.3038 | 248.1555 | G | 839.4145 | 420.2109 | | | 821.4040 | 411.2056 | 8 |
| 6 | 626.3984 | 313.7028 | 609.3719 | 305.1896 | 608.3879 | 304.6976 | I | 782.3931 | 391.7002 | | | 764.3825 | 382.6949 | 7 |
| 7 | 741.4254 | 371.2163 | 724.3988 | 362.7030 | 723.4148 | 362.2110 | D | 669.3090 | 335.1581 | | | 651.2984 | 326.1529 | 6 |
| 8 | 854.5094 | 427.7584 | 837.4829 | 419.2451 | 836.4989 | 418.7531 | L | 554.2821 | 277.6447 | | | 536.2715 | 268.6394 | 5 |
| 9 | 911.5309 | 456.2691 | 894.5043 | 447.7558 | 893.5203 | 447.2638 | G | 441.1980 | 221.1026 | | | 423.1874 | 212.0974 | 4 |
| 10 | 1012.5786 | 506.7929 | 995.5520 | 498.2796 | 994.5680 | 497.7876 | T | 384.1765 | 192.5919 | | | 366.1660 | 183.5866 | 3 |
| 11 | 1113.6263 | 557.3168 | 1096.5997 | 548.8035 | 1095.6157 | 548.3115 | T | 283.1288 | 142.0681 | | | 265.1183 | 133.0628 | 2 |
| 12 | | | | | | | Y | 182.0812 | 91.5442 | | | | | 1 |

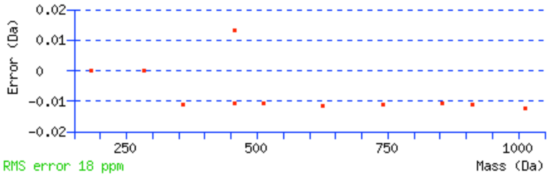


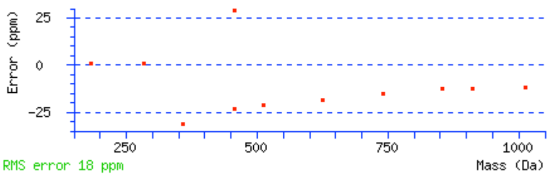

MS/MS Fragmentation of **SRAVGIDLGTTY**
Found in **HSP73_YEAST** in **Uniprot_S288C_yeast**, sp|P09435|HSP73_YEAST Heat shock protein SSA3 OS=Saccharomyces cerevisiae (strain ATCC 204508 / S288c) OX=559292 GN=SSA3 PE=1 SV=3

Match to Query 16755: 1293.682162 from(647.848357,2+) intensity(1439836.6250) rtinseconds(5467.389) index(10649)
Title: qExactive02198_+1.18888.18888.2
Data file D:\Panyue\enriched_fr.+1.mzML

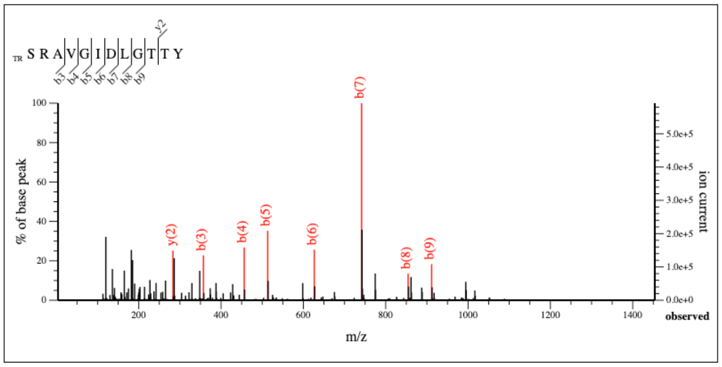


**Monoisotopic mass of neutral peptide Mr(calc):** 1293.6929
**Fixed modifications:** Carbamidomethyl (C) (apply to specified residues or termini only)
**Variable modifications:**
N-term : Trimethyl (Protein N-term)
**Ions Score:** 62 **Expect:** 2.5e-05
**Matches :** 9/108 fragment ions using 10 most intense peaks (help)

| # | b | b++ | b* | b*++ | b0 | b0++ | Seq. | y | y++ | y* | y*++ | y0 | y0++ | # |
|---|---|---|---|---|---|---|---|---|---|---|---|---|---|---|
| 1 | 130.0863 | 65.5468 | | | 112.0757 | 56.5415 | S | | | | | | | 12 |
| 2 | 286.1874 | 143.5973 | 269.1608 | 135.0840 | 268.1768 | 134.5920 | R | 1165.6212 | 583.3142 | 1148.5946 | 574.8009 | 1147.6106 | 574.3089 | 11 |
| 3 | ***357.2245*** | 179.1159 | 340.1979 | 170.6026 | 339.2139 | 170.1106 | A | 1009.5201 | 505.2637 | | | 991.5095 | 496.2584 | 10 |
| 4 | ***456.2929*** | 228.6501 | 439.2663 | 220.1368 | 438.2823 | 219.6448 | V | 938.4829 | 469.7451 | | | 920.4724 | 460.7398 | 9 |
| 5 | ***513.3144*** | 257.1608 | 496.2878 | 248.6475 | 495.3038 | 248.1555 | G | 839.4145 | 420.2109 | | | 821.4040 | 411.2056 | 8 |
| 6 | ***626.3984*** | 313.7028 | 609.3719 | 305.1896 | 608.3879 | 304.6976 | I | 782.3931 | 391.7002 | | | 764.3825 | 382.6949 | 7 |
| 7 | ***741.4254*** | 371.2163 | 724.3988 | 362.7030 | 723.4148 | 362.2110 | D | 669.3090 | 335.1581 | | | 651.2984 | 326.1529 | 6 |
| 8 | ***854.5094*** | 427.7584 | 837.4829 | 419.2451 | 836.4989 | 418.7531 | L | 554.2821 | 277.6447 | | | 536.2715 | 268.6394 | 5 |
| 9 | ***911.5309*** | **456.2691** | 894.5043 | 447.7558 | 893.5203 | 447.2638 | G | 441.1980 | 221.1026 | | | 423.1874 | 212.0974 | 4 |
| 10 | 1012.5786 | 506.7929 | 995.5520 | 498.2796 | 994.5680 | 497.7876 | T | 384.1765 | 192.5919 | | | 366.1660 | 183.5866 | 3 |
| 11 | 1113.6263 | 557.3168 | 1096.5997 | 548.8035 | 1095.6157 | 548.3115 | T | **283.1288** | 142.0681 | | | 265.1183 | 133.0628 | 2 |
| 12 | | | | | | | Y | 182.0812 | 91.5442 | | | | | 1 |

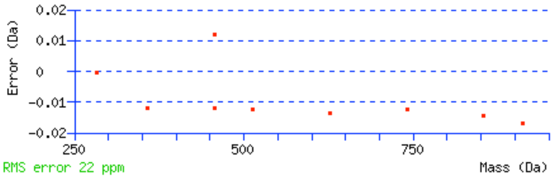


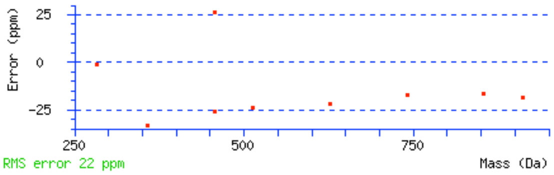

MS/MS Fragmentation of **SRAVGIDLGTTY**
Found in **HSP73_YEAST** in **Uniprot_S288C_yeast**, sp|P09435|HSP73_YEAST Heat shock protein SSA3 OS=Saccharomyces cerevisiae (strain ATCC 204508 / S288c) OX=559292 GN=SSA3 PE=1 SV=3

Match to Query 16756: 1293.682164 from(647.848358,2+) intensity(954137.2500) rtinseconds(4070.37762) index(6209)
Title: qExactive02198_+1.13909.13909.2
Data file D:\Panyue\enriched_fr.+1.mzML

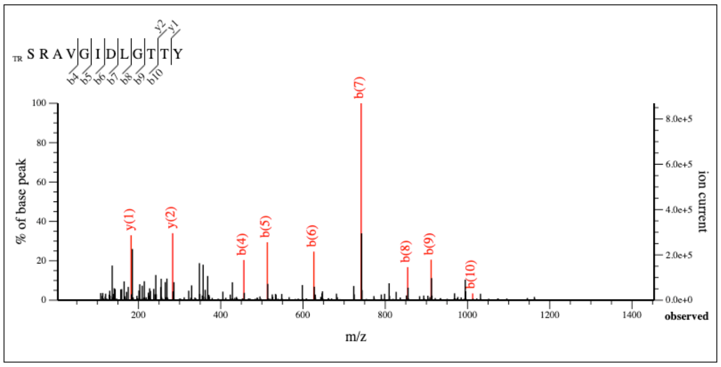


**Monoisotopic mass of neutral peptide Mr(calc):** 1293.6929
**Fixed modifications:** Carbamidomethyl (C) (apply to specified residues or termini only)
**Variable modifications:**
N-term : Trimethyl (Protein N-term)
**Ions Score:** 62 **Expect:** 2.2e-05
**Matches :** 10/108 fragment ions using 11 most intense peaks (help)

| # | b | b++ | b* | b*++ | b0 | b0++ | Seq. | y | y++ | y* | y*++ | y0 | y0++ | # |
|---|---|---|---|---|---|---|---|---|---|---|---|---|---|---|
| 1 | 130.0863 | 65.5468 | | | 112.0757 | 56.5415 | S | | | | | | | 12 |
| 2 | 286.1874 | 143.5973 | 269.1608 | 135.0840 | 268.1768 | 134.5920 | R | 1165.6212 | 583.3142 | 1148.5946 | 574.8009 | 1147.6106 | 574.3089 | 11 |
| 3 | 357.2245 | 179.1159 | 340.1979 | 170.6026 | 339.2139 | 170.1106 | A | 1009.5201 | 505.2637 | | | 991.5095 | 496.2584 | 10 |
| 4 | ***456.2929*** | 228.6501 | 439.2663 | 220.1368 | 438.2823 | 219.6448 | V | 938.4829 | 469.7451 | | | 920.4724 | 460.7398 | 9 |
| 5 | ***513.3144*** | 257.1608 | 496.2878 | 248.6475 | 495.3038 | 248.1555 | G | 839.4145 | 420.2109 | | | 821.4040 | 411.2056 | 8 |
| 6 | ***626.3984*** | 313.7028 | 609.3719 | 305.1896 | 608.3879 | 304.6976 | I | 782.3931 | 391.7002 | | | 764.3825 | 382.6949 | 7 |
| 7 | ***741.4254*** | 371.2163 | 724.3988 | 362.7030 | 723.4148 | 362.2110 | D | 669.3090 | 335.1581 | | | 651.2984 | 326.1529 | 6 |
| 8 | ***854.5094*** | 427.7584 | 837.4829 | 419.2451 | 836.4989 | 418.7531 | L | 554.2821 | 277.6447 | | | 536.2715 | 268.6394 | 5 |
| 9 | ***911.5309*** | **456.2691** | 894.5043 | 447.7558 | 893.5203 | 447.2638 | G | 441.1980 | 221.1026 | | | 423.1874 | 212.0974 | 4 |
| 10 | ***1012.5786*** | 506.7929 | 995.5520 | 498.2796 | 994.5680 | 497.7876 | T | 384.1765 | 192.5919 | | | 366.1660 | 183.5866 | 3 |
| 11 | 1113.6263 | 557.3168 | 1096.5997 | 548.8035 | 1095.6157 | 548.3115 | T | **283.1288** | 142.0681 | | | 265.1183 | 133.0628 | 2 |
| 12 | | | | | | | Y | **182.0812** | 91.5442 | | | | | 1 |

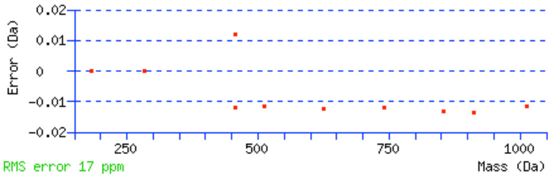


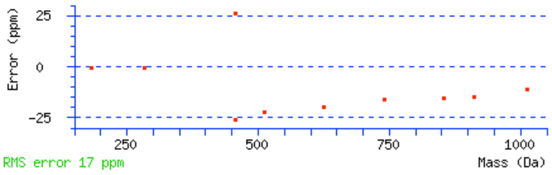

MS/MS Fragmentation of **SRAVGIDLGTTY**
Found in **HSP73_YEAST** in **Uniprot_S288C_yeast**, sp|P09435|HSP73_YEAST Heat shock protein SSA3 OS=Saccharomyces cerevisiae (strain ATCC 204508 / S288c) OX=559292 GN=SSA3 PE=1 SV=3

Match to Query 16757: 1293.682209 from(1294.689485,1+) intensity(1192507.3750) rtinseconds(5409.89064) index(10464)
Title: qExactive02198_+1.18690.18690.1
Data file D:\Panyue\enriched_fr.+1.mzML

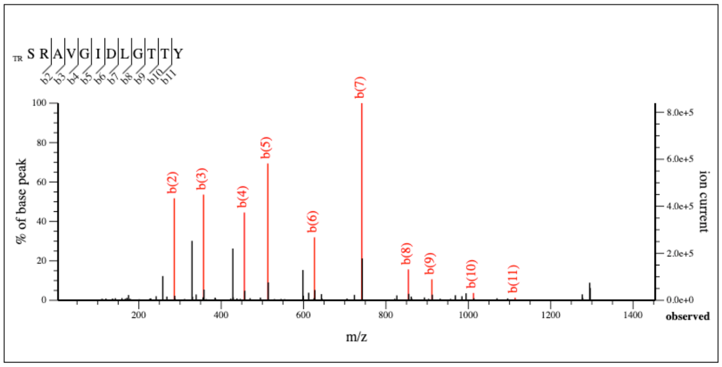


**Monoisotopic mass of neutral peptide Mr(calc):** 1293.6929
**Fixed modifications:** Carbamidomethyl (C) (apply to specified residues or termini only)
**Variable modifications:**
N-term : Trimethyl (Protein N-term)
**Ions Score:** 112 **Expect:** 2.2e-10
**Matches :** 10/54 fragment ions using 12 most intense peaks (help)

| # | b | b* | b⁰ | Seq. | y | y* | y⁰ | # |
|---|---|---|---|---|---|---|---|---|
| 1 | 130.0863 | | 112.0757 | S | | | | 12 |
| 2 | 286.1874 | 269.1608 | 268.1768 | R | 1165.6212 | 1148.5946 | 1147.6106 | 11 |
| 3 | 357.2245 | 340.1979 | 339.2139 | A | 1009.5201 | | 991.5095 | 10 |
| 4 | 456.2929 | 439.2663 | 438.2823 | V | 938.4829 | | 920.4724 | 9 |
| 5 | 513.3144 | 496.2878 | 495.3038 | G | 839.4145 | | 821.4040 | 8 |
| 6 | 626.3984 | 609.3719 | 608.3879 | I | 782.3931 | | 764.3825 | 7 |
| 7 | 741.4254 | 724.3988 | 723.4148 | D | 669.3090 | | 651.2984 | 6 |
| 8 | 854.5094 | 837.4829 | 836.4989 | L | 554.2821 | | 536.2715 | 5 |
| 9 | 911.5309 | 894.5043 | 893.5203 | G | 441.1980 | | 423.1874 | 4 |
| 10 | 1012.5786 | 995.5520 | 994.5680 | T | 384.1765 | | 366.1660 | 3 |
| 11 | 1113.6263 | 1096.5997 | 1095.6157 | T | 283.1288 | | 265.1183 | 2 |
| 12 | | | | Y | 182.0812 | | | 1 |

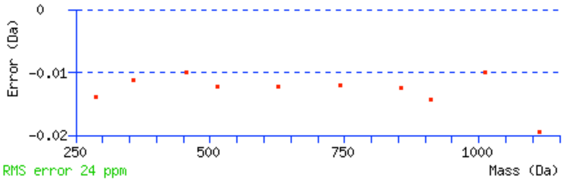


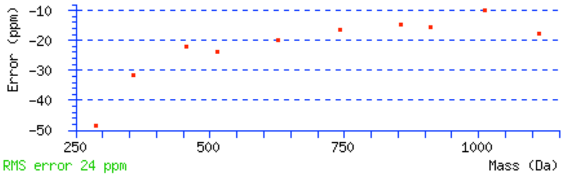

MS/MS Fragmentation of **SRAVGIDLGTTY**
Found in **HSP73_YEAST** in **Uniprot_S288C_yeast**, sp|P09435|HSP73_YEAST Heat shock protein SSA3 OS=Saccharomyces cerevisiae (strain ATCC 204508 / S288c) OX=559292 GN=SSA3 PE=1 SV=3

Match to Query 16758: 1293.682416 from(647.848484,2+) intensity(656189.4375) rtinseconds(5263.53756) index(10001)
Title: qExactive02198_+1.18187.18187.2
Data file D:\Panyue\enriched_fr.+1.mzML

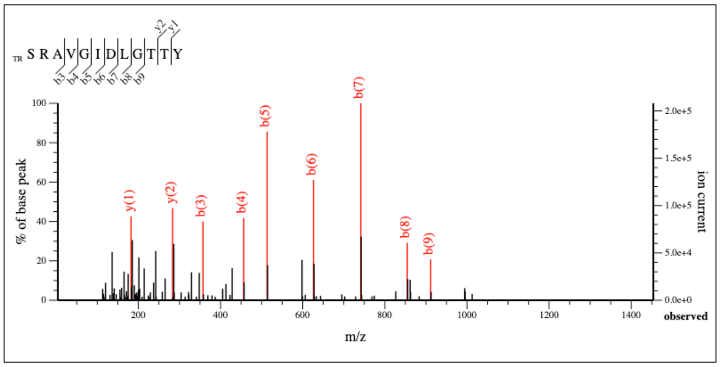


**Monoisotopic mass of neutral peptide Mr(calc):** 1293.6929
**Fixed modifications:** Carbamidomethyl (C) (apply to specified residues or termini only)
**Variable modifications:**
N-term : Trimethyl (Protein N-term)
**Ions Score:** 68 **Expect:** 5.8e-06
**Matches :** 10/108 fragment ions using 10 most intense peaks (help)

| # | b | $b^{++}$ | $b^{*}$ | $b^{*++}$ | $b^{0}$ | $b^{0++}$ | Seq. | y | $y^{++}$ | $y^{*}$ | $y^{*++}$ | $y^{0}$ | $y^{0++}$ | # |
|---|---|---|---|---|---|---|---|---|---|---|---|---|---|---|
| 1 | 130.0863 | 65.5468 | | | 112.0757 | 56.5415 | S | | | | | | | 12 |
| 2 | 286.1874 | 143.5973 | 269.1608 | 135.0840 | 268.1768 | 134.5920 | R | 1165.6212 | 583.3142 | 1148.5946 | 574.8009 | 1147.6106 | 574.3089 | 11 |
| 3 | ***357.2245*** | 179.1159 | 340.1979 | 170.6026 | 339.2139 | 170.1106 | A | 1009.5201 | 505.2637 | | | 991.5095 | 496.2584 | 10 |
| 4 | ***456.2929*** | 228.6501 | 439.2663 | 220.1368 | 438.2823 | 219.6448 | V | 938.4829 | 469.7451 | | | 920.4724 | 460.7398 | 9 |
| 5 | ***513.3144*** | 257.1608 | 496.2878 | 248.6475 | 495.3038 | 248.1555 | G | 839.4145 | 420.2109 | | | 821.4040 | 411.2056 | 8 |
| 6 | ***626.3984*** | 313.7028 | 609.3719 | 305.1896 | 608.3879 | 304.6976 | I | 782.3931 | 391.7002 | | | 764.3825 | 382.6949 | 7 |
| 7 | ***741.4254*** | 371.2163 | 724.3988 | 362.7030 | 723.4148 | 362.2110 | D | 669.3090 | 335.1581 | | | 651.2984 | 326.1529 | 6 |
| 8 | ***854.5094*** | 427.7584 | 837.4829 | 419.2451 | 836.4989 | 418.7531 | L | 554.2821 | 277.6447 | | | 536.2715 | 268.6394 | 5 |
| 9 | ***911.5309*** | **456.2691** | 894.5043 | 447.7558 | 893.5203 | 447.2638 | G | 441.1980 | 221.1026 | | | 423.1874 | 212.0974 | 4 |
| 10 | 1012.5786 | 506.7929 | 995.5520 | 498.2796 | 994.5680 | 497.7876 | T | 384.1765 | 192.5919 | | | 366.1660 | 183.5866 | 3 |
| 11 | 1113.6263 | 557.3168 | 1096.5997 | 548.8035 | 1095.6157 | 548.3115 | T | **283.1288** | 142.0681 | | | 265.1183 | 133.0628 | 2 |
| 12 | | | | | | | Y | **182.0812** | 91.5442 | | | | | 1 |

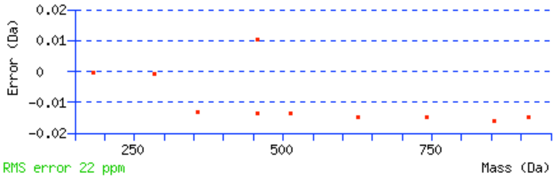


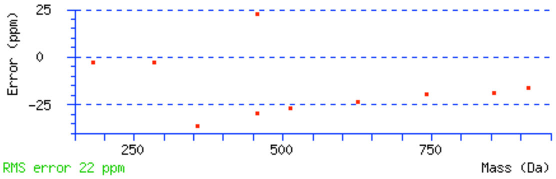

MS/MS Fragmentation of **SRAVGIDLGTTY**
Found in **HSP73_YEAST** in **Uniprot_S288C_yeast**, sp|P09435|HSP73_YEAST Heat shock protein SSA3 OS=Saccharomyces cerevisiae (strain ATCC 204508 / S288c) OX=559292 GN=SSA3 PE=1 SV=3

Match to Query 16759: 1293.682438 from(647.848495,2+) intensity(850389.0625) rtinseconds(5250.09618) index(9959)
Title: qExactive02198_+1.18138.18138.2
Data file D:\Panyue\enriched_fr.+1.mzML

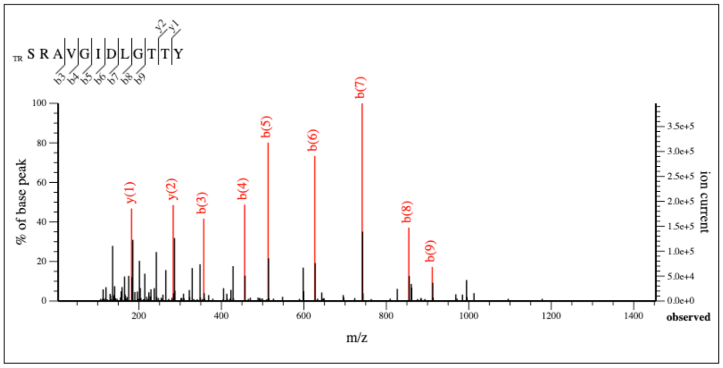


**Monoisotopic mass of neutral peptide Mr(calc):** 1293.6929
**Fixed modifications:** Carbamidomethyl (C) (apply to specified residues or termini only)
**Variable modifications:**
N-term : Trimethyl (Protein N-term)
**Ions Score:** 63 **Expect:** 1.7e-05
**Matches :** 10/108 fragment ions using 11 most intense peaks (help)

| # | b | $b^{++}$ | $b^{*}$ | $b^{*++}$ | $b^{0}$ | $b^{0++}$ | Seq. | y | $y^{++}$ | $y^{*}$ | $y^{*++}$ | $y^{0}$ | $y^{0++}$ | # |
|---|---|---|---|---|---|---|---|---|---|---|---|---|---|---|
| 1 | 130.0863 | 65.5468 | | | 112.0757 | 56.5415 | S | | | | | | | 12 |
| 2 | 286.1874 | 143.5973 | 269.1608 | 135.0840 | 268.1768 | 134.5920 | R | 1165.6212 | 583.3142 | 1148.5946 | 574.8009 | 1147.6106 | 574.3089 | 11 |
| 3 | ***357.2245*** | 179.1159 | 340.1979 | 170.6026 | 339.2139 | 170.1106 | A | 1009.5201 | 505.2637 | | | 991.5095 | 496.2584 | 10 |
| 4 | ***456.2929*** | 228.6501 | 439.2663 | 220.1368 | 438.2823 | 219.6448 | V | 938.4829 | 469.7451 | | | 920.4724 | 460.7398 | 9 |
| 5 | ***513.3144*** | 257.1608 | 496.2878 | 248.6475 | 495.3038 | 248.1555 | G | 839.4145 | 420.2109 | | | 821.4040 | 411.2056 | 8 |
| 6 | ***626.3984*** | 313.7028 | 609.3719 | 305.1896 | 608.3879 | 304.6976 | I | 782.3931 | 391.7002 | | | 764.3825 | 382.6949 | 7 |
| 7 | ***741.4254*** | 371.2163 | 724.3988 | 362.7030 | 723.4148 | 362.2110 | D | 669.3090 | 335.1581 | | | 651.2984 | 326.1529 | 6 |
| 8 | ***854.5094*** | 427.7584 | 837.4829 | 419.2451 | 836.4989 | 418.7531 | L | 554.2821 | 277.6447 | | | 536.2715 | 268.6394 | 5 |
| 9 | ***911.5309*** | **456.2691** | 894.5043 | 447.7558 | 893.5203 | 447.2638 | G | 441.1980 | 221.1026 | | | 423.1874 | 212.0974 | 4 |
| 10 | 1012.5786 | 506.7929 | 995.5520 | 498.2796 | 994.5680 | 497.7876 | T | 384.1765 | 192.5919 | | | 366.1660 | 183.5866 | 3 |
| 11 | 1113.6263 | 557.3168 | 1096.5997 | 548.8035 | 1095.6157 | 548.3115 | T | **283.1288** | 142.0681 | | | 265.1183 | 133.0628 | 2 |
| 12 | | | | | | | Y | **182.0812** | 91.5442 | | | | | 1 |

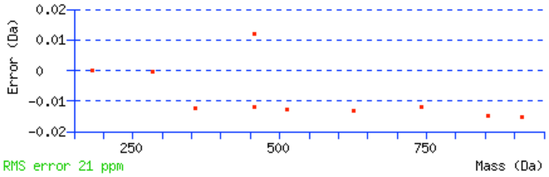


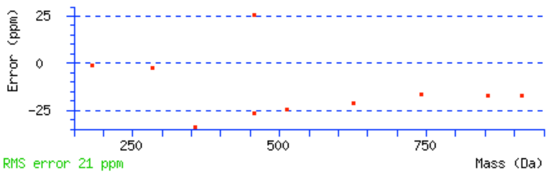

MS/MS Fragmentation of **SRAVGIDLGTTY**
Found in **HSP73_YEAST** in **Uniprot_S288C_yeast**, sp|P09435|HSP73_YEAST Heat shock protein SSA3 OS=Saccharomyces cerevisiae (strain ATCC 204508 / S288c) OX=559292 GN=SSA3 PE=1 SV=3

Match to Query 16760: 1293.682724 from(647.848638,2+) intensity(641713.0625) rtinseconds(5371.32714) index(10341)
Title: qExactive02198_+1.18556.18556.2
Data file D:\Panyue\enriched_fr.+1.mzML

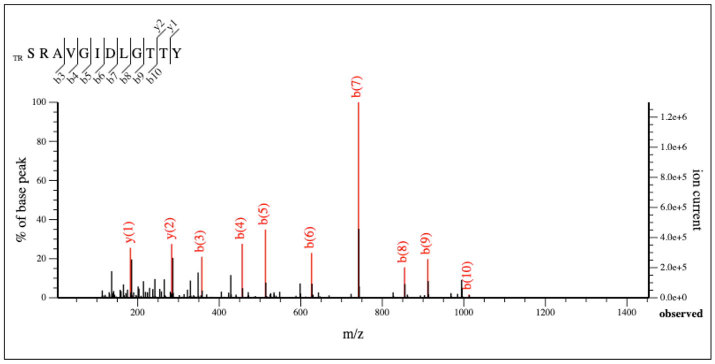


**Monoisotopic mass of neutral peptide Mr(calc):** 1293.6929
**Fixed modifications:** Carbamidomethyl (C) (apply to specified residues or termini only)
**Variable modifications:**
N-term : Trimethyl (Protein N-term)
**Ions Score:** 88 **Expect:** 6.3e-08
**Matches :** 11/108 fragment ions using 10 most intense peaks (help)

| # | b | b++ | b* | b*++ | b0 | b0++ | Seq. | y | y++ | y* | y*++ | y0 | y0++ | # |
|---|---|---|---|---|---|---|---|---|---|---|---|---|---|---|
| 1 | 130.0863 | 65.5468 | | | 112.0757 | 56.5415 | S | | | | | | | 12 |
| 2 | 286.1874 | 143.5973 | 269.1608 | 135.0840 | 268.1768 | 134.5920 | R | 1165.6212 | 583.3142 | 1148.5946 | 574.8009 | 1147.6106 | 574.3089 | 11 |
| 3 | 357.2245 | 179.1159 | 340.1979 | 170.6026 | 339.2139 | 170.1106 | A | 1009.5201 | 505.2637 | | | 991.5095 | 496.2584 | 10 |
| 4 | 456.2929 | 228.6501 | 439.2663 | 220.1368 | 438.2823 | 219.6448 | V | 938.4829 | 469.7451 | | | 920.4724 | 460.7398 | 9 |
| 5 | 513.3144 | 257.1608 | 496.2878 | 248.6475 | 495.3038 | 248.1555 | G | 839.4145 | 420.2109 | | | 821.4040 | 411.2056 | 8 |
| 6 | 626.3984 | 313.7028 | 609.3719 | 305.1896 | 608.3879 | 304.6976 | I | 782.3931 | 391.7002 | | | 764.3825 | 382.6949 | 7 |
| 7 | 741.4254 | 371.2163 | 724.3988 | 362.7030 | 723.4148 | 362.2110 | D | 669.3090 | 335.1581 | | | 651.2984 | 326.1529 | 6 |
| 8 | 854.5094 | 427.7584 | 837.4829 | 419.2451 | 836.4989 | 418.7531 | L | 554.2821 | 277.6447 | | | 536.2715 | 268.6394 | 5 |
| 9 | 911.5309 | 456.2691 | 894.5043 | 447.7558 | 893.5203 | 447.2638 | G | 441.1980 | 221.1026 | | | 423.1874 | 212.0974 | 4 |
| 10 | 1012.5786 | 506.7929 | 995.5520 | 498.2796 | 994.5680 | 497.7876 | T | 384.1765 | 192.5919 | | | 366.1660 | 183.5866 | 3 |
| 11 | 1113.6263 | 557.3168 | 1096.5997 | 548.8035 | 1095.6157 | 548.3115 | T | 283.1288 | 142.0681 | | | 265.1183 | 133.0628 | 2 |
| 12 | | | | | | | Y | 182.0812 | 91.5442 | | | | | 1 |

MS/MS Fragmentation of **SRAVGIDLGTTY**
Found in **HSP73_YEAST** in **Uniprot_S288C_yeast**, sp|P09435|HSP73_YEAST Heat shock protein SSA3 OS=Saccharomyces cerevisiae (strain ATCC 204508 / S288c) OX=559292 GN=SSA3 PE=1 SV=3

Match to Query 16761: 1293.682944 from(647.848748,2+) intensity(7808885.0000) rtinseconds(5431.0479) index(10535)
Title: qExactive02198_+1.18765.18765.2
Data file D:\Panyue\enriched_fr.+1.mzML

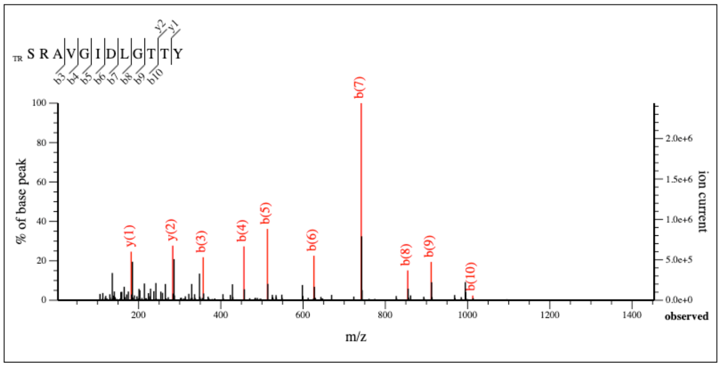


**Monoisotopic mass of neutral peptide Mr(calc):** 1293.6929
**Fixed modifications:** Carbamidomethyl (C) (apply to specified residues or termini only)
**Variable modifications:**
N-term : Trimethyl (Protein N-term)
**Ions Score:** 88 **Expect:** 6.3e-08
**Matches :** 11/108 fragment ions using 10 most intense peaks (help)

| # | b | $b^{++}$ | $b^{*}$ | $b^{*++}$ | $b^{0}$ | $b^{0++}$ | Seq. | y | $y^{++}$ | $y^{*}$ | $y^{*++}$ | $y^{0}$ | $y^{0++}$ | # |
|---|---|---|---|---|---|---|---|---|---|---|---|---|---|---|
| 1 | 130.0863 | 65.5468 | | | 112.0757 | 56.5415 | S | | | | | | | 12 |
| 2 | 286.1874 | 143.5973 | 269.1608 | 135.0840 | 268.1768 | 134.5920 | R | 1165.6212 | 583.3142 | 1148.5946 | 574.8009 | 1147.6106 | 574.3089 | 11 |
| 3 | ***357.2245*** | 179.1159 | 340.1979 | 170.6026 | 339.2139 | 170.1106 | A | 1009.5201 | 505.2637 | | | 991.5095 | 496.2584 | 10 |
| 4 | ***456.2929*** | 228.6501 | 439.2663 | 220.1368 | 438.2823 | 219.6448 | V | 938.4829 | 469.7451 | | | 920.4724 | 460.7398 | 9 |
| 5 | ***513.3144*** | 257.1608 | 496.2878 | 248.6475 | 495.3038 | 248.1555 | G | 839.4145 | 420.2109 | | | 821.4040 | 411.2056 | 8 |
| 6 | ***626.3984*** | 313.7028 | 609.3719 | 305.1896 | 608.3879 | 304.6976 | I | 782.3931 | 391.7002 | | | 764.3825 | 382.6949 | 7 |
| 7 | ***741.4254*** | 371.2163 | 724.3988 | 362.7030 | 723.4148 | 362.2110 | D | 669.3090 | 335.1581 | | | 651.2984 | 326.1529 | 6 |
| 8 | ***854.5094*** | 427.7584 | 837.4829 | 419.2451 | 836.4989 | 418.7531 | L | 554.2821 | 277.6447 | | | 536.2715 | 268.6394 | 5 |
| 9 | ***911.5309*** | **456.2691** | 894.5043 | 447.7558 | 893.5203 | 447.2638 | G | 441.1980 | 221.1026 | | | 423.1874 | 212.0974 | 4 |
| 10 | ***1012.5786*** | 506.7929 | 995.5520 | 498.2796 | 994.5680 | 497.7876 | T | 384.1765 | 192.5919 | | | 366.1660 | 183.5866 | 3 |
| 11 | 1113.6263 | 557.3168 | 1096.5997 | 548.8035 | 1095.6157 | 548.3115 | T | **283.1288** | 142.0681 | | | 265.1183 | 133.0628 | 2 |
| 12 | | | | | | | Y | **182.0812** | 91.5442 | | | | | 1 |

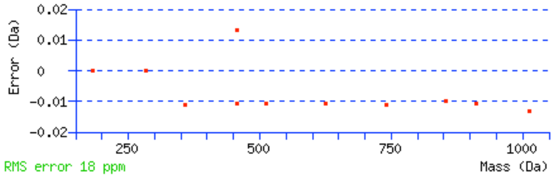


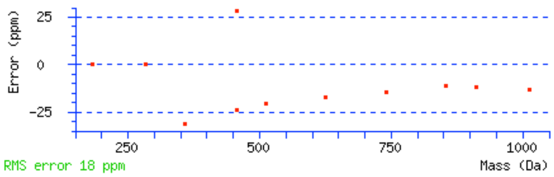

MS/MS Fragmentation of **SRAVGIDLGTTY**
Found in **HSP73_YEAST** in **Uniprot_S288C_yeast**, sp|P09435|HSP73_YEAST Heat shock protein SSA3 OS=Saccharomyces cerevisiae (strain ATCC 204508 / S288c) OX=559292 GN=SSA3 PE=1 SV=3

Match to Query 16762: 1293.684494 from(647.849523,2+) intensity(34674632.0000) rtinseconds(5412.97512) index(10476)
Title: qExactive02198_+1.18702.18702.2
Data file D:\Panyue\enriched_fr.+1.mzML

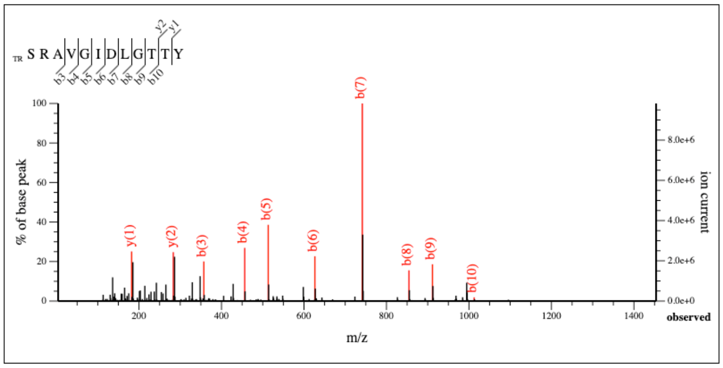


**Monoisotopic mass of neutral peptide Mr(calc):** 1293.6929
**Fixed modifications:** Carbamidomethyl (C) (apply to specified residues or termini only)
**Variable modifications:**
N-term : Trimethyl (Protein N-term)
**Ions Score:** 88 **Expect:** 6.3e-08
**Matches :** 11/108 fragment ions using 10 most intense peaks (help)

| # | b | b++ | b* | b*++ | b0 | b0++ | Seq. | y | y++ | y* | y*++ | y0 | y0++ | # |
|---|---|---|---|---|---|---|---|---|---|---|---|---|---|---|
| 1 | 130.0863 | 65.5468 | | | 112.0757 | 56.5415 | S | | | | | | | 12 |
| 2 | 286.1874 | 143.5973 | 269.1608 | 135.0840 | 268.1768 | 134.5920 | R | 1165.6212 | 583.3142 | 1148.5946 | 574.8009 | 1147.6106 | 574.3089 | 11 |
| 3 | *357.2245* | 179.1159 | 340.1979 | 170.6026 | 339.2139 | 170.1106 | A | 1009.5201 | 505.2637 | | | 991.5095 | 496.2584 | 10 |
| 4 | *456.2929* | 228.6501 | 439.2663 | 220.1368 | 438.2823 | 219.6448 | V | 938.4829 | 469.7451 | | | 920.4724 | 460.7398 | 9 |
| 5 | *513.3144* | 257.1608 | 496.2878 | 248.6475 | 495.3038 | 248.1555 | G | 839.4145 | 420.2109 | | | 821.4040 | 411.2056 | 8 |
| 6 | *626.3984* | 313.7028 | 609.3719 | 305.1896 | 608.3879 | 304.6976 | I | 782.3931 | 391.7002 | | | 764.3825 | 382.6949 | 7 |
| 7 | *741.4254* | 371.2163 | 724.3988 | 362.7030 | 723.4148 | 362.2110 | D | 669.3090 | 335.1581 | | | 651.2984 | 326.1529 | 6 |
| 8 | *854.5094* | 427.7584 | 837.4829 | 419.2451 | 836.4989 | 418.7531 | L | 554.2821 | 277.6447 | | | 536.2715 | 268.6394 | 5 |
| 9 | *911.5309* | **456.2691** | 894.5043 | 447.7558 | 893.5203 | 447.2638 | G | 441.1980 | 221.1026 | | | 423.1874 | 212.0974 | 4 |
| 10 | *1012.5786* | 506.7929 | 995.5520 | 498.2796 | 994.5680 | 497.7876 | T | 384.1765 | 192.5919 | | | 366.1660 | 183.5866 | 3 |
| 11 | 1113.6263 | 557.3168 | 1096.5997 | 548.8035 | 1095.6157 | 548.3115 | T | **283.1288** | 142.0681 | | | 265.1183 | 133.0628 | 2 |
| 12 | | | | | | | Y | **182.0812** | 91.5442 | | | | | 1 |

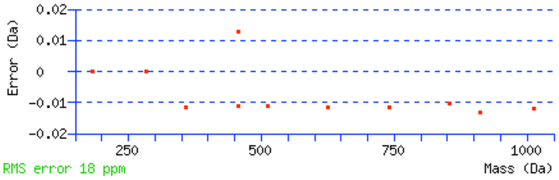


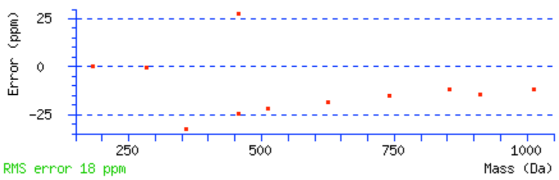

MS/MS Fragmentation of **SRAVGIDLGTTY**
Found in **HSP73_YEAST** in **Uniprot_S288C_yeast**, sp|P09435|HSP73_YEAST Heat shock protein SSA3 OS=Saccharomyces cerevisiae (strain ATCC 204508 / S288c) OX=559292 GN=SSA3 PE=1 SV=3

Match to Query 16763: 1293.687346 from(647.850949,2+) intensity(236557264.0000) rtinseconds(5388.48474) index(10396)
Title: qExactive02198_+1.18616.18616.2
Data file D:\Panyue\enriched_fr.+1.mzML

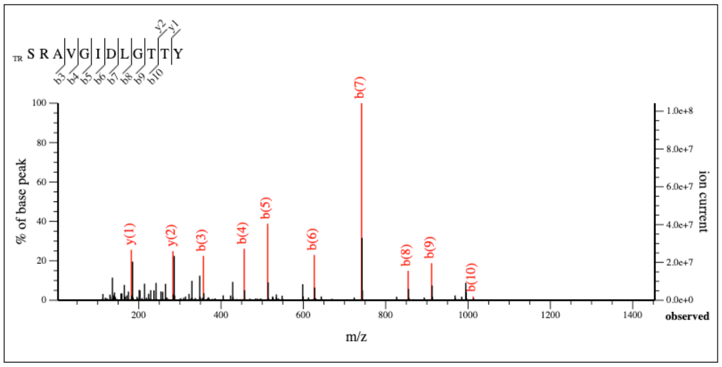


**Monoisotopic mass of neutral peptide Mr(calc):** 1293.6929
**Fixed modifications:** Carbamidomethyl (C) (apply to specified residues or termini only)
**Variable modifications:**
N-term : Trimethyl (Protein N-term)
**Ions Score:** 88 **Expect:** 6.7e-08
**Matches :** 11/108 fragment ions using 10 most intense peaks (help)

| # | b | b++ | b* | b*++ | b0 | b0++ | Seq. | y | y++ | y* | y*++ | y0 | y0++ | # |
|---|---|---|---|---|---|---|---|---|---|---|---|---|---|---|
| 1 | 130.0863 | 65.5468 | | | 112.0757 | 56.5415 | S | | | | | | | 12 |
| 2 | 286.1874 | 143.5973 | 269.1608 | 135.0840 | 268.1768 | 134.5920 | R | 1165.6212 | 583.3142 | 1148.5946 | 574.8009 | 1147.6106 | 574.3089 | 11 |
| 3 | ***357.2245*** | 179.1159 | 340.1979 | 170.6026 | 339.2139 | 170.1106 | A | 1009.5201 | 505.2637 | | | 991.5095 | 496.2584 | 10 |
| 4 | ***456.2929*** | 228.6501 | 439.2663 | 220.1368 | 438.2823 | 219.6448 | V | 938.4829 | 469.7451 | | | 920.4724 | 460.7398 | 9 |
| 5 | ***513.3144*** | 257.1608 | 496.2878 | 248.6475 | 495.3038 | 248.1555 | G | 839.4145 | 420.2109 | | | 821.4040 | 411.2056 | 8 |
| 6 | ***626.3984*** | 313.7028 | 609.3719 | 305.1896 | 608.3879 | 304.6976 | I | 782.3931 | 391.7002 | | | 764.3825 | 382.6949 | 7 |
| 7 | ***741.4254*** | 371.2163 | 724.3988 | 362.7030 | 723.4148 | 362.2110 | D | 669.3090 | 335.1581 | | | 651.2984 | 326.1529 | 6 |
| 8 | ***854.5094*** | 427.7584 | 837.4829 | 419.2451 | 836.4989 | 418.7531 | L | 554.2821 | 277.6447 | | | 536.2715 | 268.6394 | 5 |
| 9 | ***911.5309*** | **456.2691** | 894.5043 | 447.7558 | 893.5203 | 447.2638 | G | 441.1980 | 221.1026 | | | 423.1874 | 212.0974 | 4 |
| 10 | ***1012.5786*** | 506.7929 | 995.5520 | 498.2796 | 994.5680 | 497.7876 | T | 384.1765 | 192.5919 | | | 366.1660 | 183.5866 | 3 |
| 11 | 1113.6263 | 557.3168 | 1096.5997 | 548.8035 | 1095.6157 | 548.3115 | T | **283.1288** | 142.0681 | | | 265.1183 | 133.0628 | 2 |
| 12 | | | | | | | Y | **182.0812** | 91.5442 | | | | | 1 |

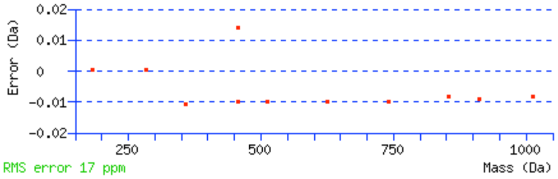


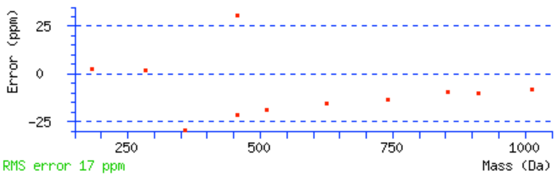

MS/MS Fragmentation of **SRAVGIDLGTTYS**
Found in **HSP73_YEAST** in **Uniprot_S288C_yeast**, sp|P09435|HSP73_YEAST Heat shock protein SSA3 OS=Saccharomyces cerevisiae (strain ATCC 204508 / S288c) OX=559292 GN=SSA3 PE=1 SV=3

Match to Query 17447: 1380.714334 from(691.364443,2+) intensity(551369.4375) rtinseconds(5175.30096) index(9746)
Title: qExactive02198_+1.17858.17858.2
Data file D:\Panyue\enriched_fr.+1.mzML

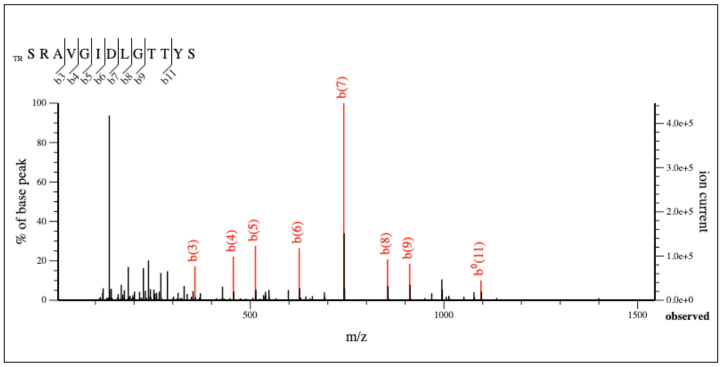


**Monoisotopic mass of neutral peptide Mr(calc):** 1380.7249
**Fixed modifications:** Carbamidomethyl (C) (apply to specified residues or termini only)
**Variable modifications:**
N-term : Trimethyl (Protein N-term)
**Ions Score:** 56 **Expect:** 8.2e-05
**Matches :** 9/120 fragment ions using 11 most intense peaks (help)

| # | b | $b^{++}$ | $b^{*}$ | $b^{*++}$ | $b^{0}$ | $b^{0++}$ | Seq. | y | $y^{++}$ | $y^{*}$ | $y^{*++}$ | $y^{0}$ | $y^{0++}$ | # |
|---|---|---|---|---|---|---|---|---|---|---|---|---|---|---|
| 1 | 130.0863 | 65.5468 | | | 112.0757 | 56.5415 | S | | | | | | | 13 |
| 2 | 286.1874 | 143.5973 | 269.1608 | 135.0840 | 268.1768 | 134.5920 | R | 1252.6532 | 626.8302 | 1235.6266 | 618.3170 | 1234.6426 | 617.8250 | 12 |
| 3 | ***357.2245*** | 179.1159 | 340.1979 | 170.6026 | 339.2139 | 170.1106 | A | 1096.5521 | 548.7797 | | | 1078.5415 | 539.7744 | 11 |
| 4 | ***456.2929*** | 228.6501 | 439.2663 | 220.1368 | 438.2823 | 219.6448 | V | 1025.5150 | 513.2611 | | | 1007.5044 | 504.2558 | 10 |
| 5 | ***513.3144*** | 257.1608 | 496.2878 | 248.6475 | 495.3038 | 248.1555 | G | 926.4466 | 463.7269 | | | 908.4360 | 454.7216 | 9 |
| 6 | ***626.3984*** | 313.7028 | 609.3719 | 305.1896 | 608.3879 | 304.6976 | I | 869.4251 | 435.2162 | | | 851.4145 | 426.2109 | 8 |
| 7 | ***741.4254*** | 371.2163 | 724.3988 | 362.7030 | 723.4148 | 362.2110 | D | 756.3410 | 378.6742 | | | 738.3305 | 369.6689 | 7 |
| 8 | ***854.5094*** | 427.7584 | 837.4829 | 419.2451 | 836.4989 | 418.7531 | L | 641.3141 | 321.1607 | | | 623.3035 | 312.1554 | 6 |
| 9 | ***911.5309*** | **456.2691** | 894.5043 | 447.7558 | 893.5203 | 447.2638 | G | 528.2300 | 264.6186 | | | 510.2195 | 255.6134 | 5 |
| 10 | 1012.5786 | 506.7929 | 995.5520 | 498.2796 | 994.5680 | 497.7876 | T | 471.2086 | 236.1079 | | | 453.1980 | 227.1026 | 4 |
| 11 | 1113.6263 | 557.3168 | 1096.5997 | 548.8035 | 1095.6157 | 548.3115 | T | 370.1609 | 185.5841 | | | 352.1503 | 176.5788 | 3 |
| 12 | 1276.6896 | 638.8484 | 1259.6630 | 630.3352 | 1258.6790 | 629.8431 | Y | 269.1132 | 135.0602 | | | 251.1026 | 126.0550 | 2 |
| 13 | | | | | | | S | 106.0499 | 53.5286 | | | 88.0393 | 44.5233 | 1 |

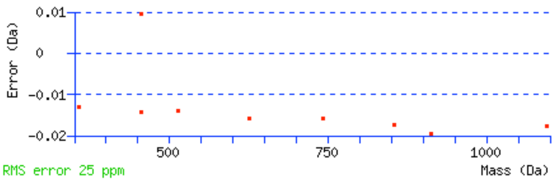


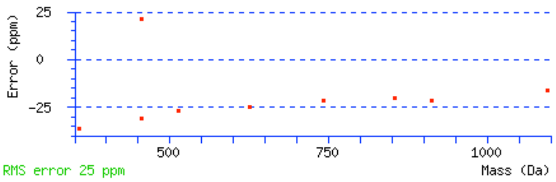

MS/MS Fragmentation of **SRAVGIDLGTTYS**
Found in **HSP73_YEAST** in **Uniprot_S288C_yeast**, sp|P09435|HSP73_YEAST Heat shock protein SSA3 OS=Saccharomyces cerevisiae (strain ATCC 204508 / S288c) OX=559292 GN=SSA3 PE=1 SV=3

Match to Query 17448: 1380.714678 from(691.364615,2+) intensity(883530.5625) rtinseconds(5162.13906) index(9708)
Title: qExactive02198_+1.17807.17807.2
Data file D:\Panyue\enriched_fr.+1.mzML

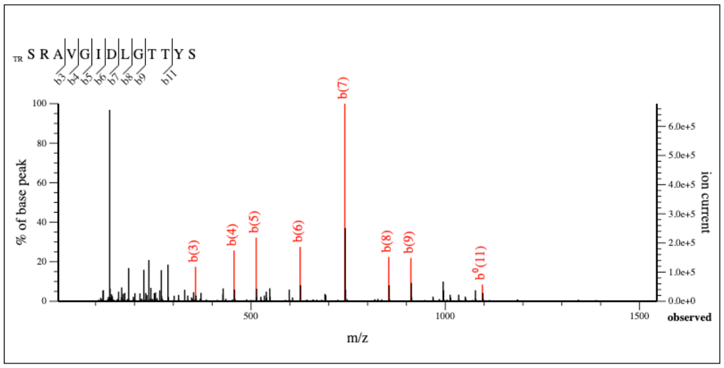


**Monoisotopic mass of neutral peptide Mr(calc):** 1380.7249
**Fixed modifications:** Carbamidomethyl (C) (apply to specified residues or termini only)
**Variable modifications:**
N-term : Trimethyl (Protein N-term)
**Ions Score:** 53 **Expect:** 0.00018
**Matches :** 9/120 fragment ions using 12 most intense peaks (help)

| # | b | b++ | b* | b*++ | b0 | b0++ | Seq. | y | y++ | y* | y*++ | y0 | y0++ | # |
|---|---|---|---|---|---|---|---|---|---|---|---|---|---|---|
| 1 | 130.0863 | 65.5468 | | | 112.0757 | 56.5415 | S | | | | | | | 13 |
| 2 | 286.1874 | 143.5973 | 269.1608 | 135.0840 | 268.1768 | 134.5920 | R | 1252.6532 | 626.8302 | 1235.6266 | 618.3170 | 1234.6426 | 617.8250 | 12 |
| 3 | ***357.2245*** | 179.1159 | 340.1979 | 170.6026 | 339.2139 | 170.1106 | A | 1096.5521 | 548.7797 | | | 1078.5415 | 539.7744 | 11 |
| 4 | ***456.2929*** | 228.6501 | 439.2663 | 220.1368 | 438.2823 | 219.6448 | V | 1025.5150 | 513.2611 | | | 1007.5044 | 504.2558 | 10 |
| 5 | ***513.3144*** | 257.1608 | 496.2878 | 248.6475 | 495.3038 | 248.1555 | G | 926.4466 | 463.7269 | | | 908.4360 | 454.7216 | 9 |
| 6 | ***626.3984*** | 313.7028 | 609.3719 | 305.1896 | 608.3879 | 304.6976 | I | 869.4251 | 435.2162 | | | 851.4145 | 426.2109 | 8 |
| 7 | ***741.4254*** | 371.2163 | 724.3988 | 362.7030 | 723.4148 | 362.2110 | D | 756.3410 | 378.6742 | | | 738.3305 | 369.6689 | 7 |
| 8 | ***854.5094*** | 427.7584 | 837.4829 | 419.2451 | 836.4989 | 418.7531 | L | 641.3141 | 321.1607 | | | 623.3035 | 312.1554 | 6 |
| 9 | ***911.5309*** | **456.2691** | 894.5043 | 447.7558 | 893.5203 | 447.2638 | G | 528.2300 | 264.6186 | | | 510.2195 | 255.6134 | 5 |
| 10 | 1012.5786 | 506.7929 | 995.5520 | 498.2796 | 994.5680 | 497.7876 | T | 471.2086 | 236.1079 | | | 453.1980 | 227.1026 | 4 |
| 11 | 1113.6263 | 557.3168 | 1096.5997 | 548.8035 | 1095.6157 | 548.3115 | T | 370.1609 | 185.5841 | | | 352.1503 | 176.5788 | 3 |
| 12 | 1276.6896 | 638.8484 | 1259.6630 | 630.3352 | 1258.6790 | 629.8431 | Y | 269.1132 | 135.0602 | | | 251.1026 | 126.0550 | 2 |
| 13 | | | | | | | S | 106.0499 | 53.5286 | | | 88.0393 | 44.5233 | 1 |

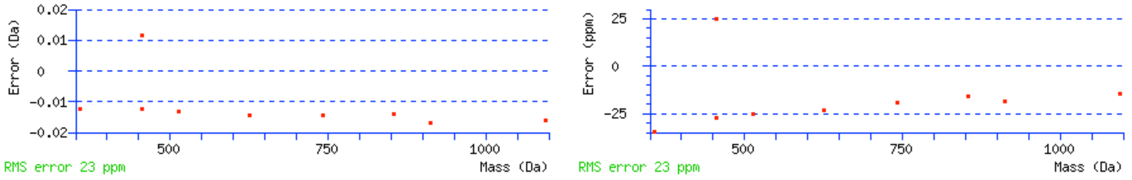


2. F031237. POR1 (1 peptide spectrum)

MS/MS Fragmentation of **SPPVYSDISR**
Found in **VDAC1_YEAST** in **Uniprot_S288C_yeast**, sp|P04840|VDAC1_YEAST Mitochondrial outer membrane protein porin 1 OS=Saccharomyces cerevisiae (strain ATCC 204508 / S288c) OX=559292 GN=POR1 PE=1 SV=4

Match to Query 3995: 1161.602428 from(581.808490,2+) intensity(924077.2500) rtinseconds(2570.56728) index(2787)
Title: qExactive02199_+2.8490.8490.2
Data file D:\Panyue\Proteomic dataset\enriched_fr.+2.mzML

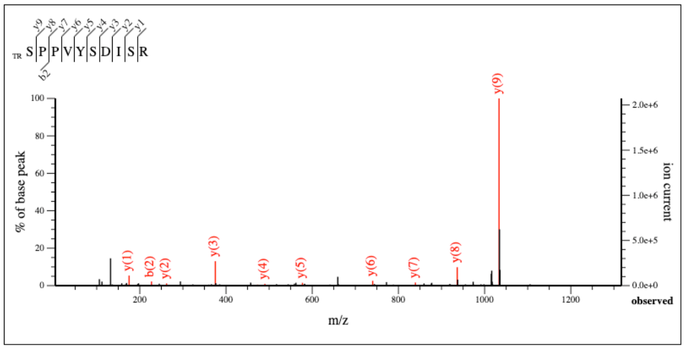


**Monoisotopic mass of neutral peptide Mr(calc):** 1161.6030
**Fixed modifications:** Carbamidomethyl (C) (apply to specified residues or termini only)
**Variable modifications:**
N-term : Trimethyl (Protein N-term)
**Ions Score:** 53 **Expect:** 1.4e-05
**Matches :** 11/88 fragment ions using 31 most intense peaks (help)

| # | b | b++ | b0 | b0++ | Seq. | y | y++ | y* | y*++ | y0 | y0++ | # |
|---|---|---|---|---|---|---|---|---|---|---|---|---|
| 1 | 130.0863 | 65.5468 | 112.0757 | 56.5415 | S | | | | | | | 10 |
| 2 | 227.1390 | 114.0731 | 209.1285 | 105.0679 | P | 1033.5313 | 517.2693 | 1016.5047 | 508.7560 | 1015.5207 | 508.2640 | 9 |
| 3 | 324.1918 | 162.5995 | 306.1812 | 153.5942 | P | 936.4785 | 468.7429 | 919.4520 | 460.2296 | 918.4680 | 459.7376 | 8 |
| 4 | 423.2602 | 212.1337 | 405.2496 | 203.1285 | V | 839.4258 | 420.2165 | 822.3992 | 411.7032 | 821.4152 | 411.2112 | 7 |
| 5 | 586.3235 | 293.6654 | 568.3130 | 284.6601 | Y | 740.3573 | 370.6823 | 723.3308 | 362.1690 | 722.3468 | 361.6770 | 6 |
| 6 | 673.3556 | 337.1814 | 655.3450 | 328.1761 | S | 577.2940 | 289.1506 | 560.2675 | 280.6374 | 559.2835 | 280.1454 | 5 |
| 7 | 788.3825 | 394.6949 | 770.3719 | 385.6896 | D | 490.2620 | 245.6346 | 473.2354 | 237.1214 | 472.2514 | 236.6293 | 4 |
| 8 | 901.4666 | 451.2369 | 883.4560 | 442.2316 | I | 375.2350 | 188.1212 | 358.2085 | 179.6079 | 357.2245 | 179.1159 | 3 |
| 9 | 988.4986 | 494.7529 | 970.4880 | 485.7476 | S | 262.1510 | 131.5791 | 245.1244 | 123.0659 | 244.1404 | 122.5738 | 2 |
| 10 | | | | | R | 175.1190 | 88.0631 | 158.0924 | 79.5498 | | | 1 |

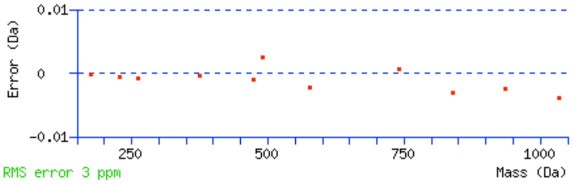


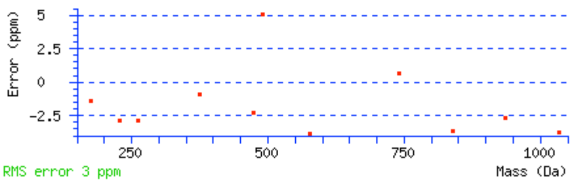


3. F001805. GPM1 (1 peptide spectrum), Rpl15A (1 peptide spectrum) and Rpl16B (1 peptide spectrum)

MS/MS Fragmentation of **PKLVLVR**
Found in **PMG1_YEAST** in **Uniprot_S288C_yeast**, sp|P00950|PMG1_YEAST Phosphoglycerate mutase 1 OS=Saccharomyces cerevisiae (strain ATCC 204508 / S288c) OX=559292 GN=GPM1 PE=1 SV=3

Match to Query 676: 879.626280 from(440.820416,2+) intensity(4672582.5000) rtinseconds(1823.72016) index(4995)
Title: qExactive02200_+3.6431.6431.2
Data file D:\Panyue\enriched_fr.+3.mzML

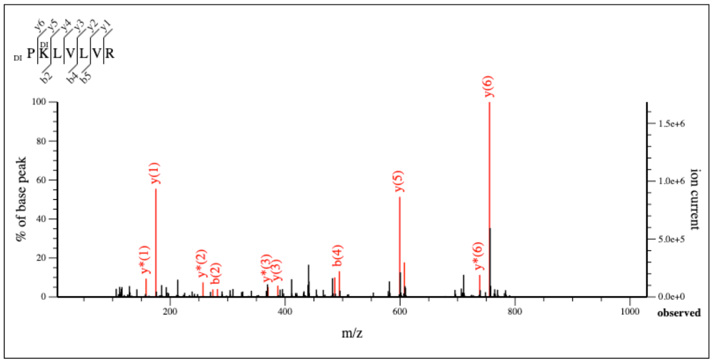


**Monoisotopic mass of neutral peptide Mr(calc):** 879.6269
**Fixed modifications:** Carbamidomethyl (C) (apply to specified residues or termini only)
**Variable modifications:**
N-term : Dimethyl (Protein N-term P)
K2 : Dimethyl (K)
**Ions Score:** 36 **Expect:** 0.00048
**Matches :** 13/46 fragment ions using 28 most intense peaks (help)

| # | b | b++ | b* | b*++ | Seq. | y | y++ | y* | y*++ | # |
|---|---|---|---|---|---|---|---|---|---|---|
| 1 | 126.0913 | 63.5493 | | | P | | | | | 7 |
| 2 | **282.2176** | 141.6124 | 265.1911 | 133.0992 | K | ***755.5502*** | 378.2787 | 738.5236 | 369.7654 | 6 |
| 3 | 395.3017 | 198.1545 | 378.2751 | 189.6412 | L | ***599.4239*** | 300.2156 | 582.3974 | 291.7023 | 5 |
| 4 | **494.3701** | 247.6887 | 477.3435 | 239.1754 | V | ***486.3398*** | 243.6736 | 469.3133 | 235.1603 | 4 |
| 5 | **607.4541** | 304.2307 | 590.4276 | 295.7174 | L | ***387.2714*** | 194.1394 | 370.2449 | 185.6261 | 3 |
| 6 | 706.5226 | 353.7649 | 689.4960 | 345.2516 | V | ***274.1874*** | 137.5973 | 257.1608 | 129.0840 | 2 |
| 7 | | | | | R | ***175.1190*** | 88.0631 | 158.0924 | 79.5498 | 1 |

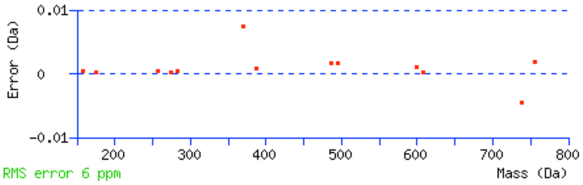


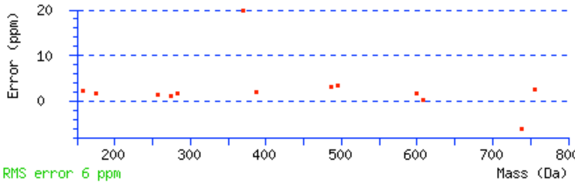


MS/MS Fragmentation of **GAYKYLEELQR**
Found in **RL15A_YEAST** in **Uniprot_S288C_yeast**, sp|P05748|RL15A_YEAST 60S ribosomal protein L15-A OS=Saccharomyces cerevisiae (strain ATCC 204508 / S288c) OX=559292 GN=RPL15A PE=1 SV=3

Match to Query 6623: 1410.750434 from(706.382493,2+) intensity(3035247.0000) rtinseconds(3106.83888) index(8973)
Title: qExactive02200_+3.10881.10881.2
Data file D:\Panyue\enriched_fr.+3.mzML

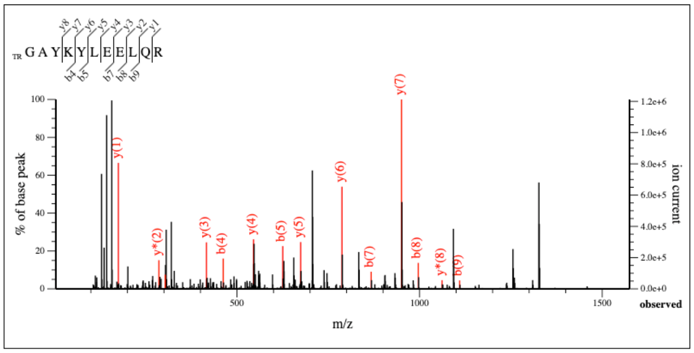

**Monoisotopic mass of neutral peptide Mr(calc):** 1410.7507
**Fixed modifications:** Carbamidomethyl (C) (apply to specified residues or termini only)
**Variable modifications:**
N-term : Trimethyl (Protein N-term)
**Ions Score:** 25 **Expect:** 0.12
**Matches :** 14/96 fragment ions using 39 most intense peaks (help)

| # | b | $b^{++}$ | b* | $b^{*++}$ | $b^0$ | $b^{0++}$ | Seq. | y | $y^{++}$ | y* | $y^{*++}$ | $y^0$ | $y^{0++}$ | # |
|---|---|---|---|---|---|---|---|---|---|---|---|---|---|---|
| 1 | 100.0757 | 50.5415 | | | | | G | | | | | | | 11 |
| 2 | 171.1128 | 86.0600 | | | | | A | 1312.6896 | 656.8484 | 1295.6630 | 648.3352 | 1294.6790 | 647.8431 | 10 |
| 3 | 334.1761 | 167.5917 | | | | | Y | 1241.6525 | 621.3299 | 1224.6259 | 612.8166 | 1223.6419 | 612.3246 | 9 |
| 4 | **462.2711** | 231.6392 | 445.2445 | 223.1259 | | | K | 1078.5891 | 539.7982 | 1061.5626 | 531.2849 | 1060.5786 | 530.7929 | 8 |
| 5 | **625.3344** | 313.1709 | 608.3079 | 304.6576 | | | Y | ***950.4942*** | 475.7507 | 933.4676 | 467.2374 | 932.4836 | 466.7454 | 7 |
| 6 | 738.4185 | 369.7129 | 721.3919 | 361.1996 | | | L | ***787.4308*** | 394.2191 | 770.4043 | 385.7058 | 769.4203 | 385.2138 | 6 |
| 7 | **867.4611** | 434.2342 | 850.4345 | 425.7209 | 849.4505 | 425.2289 | E | ***674.3468*** | 337.6770 | 657.3202 | 329.1638 | 656.3362 | 328.6717 | 5 |
| 8 | **996.5037** | 498.7555 | 979.4771 | 490.2422 | 978.4931 | 489.7502 | E | ***545.3042*** | 273.1557 | 528.2776 | 264.6425 | 527.2936 | 264.1504 | 4 |
| 9 | **1109.5877** | 555.2975 | 1092.5612 | 546.7842 | 1091.5772 | 546.2922 | L | ***416.2616*** | 208.6344 | 399.2350 | 200.1212 | | | 3 |
| 10 | 1237.6463 | 619.3268 | 1220.6198 | 610.8135 | 1219.6358 | 610.3215 | Q | ***303.1775*** | 152.0924 | 286.1510 | 143.5791 | | | 2 |
| 11 | | | | | | | R | ***175.1190*** | 88.0631 | 158.0924 | 79.5498 | | | 1 |

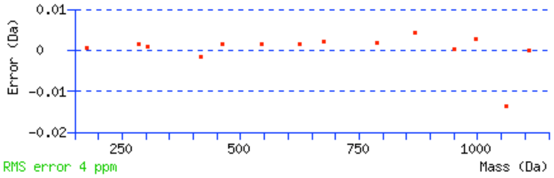


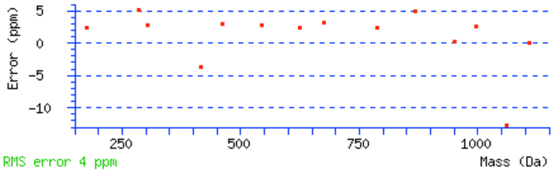


MS/MS Fragmentation of **SQPVVVIDAKDHLLGR**
Found in **RL16B_YEAST** in **Uniprot_S288C_yeast**, sp|P26785|RL16B_YEAST 60S ribosomal protein L16-B OS=Saccharomyces cerevisiae (strain ATCC 204508 / S288c) OX=559292 GN=RPL16B PE=1 SV=3

Match to Query 10140: 1816.048449 from(606.356759,3+) intensity(800303.0625) rtinseconds(5370.44106) index(15131)
Title: qExactive02200_+3.18777.18777.3
Data file D:\Panyue\enriched_fr.+3.mzML

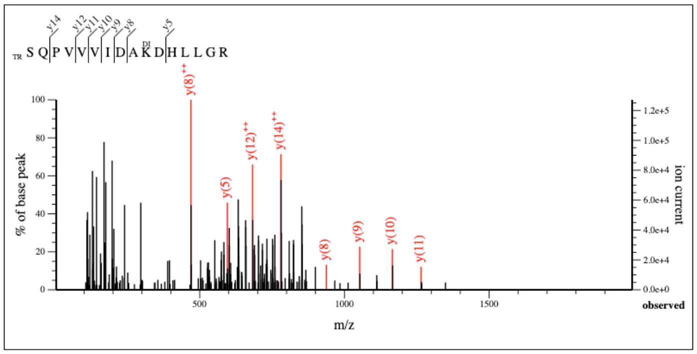

**Monoisotopic mass of neutral peptide Mr(calc):** 1816.0571
**Fixed modifications:** Carbamidomethyl (C) (apply to specified residues or termini only)
**Variable modifications:**
N-term : Trimethyl (Protein N-term)
K10 : Dimethyl (K)
**Ions Score:** 26 **Expect:** 0.15
**Matches :** 8/168 fragment ions using 13 most intense peaks (help)

| # | b | $b^{++}$ | $b^{*}$ | $b^{*++}$ | $b^{0}$ | $b^{0++}$ | Seq. | y | $y^{++}$ | $y^{*}$ | $y^{*++}$ | $y^{0}$ | $y^{0++}$ | # |
|---|---|---|---|---|---|---|---|---|---|---|---|---|---|---|
| 1 | 130.0863 | 65.5468 | | | 112.0757 | 56.5415 | S | | | | | | | 16 |
| 2 | 258.1448 | 129.5761 | 241.1183 | 121.0628 | 240.1343 | 120.5708 | Q | 1687.9854 | 844.4963 | 1670.9588 | 835.9830 | 1669.9748 | 835.4910 | 15 |
| 3 | 355.1976 | 178.1024 | 338.1710 | 169.5892 | 337.1870 | 169.0972 | P | 1559.9268 | **780.4670** | 1542.9002 | 771.9538 | 1541.9162 | 771.4618 | 14 |
| 4 | 454.2660 | 227.6366 | 437.2395 | 219.1234 | 436.2554 | 218.6314 | V | 1462.8740 | 731.9407 | 1445.8475 | 723.4274 | 1444.8635 | 722.9354 | 13 |
| 5 | 553.3344 | 277.1709 | 536.3079 | 268.6576 | 535.3239 | 268.1656 | V | 1363.8056 | **682.4064** | 1346.7791 | 673.8932 | 1345.7950 | 673.4012 | 12 |
| 6 | 652.4028 | 326.7051 | 635.3763 | 318.1918 | 634.3923 | 317.6998 | V | **1264.7372** | 632.8722 | 1247.7106 | 624.3590 | 1246.7266 | 623.8670 | 11 |
| 7 | 765.4869 | 383.2471 | 748.4604 | 374.7338 | 747.4763 | 374.2418 | I | **1165.6688** | 583.3380 | 1148.6422 | 574.8248 | 1147.6582 | 574.3327 | 10 |
| 8 | 880.5138 | 440.7606 | 863.4873 | 432.2473 | 862.5033 | 431.7553 | D | **1052.5847** | 526.7960 | 1035.5582 | 518.2827 | 1034.5742 | 517.7907 | 9 |
| 9 | 951.5510 | 476.2791 | 934.5244 | 467.7658 | 933.5404 | 467.2738 | A | **937.5578** | **469.2825** | 920.5312 | 460.7693 | 919.5472 | 460.2772 | 8 |
| 10 | 1107.6772 | 554.3422 | 1090.6507 | 545.8290 | 1089.6667 | 545.3370 | K | 866.5207 | 433.7640 | 849.4941 | 425.2507 | 848.5101 | 424.7587 | 7 |
| 11 | 1222.7042 | 611.8557 | 1205.6776 | 603.3424 | 1204.6936 | 602.8504 | D | 710.3944 | 355.7008 | 693.3678 | 347.1876 | 692.3838 | 346.6956 | 6 |
| 12 | 1359.7631 | 680.3852 | 1342.7365 | 671.8719 | 1341.7525 | 671.3799 | H | **595.3675** | 298.1874 | 578.3409 | 289.6741 | | | 5 |
| 13 | 1472.8471 | 736.9272 | 1455.8206 | 728.4139 | 1454.8366 | 727.9219 | L | 458.3085 | 229.6579 | 441.2820 | 221.1446 | | | 4 |
| 14 | 1585.9312 | 793.4692 | 1568.9047 | 784.9560 | 1567.9206 | 784.4640 | L | 345.2245 | 173.1159 | 328.1979 | 164.6026 | | | 3 |
| 15 | 1642.9527 | 821.9800 | 1625.9261 | 813.4667 | 1624.9421 | 812.9747 | G | 232.1404 | 116.5738 | 215.1139 | 108.0606 | | | 2 |
| 16 | | | | | | | R | 175.1190 | 88.0631 | 158.0924 | 79.5498 | | | 1 |

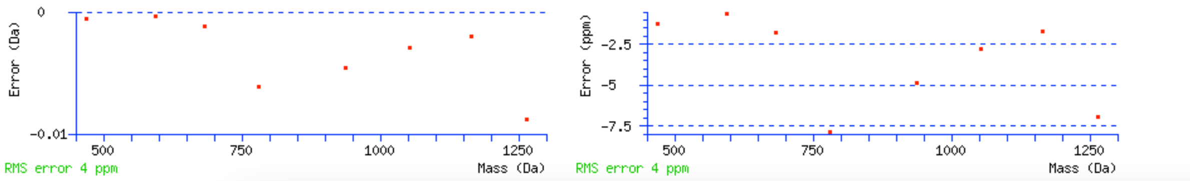


4. F031241. HOM2 (2 spectra), Rpl12A (5 peptide spectra), Rpl24A (2 peptide spectra), Rpl24B (2 peptide spectra) and Rpl26A (6 peptide spectra)

MS/MS Fragmentation of **AGKKIAGVLGATGSVGQR**
Found in **DHAS_YEAST** in **Uniprot_S288C_yeast**, sp|P13663|DHAS_YEAST Aspartate-semialdehyde dehydrogenase OS=Saccharomyces cerevisiae (strain ATCC 204508 / S288c) OX=559292 GN=HOM2 PE=1 SV=1

Match to Query 5536: 1739.041395 from(580.687741,3+) intensity(3023507.2500) rtinseconds(1942.43448) index(5140)
Title: qExactive02201_+4.6923.6923.3
Data file D:\Panyue\enriched_fr.+4.mzML

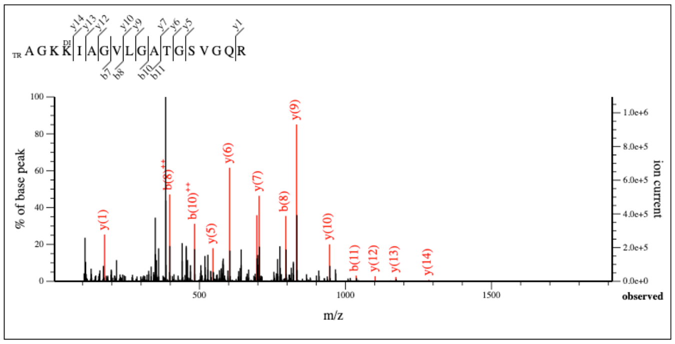

**Monoisotopic mass of neutral peptide Mr(calc):** 1739.0417
**Fixed modifications:** Carbamidomethyl (C) (apply to specified residues or termini only)
**Variable modifications:**
N-term : Trimethyl (Protein N-term)
K4 : Dimethyl (K)
**Ions Score:** 56 **Expect:** 0.0001
**Matches :** 14/170 fragment ions using 24 most intense peaks (help)

| # | b | $b^{++}$ | $b^{*}$ | $b^{*++}$ | $b^{0}$ | $b^{0++}$ | Seq. | y | $y^{++}$ | $y^{*}$ | $y^{*++}$ | $y^{0}$ | $y^{0++}$ | # |
|---|---|---|---|---|---|---|---|---|---|---|---|---|---|---|
| 1 | 114.0913 | 57.5493 | | | | | A | | | | | | | 18 |
| 2 | 171.1128 | 86.0600 | | | | | G | 1626.9650 | 813.9861 | 1609.9384 | 805.4728 | 1608.9544 | 804.9808 | 17 |
| 3 | 299.2078 | 150.1075 | 282.1812 | 141.5942 | | | K | 1569.9435 | 785.4754 | 1552.9170 | 776.9621 | 1551.9329 | 776.4701 | 16 |
| 4 | 455.3340 | 228.1707 | 438.3075 | 219.6574 | | | K | 1441.8485 | 721.4279 | 1424.8220 | 712.9146 | 1423.8380 | 712.4226 | 15 |
| 5 | 568.4181 | 284.7127 | 551.3915 | 276.1994 | | | I | ***1285.7223*** | 643.3648 | 1268.6957 | 634.8515 | 1267.7117 | 634.3595 | 14 |
| 6 | 639.4552 | 320.2312 | 622.4287 | 311.7180 | | | A | ***1172.6382*** | 586.8227 | 1155.6117 | 578.3095 | 1154.6276 | 577.8175 | 13 |
| 7 | **696.4767** | 348.7420 | 679.4501 | 340.2287 | | | G | ***1101.6011*** | 551.3042 | 1084.5745 | 542.7909 | 1083.5905 | 542.2989 | 12 |
| 8 | **795.5451** | **398.2762** | 778.5185 | 389.7629 | | | V | 1044.5796 | 522.7935 | 1027.5531 | 514.2802 | 1026.5691 | 513.7882 | 11 |
| 9 | 908.6292 | 454.8182 | 891.6026 | 446.3049 | | | L | ***945.5112*** | 473.2592 | 928.4847 | 464.7460 | 927.5007 | 464.2540 | 10 |
| 10 | 965.6506 | **483.3289** | 948.6241 | 474.8157 | | | G | ***832.4272*** | 416.7172 | 815.4006 | 408.2039 | 814.4166 | 407.7119 | 9 |
| 11 | **1036.6877** | 518.8475 | 1019.6612 | 510.3342 | | | A | 775.4057 | 388.2065 | 758.3791 | 379.6932 | 757.3951 | 379.2012 | 8 |
| 12 | 1137.7354 | 569.3713 | 1120.7089 | 560.8581 | 1119.7248 | 560.3661 | T | ***704.3686*** | 352.6879 | 687.3420 | 344.1747 | 686.3580 | 343.6826 | 7 |
| 13 | 1194.7569 | 597.8821 | 1177.7303 | 589.3688 | 1176.7463 | 588.8768 | G | ***603.3209*** | 302.1641 | 586.2944 | 293.6508 | 585.3103 | 293.1588 | 6 |
| 14 | 1281.7889 | 641.3981 | 1264.7623 | 632.8848 | 1263.7783 | 632.3928 | S | ***546.2994*** | 273.6534 | 529.2729 | 265.1401 | 528.2889 | 264.6481 | 5 |
| 15 | 1380.8573 | 690.9323 | 1363.8308 | 682.4190 | 1362.8467 | 681.9270 | V | 459.2674 | 230.1373 | 442.2409 | 221.6241 | | | 4 |
| 16 | 1437.8788 | 719.4430 | 1420.8522 | 710.9298 | 1419.8682 | 710.4377 | G | 360.1990 | 180.6031 | 343.1724 | 172.0899 | | | 3 |
| 17 | 1565.9374 | 783.4723 | 1548.9108 | 774.9590 | 1547.9268 | 774.4670 | Q | 303.1775 | 152.0924 | 286.1510 | 143.5791 | | | 2 |
| 18 | | | | | | | R | ***175.1190*** | 88.0631 | 158.0924 | 79.5498 | | | 1 |

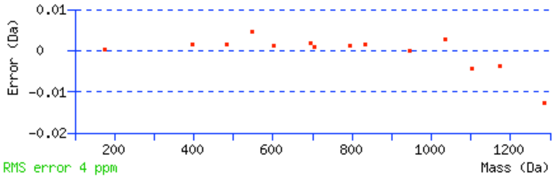


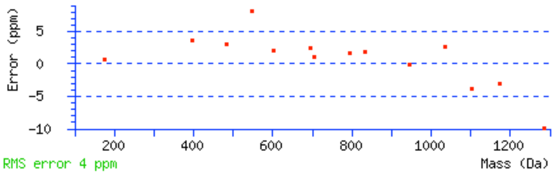


MS/MS Fragmentation of **AGKKIAGVLGATGSVGQR**
Found in **DHAS_YEAST** in **Uniprot_S288C_yeast**, sp|P13663|DHAS_YEAST Aspartate-semialdehyde dehydrogenase OS=Saccharomyces cerevisiae (strain ATCC 204508 / S288c) OX=559292 GN=HOM2 PE=1 SV=1

Match to Query 5537: 1739.043279 from(580.688369,3+) intensity(722618.5000) rtinseconds(1926.96348) index(5092)
Title: qExactive02201_+4.6866.6866.3
Data file D:\Panyue\enriched_fr.+4.mzML

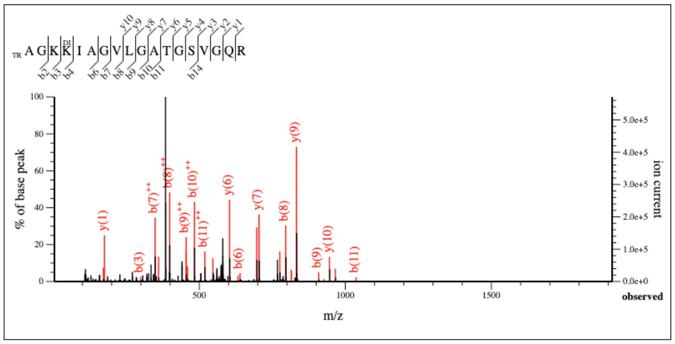

```
Monoisotopic mass of neutral peptide Mr(calc): 1739.0417
Fixed modifications: Carbamidomethyl (C) (apply to specified residues or termini only)
Variable modifications:
  N-term : Trimethyl (Protein N-term)
  K4     : Dimethyl (K)
Ions Score: 54  Expect: 0.00017
Matches : 28/170 fragment ions using 64 most intense peaks   (help)
```

| # | b | b++ | b* | b*++ | b0 | b0++ | Seq. | y | y++ | y* | y*++ | y0 | y0++ | # |
|---|---|---|---|---|---|---|---|---|---|---|---|---|---|---|
| 1 | 114.0913 | 57.5493 | | | | | A | | | | | | | 18 |
| 2 | 171.1128 | 86.0600 | | | | | G | 1626.9650 | 813.9861 | 1609.9384 | 805.4728 | 1608.9544 | 804.9808 | 17 |
| 3 | 299.2078 | 150.1075 | 282.1812 | 141.5942 | | | K | 1569.9435 | 785.4754 | 1552.9170 | 776.9621 | 1551.9329 | 776.4701 | 16 |
| 4 | 455.3340 | 228.1707 | 438.3075 | 219.6574 | | | K | 1441.8485 | 721.4279 | 1424.8220 | 712.9146 | 1423.8380 | 712.4226 | 15 |
| 5 | 568.4181 | 284.7127 | 551.3915 | 276.1994 | | | I | 1285.7223 | 643.3648 | 1268.6957 | 634.8515 | 1267.7117 | 634.3595 | 14 |
| 6 | 639.4552 | 320.2312 | 622.4287 | 311.7180 | | | A | 1172.6382 | 586.8227 | 1155.6117 | 578.3095 | 1154.6276 | 577.8175 | 13 |
| 7 | 696.4767 | 348.7420 | 679.4501 | 340.2287 | | | G | 1101.6011 | 551.3042 | 1084.5745 | 542.7909 | 1083.5905 | 542.2989 | 12 |
| 8 | 795.5451 | 398.2762 | 778.5185 | 389.7629 | | | V | 1044.5796 | 522.7935 | 1027.5531 | 514.2802 | 1026.5691 | 513.7882 | 11 |
| 9 | 908.6292 | 454.8182 | 891.6026 | 446.3049 | | | L | 945.5112 | 473.2592 | 928.4847 | 464.7460 | 927.5007 | 464.2540 | 10 |
| 10 | 965.6506 | 483.3289 | 948.6241 | 474.8157 | | | G | 832.4272 | 416.7172 | 815.4006 | 408.2039 | 814.4166 | 407.7119 | 9 |
| 11 | 1036.6877 | 518.8475 | 1019.6612 | 510.3342 | | | A | 775.4057 | 388.2065 | 758.3791 | 379.6932 | 757.3951 | 379.2012 | 8 |
| 12 | 1137.7354 | 569.3713 | 1120.7089 | 560.8581 | 1119.7248 | 560.3661 | T | 704.3686 | 352.6879 | 687.3420 | 344.1747 | 686.3580 | 343.6826 | 7 |
| 13 | 1194.7569 | 597.8821 | 1177.7303 | 589.3688 | 1176.7463 | 588.8768 | G | 603.3209 | 302.1641 | 586.2944 | 293.6508 | 585.3103 | 293.1588 | 6 |
| 14 | 1281.7889 | 641.3981 | 1264.7623 | 632.8848 | 1263.7783 | 632.3928 | S | 546.2994 | 273.6534 | 529.2729 | 265.1401 | 528.2889 | 264.6481 | 5 |
| 15 | 1380.8573 | 690.9323 | 1363.8308 | 682.4190 | 1362.8467 | 681.9270 | V | 459.2674 | 230.1373 | 442.2409 | 221.6241 | | | 4 |
| 16 | 1437.8788 | 719.4430 | 1420.8522 | 710.9298 | 1419.8682 | 710.4377 | G | 360.1990 | 180.6031 | 343.1724 | 172.0899 | | | 3 |
| 17 | 1565.9374 | 783.4723 | 1548.9108 | 774.9590 | 1547.9268 | 774.4670 | Q | 303.1775 | 152.0924 | 286.1510 | 143.5791 | | | 2 |
| 18 | | | | | | | R | 175.1190 | 88.0631 | 158.0924 | 79.5498 | | | 1 |

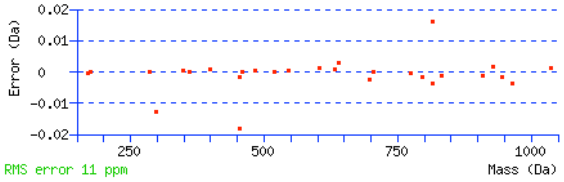


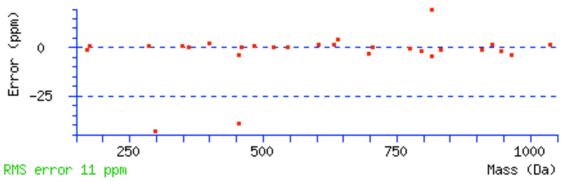


MS/MS Fragmentation of **PPKFDPNEVKYLYLR**
Found in **RL12A_YEAST** in **Uniprot_S288C_yeast**, sp|P0CX53|RL12A_YEAST 60S ribosomal protein L12-A OS=Saccharomyces cerevisiae (strain ATCC 204508 / S288c) OX=559292 GN=RPL12A PE=1 SV=1

Match to Query 7311: 1976.113776 from(495.035720,4+) intensity(814931.1250) rtinseconds(4515.52044) index(9577)
Title: qExactive02201_+4.16143.16143.4
Data file D:\Panyue\enriched_fr.+4.mzML

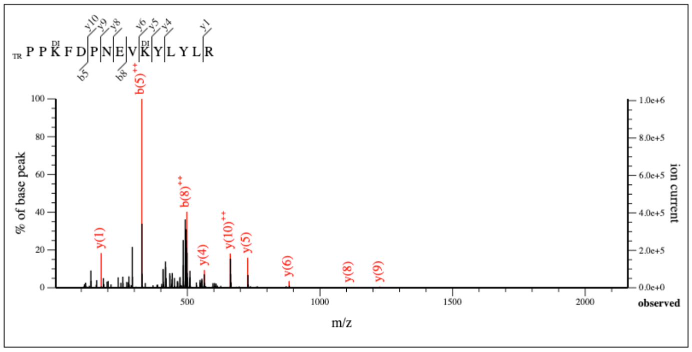

```
Monoisotopic mass of neutral peptide Mr(calc): 1976.1135
Fixed modifications: Carbamidomethyl (C) (apply to specified residues or termini only)
Variable modifications:
  N-term : Trimethyl (Protein N-term)
  K3     : Dimethyl (K)
  K10    : Dimethyl (K)
Ions Score: 44  Expect: 0.0032
Matches : 9/142 fragment ions using 11 most intense peaks   (help)
```

| # | b | b++ | b* | b*++ | b0 | b0++ | Seq. | y | y++ | y* | y*++ | y0 | y0++ | # |
|---|---|---|---|---|---|---|---|---|---|---|---|---|---|---|
| 1 | 140.1070 | 70.5571 | | | | | P | | | | | | | 15 |
| 2 | 237.1598 | 119.0835 | | | | | P | 1838.0211 | 919.5142 | 1820.9945 | 911.0009 | 1820.0105 | 910.5089 | 14 |
| 3 | 393.2860 | 197.1466 | 376.2595 | 188.6334 | | | K | 1740.9683 | 870.9878 | 1723.9418 | 862.4745 | 1722.9578 | 861.9825 | 13 |
| 4 | 540.3544 | 270.6809 | 523.3279 | 262.1676 | | | F | 1584.8421 | 792.9247 | 1567.8155 | 784.4114 | 1566.8315 | 783.9194 | 12 |
| 5 | 655.3814 | **328.1943** | 638.3548 | 319.6811 | 637.3708 | 319.1890 | D | 1437.7736 | 719.3905 | 1420.7471 | 710.8772 | 1419.7631 | 710.3852 | 11 |
| 6 | 752.4341 | 376.7207 | 735.4076 | 368.2074 | 734.4236 | 367.7154 | P | 1322.7467 | **661.8770** | 1305.7201 | 653.3637 | 1304.7361 | 652.8717 | 10 |
| 7 | 866.4771 | 433.7422 | 849.4505 | 425.2289 | 848.4665 | 424.7369 | N | ***1225.6939*** | 613.3506 | 1208.6674 | 604.8373 | 1207.6834 | 604.3453 | 9 |
| 8 | 995.5197 | **498.2635** | 978.4931 | 489.7502 | 977.5091 | 489.2582 | E | ***1111.6510*** | 556.3291 | 1094.6245 | 547.8159 | 1093.6404 | 547.3239 | 8 |
| 9 | 1094.5881 | 547.7977 | 1077.5615 | 539.2844 | 1076.5775 | 538.7924 | V | 982.6084 | 491.8078 | 965.5819 | 483.2946 | | | 7 |
| 10 | 1250.7143 | 625.8608 | 1233.6878 | 617.3475 | 1232.7038 | 616.8555 | K | ***883.5400*** | 442.2736 | 866.5135 | 433.7604 | | | 6 |
| 11 | 1413.7777 | 707.3925 | 1396.7511 | 698.8792 | 1395.7671 | 698.3872 | Y | ***727.4137*** | 364.2105 | 710.3872 | 355.6972 | | | 5 |
| 12 | 1526.8617 | 763.9345 | 1509.8352 | 755.4212 | 1508.8512 | 754.9292 | L | ***564.3504*** | 282.6788 | 547.3239 | 274.1656 | | | 4 |
| 13 | 1689.9251 | 845.4662 | 1672.8985 | 836.9529 | 1671.9145 | 836.4609 | Y | 451.2663 | 226.1368 | 434.2398 | 217.6235 | | | 3 |
| 14 | 1803.0091 | 902.0082 | 1785.9826 | 893.4949 | 1784.9986 | 893.0029 | L | 288.2030 | 144.6051 | 271.1765 | 136.0919 | | | 2 |
| 15 | | | | | | | R | ***175.1190*** | 88.0631 | 158.0924 | 79.5498 | | | 1 |

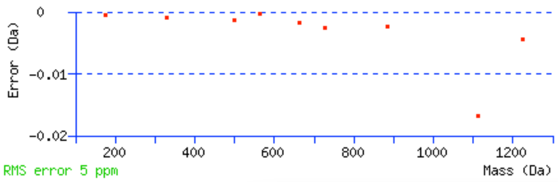


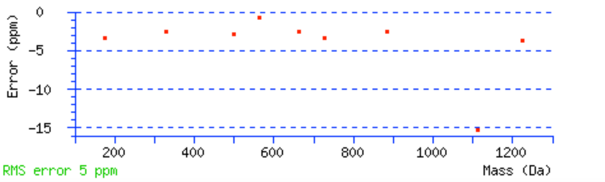


MS/MS Fragmentation of **PPKFDPNEVKYLYLR**
Found in **RL12A_YEAST** in **Uniprot_S288C_yeast**, sp|P0CX53|RL12A_YEAST 60S ribosomal protein L12-A OS=Saccharomyces cerevisiae (strain ATCC 204508 / S288c) OX=559292 GN=RPL12A PE=1 SV=1

Match to Query 7314: 1976.114361 from(659.712063,3+) intensity(4635530.0000) rtinseconds(4513.33662) index(9576)
Title: qExactive02201_+4.16136.16136.3
Data file D:\Panyue\enriched_fr.+4.mzML

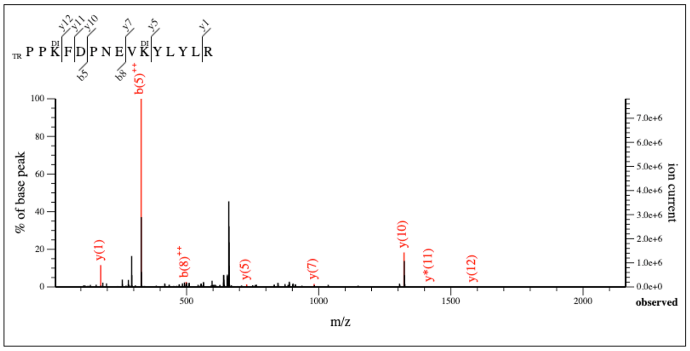

**Monoisotopic mass of neutral peptide Mr(calc):** 1976.1135
**Fixed modifications:** Carbamidomethyl (C) (apply to specified residues or termini only)
**Variable modifications:**
N-term : Trimethyl (Protein N-term)
K3 : Dimethyl (K)
K10 : Dimethyl (K)
**Ions Score:** 22 **Expect:** 0.45
**Matches :** 8/142 fragment ions using 15 most intense peaks (help)

| # | b | b++ | b* | b*++ | b0 | b0++ | Seq. | y | y++ | y* | y*++ | y0 | y0++ | # |
|---|---|---|---|---|---|---|---|---|---|---|---|---|---|---|
| 1 | 140.1070 | 70.5571 | | | | | P | | | | | | | 15 |
| 2 | 237.1598 | 119.0835 | | | | | P | 1838.0211 | 919.5142 | 1820.9945 | 911.0009 | 1820.0105 | 910.5089 | 14 |
| 3 | 393.2860 | 197.1466 | 376.2595 | 188.6334 | | | K | 1740.9683 | 870.9878 | 1723.9418 | 862.4745 | 1722.9578 | 861.9825 | 13 |
| 4 | 540.3544 | 270.6809 | 523.3279 | 262.1676 | | | F | 1584.8421 | 792.9247 | 1567.8155 | 784.4114 | 1566.8315 | 783.9194 | 12 |
| 5 | 655.3814 | 328.1943 | 638.3548 | 319.6811 | 637.3708 | 319.1890 | D | 1437.7736 | 719.3905 | 1420.7471 | 710.8772 | 1419.7631 | 710.3852 | 11 |
| 6 | 752.4341 | 376.7207 | 735.4076 | 368.2074 | 734.4236 | 367.7154 | P | 1322.7467 | 661.8770 | 1305.7201 | 653.3637 | 1304.7361 | 652.8717 | 10 |
| 7 | 866.4771 | 433.7422 | 849.4505 | 425.2289 | 848.4665 | 424.7369 | N | 1225.6939 | 613.3506 | 1208.6674 | 604.8373 | 1207.6834 | 604.3453 | 9 |
| 8 | 995.5197 | 498.2635 | 978.4931 | 489.7502 | 977.5091 | 489.2582 | E | 1111.6510 | 556.3291 | 1094.6245 | 547.8159 | 1093.6404 | 547.3239 | 8 |
| 9 | 1094.5881 | 547.7977 | 1077.5615 | 539.2844 | 1076.5775 | 538.7924 | V | 982.6084 | 491.8078 | 965.5819 | 483.2946 | | | 7 |
| 10 | 1250.7143 | 625.8608 | 1233.6878 | 617.3475 | 1232.7038 | 616.8555 | K | 883.5400 | 442.2736 | 866.5135 | 433.7604 | | | 6 |
| 11 | 1413.7777 | 707.3925 | 1396.7511 | 698.8792 | 1395.7671 | 698.3872 | Y | 727.4137 | 364.2105 | 710.3872 | 355.6972 | | | 5 |
| 12 | 1526.8617 | 763.9345 | 1509.8352 | 755.4212 | 1508.8512 | 754.9292 | L | 564.3504 | 282.6788 | 547.3239 | 274.1656 | | | 4 |
| 13 | 1689.9251 | 845.4662 | 1672.8985 | 836.9529 | 1671.9145 | 836.4609 | Y | 451.2663 | 226.1368 | 434.2398 | 217.6235 | | | 3 |
| 14 | 1803.0091 | 902.0082 | 1785.9826 | 893.4949 | 1784.9986 | 893.0029 | L | 288.2030 | 144.6051 | 271.1765 | 136.0919 | | | 2 |
| 15 | | | | | | | R | 175.1190 | 88.0631 | 158.0924 | 79.5498 | | | 1 |

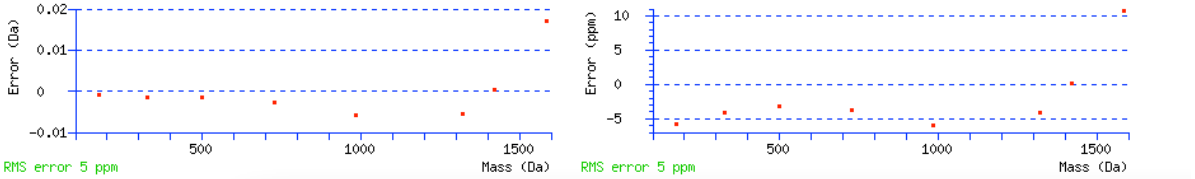


MS/MS Fragmentation of PPKFDPNEVKYLYLR
Found in RL12A_YEAST in Uniprot_S288C_yeast, sp|P0CX53|RL12A_YEAST 60S ribosomal protein L12-A OS=Saccharomyces cerevisiae (strain ATCC 204508 / S288c) OX=559292 GN=RPL12A PE=1 SV=1

Match to Query 7315: 1976.114744 from(495.035962,4+) intensity(2758850.0000) rtinseconds(4503.10422) index(9573)
Title: qExactive02201_+4.16105.16105.4
Data file D:\Panyue\enriched_fr.+4.mzML

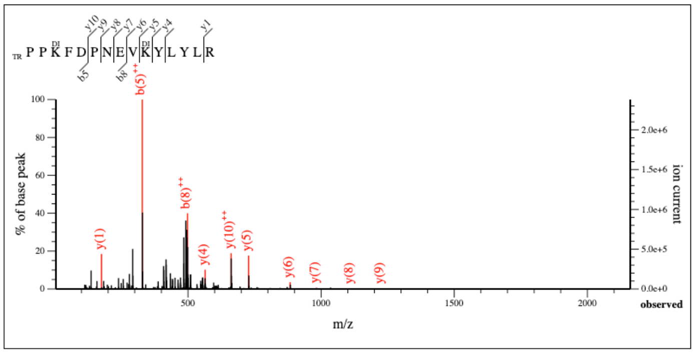

**Monoisotopic mass of neutral peptide Mr(calc):** 1976.1135
**Fixed modifications:** Carbamidomethyl (C) (apply to specified residues or termini only)
**Variable modifications:**
N-term : Trimethyl (Protein N-term)
K3 : Dimethyl (K)
K10 : Dimethyl (K)
**Ions Score:** 55 **Expect:** 0.00026
**Matches :** 10/142 fragment ions using 12 most intense peaks (help)

| # | b | b++ | b* | b*++ | b0 | b0++ | Seq. | y | y++ | y* | y*++ | y0 | y0++ | # |
|---|---|---|---|---|---|---|---|---|---|---|---|---|---|---|
| 1 | 140.1070 | 70.5571 | | | | | P | | | | | | | 15 |
| 2 | 237.1598 | 119.0835 | | | | | P | 1838.0211 | 919.5142 | 1820.9945 | 911.0009 | 1820.0105 | 910.5089 | 14 |
| 3 | 393.2860 | 197.1466 | 376.2595 | 188.6334 | | | K | 1740.9683 | 870.9878 | 1723.9418 | 862.4745 | 1722.9578 | 861.9825 | 13 |
| 4 | 540.3544 | 270.6809 | 523.3279 | 262.1676 | | | F | 1584.8421 | 792.9247 | 1567.8155 | 784.4114 | 1566.8315 | 783.9194 | 12 |
| 5 | 655.3814 | **328.1943** | 638.3548 | 319.6811 | 637.3708 | 319.1890 | D | 1437.7736 | 719.3905 | 1420.7471 | 710.8772 | 1419.7631 | 710.3852 | 11 |
| 6 | 752.4341 | 376.7207 | 735.4076 | 368.2074 | 734.4236 | 367.7154 | P | 1322.7467 | **661.8770** | 1305.7201 | 653.3637 | 1304.7361 | 652.8717 | 10 |
| 7 | 866.4771 | 433.7422 | 849.4505 | 425.2289 | 848.4665 | 424.7369 | N | ***1225.6939*** | 613.3506 | 1208.6674 | 604.8373 | 1207.6834 | 604.3453 | 9 |
| 8 | 995.5197 | **498.2635** | 978.4931 | 489.7502 | 977.5091 | 489.2582 | E | ***1111.6510*** | 556.3291 | 1094.6245 | 547.8159 | 1093.6404 | 547.3239 | 8 |
| 9 | 1094.5881 | 547.7977 | 1077.5615 | 539.2844 | 1076.5775 | 538.7924 | V | ***982.6084*** | 491.8078 | 965.5819 | 483.2946 | | | 7 |
| 10 | 1250.7143 | 625.8608 | 1233.6878 | 617.3475 | 1232.7038 | 616.8555 | K | ***883.5400*** | 442.2736 | 866.5135 | 433.7604 | | | 6 |
| 11 | 1413.7777 | 707.3925 | 1396.7511 | 698.8792 | 1395.7671 | 698.3872 | Y | ***727.4137*** | 364.2105 | 710.3872 | 355.6972 | | | 5 |
| 12 | 1526.8617 | 763.9345 | 1509.8352 | 755.4212 | 1508.8512 | 754.9292 | L | ***564.3504*** | 282.6788 | 547.3239 | 274.1656 | | | 4 |
| 13 | 1689.9251 | 845.4662 | 1672.8985 | 836.9529 | 1671.9145 | 836.4609 | Y | 451.2663 | 226.1368 | 434.2398 | 217.6235 | | | 3 |
| 14 | 1803.0091 | 902.0082 | 1785.9826 | 893.4949 | 1784.9986 | 893.0029 | L | 288.2030 | 144.6051 | 271.1765 | 136.0919 | | | 2 |
| 15 | | | | | | | R | ***175.1190*** | 88.0631 | 158.0924 | 79.5498 | | | 1 |

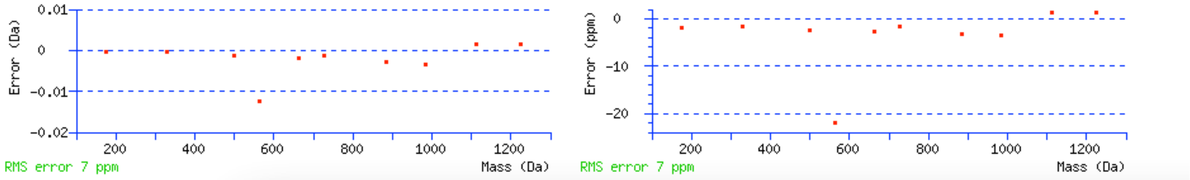


MS/MS Fragmentation of **PPKFDPNEVKYLYLR**
Found in **RL12A_YEAST** in **Uniprot_S288C_yeast**, sp|P0CX53|RL12A_YEAST 60S ribosomal protein L12-A OS=Saccharomyces cerevisiae (strain ATCC 204508 / S288c) OX=559292 GN=RPL12A PE=1 SV=1

Match to Query 7317: 1976.115484 from(495.036147,4+) intensity(10307830.0000) rtinseconds(4490.51304) index(9568)
Title: qExactive02201_+4.16065.16065.4
Data file D:\Panyue\enriched_fr.+4.mzML

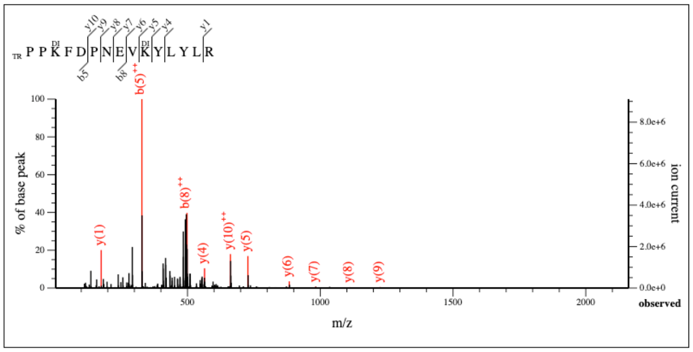

**Monoisotopic mass of neutral peptide Mr(calc):** 1976.1135
**Fixed modifications:** Carbamidomethyl (C) (apply to specified residues or termini only)
**Variable modifications:**
N-term : Trimethyl (Protein N-term)
K3 : Dimethyl (K)
K10 : Dimethyl (K)
**Ions Score:** 51 **Expect:** 0.00058
**Matches :** 10/142 fragment ions using 13 most intense peaks (help)

| # | b | b++ | b* | b*++ | b0 | b0++ | Seq. | y | y++ | y* | y*++ | y0 | y0++ | # |
|---|---|---|---|---|---|---|---|---|---|---|---|---|---|---|
| 1 | 140.1070 | 70.5571 | | | | | P | | | | | | | 15 |
| 2 | 237.1598 | 119.0835 | | | | | P | 1838.0211 | 919.5142 | 1820.9945 | 911.0009 | 1820.0105 | 910.5089 | 14 |
| 3 | 393.2860 | 197.1466 | 376.2595 | 188.6334 | | | K | 1740.9683 | 870.9878 | 1723.9418 | 862.4745 | 1722.9578 | 861.9825 | 13 |
| 4 | 540.3544 | 270.6809 | 523.3279 | 262.1676 | | | F | 1584.8421 | 792.9247 | 1567.8155 | 784.4114 | 1566.8315 | 783.9194 | 12 |
| 5 | 655.3814 | **328.1943** | 638.3548 | 319.6811 | 637.3708 | 319.1890 | D | 1437.7736 | 719.3905 | 1420.7471 | 710.8772 | 1419.7631 | 710.3852 | 11 |
| 6 | 752.4341 | 376.7207 | 735.4076 | 368.2074 | 734.4236 | 367.7154 | P | 1322.7467 | **661.8770** | 1305.7201 | 653.3637 | 1304.7361 | 652.8717 | 10 |
| 7 | 866.4771 | 433.7422 | 849.4505 | 425.2289 | 848.4665 | 424.7369 | N | ***1225.6939*** | 613.3506 | 1208.6674 | 604.8373 | 1207.6834 | 604.3453 | 9 |
| 8 | 995.5197 | **498.2635** | 978.4931 | 489.7502 | 977.5091 | 489.2582 | E | ***1111.6510*** | 556.3291 | 1094.6245 | 547.8159 | 1093.6404 | 547.3239 | 8 |
| 9 | 1094.5881 | 547.7977 | 1077.5615 | 539.2844 | 1076.5775 | 538.7924 | V | ***982.6084*** | 491.8078 | 965.5819 | 483.2946 | | | 7 |
| 10 | 1250.7143 | 625.8608 | 1233.6878 | 617.3475 | 1232.7038 | 616.8555 | K | ***883.5400*** | 442.2736 | 866.5135 | 433.7604 | | | 6 |
| 11 | 1413.7777 | 707.3925 | 1396.7511 | 698.8792 | 1395.7671 | 698.3872 | Y | ***727.4137*** | 364.2105 | 710.3872 | 355.6972 | | | 5 |
| 12 | 1526.8617 | 763.9345 | 1509.8352 | 755.4212 | 1508.8512 | 754.9292 | L | ***564.3504*** | 282.6788 | 547.3239 | 274.1656 | | | 4 |
| 13 | 1689.9251 | 845.4662 | 1672.8985 | 836.9529 | 1671.9145 | 836.4609 | Y | 451.2663 | 226.1368 | 434.2398 | 217.6235 | | | 3 |
| 14 | 1803.0091 | 902.0082 | 1785.9826 | 893.4949 | 1784.9986 | 893.0029 | L | 288.2030 | 144.6051 | 271.1765 | 136.0919 | | | 2 |
| 15 | | | | | | | R | ***175.1190*** | 88.0631 | 158.0924 | 79.5498 | | | 1 |

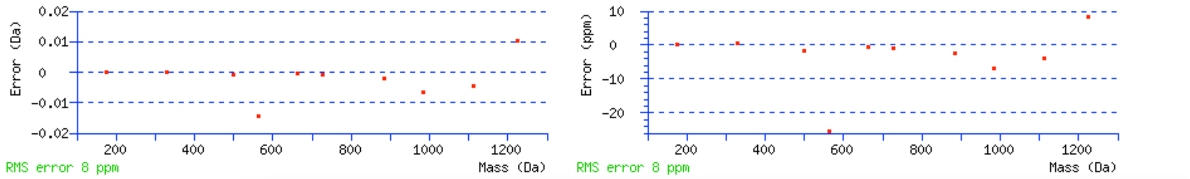


MS/MS Fragmentation of **PPKFDPNEVKYLYLR**
Found in **RL12A_YEAST** in **Uniprot_S288C_yeast**, sp|P0CX53|RL12A_YEAST 60S ribosomal protein L12-A OS=Saccharomyces cerevisiae (strain ATCC 204508 / S288c) OX=559292 GN=RPL12A PE=1 SV=1

Match to Query 7318: 1976.116494 from(659.712774,3+) intensity(17437658.0000) rtinseconds(4500.79248) index(9572)
Title: qExactive02201_+4.16098.16098.3
Data file D:\Panyue\enriched_fr.+4.mzML

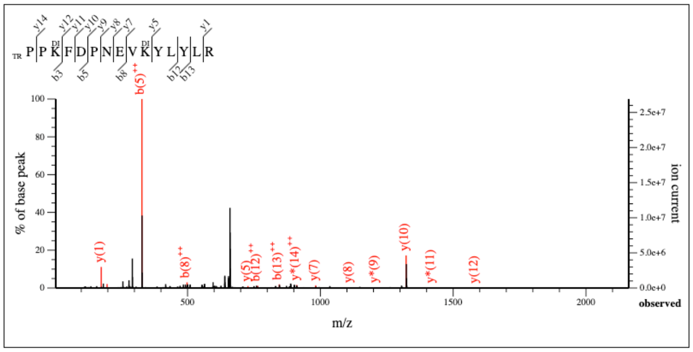

**Monoisotopic mass of neutral peptide Mr(calc):** 1976.1135
**Fixed modifications:** Carbamidomethyl (C) (apply to specified residues or termini only)
**Variable modifications:**
N-term : Trimethyl (Protein N-term)
K3 : Dimethyl (K)
K10 : Dimethyl (K)
**Ions Score:** 22 **Expect:** 0.47
**Matches :** 16/142 fragment ions using 45 most intense peaks (help)

| # | b | b++ | b* | b*++ | b0 | b0++ | Seq. | y | y++ | y* | y*++ | y0 | y0++ | # |
|---|---|---|---|---|---|---|---|---|---|---|---|---|---|---|
| 1 | 140.1070 | 70.5571 | | | | | P | | | | | | | 15 |
| 2 | 237.1598 | 119.0835 | | | | | P | 1838.0211 | 919.5142 | 1820.9945 | 911.0009 | 1820.0105 | 910.5089 | 14 |
| 3 | 393.2860 | ***197.1466*** | 376.2595 | 188.6334 | | | K | 1740.9683 | 870.9878 | 1723.9418 | 862.4745 | 1722.9578 | 861.9825 | 13 |
| 4 | 540.3544 | 270.6809 | 523.3279 | 262.1676 | | | F | ***1584.8421*** | 792.9247 | 1567.8155 | 784.4114 | 1566.8315 | 783.9194 | 12 |
| 5 | 655.3814 | ***328.1943*** | 638.3548 | 319.6811 | 637.3708 | 319.1890 | D | ***1437.7736*** | 719.3905 | 1420.7471 | 710.8772 | 1419.7631 | 710.3852 | 11 |
| 6 | 752.4341 | 376.7207 | 735.4076 | 368.2074 | 734.4236 | 367.7154 | P | ***1322.7467*** | 661.8770 | 1305.7201 | 653.3637 | 1304.7361 | 652.8717 | 10 |
| 7 | 866.4771 | 433.7422 | 849.4505 | 425.2289 | 848.4665 | 424.7369 | N | 1225.6939 | 613.3506 | 1208.6674 | 604.8373 | 1207.6834 | 604.3453 | 9 |
| 8 | 995.5197 | ***498.2635*** | 978.4931 | 489.7502 | 977.5091 | 489.2582 | E | ***1111.6510*** | 556.3291 | 1094.6245 | 547.8159 | 1093.6404 | 547.3239 | 8 |
| 9 | 1094.5881 | 547.7977 | 1077.5615 | 539.2844 | 1076.5775 | 538.7924 | V | ***982.6084*** | 491.8078 | 965.5819 | 483.2946 | | | 7 |
| 10 | 1250.7143 | 625.8608 | 1233.6878 | 617.3475 | 1232.7038 | 616.8555 | K | 883.5400 | 442.2736 | 866.5135 | 433.7604 | | | 6 |
| 11 | 1413.7777 | 707.3925 | 1396.7511 | 698.8792 | 1395.7671 | 698.3872 | Y | ***727.4137*** | 364.2105 | 710.3872 | 355.6972 | | | 5 |
| 12 | 1526.8617 | ***763.9345*** | 1509.8352 | 755.4212 | 1508.8512 | 754.9292 | L | 564.3504 | 282.6788 | 547.3239 | 274.1656 | | | 4 |
| 13 | 1689.9251 | ***845.4662*** | 1672.8985 | 836.9529 | 1671.9145 | 836.4609 | Y | 451.2663 | 226.1368 | 434.2398 | 217.6235 | | | 3 |
| 14 | 1803.0091 | 902.0082 | 1785.9826 | 893.4949 | 1784.9986 | 893.0029 | L | 288.2030 | 144.6051 | 271.1765 | 136.0919 | | | 2 |
| 15 | | | | | | | R | ***175.1190*** | 88.0631 | 158.0924 | 79.5498 | | | 1 |

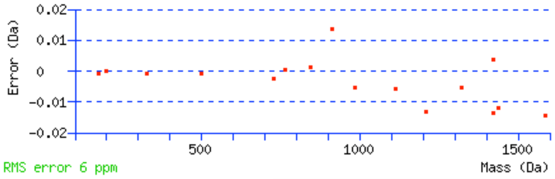


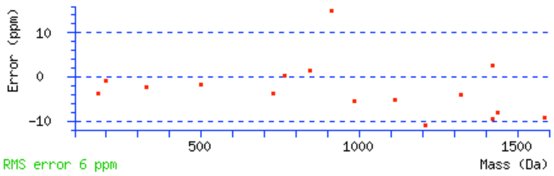


MS/MS Fragmentation of **MKVEIDSFSGAKIYPGR**
Found in **RL24A_YEAST** in **Uniprot_S288C_yeast**, sp|P04449|RL24A_YEAST 60S ribosomal protein L24-A OS=Saccharomyces cerevisiae (strain ATCC 204508 / S288c) OX=559292 GN=RPL24A PE=1 SV=1

Match to Query 7260: 1967.056280 from(984.535416,2+) intensity(719955.1250) rtinseconds(3638.54208) index(9100)
Title: qExactive02201_+4.13313.13313.2
Data file D:\Panyue\enriched_fr.+4.mzML

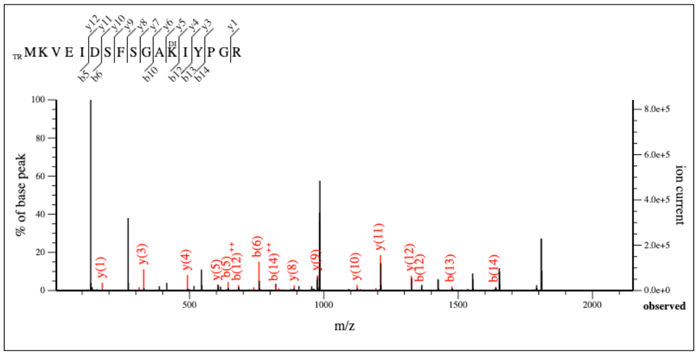

**Monoisotopic mass of neutral peptide Mr(calc):** 1967.0550
**Fixed modifications:** Carbamidomethyl (C) (apply to specified residues or termini only)
**Variable modifications:**
N-term : Trimethyl (Protein N-term)
K12 : Dimethyl (K)
**Ions Score:** 39 **Expect:** 0.009
**Matches :** 22/168 fragment ions using 66 most intense peaks (help)

| # | b | $b^{++}$ | $b^{*}$ | $b^{*++}$ | $b^{0}$ | $b^{0++}$ | Seq. | y | $y^{++}$ | $y^{*}$ | $y^{*++}$ | $y^{0}$ | $y^{0++}$ | # |
|---|---|---|---|---|---|---|---|---|---|---|---|---|---|---|
| 1 | 174.0947 | 87.5510 | | | | | M | | | | | | | 17 |
| 2 | 302.1897 | 151.5985 | 285.1631 | 143.0852 | | | K | 1794.9749 | 897.9911 | 1777.9483 | 889.4778 | 1776.9643 | 888.9858 | 16 |
| 3 | 401.2581 | 201.1327 | 384.2315 | 192.6194 | | | V | 1666.8799 | 833.9436 | 1649.8533 | 825.4303 | 1648.8693 | 824.9383 | 15 |
| 4 | 530.3007 | 265.6540 | 513.2741 | 257.1407 | 512.2901 | 256.6487 | E | 1567.8115 | 784.4094 | 1550.7849 | 775.8961 | 1549.8009 | 775.4041 | 14 |
| 5 | **643.3847** | 322.1960 | 626.3582 | 313.6827 | 625.3742 | 313.1907 | I | 1438.7689 | 719.8881 | 1421.7423 | 711.3748 | 1420.7583 | 710.8828 | 13 |
| 6 | **758.4117** | 379.7095 | 741.3851 | 371.1962 | 740.4011 | 370.7042 | D | ***1325.6848*** | 663.3461 | 1308.6583 | 654.8328 | 1307.6743 | 654.3408 | 12 |
| 7 | 845.4437 | 423.2255 | 828.4172 | 414.7122 | 827.4332 | 414.2202 | S | ***1210.6579*** | 605.8326 | 1193.6313 | 597.3193 | 1192.6473 | 596.8273 | 11 |
| 8 | 992.5121 | 496.7597 | 975.4856 | 488.2464 | 974.5016 | 487.7544 | F | ***1123.6259*** | 562.3166 | 1106.5993 | 553.8033 | 1105.6153 | 553.3113 | 10 |
| 9 | 1079.5442 | 540.2757 | 1062.5176 | 531.7624 | 1061.5336 | 531.2704 | S | ***976.5574*** | 488.7824 | 959.5309 | 480.2691 | 958.5469 | 479.7771 | 9 |
| 10 | **1136.5656** | 568.7864 | 1119.5391 | 560.2732 | 1118.5551 | 559.7812 | G | ***889.5254*** | 445.2663 | 872.4989 | 436.7531 | | | 8 |
| 11 | 1207.6027 | 604.3050 | 1190.5762 | 595.7917 | 1189.5922 | 595.2997 | A | ***832.5039*** | 416.7556 | 815.4774 | 408.2423 | | | 7 |
| 12 | **1363.7290** | **682.3681** | 1346.7024 | 673.8549 | 1345.7184 | 673.3629 | K | ***761.4668*** | 381.2371 | 744.4403 | 372.7238 | | | 6 |
| 13 | **1476.8131** | **738.9102** | 1459.7865 | 730.3969 | 1458.8025 | 729.9049 | I | ***605.3406*** | 303.1739 | 588.3140 | 294.6606 | | | 5 |
| 14 | **1639.8764** | **820.4418** | 1622.8498 | 811.9286 | 1621.8658 | 811.4366 | Y | ***492.2565*** | 246.6319 | 475.2300 | 238.1186 | | | 4 |
| 15 | 1736.9292 | 868.9682 | 1719.9026 | 860.4549 | 1718.9186 | 859.9629 | P | ***329.1932*** | 165.1002 | 312.1666 | 156.5870 | | | 3 |
| 16 | 1793.9506 | 897.4789 | 1776.9241 | 888.9657 | 1775.9401 | 888.4737 | G | 232.1404 | 116.5738 | 215.1139 | 108.0606 | | | 2 |
| 17 | | | | | | | R | ***175.1190*** | 88.0631 | 158.0924 | 79.5498 | | | 1 |

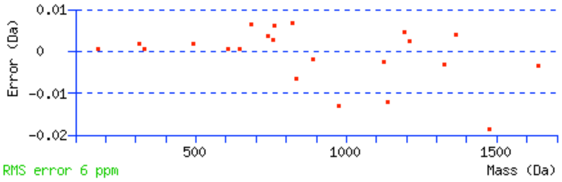


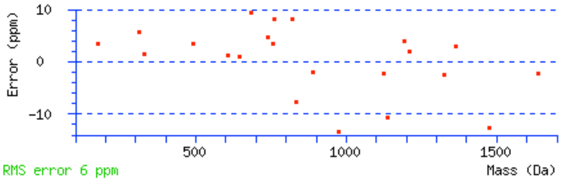


MS/MS Fragmentation of **MKVEIDSFSGAKIYPGR**
Found in **RL24A_YEAST** in **Uniprot_S288C_yeast**, sp|P04449|RL24A_YEAST 60S ribosomal protein L24-A OS=Saccharomyces cerevisiae (strain ATCC 204508 / S288c) OX=559292 GN=RPL24A PE=1 SV=1

Match to Query 7261: 1967.056328 from(492.771358,4+) intensity(606008.1875) rtinseconds(3640.02366) index(9102)
Title: qExactive02201_+4.13319.13319.4
Data file D:\Panyue\enriched_fr.+4.mzML

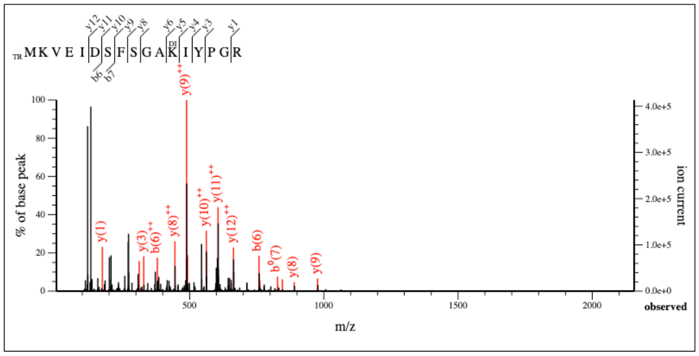

**Monoisotopic mass of neutral peptide Mr(calc):** 1967.0550
**Fixed modifications:** Carbamidomethyl (C) (apply to specified residues or termini only)
**Variable modifications:**
N-term : Trimethyl (Protein N-term)
K12 : Dimethyl (K)
**Ions Score:** 35 **Expect:** 0.021
**Matches :** 18/168 fragment ions using 36 most intense peaks (help)

| # | b | $b^{++}$ | b* | $b^{*++}$ | $b^0$ | $b^{0++}$ | Seq. | y | $y^{++}$ | y* | $y^{*++}$ | $y^0$ | $y^{0++}$ | # |
|---|---|---|---|---|---|---|---|---|---|---|---|---|---|---|
| 1 | 174.0947 | 87.5510 | | | | | M | | | | | | | 17 |
| 2 | 302.1897 | 151.5985 | 285.1631 | 143.0852 | | | K | 1794.9749 | 897.9911 | 1777.9483 | 889.4778 | 1776.9643 | 888.9858 | 16 |
| 3 | 401.2581 | 201.1327 | 384.2315 | 192.6194 | | | V | 1666.8799 | 833.9436 | 1649.8533 | 825.4303 | 1648.8693 | 824.9383 | 15 |
| 4 | 530.3007 | 265.6540 | 513.2741 | 257.1407 | 512.2901 | 256.6487 | E | 1567.8115 | 784.4094 | 1550.7849 | 775.8961 | 1549.8009 | 775.4041 | 14 |
| 5 | 643.3847 | 322.1960 | 626.3582 | 313.6827 | 625.3742 | 313.1907 | I | 1438.7689 | 719.8881 | 1421.7423 | 711.3748 | 1420.7583 | 710.8828 | 13 |
| 6 | **758.4117** | **379.7095** | 741.3851 | 371.1962 | 740.4011 | 370.7042 | D | 1325.6848 | ***663.3461*** | 1308.6583 | 654.8328 | 1307.6743 | 654.3408 | 12 |
| 7 | **845.4437** | 423.2255 | 828.4172 | 414.7122 | 827.4332 | 414.2202 | S | 1210.6579 | ***605.8326*** | 1193.6313 | 597.3193 | 1192.6473 | 596.8273 | 11 |
| 8 | 992.5121 | 496.7597 | 975.4856 | 488.2464 | 974.5016 | 487.7544 | F | 1123.6259 | ***562.3166*** | 1106.5993 | 553.8033 | 1105.6153 | 553.3113 | 10 |
| 9 | 1079.5442 | 540.2757 | 1062.5176 | 531.7624 | 1061.5336 | 531.2704 | S | ***976.5574*** | ***488.7824*** | 959.5309 | 480.2691 | 958.5469 | 479.7771 | 9 |
| 10 | 1136.5656 | 568.7864 | 1119.5391 | 560.2732 | 1118.5551 | 559.7812 | G | ***889.5254*** | ***445.2663*** | 872.4989 | 436.7531 | | | 8 |
| 11 | 1207.6027 | 604.3050 | 1190.5762 | 595.7917 | 1189.5922 | 595.2997 | A | 832.5039 | 416.7556 | 815.4774 | 408.2423 | | | 7 |
| 12 | 1363.7290 | 682.3681 | 1346.7024 | 673.8549 | 1345.7184 | 673.3629 | K | ***761.4668*** | 381.2371 | 744.4403 | 372.7238 | | | 6 |
| 13 | 1476.8131 | 738.9102 | 1459.7865 | 730.3969 | 1458.8025 | 729.9049 | I | ***605.3406*** | 303.1739 | 588.3140 | 294.6606 | | | 5 |
| 14 | 1639.8764 | 820.4418 | 1622.8498 | 811.9286 | 1621.8658 | 811.4366 | Y | ***492.2565*** | 246.6319 | 475.2300 | 238.1186 | | | 4 |
| 15 | 1736.9292 | 868.9682 | 1719.9026 | 860.4549 | 1718.9186 | 859.9629 | P | ***329.1932*** | 165.1002 | 312.1666 | 156.5870 | | | 3 |
| 16 | 1793.9506 | 897.4789 | 1776.9241 | 888.9657 | 1775.9401 | 888.4737 | G | 232.1404 | 116.5738 | 215.1139 | 108.0606 | | | 2 |
| 17 | | | | | | | R | ***175.1190*** | 88.0631 | 158.0924 | 79.5498 | | | 1 |

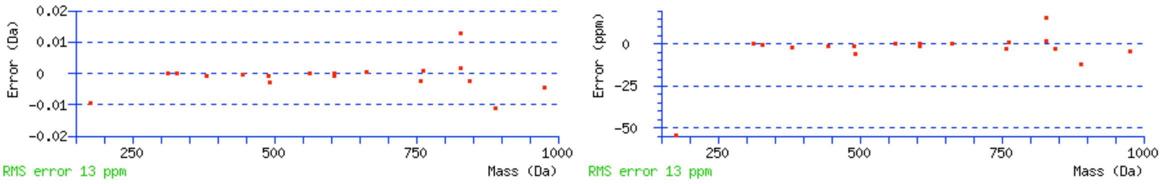


MS/MS Fragmentation of **MKVEVDSFSGAKIYPGR**
Found in **RL24B_YEAST** in **Uniprot_S288C_yeast**, sp|P24000|RL24B_YEAST 60S ribosomal protein L24-B OS=Saccharomyces cerevisiae (strain ATCC 204508 / S288c) OX=559292 GN=RPL24B PE=1 SV=1

Match to Query 7344: 1981.071681 from(661.364503,3+) intensity(1790589.7500) rtinseconds(3978.66768) index(9384)
Title: qExactive02201_+4.14462.14462.3
Data file D:\Panyue\enriched_fr.+4.mzML

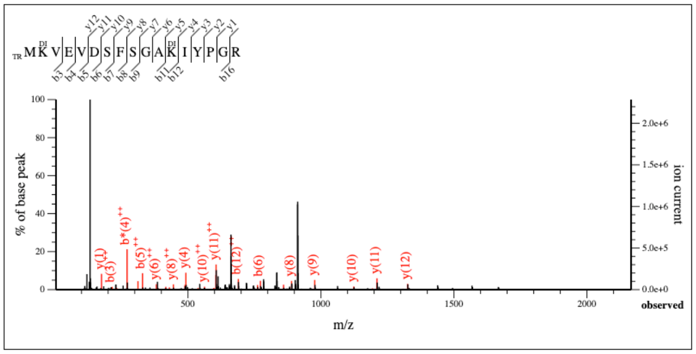

**Monoisotopic mass of neutral peptide Mr(calc):** 1981.0706
**Fixed modifications:** Carbamidomethyl (C) (apply to specified residues or termini only)
**Variable modifications:**
N-term : Trimethyl (Protein N-term)
K2 : Dimethyl (K)
K12 : Dimethyl (K)
**Ions Score:** 24 **Expect:** 0.31
**Matches :** 35/168 fragment ions using 126 most intense peaks (help)

| # | b | b++ | b* | b*++ | b0 | b0++ | Seq. | y | y++ | y* | y*++ | y0 | y0++ | # |
|---|---|---|---|---|---|---|---|---|---|---|---|---|---|---|
| 1 | 174.0947 | 87.5510 | | | | | M | | | | | | | 17 |
| 2 | 330.2210 | 165.6141 | 313.1944 | 157.1009 | | | K | 1808.9905 | 904.9989 | 1791.9640 | 896.4856 | 1790.9799 | 895.9936 | 16 |
| 3 | 429.2894 | **215.1483** | 412.2628 | 206.6351 | | | V | 1652.8642 | 826.9358 | 1635.8377 | 818.4225 | 1634.8537 | 817.9305 | 15 |
| 4 | 558.3320 | 279.6696 | 541.3054 | 271.1564 | 540.3214 | 270.6643 | E | 1553.7958 | 777.4016 | 1536.7693 | 768.8883 | 1535.7853 | 768.3963 | 14 |
| 5 | 657.4004 | **329.2038** | 640.3738 | 320.6906 | 639.3898 | 320.1986 | V | 1424.7532 | 712.8803 | 1407.7267 | 704.3670 | 1406.7427 | 703.8750 | 13 |
| 6 | **772.4273** | **386.7173** | 755.4008 | 378.2040 | 754.4168 | 377.7120 | D | ***1325.6848*** | 663.3461 | 1308.6583 | 654.8328 | 1307.6743 | 654.3408 | 12 |
| 7 | **859.4594** | **430.2333** | 842.4328 | 421.7200 | 841.4488 | 421.2280 | S | ***1210.6579*** | **605.8326** | 1193.6313 | 597.3193 | 1192.6473 | 596.8273 | 11 |
| 8 | **1006.5278** | 503.7675 | 989.5012 | 495.2543 | 988.5172 | 494.7622 | F | ***1123.6259*** | **562.3166** | 1106.5993 | 553.8033 | 1105.6153 | 553.3113 | 10 |
| 9 | 1093.5598 | 547.2835 | 1076.5333 | 538.7703 | 1075.5492 | 538.2783 | S | ***976.5574*** | **488.7824** | 959.5309 | 480.2691 | 958.5469 | 479.7771 | 9 |
| 10 | 1150.5813 | 575.7943 | 1133.5547 | 567.2810 | 1132.5707 | 566.7890 | G | ***889.5254*** | **445.2663** | 872.4989 | 436.7531 | | | 8 |
| 11 | **1221.6184** | 611.3128 | 1204.5918 | 602.7996 | 1203.6078 | 602.3075 | A | 832.5039 | **416.7556** | 815.4774 | 408.2423 | | | 7 |
| 12 | 1377.7446 | **689.3760** | 1360.7181 | 680.8627 | 1359.7341 | 680.3707 | K | ***761.4668*** | **381.2371** | 744.4403 | 372.7238 | | | 6 |
| 13 | 1490.8287 | 745.9180 | 1473.8022 | 737.4047 | 1472.8181 | 736.9127 | I | ***605.3406*** | 303.1739 | 588.3140 | 294.6606 | | | 5 |
| 14 | 1653.8920 | 827.4497 | 1636.8655 | 818.9364 | 1635.8815 | 818.4444 | Y | ***492.2565*** | 246.6319 | 475.2300 | 238.1186 | | | 4 |
| 15 | 1750.9448 | 875.9760 | 1733.9183 | 867.4628 | 1732.9342 | 866.9708 | P | ***329.1932*** | 165.1002 | 312.1666 | 156.5870 | | | 3 |
| 16 | 1807.9663 | **904.4868** | 1790.9397 | 895.9735 | 1789.9557 | 895.4815 | G | ***232.1404*** | 116.5738 | 215.1139 | 108.0606 | | | 2 |
| 17 | | | | | | | R | ***175.1190*** | 88.0631 | 158.0924 | 79.5498 | | | 1 |

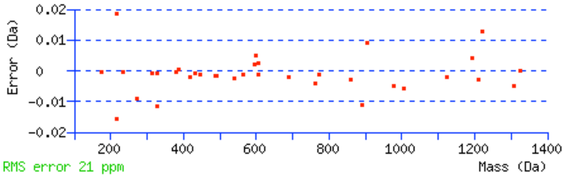


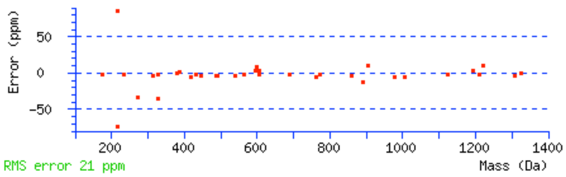


MS/MS Fragmentation of **AKQSLDVSSDRR**
Found in **RL26A_YEAST** in **Uniprot_S288C_yeast**, sp|P05743|RL26A_YEAST 60S ribosomal protein L26-A OS=Saccharomyces cerevisiae (strain ATCC 204508 / S288c) OX=559292 GN=RPL26A PE=1 SV=3

Match to Query 2203: 1402.745448 from(702.380000,2+) intensity(1231792.6250) rtinseconds(370.442946) index(357)
Title: qExactive02201_+4.1159.1159.2
Data file D:\Panyue\enriched_fr.+4.mzML

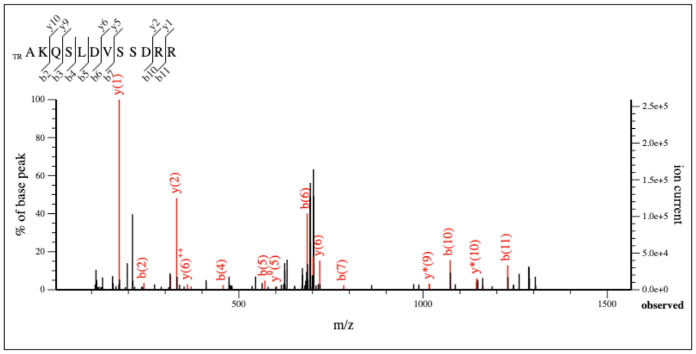

**Monoisotopic mass of neutral peptide Mr(calc):** 1402.7528
**Fixed modifications:** Carbamidomethyl (C) (apply to specified residues or termini only)
**Variable modifications:**
N-term : Trimethyl (Protein N-term)
**Ions Score:** 21 **Expect:** 0.36
**Matches :** 20/120 fragment ions using 78 most intense peaks (help)

| # | b | b++ | b* | b*++ | b0 | b0++ | Seq. | y | y++ | y* | y*++ | y0 | y0++ | # |
|---|---|---|---|---|---|---|---|---|---|---|---|---|---|---|
| 1 | 114.0913 | 57.5493 | | | | | A | | | | | | | 12 |
| 2 | 242.1863 | 121.5968 | 225.1598 | 113.0835 | | | K | 1290.6761 | 645.8417 | 1273.6495 | 637.3284 | 1272.6655 | 636.8364 | 11 |
| 3 | 370.2449 | 185.6261 | 353.2183 | 177.1128 | | | Q | 1162.5811 | 581.7942 | 1145.5545 | 573.2809 | 1144.5705 | 572.7889 | 10 |
| 4 | 457.2769 | 229.1421 | 440.2504 | 220.6288 | 439.2663 | 220.1368 | S | 1034.5225 | 517.7649 | 1017.4960 | 509.2516 | 1016.5119 | 508.7596 | 9 |
| 5 | 570.3610 | 285.6841 | 553.3344 | 277.1708 | 552.3504 | 276.6788 | L | 947.4905 | 474.2489 | 930.4639 | 465.7356 | 929.4799 | 465.2436 | 8 |
| 6 | 685.3879 | 343.1976 | 668.3614 | 334.6843 | 667.3774 | 334.1923 | D | 834.4064 | 417.7068 | 817.3799 | 409.1936 | 816.3959 | 408.7016 | 7 |
| 7 | 784.4563 | 392.7318 | 767.4298 | 384.2185 | 766.4458 | 383.7265 | V | 719.3795 | 360.1934 | 702.3529 | 351.6801 | 701.3689 | 351.1881 | 6 |
| 8 | 871.4884 | 436.2478 | 854.4618 | 427.7345 | 853.4778 | 427.2425 | S | 620.3111 | 310.6592 | 603.2845 | 302.1459 | 602.3005 | 301.6539 | 5 |
| 9 | 958.5204 | 479.7638 | 941.4938 | 471.2506 | 940.5098 | 470.7585 | S | 533.2790 | 267.1432 | 516.2525 | 258.6299 | 515.2685 | 258.1379 | 4 |
| 10 | 1073.5473 | 537.2773 | 1056.5208 | 528.7640 | 1055.5368 | 528.2720 | D | 446.2470 | 223.6271 | 429.2205 | 215.1139 | 428.2364 | 214.6219 | 3 |
| 11 | 1229.6484 | 615.3279 | 1212.6219 | 606.8146 | 1211.6379 | 606.3226 | R | 331.2201 | 166.1137 | 314.1935 | 157.6004 | | | 2 |
| 12 | | | | | | | R | 175.1190 | 88.0631 | 158.0924 | 79.5498 | | | 1 |

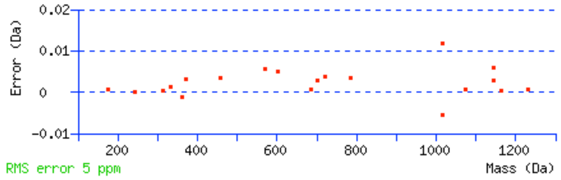


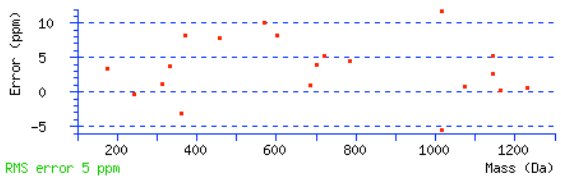


MS/MS Fragmentation of AKQSLDVSSDRR
Found in RL26A_YEAST in Uniprot_S288C_yeast, sp|P05743|RL26A_YEAST 60S ribosomal protein L26-A OS=Saccharomyces cerevisiae (strain ATCC 204508 / S288c) OX=559292 GN=RPL26A PE=1 SV=3

Match to Query 2204: 1402.752978 from(702.383765,2+) intensity(16009216.0000) rtinseconds(344.395104) index(279)
Title: qExactive02201_+4.1059.1059.2
Data file D:\Panyue\enriched_fr.+4.mzML

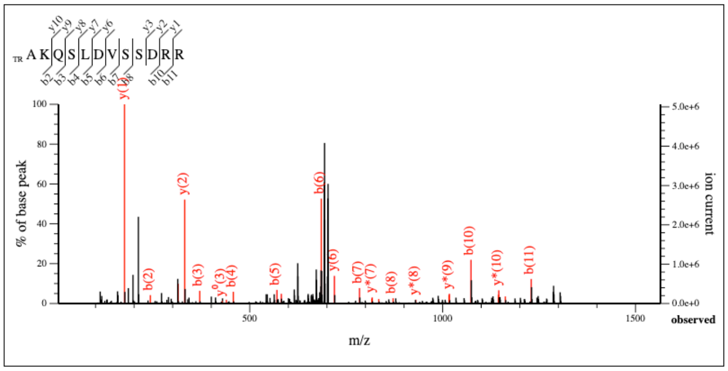

**Monoisotopic mass of neutral peptide Mr(calc):** 1402.7528
**Fixed modifications:** Carbamidomethyl (C) (apply to specified residues or termini only)
**Variable modifications:**
N-term : Trimethyl (Protein N-term)
**Ions Score:** 39 **Expect:** 0.0055
**Matches :** 25/120 fragment ions using 65 most intense peaks (help)

| # | b | b++ | b* | b*++ | b0 | b0++ | Seq. | y | y++ | y* | y*++ | y0 | y0++ | # |
|---|---|---|---|---|---|---|---|---|---|---|---|---|---|---|
| 1 | 114.0913 | 57.5493 | | | | | A | | | | | | | 12 |
| 2 | 242.1863 | 121.5968 | 225.1598 | 113.0835 | | | K | 1290.6761 | 645.8417 | 1273.6495 | 637.3284 | 1272.6655 | 636.8364 | 11 |
| 3 | 370.2449 | 185.6261 | 353.2183 | 177.1128 | | | Q | 1162.5811 | 581.7942 | 1145.5545 | 573.2809 | 1144.5705 | 572.7889 | 10 |
| 4 | 457.2769 | 229.1421 | 440.2504 | 220.6288 | 439.2663 | 220.1368 | S | 1034.5225 | 517.7649 | 1017.4960 | 509.2516 | 1016.5119 | 508.7596 | 9 |
| 5 | 570.3610 | 285.6841 | 553.3344 | 277.1708 | 552.3504 | 276.6788 | L | 947.4905 | 474.2489 | 930.4639 | 465.7356 | 929.4799 | 465.2436 | 8 |
| 6 | 685.3879 | 343.1976 | 668.3614 | 334.6843 | 667.3774 | 334.1923 | D | 834.4064 | 417.7068 | 817.3799 | 409.1936 | 816.3959 | 408.7016 | 7 |
| 7 | 784.4563 | 392.7318 | 767.4298 | 384.2185 | 766.4458 | 383.7265 | V | 719.3795 | 360.1934 | 702.3529 | 351.6801 | 701.3689 | 351.1881 | 6 |
| 8 | 871.4884 | 436.2478 | 854.4618 | 427.7345 | 853.4778 | 427.2425 | S | 620.3111 | 310.6592 | 603.2845 | 302.1459 | 602.3005 | 301.6539 | 5 |
| 9 | 958.5204 | 479.7638 | 941.4938 | 471.2506 | 940.5098 | 470.7585 | S | 533.2790 | 267.1432 | 516.2525 | 258.6299 | 515.2685 | 258.1379 | 4 |
| 10 | 1073.5473 | 537.2773 | 1056.5208 | 528.7640 | 1055.5368 | 528.2720 | D | 446.2470 | 223.6271 | 429.2205 | 215.1139 | 428.2364 | 214.6219 | 3 |
| 11 | 1229.6484 | 615.3279 | 1212.6219 | 606.8146 | 1211.6379 | 606.3226 | R | 331.2201 | 166.1137 | 314.1935 | 157.6004 | | | 2 |
| 12 | | | | | | | R | 175.1190 | 88.0631 | 158.0924 | 79.5498 | | | 1 |

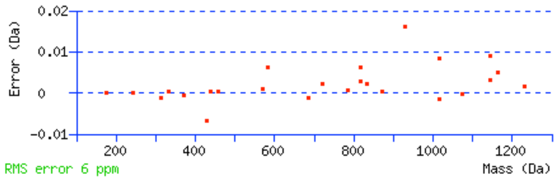


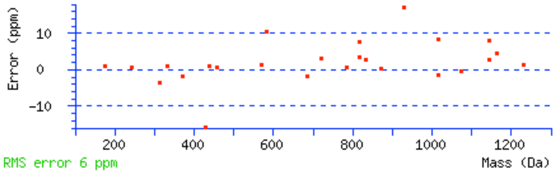


MS/MS Fragmentation of **AKQSLDVSSDRR**
Found in **RL26A_YEAST** in **Uniprot_S288C_yeast**, sp|P05743|RL26A_YEAST 60S ribosomal protein L26-A OS=Saccharomyces cerevisiae (strain ATCC 204508 / S288c) OX=559292 GN=RPL26A PE=1 SV=3

Match to Query 2205: 1402.753124 from(702.383838,2+) intensity(1219017.6250) rtinseconds(331.87272) index(245)
Title: qExactive02201_+4.1011.1011.2
Data file D:\Panyue\enriched_fr.+4.mzML

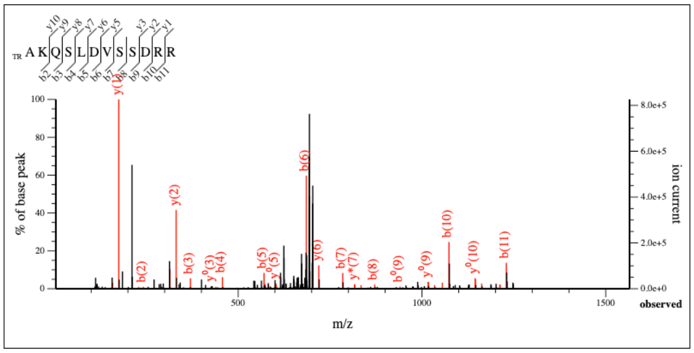

**Monoisotopic mass of neutral peptide Mr(calc):** 1402.7528
**Fixed modifications:** Carbamidomethyl (C) (apply to specified residues or termini only)
**Variable modifications:**
N-term : Trimethyl (Protein N-term)
**Ions Score:** 39 **Expect:** 0.0051
**Matches :** 38/120 fragment ions using 114 most intense peaks (help)

| # | b | b++ | b* | b*++ | b0 | b0++ | Seq. | y | y++ | y* | y*++ | y0 | y0++ | # |
|---|---|---|---|---|---|---|---|---|---|---|---|---|---|---|
| 1 | 114.0913 | 57.5493 | | | | | A | | | | | | | 12 |
| 2 | 242.1863 | 121.5968 | 225.1598 | 113.0835 | | | K | 1290.6761 | 645.8417 | 1273.6495 | 637.3284 | 1272.6655 | 636.8364 | 11 |
| 3 | 370.2449 | 185.6261 | 353.2183 | 177.1128 | | | Q | 1162.5811 | 581.7942 | 1145.5545 | 573.2809 | 1144.5705 | 572.7889 | 10 |
| 4 | 457.2769 | 229.1421 | 440.2504 | 220.6288 | 439.2663 | 220.1368 | S | 1034.5225 | 517.7649 | 1017.4960 | 509.2516 | 1016.5119 | 508.7596 | 9 |
| 5 | 570.3610 | 285.6841 | 553.3344 | 277.1708 | 552.3504 | 276.6788 | L | 947.4905 | 474.2489 | 930.4639 | 465.7356 | 929.4799 | 465.2436 | 8 |
| 6 | 685.3879 | 343.1976 | 668.3614 | 334.6843 | 667.3774 | 334.1923 | D | 834.4064 | 417.7068 | 817.3799 | 409.1936 | 816.3959 | 408.7016 | 7 |
| 7 | 784.4563 | 392.7318 | 767.4298 | 384.2185 | 766.4458 | 383.7265 | V | 719.3795 | 360.1934 | 702.3529 | 351.6801 | 701.3689 | 351.1881 | 6 |
| 8 | 871.4884 | 436.2478 | 854.4618 | 427.7345 | 853.4778 | 427.2425 | S | 620.3111 | 310.6592 | 603.2845 | 302.1459 | 602.3005 | 301.6539 | 5 |
| 9 | 958.5204 | 479.7638 | 941.4938 | 471.2506 | 940.5098 | 470.7585 | S | 533.2790 | 267.1432 | 516.2525 | 258.6299 | 515.2685 | 258.1379 | 4 |
| 10 | 1073.5473 | 537.2773 | 1056.5208 | 528.7640 | 1055.5368 | 528.2720 | D | 446.2470 | 223.6271 | 429.2205 | 215.1139 | 428.2364 | 214.6219 | 3 |
| 11 | 1229.6484 | 615.3279 | 1212.6219 | 606.8146 | 1211.6379 | 606.3226 | R | 331.2201 | 166.1137 | 314.1935 | 157.6004 | | | 2 |
| 12 | | | | | | | R | 175.1190 | 88.0631 | 158.0924 | 79.5498 | | | 1 |

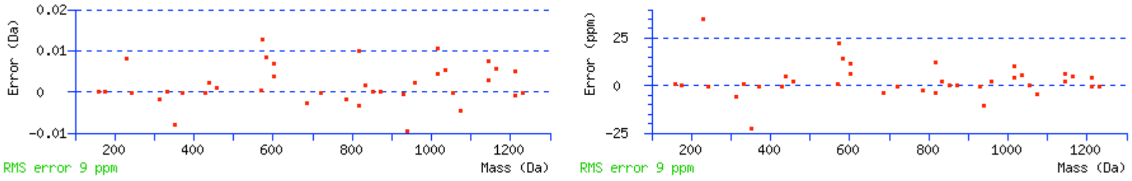


MS/MS Fragmentation of **AKQSLDVSSDRR**
Found in **RL26A_YEAST** in **Uniprot_S288C_yeast**, sp|P05743|RL26A_YEAST 60S ribosomal protein L26-A OS=Saccharomyces cerevisiae (strain ATCC 204508 / S288c) OX=559292 GN=RPL26A PE=1 SV=3

Match to Query 2206: 1402.753468 from(702.384010,2+) intensity(2944040.0000) rtinseconds(357.001128) index(313)
Title: qExactive02201_+4.1109.1109.2
Data file D:\Panyue\enriched_fr.+4.mzML

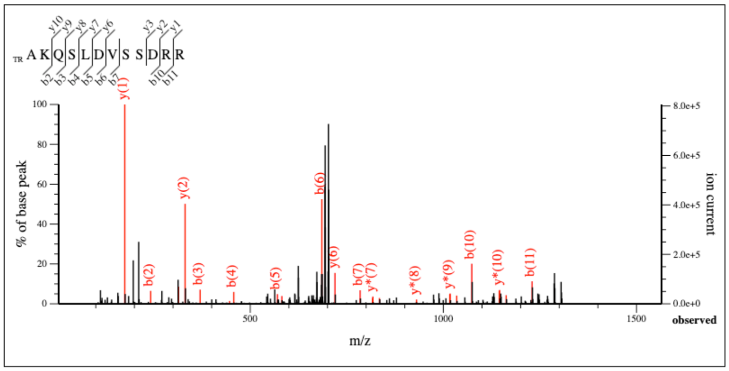

**Monoisotopic mass of neutral peptide Mr(calc):** 1402.7528
**Fixed modifications:** Carbamidomethyl (C) (apply to specified residues or termini only)
**Variable modifications:**
N-term : Trimethyl (Protein N-term)
**Ions Score:** 30 **Expect:** 0.045
**Matches :** 24/120 fragment ions using 65 most intense peaks (help)

| # | b | b++ | b* | b*++ | b0 | b0++ | Seq. | y | y++ | y* | y*++ | y0 | y0++ | # |
|---|---|---|---|---|---|---|---|---|---|---|---|---|---|---|
| 1 | 114.0913 | 57.5493 | | | | | A | | | | | | | 12 |
| 2 | ***242.1863*** | 121.5968 | 225.1598 | 113.0835 | | | K | 1290.6761 | 645.8417 | 1273.6495 | 637.3284 | 1272.6655 | 636.8364 | 11 |
| 3 | ***370.2449*** | 185.6261 | 353.2183 | 177.1128 | | | Q | **1162.5811** | **581.7942** | 1145.5545 | 573.2809 | 1144.5705 | 572.7889 | 10 |
| 4 | ***457.2769*** | 229.1421 | 440.2504 | 220.6288 | 439.2663 | 220.1368 | S | **1034.5225** | 517.7649 | 1017.4960 | 509.2516 | 1016.5119 | 508.7596 | 9 |
| 5 | ***570.3610*** | 285.6841 | 553.3344 | 277.1708 | 552.3504 | 276.6788 | L | 947.4905 | 474.2489 | 930.4639 | 465.7356 | 929.4799 | 465.2436 | 8 |
| 6 | ***685.3879*** | 343.1976 | 668.3614 | 334.6843 | 667.3774 | 334.1923 | D | **834.4064** | 417.7068 | 817.3799 | 409.1936 | 816.3959 | 408.7016 | 7 |
| 7 | ***784.4563*** | 392.7318 | 767.4298 | 384.2185 | 766.4458 | 383.7265 | V | **719.3795** | 360.1934 | 702.3529 | 351.6801 | 701.3689 | 351.1881 | 6 |
| 8 | 871.4884 | 436.2478 | 854.4618 | 427.7345 | 853.4778 | 427.2425 | S | 620.3111 | 310.6592 | 603.2845 | 302.1459 | 602.3005 | 301.6539 | 5 |
| 9 | 958.5204 | 479.7638 | 941.4938 | 471.2506 | 940.5098 | 470.7585 | S | 533.2790 | 267.1432 | 516.2525 | 258.6299 | 515.2685 | 258.1379 | 4 |
| 10 | ***1073.5473*** | 537.2773 | 1056.5208 | 528.7640 | 1055.5368 | 528.2720 | D | **446.2470** | 223.6271 | 429.2205 | 215.1139 | 428.2364 | 214.6219 | 3 |
| 11 | ***1229.6484*** | 615.3279 | 1212.6219 | 606.8146 | 1211.6379 | 606.3226 | R | **331.2201** | 166.1137 | 314.1935 | 157.6004 | | | 2 |
| 12 | | | | | | | R | **175.1190** | 88.0631 | 158.0924 | 79.5498 | | | 1 |

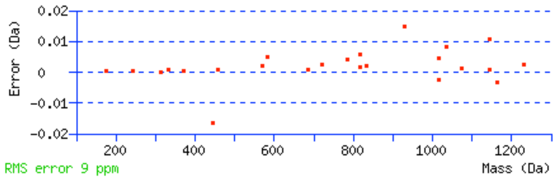


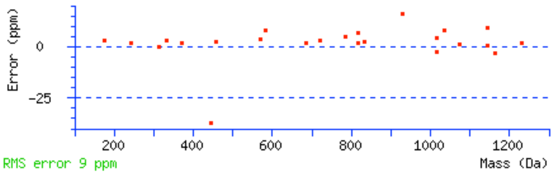


MS/MS Fragmentation of **AKQSLDVSSDRR**
Found in **RL26A_YEAST** in **Uniprot_S288C_yeast**, sp|P05743|RL26A_YEAST 60S ribosomal protein L26-A OS=Saccharomyces cerevisiae (strain ATCC 204508 / S288c) OX=559292 GN=RPL26A PE=1 SV=3

Match to Query 2207: 1402.753869 from(468.591899,3+) intensity(8603326.0000) rtinseconds(346.091064) index(284)
Title: qExactive02201_+4.1066.1066.3
Data file D:\Panyue\enriched_fr.+4.mzML

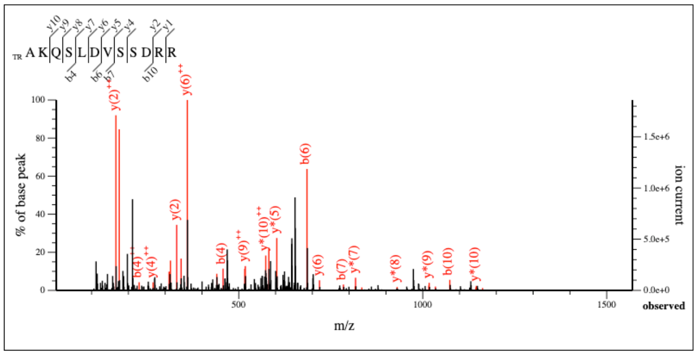

**Monoisotopic mass of neutral peptide Mr(calc):** 1402.7528
**Fixed modifications:** Carbamidomethyl (C) (apply to specified residues or termini only)
**Variable modifications:**
N-term : Trimethyl (Protein N-term)
**Ions Score:** 21 **Expect:** 0.33
**Matches :** 30/120 fragment ions using 77 most intense peaks (help)

| # | b | b++ | b* | b*++ | b0 | b0++ | Seq. | y | y++ | y* | y*++ | y0 | y0++ | # |
|---|---|---|---|---|---|---|---|---|---|---|---|---|---|---|
| 1 | 114.0913 | 57.5493 | | | | | A | | | | | | | 12 |
| 2 | 242.1863 | 121.5968 | 225.1598 | 113.0835 | | | K | 1290.6761 | 645.8417 | 1273.6495 | 637.3284 | 1272.6655 | 636.8364 | 11 |
| 3 | 370.2449 | 185.6261 | 353.2183 | 177.1128 | | | Q | 1162.5811 | 581.7942 | 1145.5545 | 573.2809 | 1144.5705 | 572.7889 | 10 |
| 4 | 457.2769 | 229.1421 | 440.2504 | 220.6288 | 439.2663 | 220.1368 | S | 1034.5225 | 517.7649 | 1017.4960 | 509.2516 | 1016.5119 | 508.7596 | 9 |
| 5 | 570.3610 | 285.6841 | 553.3344 | 277.1708 | 552.3504 | 276.6788 | L | 947.4905 | 474.2489 | 930.4639 | 465.7356 | 929.4799 | 465.2436 | 8 |
| 6 | 685.3879 | 343.1976 | 668.3614 | 334.6843 | 667.3774 | 334.1923 | D | 834.4064 | 417.7068 | 817.3799 | 409.1936 | 816.3959 | 408.7016 | 7 |
| 7 | 784.4563 | 392.7318 | 767.4298 | 384.2185 | 766.4458 | 383.7265 | V | 719.3795 | 360.1934 | 702.3529 | 351.6801 | 701.3689 | 351.1881 | 6 |
| 8 | 871.4884 | 436.2478 | 854.4618 | 427.7345 | 853.4778 | 427.2425 | S | 620.3111 | 310.6592 | 603.2845 | 302.1459 | 602.3005 | 301.6539 | 5 |
| 9 | 958.5204 | 479.7638 | 941.4938 | 471.2506 | 940.5098 | 470.7585 | S | 533.2790 | 267.1432 | 516.2525 | 258.6299 | 515.2685 | 258.1379 | 4 |
| 10 | 1073.5473 | 537.2773 | 1056.5208 | 528.7640 | 1055.5368 | 528.2720 | D | 446.2470 | 223.6271 | 429.2205 | 215.1139 | 428.2364 | 214.6219 | 3 |
| 11 | 1229.6484 | 615.3279 | 1212.6219 | 606.8146 | 1211.6379 | 606.3226 | R | 331.2201 | 166.1137 | 314.1935 | 157.6004 | | | 2 |
| 12 | | | | | | | R | 175.1190 | 88.0631 | 158.0924 | 79.5498 | | | 1 |

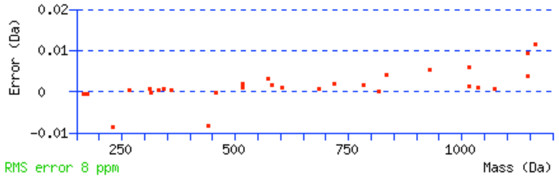


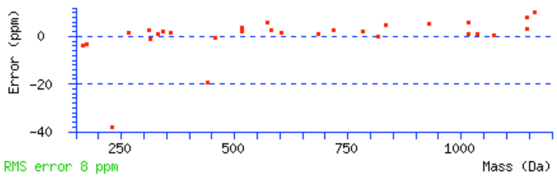


MS/MS Fragmentation of **AKQSLDVSSDRR**
Found in **RL26A_YEAST** in **Uniprot_S288C_yeast**, sp|P05743|RL26A_YEAST 60S ribosomal protein L26-A OS=Saccharomyces cerevisiae (strain ATCC 204508 / S288c) OX=559292 GN=RPL26A PE=1 SV=3

Match to Query 2208: 1402.754973 from(468.592267,3+) intensity(1676556.2500) rtinseconds(332.859792) index(248)
Title: qExactive02201_+4.1015.1015.3
Data file D:\Panyue\enriched_fr.+4.mzML

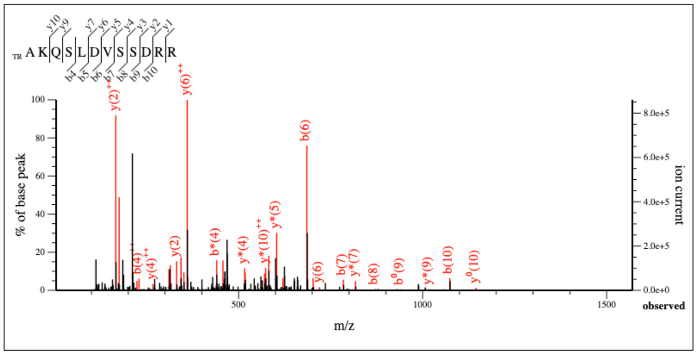

```
Monoisotopic mass of neutral peptide Mr(calc): 1402.7528
Fixed modifications: Carbamidomethyl (C) (apply to specified residues or termini only)
Variable modifications:
  N-term : Trimethyl (Protein N-term)
Ions Score: 28  Expect: 0.074
Matches : 32/120 fragment ions using 77 most intense peaks   (help)
```

| # | b | $b^{++}$ | b* | $b^{*++}$ | $b^0$ | $b^{0++}$ | Seq. | y | $y^{++}$ | y* | $y^{*++}$ | $y^0$ | $y^{0++}$ | # |
|---|---|---|---|---|---|---|---|---|---|---|---|---|---|---|
| 1 | 114.0913 | 57.5493 | | | | | A | | | | | | | 12 |
| 2 | 242.1863 | 121.5968 | 225.1598 | 113.0835 | | | K | 1290.6761 | 645.8417 | 1273.6495 | 637.3284 | 1272.6655 | 636.8364 | 11 |
| 3 | 370.2449 | 185.6261 | 353.2183 | 177.1128 | | | Q | 1162.5811 | ***581.7942*** | 1145.5545 | 573.2809 | 1144.5705 | 572.7889 | 10 |
| 4 | ***457.2769*** | **229.1421** | 440.2504 | 220.6288 | 439.2663 | 220.1368 | S | 1034.5225 | ***517.7649*** | 1017.4960 | 509.2516 | 1016.5119 | 508.7596 | 9 |
| 5 | ***570.3610*** | 285.6841 | 553.3344 | 277.1708 | 552.3504 | 276.6788 | L | 947.4905 | 474.2489 | 930.4639 | 465.7356 | 929.4799 | 465.2436 | 8 |
| 6 | ***685.3879*** | **343.1976** | 668.3614 | 334.6843 | 667.3774 | 334.1923 | D | 834.4064 | 417.7068 | 817.3799 | 409.1936 | 816.3959 | 408.7016 | 7 |
| 7 | ***784.4563*** | 392.7318 | 767.4298 | 384.2185 | 766.4458 | 383.7265 | V | **719.3795** | ***360.1934*** | 702.3529 | 351.6801 | 701.3689 | 351.1881 | 6 |
| 8 | ***871.4884*** | 436.2478 | 854.4618 | 427.7345 | 853.4778 | 427.2425 | S | **620.3111** | ***310.6592*** | 603.2845 | 302.1459 | 602.3005 | 301.6539 | 5 |
| 9 | 958.5204 | 479.7638 | 941.4938 | 471.2506 | 940.5098 | 470.7585 | S | 533.2790 | ***267.1432*** | 516.2525 | 258.6299 | 515.2685 | 258.1379 | 4 |
| 10 | ***1073.5473*** | 537.2773 | 1056.5208 | 528.7640 | 1055.5368 | 528.2720 | D | 446.2470 | ***223.6271*** | 429.2205 | 215.1139 | 428.2364 | 214.6219 | 3 |
| 11 | 1229.6484 | 615.3279 | 1212.6219 | 606.8146 | 1211.6379 | 606.3226 | R | **331.2201** | ***166.1137*** | 314.1935 | 157.6004 | | | 2 |
| 12 | | | | | | | R | **175.1190** | 88.0631 | 158.0924 | 79.5498 | | | 1 |

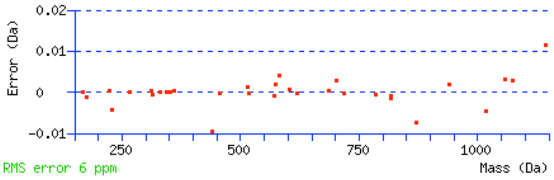


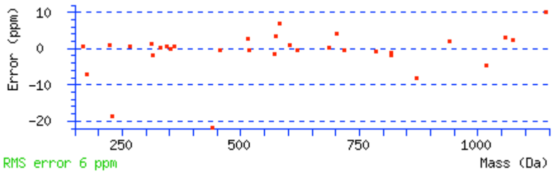


5. F031238. TEF1 (1 peptide spectrum) and Rpl27A/B (8 peptide spectra)

MS/MS Fragmentation of **GKEKSHINVVVIGHVDSGKSTTT**
Found in **EF1A_YEAST** in **Uniprot_S288C_yeast**, sp|P02994|EF1A_YEAST Elongation factor 1-alpha OS=Saccharomyces cerevisiae (strain ATCC 204508 / S288c) OX=559292 GN=TEF1 PE=1 SV=1

Match to Query 6620: 2518.410801 from(840.477543,3+) intensity(8175608.5000) rtinseconds(1754.40684) index(3969)
Title: qExactive02202_+5.6810.6810.3
Data file D:\Panyue\enriched_fr.+5.mzML

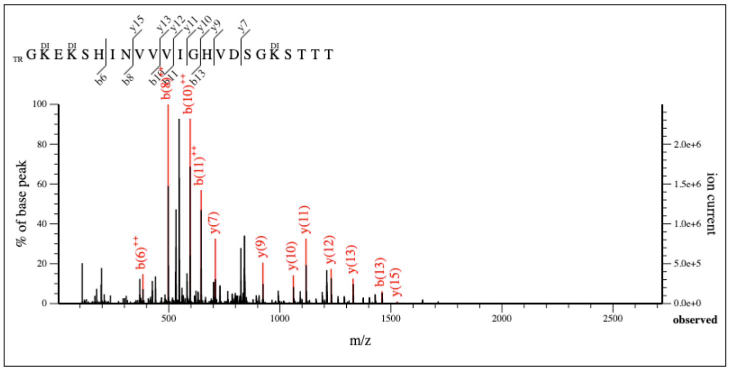


```
Monoisotopic mass of neutral peptide Mr(calc): 2518.4119
Fixed modifications: Carbamidomethyl (C) (apply to specified residues or termini only)
Variable modifications:
  N-term : Trimethyl (Protein N-term)
  K2     : Dimethyl (K)
  K4     : Dimethyl (K)
  K19    : Dimethyl (K)
Ions Score: 41  Expect: 0.0072
Matches : 12/250 fragment ions using 17 most intense peaks   (help)
```

| # | b | b++ | b* | b*++ | b0 | b0++ | Seq. | y | y++ | y* | y*++ | y0 | y0++ | # |
|---|---|---|---|---|---|---|---|---|---|---|---|---|---|---|
| **1** | 100.0757 | 50.5415 | | | | | **G** | | | | | | | **23** |
| **2** | 256.2020 | 128.6046 | 239.1754 | 120.0913 | | | **K** | 2420.3507 | 1210.6790 | 2403.3242 | 1202.1657 | 2402.3402 | 1201.6737 | **22** |
| **3** | 385.2445 | 193.1259 | 368.2180 | 184.6126 | 367.2340 | 184.1206 | **E** | 2264.2245 | 1132.6159 | 2247.1979 | 1124.1026 | 2246.2139 | 1123.6106 | **21** |
| **4** | 541.3708 | 271.1890 | 524.3443 | 262.6758 | 523.3602 | 262.1838 | **K** | 2135.1819 | 1068.0946 | 2118.1553 | 1059.5813 | 2117.1713 | 1059.0893 | **20** |
| **5** | 628.4028 | 314.7051 | 611.3763 | 306.1918 | 610.3923 | 305.6998 | **S** | 1979.0556 | 990.0315 | 1962.0291 | 981.5182 | 1961.0451 | 981.0262 | **19** |
| **6** | 765.4617 | **383.2345** | 748.4352 | 374.7212 | 747.4512 | 374.2292 | **H** | 1892.0236 | 946.5154 | 1874.9971 | 938.0022 | 1874.0130 | 937.5102 | **18** |
| **7** | 878.5458 | 439.7765 | 861.5193 | 431.2633 | 860.5352 | 430.7713 | **I** | 1754.9647 | 877.9860 | 1737.9381 | 869.4727 | 1736.9541 | 868.9807 | **17** |
| **8** | 992.5887 | **496.7980** | 975.5622 | 488.2847 | 974.5782 | 487.7927 | **N** | 1641.8806 | 821.4440 | 1624.8541 | 812.9307 | 1623.8701 | 812.4387 | **16** |
| **9** | 1091.6572 | 546.3322 | 1074.6306 | 537.8189 | 1073.6466 | 537.3269 | **V** | ***1527.8377*** | 764.4225 | 1510.8112 | 755.9092 | 1509.8271 | 755.4172 | **15** |
| **10** | 1190.7256 | **595.8664** | 1173.6990 | 587.3531 | 1172.7150 | 586.8611 | **V** | 1428.7693 | 714.8883 | 1411.7427 | 706.3750 | 1410.7587 | 705.8830 | **14** |
| **11** | 1289.7940 | **645.4006** | 1272.7674 | 636.8874 | 1271.7834 | 636.3953 | **V** | ***1329.7009*** | 665.3541 | 1312.6743 | 656.8408 | 1311.6903 | 656.3488 | **13** |
| **12** | 1402.8780 | 701.9427 | 1385.8515 | 693.4294 | 1384.8675 | 692.9374 | **I** | ***1230.6325*** | 615.8199 | 1213.6059 | 607.3066 | 1212.6219 | 606.8146 | **12** |
| **13** | **1459.8995** | 730.4534 | 1442.8730 | 721.9401 | 1441.8889 | 721.4481 | **G** | ***1117.5484*** | 559.2778 | 1100.5218 | 550.7646 | 1099.5378 | 550.2726 | **11** |
| **14** | 1596.9584 | 798.9828 | 1579.9319 | 790.4696 | 1578.9479 | 789.9776 | **H** | ***1060.5269*** | 530.7671 | 1043.5004 | 522.2538 | 1042.5164 | 521.7618 | **10** |
| **15** | 1696.0268 | 848.5171 | 1679.0003 | 840.0038 | 1678.0163 | 839.5118 | **V** | ***923.4680*** | 462.2376 | 906.4415 | 453.7244 | 905.4575 | 453.2324 | **9** |
| **16** | 1811.0538 | 906.0305 | 1794.0272 | 897.5173 | 1793.0432 | 897.0252 | **D** | 824.3996 | 412.7034 | 807.3731 | 404.1902 | 806.3890 | 403.6982 | **8** |
| **17** | 1898.0858 | 949.5465 | 1881.0593 | 941.0333 | 1880.0752 | 940.5413 | **S** | ***709.3727*** | 355.1900 | 692.3461 | 346.6767 | 691.3621 | 346.1847 | **7** |
| **18** | 1955.1073 | 978.0573 | 1938.0807 | 969.5440 | 1937.0967 | 969.0520 | **G** | 622.3406 | 311.6740 | 605.3141 | 303.1607 | 604.3301 | 302.6687 | **6** |
| **19** | 2111.2335 | 1056.1204 | 2094.2070 | 1047.6071 | 2093.2230 | 1047.1151 | **K** | 565.3192 | 283.1632 | 548.2926 | 274.6499 | 547.3086 | 274.1579 | **5** |
| **20** | 2198.2656 | 1099.6364 | 2181.2390 | 1091.1231 | 2180.2550 | 1090.6311 | **S** | 409.1929 | 205.1001 | | | 391.1823 | 196.0948 | **4** |
| **21** | 2299.3132 | 1150.1603 | 2282.2867 | 1141.6470 | 2281.3027 | 1141.1550 | **T** | 322.1609 | 161.5841 | | | 304.1503 | 152.5788 | **3** |
| **22** | 2400.3609 | 1200.6841 | 2383.3344 | 1192.1708 | 2382.3504 | 1191.6788 | **T** | 221.1132 | 111.0602 | | | 203.1026 | 102.0550 | **2** |
| **23** | | | | | | | **T** | 120.0655 | 60.5364 | | | 102.0550 | 51.5311 | **1** |

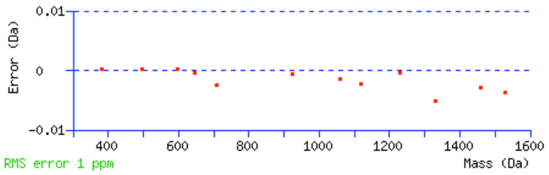


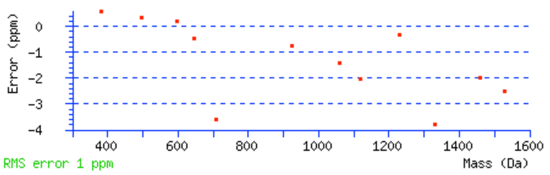


MS/MS Fragmentation of **AKFLKAGKVAVVVR**
Found in **RL27A_YEAST** in **Uniprot_S288C_yeast**, sp|P0C2H6|RL27A_YEAST 60S ribosomal protein L27-A OS=Saccharomyces cerevisiae (strain ATCC 204508 / S288c) OX=559292 GN=RPL27A PE=1 SV=1

Match to Query 1662: 1583.062570 from(792.538561,2+) intensity(107463960.0000) rtinseconds(1895.9883) index(4252)
Title: qExactive02202_+5.7369.7369.2
Data file D:\Panyue\enriched_fr.+5.mzML

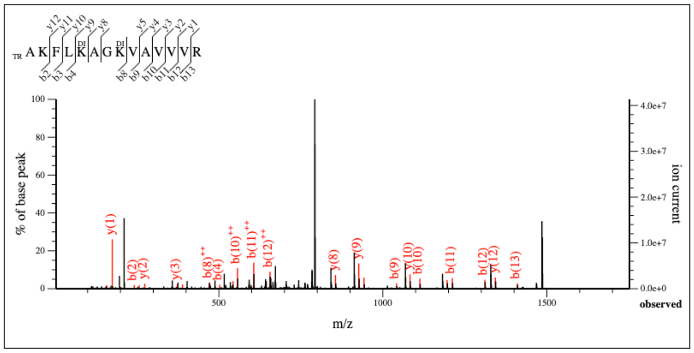

**Monoisotopic mass of neutral peptide Mr(calc):** 1583.0650
**Fixed modifications:** Carbamidomethyl (C) (apply to specified residues or termini only)
**Variable modifications:**
N-term : Trimethyl (Protein N-term)
K5 : Dimethyl (K)
K8 : Dimethyl (K)
**Ions Score:** 35 **Expect:** 0.0028
**Matches :** 25/102 fragment ions using 75 most intense peaks (help)

| # | b | b++ | b* | b*++ | Seq. | y | y++ | y* | y*++ | # |
|---|---|---|---|---|---|---|---|---|---|---|
| 1 | 114.0913 | 57.5493 | | | A | | | | | 14 |
| 2 | **242.1863** | 121.5968 | 225.1598 | 113.0835 | K | 1470.9883 | 735.9978 | 1453.9617 | 727.4845 | 13 |
| 3 | **389.2547** | 195.1310 | 372.2282 | 186.6177 | F | ***1342.8933*** | 671.9503 | 1325.8668 | 663.4370 | 12 |
| 4 | **502.3388** | 251.6730 | 485.3122 | 243.1598 | L | ***1195.8249*** | 598.4161 | 1178.7983 | 589.9028 | 11 |
| 5 | 658.4650 | 329.7362 | 641.4385 | 321.2229 | K | ***1082.7408*** | 541.8741 | 1065.7143 | 533.3608 | 10 |
| 6 | 729.5022 | 365.2547 | 712.4756 | 356.7414 | A | ***926.6146*** | 463.8109 | 909.5880 | 455.2976 | 9 |
| 7 | 786.5236 | 393.7654 | 769.4971 | 385.2522 | G | ***855.5774*** | 428.2924 | 838.5509 | 419.7791 | 8 |
| 8 | **942.6499** | **471.8286** | 925.6233 | 463.3153 | K | 798.5560 | 399.7816 | 781.5294 | 391.2684 | 7 |
| 9 | **1041.7183** | 521.3628 | 1024.6917 | 512.8495 | V | 642.4297 | 321.7185 | 625.4032 | 313.2052 | 6 |
| 10 | **1112.7554** | **556.8813** | 1095.7289 | 548.3681 | A | ***543.3613*** | 272.1843 | 526.3348 | 263.6710 | 5 |
| 11 | **1211.8238** | **606.4156** | 1194.7973 | 597.9023 | V | ***472.3242*** | 236.6657 | 455.2976 | 228.1525 | 4 |
| 12 | **1310.8922** | **655.9498** | 1293.8657 | 647.4365 | V | ***373.2558*** | 187.1315 | 356.2292 | 178.6183 | 3 |
| 13 | **1409.9607** | 705.4840 | 1392.9341 | 696.9707 | V | ***274.1874*** | 137.5973 | 257.1608 | 129.0840 | 2 |
| 14 | | | | | R | ***175.1190*** | 88.0631 | 158.0924 | 79.5498 | 1 |

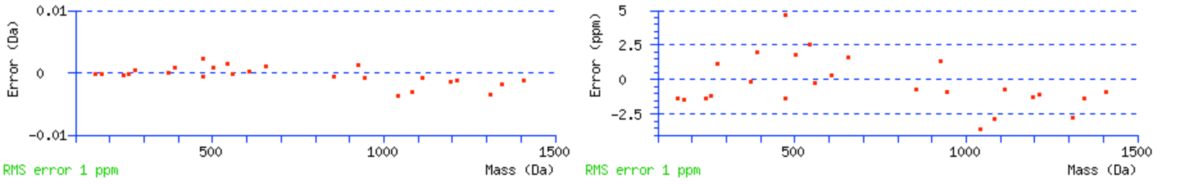


MS/MS Fragmentation of **AKFLKAGKVAVVVR**
Found in **RL27A_YEAST** in **Uniprot_S288C_yeast**, sp|P0C2H6|RL27A_YEAST 60S ribosomal protein L27-A OS=Saccharomyces cerevisiae (strain ATCC 204508 / S288c) OX=559292 GN=RPL27A PE=1 SV=1

Match to Query 1664: 1583.065176 from(528.695668,3+) intensity(60101048.0000) rtinseconds(1889.64402) index(4238)
Title: qExactive02202_+5.7336.7336.3
Data file D:\Panyue\enriched_fr.+5.mzML

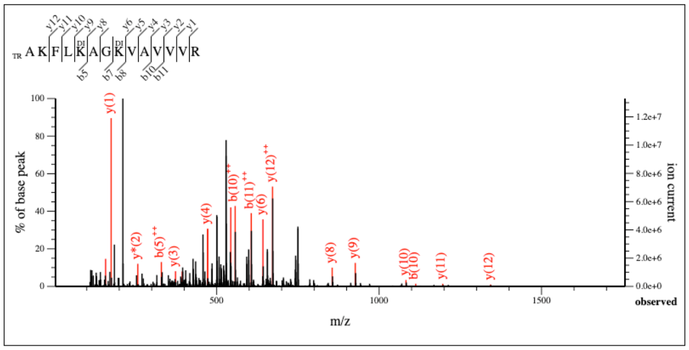

**Monoisotopic mass of neutral peptide Mr(calc):** 1583.0650
**Fixed modifications:** Carbamidomethyl (C) (apply to specified residues or termini only)
**Variable modifications:**
N-term : Trimethyl (Protein N-term)
K5 : Dimethyl (K)
K8 : Dimethyl (K)
**Ions Score:** 46 **Expect:** 0.00019
**Matches :** 20/102 fragment ions using 52 most intense peaks (help)

| # | b | b++ | b* | b*++ | Seq. | y | y++ | y* | y*++ | # |
|---|---|---|---|---|---|---|---|---|---|---|
| 1 | 114.0913 | 57.5493 | | | A | | | | | 14 |
| 2 | 242.1863 | 121.5968 | 225.1598 | 113.0835 | K | 1470.9883 | 735.9978 | 1453.9617 | 727.4845 | 13 |
| 3 | 389.2547 | 195.1310 | 372.2282 | 186.6177 | F | ***1342.8933*** | **671.9503** | 1325.8668 | 663.4370 | 12 |
| 4 | 502.3388 | 251.6730 | 485.3122 | 243.1598 | L | ***1195.8249*** | 598.4161 | 1178.7983 | 589.9028 | 11 |
| 5 | 658.4650 | **329.7362** | 641.4385 | 321.2229 | K | ***1082.7408*** | 541.8741 | 1065.7143 | 533.3608 | 10 |
| 6 | 729.5022 | 365.2547 | 712.4756 | 356.7414 | A | ***926.6146*** | 463.8109 | 909.5880 | 455.2976 | 9 |
| 7 | 786.5236 | **393.7654** | 769.4971 | 385.2522 | G | ***855.5774*** | 428.2924 | 838.5509 | 419.7791 | 8 |
| 8 | 942.6499 | **471.8286** | 925.6233 | 463.3153 | K | 798.5560 | 399.7816 | 781.5294 | 391.2684 | 7 |
| 9 | 1041.7183 | 521.3628 | 1024.6917 | 512.8495 | V | ***642.4297*** | 321.7185 | 625.4032 | 313.2052 | 6 |
| 10 | **1112.7554** | **556.8813** | 1095.7289 | 548.3681 | A | ***543.3613*** | 272.1843 | 526.3348 | 263.6710 | 5 |
| 11 | **1211.8238** | **606.4156** | 1194.7973 | 597.9023 | V | ***472.3242*** | 236.6657 | 455.2976 | 228.1525 | 4 |
| 12 | 1310.8922 | 655.9498 | 1293.8657 | 647.4365 | V | ***373.2558*** | 187.1315 | 356.2292 | 178.6183 | 3 |
| 13 | 1409.9607 | 705.4840 | 1392.9341 | 696.9707 | V | 274.1874 | 137.5973 | 257.1608 | 129.0840 | 2 |
| 14 | | | | | R | ***175.1190*** | 88.0631 | 158.0924 | 79.5498 | 1 |

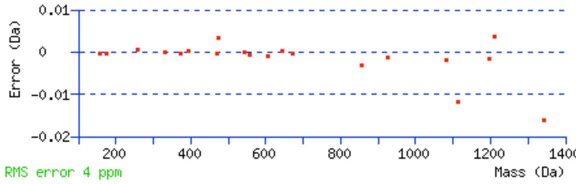


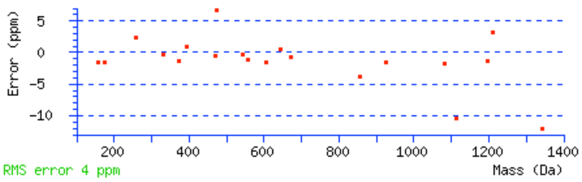


MS/MS Fragmentation of **AKFLKAGKVAVVVR**
Found in **RL27A_YEAST** in **Uniprot_S288C_yeast**, sp|P0C2H6|RL27A_YEAST 60S ribosomal protein L27-A OS=Saccharomyces cerevisiae (strain ATCC 204508 / S288c) OX=559292 GN=RPL27A PE=1 SV=1

Match to Query 1665: 1583.065630 from(792.540091,2+) intensity(48775780.0000) rtinseconds(1909.81716) index(4276)
Title: qExactive02202_+5.7431.7431.2
Data file D:\Panyue\enriched_fr.+5.mzML

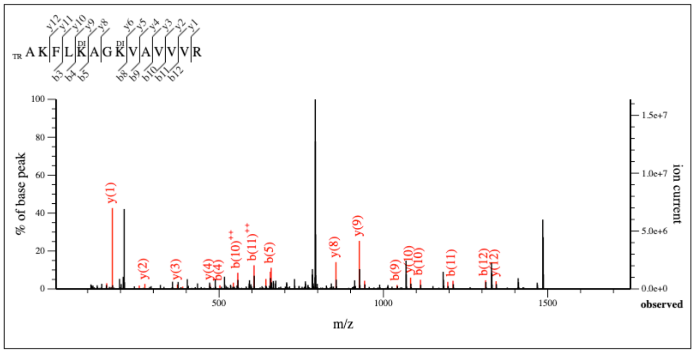

**Monoisotopic mass of neutral peptide Mr(calc):** 1583.0650
**Fixed modifications:** Carbamidomethyl (C) (apply to specified residues or termini only)
**Variable modifications:**
  N-term : Trimethyl (Protein N-term)
  K5     : Dimethyl (K)
  K8     : Dimethyl (K)
**Ions Score:** 35  **Expect:** 0.0024
**Matches :** 24/102 fragment ions using 89 most intense peaks  (help)

| # | b | b++ | b* | b*++ | Seq. | y | y++ | y* | y*++ | # |
|---|---|---|---|---|---|---|---|---|---|---|
| 1 | 114.0913 | 57.5493 | | | A | | | | | 14 |
| 2 | 242.1863 | 121.5968 | 225.1598 | 113.0835 | K | 1470.9883 | 735.9978 | 1453.9617 | 727.4845 | 13 |
| 3 | 389.2547 | 195.1310 | 372.2282 | 186.6177 | F | 1342.8933 | 671.9503 | 1325.8668 | 663.4370 | 12 |
| 4 | 502.3388 | 251.6730 | 485.3122 | 243.1598 | L | 1195.8249 | 598.4161 | 1178.7983 | 589.9028 | 11 |
| 5 | 658.4650 | 329.7362 | 641.4385 | 321.2229 | K | 1082.7408 | 541.8741 | 1065.7143 | 533.3608 | 10 |
| 6 | 729.5022 | 365.2547 | 712.4756 | 356.7414 | A | 926.6146 | 463.8109 | 909.5880 | 455.2976 | 9 |
| 7 | 786.5236 | 393.7654 | 769.4971 | 385.2522 | G | 855.5774 | 428.2924 | 838.5509 | 419.7791 | 8 |
| 8 | 942.6499 | 471.8286 | 925.6233 | 463.3153 | K | 798.5560 | 399.7816 | 781.5294 | 391.2684 | 7 |
| 9 | 1041.7183 | 521.3628 | 1024.6917 | 512.8495 | V | 642.4297 | 321.7185 | 625.4032 | 313.2052 | 6 |
| 10 | 1112.7554 | 556.8813 | 1095.7289 | 548.3681 | A | 543.3613 | 272.1843 | 526.3348 | 263.6710 | 5 |
| 11 | 1211.8238 | 606.4156 | 1194.7973 | 597.9023 | V | 472.3242 | 236.6657 | 455.2976 | 228.1525 | 4 |
| 12 | 1310.8922 | 655.9498 | 1293.8657 | 647.4365 | V | 373.2558 | 187.1315 | 356.2292 | 178.6183 | 3 |
| 13 | 1409.9607 | 705.4840 | 1392.9341 | 696.9707 | V | 274.1874 | 137.5973 | 257.1608 | 129.0840 | 2 |
| 14 | | | | | R | 175.1190 | 88.0631 | 158.0924 | 79.5498 | 1 |

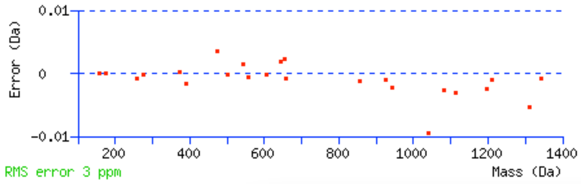


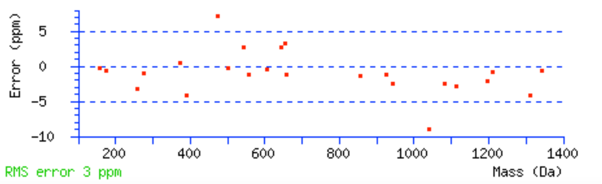


MS/MS Fragmentation of **AKFLKAGKVAVVVR**
Found in **RL27A_YEAST** in **Uniprot_S288C_yeast**, sp|P0C2H6|RL27A_YEAST 60S ribosomal protein L27-A OS=Saccharomyces cerevisiae (strain ATCC 204508 / S288c) OX=559292 GN=RPL27A PE=1 SV=1

Match to Query 1666: 1583.066007 from(528.695945,3+) intensity(83419600.0000) rtinseconds(1904.30082) index(4265)
Title: qExactive02202_+5.7405.7405.3
Data file D:\Panyue\enriched_fr.+5.mzML

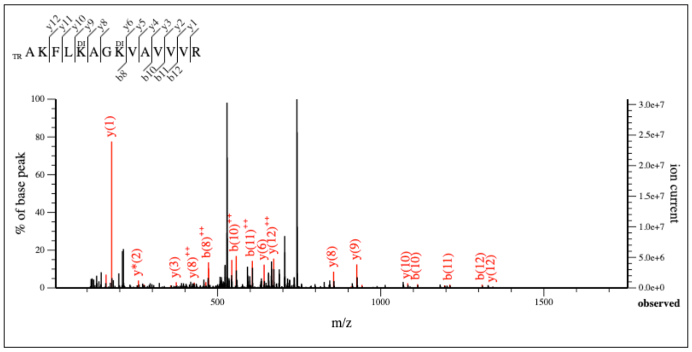

**Monoisotopic mass of neutral peptide Mr(calc):** 1583.0650
**Fixed modifications:** Carbamidomethyl (C) (apply to specified residues or termini only)
**Variable modifications:**
N-term : Trimethyl (Protein N-term)
K5 : Dimethyl (K)
K8 : Dimethyl (K)
**Ions Score:** 30 **Expect:** 0.0078
**Matches :** 25/102 fragment ions using 104 most intense peaks (help)

| # | b | b++ | b* | b*++ | Seq. | y | y++ | y* | y*++ | # |
|---|---|---|---|---|---|---|---|---|---|---|
| 1 | 114.0913 | 57.5493 | | | A | | | | | 14 |
| 2 | 242.1863 | 121.5968 | 225.1598 | 113.0835 | K | 1470.9883 | 735.9978 | 1453.9617 | 727.4845 | 13 |
| 3 | 389.2547 | 195.1310 | 372.2282 | 186.6177 | F | 1342.8933 | 671.9503 | 1325.8668 | 663.4370 | 12 |
| 4 | 502.3388 | 251.6730 | 485.3122 | 243.1598 | L | 1195.8249 | 598.4161 | 1178.7983 | 589.9028 | 11 |
| 5 | 658.4650 | 329.7362 | 641.4385 | 321.2229 | K | 1082.7408 | 541.8741 | 1065.7143 | 533.3608 | 10 |
| 6 | 729.5022 | 365.2547 | 712.4756 | 356.7414 | A | 926.6146 | 463.8109 | 909.5880 | 455.2976 | 9 |
| 7 | 786.5236 | 393.7654 | 769.4971 | 385.2522 | G | 855.5774 | 428.2924 | 838.5509 | 419.7791 | 8 |
| 8 | 942.6499 | 471.8286 | 925.6233 | 463.3153 | K | 798.5560 | 399.7816 | 781.5294 | 391.2684 | 7 |
| 9 | 1041.7183 | 521.3628 | 1024.6917 | 512.8495 | V | 642.4297 | 321.7185 | 625.4032 | 313.2052 | 6 |
| 10 | 1112.7554 | 556.8813 | 1095.7289 | 548.3681 | A | 543.3613 | 272.1843 | 526.3348 | 263.6710 | 5 |
| 11 | 1211.8238 | 606.4156 | 1194.7973 | 597.9023 | V | 472.3242 | 236.6657 | 455.2976 | 228.1525 | 4 |
| 12 | 1310.8922 | 655.9498 | 1293.8657 | 647.4365 | V | 373.2558 | 187.1315 | 356.2292 | 178.6183 | 3 |
| 13 | 1409.9607 | 705.4840 | 1392.9341 | 696.9707 | V | 274.1874 | 137.5973 | 257.1608 | 129.0840 | 2 |
| 14 | | | | | R | 175.1190 | 88.0631 | 158.0924 | 79.5498 | 1 |

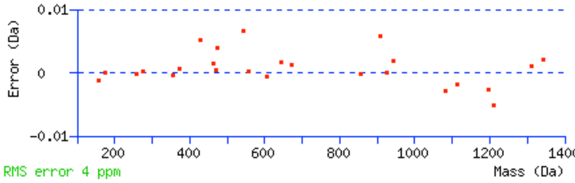


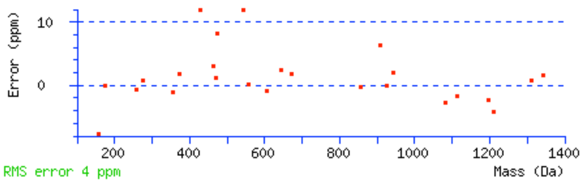


MS/MS Fragmentation of **AKFLKAGKVAVVVR**
Found in **RL27A_YEAST** in **Uniprot_S288C_yeast**, sp|P0C2H6|RL27A_YEAST 60S ribosomal protein L27-A OS=Saccharomyces cerevisiae (strain ATCC 204508 / S288c) OX=559292 GN=RPL27A PE=1 SV=1

Match to Query 1667: 1583.066196 from(396.773825,4+) intensity(10738310.0000) rtinseconds(1897.61472) index(4258)
Title: qExactive02202_+5.7378.7378.4
Data file D:\Panyue\enriched_fr.+5.mzML

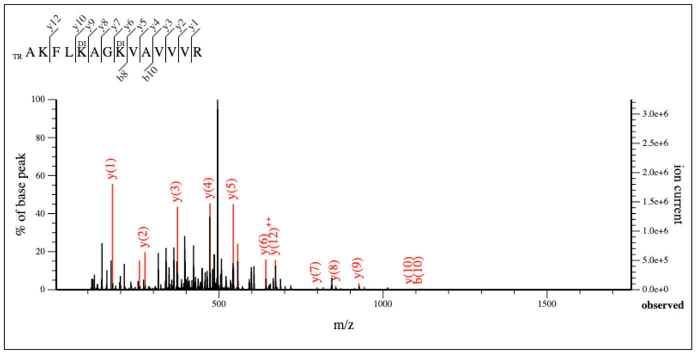

**Monoisotopic mass of neutral peptide Mr(calc):** 1583.0650
**Fixed modifications:** Carbamidomethyl (C) (apply to specified residues or termini only)
**Variable modifications:**
N-term : Trimethyl (Protein N-term)
K5 : Dimethyl (K)
K8 : Dimethyl (K)
**Ions Score:** 59 **Expect:** 9.2e-06
**Matches :** 15/102 fragment ions using 32 most intense peaks (help)

| # | b | b++ | b* | b*++ | Seq. | y | y++ | y* | y*++ | # |
|---|---|---|---|---|---|---|---|---|---|---|
| 1 | 114.0913 | 57.5493 | | | A | | | | | 14 |
| 2 | 242.1863 | 121.5968 | 225.1598 | 113.0835 | K | 1470.9883 | 735.9978 | 1453.9617 | 727.4845 | 13 |
| 3 | 389.2547 | 195.1310 | 372.2282 | 186.6177 | F | 1342.8933 | **671.9503** | 1325.8668 | 663.4370 | 12 |
| 4 | 502.3388 | 251.6730 | 485.3122 | 243.1598 | L | 1195.8249 | 598.4161 | 1178.7983 | 589.9028 | 11 |
| 5 | 658.4650 | 329.7362 | 641.4385 | 321.2229 | K | ***1082.7408*** | 541.8741 | 1065.7143 | 533.3608 | 10 |
| 6 | 729.5022 | 365.2547 | 712.4756 | 356.7414 | A | ***926.6146*** | 463.8109 | 909.5880 | 455.2976 | 9 |
| 7 | 786.5236 | 393.7654 | 769.4971 | 385.2522 | G | ***855.5774*** | 428.2924 | 838.5509 | 419.7791 | 8 |
| 8 | **942.6499** | 471.8286 | 925.6233 | 463.3153 | K | ***798.5560*** | 399.7816 | 781.5294 | 391.2684 | 7 |
| 9 | 1041.7183 | 521.3628 | 1024.6917 | 512.8495 | V | ***642.4297*** | 321.7185 | 625.4032 | 313.2052 | 6 |
| 10 | **1112.7554** | **556.8813** | 1095.7289 | 548.3681 | A | ***543.3613*** | 272.1843 | 526.3348 | 263.6710 | 5 |
| 11 | 1211.8238 | 606.4156 | 1194.7973 | 597.9023 | V | ***472.3242*** | 236.6657 | 455.2976 | 228.1525 | 4 |
| 12 | 1310.8922 | 655.9498 | 1293.8657 | 647.4365 | V | ***373.2558*** | 187.1315 | 356.2292 | 178.6183 | 3 |
| 13 | 1409.9607 | 705.4840 | 1392.9341 | 696.9707 | V | ***274.1874*** | 137.5973 | 257.1608 | 129.0840 | 2 |
| 14 | | | | | R | ***175.1190*** | 88.0631 | 158.0924 | 79.5498 | 1 |

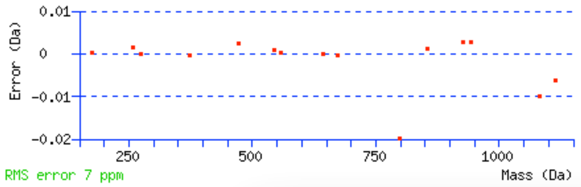


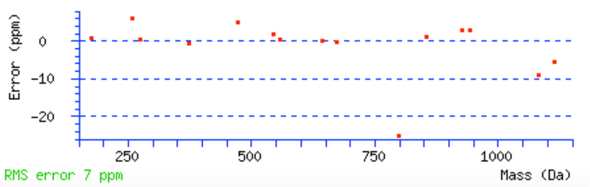


MS/MS Fragmentation of **AKFLKAGKVAVVVR**
Found in **RL27A_YEAST** in **Uniprot_S288C_yeast**, sp|P0C2H6|RL27A_YEAST 60S ribosomal protein L27-A OS=Saccharomyces cerevisiae (strain ATCC 204508 / S288c) OX=559292 GN=RPL27A PE=1 SV=1

Match to Query 1669: 1583.067504 from(528.696444,3+) intensity(3420148.7500) rtinseconds(1956.40164) index(4389)
Title: qExactive02202_+5.7630.7630.3
Data file D:\Panyue\enriched_fr.+5.mzML

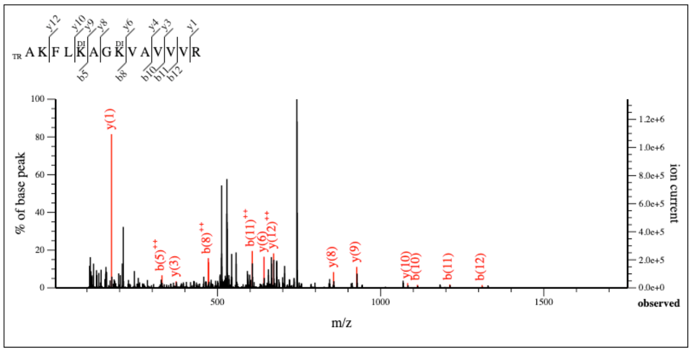

**Monoisotopic mass of neutral peptide Mr(calc):** 1583.0650
**Fixed modifications:** Carbamidomethyl (C) (apply to specified residues or termini only)
**Variable modifications:**
N-term : Trimethyl (Protein N-term)
K5 : Dimethyl (K)
K8 : Dimethyl (K)
**Ions Score:** 22 **Expect:** 0.05
**Matches :** 14/102 fragment ions using 39 most intense peaks (help)

| # | b | b++ | b* | b*++ | Seq. | y | y++ | y* | y*++ | # |
|---|---|---|---|---|---|---|---|---|---|---|
| 1 | 114.0913 | 57.5493 | | | A | | | | | 14 |
| 2 | 242.1863 | 121.5968 | 225.1598 | 113.0835 | K | 1470.9883 | 735.9978 | 1453.9617 | 727.4845 | 13 |
| 3 | 389.2547 | 195.1310 | 372.2282 | 186.6177 | F | 1342.8933 | **671.9503** | 1325.8668 | 663.4370 | 12 |
| 4 | 502.3388 | 251.6730 | 485.3122 | 243.1598 | L | 1195.8249 | 598.4161 | 1178.7983 | 589.9028 | 11 |
| 5 | 658.4650 | **329.7362** | 641.4385 | 321.2229 | K | ***1082.7408*** | 541.8741 | 1065.7143 | 533.3608 | 10 |
| 6 | 729.5022 | 365.2547 | 712.4756 | 356.7414 | A | ***926.6146*** | 463.8109 | 909.5880 | 455.2976 | 9 |
| 7 | 786.5236 | 393.7654 | 769.4971 | 385.2522 | G | ***855.5774*** | 428.2924 | 838.5509 | 419.7791 | 8 |
| 8 | 942.6499 | **471.8286** | 925.6233 | 463.3153 | K | 798.5560 | 399.7816 | 781.5294 | 391.2684 | 7 |
| 9 | 1041.7183 | 521.3628 | 1024.6917 | 512.8495 | V | ***642.4297*** | 321.7185 | 625.4032 | 313.2052 | 6 |
| 10 | **1112.7554** | 556.8813 | 1095.7289 | 548.3681 | A | 543.3613 | 272.1843 | 526.3348 | 263.6710 | 5 |
| 11 | **1211.8238** | **606.4156** | 1194.7973 | 597.9023 | V | ***472.3242*** | 236.6657 | 455.2976 | 228.1525 | 4 |
| 12 | **1310.8922** | 655.9498 | 1293.8657 | 647.4365 | V | ***373.2558*** | 187.1315 | 356.2292 | 178.6183 | 3 |
| 13 | 1409.9607 | 705.4840 | 1392.9341 | 696.9707 | V | 274.1874 | 137.5973 | 257.1608 | 129.0840 | 2 |
| 14 | | | | | R | ***175.1190*** | 88.0631 | 158.0924 | 79.5498 | 1 |

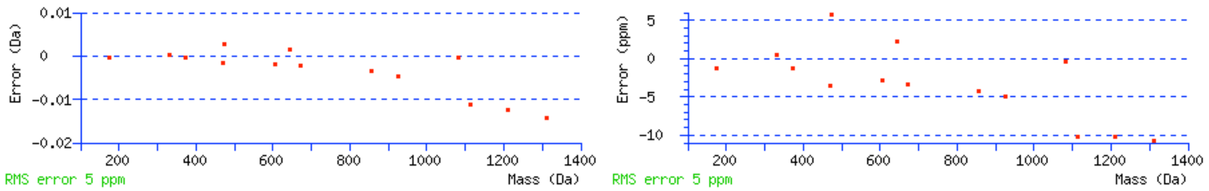


MS/MS Fragmentation of **AKFLKAGKVAVVVR**
Found in **RL27A_YEAST** in **Uniprot_S288C_yeast**, sp|P0C2H6|RL27A_YEAST 60S ribosomal protein L27-A OS=Saccharomyces cerevisiae (strain ATCC 204508 / S288c) OX=559292 GN=RPL27A PE=1 SV=1

Match to Query 1670: 1583.067753 from(528.696527,3+) intensity(21590696.0000) rtinseconds(1921.49454) index(4308)
Title: qExactive02202_+5.7486.7486.3
Data file D:\Panyue\enriched_fr.+5.mzML

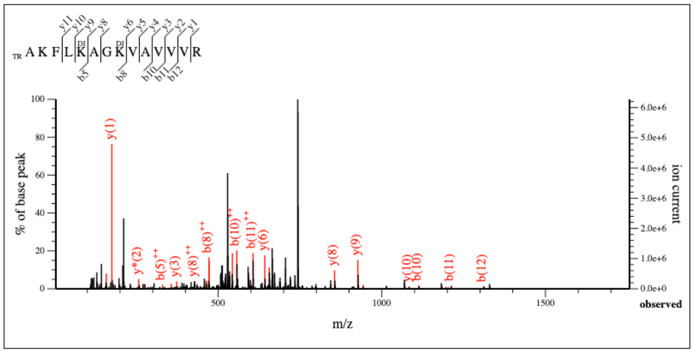

**Monoisotopic mass of neutral peptide Mr(calc):** 1583.0650
**Fixed modifications:** Carbamidomethyl (C) (apply to specified residues or termini only)
**Variable modifications:**
N-term : Trimethyl (Protein N-term)
K5 : Dimethyl (K)
K8 : Dimethyl (K)
**Ions Score:** 27 **Expect:** 0.014
**Matches :** 24/102 fragment ions using 91 most intense peaks (help)

| # | b | b++ | b* | b*++ | Seq. | y | y++ | y* | y*++ | # |
|---|---|---|---|---|---|---|---|---|---|---|
| 1 | 114.0913 | 57.5493 | | | A | | | | | 14 |
| 2 | 242.1863 | 121.5968 | 225.1598 | 113.0835 | K | 1470.9883 | 735.9978 | 1453.9617 | 727.4845 | 13 |
| 3 | 389.2547 | 195.1310 | 372.2282 | 186.6177 | F | 1342.8933 | 671.9503 | 1325.8668 | 663.4370 | 12 |
| 4 | 502.3388 | 251.6730 | 485.3122 | 243.1598 | L | 1195.8249 | 598.4161 | 1178.7983 | 589.9028 | 11 |
| 5 | 658.4650 | 329.7362 | 641.4385 | 321.2229 | K | 1082.7408 | 541.8741 | 1065.7143 | 533.3608 | 10 |
| 6 | 729.5022 | 365.2547 | 712.4756 | 356.7414 | A | 926.6146 | 463.8109 | 909.5880 | 455.2976 | 9 |
| 7 | 786.5236 | 393.7654 | 769.4971 | 385.2522 | G | 855.5774 | 428.2924 | 838.5509 | 419.7791 | 8 |
| 8 | 942.6499 | 471.8286 | 925.6233 | 463.3153 | K | 798.5560 | 399.7816 | 781.5294 | 391.2684 | 7 |
| 9 | 1041.7183 | 521.3628 | 1024.6917 | 512.8495 | V | 642.4297 | 321.7185 | 625.4032 | 313.2052 | 6 |
| 10 | 1112.7554 | 556.8813 | 1095.7289 | 548.3681 | A | 543.3613 | 272.1843 | 526.3348 | 263.6710 | 5 |
| 11 | 1211.8238 | 606.4156 | 1194.7973 | 597.9023 | V | 472.3242 | 236.6657 | 455.2976 | 228.1525 | 4 |
| 12 | 1310.8922 | 655.9498 | 1293.8657 | 647.4365 | V | 373.2558 | 187.1315 | 356.2292 | 178.6183 | 3 |
| 13 | 1409.9607 | 705.4840 | 1392.9341 | 696.9707 | V | 274.1874 | 137.5973 | 257.1608 | 129.0840 | 2 |
| 14 | | | | | R | 175.1190 | 88.0631 | 158.0924 | 79.5498 | 1 |

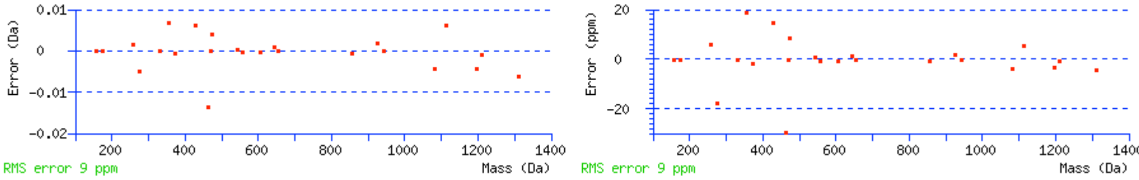


MS/MS Fragmentation of AKFLKAGKVAVVVR
Found in RL27A_YEAST in Uniprot_S288C_yeast, sp|P0C2H6|RL27A_YEAST 60S ribosomal protein L27-A OS=Saccharomyces cerevisiae (strain ATCC 204508 / S288c) OX=559292 GN=RPL27A PE=1 SV=1

Match to Query 1846: 1611.102816 from(538.041548,3+) intensity(1690707.2500) rtinseconds(2029.00464) index(4513)
Title: qExactive02202_+5.7900.7900.3
Data file D:\Panyue\enriched_fr.+5.mzML

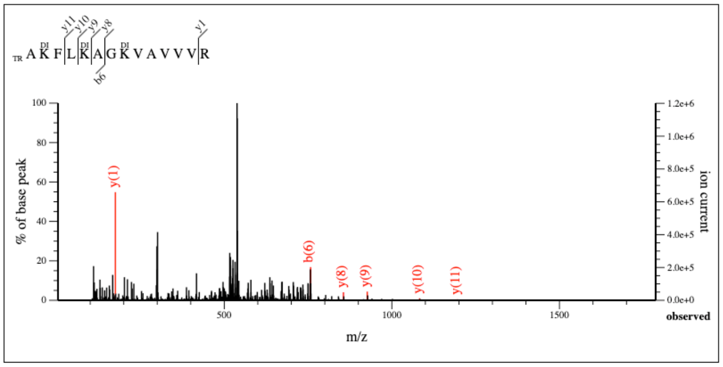

**Monoisotopic mass of neutral peptide Mr(calc):** 1611.0963
**Fixed modifications:** Carbamidomethyl (C) (apply to specified residues or termini only)
**Variable modifications:**
N-term : Trimethyl (Protein N-term)
K2 : Dimethyl (K)
K5 : Dimethyl (K)
K8 : Dimethyl (K)
**Ions Score:** 21 **Expect:** 0.037
**Matches :** 6/102 fragment ions using 11 most intense peaks (help)

| # | b | $b^{++}$ | b* | $b^{*++}$ | Seq. | y | $y^{++}$ | y* | $y^{*++}$ | # |
|---|---|---|---|---|---|---|---|---|---|---|
| 1 | 114.0913 | 57.5493 | | | A | | | | | 14 |
| 2 | 270.2176 | 135.6124 | 253.1911 | 127.0992 | K | 1499.0196 | 750.0134 | 1481.9930 | 741.5001 | 13 |
| 3 | 417.2860 | 209.1466 | 400.2595 | 200.6334 | F | 1342.8933 | 671.9503 | 1325.8668 | 663.4370 | 12 |
| 4 | 530.3701 | 265.6887 | 513.3435 | 257.1754 | L | **1195.8249** | 598.4161 | 1178.7983 | 589.9028 | 11 |
| 5 | 686.4963 | 343.7518 | 669.4698 | 335.2385 | K | **1082.7408** | 541.8741 | 1065.7143 | 533.3608 | 10 |
| 6 | **757.5335** | 379.2704 | 740.5069 | 370.7571 | A | **926.6146** | 463.8109 | 909.5880 | 455.2976 | 9 |
| 7 | 814.5549 | 407.7811 | 797.5284 | 399.2678 | G | **855.5774** | 428.2924 | 838.5509 | 419.7791 | 8 |
| 8 | 970.6812 | 485.8442 | 953.6546 | 477.3310 | K | 798.5560 | 399.7816 | 781.5294 | 391.2684 | 7 |
| 9 | 1069.7496 | 535.3784 | 1052.7230 | 526.8652 | V | 642.4297 | 321.7185 | 625.4032 | 313.2052 | 6 |
| 10 | 1140.7867 | 570.8970 | 1123.7602 | 562.3837 | A | 543.3613 | 272.1843 | 526.3348 | 263.6710 | 5 |
| 11 | 1239.8551 | 620.4312 | 1222.8286 | 611.9179 | V | 472.3242 | 236.6657 | 455.2976 | 228.1525 | 4 |
| 12 | 1338.9235 | 669.9654 | 1321.8970 | 661.4521 | V | 373.2558 | 187.1315 | 356.2292 | 178.6183 | 3 |
| 13 | 1437.9920 | 719.4996 | 1420.9654 | 710.9863 | V | 274.1874 | 137.5973 | 257.1608 | 129.0840 | 2 |
| 14 | | | | | R | **175.1190** | 88.0631 | 158.0924 | 79.5498 | 1 |

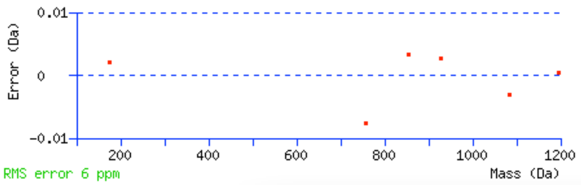


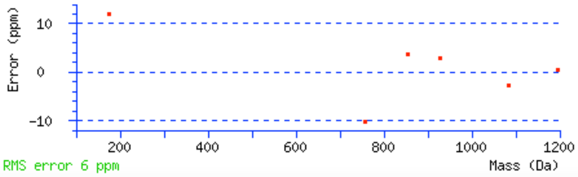

MS/MS spectra for hits from iProx dataset:
*Only spectra of peptide score >20 will be listed here. Peptides have initial M cleaved.

1. Arb1 from yeast fraction 2.

**Peptide View**

MS/MS Fragmentation of **PPVSASK**
Found in **sp|P40024|ARB1_YEAST** in **uniprot-yeast288C**, sp|P40024|ARB1_YEAST ABC transporter ATP-binding protein ARB1 OS=Saccharomyces cerevisiae (strain ATCC 204508 / S288c) OX=559292 GN=ARB1 PE=1 SV=1

Match to Query 241: 720.456248 from(361.235400,2+) intensity(429192.5312) rtinseconds(317.8953028) index(846)
Title: zm-yeast-protein-20151023-fr2.1522.1522.2
Data file D:\Panyue\Proteomic dataset\iprox yeast\zm-yeast-protein-20151023-fr2.raw

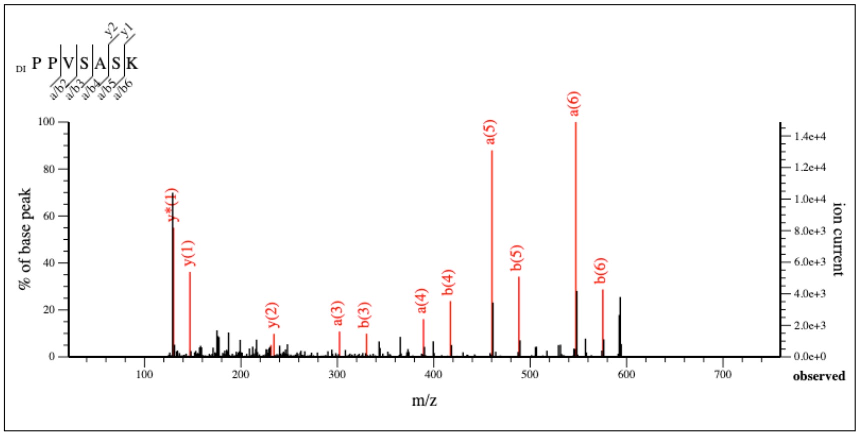


**Monoisotopic mass of neutral peptide Mr(calc):** 720.4563
**Fixed modifications:** Carbamidomethyl (C),Label:13C(1)2H(3) (M) (apply to specified residues or termini only)
**Variable modifications:**
N-term : Dimethyl:2H(6)13C(2) (Protein N-term)
**Ions Score:** 33 **Expect:** 0.00049
**Matches :** 12/48 fragment ions using 22 most intense peaks (help)

| # | a | a++ | b | b++ | Seq. | y | y++ | y* | y*++ | # |
|---|---|---|---|---|---|---|---|---|---|---|
| 1 | 106.1408 | 53.5740 | 134.1357 | 67.5715 | P | | | | | 7 |
| 2 | 203.1936 | 102.1004 | **231.1885** | 116.0979 | P | 588.3352 | 294.6712 | 571.3086 | 286.1579 | 6 |
| 3 | **302.2620** | 151.6346 | **330.2569** | 165.6321 | V | 491.2824 | 246.1448 | 474.2558 | 237.6316 | 5 |
| 4 | **389.2940** | 195.1506 | **417.2889** | 209.1481 | S | 392.2140 | 196.6106 | 375.1874 | 188.0974 | 4 |
| 5 | **460.3311** | 230.6692 | **488.3260** | 244.6667 | A | 305.1819 | 153.0946 | 288.1554 | 144.5813 | 3 |
| 6 | **547.3631** | 274.1852 | **575.3581** | 288.1827 | S | **234.1448** | 117.5761 | 217.1183 | 109.0628 | 2 |
| 7 | | | | | K | **147.1128** | 74.0600 | 130.0863 | 65.5468 | 1 |

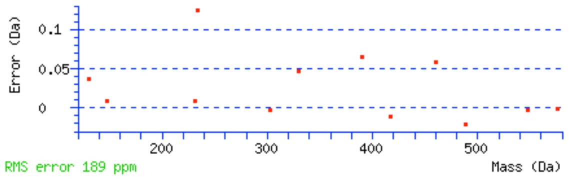


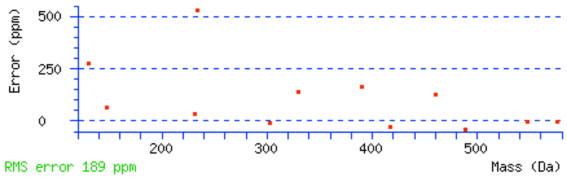


2. Arb1 from yeast fraction 3.

**Peptide View**

MS/MS Fragmentation of **PPVSASK**
Found in **sp|P40024|ARB1_YEAST** in **uniprot-yeast288C**, sp|P40024|ARB1_YEAST ABC transporter ATP-binding protein ARB1 OS=Saccharomyces cerevisiae (strain ATCC 204508 / S288c) OX=559292 GN=ARB1 PE=1 SV=1

Match to Query 211: 720.455048 from(361.234800,2+) intensity(4255985.5000) rtinseconds(292.7093092) index(787)
Title: zm-yeast-protein-20151023-fr3.1400.1400.2
Data file D:\Panyue\Proteomic dataset\iprox yeast\zm-yeast-protein-20151023-fr3.raw

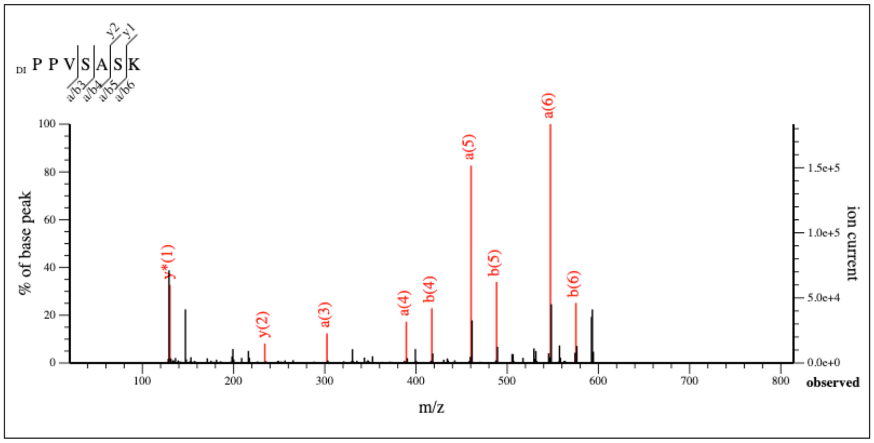


**Monoisotopic mass of neutral peptide Mr(calc):** 720.4563
**Fixed modifications:** Carbamidomethyl (C),Label:13C(1)2H(3) (M) (apply to specified residues or termini only)
**Variable modifications:**
N-term : Dimethyl:2H(6)13C(2) (Protein N-term)
**Ions Score:** 28 **Expect:** 0.0017
**Matches :** 9/48 fragment ions using 12 most intense peaks (help)

| # | a | a++ | b | b++ | Seq. | y | y++ | y* | y*++ | # |
|---|---|---|---|---|---|---|---|---|---|---|
| 1 | 106.1408 | 53.5740 | 134.1357 | 67.5715 | P | | | | | 7 |
| 2 | 203.1936 | 102.1004 | 231.1885 | 116.0979 | P | 588.3352 | 294.6712 | 571.3086 | 286.1579 | 6 |
| 3 | ***302.2620*** | 151.6346 | 330.2569 | 165.6321 | V | 491.2824 | 246.1448 | 474.2558 | 237.6316 | 5 |
| 4 | ***389.2940*** | 195.1506 | **417.2889** | 209.1481 | S | 392.2140 | 196.6106 | 375.1874 | 188.0974 | 4 |
| 5 | ***460.3311*** | 230.6692 | **488.3260** | 244.6667 | A | 305.1819 | 153.0946 | 288.1554 | 144.5813 | 3 |
| 6 | ***547.3631*** | 274.1852 | **575.3581** | 288.1827 | S | 234.1448 | 117.5761 | 217.1183 | 109.0628 | 2 |
| 7 | | | | | K | 147.1128 | 74.0600 | 130.0863 | 65.5468 | 1 |

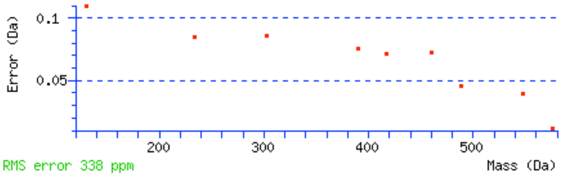


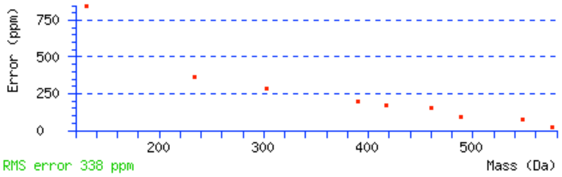


3. Arb1 from yeast fraction 4 (2 spectrums).

#### Peptide View

MS/MS Fragmentation of **PPVSASK**
Found in **sp|P40024|ARB1_YEAST** in **uniprot-yeast288C**, sp|P40024|ARB1_YEAST ABC transporter ATP-binding protein ARB1 OS=Saccharomyces cerevisiae (strain ATCC 204508 / S288c) OX=559292 GN=ARB1 PE=1 SV=1

Match to Query 217: 720.458410 from(361.236481,2+) intensity(155435.4844) rtinseconds(307.0990575) index(690)
Title: zm-yeast-protein-20151023-fr4.1338.1338.2
Data file D:\Panyue\Proteomic dataset\iprox yeast\zm-yeast-protein-20151023-fr4.raw

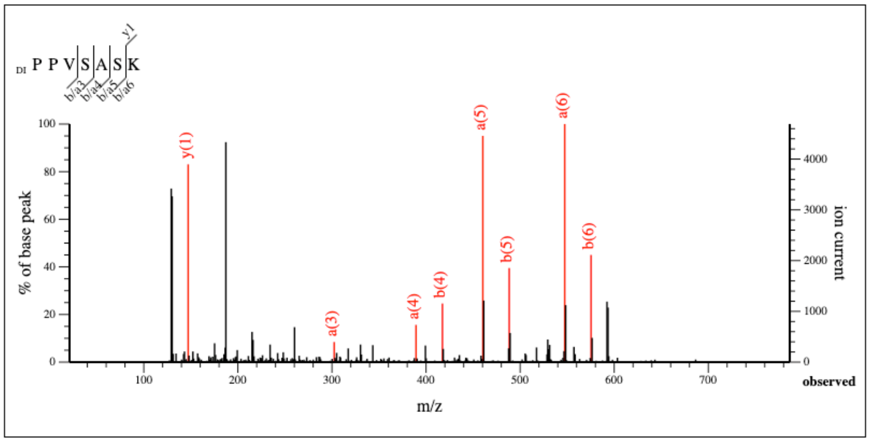


**Monoisotopic mass of neutral peptide Mr(calc):** 720.4563
**Fixed modifications:** Carbamidomethyl (C),Label:13C(1)2H(3) (M) (apply to specified residues or termini only)
**Variable modifications:**
N-term : Dimethyl:2H(6)13C(2) (Protein N-term)
**Ions Score:** 29 **Expect:** 0.0014
**Matches :** 8/48 fragment ions using 12 most intense peaks (help)

| # | a | a++ | b | b++ | Seq. | y | y++ | y* | y*++ | # |
|---|---|---|---|---|---|---|---|---|---|---|
| 1 | 106.1408 | 53.5740 | 134.1357 | 67.5715 | P | | | | | 7 |
| 2 | 203.1936 | 102.1004 | 231.1885 | 116.0979 | P | 588.3352 | 294.6712 | 571.3086 | 286.1579 | 6 |
| 3 | *302.2620* | 151.6346 | 330.2569 | 165.6321 | V | 491.2824 | 246.1448 | 474.2558 | 237.6316 | 5 |
| 4 | *389.2940* | 195.1506 | **417.2889** | 209.1481 | S | 392.2140 | 196.6106 | 375.1874 | 188.0974 | 4 |
| 5 | *460.3311* | 230.6692 | **488.3260** | 244.6667 | A | 305.1819 | 153.0946 | 288.1554 | 144.5813 | 3 |
| 6 | *547.3631* | 274.1852 | **575.3581** | 288.1827 | S | 234.1448 | 117.5761 | 217.1183 | 109.0628 | 2 |
| 7 | | | | | K | 147.1128 | 74.0600 | 130.0863 | 65.5468 | 1 |

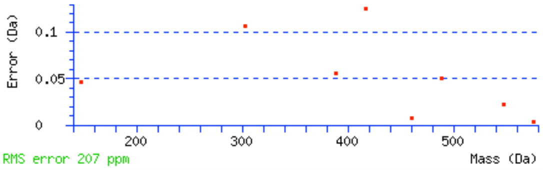


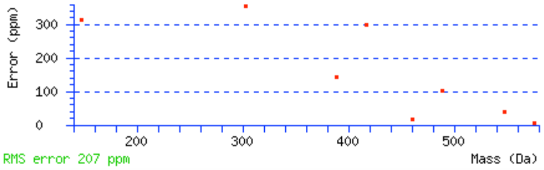

**Peptide View**

MS/MS Fragmentation of **PPVSASK**
Found in **sp|P40024|ARB1_YEAST** in **uniprot-yeast288C**, sp|P40024|ARB1_YEAST ABC transporter ATP-binding protein ARB1 OS=Saccharomyces cerevisiae (strain ATCC 204508 / S288c) OX=559292 GN=ARB1 PE=1 SV=1

Match to Query 216: 720.456030 from(361.235291,2+) intensity(592528.5625) rtinseconds(293.9832653) index(544)
Title: zm-yeast-protein-20151023-fr4.1163.1163.2
Data file D:\Panyue\Proteomic dataset\iprox yeast\zm-yeast-protein-20151023-fr4.raw

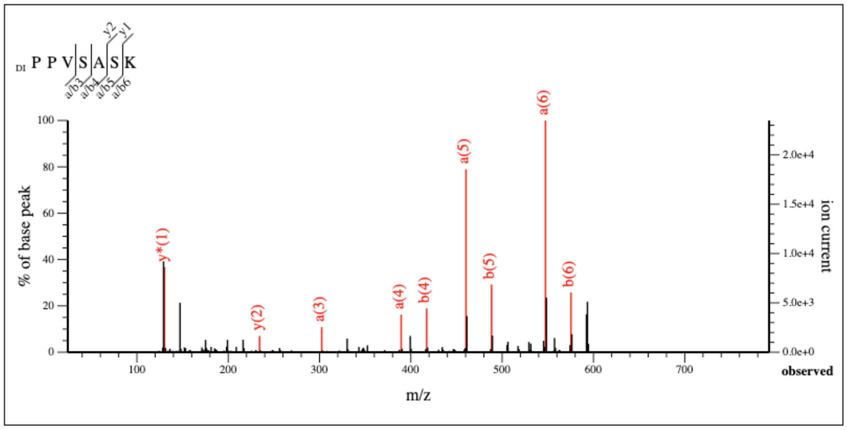


```
Monoisotopic mass of neutral peptide Mr(calc): 720.4563
Fixed modifications: Carbamidomethyl (C),Label:13C(1)2H(3) (M) (apply to specified residues or termini only)
Variable modifications:
  N-term : Dimethyl:2H(6)13C(2) (Protein N-term)
Ions Score: 28  Expect: 0.0018
Matches : 9/48 fragment ions using 12 most intense peaks   (help)
```

| # | a | a++ | b | b++ | Seq. | y | y++ | y* | y*++ | # |
|---|---|---|---|---|---|---|---|---|---|---|
| 1 | 106.1408 | 53.5740 | 134.1357 | 67.5715 | P | | | | | 7 |
| 2 | 203.1936 | 102.1004 | 231.1885 | 116.0979 | P | 588.3352 | 294.6712 | 571.3086 | 286.1579 | 6 |
| 3 | 302.2620 | 151.6346 | 330.2569 | 165.6321 | V | 491.2824 | 246.1448 | 474.2558 | 237.6316 | 5 |
| 4 | 389.2940 | 195.1506 | 417.2889 | 209.1481 | S | 392.2140 | 196.6106 | 375.1874 | 188.0974 | 4 |
| 5 | 460.3311 | 230.6692 | 488.3260 | 244.6667 | A | 305.1819 | 153.0946 | 288.1554 | 144.5813 | 3 |
| 6 | 547.3631 | 274.1852 | 575.3581 | 288.1827 | S | 234.1448 | 117.5761 | 217.1183 | 109.0628 | 2 |
| 7 | | | | | K | 147.1128 | 74.0600 | 130.0863 | 65.5468 | 1 |

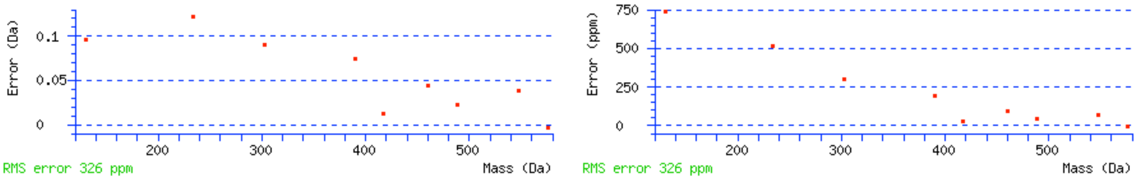


4. Arb1 from yeast fraction 5.

**Peptide View**

MS/MS Fragmentation of **PPVSASK**
Found in **sp|P40024|ARB1_YEAST** in **uniprot-yeast288C**, sp|P40024|ARB1_YEAST ABC transporter ATP-binding protein ARB1 OS=Saccharomyces cerevisiae (strain ATCC 204508 / S288c) OX=559292 GN=ARB1 PE=1 SV=1

Match to Query 224: 720.455784 from(361.235168,2+) intensity(957666.5625) rtinseconds(309.0344023) index(917)
Title: zm-yeast-protein-20151023-fr5.1571.1571.2
Data file D:\Panyue\Proteomic dataset\iprox yeast\zm-yeast-protein-20151023-fr5.raw

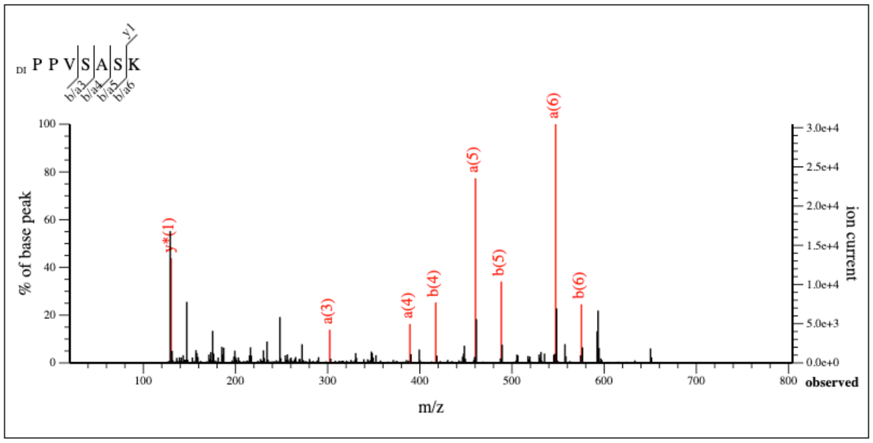


**Monoisotopic mass of neutral peptide Mr(calc):** 720.4563
**Fixed modifications:** Carbamidomethyl (C),Label:13C(1)2H(3) (M) (apply to specified residues or termini only)
**Variable modifications:**
N-term : Dimethyl:2H(6)13C(2) (Protein N-term)
**Ions Score:** 27 **Expect:** 0.0021
**Matches :** 8/48 fragment ions using 12 most intense peaks (help)

| # | a | a++ | b | b++ | Seq. | y | y++ | y* | y*++ | # |
|---|---|---|---|---|---|---|---|---|---|---|
| 1 | 106.1408 | 53.5740 | 134.1357 | 67.5715 | P | | | | | 7 |
| 2 | 203.1936 | 102.1004 | 231.1885 | 116.0979 | P | 588.3352 | 294.6712 | 571.3086 | 286.1579 | 6 |
| 3 | *302.2620* | 151.6346 | 330.2569 | 165.6321 | V | 491.2824 | 246.1448 | 474.2558 | 237.6316 | 5 |
| 4 | *389.2940* | 195.1506 | **417.2889** | 209.1481 | S | 392.2140 | 196.6106 | 375.1874 | 188.0974 | 4 |
| 5 | *460.3311* | 230.6692 | **488.3260** | 244.6667 | A | 305.1819 | 153.0946 | 288.1554 | 144.5813 | 3 |
| 6 | *547.3631* | 274.1852 | **575.3581** | 288.1827 | S | 234.1448 | 117.5761 | 217.1183 | 109.0628 | 2 |
| 7 | | | | | K | 147.1128 | 74.0600 | 130.0863 | 65.5468 | 1 |

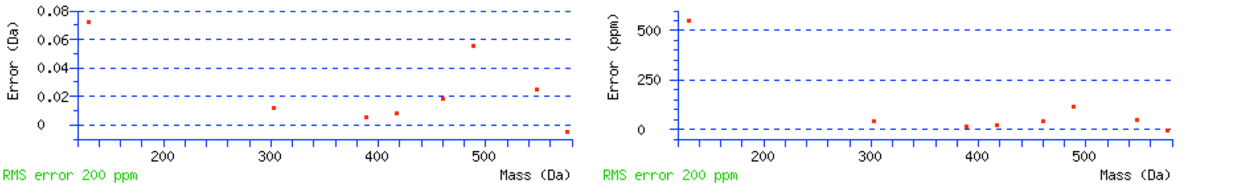


5. Arb1 from yeast fraction 6.

**Peptide View**

MS/MS Fragmentation of PPVSASK
Found in sp|P40024|ARB1_YEAST in uniprot-yeast288C, sp|P40024|ARB1_YEAST ABC transporter ATP-binding protein ARB1 OS=Saccharomyces cerevisiae (strain ATCC 204508 / S288c) OX=559292 GN=ARB1 PE=1 SV=1

Match to Query 232: 720.456640 from(361.235596,2+) intensity(380683.7812) rtinseconds(352.1012941) index(911)
Title: zm-yeast-protein-20151023-fr6.1655.1655.2
Data file D:\Panyue\Proteomic dataset\iprox yeast\zm-yeast-protein-20151023-fr6.raw

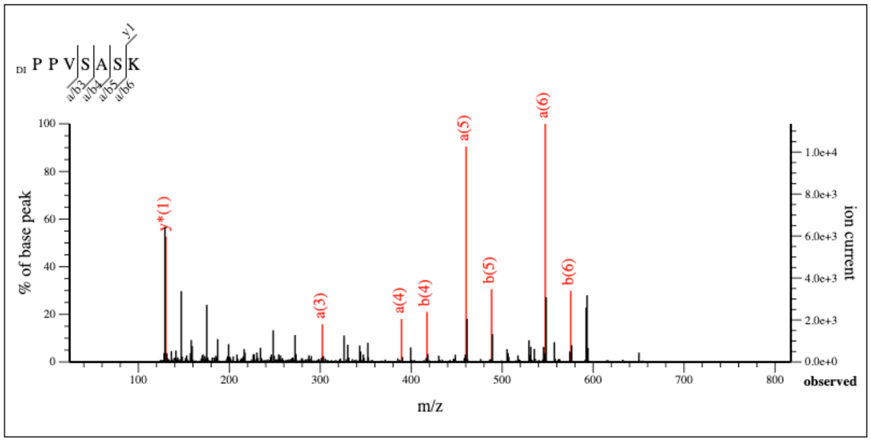


**Monoisotopic mass of neutral peptide Mr(calc):** 720.4563
**Fixed modifications:** Carbamidomethyl (C),Label:13C(1)2H(3) (M) (apply to specified residues or termini only)
**Variable modifications:**
N-term : Dimethyl:2H(6)13C(2) (Protein N-term)
**Ions Score:** 27 **Expect:** 0.0022
**Matches :** 8/48 fragment ions using 12 most intense peaks (help)

| # | a | a++ | b | b++ | Seq. | y | y++ | y* | y*++ | # |
|---|---|---|---|---|---|---|---|---|---|---|
| 1 | 106.1408 | 53.5740 | 134.1357 | 67.5715 | P | | | | | 7 |
| 2 | 203.1936 | 102.1004 | 231.1885 | 116.0979 | P | 588.3352 | 294.6712 | 571.3086 | 286.1579 | 6 |
| 3 | ***302.2620*** | 151.6346 | 330.2569 | 165.6321 | V | 491.2824 | 246.1448 | 474.2558 | 237.6316 | 5 |
| 4 | ***389.2940*** | 195.1506 | **417.2889** | 209.1481 | S | 392.2140 | 196.6106 | 375.1874 | 188.0974 | 4 |
| 5 | ***460.3311*** | 230.6692 | **488.3260** | 244.6667 | A | 305.1819 | 153.0946 | 288.1554 | 144.5813 | 3 |
| 6 | ***547.3631*** | 274.1852 | **575.3581** | 288.1827 | S | 234.1448 | 117.5761 | 217.1183 | 109.0628 | 2 |
| 7 | | | | | K | 147.1128 | 74.0600 | 130.0863 | 65.5468 | 1 |

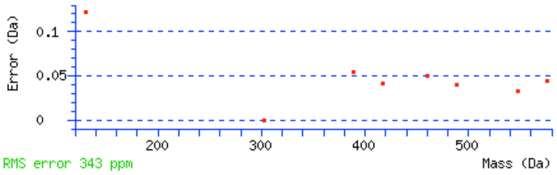


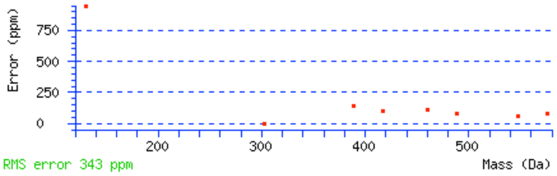


6. Arb1 from yeast fraction 7.

**Peptide View**

MS/MS Fragmentation of **PPVSASK**
Found in **sp|P40024|ARB1_YEAST** in **uniprot-yeast288C**, sp|P40024|ARB1_YEAST ABC transporter ATP-binding protein ARB1 OS=Saccharomyces cerevisiae (strain ATCC 204508 / S288c) OX=559292 GN=ARB1 PE=1 SV=1

Match to Query 237: 720.457372 from(361.235962,2+) intensity(1765692.6250) rtinseconds(315.7011797) index(925)
Title: zm-yeast-protein-20151023-fr7.1581.1581.2
Data file D:\Panyue\Proteomic dataset\iprox yeast\zm-yeast-protein-20151023-fr7.raw

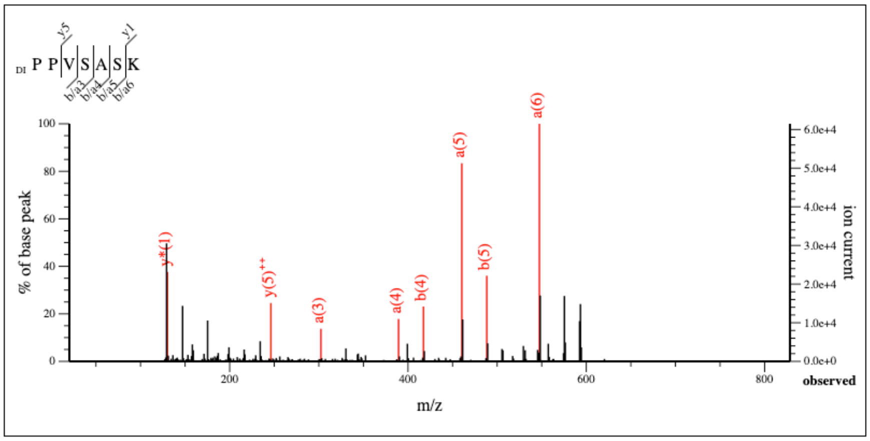


**Monoisotopic mass of neutral peptide Mr(calc):** 720.4563
**Fixed modifications:** Carbamidomethyl (C),Label:13C(1)2H(3) (M) (apply to specified residues or termini only)
**Variable modifications:**
N-term : Dimethyl:2H(6)13C(2) (Protein N-term)
**Ions Score:** 25 **Expect:** 0.0034
**Matches :** 8/48 fragment ions using 12 most intense peaks (help)

| # | a | a++ | b | b++ | Seq. | y | y++ | y* | y*++ | # |
|---|---|---|---|---|---|---|---|---|---|---|
| 1 | 106.1408 | 53.5740 | 134.1357 | 67.5715 | P | | | | | 7 |
| 2 | 203.1936 | 102.1004 | 231.1885 | 116.0979 | P | 588.3352 | 294.6712 | 571.3086 | 286.1579 | 6 |
| 3 | ***302.2620*** | 151.6346 | 330.2569 | 165.6321 | V | 491.2824 | 246.1448 | 474.2558 | 237.6316 | 5 |
| 4 | ***389.2940*** | 195.1506 | **417.2889** | 209.1481 | S | 392.2140 | 196.6106 | 375.1874 | 188.0974 | 4 |
| 5 | ***460.3311*** | 230.6692 | **488.3260** | 244.6667 | A | 305.1819 | 153.0946 | 288.1554 | 144.5813 | 3 |
| 6 | ***547.3631*** | 274.1852 | 575.3581 | 288.1827 | S | 234.1448 | 117.5761 | 217.1183 | 109.0628 | 2 |
| 7 | | | | | K | 147.1128 | 74.0600 | 130.0863 | 65.5468 | 1 |

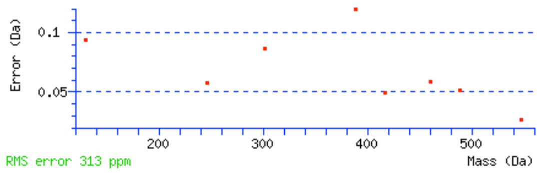


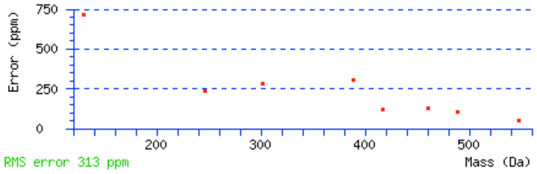


7. Arb1 from yeast fraction 8.

**Peptide View**

MS/MS Fragmentation of **PPVSASK**
Found in **sp|P40024|ARB1_YEAST** in **uniprot-yeast288C**, sp|P40024|ARB1_YEAST ABC transporter ATP-binding protein ARB1 OS=Saccharomyces cerevisiae (strain ATCC 204508 / S288c) OX=559292 GN=ARB1 PE=1 SV=1

Match to Query 225: 720.456456 from(361.235504,2+) intensity(370763.4688) rtinseconds(320.0170276) index(1046)
Title: zm-yeast-protein-20151023-fr8.1716.1716.2
Data file D:\Panyue\Proteomic dataset\iprox yeast\zm-yeast-protein-20151023-fr8.raw

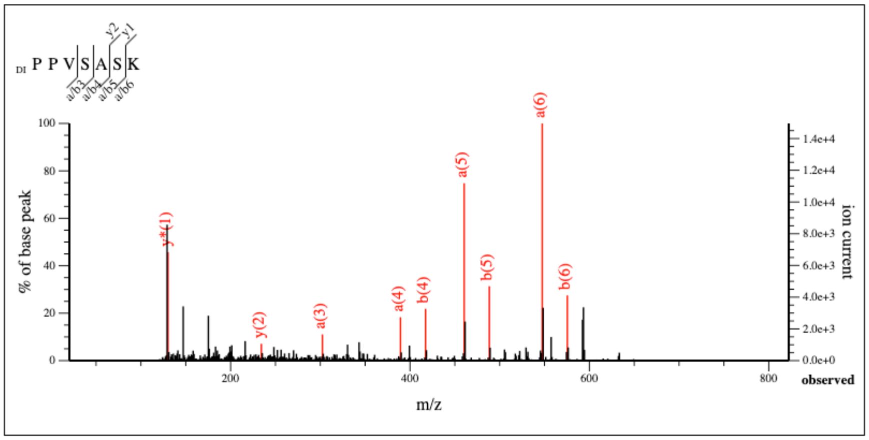

**Monoisotopic mass of neutral peptide Mr(calc):** 720.4563
**Fixed modifications:** Carbamidomethyl (C),Label:13C(1)2H(3) (M) (apply to specified residues or termini only)
**Variable modifications:**
N-term : Dimethyl:2H(6)13C(2) (Protein N-term)
**Ions Score:** 27 **Expect:** 0.002
**Matches :** 9/48 fragment ions using 12 most intense peaks (help)

| # | a | a++ | b | b++ | Seq. | y | y++ | y* | y*++ | # |
|---|---|---|---|---|---|---|---|---|---|---|
| 1 | 106.1408 | 53.5740 | 134.1357 | 67.5715 | P | | | | | 7 |
| 2 | 203.1936 | 102.1004 | 231.1885 | 116.0979 | P | 588.3352 | 294.6712 | 571.3086 | 286.1579 | 6 |
| 3 | 302.2620 | 151.6346 | 330.2569 | 165.6321 | V | 491.2824 | 246.1448 | 474.2558 | 237.6316 | 5 |
| 4 | 389.2940 | 195.1506 | 417.2889 | 209.1481 | S | 392.2140 | 196.6106 | 375.1874 | 188.0974 | 4 |
| 5 | 460.3311 | 230.6692 | 488.3260 | 244.6667 | A | 305.1819 | 153.0946 | 288.1554 | 144.5813 | 3 |
| 6 | 547.3631 | 274.1852 | 575.3581 | 288.1827 | S | 234.1448 | 117.5761 | 217.1183 | 109.0628 | 2 |
| 7 | | | | | K | 147.1128 | 74.0600 | 130.0863 | 65.5468 | 1 |

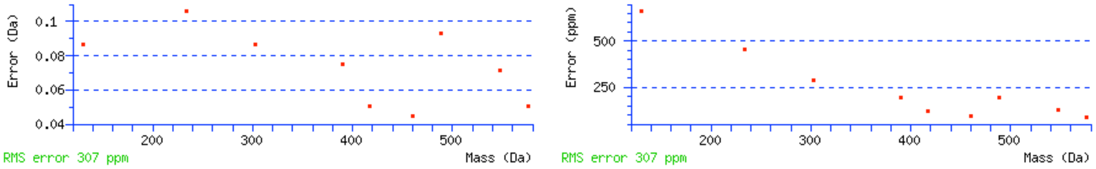

8. Bdh2 from yeast fraction 12.

### Peptide View

MS/MS Fragmentation of **RALAYFGK**
Found in **sp|P39713|BDH2_YEAST** in **uniprot-yeast288C**, sp|P39713|BDH2_YEAST Probable diacetyl reductase [(R)-acetoin forming] 2 OS=Saccharomyces cerevisiae (strain ATCC 204508 / S288c) OX=559292 GN=BDH2 PE=1 SV=1

Match to Query 3299: 942.550024 from(472.282288,2+) intensity(1785350.1250) rtinseconds(1002.087713) index(7446)
Title: zm-yeast-protein-20151023-fr12.9616.9616.2
Data file D:\Panyue\Proteomic dataset\iprox yeast\zm-yeast-protein-20151023-fr12.raw

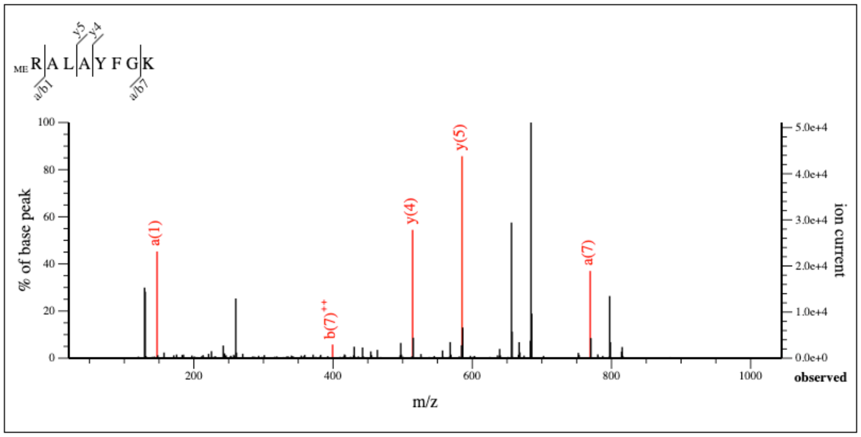


**Monoisotopic mass of neutral peptide Mr(calc):** 942.5559
**Fixed modifications:** Carbamidomethyl (C),Label:13C(1)2H(3) (M) (apply to specified residues or termini only)
**Variable modifications:**
N-term : Methyl:2H(3)13C(1) (Protein N-term)
**Ions Score:** 21 **Expect:** 0.022
**Matches :** 7/84 fragment ions using 9 most intense peaks (help)

| # | a | a++ | a* | a*++ | b | b++ | b* | b*++ | Seq. | y | y++ | y* | y*++ | # |
|---|---|---|---|---|---|---|---|---|---|---|---|---|---|---|
| 1 | **147.1513** | 74.0793 | 130.1248 | 65.5660 | 175.1462 | 88.0767 | 158.1197 | 79.5635 | R | | | | | 8 |
| 2 | 218.1884 | 109.5978 | 201.1619 | 101.0846 | 246.1833 | 123.5953 | 229.1568 | 115.0820 | A | ***769.4243*** | 385.2158 | 752.3978 | 376.7025 | 7 |
| 3 | 331.2725 | 166.1399 | 314.2459 | 157.6266 | 359.2674 | 180.1373 | 342.2409 | 171.6241 | L | 698.3872 | 349.6972 | 681.3606 | 341.1840 | 6 |
| 4 | 402.3096 | 201.6584 | 385.2831 | 193.1452 | 430.3045 | 215.6559 | 413.2780 | 207.1426 | A | ***585.3031*** | 293.1552 | 568.2766 | 284.6419 | 5 |
| 5 | 565.3729 | 283.1901 | 548.3464 | 274.6768 | 593.3678 | 297.1876 | 576.3413 | 288.6743 | Y | ***514.2660*** | 257.6366 | 497.2395 | 249.1234 | 4 |
| 6 | 712.4413 | 356.7243 | 695.4148 | 348.2110 | 740.4363 | 370.7218 | 723.4097 | 362.2085 | F | 351.2027 | 176.1050 | 334.1761 | 167.5917 | 3 |
| 7 | **769.4628** | 385.2350 | 752.4363 | 376.7218 | 797.4577 | 399.2325 | 780.4312 | 390.7192 | G | 204.1343 | 102.5708 | 187.1077 | 94.0575 | 2 |
| 8 | | | | | | | | | K | ***147.1128*** | 74.0600 | 130.0863 | 65.5468 | 1 |

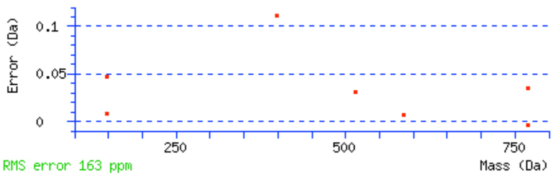


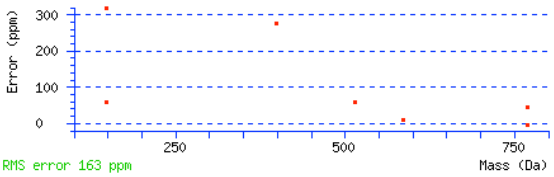


9. Arb1 from yeast fraction 13

### Peptide View

MS/MS Fragmentation of **PPVSASK**
Found in **sp|P40024|ARB1_YEAST** in **uniprot-yeast288C**, sp|P40024|ARB1_YEAST ABC transporter ATP-binding protein ARB1 OS=Saccharomyces cerevisiae (strain ATCC 204508 / S288c) OX=559292 GN=ARB1 PE=1 SV=1

Match to Query 81: 702.417822 from(352.216187,2+) intensity(2060553.1250) rtinseconds(292.7792587) index(647)
Title: zm-yeast-protein-20151023-fr13.1257.1257.2
Data file D:\Panyue\Proteomic dataset\iprox yeast\zm-yeast-protein-20151023-fr13.raw

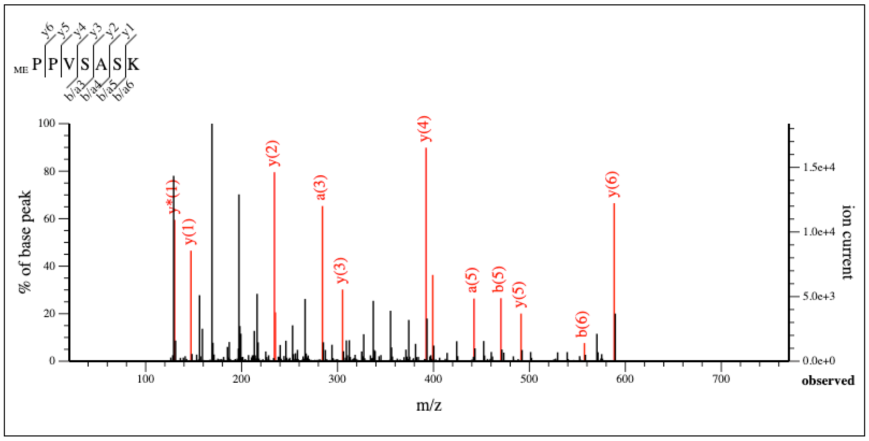


**Monoisotopic mass of neutral peptide Mr(calc):** 702.4185
**Fixed modifications:** Carbamidomethyl (C),Label:13C(1)2H(3) (M) (apply to specified residues or termini only)
**Variable modifications:**
N-term : Methyl:2H(3)13C(1) (Protein N-term)
**Ions Score:** 36 **Expect:** 0.00026
**Matches :** 13/48 fragment ions using 29 most intense peaks (help)

| # | a | a++ | b | b++ | Seq. | y | y++ | y* | y*++ | # |
|---|---|---|---|---|---|---|---|---|---|---|
| 1 | 88.1030 | 44.5551 | 116.0979 | 58.5526 | P | | | | | 7 |
| 2 | 185.1557 | 93.0815 | 213.1506 | 107.0790 | P | ***588.3352*** | 294.6712 | 571.3086 | 286.1579 | 6 |
| 3 | **284.2241** | 142.6157 | 312.2191 | 156.6132 | V | ***491.2824*** | 246.1448 | 474.2558 | 237.6316 | 5 |
| 4 | 371.2562 | 186.1317 | **399.2511** | 200.1292 | S | ***392.2140*** | 196.6106 | 375.1874 | 188.0974 | 4 |
| 5 | **442.2933** | 221.6503 | **470.2882** | 235.6477 | A | ***305.1819*** | 153.0946 | 288.1554 | 144.5813 | 3 |
| 6 | 529.3253 | 265.1663 | **557.3202** | 279.1637 | S | ***234.1448*** | 117.5761 | 217.1183 | 109.0628 | 2 |
| 7 | | | | | K | ***147.1128*** | 74.0600 | 130.0863 | 65.5468 | 1 |

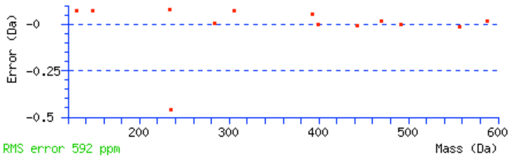


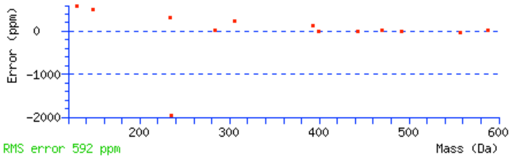


10. Arb1 from yeast fraction 14

### Peptide View

MS/MS Fragmentation of **PPVSASK**
Found in **sp|P40024|ARB1_YEAST** in **uniprot-yeast288C**, sp|P40024|ARB1_YEAST ABC transporter ATP-binding protein ARB1 OS=Saccharomyces cerevisiae (strain ATCC 204508 / S288c) OX=559292 GN=ARB1 PE=1 SV=1

Match to Query 82: 702.419468 from(352.217010,2+) intensity(108532.0391) rtinseconds(313.0045996) index(632)
Title: zm-yeast-protein-20151023-fr14.1294.1294.2
Data file D:\Panyue\Proteomic dataset\iprox yeast\zm-yeast-protein-20151023-fr14.raw

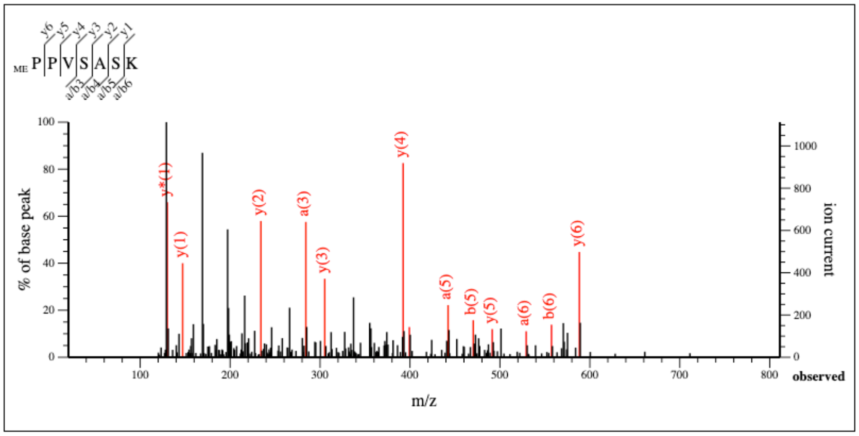

```
Monoisotopic mass of neutral peptide Mr(calc): 702.4185
Fixed modifications: Carbamidomethyl (C),Label:13C(1)2H(3) (M) (apply to specified residues or termini only)
Variable modifications:
  N-term : Methyl:2H(3)13C(1) (Protein N-term)
Ions Score: 39  Expect: 0.00013
Matches : 13/48 fragment ions using 27 most intense peaks   (help)
```

| # | a | a++ | b | b++ | Seq. | y | y++ | y* | y*++ | # |
|---|---|---|---|---|---|---|---|---|---|---|
| 1 | 88.1030 | 44.5551 | 116.0979 | 58.5526 | P | | | | | 7 |
| 2 | 185.1557 | 93.0815 | 213.1506 | 107.0790 | P | 588.3352 | 294.6712 | 571.3086 | 286.1579 | 6 |
| 3 | 284.2241 | 142.6157 | 312.2191 | 156.6132 | V | 491.2824 | 246.1448 | 474.2558 | 237.6316 | 5 |
| 4 | 371.2562 | 186.1317 | 399.2511 | 200.1292 | S | 392.2140 | 196.6106 | 375.1874 | 188.0974 | 4 |
| 5 | 442.2933 | 221.6503 | 470.2882 | 235.6477 | A | 305.1819 | 153.0946 | 288.1554 | 144.5813 | 3 |
| 6 | 529.3253 | 265.1663 | 557.3202 | 279.1637 | S | 234.1448 | 117.5761 | 217.1183 | 109.0628 | 2 |
| 7 | | | | | K | 147.1128 | 74.0600 | 130.0863 | 65.5468 | 1 |

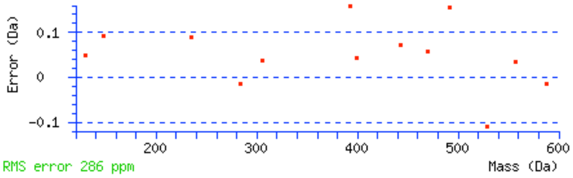

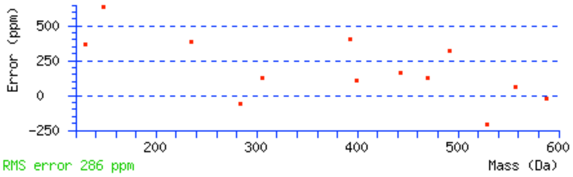

MS/MS spectra for MISL hits (heavy labeled, iMet excluded):

1. Rps25B (1 peptide in 200mM fraction 1)

MS/MS Fragmentation of **PPKQQLSKAAK**
Found in **P0C0T4** in **Uniprot_yeastS288C**, 40S ribosomal protein S25-B OS=Saccharomyces cerevisiae (strain ATCC 204508 / S288c) OX=559292 GN=RPS25B PE=1 SV=1

Match to Query 7087: 1211.711766 from(606.863159,2+) intensity(928242.0625) rtinseconds(3381.1665) index(9066)
Title: 20100530_methyl200mM_1.10088.10088.2
Data file D:\Panyue\Proteomic dataset\20100530_methyl200mM_1.RAW

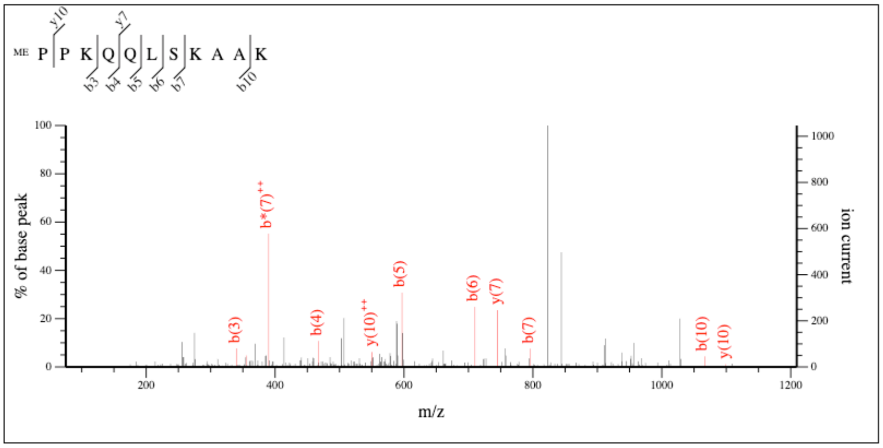


**Monoisotopic mass of neutral peptide Mr(calc):** 1211.7429
**Fixed modifications:** Carbamidomethyl (C) (apply to specified residues or termini only)
**Variable modifications:**
**N-term :** Methyl:2H(3) (Protein N-term)
**Ions Score:** 21 **Expect:** 0.1
**Matches :** 12/96 fragment ions using 29 most intense peaks (help)

| # | b | $b^{++}$ | b* | $b^{*++}$ | $b^0$ | $b^{0++}$ | Seq. | y | $y^{++}$ | y* | $y^{*++}$ | $y^0$ | $y^{0++}$ | # |
|---|---|---|---|---|---|---|---|---|---|---|---|---|---|---|
| 1 | 115.0945 | 58.0509 | | | | | P | | | | | | | 11 |
| 2 | 212.1473 | 106.5773 | | | | | P | 1098.6630 | 549.8351 | 1081.6364 | 541.3218 | 1080.6524 | 540.8298 | 10 |
| 3 | *340.2422* | 170.6248 | 323.2157 | 162.1115 | | | K | 1001.6102 | 501.3087 | 984.5837 | 492.7955 | 983.5996 | 492.3035 | 9 |
| 4 | *468.3008* | 234.6541 | 451.2743 | 226.1408 | | | Q | 873.5152 | 437.2613 | 856.4887 | 428.7480 | 855.5047 | 428.2560 | 8 |
| 5 | *596.3594* | 298.6833 | 579.3329 | 290.1701 | | | Q | 745.4567 | 373.2320 | 728.4301 | 364.7187 | 727.4461 | 364.2267 | 7 |
| 6 | *709.4435* | 355.2254 | 692.4169 | 346.7121 | | | L | 617.3981 | 309.2027 | 600.3715 | 300.6894 | 599.3875 | 300.1974 | 6 |
| 7 | *796.4755* | 398.7414 | 779.4489 | 390.2281 | 778.4649 | 389.7361 | S | 504.3140 | 252.6606 | 487.2875 | 244.1474 | 486.3035 | 243.6554 | 5 |
| 8 | 924.5705 | 462.7889 | 907.5439 | 454.2756 | 906.5599 | 453.7836 | K | 417.2820 | 209.1446 | 400.2554 | 200.6314 | | | 4 |
| 9 | 995.6076 | 498.3074 | 978.5810 | 489.7941 | 977.5970 | 489.3021 | A | 289.1870 | 145.0972 | 272.1605 | 136.5839 | | | 3 |
| 10 | *1066.6447* | 533.8260 | 1049.6181 | 525.3127 | 1048.6341 | 524.8207 | A | 218.1499 | 109.5786 | 201.1234 | 101.0653 | | | 2 |
| 11 | | | | | | | K | 147.1128 | 74.0600 | 130.0863 | 65.5468 | | | 1 |

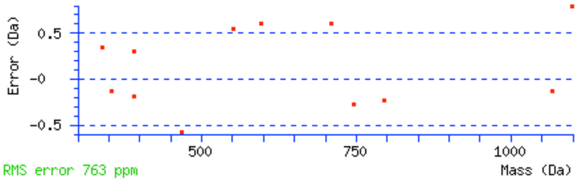


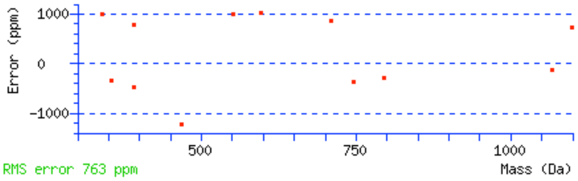


2. Pnc1 (1 peptide in 200mM fraction 1)

MS/MS Fragmentation of **KTLIVVDMQNDFISPLGSLTVPK**
Found in **P53184** in **Uniprot_yeastS288C**, Nicotinamidase OS=Saccharomyces cerevisiae (strain ATCC 204508 / S288c) OX=559292 GN=PNC1 PE=1 SV=1

Match to Query 7565: 2547.410896 from(637.860000,4+) intensity(421356.2188) rtinseconds(4093.7432) index(10948)
Title: 20100530_methyl200mM_1.12181.12181.
Data file D:\Panyue\Proteomic dataset\20100530_methyl200mM_1.RAW

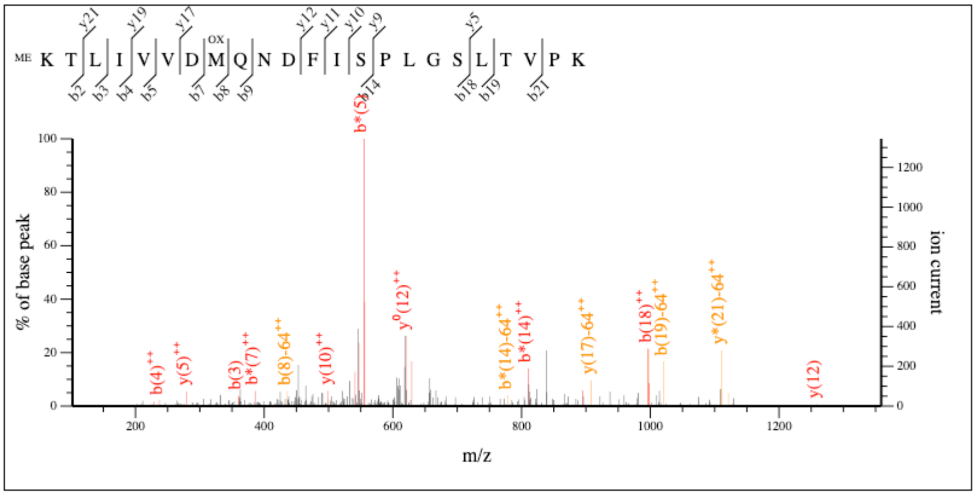


**Monoisotopic mass of neutral peptide Mr(calc):** 2547.4062
**Fixed modifications:** Carbamidomethyl (C) (apply to specified residues or termini only)
**Variable modifications:**
**N-term :** Methyl:2H(3) (Protein N-term)
**M8** : Oxidation (M), with neutral losses 63.9983(shown in table), 0.0000
**Ions Score:** 21 **Expect:** 0.1
**Matches :** 36/388 fragment ions using 51 most intense peaks (help)

| # | b | $b^{++}$ | $b^{*}$ | $b^{*++}$ | $b^{0}$ | $b^{0++}$ | Seq. | y | $y^{++}$ | $y^{*}$ | $y^{*++}$ | $y^{0}$ | $y^{0++}$ | # |
|---|---|---|---|---|---|---|---|---|---|---|---|---|---|---|
| 1 | 146.1367 | 73.5720 | 129.1102 | 65.0587 | | | K | | | | | | | 23 |
| 2 | 247.1844 | 124.0958 | 230.1578 | 115.5826 | 229.1738 | 115.0906 | T | 2339.2857 | 1170.1465 | 2322.2591 | 1161.6332 | 2321.2751 | 1161.1412 | 22 |
| 3 | **360.2685** | 180.6379 | 343.2419 | 172.1246 | 342.2579 | 171.6326 | L | 2238.2380 | 1119.6226 | 2221.2115 | 1111.1094 | 2220.2274 | 1110.6174 | 21 |
| 4 | 473.3525 | ***237.1799*** | 456.3260 | 228.6666 | 455.3420 | 228.1746 | I | 2125.1539 | 1063.0806 | 2108.1274 | 1054.5673 | 2107.1434 | 1054.0753 | 20 |
| 5 | 572.4209 | 286.7141 | 555.3944 | 278.2008 | 554.4104 | 277.7088 | V | 2012.0699 | 1006.5386 | 1995.0433 | 998.0253 | 1994.0593 | 997.5333 | 19 |
| 6 | 671.4894 | 336.2483 | 654.4628 | 327.7350 | 653.4788 | 327.2430 | V | 1913.0015 | 957.0044 | 1895.9749 | 948.4911 | 1894.9909 | 947.9991 | 18 |
| 7 | 786.5163 | 393.7618 | 769.4897 | 385.2485 | 768.5057 | 384.7565 | D | 1813.9331 | ***907.4702*** | 1796.9065 | 898.9569 | 1795.9225 | 898.4649 | 17 |
| 8 | 869.5534 | ***435.2803*** | 852.5269 | 426.7671 | 851.5428 | 426.2751 | M | 1698.9061 | 849.9567 | 1681.8796 | 841.4434 | 1680.8955 | 840.9514 | 16 |
| 9 | **997.6120** | ***499.3096*** | 980.5854 | 490.7964 | 979.6014 | 490.3044 | Q | 1615.8690 | 808.4381 | 1598.8425 | 799.9249 | 1597.8584 | 799.4329 | 15 |
| 10 | **1111.6549** | ***556.3311*** | 1094.6284 | 547.8178 | 1093.6444 | 547.3258 | N | 1487.8104 | 744.4088 | 1470.7839 | 735.8956 | 1469.7999 | 735.4036 | 14 |
| 11 | 1226.6819 | 613.8446 | 1209.6553 | 605.3313 | 1208.6713 | 604.8393 | D | 1373.7675 | 687.3874 | 1356.7409 | 678.8741 | 1355.7569 | 678.3821 | 13 |
| 12 | 1373.7503 | 687.3788 | 1356.7237 | 678.8655 | 1355.7397 | 678.3735 | F | **1258.7406** | ***629.8739*** | 1241.7140 | 621.3606 | 1240.7300 | 620.8686 | 12 |
| 13 | 1486.8343 | 743.9208 | 1469.8078 | 735.4075 | 1468.8238 | 734.9155 | I | **1111.6721** | ***556.3397*** | 1094.6456 | 547.8264 | 1093.6616 | 547.3344 | 11 |
| 14 | 1573.8664 | 787.4368 | 1556.8398 | 778.9235 | 1555.8558 | 778.4315 | S | **998.5881** | ***499.7977*** | 981.5615 | 491.2844 | 980.5775 | 490.7924 | 10 |
| 15 | 1670.9191 | 835.9632 | 1653.8926 | 827.4499 | 1652.9086 | 826.9579 | P | 911.5560 | 456.2817 | 894.5295 | 447.7684 | 893.5455 | 447.2764 | 9 |
| 16 | 1784.0032 | 892.5052 | 1766.9766 | 883.9920 | 1765.9926 | 883.5000 | L | 814.5033 | 407.7553 | 797.4767 | 399.2420 | 796.4927 | 398.7500 | 8 |
| 17 | 1841.0247 | 921.0160 | 1823.9981 | 912.5027 | 1823.0141 | 912.0107 | G | 701.4192 | 351.2132 | 684.3927 | 342.7000 | 683.4087 | 342.2080 | 7 |
| 18 | 1928.0567 | 964.5320 | 1911.0301 | 956.0187 | 1910.0461 | 955.5267 | S | 644.3978 | 322.7025 | 627.3712 | 314.1892 | 626.3872 | 313.6972 | 6 |
| 19 | 2041.1408 | ***1021.0740*** | 2024.1142 | 1012.5607 | 2023.1302 | 1012.0687 | L | 557.3657 | ***279.1865*** | 540.3392 | 270.6732 | 539.3552 | 270.1812 | 5 |
| 20 | 2142.1884 | 1071.5979 | 2125.1619 | 1063.0846 | 2124.1779 | 1062.5926 | T | 444.2817 | 222.6445 | 427.2551 | 214.1312 | 426.2711 | 213.6392 | 4 |
| 21 | 2241.2568 | ***1121.1321*** | 2224.2303 | 1112.6188 | 2223.2463 | 1112.1268 | V | 343.2340 | 172.1206 | 326.2074 | 163.6074 | | | 3 |
| 22 | 2338.3096 | 1169.6584 | 2321.2831 | 1161.1452 | 2320.2990 | 1160.6532 | P | 244.1656 | 122.5864 | 227.1390 | 114.0731 | | | 2 |
| 23 | | | | | | | K | 147.1128 | 74.0600 | 130.0863 | 65.5468 | | | 1 |

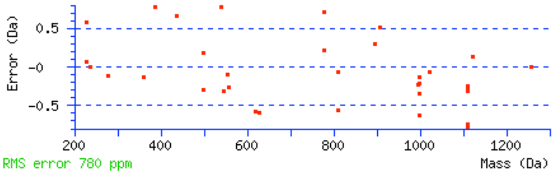


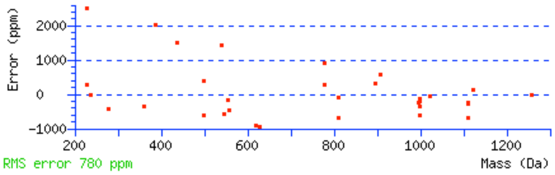


3. Rsc58 (2 peptides in 200mM fraction 1)

MS/MS Fragmentation of **TESVGGNK**
Found in **Q07979** in **Uniprot_yeastS288C**, Chromatin structure-remodeling complex protein RSC58 OS=Saccharomyces cerevisiae (strain ATCC 204508 / S288c) OX=559292 GN=RSC58 PE=1 SV=1

Match to Query 2744: 807.413195 from(808.420471,1+) intensity(253010.4844) rtinseconds(1335.2454) index(3406)
Title: 20100530_methyl200mM_1.3790.3790.1
Data file D:\Panyue\Proteomic dataset\20100530_methyl200mM_1.RAW

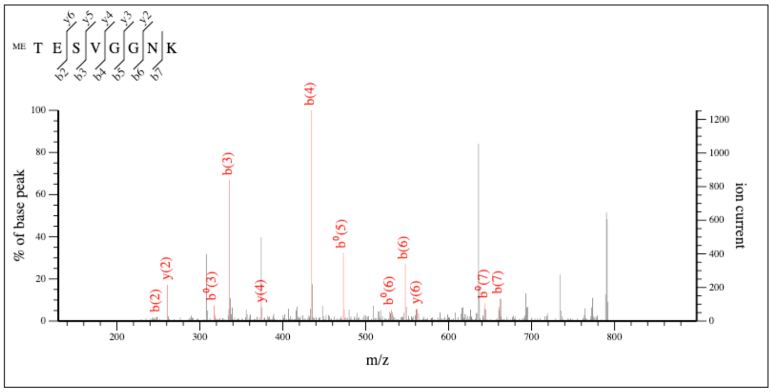


**Monoisotopic mass of neutral peptide Mr(calc):** 807.4166
**Fixed modifications:** Carbamidomethyl (C) (apply to specified residues or termini only)
**Variable modifications:**
**N-term :** Methyl:2H(3) (Protein N-term)
**Ions Score:** 25 **Expect:** 0.044
**Matches :** 14/31 fragment ions using 48 most intense peaks (help)

| # | b | b* | $b^0$ | Seq. | y | y* | $y^0$ | # |
|---|---|---|---|---|---|---|---|---|
| 1 | 119.0894 | | 101.0789 | T | | | | 8 |
| 2 | *248.1320* | | 230.1215 | E | 690.3417 | 673.3151 | 672.3311 | 7 |
| 3 | *335.1641* | | 317.1535 | S | *561.2991* | 544.2726 | 543.2885 | 6 |
| 4 | *434.2325* | | 416.2219 | V | *474.2671* | 457.2405 | | 5 |
| 5 | 491.2539 | | 473.2434 | G | *375.1987* | 358.1721 | | 4 |
| 6 | *548.2754* | | 530.2648 | G | *318.1772* | 301.1506 | | 3 |
| 7 | *662.3183* | 645.2918 | 644.3078 | N | *261.1557* | 244.1292 | | 2 |
| 8 | | | | K | 147.1128 | 130.0863 | | 1 |

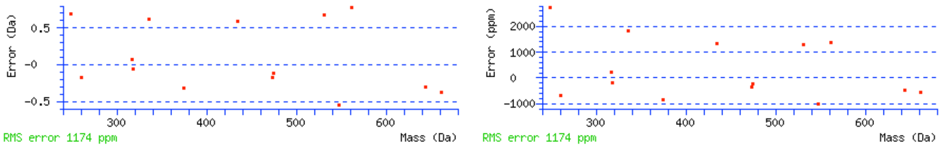


MS/MS Fragmentation of **TESVGGNK**
Found in **Q07979** in **Uniprot_yeastS288C**, Chromatin structure-remodeling complex protein RSC58 OS=Saccharomyces cerevisiae (strain ATCC 204508 / S288c) OX=559292 GN=RSC58 PE=1 SV=1

Match to Query 2745: 807.413745 from(808.421021,1+) intensity(221410.3125) rtinseconds(1331.9284) index(3397)
Title: 20100530_methyl200mM_1.3780.3780.1
Data file D:\Panyue\Proteomic dataset\20100530_methyl200mM_1.RAW

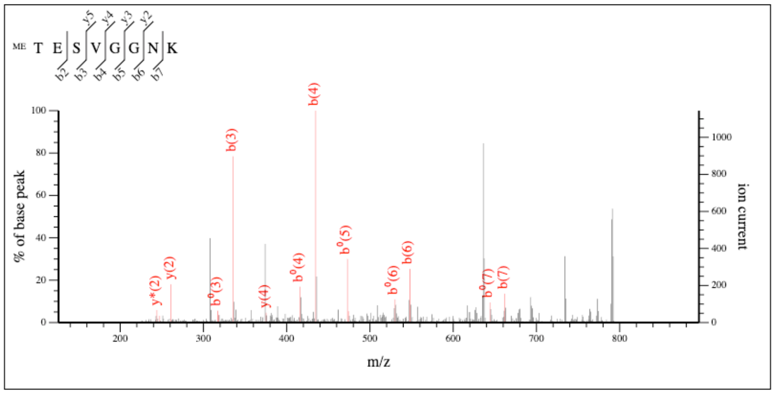

**Monoisotopic mass of neutral peptide Mr(calc):** 807.4166
**Fixed modifications:** Carbamidomethyl (C) (apply to specified residues or termini only)
**Variable modifications:**
**N-term :** Methyl:2H(3) (Protein N-term)
**Ions Score:** 21 **Expect:** 0.13
**Matches :** 15/31 fragment ions using 54 most intense peaks (help)

| # | b | b* | $b^0$ | Seq. | y | y* | $y^0$ | # |
|---|---|---|---|---|---|---|---|---|
| 1 | 119.0894 | | 101.0789 | T | | | | 8 |
| 2 | ***248.1320*** | | 230.1215 | E | 690.3417 | 673.3151 | 672.3311 | 7 |
| 3 | ***335.1641*** | | 317.1535 | S | 561.2991 | 544.2726 | 543.2885 | 6 |
| 4 | ***434.2325*** | | 416.2219 | V | ***474.2671*** | 457.2405 | | 5 |
| 5 | 491.2539 | | 473.2434 | G | ***375.1987*** | 358.1721 | | 4 |
| 6 | ***548.2754*** | | 530.2648 | G | ***318.1772*** | 301.1506 | | 3 |
| 7 | ***662.3183*** | 645.2918 | 644.3078 | N | ***261.1557*** | 244.1292 | | 2 |
| 8 | | | | K | 147.1128 | 130.0863 | | 1 |

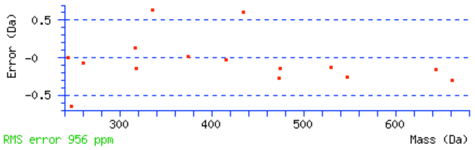


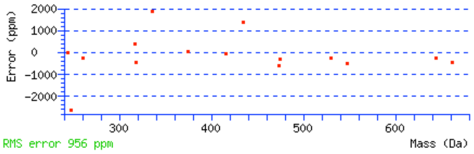


4. Eft1 (1 peptide in 200mM fraction 1)

MS/MS Fragmentation of **VAFTVDQMRSLMDK**
Found in **P32324** in **Uniprot_yeastS288C**, Elongation factor 2 OS=Saccharomyces cerevisiae (strain ATCC 204508 / S288c) OX=559292 GN=EFT1 PE=1 SV=1

Match to Query 10310: 1706.869848 from(854.442200,2+) intensity(2487614.2500) rtinseconds(2705.1608) index(7228)
Title: 20100530_methyl200mM_1.8044.8044.2
Data file D:\Panyue\Proteomic dataset\20100530_methyl200mM_1.RAW

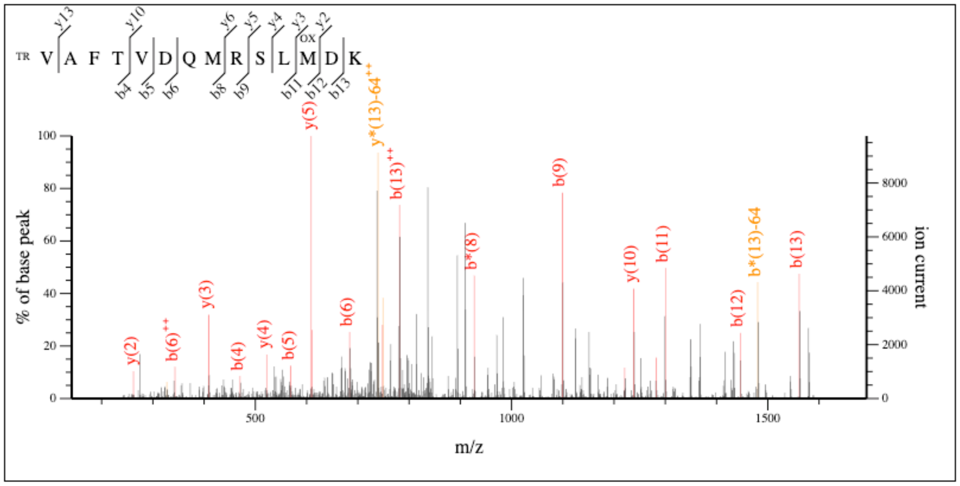

**Monoisotopic mass of neutral peptide Mr(calc):** 1706.9046
**Fixed modifications:** Carbamidomethyl (C) (apply to specified residues or termini only)
**Variable modifications:**
**N-term :** Trimethyl:2H(9) (Protein N-term)
**M12** : Oxidation (M), with neutral losses 0.0000(shown in table), 63.9983
**Ions Score:** 21 **Expect:** 0.13
**Matches :** 30/214 fragment ions using 56 most intense peaks (help)

| # | b | b++ | b* | b*++ | b0 | b0++ | Seq. | y | y++ | y* | y*++ | y0 | y0++ | # |
|---|---|---|---|---|---|---|---|---|---|---|---|---|---|---|
| 1 | 151.1791 | 76.0932 | | | | | V | | | | | | | 14 |
| 2 | 222.2162 | 111.6118 | | | | | A | 1557.7400 | 779.3736 | 1540.7134 | 770.8604 | 1539.7294 | 770.3683 | 13 |
| 3 | 369.2847 | 185.1460 | | | | | F | 1486.7029 | 743.8551 | 1469.6763 | 735.3418 | 1468.6923 | 734.8498 | 12 |
| 4 | 470.3323 | 235.6698 | | | 452.3218 | 226.6645 | T | 1339.6344 | 670.3209 | 1322.6079 | 661.8076 | 1321.6239 | 661.3156 | 11 |
| 5 | 569.4008 | 285.2040 | | | 551.3902 | 276.1987 | V | 1238.5868 | 619.7970 | 1221.5602 | 611.2837 | 1220.5762 | 610.7917 | 10 |
| 6 | 684.4277 | 342.7175 | | | 666.4171 | 333.7122 | D | 1139.5184 | 570.2628 | 1122.4918 | 561.7495 | 1121.5078 | 561.2575 | 9 |
| 7 | 812.4863 | 406.7468 | 795.4597 | 398.2335 | 794.4757 | 397.7415 | Q | 1024.4914 | 512.7493 | 1007.4649 | 504.2361 | 1006.4808 | 503.7441 | 8 |
| 8 | 943.5268 | 472.2670 | 926.5002 | 463.7537 | 925.5162 | 463.2617 | M | 896.4328 | 448.7201 | 879.4063 | 440.2068 | 878.4223 | 439.7148 | 7 |
| 9 | 1099.6279 | 550.3176 | 1082.6013 | 541.8043 | 1081.6173 | 541.3123 | R | 765.3924 | 383.1998 | 748.3658 | 374.6865 | 747.3818 | 374.1945 | 6 |
| 10 | 1186.6599 | 593.8336 | 1169.6333 | 585.3203 | 1168.6493 | 584.8283 | S | 609.2912 | 305.1493 | 592.2647 | 296.6360 | 591.2807 | 296.1440 | 5 |
| 11 | 1299.7440 | 650.3756 | 1282.7174 | 641.8623 | 1281.7334 | 641.3703 | L | 522.2592 | 261.6332 | 505.2327 | 253.1200 | 504.2486 | 252.6280 | 4 |
| 12 | 1446.7794 | 723.8933 | 1429.7528 | 715.3800 | 1428.7688 | 714.8880 | M | 409.1751 | 205.0912 | 392.1486 | 196.5779 | 391.1646 | 196.0859 | 3 |
| 13 | 1561.8063 | 781.4068 | 1544.7798 | 772.8935 | 1543.7957 | 772.4015 | D | 262.1397 | 131.5735 | 245.1132 | 123.0602 | 244.1292 | 122.5682 | 2 |
| 14 | | | | | | | K | 147.1128 | 74.0600 | 130.0863 | 65.5468 | | | 1 |

### 5. Rsc58 (1 peptide in 200mM fraction 2)

MS/MS Fragmentation of TESVGGNK
Found in Q07979 in Uniprot_yeastS288C, Chromatin structure-remodeling complex protein RSC58 OS=Saccharomyces cerevisiae (strain ATCC 204508 / S288c) OX=559292 GN=RSC58 PE=1 SV=1

Match to Query 2922: 807.412951 from(808.420227,1+) intensity(266006.5938) rtinseconds(1363.7524) index(3471)
Title: 20100530_methyl200mM_2.3867.3867.1
Data file D:\Panyue\Proteomic dataset\20100530_methyl200mM_2.RAW

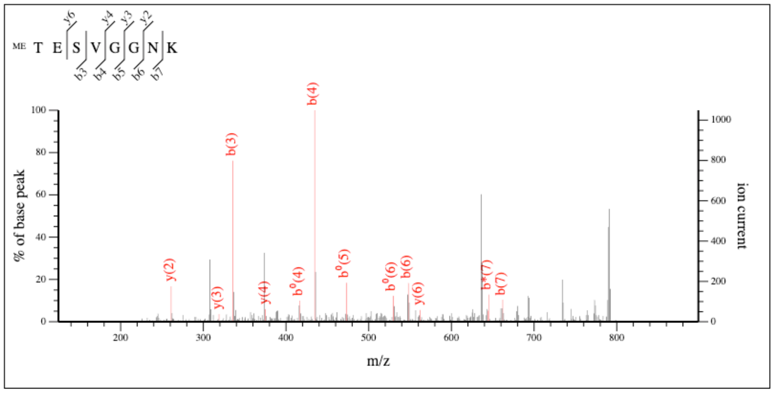


**Monoisotopic mass of neutral peptide Mr(calc):** 807.4166
**Fixed modifications:** Carbamidomethyl (C) (apply to specified residues or termini only)
**Variable modifications:**
**N-term :** Methyl:2H(3) (Protein N-term)
**Ions Score:** 24 **Expect:** 0.052
**Matches :** 13/31 fragment ions using 36 most intense peaks (help)

| # | b | b* | b0 | Seq. | y | y* | y0 | # |
|---|---|---|---|---|---|---|---|---|
| 1 | 119.0894 | | 101.0789 | T | | | | 8 |
| 2 | 248.1320 | | 230.1215 | E | 690.3417 | 673.3151 | 672.3311 | 7 |
| 3 | 335.1641 | | 317.1535 | S | 561.2991 | 544.2726 | 543.2885 | 6 |
| 4 | 434.2325 | | 416.2219 | V | 474.2671 | 457.2405 | | 5 |
| 5 | 491.2539 | | 473.2434 | G | 375.1987 | 358.1721 | | 4 |
| 6 | 548.2754 | | 530.2648 | G | 318.1772 | 301.1506 | | 3 |
| 7 | 662.3183 | 645.2918 | 644.3078 | N | 261.1557 | 244.1292 | | 2 |
| 8 | | | | K | 147.1128 | 130.0863 | | 1 |

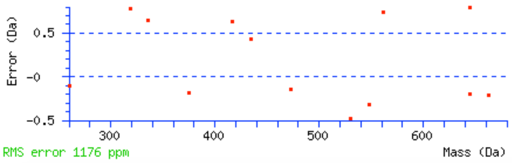


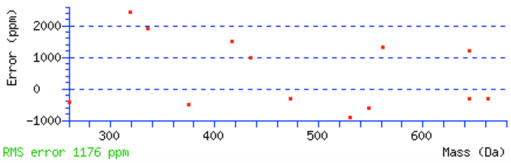

6. Rpn13 (1 peptide in 200mM fraction2)

MS/MS Fragmentation of **SSTVIK**
Found in **O13563-2** in **Uniprot_yeastS288C**, Isoform 2 of 26S proteasome regulatory subunit RPN13 OS=Saccharomyces cerevisiae (strain ATCC 204508 / S288c) OX=559292 GN=RPN13

Match to Query 1977: 684.453784 from(685.461060,1+) intensity(685624.8750) rtinseconds(1458.6093) index(3736)
Title: 20100530_methyl200mM_2.4162.4162.1
Data file D:\Panyue\Proteomic dataset\20100530_methyl200mM_2.RAW

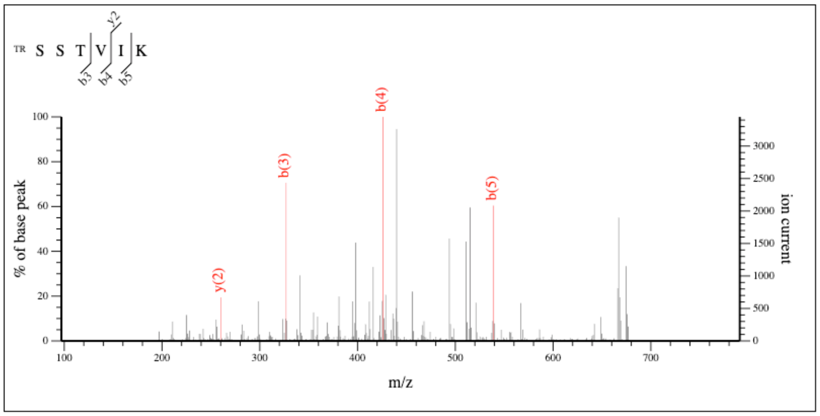


**Monoisotopic mass of neutral peptide Mr(calc):** 684.4732
**Fixed modifications:** Carbamidomethyl (C) (apply to specified residues or termini only)
**Variable modifications:**
**N-term :** Trimethyl:2H(9) (Protein N-term)
**Ions Score:** 21 **Expect:** 0.069
**Matches :** 4/22 fragment ions using 5 most intense peaks (help)

| # | b | $b^0$ | Seq. | y | y* | $y^0$ | # |
|---|---|---|---|---|---|---|---|
| 1 | 139.1427 | 121.1322 | S | | | | 6 |
| 2 | 226.1748 | 208.1642 | S | 547.3450 | 530.3184 | 529.3344 | 5 |
| 3 | *327.2225* | 309.2119 | T | 460.3130 | 443.2864 | 442.3024 | 4 |
| 4 | *426.2909* | 408.2803 | V | 359.2653 | 342.2387 | | 3 |
| 5 | *539.3749* | 521.3644 | I | 260.1969 | 243.1703 | | 2 |
| 6 | | | K | 147.1128 | 130.0863 | | 1 |

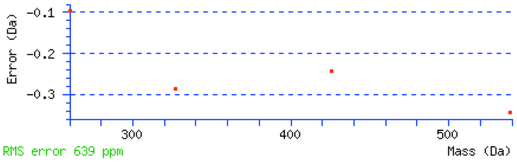


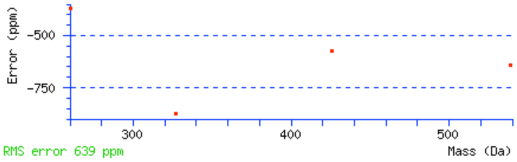


7. L3153 (1 peptide in 200mM fraction 2)

MS/MS Fragmentation of **IEVTMK**
Found in **E9PAA3** in **Uniprot_yeastS288C**, L3153 protein OS=Saccharomyces cerevisiae OX=4932 GN=L3153 PE=4 SV=1

Match to Query 2330: 736.393054 from(737.400330,1+) intensity(617378.8125) rtinseconds(1985.2577) index(5195)
Title: 20100530_methyl200mM_2.5786.5786.1
Data file D:\Panyue\Proteomic dataset\20100530_methyl200mM_2.RAW

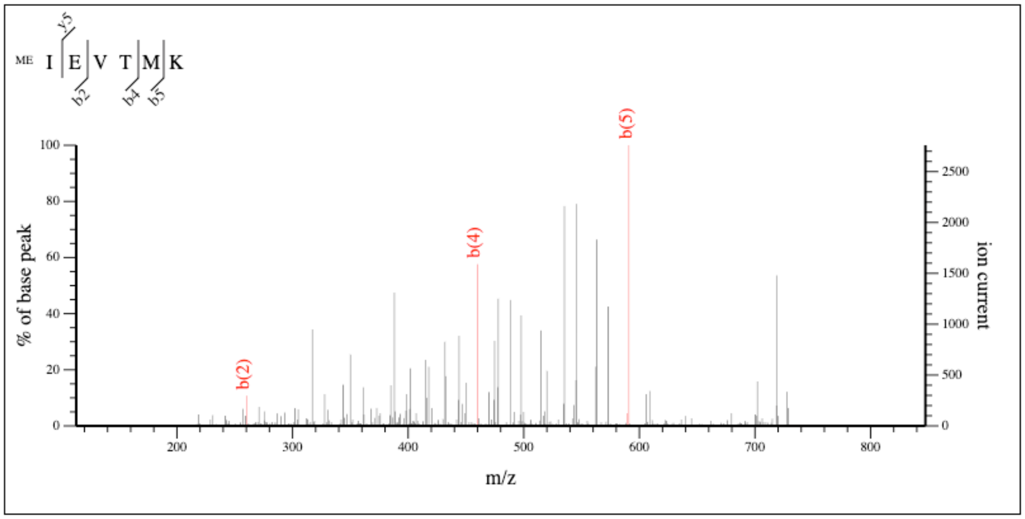


**Monoisotopic mass of neutral peptide Mr(calc):** 736.4232
**Fixed modifications:** Carbamidomethyl (C) (apply to specified residues or termini only)
**Variable modifications:**
**N-term :** Methyl:2H(3) (Protein N-term)
**Ions Score:** 20 **Expect:** 0.13
**Matches :** 5/22 fragment ions using 6 most intense peaks (help)

| # | b | $b^0$ | Seq. | y | y* | $y^0$ | # |
|---|---|---|---|---|---|---|---|
| 1 | 131.1258 | | I | | | | 6 |
| 2 | *260.1684* | 242.1578 | E | 607.3120 | 590.2854 | 589.3014 | 5 |
| 3 | 359.2368 | 341.2263 | V | 478.2694 | 461.2428 | 460.2588 | 4 |
| 4 | *460.2845* | 442.2739 | T | 379.2010 | 362.1744 | 361.1904 | 3 |
| 5 | *591.3250* | 573.3144 | M | 278.1533 | 261.1267 | | 2 |
| 6 | | | K | 147.1128 | 130.0863 | | 1 |

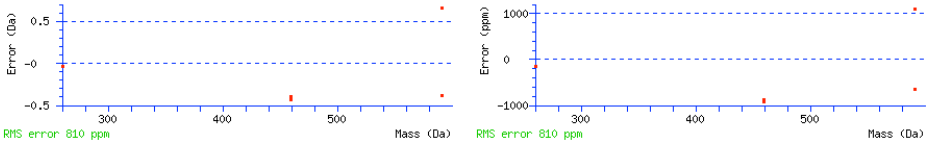


### 8. Asc1 (1 peptide in 200mM fraction3)

MS/MS Fragmentation of **ASNEVLVLR**
Found in **P38011** in **Uniprot_yeastS288C**, Guanine nucleotide-binding protein subunit beta-like protein OS=Saccharomyces cerevisiae (strain ATCC 204508 / S288c) OX=559292 GN=ASC1 PE=1 SV=4

Match to Query 5648: 1050.681323 from(1051.688599,1+) intensity(761627.3750) rtinseconds(3404.0466) index(9100)
Title: 20100530_methyl200mM_3.10126.10126.1
Data file D:\Panyue\Proteomic dataset\20100530_methyl200mM_3.RAW

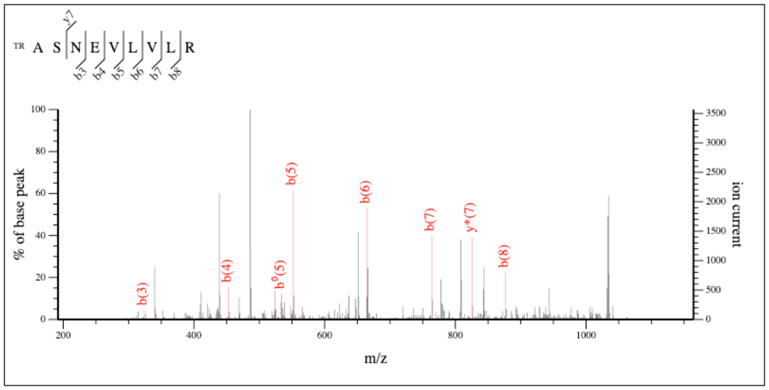

**Monoisotopic mass of neutral peptide Mr(calc):** 1050.6747
**Fixed modifications:** Carbamidomethyl (C) (apply to specified residues or termini only)
**Variable modifications:**
**N-term :** Trimethyl:2H(9) (Protein N-term)
**Ions Score:** 23 **Expect:** 0.011
**Matches :** 8/40 fragment ions using 32 most intense peaks (help)

| # | b | b* | b0 | Seq. | y | y* | y0 | # |
|---|---|---|---|---|---|---|---|---|
| 1 | 123.1478 | | | A | | | | 9 |
| 2 | 210.1799 | | 192.1693 | S | 929.5415 | 912.5149 | 911.5309 | 8 |
| 3 | ***324.2228*** | 307.1962 | 306.2122 | N | 842.5094 | 825.4829 | 824.4989 | 7 |
| 4 | ***453.2654*** | 436.2388 | 435.2548 | E | 728.4665 | 711.4400 | 710.4559 | 6 |
| 5 | ***552.3338*** | 535.3072 | 534.3232 | V | 599.4239 | 582.3974 | | 5 |
| 6 | ***665.4179*** | 648.3913 | 647.4073 | L | 500.3555 | 483.3289 | | 4 |
| 7 | ***764.4863*** | 747.4597 | 746.4757 | V | 387.2714 | 370.2449 | | 3 |
| 8 | ***877.5703*** | 860.5438 | 859.5598 | L | 288.2030 | 271.1765 | | 2 |
| 9 | | | | R | 175.1190 | 158.0924 | | 1 |

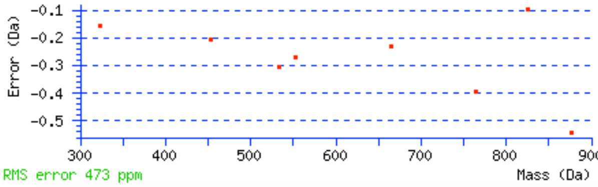


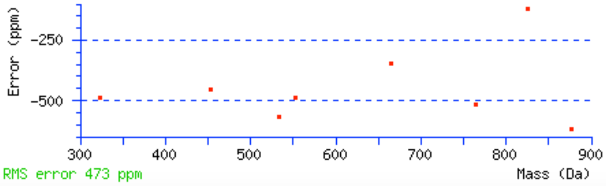


### 9. Pdc6 (1 peptide in 200mM fraction3)

MS/MS Fragmentation of **SEITLGK**
Found in **P26263** in **Uniprot_yeastS288C**, Pyruvate decarboxylase isozyme 3 OS=Saccharomyces cerevisiae (strain ATCC 204508 / S288c) OX=559292 GN=PDC6 PE=1 SV=3

Match to Query 2409: 780.448596 from(781.455872,1+) intensity(231849.4844) rtinseconds(1069.2189) index(2672)
Title: 20100530_methyl200mM_3.2973.2973.1
Data file D:\Panyue\Proteomic dataset\20100530_methyl200mM_3.RAW

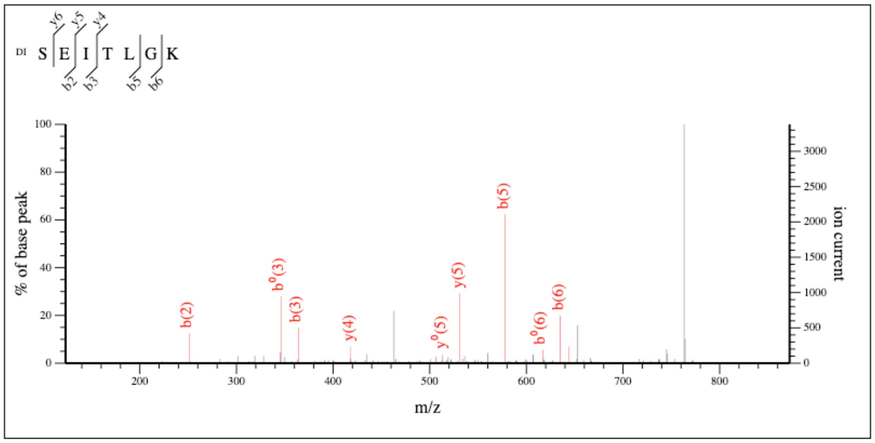

**Monoisotopic mass of neutral peptide Mr(calc):** 780.4864
**Fixed modifications:** Carbamidomethyl (C) (apply to specified residues or termini only)
**Variable modifications:**
**N-term :** Dimethyl:2H(6) (Protein N-term)
**Ions Score:** 29 **Expect:** 0.017
**Matches :** 10/27 fragment ions using 18 most intense peaks (help)

| # | b | $b^0$ | Seq. | y | y* | $y^0$ | # |
|---|---|---|---|---|---|---|---|
| 1 | 122.1083 | 104.0977 | S | | | | 7 |
| 2 | 251.1509 | 233.1403 | E | 660.3927 | 643.3661 | 642.3821 | 6 |
| 3 | 364.2349 | 346.2244 | I | 531.3501 | 514.3235 | 513.3395 | 5 |
| 4 | 465.2826 | 447.2720 | T | 418.2660 | 401.2395 | 400.2554 | 4 |
| 5 | 578.3667 | 560.3561 | L | 317.2183 | 300.1918 | | 3 |
| 6 | 635.3881 | 617.3776 | G | 204.1343 | 187.1077 | | 2 |
| 7 | | | K | 147.1128 | 130.0863 | | 1 |

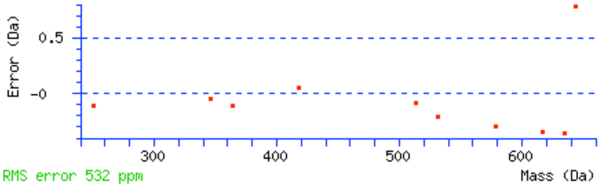


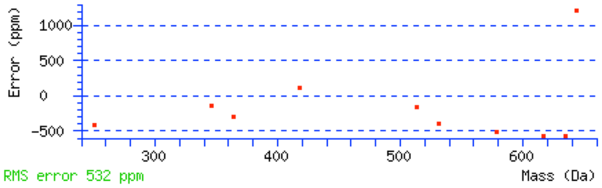


### 10. Pdc5 (1 peptide in 200mM fraction3)

MS/MS Fragmentation of **SEITLGK**
Found in **P16467** in **Uniprot_yeastS288C**, Pyruvate decarboxylase isozyme 2 OS=Saccharomyces cerevisiae (strain ATCC 204508 / S288c) OX=559292 GN=PDC5 PE=1 SV=4

Match to Query 2409: 780.448596 from(781.455872,1+) intensity(231849.4844) rtinseconds(1069.2189) index(2672)
Title: 20100530_methyl200mM_3.2973.2973.1
Data file D:\Panyue\Proteomic dataset\20100530_methyl200mM_3.RAW

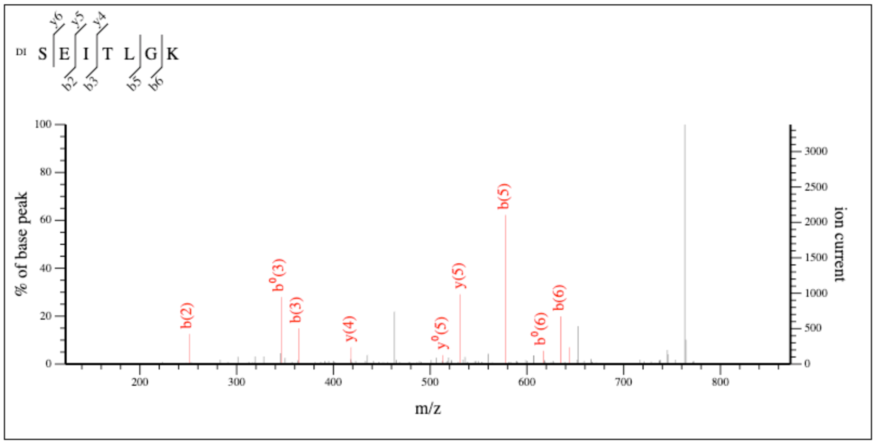

**Monoisotopic mass of neutral peptide Mr(calc):** 780.4864
**Fixed modifications:** Carbamidomethyl (C) (apply to specified residues or termini only)
**Variable modifications:**
**N-term :** Dimethyl:2H(6) (Protein N-term)
**Ions Score:** 29 **Expect:** 0.017
**Matches :** 10/27 fragment ions using 18 most intense peaks (help)

| # | b | $b^0$ | Seq. | y | y* | $y^0$ | # |
|---|---|---|---|---|---|---|---|
| 1 | 122.1083 | 104.0977 | S | | | | 7 |
| 2 | ***251.1509*** | 233.1403 | E | 660.3927 | 643.3661 | 642.3821 | 6 |
| 3 | ***364.2349*** | 346.2244 | I | **531.3501** | 514.3235 | 513.3395 | 5 |
| 4 | 465.2826 | 447.2720 | T | **418.2660** | 401.2395 | 400.2554 | 4 |
| 5 | ***578.3667*** | 560.3561 | L | 317.2183 | 300.1918 | | 3 |
| 6 | ***635.3881*** | 617.3776 | G | 204.1343 | 187.1077 | | 2 |
| 7 | | | K | 147.1128 | 130.0863 | | 1 |

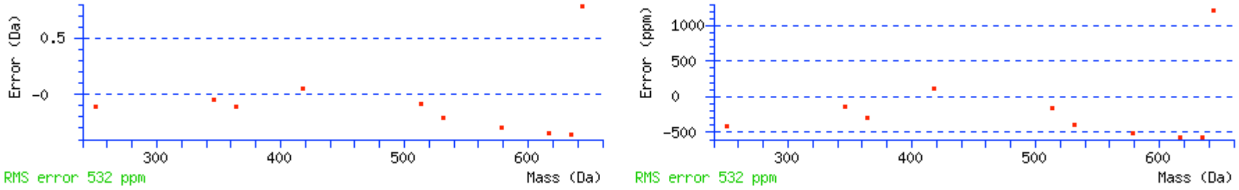


11. Pdc1 (1 peptide in 200mM fraction3)

MS/MS Fragmentation of SEITLGK
Found in P06169 in Uniprot_yeastS288C, Pyruvate decarboxylase isozyme 1 OS=Saccharomyces cerevisiae (strain ATCC 204508 / S288c) OX=559292 GN=PDC1 PE=1 SV=7

Match to Query 2409: 780.448596 from(781.455872,1+) intensity(231849.4844) rtinseconds(1069.2189) index(2672)
Title: 20100530_methyl200mM_3.2973.2973.1
Data file D:\Panyue\Proteomic dataset\20100530_methyl200mM_3.RAW

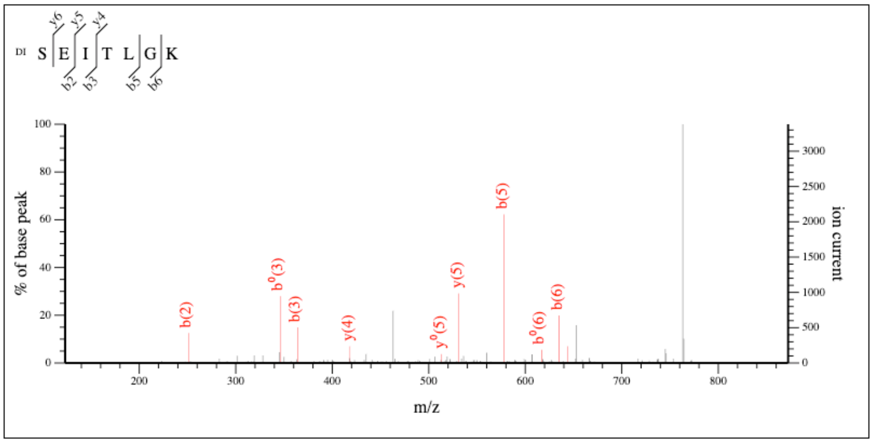

**Monoisotopic mass of neutral peptide Mr(calc):** 780.4864
**Fixed modifications:** Carbamidomethyl (C) (apply to specified residues or termini only)
**Variable modifications:**
**N-term :** Dimethyl:2H(6) (Protein N-term)
**Ions Score:** 29 **Expect:** 0.017
**Matches :** 10/27 fragment ions using 18 most intense peaks (help)

| # | b | b⁰ | Seq. | y | y* | y⁰ | # |
|---|---|---|---|---|---|---|---|
| 1 | 122.1083 | 104.0977 | S | | | | 7 |
| 2 | 251.1509 | 233.1403 | E | 660.3927 | 643.3661 | 642.3821 | 6 |
| 3 | 364.2349 | 346.2244 | I | 531.3501 | 514.3235 | 513.3395 | 5 |
| 4 | 465.2826 | 447.2720 | T | 418.2660 | 401.2395 | 400.2554 | 4 |
| 5 | 578.3667 | 560.3561 | L | 317.2183 | 300.1918 | | 3 |
| 6 | 635.3881 | 617.3776 | G | 204.1343 | 187.1077 | | 2 |
| 7 | | | K | 147.1128 | 130.0863 | | 1 |

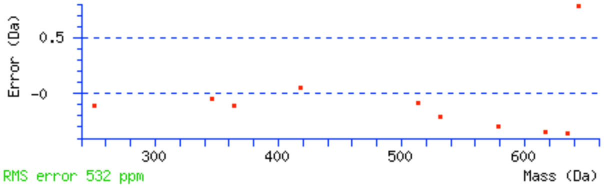


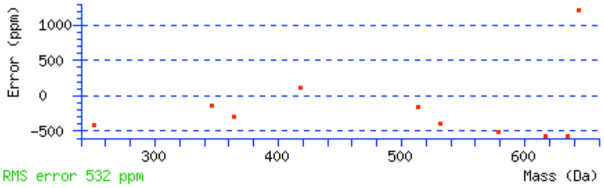


### 12. Ahp1 (1 peptide in 200mM fraction 3)

MS/MS Fragmentation of **SDLVNK**
Found in **P38013** in **Uniprot_yeastS288C**, Peroxiredoxin AHP1 OS=Saccharomyces cerevisiae (strain ATCC 204508 / S288c) OX=559292 GN=AHP1 PE=1 SV=4

Match to Query 2016: 725.443347 from(726.450623,1+) intensity(468986.4375) rtinseconds(1402.1582) index(3570)
Title: 20100530_methyl200mM_3.3974.3974.1
Data file D:\Panyue\Proteomic dataset\20100530_methyl200mM_3.RAW

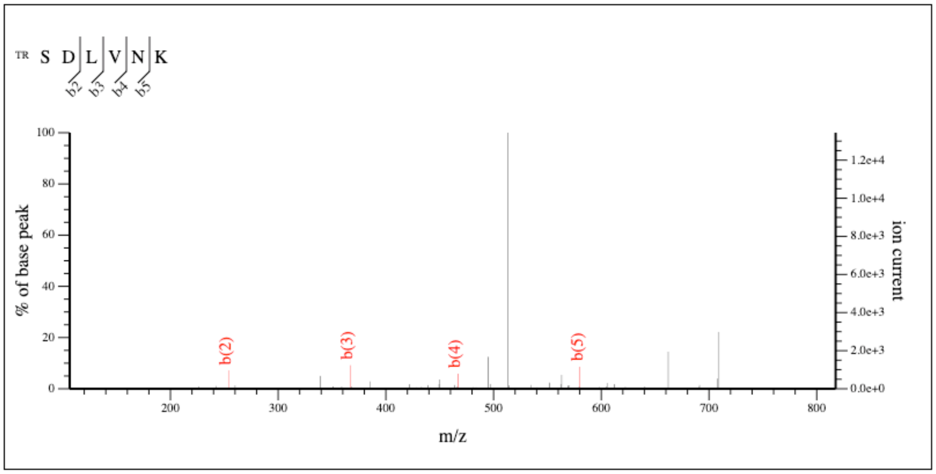

**Monoisotopic mass of neutral peptide Mr(calc):** 725.4633
**Fixed modifications:** Carbamidomethyl (C) (apply to specified residues or termini only)
**Variable modifications:**
**N-term :** Trimethyl:2H(9) (Protein N-term)
**Ions Score:** 21 **Expect:** 0.058
**Matches :** 4/22 fragment ions using 12 most intense peaks (help)

| # | b | b* | b⁰ | Seq. | y | y* | y⁰ | # |
|---|---|---|---|---|---|---|---|---|
| 1 | 139.1427 | | 121.1322 | S | | | | 6 |
| 2 | *254.1697* | | 236.1591 | D | 588.3352 | 571.3086 | 570.3246 | 5 |
| 3 | *367.2538* | | 349.2432 | L | 473.3082 | 456.2817 | | 4 |
| 4 | *466.3222* | | 448.3116 | V | 360.2241 | 343.1976 | | 3 |
| 5 | *580.3651* | 563.3385 | 562.3545 | N | 261.1557 | 244.1292 | | 2 |
| 6 | | | | K | 147.1128 | 130.0863 | | 1 |

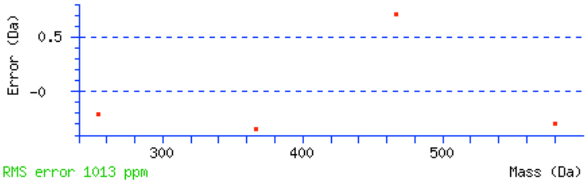


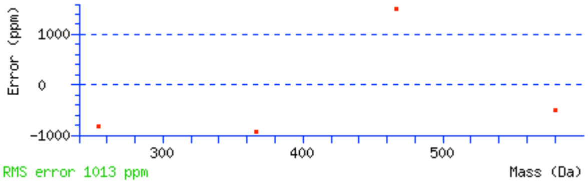


### 13. Rps25A (1 peptide in 200mM fraction 3)

MS/MS Fragmentation of **PPKQQLSKAAK**
Found in **Q3E792** in **Uniprot_yeastS288C**, 40S ribosomal protein S25-A OS=Saccharomyces cerevisiae (strain ATCC 204508 / S288c) OX=559292 GN=RPS25A PE=1 SV=1

Match to Query 7081: 1211.712254 from(606.863403,2+) intensity(3001675.2500) rtinseconds(3277.1282) index(8765)
Title: 20100530_methyl200mM_3.9754.9754.2
Data file D:\Panyue\Proteomic dataset\20100530_methyl200mM_3.RAW

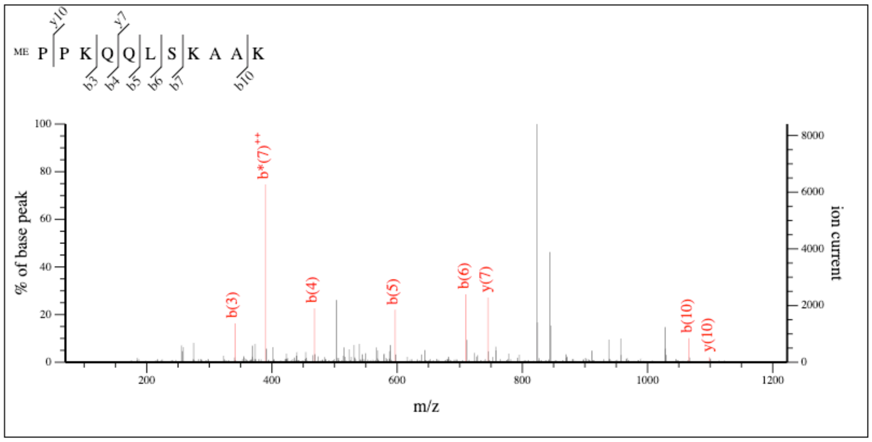

**Monoisotopic mass of neutral peptide Mr(calc):** 1211.7429
**Fixed modifications:** Carbamidomethyl (C) (apply to specified residues or termini only)
**Variable modifications:**
**N-term :** Methyl:2H(3) (Protein N-term)
**Ions Score:** 22 **Expect:** 0.085
**Matches :** 9/96 fragment ions using 20 most intense peaks (help)

| # | b | b++ | b* | b*++ | b0 | b0++ | Seq. | y | y++ | y* | y*++ | y0 | y0++ | # |
|---|---|---|---|---|---|---|---|---|---|---|---|---|---|---|
| 1 | 115.0945 | 58.0509 | | | | | P | | | | | | | 11 |
| 2 | 212.1473 | 106.5773 | | | | | P | **1098.6630** | 549.8351 | 1081.6364 | 541.3218 | 1080.6524 | 540.8298 | 10 |
| 3 | ***340.2422*** | 170.6248 | 323.2157 | 162.1115 | | | K | 1001.6102 | 501.3087 | 984.5837 | 492.7955 | 983.5996 | 492.3035 | 9 |
| 4 | ***468.3008*** | 234.6541 | 451.2743 | 226.1408 | | | Q | 873.5152 | 437.2613 | 856.4887 | 428.7480 | 855.5047 | 428.2560 | 8 |
| 5 | ***596.3594*** | 298.6833 | 579.3329 | 290.1701 | | | Q | **745.4567** | 373.2320 | 728.4301 | 364.7187 | 727.4461 | 364.2267 | 7 |
| 6 | ***709.4435*** | 355.2254 | 692.4169 | 346.7121 | | | L | 617.3981 | 309.2027 | 600.3715 | 300.6894 | 599.3875 | 300.1974 | 6 |
| 7 | 796.4755 | 398.7414 | 779.4489 | 390.2281 | 778.4649 | 389.7361 | S | 504.3140 | 252.6606 | 487.2875 | 244.1474 | 486.3035 | 243.6554 | 5 |
| 8 | 924.5705 | 462.7889 | 907.5439 | 454.2756 | 906.5599 | 453.7836 | K | 417.2820 | 209.1446 | 400.2554 | 200.6314 | | | 4 |
| 9 | 995.6076 | 498.3074 | 978.5810 | 489.7941 | 977.5970 | 489.3021 | A | 289.1870 | 145.0972 | 272.1605 | 136.5839 | | | 3 |
| 10 | ***1066.6447*** | 533.8260 | 1049.6181 | 525.3127 | 1048.6341 | 524.8207 | A | 218.1499 | 109.5786 | 201.1234 | 101.0653 | | | 2 |
| 11 | | | | | | | K | 147.1128 | 74.0600 | 130.0863 | 65.5468 | | | 1 |

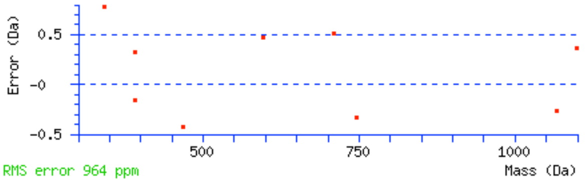


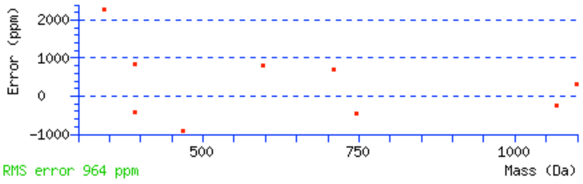


### 14. Pdc5 (2 peptides in 250mM fraction 1)

MS/MS Fragmentation of **SEITLGK**
Found in **P16467** in **Uniprot_yeastS288C**, Pyruvate decarboxylase isozyme 2 OS=Saccharomyces cerevisiae (strain ATCC 204508 / S288c) OX=559292 GN=PDC5 PE=1 SV=4

Match to Query 2434: 780.449267 from(781.456543,1+) intensity(563499.6875) rtinseconds(1029.388) index(2565)
Title: 20100530_methyl250mM_1.2854.2854.1
Data file D:\Panyue\Proteomic dataset\20100530_methyl250mM_1.RAW

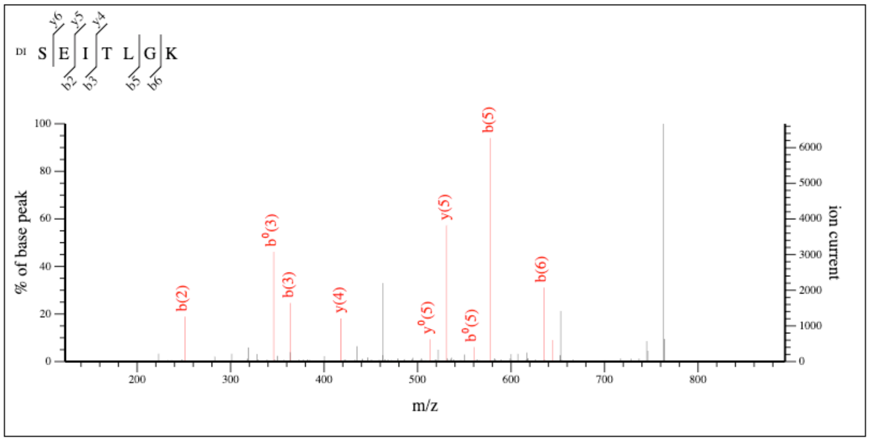

**Monoisotopic mass of neutral peptide Mr(calc):** 780.4864
**Fixed modifications:** Carbamidomethyl (C) (apply to specified residues or termini only)
**Variable modifications:**
**N-term :** Dimethyl:2H(6) (Protein N-term)
**Ions Score:** 29 **Expect:** 0.016
**Matches :** 10/27 fragment ions using 18 most intense peaks (help)

| # | b | $b^0$ | Seq. | y | y* | $y^0$ | # |
|---|---|---|---|---|---|---|---|
| 1 | 122.1083 | 104.0977 | S | | | | 7 |
| 2 | ***251.1509*** | 233.1403 | E | 660.3927 | 643.3661 | 642.3821 | 6 |
| 3 | ***364.2349*** | 346.2244 | I | **531.3501** | 514.3235 | 513.3395 | 5 |
| 4 | 465.2826 | 447.2720 | T | **418.2660** | 401.2395 | 400.2554 | 4 |
| 5 | ***578.3667*** | 560.3561 | L | 317.2183 | 300.1918 | | 3 |
| 6 | ***635.3881*** | 617.3776 | G | 204.1343 | 187.1077 | | 2 |
| 7 | | | K | 147.1128 | 130.0863 | | 1 |

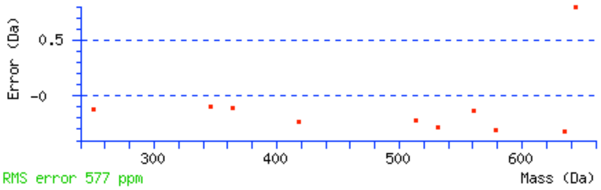


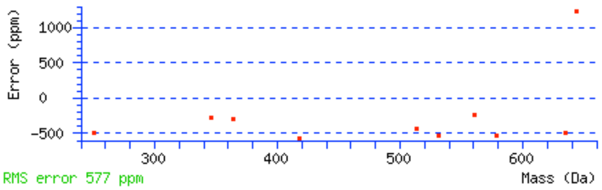


MS/MS Fragmentation of **SEITLGK**
Found in **P16467** in **Uniprot_yeastS288C**, Pyruvate decarboxylase isozyme 2 OS=Saccharomyces cerevisiae (strain ATCC 204508 / S288c) OX=559292 GN=PDC5 PE=1 SV=4

Match to Query 2435: 780.449755 from(781.457031,1+) intensity(273253.8438) rtinseconds(1027.5334) index(2561)
Title: 20100530_methyl250mM_1.2849.2849.1
Data file D:\Panyue\Proteomic dataset\20100530_methyl250mM_1.RAW

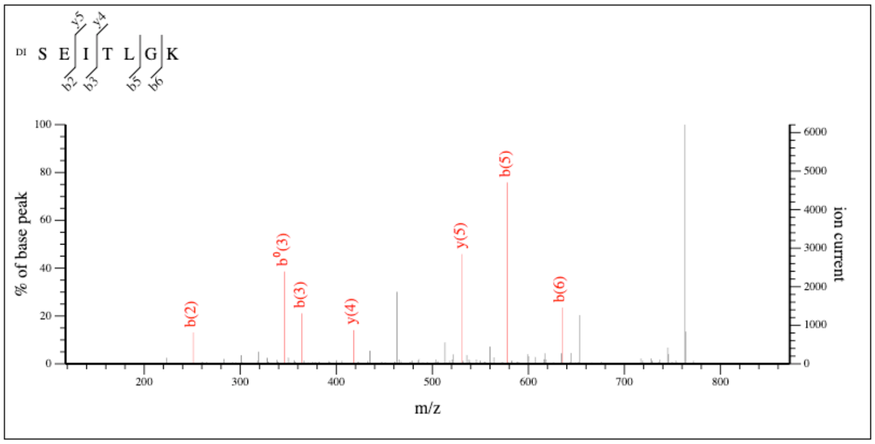

**Monoisotopic mass of neutral peptide Mr(calc):** 780.4864
**Fixed modifications:** Carbamidomethyl (C) (apply to specified residues or termini only)
**Variable modifications:**
**N-term :** Dimethyl:2H(6) (Protein N-term)
**Ions Score:** 31 **Expect:** 0.0096
**Matches :** 7/27 fragment ions using 12 most intense peaks (help)

| # | b | $b^0$ | Seq. | y | y* | $y^0$ | # |
|---|---|---|---|---|---|---|---|
| 1 | 122.1083 | 104.0977 | S | | | | 7 |
| 2 | 251.1509 | 233.1403 | E | 660.3927 | 643.3661 | 642.3821 | 6 |
| 3 | 364.2349 | 346.2244 | I | 531.3501 | 514.3235 | 513.3395 | 5 |
| 4 | 465.2826 | 447.2720 | T | 418.2660 | 401.2395 | 400.2554 | 4 |
| 5 | 578.3667 | 560.3561 | L | 317.2183 | 300.1918 | | 3 |
| 6 | 635.3881 | 617.3776 | G | 204.1343 | 187.1077 | | 2 |
| 7 | | | K | 147.1128 | 130.0863 | | 1 |

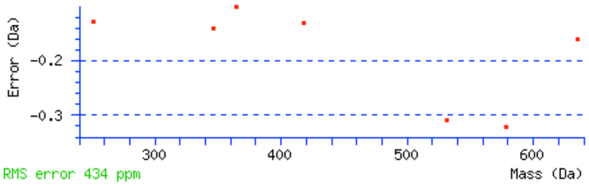


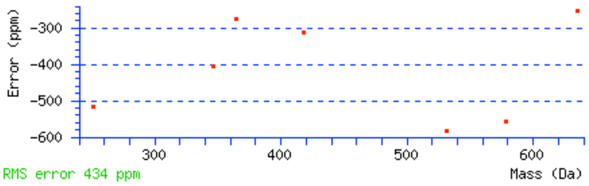


### 15. Pdc1 (2 peptides in 250mM fraction 1)

MS/MS Fragmentation of **SEITLGK**
Found in **P06169** in **Uniprot_yeastS288C**, Pyruvate decarboxylase isozyme 1 OS=Saccharomyces cerevisiae (strain ATCC 204508 / S288c) OX=559292 GN=PDC1 PE=1 SV=7

Match to Query 2434: 780.449267 from(781.456543,1+) intensity(563499.6875) rtinseconds(1029.388) index(2565)
Title: 20100530_methyl250mM_1.2854.2854.1
Data file D:\Panyue\Proteomic dataset\20100530_methyl250mM_1.RAW

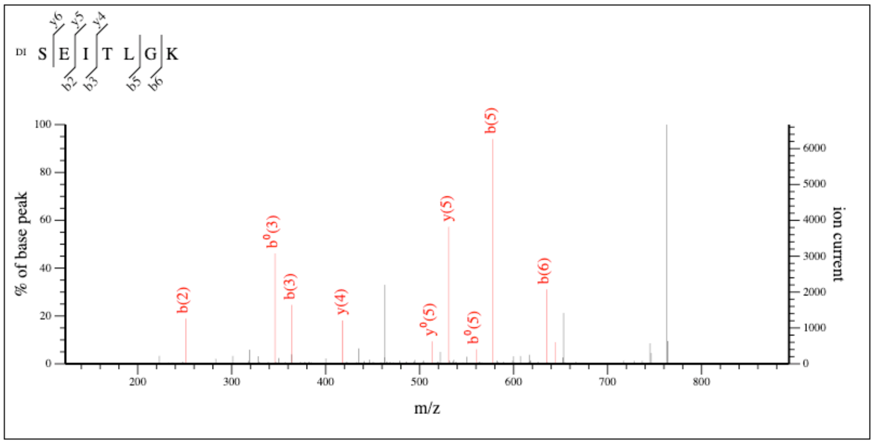

**Monoisotopic mass of neutral peptide Mr(calc):** 780.4864
**Fixed modifications:** Carbamidomethyl (C) (apply to specified residues or termini only)
**Variable modifications:**
**N-term :** Dimethyl:2H(6) (Protein N-term)
**Ions Score:** 29 **Expect:** 0.016
**Matches :** 10/27 fragment ions using 18 most intense peaks (help)

| # | b | $b^0$ | Seq. | y | y* | $y^0$ | # |
|---|---|---|---|---|---|---|---|
| 1 | 122.1083 | 104.0977 | S | | | | 7 |
| 2 | 251.1509 | 233.1403 | E | 660.3927 | 643.3661 | 642.3821 | 6 |
| 3 | 364.2349 | 346.2244 | I | 531.3501 | 514.3235 | 513.3395 | 5 |
| 4 | 465.2826 | 447.2720 | T | 418.2660 | 401.2395 | 400.2554 | 4 |
| 5 | 578.3667 | 560.3561 | L | 317.2183 | 300.1918 | | 3 |
| 6 | 635.3881 | 617.3776 | G | 204.1343 | 187.1077 | | 2 |
| 7 | | | K | 147.1128 | 130.0863 | | 1 |

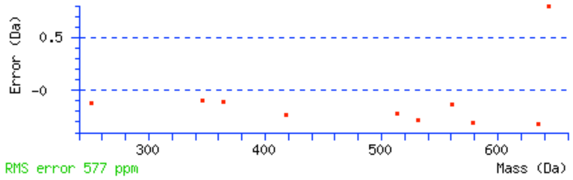


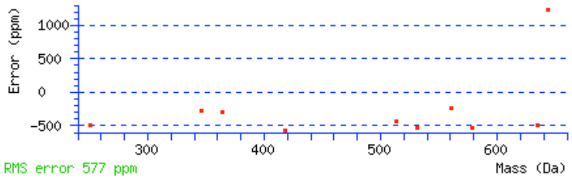


MS/MS Fragmentation of **SEITLGK**
Found in **P06169** in **Uniprot_yeastS288C**, Pyruvate decarboxylase isozyme 1 OS=Saccharomyces cerevisiae (strain ATCC 204508 / S288c) OX=559292 GN=PDC1 PE=1 SV=7

Match to Query 2435: 780.449755 from(781.457031,1+) intensity(273253.8438) rtinseconds(1027.5334) index(2561)
Title: 20100530_methyl250mM_1.2849.2849.1
Data file D:\Panyue\Proteomic dataset\20100530_methyl250mM_1.RAW

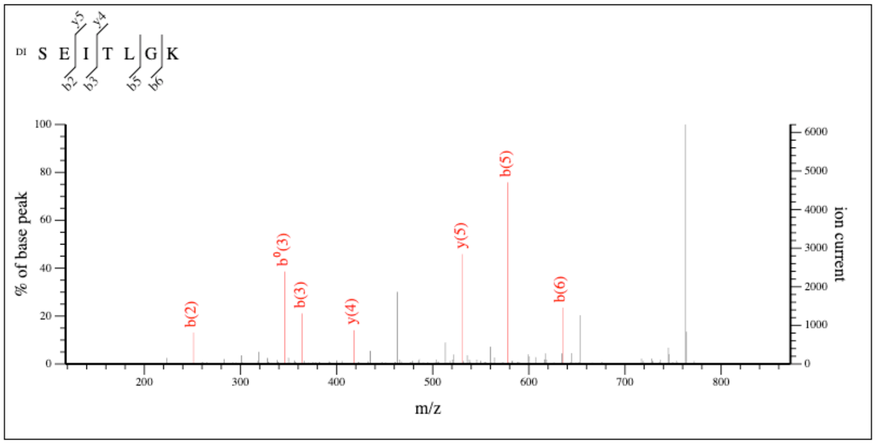

**Monoisotopic mass of neutral peptide Mr(calc):** 780.4864
**Fixed modifications:** Carbamidomethyl (C) (apply to specified residues or termini only)
**Variable modifications:**
**N-term :** Dimethyl:2H(6) (Protein N-term)
**Ions Score:** 31 **Expect:** 0.0096
**Matches :** 7/27 fragment ions using 12 most intense peaks (help)

| # | b | $b^0$ | Seq. | y | y* | $y^0$ | # |
|---|---|---|---|---|---|---|---|
| 1 | 122.1083 | 104.0977 | S | | | | 7 |
| 2 | ***251.1509*** | 233.1403 | E | 660.3927 | 643.3661 | 642.3821 | 6 |
| 3 | ***364.2349*** | 346.2244 | I | **531.3501** | 514.3235 | 513.3395 | 5 |
| 4 | 465.2826 | 447.2720 | T | **418.2660** | 401.2395 | 400.2554 | 4 |
| 5 | ***578.3667*** | 560.3561 | L | 317.2183 | 300.1918 | | 3 |
| 6 | ***635.3881*** | 617.3776 | G | 204.1343 | 187.1077 | | 2 |
| 7 | | | K | 147.1128 | 130.0863 | | 1 |

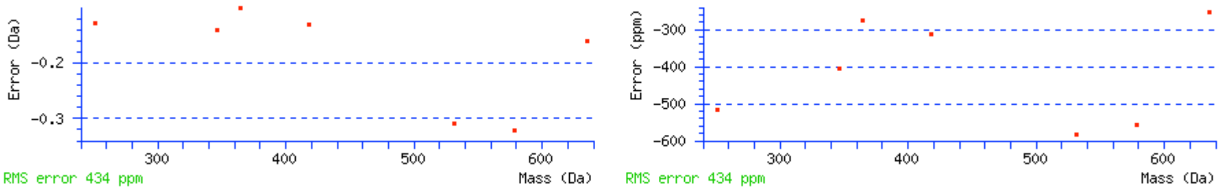


### 16. Nop14 (2 peptides in 250mM fraction 1)

MS/MS Fragmentation of AGSQLKNLK
Found in Q99207 in Uniprot_yeastS288C, Nucleolar complex protein 14 OS=Saccharomyces cerevisiae (strain ATCC 204508 / S288c) OX=559292 GN=NOP14 PE=1 SV=1

Match to Query 4835: 1008.670825 from(1009.678101,1+) intensity(306642.0938) rtinseconds(1644.3817) index(4258)
Title: 20100530_methyl250mM_1.4737.4737.1
Data file D:\Panyue\Proteomic dataset\20100530_methyl250mM_1.RAW

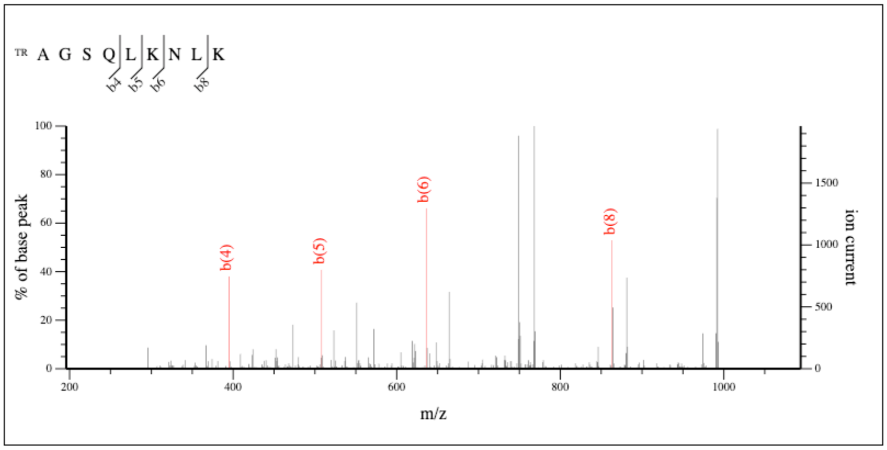

**Monoisotopic mass of neutral peptide Mr(calc):** 1008.6641
**Fixed modifications:** Carbamidomethyl (C) (apply to specified residues or termini only)
**Variable modifications:**
**N-term :** Trimethyl:2H(9) (Protein N-term)
**Ions Score:** 26 **Expect:** 0.0044
**Matches :** 4/37 fragment ions using 7 most intense peaks (help)

| # | b | b* | b$^0$ | Seq. | y | y* | y$^0$ | # |
|---|---|---|---|---|---|---|---|---|
| 1 | 123.1478 | | | A | | | | 9 |
| 2 | 180.1693 | | | G | 887.5309 | 870.5043 | 869.5203 | 8 |
| 3 | 267.2013 | | 249.1908 | S | 830.5094 | 813.4829 | 812.4989 | 7 |
| 4 | ***395.2599*** | 378.2334 | 377.2493 | Q | 743.4774 | 726.4509 | | 6 |
| 5 | ***508.3440*** | 491.3174 | 490.3334 | L | 615.4188 | 598.3923 | | 5 |
| 6 | ***636.4389*** | 619.4124 | 618.4284 | K | 502.3348 | 485.3082 | | 4 |
| 7 | 750.4819 | 733.4553 | 732.4713 | N | 374.2398 | 357.2132 | | 3 |
| 8 | ***863.5659*** | 846.5394 | 845.5554 | L | 260.1969 | 243.1703 | | 2 |
| 9 | | | | K | 147.1128 | 130.0863 | | 1 |

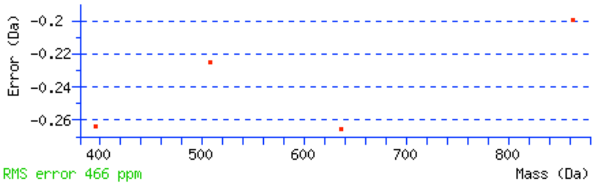


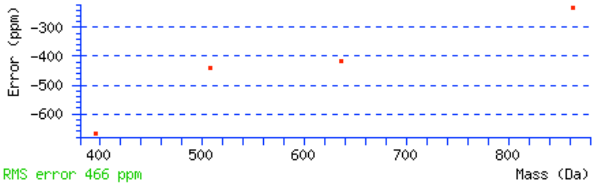


MS/MS Fragmentation of **AGSQLKNLK**
Found in **Q99207** in **Uniprot_yeastS288C**, Nucleolar complex protein 14 OS=Saccharomyces cerevisiae (strain ATCC 204508 / S288c) OX=559292 GN=NOP14 PE=1 SV=1

Match to Query 4836: 1008.671191 from(1009.678467,1+) intensity(239066.4844) rtinseconds(1642.6625) index(4254)
Title: 20100530_methyl250mM_1.4732.4732.1
Data file D:\Panyue\Proteomic dataset\20100530_methyl250mM_1.RAW

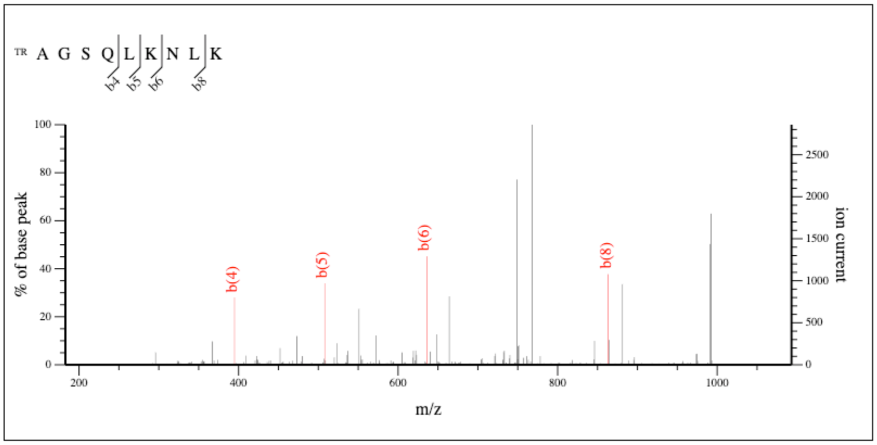

**Monoisotopic mass of neutral peptide Mr(calc):** 1008.6641
**Fixed modifications:** Carbamidomethyl (C) (apply to specified residues or termini only)
**Variable modifications:**
**N-term :** Trimethyl:2H(9) (Protein N-term)
**Ions Score:** 27 **Expect:** 0.0039
**Matches :** 4/37 fragment ions using 8 most intense peaks (help)

| # | b | b* | $b^0$ | Seq. | y | y* | $y^0$ | # |
|---|---|---|---|---|---|---|---|---|
| 1 | 123.1478 | | | A | | | | 9 |
| 2 | 180.1693 | | | G | 887.5309 | 870.5043 | 869.5203 | 8 |
| 3 | 267.2013 | | 249.1908 | S | 830.5094 | 813.4829 | 812.4989 | 7 |
| 4 | *395.2599* | 378.2334 | 377.2493 | Q | 743.4774 | 726.4509 | | 6 |
| 5 | *508.3440* | 491.3174 | 490.3334 | L | 615.4188 | 598.3923 | | 5 |
| 6 | *636.4389* | 619.4124 | 618.4284 | K | 502.3348 | 485.3082 | | 4 |
| 7 | 750.4819 | 733.4553 | 732.4713 | N | 374.2398 | 357.2132 | | 3 |
| 8 | *863.5659* | 846.5394 | 845.5554 | L | 260.1969 | 243.1703 | | 2 |
| 9 | | | | K | 147.1128 | 130.0863 | | 1 |

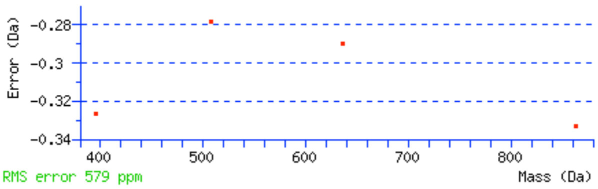


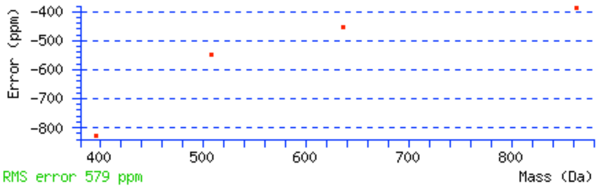


### 17. Eft1 (1 peptide in 250mM fraction 1)

MS/MS Fragmentation of **VAFTVDQMRSLMDK**
Found in **P32324** in **Uniprot_yeastS288C**, Elongation factor 2 OS=Saccharomyces cerevisiae (strain ATCC 204508 / S288c) OX=559292 GN=EFT1 PE=1 SV=1

Match to Query 9664: 1706.870824 from(854.442688,2+) intensity(3001281.5000) rtinseconds(2658.2495) index(7034)
Title: 20100530_methyl250mM_1.7825.7825.2
Data file D:\Panyue\Proteomic dataset\20100530_methyl250mM_1.RAW

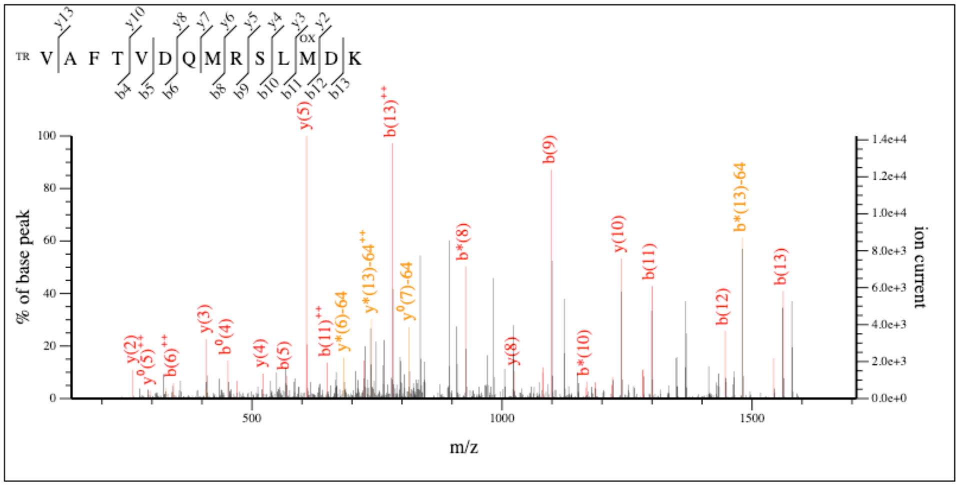

**Monoisotopic mass of neutral peptide Mr(calc):** 1706.9046
**Fixed modifications:** Carbamidomethyl (C) (apply to specified residues or termini only)
**Variable modifications:**
**N-term :** Trimethyl:2H(9) (Protein N-term)
**M12** : Oxidation (M), with neutral losses 0.0000(shown in table), 63.9983
**Ions Score:** 22 **Expect:** 0.11
**Matches :** 44/214 fragment ions using 84 most intense peaks (help)

| # | b | $b^{++}$ | b* | $b^{*++}$ | $b^0$ | $b^{0++}$ | Seq. | y | $y^{++}$ | y* | $y^{*++}$ | $y^0$ | $y^{0++}$ | # |
|---|---|---|---|---|---|---|---|---|---|---|---|---|---|---|
| 1 | 151.1791 | 76.0932 | | | | | V | | | | | | | 14 |
| 2 | 222.2162 | 111.6118 | | | | | A | 1557.7400 | 779.3736 | 1540.7134 | 770.8604 | 1539.7294 | 770.3683 | 13 |
| 3 | 369.2847 | 185.1460 | | | | | F | 1486.7029 | 743.8551 | 1469.6763 | 735.3418 | 1468.6923 | 734.8498 | 12 |
| 4 | ***470.3323*** | 235.6698 | | | 452.3218 | 226.6645 | T | 1339.6344 | 670.3209 | 1322.6079 | 661.8076 | 1321.6239 | 661.3156 | 11 |
| 5 | ***569.4008*** | 285.2040 | | | 551.3902 | 276.1987 | V | **1238.5868** | 619.7970 | 1221.5602 | 611.2837 | 1220.5762 | 610.7917 | 10 |
| 6 | 684.4277 | **342.7175** | | | 666.4171 | 333.7122 | D | 1139.5184 | 570.2628 | 1122.4918 | 561.7495 | 1121.5078 | 561.2575 | 9 |
| 7 | 812.4863 | 406.7468 | 795.4597 | 398.2335 | 794.4757 | 397.7415 | Q | **1024.4914** | 512.7493 | 1007.4649 | 504.2361 | 1006.4808 | 503.7441 | 8 |
| 8 | 943.5268 | 472.2670 | 926.5002 | 463.7537 | 925.5162 | 463.2617 | M | 896.4328 | 448.7201 | 879.4063 | 440.2068 | 878.4223 | 439.7148 | 7 |
| 9 | ***1099.6279*** | 550.3176 | 1082.6013 | 541.8043 | 1081.6173 | 541.3123 | R | 765.3924 | 383.1998 | 748.3658 | 374.6865 | 747.3818 | 374.1945 | 6 |
| 10 | ***1186.6599*** | 593.8336 | 1169.6333 | 585.3203 | 1168.6493 | 584.8283 | S | **609.2912** | 305.1493 | 592.2647 | 296.6360 | 591.2807 | 296.1440 | 5 |
| 11 | ***1299.7440*** | **650.3756** | 1282.7174 | 641.8623 | 1281.7334 | 641.3703 | L | **522.2592** | 261.6332 | 505.2327 | 253.1200 | 504.2486 | 252.6280 | 4 |
| 12 | ***1446.7794*** | **723.8933** | 1429.7528 | 715.3800 | 1428.7688 | 714.8880 | M | **409.1751** | 205.0912 | 392.1486 | 196.5779 | 391.1646 | 196.0859 | 3 |
| 13 | ***1561.8063*** | **781.4068** | 1544.7798 | 772.8935 | 1543.7957 | 772.4015 | D | **262.1397** | 131.5735 | 245.1132 | 123.0602 | 244.1292 | 122.5682 | 2 |
| 14 | | | | | | | K | 147.1128 | 74.0600 | 130.0863 | 65.5468 | | | 1 |

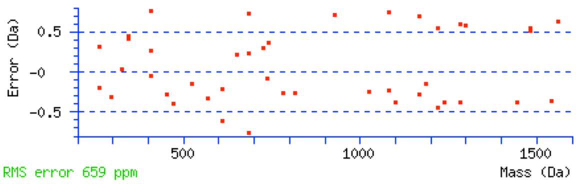


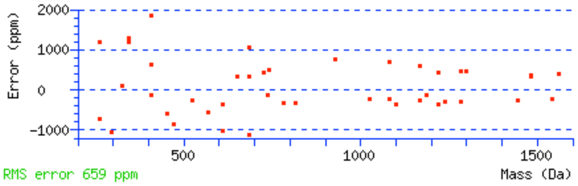


### 18. Asc1 (2 peptides in 250mM fraction 1)

MS/MS Fragmentation of **ASNEVLVLR**
Found in **P38011** in **Uniprot_yeastS288C**, Guanine nucleotide-binding protein subunit beta-like protein OS=Saccharomyces cerevisiae (strain ATCC 204508 / S288c) OX=559292 GN=ASC1 PE=1 SV=4

Match to Query 5198: 1050.680712 from(1051.687988,1+) intensity(782943.0000) rtinseconds(3331.8371) index(8866)
Title: 20100530_methyl250mM_1.9862.9862.1
Data file D:\Panyue\Proteomic dataset\20100530_methyl250mM_1.RAW

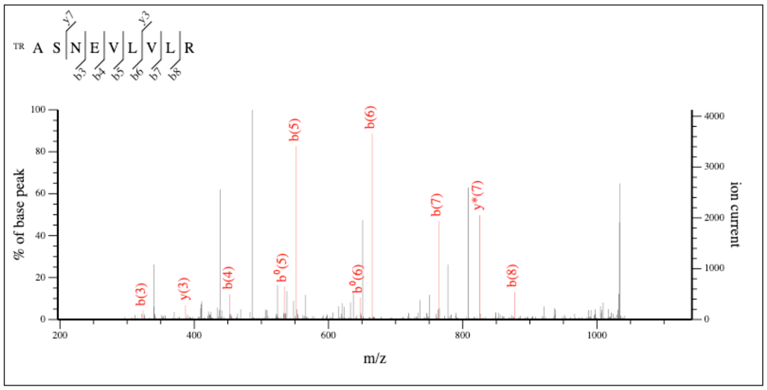

**Monoisotopic mass of neutral peptide Mr(calc):** 1050.6747
**Fixed modifications:** Carbamidomethyl (C) (apply to specified residues or termini only)
**Variable modifications:**
**N-term :** Trimethyl:2H(9) (Protein N-term)
**Ions Score:** 22 **Expect:** 0.015
**Matches :** 11/40 fragment ions using 32 most intense peaks (help)

| # | b | b* | $b^0$ | Seq. | y | y* | $y^0$ | # |
|---|---|---|---|---|---|---|---|---|
| 1 | 123.1478 | | | A | | | | 9 |
| 2 | 210.1799 | | 192.1693 | S | 929.5415 | 912.5149 | 911.5309 | 8 |
| 3 | ***324.2228*** | 307.1962 | 306.2122 | N | 842.5094 | 825.4829 | 824.4989 | 7 |
| 4 | ***453.2654*** | 436.2388 | 435.2548 | E | 728.4665 | 711.4400 | 710.4559 | 6 |
| 5 | ***552.3338*** | 535.3072 | 534.3232 | V | 599.4239 | 582.3974 | | 5 |
| 6 | ***665.4179*** | 648.3913 | 647.4073 | L | 500.3555 | 483.3289 | | 4 |
| 7 | ***764.4863*** | 747.4597 | 746.4757 | V | 387.2714 | 370.2449 | | 3 |
| 8 | ***877.5703*** | 860.5438 | 859.5598 | L | 288.2030 | 271.1765 | | 2 |
| 9 | | | | R | 175.1190 | 158.0924 | | 1 |

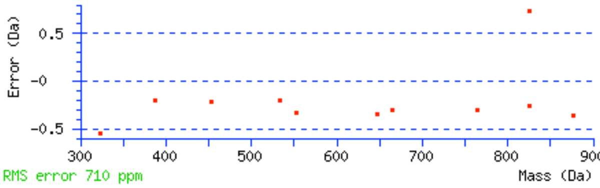


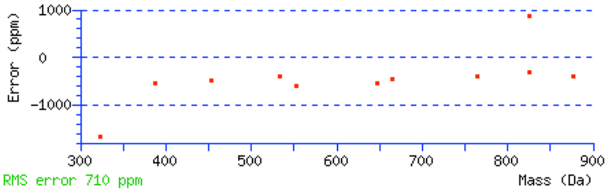


MS/MS Fragmentation of **ASNEVLVLR**
Found in **P38011** in **Uniprot_yeastS288C**, Guanine nucleotide-binding protein subunit beta-like protein OS=Saccharomyces cerevisiae (strain ATCC 204508 / S288c) OX=559292 GN=ASC1 PE=1 SV=4

Match to Query 5199: 1050.681445 from(1051.688721,1+) intensity(1331435.8750) rtinseconds(3334.154) index(8872)
Title: 20100530_methyl250mM_1.9869.9869.1
Data file D:\Panyue\Proteomic dataset\20100530_methyl250mM_1.RAW

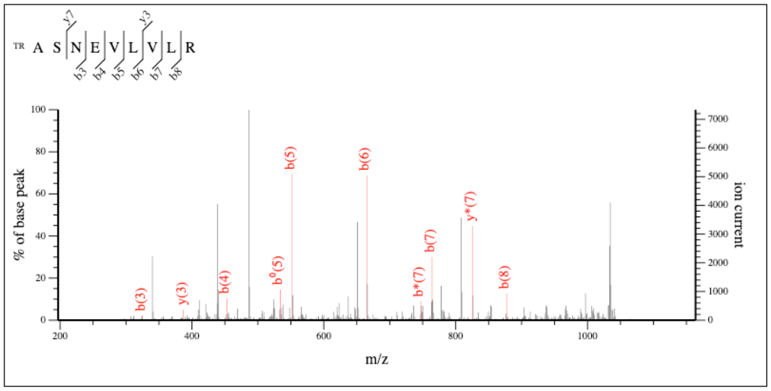

**Monoisotopic mass of neutral peptide Mr(calc):** 1050.6747
**Fixed modifications:** Carbamidomethyl (C) (apply to specified residues or termini only)
**Variable modifications:**
**N-term :** Trimethyl:2H(9) (Protein N-term)
**Ions Score:** 20 **Expect:** 0.022
**Matches :** 11/40 fragment ions using 40 most intense peaks (help)

| # | b | b* | b0 | Seq. | y | y* | y0 | # |
|---|---|---|---|---|---|---|---|---|
| 1 | 123.1478 | | | A | | | | 9 |
| 2 | 210.1799 | | 192.1693 | S | 929.5415 | 912.5149 | 911.5309 | 8 |
| 3 | ***324.2228*** | 307.1962 | 306.2122 | N | 842.5094 | 825.4829 | 824.4989 | 7 |
| 4 | ***453.2654*** | 436.2388 | 435.2548 | E | 728.4665 | 711.4400 | 710.4559 | 6 |
| 5 | ***552.3338*** | 535.3072 | 534.3232 | V | 599.4239 | 582.3974 | | 5 |
| 6 | ***665.4179*** | 648.3913 | 647.4073 | L | 500.3555 | 483.3289 | | 4 |
| 7 | ***764.4863*** | 747.4597 | 746.4757 | V | 387.2714 | 370.2449 | | 3 |
| 8 | ***877.5703*** | 860.5438 | 859.5598 | L | 288.2030 | 271.1765 | | 2 |
| 9 | | | | R | 175.1190 | 158.0924 | | 1 |

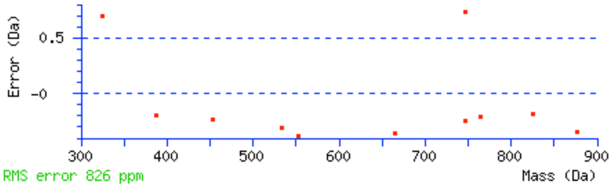


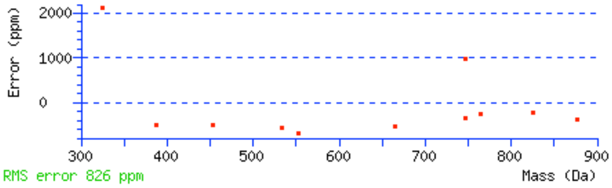


### 19. Pdc5 (2 peptides in 250mM fraction 2)

MS/MS Fragmentation of **SEITLGK**
Found in **P16467** in **Uniprot_yeastS288C**, Pyruvate decarboxylase isozyme 2 OS=Saccharomyces cerevisiae (strain ATCC 204508 / S288c) OX=559292 GN=PDC5 PE=1 SV=4

Match to Query 2505: 780.449206 from(781.456482,1+) intensity(571654.5000) rtinseconds(1067.6832) index(2638)
Title: 20100530_methyl250mM_2.2937.2937.1
Data file D:\Panyue\Proteomic dataset\20100530_methyl250mM_2.RAW

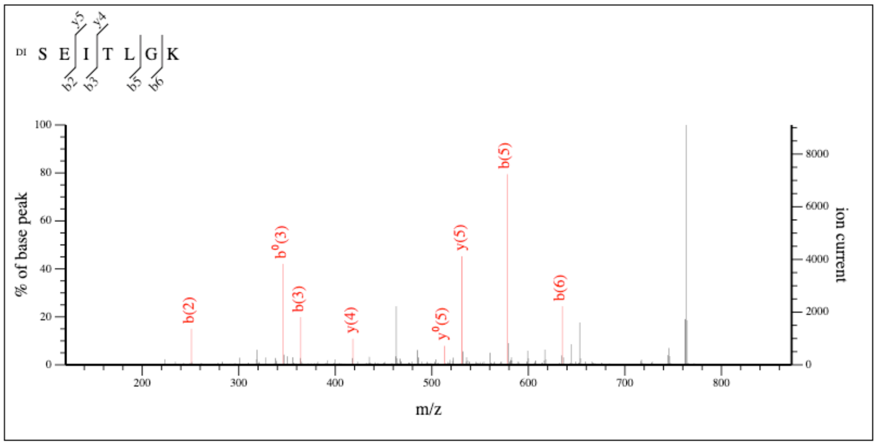

**Monoisotopic mass of neutral peptide Mr(calc):** 780.4864
**Fixed modifications:** Carbamidomethyl (C) (apply to specified residues or termini only)
**Variable modifications:**
**N-term :** Dimethyl:2H(6) (Protein N-term)
**Ions Score:** 25 **Expect:** 0.04
**Matches :** 8/27 fragment ions using 18 most intense peaks (help)

| # | b | b$^0$ | Seq. | y | y* | y$^0$ | # |
|---|---|---|---|---|---|---|---|
| 1 | 122.1083 | 104.0977 | S | | | | 7 |
| 2 | ***251.1509*** | 233.1403 | E | 660.3927 | 643.3661 | 642.3821 | 6 |
| 3 | ***364.2349*** | 346.2244 | I | **531.3501** | 514.3235 | 513.3395 | 5 |
| 4 | 465.2826 | 447.2720 | T | **418.2660** | 401.2395 | 400.2554 | 4 |
| 5 | ***578.3667*** | 560.3561 | L | 317.2183 | 300.1918 | | 3 |
| 6 | ***635.3881*** | 617.3776 | G | 204.1343 | 187.1077 | | 2 |
| 7 | | | K | 147.1128 | 130.0863 | | 1 |

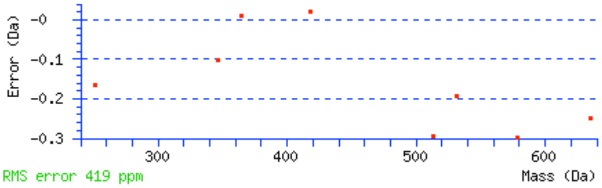


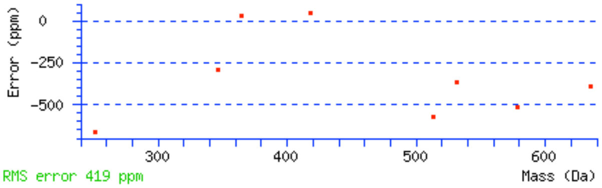


MS/MS Fragmentation of **SEITLGK**
Found in **P16467** in **Uniprot_yeastS288C**, Pyruvate decarboxylase isozyme 2 OS=Saccharomyces cerevisiae (strain ATCC 204508 / S288c) OX=559292 GN=PDC5 PE=1 SV=4

Match to Query 2506: 780.449877 from(781.457153,1+) intensity(275676.1250) rtinseconds(1066.1433) index(2635)
Title: 20100530_methyl250mM_2.2933.2933.1
Data file D:\Panyue\Proteomic dataset\20100530_methyl250mM_2.RAW

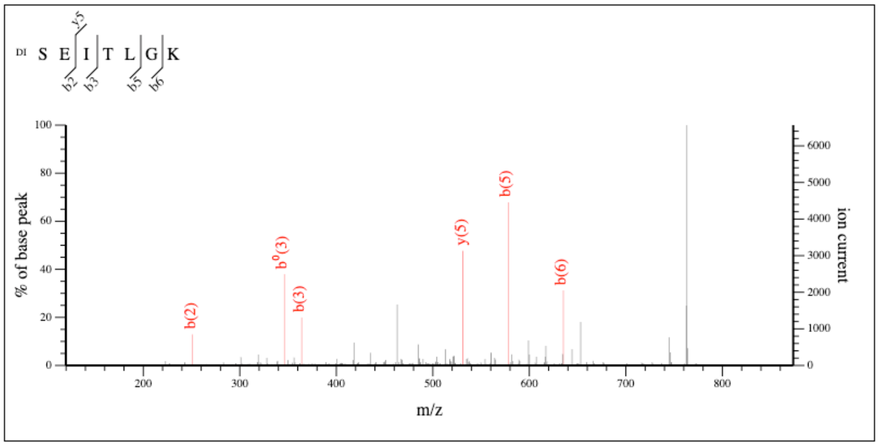

**Monoisotopic mass of neutral peptide Mr(calc):** 780.4864
**Fixed modifications:** Carbamidomethyl (C) (apply to specified residues or termini only)
**Variable modifications:**
**N-term :** Dimethyl:2H(6) (Protein N-term)
**Ions Score:** 24 **Expect:** 0.053
**Matches :** 6/27 fragment ions using 12 most intense peaks (help)

| # | b | $b^0$ | Seq. | y | y* | $y^0$ | # |
|---|---|---|---|---|---|---|---|
| 1 | 122.1083 | 104.0977 | S | | | | 7 |
| 2 | 251.1509 | 233.1403 | E | 660.3927 | 643.3661 | 642.3821 | 6 |
| 3 | 364.2349 | 346.2244 | I | 531.3501 | 514.3235 | 513.3395 | 5 |
| 4 | 465.2826 | 447.2720 | T | 418.2660 | 401.2395 | 400.2554 | 4 |
| 5 | 578.3667 | 560.3561 | L | 317.2183 | 300.1918 | | 3 |
| 6 | 635.3881 | 617.3776 | G | 204.1343 | 187.1077 | | 2 |
| 7 | | | K | 147.1128 | 130.0863 | | 1 |

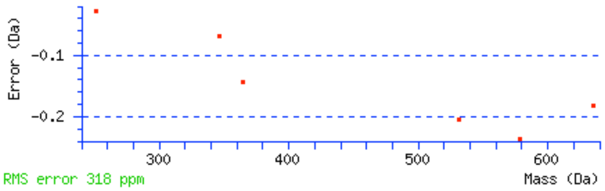


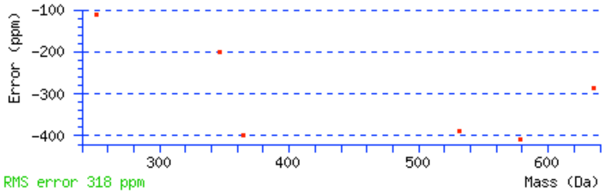


### 20. Pdc1 (2 peptides in 250mM fraction 2)

MS/MS Fragmentation of **SEITLGK**
Found in **P06169** in **Uniprot_yeastS288C**, Pyruvate decarboxylase isozyme 1 OS=Saccharomyces cerevisiae (strain ATCC 204508 / S288c) OX=559292 GN=PDC1 PE=1 SV=7

Match to Query 2506: 780.449877 from(781.457153,1+) intensity(275676.1250) rtinseconds(1066.1433) index(2635)
Title: 20100530_methyl250mM_2.2933.2933.1
Data file D:\Panyue\Proteomic dataset\20100530_methyl250mM_2.RAW

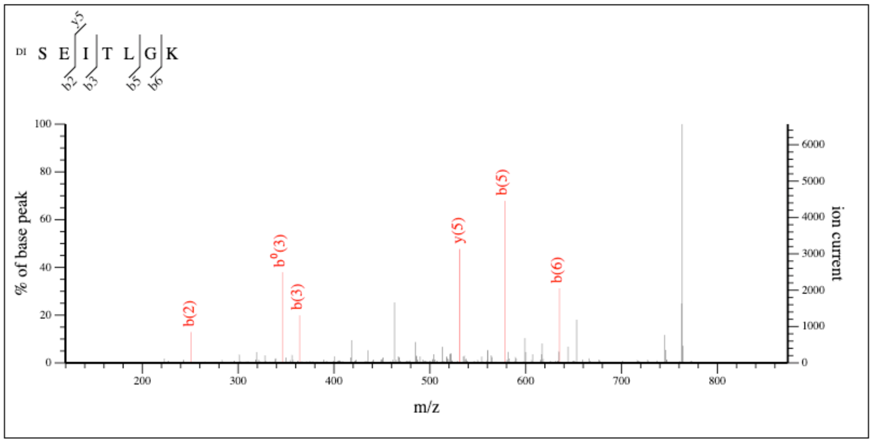

**Monoisotopic mass of neutral peptide Mr(calc):** 780.4864
**Fixed modifications:** Carbamidomethyl (C) (apply to specified residues or termini only)
**Variable modifications:**
**N-term :** Dimethyl:2H(6) (Protein N-term)
**Ions Score:** 24 **Expect:** 0.053
**Matches :** 6/27 fragment ions using 12 most intense peaks (help)

| # | b | $b^0$ | Seq. | y | y* | $y^0$ | # |
|---|---|---|---|---|---|---|---|
| 1 | 122.1083 | 104.0977 | S | | | | 7 |
| 2 | ***251.1509*** | 233.1403 | E | 660.3927 | 643.3661 | 642.3821 | 6 |
| 3 | ***364.2349*** | 346.2244 | I | 531.3501 | 514.3235 | 513.3395 | 5 |
| 4 | 465.2826 | 447.2720 | T | 418.2660 | 401.2395 | 400.2554 | 4 |
| 5 | ***578.3667*** | 560.3561 | L | 317.2183 | 300.1918 | | 3 |
| 6 | ***635.3881*** | 617.3776 | G | 204.1343 | 187.1077 | | 2 |
| 7 | | | K | 147.1128 | 130.0863 | | 1 |

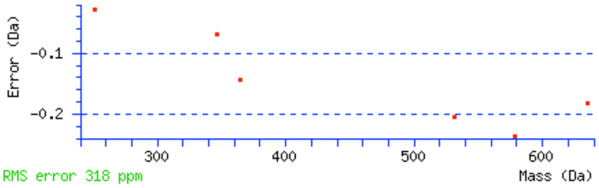


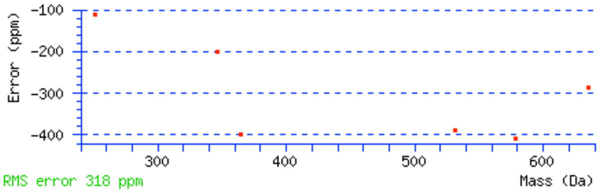


MS/MS Fragmentation of **SEITLGK**
Found in **P06169** in **Uniprot_yeastS288C**, Pyruvate decarboxylase isozyme 1 OS=Saccharomyces cerevisiae (strain ATCC 204508 / S288c) OX=559292 GN=PDC1 PE=1 SV=7

Match to Query 2505: 780.449206 from(781.456482,1+) intensity(571654.5000) rtinseconds(1067.6832) index(2638)
Title: 20100530_methyl250mM_2.2937.2937.1
Data file D:\Panyue\Proteomic dataset\20100530_methyl250mM_2.RAW

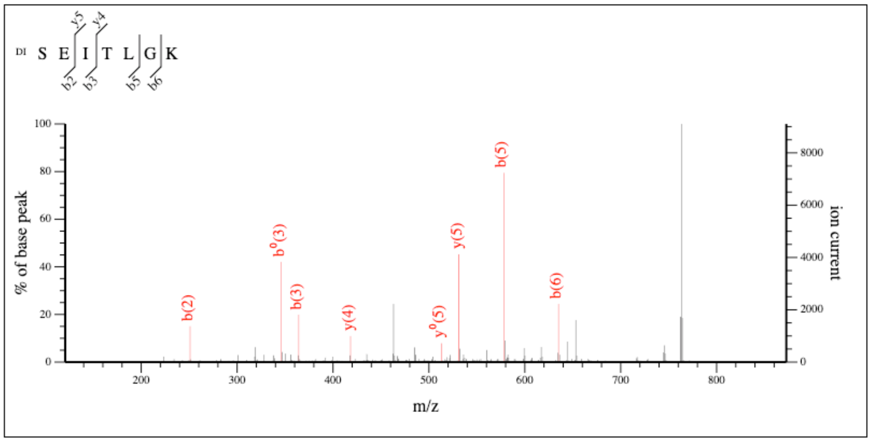

**Monoisotopic mass of neutral peptide Mr(calc):** 780.4864
**Fixed modifications:** Carbamidomethyl (C) (apply to specified residues or termini only)
**Variable modifications:**
**N-term :** Dimethyl:2H(6) (Protein N-term)
**Ions Score:** 25 **Expect:** 0.04
**Matches :** 8/27 fragment ions using 18 most intense peaks (help)

| # | b | $b^0$ | Seq. | y | y* | $y^0$ | # |
|---|---|---|---|---|---|---|---|
| 1 | 122.1083 | 104.0977 | S | | | | 7 |
| 2 | ***251.1509*** | 233.1403 | E | 660.3927 | 643.3661 | 642.3821 | 6 |
| 3 | ***364.2349*** | 346.2244 | I | **531.3501** | 514.3235 | 513.3395 | 5 |
| 4 | 465.2826 | 447.2720 | T | **418.2660** | 401.2395 | 400.2554 | 4 |
| 5 | ***578.3667*** | 560.3561 | L | 317.2183 | 300.1918 | | 3 |
| 6 | ***635.3881*** | 617.3776 | G | 204.1343 | 187.1077 | | 2 |
| 7 | | | K | 147.1128 | 130.0863 | | 1 |

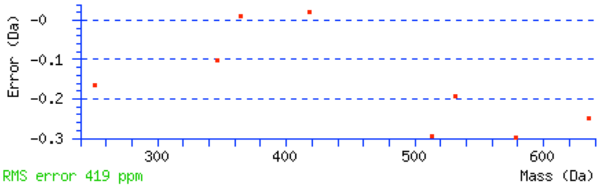


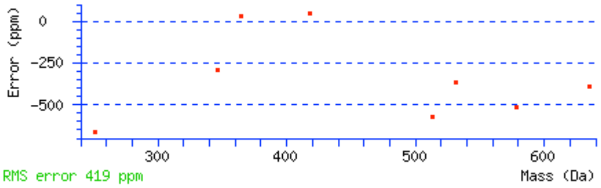


### 21. Nop14 (2 peptides in 250mM fraction2)

MS/MS Fragmentation of **AGSQLKNLK**
Found in **Q99207** in **Uniprot_yeastS288C**, Nucleolar complex protein 14 OS=Saccharomyces cerevisiae (strain ATCC 204508 / S288c) OX=559292 GN=NOP14 PE=1 SV=1

Match to Query 4909: 1008.670336 from(1009.677612,1+) intensity(1310980.8750) rtinseconds(1708.0658) index(4363)
Title: 20100530_methyl250mM_2.4857.4857.1
Data file D:\Panyue\Proteomic dataset\20100530_methyl250mM_2.RAW

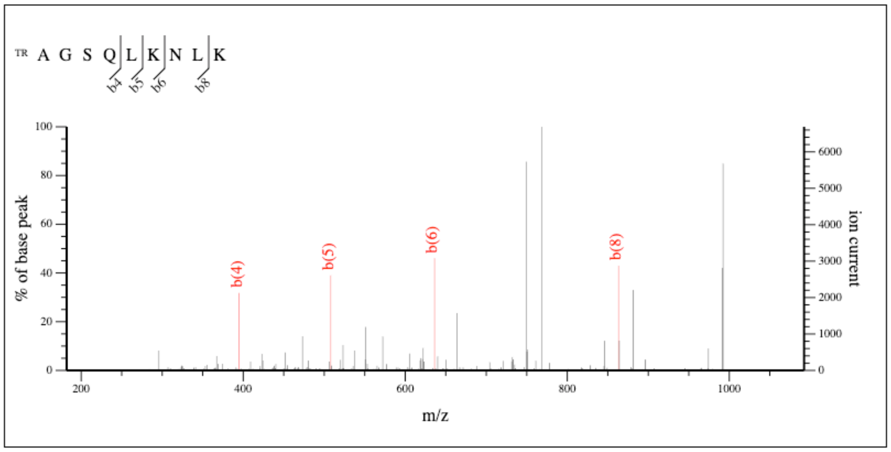

**Monoisotopic mass of neutral peptide Mr(calc):** 1008.6641
**Fixed modifications:** Carbamidomethyl (C) (apply to specified residues or termini only)
**Variable modifications:**
**N-term :** Trimethyl:2H(9) (Protein N-term)
**Ions Score:** 28 **Expect:** 0.0035
**Matches :** 4/37 fragment ions using 8 most intense peaks (help)

| # | b | b* | b$^{0}$ | Seq. | y | y* | y$^{0}$ | # |
|---|---|---|---|---|---|---|---|---|
| 1 | 123.1478 | | | A | | | | 9 |
| 2 | 180.1693 | | | G | 887.5309 | 870.5043 | 869.5203 | 8 |
| 3 | 267.2013 | | 249.1908 | S | 830.5094 | 813.4829 | 812.4989 | 7 |
| 4 | ***395.2599*** | 378.2334 | 377.2493 | Q | 743.4774 | 726.4509 | | 6 |
| 5 | ***508.3440*** | 491.3174 | 490.3334 | L | 615.4188 | 598.3923 | | 5 |
| 6 | ***636.4389*** | 619.4124 | 618.4284 | K | 502.3348 | 485.3082 | | 4 |
| 7 | 750.4819 | 733.4553 | 732.4713 | N | 374.2398 | 357.2132 | | 3 |
| 8 | ***863.5659*** | 846.5394 | 845.5554 | L | 260.1969 | 243.1703 | | 2 |
| 9 | | | | K | 147.1128 | 130.0863 | | 1 |

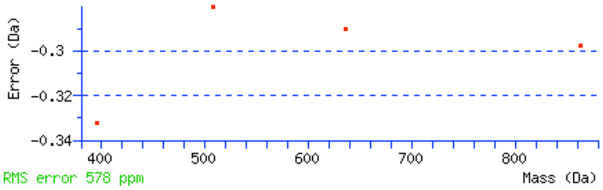


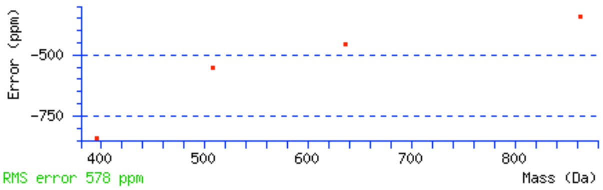


MS/MS Fragmentation of AGSQLKNLK
Found in Q99207 in Uniprot_yeastS288C, Nucleolar complex protein 14 OS=Saccharomyces cerevisiae (strain ATCC 204508 / S288c) OX=559292 GN=NOP14 PE=1 SV=1

Match to Query 4911: 1008.670458 from(1009.677734,1+) intensity(1486084.8750) rtinseconds(1705.1361) index(4355)
Title: 20100530_methyl250mM_2.4848.4848.1
Data file D:\Panyue\Proteomic dataset\20100530_methyl250mM_2.RAW

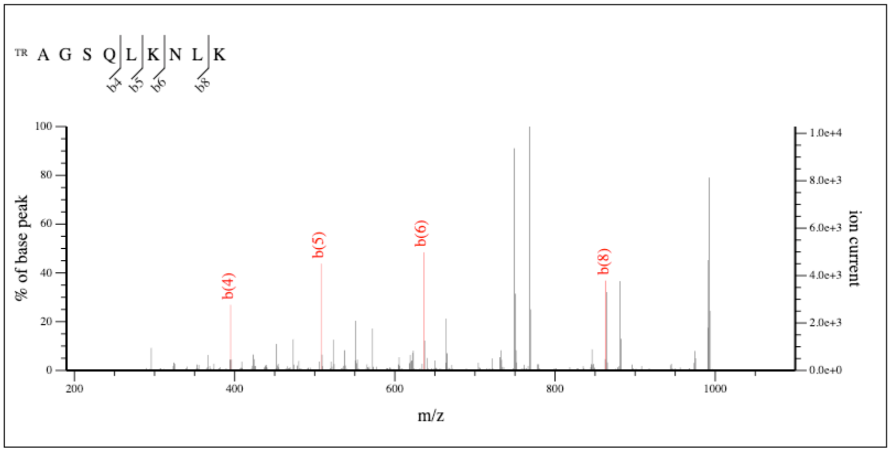

**Monoisotopic mass of neutral peptide Mr(calc):** 1008.6641
**Fixed modifications:** Carbamidomethyl (C) (apply to specified residues or termini only)
**Variable modifications:**
**N-term :** Trimethyl:2H(9) (Protein N-term)
**Ions Score:** 26 **Expect:** 0.0044
**Matches :** 4/37 fragment ions using 8 most intense peaks (help)

| # | b | b* | $b^0$ | Seq. | y | y* | $y^0$ | # |
|---|---|---|---|---|---|---|---|---|
| 1 | 123.1478 | | | A | | | | 9 |
| 2 | 180.1693 | | | G | 887.5309 | 870.5043 | 869.5203 | 8 |
| 3 | 267.2013 | | 249.1908 | S | 830.5094 | 813.4829 | 812.4989 | 7 |
| 4 | *395.2599* | 378.2334 | 377.2493 | Q | 743.4774 | 726.4509 | | 6 |
| 5 | *508.3440* | 491.3174 | 490.3334 | L | 615.4188 | 598.3923 | | 5 |
| 6 | *636.4389* | 619.4124 | 618.4284 | K | 502.3348 | 485.3082 | | 4 |
| 7 | 750.4819 | 733.4553 | 732.4713 | N | 374.2398 | 357.2132 | | 3 |
| 8 | *863.5659* | 846.5394 | 845.5554 | L | 260.1969 | 243.1703 | | 2 |
| 9 | | | | K | 147.1128 | 130.0863 | | 1 |

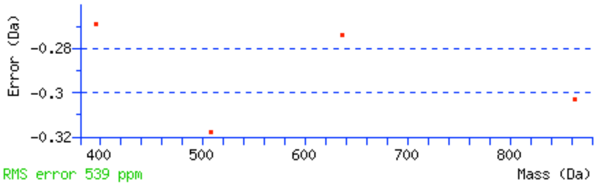


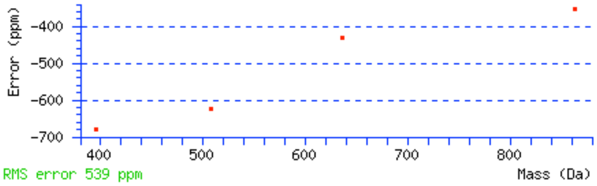


### 22. Jen1 (1 peptide in 250mM fraction2)

MS/MS Fragmentation of **SSSITDEK**
Found in **P36035** in **Uniprot_yeastS288C**, Carboxylic acid transporter protein homolog OS=Saccharomyces cerevisiae (strain ATCC 204508 / S288c) OX=559292 GN=JEN1 PE=1 SV=1

Match to Query 3601: 882.453845 from(883.461121,1+) intensity(481645.2812) rtinseconds(1949.973) index(5021)
Title: 20100530_methyl250mM_2.5589.5589.1
Data file D:\Panyue\Proteomic dataset\20100530_methyl250mM_2.RAW

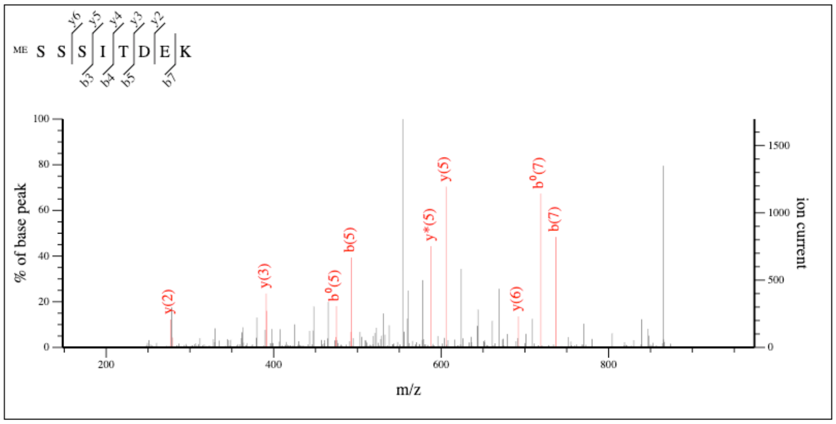

**Monoisotopic mass of neutral peptide Mr(calc):** 882.4374
**Fixed modifications:** Carbamidomethyl (C) (apply to specified residues or termini only)
**Variable modifications:**
**N-term :** Methyl:2H(3) (Protein N-term)
**Ions Score:** 30 **Expect:** 0.012
**Matches :** 15/34 fragment ions using 28 most intense peaks (help)

| # | b | $b^0$ | Seq. | y | y* | $y^0$ | # |
|---|---|---|---|---|---|---|---|
| 1 | 105.0738 | 87.0632 | S | | | | 8 |
| 2 | 192.1058 | 174.0952 | S | 779.3781 | 762.3516 | 761.3676 | 7 |
| 3 | 279.1378 | 261.1273 | S | 692.3461 | 675.3196 | 674.3355 | 6 |
| 4 | 392.2219 | 374.2113 | I | 605.3141 | 588.2875 | 587.3035 | 5 |
| 5 | 493.2696 | 475.2590 | T | 492.2300 | 475.2035 | 474.2195 | 4 |
| 6 | 608.2965 | 590.2860 | D | 391.1823 | 374.1558 | 373.1718 | 3 |
| 7 | 737.3391 | 719.3286 | E | 276.1554 | 259.1288 | 258.1448 | 2 |
| 8 | | | K | 147.1128 | 130.0863 | | 1 |

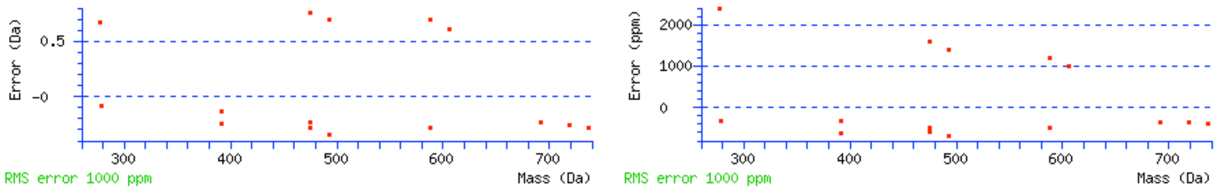


### 23. Eft1 (1 peptide in 250mM fraction2)

MS/MS Fragmentation of **VAFTVDQMRSLMDK**
Found in **P32324** in **Uniprot_yeastS288C**, Elongation factor 2 OS=Saccharomyces cerevisiae (strain ATCC 204508 / S288c) OX=559292 GN=EFT1 PE=1 SV=1

Match to Query 9765: 1706.871434 from(854.442993,2+) intensity(1598736.1250) rtinseconds(2677.8771) index(7003)
Title: 20100530_methyl250mM_2.7794.7794.2
Data file D:\Panyue\Proteomic dataset\20100530_methyl250mM_2.RAW

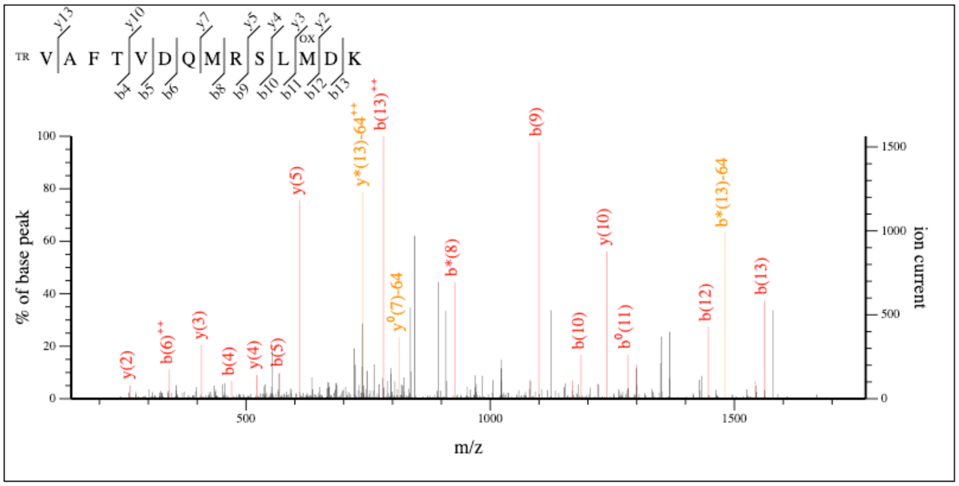

**Monoisotopic mass of neutral peptide Mr(calc):** 1706.9046
**Fixed modifications:** Carbamidomethyl (C) (apply to specified residues or termini only)
**Variable modifications:**
**N-term :** Trimethyl:2H(9) (Protein N-term)
**M12** : Oxidation (M), with neutral losses 0.0000(shown in table), 63.9983
**Ions Score:** 28 **Expect:** 0.027
**Matches :** 27/214 fragment ions using 43 most intense peaks (help)

| # | b | b++ | b* | b*++ | b0 | b0++ | Seq. | y | y++ | y* | y*++ | y0 | y0++ | # |
|---|---|---|---|---|---|---|---|---|---|---|---|---|---|---|
| 1 | 151.1791 | 76.0932 | | | | | V | | | | | | | 14 |
| 2 | 222.2162 | 111.6118 | | | | | A | 1557.7400 | 779.3736 | 1540.7134 | 770.8604 | 1539.7294 | 770.3683 | 13 |
| 3 | 369.2847 | 185.1460 | | | | | F | 1486.7029 | 743.8551 | 1469.6763 | 735.3418 | 1468.6923 | 734.8498 | 12 |
| 4 | 470.3323 | 235.6698 | | | 452.3218 | 226.6645 | T | 1339.6344 | 670.3209 | 1322.6079 | 661.8076 | 1321.6239 | 661.3156 | 11 |
| 5 | 569.4008 | 285.2040 | | | 551.3902 | 276.1987 | V | 1238.5868 | 619.7970 | 1221.5602 | 611.2837 | 1220.5762 | 610.7917 | 10 |
| 6 | 684.4277 | 342.7175 | | | 666.4171 | 333.7122 | D | 1139.5184 | 570.2628 | 1122.4918 | 561.7495 | 1121.5078 | 561.2575 | 9 |
| 7 | 812.4863 | 406.7468 | 795.4597 | 398.2335 | 794.4757 | 397.7415 | Q | 1024.4914 | 512.7493 | 1007.4649 | 504.2361 | 1006.4808 | 503.7441 | 8 |
| 8 | 943.5268 | 472.2670 | 926.5002 | 463.7537 | 925.5162 | 463.2617 | M | 896.4328 | 448.7201 | 879.4063 | 440.2068 | 878.4223 | 439.7148 | 7 |
| 9 | 1099.6279 | 550.3176 | 1082.6013 | 541.8043 | 1081.6173 | 541.3123 | R | 765.3924 | 383.1998 | 748.3658 | 374.6865 | 747.3818 | 374.1945 | 6 |
| 10 | 1186.6599 | 593.8336 | 1169.6333 | 585.3203 | 1168.6493 | 584.8283 | S | 609.2912 | 305.1493 | 592.2647 | 296.6360 | 591.2807 | 296.1440 | 5 |
| 11 | 1299.7440 | 650.3756 | 1282.7174 | 641.8623 | 1281.7334 | 641.3703 | L | 522.2592 | 261.6332 | 505.2327 | 253.1200 | 504.2486 | 252.6280 | 4 |
| 12 | 1446.7794 | 723.8933 | 1429.7528 | 715.3800 | 1428.7688 | 714.8880 | M | 409.1751 | 205.0912 | 392.1486 | 196.5779 | 391.1646 | 196.0859 | 3 |
| 13 | 1561.8063 | 781.4068 | 1544.7798 | 772.8935 | 1543.7957 | 772.4015 | D | 262.1397 | 131.5735 | 245.1132 | 123.0602 | 244.1292 | 122.5682 | 2 |
| 14 | | | | | | | K | 147.1128 | 74.0600 | 130.0863 | 65.5468 | | | 1 |

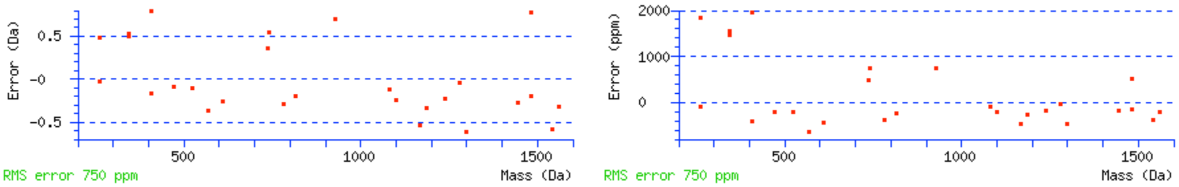


### 24. Pdc6 (2 peptides in 250mM fraction 3)

MS/MS Fragmentation of SEITLGK
Found in P26263 in Uniprot_yeastS288C, Pyruvate decarboxylase isozyme 3 OS=Saccharomyces cerevisiae (strain ATCC 204508 / S288c) OX=559292 GN=PDC6 PE=1 SV=3

Match to Query 2334: 780.449877 from(781.457153,1+) intensity(667905.8750) rtinseconds(1062.2886) index(2616)
Title: 20100530_methyl250mM_3.2911.2911.1
Data file D:\Panyue\Proteomic dataset\20100530_methyl250mM_3.RAW

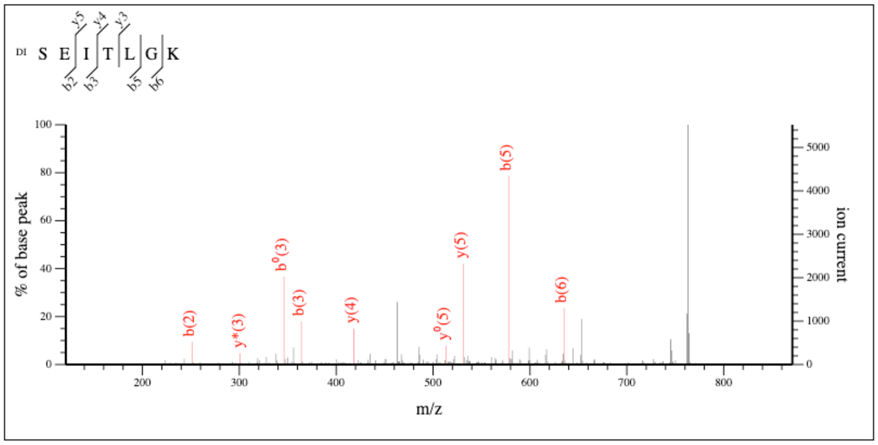

**Monoisotopic mass of neutral peptide Mr(calc):** 780.4864
**Fixed modifications:** Carbamidomethyl (C) (apply to specified residues or termini only)
**Variable modifications:**
**N-term :** Dimethyl:2H(6) (Protein N-term)
**Ions Score:** 25 **Expect:** 0.042
**Matches :** 9/27 fragment ions using 18 most intense peaks (help)

| # | b | $b^0$ | Seq. | y | y* | $y^0$ | # |
|---|---|---|---|---|---|---|---|
| 1 | 122.1083 | 104.0977 | S | | | | 7 |
| 2 | 251.1509 | 233.1403 | E | 660.3927 | 643.3661 | 642.3821 | 6 |
| 3 | 364.2349 | 346.2244 | I | 531.3501 | 514.3235 | 513.3395 | 5 |
| 4 | 465.2826 | 447.2720 | T | 418.2660 | 401.2395 | 400.2554 | 4 |
| 5 | 578.3667 | 560.3561 | L | 317.2183 | 300.1918 | | 3 |
| 6 | 635.3881 | 617.3776 | G | 204.1343 | 187.1077 | | 2 |
| 7 | | | K | 147.1128 | 130.0863 | | 1 |

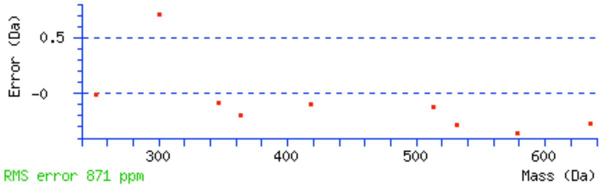


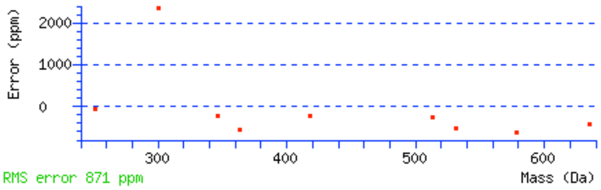


MS/MS Fragmentation of **SEITLGK**
Found in **P26263** in **Uniprot_yeastS288C**, Pyruvate decarboxylase isozyme 3 OS=Saccharomyces cerevisiae (strain ATCC 204508 / S288c) OX=559292 GN=PDC6 PE=1 SV=3

Match to Query 2335: 780.450182 from(781.457458,1+) intensity(291644.1250) rtinseconds(1059.8585) index(2610)
Title: 20100530_methyl250mM_3.2904.2904.1
Data file D:\Panyue\Proteomic dataset\20100530_methyl250mM_3.RAW

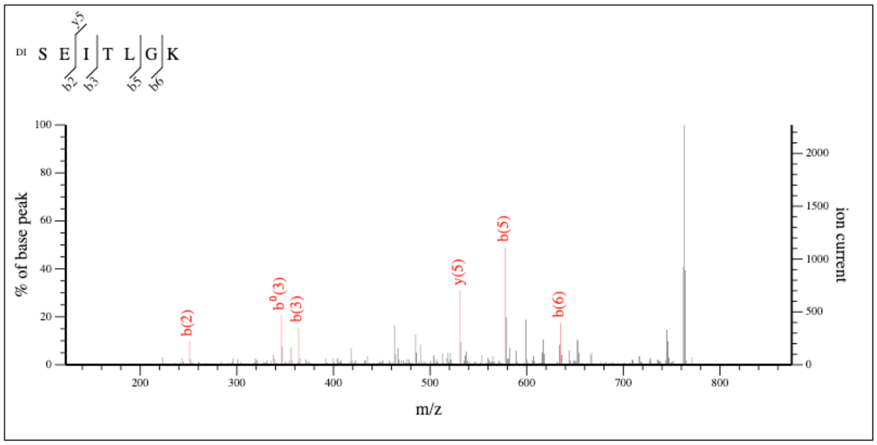

**Monoisotopic mass of neutral peptide Mr(calc):** 780.4864
**Fixed modifications:** Carbamidomethyl (C) (apply to specified residues or termini only)
**Variable modifications:**
**N-term :** Dimethyl:2H(6) (Protein N-term)
**Ions Score:** 22 **Expect:** 0.078
**Matches :** 6/27 fragment ions using 12 most intense peaks (help)

| # | b | $b^0$ | Seq. | y | y* | $y^0$ | # |
|---|---|---|---|---|---|---|---|
| 1 | 122.1083 | 104.0977 | S | | | | 7 |
| 2 | 251.1509 | 233.1403 | E | 660.3927 | 643.3661 | 642.3821 | 6 |
| 3 | 364.2349 | 346.2244 | I | 531.3501 | 514.3235 | 513.3395 | 5 |
| 4 | 465.2826 | 447.2720 | T | 418.2660 | 401.2395 | 400.2554 | 4 |
| 5 | 578.3667 | 560.3561 | L | 317.2183 | 300.1918 | | 3 |
| 6 | 635.3881 | 617.3776 | G | 204.1343 | 187.1077 | | 2 |
| 7 | | | K | 147.1128 | 130.0863 | | 1 |

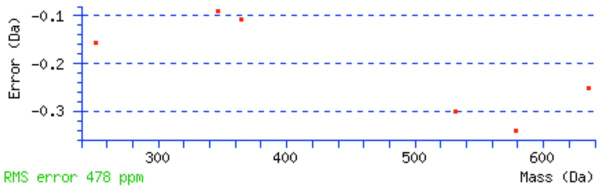


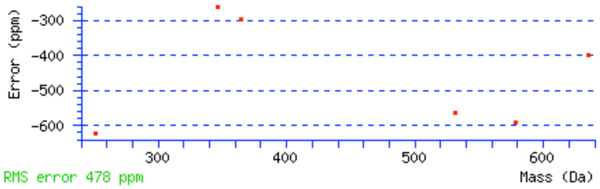


### 25. Pdc5 (2 peptides in 250mM fraction 3)

MS/MS Fragmentation of **SEITLGK**
Found in **P16467** in **Uniprot_yeastS288C**, Pyruvate decarboxylase isozyme 2 OS=Saccharomyces cerevisiae (strain ATCC 204508 / S288c) OX=559292 GN=PDC5 PE=1 SV=4

Match to Query 2334: 780.449877 from(781.457153,1+) intensity(667905.8750) rtinseconds(1062.2886) index(2616)
Title: 20100530_methyl250mM_3.2911.2911.1
Data file D:\Panyue\Proteomic dataset\20100530_methyl250mM_3.RAW

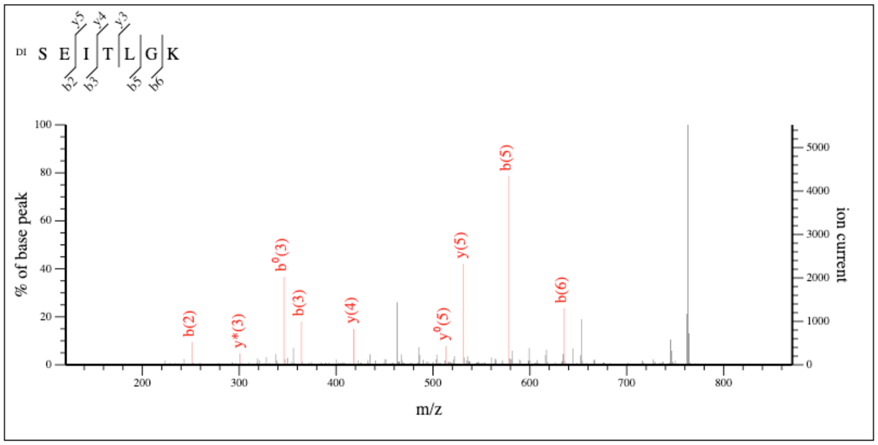

**Monoisotopic mass of neutral peptide Mr(calc):** 780.4864
**Fixed modifications:** Carbamidomethyl (C) (apply to specified residues or termini only)
**Variable modifications:**
**N-term :** Dimethyl:2H(6) (Protein N-term)
**Ions Score:** 25 **Expect:** 0.042
**Matches :** 9/27 fragment ions using 18 most intense peaks (help)

| # | b | $b^0$ | Seq. | y | y* | $y^0$ | # |
|---|---|---|---|---|---|---|---|
| 1 | 122.1083 | 104.0977 | S | | | | 7 |
| 2 | *251.1509* | 233.1403 | E | 660.3927 | 643.3661 | 642.3821 | 6 |
| 3 | *364.2349* | 346.2244 | I | **531.3501** | 514.3235 | 513.3395 | 5 |
| 4 | 465.2826 | 447.2720 | T | **418.2660** | 401.2395 | 400.2554 | 4 |
| 5 | *578.3667* | 560.3561 | L | 317.2183 | 300.1918 | | 3 |
| 6 | *635.3881* | 617.3776 | G | 204.1343 | 187.1077 | | 2 |
| 7 | | | K | 147.1128 | 130.0863 | | 1 |

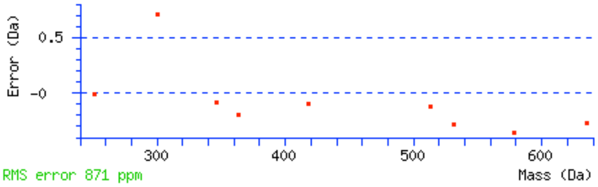


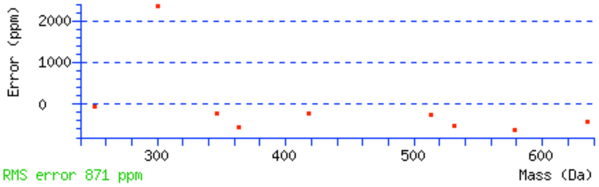


MS/MS Fragmentation of **SEITLGK**
Found in **P16467** in **Uniprot_yeastS288C**, Pyruvate decarboxylase isozyme 2 OS=Saccharomyces cerevisiae (strain ATCC 204508 / S288c) OX=559292 GN=PDC5 PE=1 SV=4

Match to Query 2335: 780.450182 from(781.457458,1+) intensity(291644.1250) rtinseconds(1059.8585) index(2610)
Title: 20100530_methyl250mM_3.2904.2904.1
Data file D:\Panyue\Proteomic dataset\20100530_methyl250mM_3.RAW

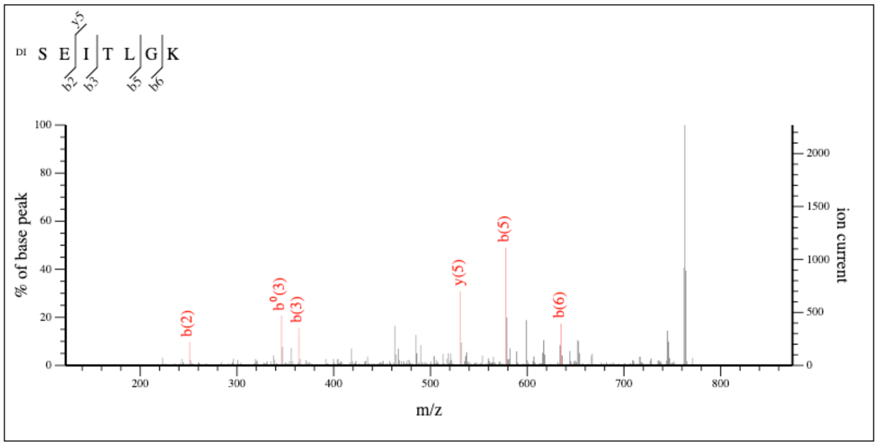

**Monoisotopic mass of neutral peptide Mr(calc):** 780.4864
**Fixed modifications:** Carbamidomethyl (C) (apply to specified residues or termini only)
**Variable modifications:**
**N-term :** Dimethyl:2H(6) (Protein N-term)
**Ions Score:** 22 **Expect:** 0.078
**Matches :** 6/27 fragment ions using 12 most intense peaks (help)

| # | b | $b^0$ | Seq. | y | y* | $y^0$ | # |
|---|---|---|---|---|---|---|---|
| 1 | 122.1083 | 104.0977 | S | | | | 7 |
| 2 | ***251.1509*** | 233.1403 | E | 660.3927 | 643.3661 | 642.3821 | 6 |
| 3 | ***364.2349*** | 346.2244 | I | 531.3501 | 514.3235 | 513.3395 | 5 |
| 4 | 465.2826 | 447.2720 | T | 418.2660 | 401.2395 | 400.2554 | 4 |
| 5 | ***578.3667*** | 560.3561 | L | 317.2183 | 300.1918 | | 3 |
| 6 | ***635.3881*** | 617.3776 | G | 204.1343 | 187.1077 | | 2 |
| 7 | | | K | 147.1128 | 130.0863 | | 1 |

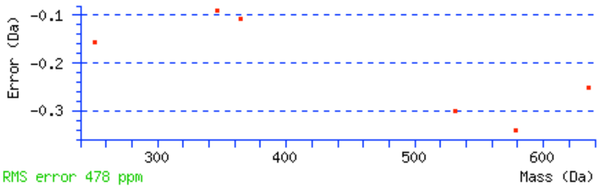


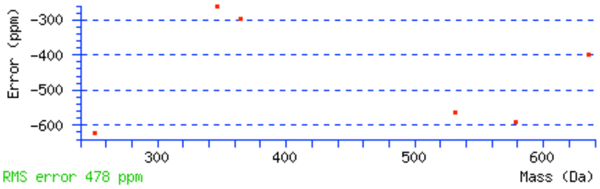


### 26. Pdc1 (2 peptides in 250mM fraction 3)

MS/MS Fragmentation of **SEITLGK**
Found in **P06169** in **Uniprot_yeastS288C**, Pyruvate decarboxylase isozyme 1 OS=Saccharomyces cerevisiae (strain ATCC 204508 / S288c) OX=559292 GN=PDC1 PE=1 SV=7

Match to Query 2335: 780.450182 from(781.457458,1+) intensity(291644.1250) rtinseconds(1059.8585) index(2610)
Title: 20100530_methyl250mM_3.2904.2904.1
Data file D:\Panyue\Proteomic dataset\20100530_methyl250mM_3.RAW

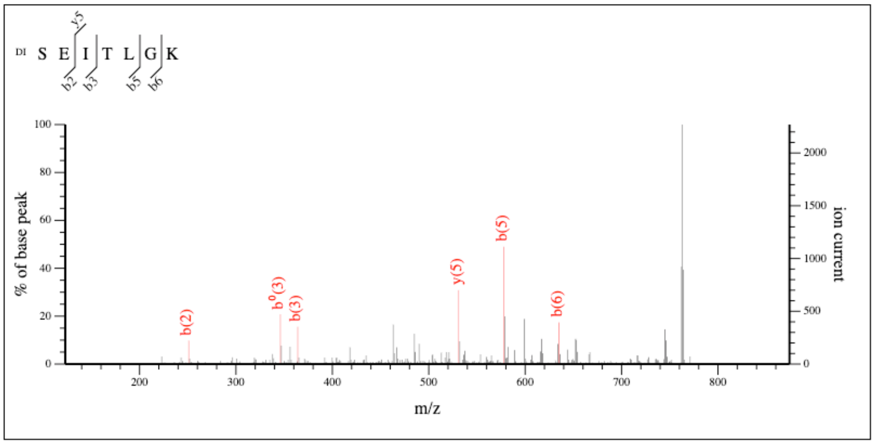

**Monoisotopic mass of neutral peptide Mr(calc):** 780.4864
**Fixed modifications:** Carbamidomethyl (C) (apply to specified residues or termini only)
**Variable modifications:**
**N-term :** Dimethyl:2H(6) (Protein N-term)
**Ions Score:** 22 **Expect:** 0.078
**Matches :** 6/27 fragment ions using 12 most intense peaks (help)

| # | b | $b^0$ | Seq. | y | y* | $y^0$ | # |
|---|---|---|---|---|---|---|---|
| 1 | 122.1083 | 104.0977 | S | | | | 7 |
| 2 | ***251.1509*** | 233.1403 | E | 660.3927 | 643.3661 | 642.3821 | 6 |
| 3 | ***364.2349*** | 346.2244 | I | 531.3501 | 514.3235 | 513.3395 | 5 |
| 4 | 465.2826 | 447.2720 | T | 418.2660 | 401.2395 | 400.2554 | 4 |
| 5 | ***578.3667*** | 560.3561 | L | 317.2183 | 300.1918 | | 3 |
| 6 | ***635.3881*** | 617.3776 | G | 204.1343 | 187.1077 | | 2 |
| 7 | | | K | 147.1128 | 130.0863 | | 1 |

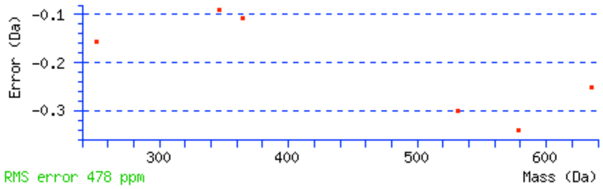


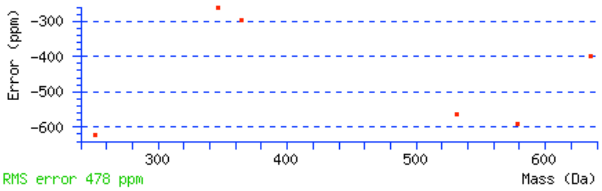


MS/MS Fragmentation of **SEITLGK**
Found in **P06169** in **Uniprot_yeastS288C**, Pyruvate decarboxylase isozyme 1 OS=Saccharomyces cerevisiae (strain ATCC 204508 / S288c) OX=559292 GN=PDC1 PE=1 SV=7

Match to Query 2334: 780.449877 from(781.457153,1+) intensity(667905.8750) rtinseconds(1062.2886) index(2616)
Title: 20100530_methyl250mM_3.2911.2911.1
Data file D:\Panyue\Proteomic dataset\20100530_methyl250mM_3.RAW

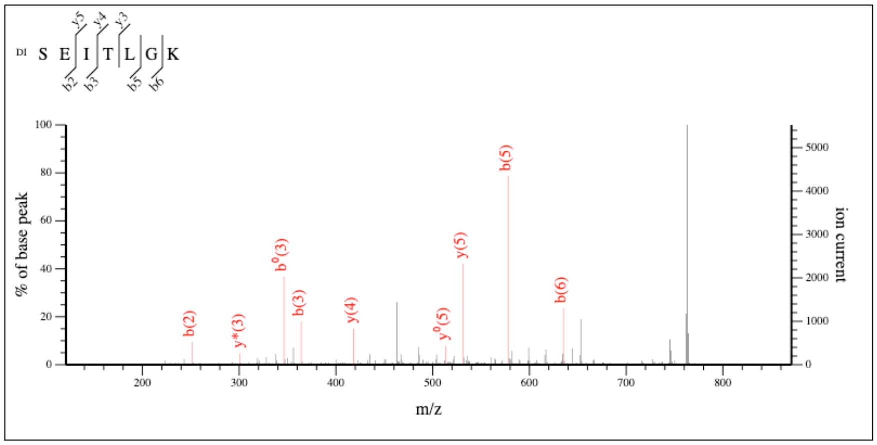

**Monoisotopic mass of neutral peptide Mr(calc):** 780.4864
**Fixed modifications:** Carbamidomethyl (C) (apply to specified residues or termini only)
**Variable modifications:**
**N-term :** Dimethyl:2H(6) (Protein N-term)
**Ions Score:** 25 **Expect:** 0.042
**Matches :** 9/27 fragment ions using 18 most intense peaks (help)

| # | b | b0 | Seq. | y | y* | y0 | # |
|---|---|---|---|---|---|---|---|
| 1 | 122.1083 | 104.0977 | S | | | | 7 |
| 2 | *251.1509* | 233.1403 | E | 660.3927 | 643.3661 | 642.3821 | 6 |
| 3 | *364.2349* | 346.2244 | I | **531.3501** | 514.3235 | 513.3395 | 5 |
| 4 | 465.2826 | 447.2720 | T | **418.2660** | 401.2395 | 400.2554 | 4 |
| 5 | *578.3667* | 560.3561 | L | 317.2183 | 300.1918 | | 3 |
| 6 | *635.3881* | 617.3776 | G | 204.1343 | 187.1077 | | 2 |
| 7 | | | K | 147.1128 | 130.0863 | | 1 |

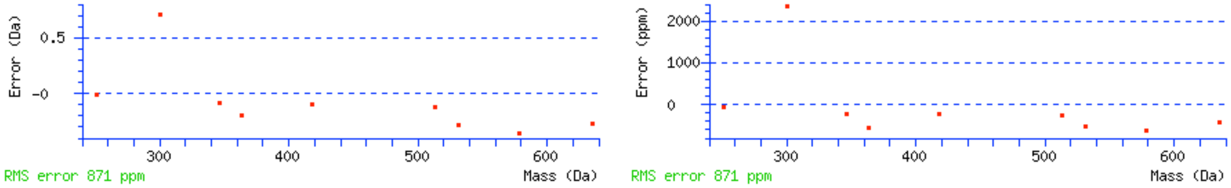


### 27. Nop14 (2 peptides in 250mM fraction 3)

MS/MS Fragmentation of **AGSQLKNLK**
Found in **Q99207** in **Uniprot_yeastS288C**, Nucleolar complex protein 14 OS=Saccharomyces cerevisiae (strain ATCC 204508 / S288c) OX=559292 GN=NOP14 PE=1 SV=1

Match to Query 4755: 1008.669787 from(1009.677063,1+) intensity(593260.6250) rtinseconds(1660.6244) index(4253)
Title: 20100530_methyl250mM_3.4733.4733.1
Data file D:\Panyue\Proteomic dataset\20100530_methyl250mM_3.RAW

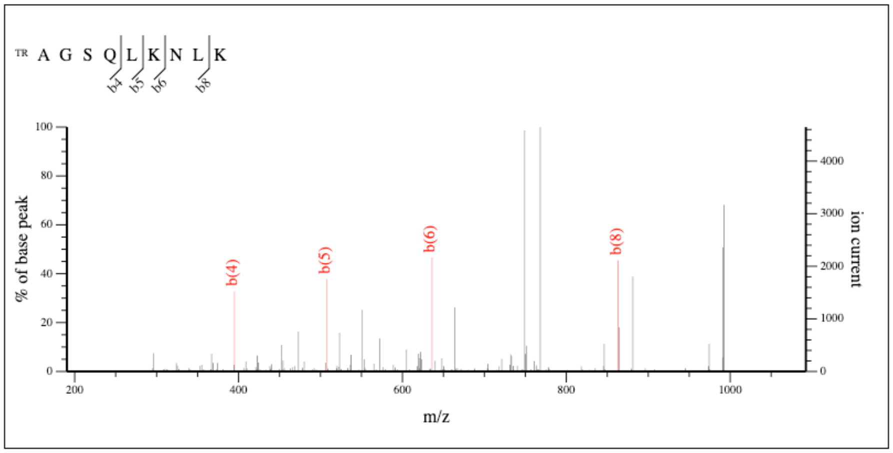

**Monoisotopic mass of neutral peptide Mr(calc):** 1008.6641
**Fixed modifications:** Carbamidomethyl (C) (apply to specified residues or termini only)
**Variable modifications:**
**N-term :** Trimethyl:2H(9) (Protein N-term)
**Ions Score:** 27 **Expect:** 0.0039
**Matches :** 4/37 fragment ions using 8 most intense peaks (help)

| # | b | b* | b$^0$ | Seq. | y | y* | y$^0$ | # |
|---|---|---|---|---|---|---|---|---|
| 1 | 123.1478 | | | A | | | | 9 |
| 2 | 180.1693 | | | G | 887.5309 | 870.5043 | 869.5203 | 8 |
| 3 | 267.2013 | | 249.1908 | S | 830.5094 | 813.4829 | 812.4989 | 7 |
| 4 | *395.2599* | 378.2334 | 377.2493 | Q | 743.4774 | 726.4509 | | 6 |
| 5 | *508.3440* | 491.3174 | 490.3334 | L | 615.4188 | 598.3923 | | 5 |
| 6 | *636.4389* | 619.4124 | 618.4284 | K | 502.3348 | 485.3082 | | 4 |
| 7 | 750.4819 | 733.4553 | 732.4713 | N | 374.2398 | 357.2132 | | 3 |
| 8 | *863.5659* | 846.5394 | 845.5554 | L | 260.1969 | 243.1703 | | 2 |
| 9 | | | | K | 147.1128 | 130.0863 | | 1 |

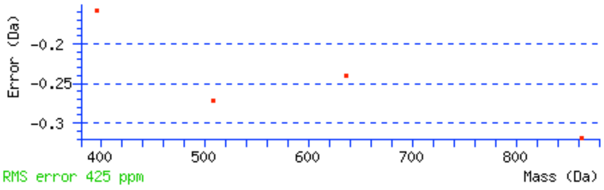


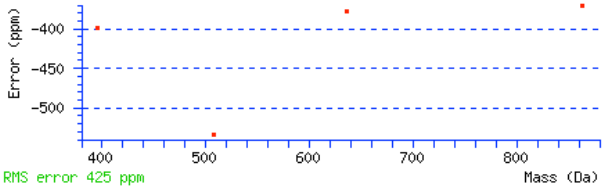


MS/MS Fragmentation of **AGSQLKNLK**
Found in **Q99207** in **Uniprot_yeastS288C**, Nucleolar complex protein 14 OS=Saccharomyces cerevisiae (strain ATCC 204508 / S288c) OX=559292 GN=NOP14 PE=1 SV=1

Match to Query 4756: 1008.669787 from(1009.677063,1+) intensity(1274352.8750) rtinseconds(1663.8592) index(4262)
Title: 20100530_methyl250mM_3.4743.4743.1
Data file D:\Panyue\Proteomic dataset\20100530_methyl250mM_3.RAW

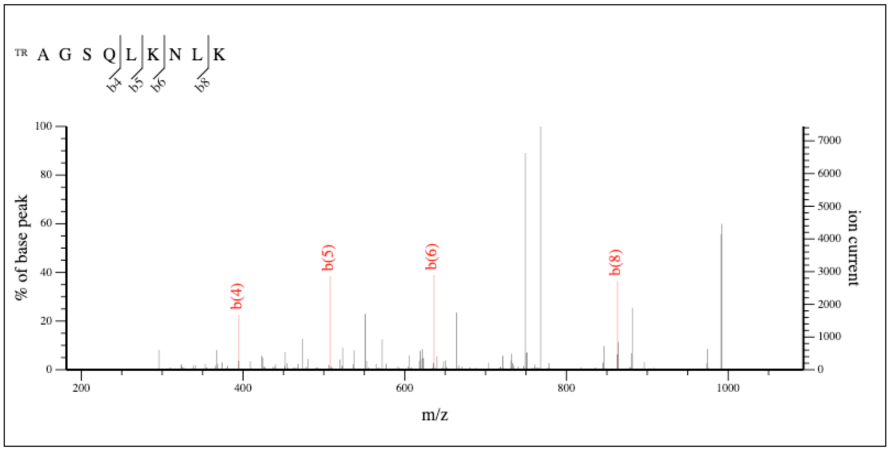

**Monoisotopic mass of neutral peptide Mr(calc):** 1008.6641
**Fixed modifications:** Carbamidomethyl (C) (apply to specified residues or termini only)
**Variable modifications:**
**N-term :** Trimethyl:2H(9) (Protein N-term)
**Ions Score:** 27 **Expect:** 0.0041
**Matches :** 4/37 fragment ions using 8 most intense peaks (help)

| # | b | b* | b⁰ | Seq. | y | y* | y⁰ | # |
|---|---|---|---|---|---|---|---|---|
| 1 | 123.1478 | | | A | | | | 9 |
| 2 | 180.1693 | | | G | 887.5309 | 870.5043 | 869.5203 | 8 |
| 3 | 267.2013 | | 249.1908 | S | 830.5094 | 813.4829 | 812.4989 | 7 |
| 4 | ***395.2599*** | 378.2334 | 377.2493 | Q | 743.4774 | 726.4509 | | 6 |
| 5 | ***508.3440*** | 491.3174 | 490.3334 | L | 615.4188 | 598.3923 | | 5 |
| 6 | ***636.4389*** | 619.4124 | 618.4284 | K | 502.3348 | 485.3082 | | 4 |
| 7 | 750.4819 | 733.4553 | 732.4713 | N | 374.2398 | 357.2132 | | 3 |
| 8 | ***863.5659*** | 846.5394 | 845.5554 | L | 260.1969 | 243.1703 | | 2 |
| 9 | | | | K | 147.1128 | 130.0863 | | 1 |

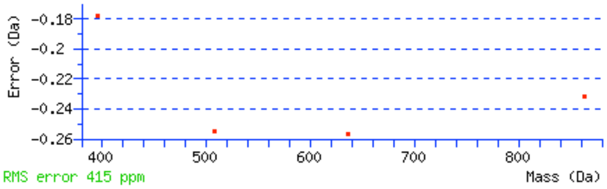


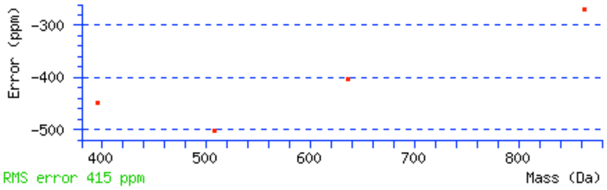


### 28. Eft1 (2 peptides in 250mM fraction 3)

MS/MS Fragmentation of **VAFTVDQMRSLMDK**
Found in **P32324** in **Uniprot_yeastS288C**, Elongation factor 2 OS=Saccharomyces cerevisiae (strain ATCC 204508 / S288c) OX=559292 GN=EFT1 PE=1 SV=1

Match to Query 9780: 1706.865448 from(854.440000,2+) intensity(2183071.5000) rtinseconds(2698.8157) index(7096)
Title: 20100530_methyl250mM_3.7895.7895.2
Data file D:\Panyue\Proteomic dataset\20100530_methyl250mM_3.RAW

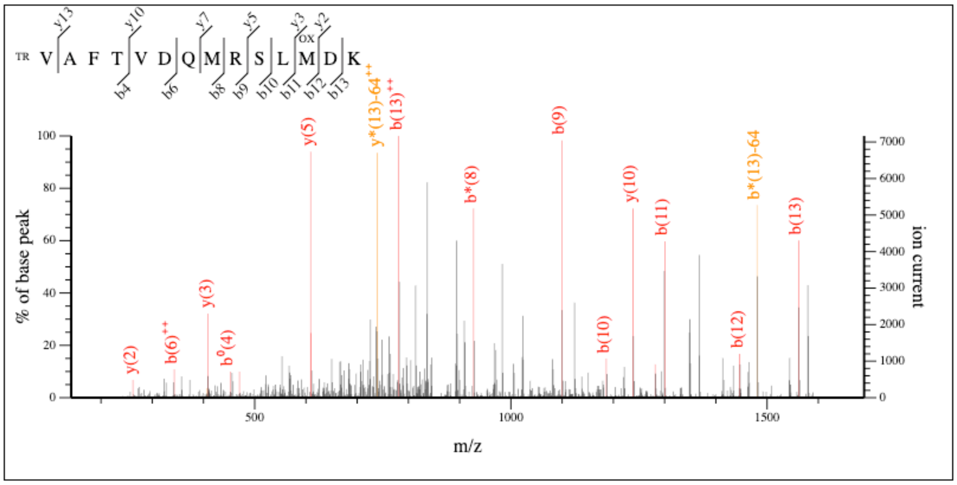

**Monoisotopic mass of neutral peptide Mr(calc):** 1706.9046
**Fixed modifications:** Carbamidomethyl (C) (apply to specified residues or termini only)
**Variable modifications:**
**N-term :** Trimethyl:2H(9) (Protein N-term)
**M12** : Oxidation (M), with neutral losses 0.0000(shown in table), 63.9983
**Ions Score:** 21 **Expect:** 0.14
**Matches :** 23/214 fragment ions using 42 most intense peaks (help)

| # | b | b++ | b* | b*++ | b0 | b0++ | Seq. | y | y++ | y* | y*++ | y0 | y0++ | # |
|---|---|---|---|---|---|---|---|---|---|---|---|---|---|---|
| 1 | 151.1791 | 76.0932 | | | | | V | | | | | | | 14 |
| 2 | 222.2162 | 111.6118 | | | | | A | 1557.7400 | 779.3736 | 1540.7134 | 770.8604 | 1539.7294 | 770.3683 | 13 |
| 3 | 369.2847 | 185.1460 | | | | | F | 1486.7029 | 743.8551 | 1469.6763 | 735.3418 | 1468.6923 | 734.8498 | 12 |
| 4 | ***470.3323*** | 235.6698 | | | 452.3218 | 226.6645 | T | 1339.6344 | 670.3209 | 1322.6079 | 661.8076 | 1321.6239 | 661.3156 | 11 |
| 5 | 569.4008 | 285.2040 | | | 551.3902 | 276.1987 | V | **1238.5868** | 619.7970 | 1221.5602 | 611.2837 | 1220.5762 | 610.7917 | 10 |
| 6 | 684.4277 | 342.7175 | | | 666.4171 | 333.7122 | D | 1139.5184 | 570.2628 | 1122.4918 | 561.7495 | 1121.5078 | 561.2575 | 9 |
| 7 | 812.4863 | 406.7468 | 795.4597 | 398.2335 | 794.4757 | 397.7415 | Q | 1024.4914 | 512.7493 | 1007.4649 | 504.2361 | 1006.4808 | 503.7441 | 8 |
| 8 | 943.5268 | 472.2670 | 926.5002 | 463.7537 | 925.5162 | 463.2617 | M | 896.4328 | 448.7201 | 879.4063 | 440.2068 | 878.4223 | 439.7148 | 7 |
| 9 | ***1099.6279*** | 550.3176 | 1082.6013 | 541.8043 | 1081.6173 | 541.3123 | R | 765.3924 | 383.1998 | 748.3658 | 374.6865 | 747.3818 | 374.1945 | 6 |
| 10 | ***1186.6599*** | 593.8336 | 1169.6333 | 585.3203 | 1168.6493 | 584.8283 | S | **609.2912** | 305.1493 | 592.2647 | 296.6360 | 591.2807 | 296.1440 | 5 |
| 11 | ***1299.7440*** | 650.3756 | 1282.7174 | 641.8623 | 1281.7334 | 641.3703 | L | 522.2592 | 261.6332 | 505.2327 | 253.1200 | 504.2486 | 252.6280 | 4 |
| 12 | ***1446.7794*** | 723.8933 | 1429.7528 | 715.3800 | 1428.7688 | 714.8880 | M | **409.1751** | 205.0912 | 392.1486 | 196.5779 | 391.1646 | 196.0859 | 3 |
| 13 | ***1561.8063*** | 781.4068 | 1544.7798 | 772.8935 | 1543.7957 | 772.4015 | D | **262.1397** | 131.5735 | 245.1132 | 123.0602 | 244.1292 | 122.5682 | 2 |
| 14 | | | | | | | K | 147.1128 | 74.0600 | 130.0863 | 65.5468 | | | 1 |

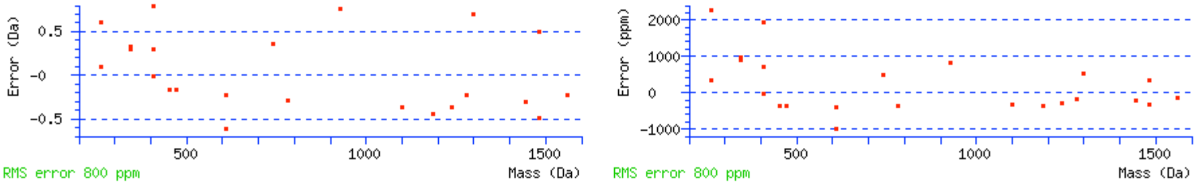


MS/MS Fragmentation of **VAFTVDQMRSLMDK**
Found in **P32324** in **Uniprot_yeastS288C**, Elongation factor 2 OS=Saccharomyces cerevisiae (strain ATCC 204508 / S288c) OX=559292 GN=EFT1 PE=1 SV=1

Match to Query 9781: 1706.872778 from(854.443665,2+) intensity(3745608.5000) rtinseconds(2701.7253) index(7105)
Title: 20100530_methyl250mM_3.7905.7905.2
Data file D:\Panyue\Proteomic dataset\20100530_methyl250mM_3.RAW

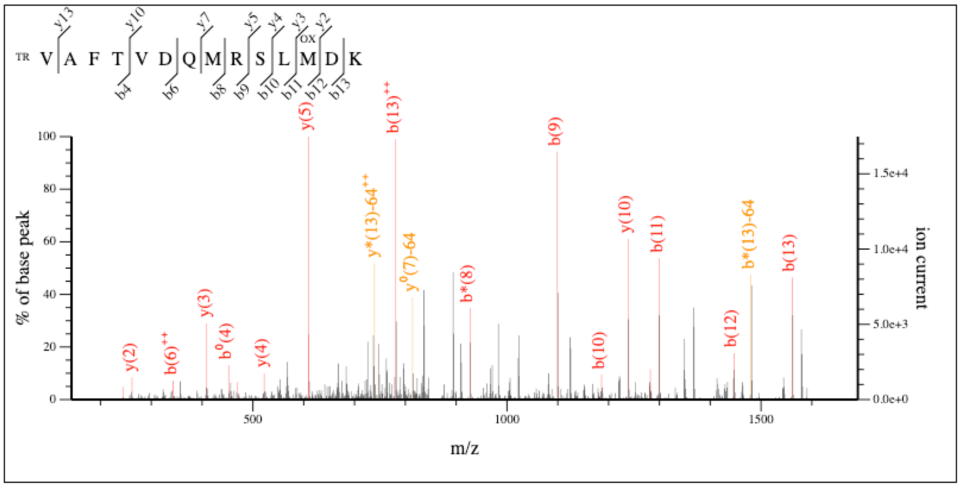

**Monoisotopic mass of neutral peptide Mr(calc):** 1706.9046
**Fixed modifications:** Carbamidomethyl (C) (apply to specified residues or termini only)
**Variable modifications:**
**N-term :** Trimethyl:2H(9) (Protein N-term)
**M12 :** Oxidation (M), with neutral losses 0.0000(shown in table), 63.9983
**Ions Score:** 23 **Expect:** 0.081
**Matches :** 25/214 fragment ions using 42 most intense peaks (help)

| # | b | b++ | b* | b*++ | b0 | b0++ | Seq. | y | y++ | y* | y*++ | y0 | y0++ | # |
|---|---|---|---|---|---|---|---|---|---|---|---|---|---|---|
| 1 | 151.1791 | 76.0932 | | | | | V | | | | | | | 14 |
| 2 | 222.2162 | 111.6118 | | | | | A | 1557.7400 | 779.3736 | 1540.7134 | 770.8604 | 1539.7294 | 770.3683 | 13 |
| 3 | 369.2847 | 185.1460 | | | | | F | 1486.7029 | 743.8551 | 1469.6763 | 735.3418 | 1468.6923 | 734.8498 | 12 |
| 4 | ***470.3323*** | 235.6698 | | | 452.3218 | 226.6645 | T | 1339.6344 | 670.3209 | 1322.6079 | 661.8076 | 1321.6239 | 661.3156 | 11 |
| 5 | 569.4008 | 285.2040 | | | 551.3902 | 276.1987 | V | **1238.5868** | 619.7970 | 1221.5602 | 611.2837 | 1220.5762 | 610.7917 | 10 |
| 6 | 684.4277 | 342.7175 | | | 666.4171 | 333.7122 | D | 1139.5184 | 570.2628 | 1122.4918 | 561.7495 | 1121.5078 | 561.2575 | 9 |
| 7 | 812.4863 | 406.7468 | 795.4597 | 398.2335 | 794.4757 | 397.7415 | Q | 1024.4914 | 512.7493 | 1007.4649 | 504.2361 | 1006.4808 | 503.7441 | 8 |
| 8 | 943.5268 | 472.2670 | 926.5002 | 463.7537 | 925.5162 | 463.2617 | M | 896.4328 | 448.7201 | 879.4063 | 440.2068 | 878.4223 | 439.7148 | 7 |
| 9 | ***1099.6279*** | 550.3176 | 1082.6013 | 541.8043 | 1081.6173 | 541.3123 | R | 765.3924 | 383.1998 | 748.3658 | 374.6865 | 747.3818 | 374.1945 | 6 |
| 10 | ***1186.6599*** | 593.8336 | 1169.6333 | 585.3203 | 1168.6493 | 584.8283 | S | **609.2912** | 305.1493 | 592.2647 | 296.6360 | 591.2807 | 296.1440 | 5 |
| 11 | ***1299.7440*** | 650.3756 | 1282.7174 | 641.8623 | 1281.7334 | 641.3703 | L | **522.2592** | 261.6332 | 505.2327 | 253.1200 | 504.2486 | 252.6280 | 4 |
| 12 | ***1446.7794*** | 723.8933 | 1429.7528 | 715.3800 | 1428.7688 | 714.8880 | M | **409.1751** | 205.0912 | 392.1486 | 196.5779 | 391.1646 | 196.0859 | 3 |
| 13 | ***1561.8063*** | 781.4068 | 1544.7798 | 772.8935 | 1543.7957 | 772.4015 | D | **262.1397** | 131.5735 | 245.1132 | 123.0602 | 244.1292 | 122.5682 | 2 |
| 14 | | | | | | | K | 147.1128 | 74.0600 | 130.0863 | 65.5468 | | | 1 |

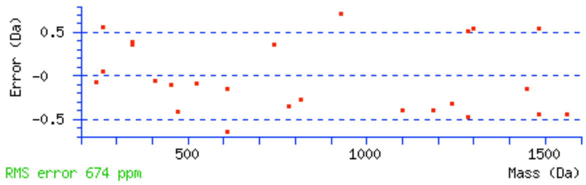


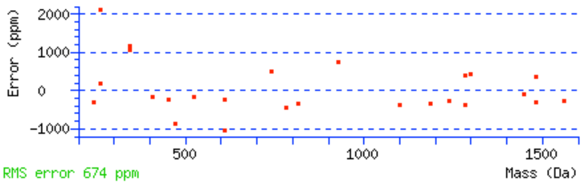


### 29. Mon2 (6 peptides in 300mM fraction 1)

MS/MS Fragmentation of **AMNTGGFDSMQRQLEAELRSLSSESK**
Found in **P48563** in **Uniprot_yeastS288C**, Protein MON2 OS=Saccharomyces cerevisiae (strain ATCC 204508 / S288c) OX=559292 GN=MON2 PE=1 SV=1

Match to Query 16252: 2938.443198 from(980.488342,3+) intensity(483540.4375) rtinseconds(6826.1792) index(17421)
Title: 20100530_methyl300mM_1.19379.19379.3
Data file D:\Panyue\Proteomic dataset\20100530_methyl300mM_1.RAW

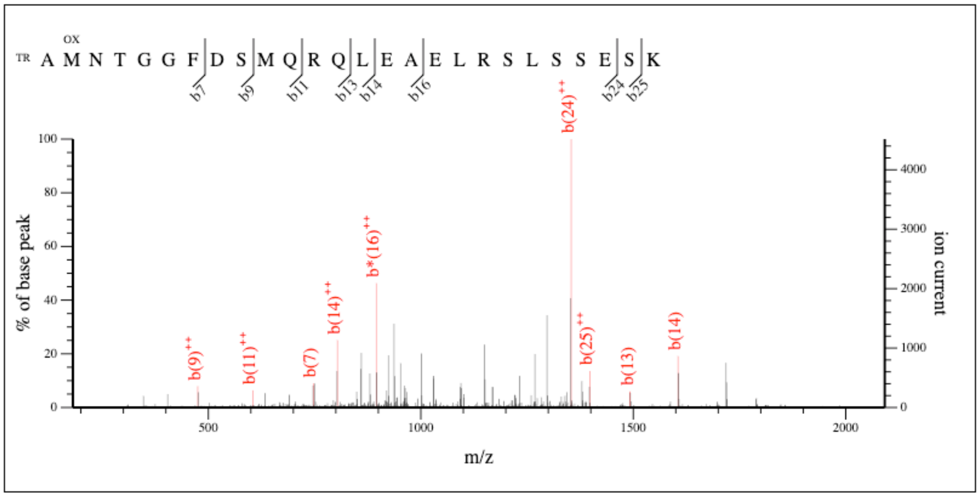

**Monoisotopic mass of neutral peptide Mr(calc):** 2938.4474
**Fixed modifications:** Carbamidomethyl (C) (apply to specified residues or termini only)
**Variable modifications:**
**N-term :** Trimethyl:2H(9) (Protein N-term)
**M2 :** Oxidation (M), with neutral losses 0.0000(shown in table), 63.9983
**Ions Score:** 22 **Expect:** 0.066
**Matches :** 11/432 fragment ions using 18 most intense peaks (help)

| # | b | $b^{++}$ | b* | $b^{*++}$ | $b^{0}$ | $b^{0++}$ | Seq. | y | $y^{++}$ | y* | $y^{*++}$ | $y^{0}$ | $y^{0++}$ | # |
|---|---|---|---|---|---|---|---|---|---|---|---|---|---|---|
| 1 | 123.1478 | 62.0776 | | | | | A | | | | | | | 26 |
| 2 | 270.1832 | 135.5953 | | | | | M | 2817.3142 | 1409.1607 | 2800.2876 | 1400.6475 | 2799.3036 | 1400.1554 | 25 |
| 3 | 384.2262 | 192.6167 | 367.1996 | 184.1034 | | | N | 2670.2788 | 1335.6430 | 2653.2522 | 1327.1298 | 2652.2682 | 1326.6377 | 24 |
| 4 | 485.2738 | 243.1406 | 468.2473 | 234.6273 | 467.2633 | 234.1353 | T | 2556.2359 | 1278.6216 | 2539.2093 | 1270.1083 | 2538.2253 | 1269.6163 | 23 |
| 5 | 542.2953 | 271.6513 | 525.2688 | 263.1380 | 524.2847 | 262.6460 | G | 2455.1882 | 1228.0977 | 2438.1616 | 1219.5845 | 2437.1776 | 1219.0924 | 22 |
| 6 | 599.3168 | 300.1620 | 582.2902 | 291.6487 | 581.3062 | 291.1567 | G | 2398.1667 | 1199.5870 | 2381.1402 | 1191.0737 | 2380.1561 | 1190.5817 | 21 |
| 7 | **746.3852** | 373.6962 | 729.3586 | 365.1830 | 728.3746 | 364.6909 | F | 2341.1452 | 1171.0763 | 2324.1187 | 1162.5630 | 2323.1347 | 1162.0710 | 20 |
| 8 | 861.4121 | 431.2097 | 844.3856 | 422.6964 | 843.4016 | 422.2044 | D | 2194.0768 | 1097.5421 | 2177.0503 | 1089.0288 | 2176.0663 | 1088.5368 | 19 |
| 9 | 948.4441 | ***474.7257*** | 931.4176 | 466.2124 | 930.4336 | 465.7204 | S | 2079.0499 | 1040.0286 | 2062.0233 | 1031.5153 | 2061.0393 | 1031.0233 | 18 |
| 10 | 1079.4846 | 540.2460 | 1062.4581 | 531.7327 | 1061.4741 | 531.2407 | M | 1992.0179 | 996.5126 | 1974.9913 | 987.9993 | 1974.0073 | 987.5073 | 17 |
| 11 | 1207.5432 | ***604.2752*** | 1190.5167 | 595.7620 | 1189.5326 | 595.2700 | Q | 1860.9774 | 930.9923 | 1843.9508 | 922.4791 | 1842.9668 | 921.9870 | 16 |
| 12 | 1363.6443 | 682.3258 | 1346.6178 | 673.8125 | 1345.6338 | 673.3205 | R | 1732.9188 | 866.9630 | 1715.8923 | 858.4498 | 1714.9082 | 857.9578 | 15 |
| 13 | **1491.7029** | ***746.3551*** | 1474.6764 | 737.8418 | 1473.6923 | 737.3498 | Q | 1576.8177 | 788.9125 | 1559.7911 | 780.3992 | 1558.8071 | 779.9072 | 14 |
| 14 | **1604.7870** | ***802.8971*** | 1587.7604 | 794.3838 | 1586.7764 | 793.8918 | L | 1448.7591 | 724.8832 | 1431.7326 | 716.3699 | 1430.7485 | 715.8779 | 13 |
| 15 | 1733.8296 | 867.4184 | 1716.8030 | 858.9051 | 1715.8190 | 858.4131 | E | 1335.6750 | 668.3412 | 1318.6485 | 659.8279 | 1317.6645 | 659.3359 | 12 |
| 16 | 1804.8667 | 902.9370 | 1787.8401 | 894.4237 | 1786.8561 | 893.9317 | A | 1206.6325 | 603.8199 | 1189.6059 | 595.3066 | 1188.6219 | 594.8146 | 11 |
| 17 | 1933.9093 | 967.4583 | 1916.8827 | 958.9450 | 1915.8987 | 958.4530 | E | 1135.5953 | 568.3013 | 1118.5688 | 559.7880 | 1117.5848 | 559.2960 | 10 |
| 18 | 2046.9933 | 1024.0003 | 2029.9668 | 1015.4870 | 2028.9828 | 1014.9950 | L | 1006.5527 | 503.7800 | 989.5262 | 495.2667 | 988.5422 | 494.7747 | 9 |
| 19 | 2203.0944 | 1102.0509 | 2186.0679 | 1093.5376 | 2185.0839 | 1093.0456 | R | 893.4687 | 447.2380 | 876.4421 | 438.7247 | 875.4581 | 438.2327 | 8 |
| 20 | 2290.1265 | 1145.5669 | 2273.0999 | 1137.0536 | 2272.1159 | 1136.5616 | S | 737.3676 | 369.1874 | 720.3410 | 360.6742 | 719.3570 | 360.1821 | 7 |
| 21 | 2403.2105 | 1202.1089 | 2386.1840 | 1193.5956 | 2385.2000 | 1193.1036 | L | 650.3355 | 325.6714 | 633.3090 | 317.1581 | 632.3250 | 316.6661 | 6 |
| 22 | 2490.2426 | 1245.6249 | 2473.2160 | 1237.1116 | 2472.2320 | 1236.6196 | S | 537.2515 | 269.1294 | 520.2249 | 260.6161 | 519.2409 | 260.1241 | 5 |
| 23 | 2577.2746 | 1289.1409 | 2560.2480 | 1280.6277 | 2559.2640 | 1280.1356 | S | 450.2195 | 225.6134 | 433.1929 | 217.1001 | 432.2089 | 216.6081 | 4 |
| 24 | 2706.3172 | ***1353.6622*** | 2689.2906 | 1345.1490 | 2688.3066 | 1344.6569 | E | 363.1874 | 182.0974 | 346.1609 | 173.5841 | 345.1769 | 173.0921 | 3 |
| 25 | 2793.3492 | ***1397.1782*** | 2776.3227 | 1388.6650 | 2775.3386 | 1388.1730 | S | 234.1448 | 117.5761 | 217.1183 | 109.0628 | 216.1343 | 108.5708 | 2 |
| 26 | | | | | | | K | 147.1128 | 74.0600 | 130.0863 | 65.5468 | | | 1 |

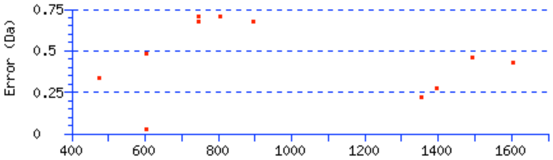

RMS error 634 ppm

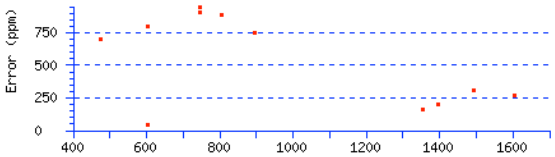

RMS error 634 ppm

MS/MS Fragmentation of **AMNTGGFDSMQRQLEAELRSLSSESK**
Found in **P48563** in **Uniprot_yeastS288C**, Protein MON2 OS=Saccharomyces cerevisiae (strain ATCC 204508 / S288c) OX=559292 GN=MON2 PE=1 SV=1

Match to Query 16253: 2938.443198 from(980.488342,3+) intensity(325277.6875) rtinseconds(5851.3603) index(15013)
Title: 20100530_methyl300mM_1.16701.16701.3
Data file D:\Panyue\Proteomic dataset\20100530_methyl300mM_1.RAW

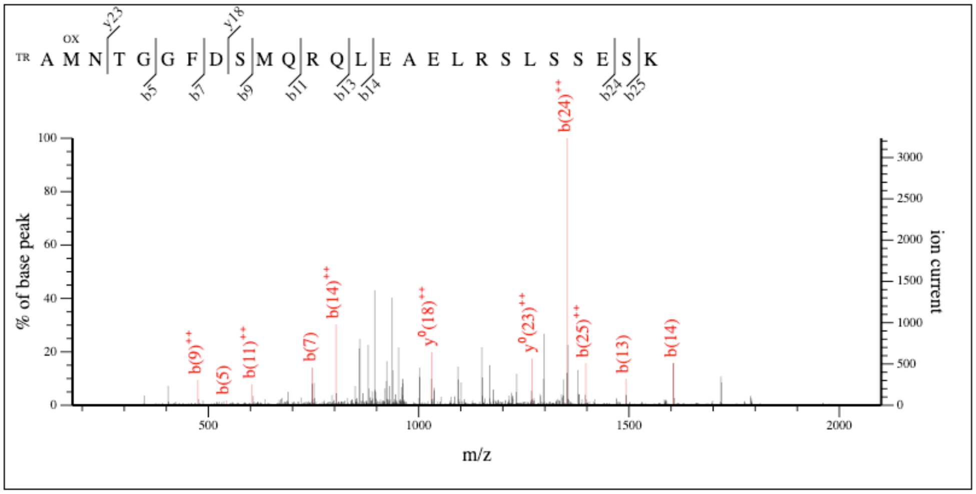


**Monoisotopic mass of neutral peptide Mr(calc):** 2938.4474
**Fixed modifications:** Carbamidomethyl (C) (apply to specified residues or termini only)
**Variable modifications:**
**N-term :** Trimethyl:2H(9) (Protein N-term)
**M2** : Oxidation (M), with neutral losses 0.0000(shown in table), 63.9983
**Ions Score:** 22 **Expect:** 0.073
**Matches :** 13/432 fragment ions using 18 most intense peaks (help)

| # | b | b++ | b* | b*++ | b0 | b0++ | Seq. | y | y++ | y* | y*++ | y0 | y0++ | # |
|---|---|---|---|---|---|---|---|---|---|---|---|---|---|---|
| 1 | 123.1478 | 62.0776 | | | | | A | | | | | | | 26 |
| 2 | 270.1832 | 135.5953 | | | | | M | 2817.3142 | 1409.1607 | 2800.2876 | 1400.6475 | 2799.3036 | 1400.1554 | 25 |
| 3 | 384.2262 | 192.6167 | 367.1996 | 184.1034 | | | N | 2670.2788 | 1335.6430 | 2653.2522 | 1327.1298 | 2652.2682 | 1326.6377 | 24 |
| 4 | 485.2738 | 243.1406 | 468.2473 | 234.6273 | 467.2633 | 234.1353 | T | 2556.2359 | 1278.6216 | 2539.2093 | 1270.1083 | 2538.2253 | 1269.6163 | 23 |
| 5 | ***542.2953*** | 271.6513 | 525.2688 | 263.1380 | 524.2847 | 262.6460 | G | 2455.1882 | 1228.0977 | 2438.1616 | 1219.5845 | 2437.1776 | 1219.0924 | 22 |
| 6 | 599.3168 | 300.1620 | 582.2902 | 291.6487 | 581.3062 | 291.1567 | G | 2398.1667 | 1199.5870 | 2381.1402 | 1191.0737 | 2380.1561 | 1190.5817 | 21 |
| 7 | ***746.3852*** | 373.6962 | 729.3586 | 365.1830 | 728.3746 | 364.6909 | F | 2341.1452 | 1171.0763 | 2324.1187 | 1162.5630 | 2323.1347 | 1162.0710 | 20 |
| 8 | 861.4121 | 431.2097 | 844.3856 | 422.6964 | 843.4016 | 422.2044 | D | 2194.0768 | 1097.5421 | 2177.0503 | 1089.0288 | 2176.0663 | 1088.5368 | 19 |
| 9 | 948.4441 | ***474.7257*** | 931.4176 | 466.2124 | 930.4336 | 465.7204 | S | 2079.0499 | 1040.0286 | 2062.0233 | 1031.5153 | 2061.0393 | 1031.0233 | 18 |
| 10 | 1079.4846 | 540.2460 | 1062.4581 | 531.7327 | 1061.4741 | 531.2407 | M | 1992.0179 | 996.5126 | 1974.9913 | 987.9993 | 1974.0073 | 987.5073 | 17 |
| 11 | 1207.5432 | ***604.2752*** | 1190.5167 | 595.7620 | 1189.5326 | 595.2700 | Q | 1860.9774 | 930.9923 | 1843.9508 | 922.4791 | 1842.9668 | 921.9870 | 16 |
| 12 | 1363.6443 | 682.3258 | 1346.6178 | 673.8125 | 1345.6338 | 673.3205 | R | 1732.9188 | 866.9630 | 1715.8923 | 858.4498 | 1714.9082 | 857.9578 | 15 |
| 13 | ***1491.7029*** | ***746.3551*** | 1474.6764 | 737.8418 | 1473.6923 | 737.3498 | Q | 1576.8177 | 788.9125 | 1559.7911 | 780.3992 | 1558.8071 | 779.9072 | 14 |
| 14 | ***1604.7870*** | ***802.8971*** | 1587.7604 | 794.3838 | 1586.7764 | 793.8918 | L | 1448.7591 | 724.8832 | 1431.7326 | 716.3699 | 1430.7485 | 715.8779 | 13 |
| 15 | 1733.8296 | 867.4184 | 1716.8030 | 858.9051 | 1715.8190 | 858.4131 | E | 1335.6750 | 668.3412 | 1318.6485 | 659.8279 | 1317.6645 | 659.3359 | 12 |
| 16 | 1804.8667 | 902.9370 | 1787.8401 | 894.4237 | 1786.8561 | 893.9317 | A | 1206.6325 | 603.8199 | 1189.6059 | 595.3066 | 1188.6219 | 594.8146 | 11 |
| 17 | 1933.9093 | 967.4583 | 1916.8827 | 958.9450 | 1915.8987 | 958.4530 | E | 1135.5953 | 568.3013 | 1118.5688 | 559.7880 | 1117.5848 | 559.2960 | 10 |
| 18 | 2046.9933 | 1024.0003 | 2029.9668 | 1015.4870 | 2028.9828 | 1014.9950 | L | 1006.5527 | 503.7800 | 989.5262 | 495.2667 | 988.5422 | 494.7747 | 9 |
| 19 | 2203.0944 | 1102.0509 | 2186.0679 | 1093.5376 | 2185.0839 | 1093.0456 | R | 893.4687 | 447.2380 | 876.4421 | 438.7247 | 875.4581 | 438.2327 | 8 |
| 20 | 2290.1265 | 1145.5669 | 2273.0999 | 1137.0536 | 2272.1159 | 1136.5616 | S | 737.3676 | 369.1874 | 720.3410 | 360.6742 | 719.3570 | 360.1821 | 7 |
| 21 | 2403.2105 | 1202.1089 | 2386.1840 | 1193.5956 | 2385.2000 | 1193.1036 | L | 650.3355 | 325.6714 | 633.3090 | 317.1581 | 632.3250 | 316.6661 | 6 |
| 22 | 2490.2426 | 1245.6249 | 2473.2160 | 1237.1116 | 2472.2320 | 1236.6196 | S | 537.2515 | 269.1294 | 520.2249 | 260.6161 | 519.2409 | 260.1241 | 5 |
| 23 | 2577.2746 | 1289.1409 | 2560.2480 | 1280.6277 | 2559.2640 | 1280.1356 | S | 450.2195 | 225.6134 | 433.1929 | 217.1001 | 432.2089 | 216.6081 | 4 |
| 24 | 2706.3172 | ***1353.6622*** | 2689.2906 | 1345.1490 | 2688.3066 | 1344.6569 | E | 363.1874 | 182.0974 | 346.1609 | 173.5841 | 345.1769 | 173.0921 | 3 |
| 25 | 2793.3492 | ***1397.1782*** | 2776.3227 | 1388.6650 | 2775.3386 | 1388.1730 | S | 234.1448 | 117.5761 | 217.1183 | 109.0628 | 216.1343 | 108.5708 | 2 |
| 26 | | | | | | | K | 147.1128 | 74.0600 | 130.0863 | 65.5468 | | | 1 |

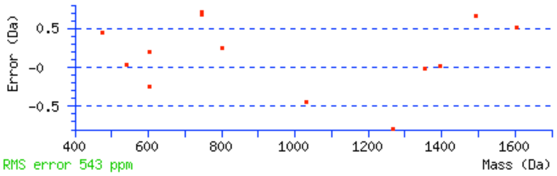

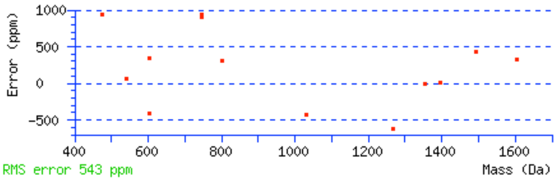

MS/MS Fragmentation of **AMNTGGFDSMQRQLEAELRSLSSESK**
Found in **P48563** in **Uniprot_yeastS288C**, Protein MON2 OS=Saccharomyces cerevisiae (strain ATCC 204508 / S288c) OX=559292 GN=MON2 PE=1 SV=1

Match to Query 16216: 2938.438071 from(980.486633,3+) intensity(256311.7344) rtinseconds(5349.1153) index(13834)
Title: 20100530_methyl300mM_1.15390.15390.3
Data file D:\Panyue\Proteomic dataset\20100530_methyl300mM_1.RAW

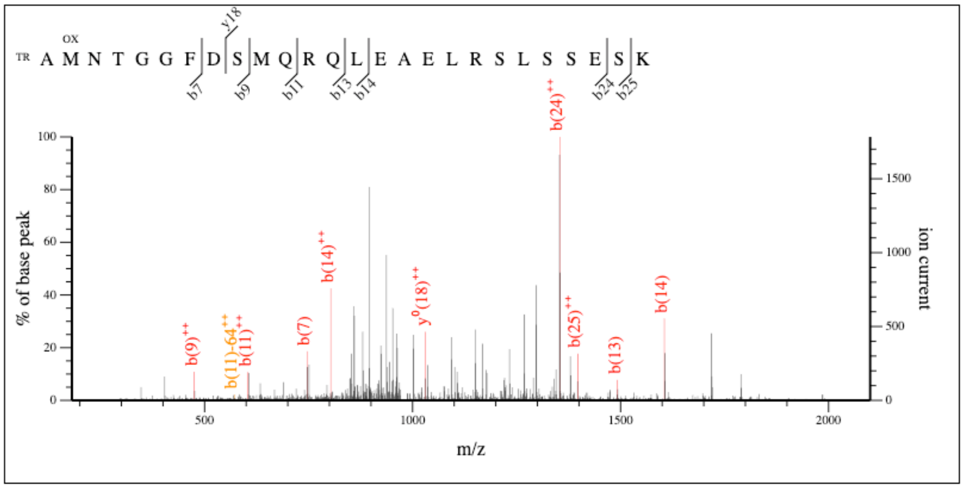

**Monoisotopic mass of neutral peptide Mr(calc):** 2938.4474
**Fixed modifications:** Carbamidomethyl (C) (apply to specified residues or termini only)
**Variable modifications:**
**N-term :** Trimethyl:2H(9) (Protein N-term)
**M2 :** Oxidation (M), with neutral losses 0.0000(shown in table), 63.9983
**Ions Score:** 20 **Expect:** 0.11
**Matches :** 12/432 fragment ions using 18 most intense peaks (help)

| # | b | b++ | b* | b*++ | b0 | b0++ | Seq. | y | y++ | y* | y*++ | y0 | y0++ | # |
|---|---|---|---|---|---|---|---|---|---|---|---|---|---|---|
| 1 | 123.1478 | 62.0776 | | | | | A | | | | | | | 26 |
| 2 | 270.1832 | 135.5953 | | | | | M | 2817.3142 | 1409.1607 | 2800.2876 | 1400.6475 | 2799.3036 | 1400.1554 | 25 |
| 3 | 384.2262 | 192.6167 | 367.1996 | 184.1034 | | | N | 2670.2788 | 1335.6430 | 2653.2522 | 1327.1298 | 2652.2682 | 1326.6377 | 24 |
| 4 | 485.2738 | 243.1406 | 468.2473 | 234.6273 | 467.2633 | 234.1353 | T | 2556.2359 | 1278.6216 | 2539.2093 | 1270.1083 | 2538.2253 | 1269.6163 | 23 |
| 5 | 542.2953 | 271.6513 | 525.2688 | 263.1380 | 524.2847 | 262.6460 | G | 2455.1882 | 1228.0977 | 2438.1616 | 1219.5845 | 2437.1776 | 1219.0924 | 22 |
| 6 | 599.3168 | 300.1620 | 582.2902 | 291.6487 | 581.3062 | 291.1567 | G | 2398.1667 | 1199.5870 | 2381.1402 | 1191.0737 | 2380.1561 | 1190.5817 | 21 |
| 7 | **746.3852** | 373.6962 | 729.3586 | 365.1830 | 728.3746 | 364.6909 | F | 2341.1452 | 1171.0763 | 2324.1187 | 1162.5630 | 2323.1347 | 1162.0710 | 20 |
| 8 | 861.4121 | 431.2097 | 844.3856 | 422.6964 | 843.4016 | 422.2044 | D | 2194.0768 | 1097.5421 | 2177.0503 | 1089.0288 | 2176.0663 | 1088.5368 | 19 |
| 9 | 948.4441 | ***474.7257*** | 931.4176 | 466.2124 | 930.4336 | 465.7204 | S | 2079.0499 | 1040.0286 | 2062.0233 | 1031.5153 | 2061.0393 | 1031.0233 | 18 |
| 10 | 1079.4846 | 540.2460 | 1062.4581 | 531.7327 | 1061.4741 | 531.2407 | M | 1992.0179 | 996.5126 | 1974.9913 | 987.9993 | 1974.0073 | 987.5073 | 17 |
| 11 | 1207.5432 | ***604.2752*** | 1190.5167 | 595.7620 | 1189.5326 | 595.2700 | Q | 1860.9774 | 930.9923 | 1843.9508 | 922.4791 | 1842.9668 | 921.9870 | 16 |
| 12 | 1363.6443 | 682.3258 | 1346.6178 | 673.8125 | 1345.6338 | 673.3205 | R | 1732.9188 | 866.9630 | 1715.8923 | 858.4498 | 1714.9082 | 857.9578 | 15 |
| 13 | **1491.7029** | ***746.3551*** | 1474.6764 | 737.8418 | 1473.6923 | 737.3498 | Q | 1576.8177 | 788.9125 | 1559.7911 | 780.3992 | 1558.8071 | 779.9072 | 14 |
| 14 | **1604.7870** | ***802.8971*** | 1587.7604 | 794.3838 | 1586.7764 | 793.8918 | L | 1448.7591 | 724.8832 | 1431.7326 | 716.3699 | 1430.7485 | 715.8779 | 13 |
| 15 | 1733.8296 | 867.4184 | 1716.8030 | 858.9051 | 1715.8190 | 858.4131 | E | 1335.6750 | 668.3412 | 1318.6485 | 659.8279 | 1317.6645 | 659.3359 | 12 |
| 16 | 1804.8667 | 902.9370 | 1787.8401 | 894.4237 | 1786.8561 | 893.9317 | A | 1206.6325 | 603.8199 | 1189.6059 | 595.3066 | 1188.6219 | 594.8146 | 11 |
| 17 | 1933.9093 | 967.4583 | 1916.8827 | 958.9450 | 1915.8987 | 958.4530 | E | 1135.5953 | 568.3013 | 1118.5688 | 559.7880 | 1117.5848 | 559.2960 | 10 |
| 18 | 2046.9933 | 1024.0003 | 2029.9668 | 1015.4870 | 2028.9828 | 1014.9950 | L | 1006.5527 | 503.7800 | 989.5262 | 495.2667 | 988.5422 | 494.7747 | 9 |
| 19 | 2203.0944 | 1102.0509 | 2186.0679 | 1093.5376 | 2185.0839 | 1093.0456 | R | 893.4687 | 447.2380 | 876.4421 | 438.7247 | 875.4581 | 438.2327 | 8 |
| 20 | 2290.1265 | 1145.5669 | 2273.0999 | 1137.0536 | 2272.1159 | 1136.5616 | S | 737.3676 | 369.1874 | 720.3410 | 360.6742 | 719.3570 | 360.1821 | 7 |
| 21 | 2403.2105 | 1202.1089 | 2386.1840 | 1193.5956 | 2385.2000 | 1193.1036 | L | 650.3355 | 325.6714 | 633.3090 | 317.1581 | 632.3250 | 316.6661 | 6 |
| 22 | 2490.2426 | 1245.6249 | 2473.2160 | 1237.1116 | 2472.2320 | 1236.6196 | S | 537.2515 | 269.1294 | 520.2249 | 260.6161 | 519.2409 | 260.1241 | 5 |
| 23 | 2577.2746 | 1289.1409 | 2560.2480 | 1280.6277 | 2559.2640 | 1280.1356 | S | 450.2195 | 225.6134 | 433.1929 | 217.1001 | 432.2089 | 216.6081 | 4 |
| 24 | 2706.3172 | ***1353.6622*** | 2689.2906 | 1345.1490 | 2688.3066 | 1344.6569 | E | 363.1874 | 182.0974 | 346.1609 | 173.5841 | 345.1769 | 173.0921 | 3 |
| 25 | 2793.3492 | ***1397.1782*** | 2776.3227 | 1388.6650 | 2775.3386 | 1388.1730 | S | 234.1448 | 117.5761 | 217.1183 | 109.0628 | 216.1343 | 108.5708 | 2 |
| 26 | | | | | | | K | 147.1128 | 74.0600 | 130.0863 | 65.5468 | | | 1 |

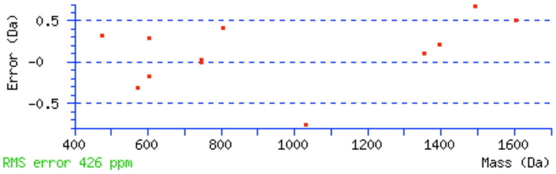


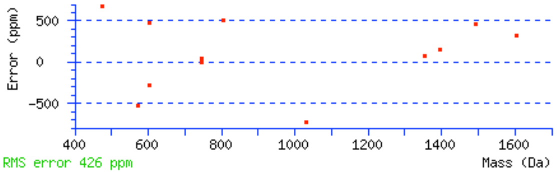


MS/MS Fragmentation of **AMNTGGFDSMQRQLEAELRSLSSESK**
Found in **P48563** in **Uniprot_yeastS288C**, Protein MON2 OS=Saccharomyces cerevisiae (strain ATCC 204508 / S288c) OX=559292 GN=MON2 PE=1 SV=1

Match to Query 16215: 2938.437888 from(980.486572,3+) intensity(264404.9688) rtinseconds(5846.8823) index(15002)
Title: 20100530_methyl300mM_1.16689.16689.3
Data file D:\Panyue\Proteomic dataset\20100530_methyl300mM_1.RAW

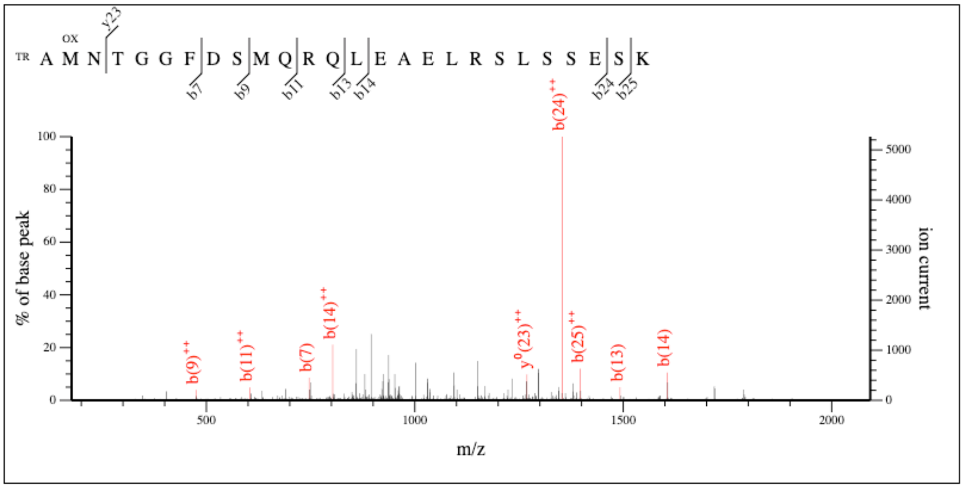

**Monoisotopic mass of neutral peptide Mr(calc):** 2938.4474
**Fixed modifications:** Carbamidomethyl (C) (apply to specified residues or termini only)
**Variable modifications:**
**N-term :** Trimethyl:2H(9) (Protein N-term)
**M2 :** Oxidation (M), with neutral losses 0.0000(shown in table), 63.9983
**Ions Score:** 21 **Expect:** 0.093
**Matches :** 11/432 fragment ions using 18 most intense peaks (help)

| # | b | b++ | b* | b*++ | b0 | b0++ | Seq. | y | y++ | y* | y*++ | y0 | y0++ | # |
|---|---|---|---|---|---|---|---|---|---|---|---|---|---|---|
| 1 | 123.1478 | 62.0776 | | | | | A | | | | | | | 26 |
| 2 | 270.1832 | 135.5953 | | | | | M | 2817.3142 | 1409.1607 | 2800.2876 | 1400.6475 | 2799.3036 | 1400.1554 | 25 |
| 3 | 384.2262 | 192.6167 | 367.1996 | 184.1034 | | | N | 2670.2788 | 1335.6430 | 2653.2522 | 1327.1298 | 2652.2682 | 1326.6377 | 24 |
| 4 | 485.2738 | 243.1406 | 468.2473 | 234.6273 | 467.2633 | 234.1353 | T | 2556.2359 | 1278.6216 | 2539.2093 | 1270.1083 | 2538.2253 | 1269.6163 | 23 |
| 5 | 542.2953 | 271.6513 | 525.2688 | 263.1380 | 524.2847 | 262.6460 | G | 2455.1882 | 1228.0977 | 2438.1616 | 1219.5845 | 2437.1776 | 1219.0924 | 22 |
| 6 | 599.3168 | 300.1620 | 582.2902 | 291.6487 | 581.3062 | 291.1567 | G | 2398.1667 | 1199.5870 | 2381.1402 | 1191.0737 | 2380.1561 | 1190.5817 | 21 |
| 7 | **746.3852** | 373.6962 | 729.3586 | 365.1830 | 728.3746 | 364.6909 | F | 2341.1452 | 1171.0763 | 2324.1187 | 1162.5630 | 2323.1347 | 1162.0710 | 20 |
| 8 | 861.4121 | 431.2097 | 844.3856 | 422.6964 | 843.4016 | 422.2044 | D | 2194.0768 | 1097.5421 | 2177.0503 | 1089.0288 | 2176.0663 | 1088.5368 | 19 |
| 9 | 948.4441 | ***474.7257*** | 931.4176 | 466.2124 | 930.4336 | 465.7204 | S | 2079.0499 | 1040.0286 | 2062.0233 | 1031.5153 | 2061.0393 | 1031.0233 | 18 |
| 10 | 1079.4846 | 540.2460 | 1062.4581 | 531.7327 | 1061.4741 | 531.2407 | M | 1992.0179 | 996.5126 | 1974.9913 | 987.9993 | 1974.0073 | 987.5073 | 17 |
| 11 | 1207.5432 | ***604.2752*** | 1190.5167 | 595.7620 | 1189.5326 | 595.2700 | Q | 1860.9774 | 930.9923 | 1843.9508 | 922.4791 | 1842.9668 | 921.9870 | 16 |
| 12 | 1363.6443 | 682.3258 | 1346.6178 | 673.8125 | 1345.6338 | 673.3205 | R | 1732.9188 | 866.9630 | 1715.8923 | 858.4498 | 1714.9082 | 857.9578 | 15 |
| 13 | **1491.7029** | ***746.3551*** | 1474.6764 | 737.8418 | 1473.6923 | 737.3498 | Q | 1576.8177 | 788.9125 | 1559.7911 | 780.3992 | 1558.8071 | 779.9072 | 14 |
| 14 | **1604.7870** | ***802.8971*** | 1587.7604 | 794.3838 | 1586.7764 | 793.8918 | L | 1448.7591 | 724.8832 | 1431.7326 | 716.3699 | 1430.7485 | 715.8779 | 13 |
| 15 | 1733.8296 | 867.4184 | 1716.8030 | 858.9051 | 1715.8190 | 858.4131 | E | 1335.6750 | 668.3412 | 1318.6485 | 659.8279 | 1317.6645 | 659.3359 | 12 |
| 16 | 1804.8667 | 902.9370 | 1787.8401 | 894.4237 | 1786.8561 | 893.9317 | A | 1206.6325 | 603.8199 | 1189.6059 | 595.3066 | 1188.6219 | 594.8146 | 11 |
| 17 | 1933.9093 | 967.4583 | 1916.8827 | 958.9450 | 1915.8987 | 958.4530 | E | 1135.5953 | 568.3013 | 1118.5688 | 559.7880 | 1117.5848 | 559.2960 | 10 |
| 18 | 2046.9933 | 1024.0003 | 2029.9668 | 1015.4870 | 2028.9828 | 1014.9950 | L | 1006.5527 | 503.7800 | 989.5262 | 495.2667 | 988.5422 | 494.7747 | 9 |
| 19 | 2203.0944 | 1102.0509 | 2186.0679 | 1093.5376 | 2185.0839 | 1093.0456 | R | 893.4687 | 447.2380 | 876.4421 | 438.7247 | 875.4581 | 438.2327 | 8 |
| 20 | 2290.1265 | 1145.5669 | 2273.0999 | 1137.0536 | 2272.1159 | 1136.5616 | S | 737.3676 | 369.1874 | 720.3410 | 360.6742 | 719.3570 | 360.1821 | 7 |
| 21 | 2403.2105 | 1202.1089 | 2386.1840 | 1193.5956 | 2385.2000 | 1193.1036 | L | 650.3355 | 325.6714 | 633.3090 | 317.1581 | 632.3250 | 316.6661 | 6 |
| 22 | 2490.2426 | 1245.6249 | 2473.2160 | 1237.1116 | 2472.2320 | 1236.6196 | S | 537.2515 | 269.1294 | 520.2249 | 260.6161 | 519.2409 | 260.1241 | 5 |
| 23 | 2577.2746 | 1289.1409 | 2560.2480 | 1280.6277 | 2559.2640 | 1280.1356 | S | 450.2195 | 225.6134 | 433.1929 | 217.1001 | 432.2089 | 216.6081 | 4 |
| 24 | 2706.3172 | ***1353.6622*** | 2689.2906 | 1345.1490 | 2688.3066 | 1344.6569 | E | 363.1874 | 182.0974 | 346.1609 | 173.5841 | 345.1769 | 173.0921 | 3 |
| 25 | 2793.3492 | ***1397.1782*** | 2776.3227 | 1388.6650 | 2775.3386 | 1388.1730 | S | 234.1448 | 117.5761 | 217.1183 | 109.0628 | 216.1343 | 108.5708 | 2 |
| 26 | | | | | | | K | 147.1128 | 74.0600 | 130.0863 | 65.5468 | | | 1 |

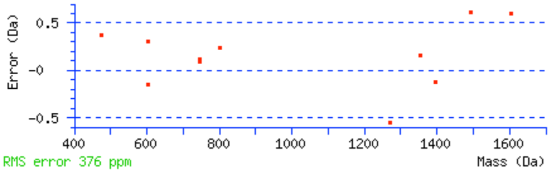


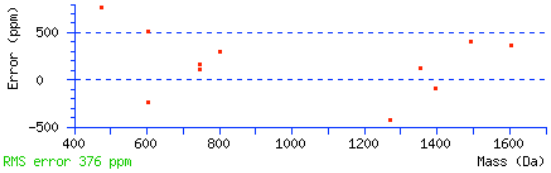

MS/MS Fragmentation of **AMNTGGFDSMQRQLEAELRSLSSESK**
Found in **P48563** in **Uniprot_yeastS288C**, Protein MON2 OS=Saccharomyces cerevisiae (strain ATCC 204508 / S288c) OX=559292 GN=MON2 PE=1 SV=1

Match to Query 16212: 2938.433859 from(980.485229,3+) intensity(915230.3125) rtinseconds(6464.8037) index(16493)
Title: 20100530_methyl300mM_1.18347.18347.3
Data file D:\Panyue\Proteomic dataset\20100530_methyl300mM_1.RAW

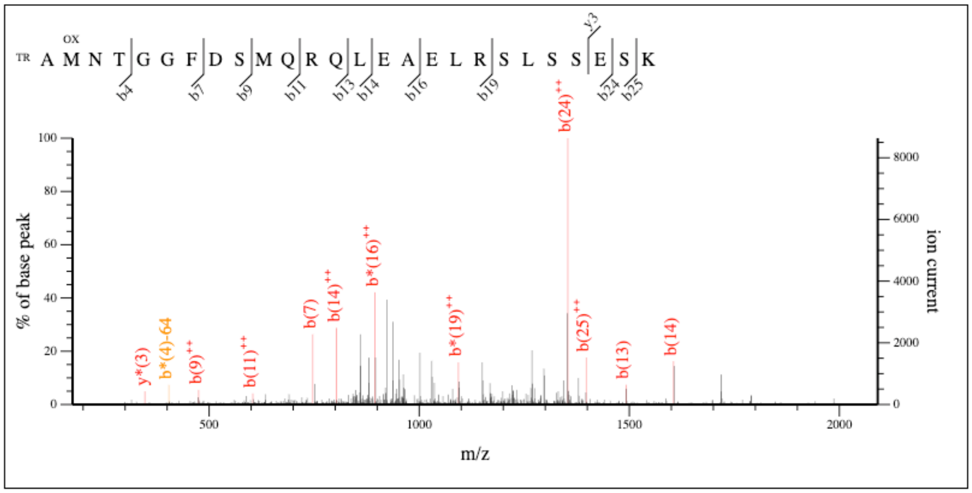


**Monoisotopic mass of neutral peptide Mr(calc):** 2938.4474
**Fixed modifications:** Carbamidomethyl (C) (apply to specified residues or termini only)
**Variable modifications:**
**N-term :** Trimethyl:2H(9) (Protein N-term)
**M2 :** Oxidation (M), with neutral losses 0.0000(shown in table), 63.9983
**Ions Score:** 24 **Expect:** 0.039
**Matches :** 16/432 fragment ions using 18 most intense peaks (help)

| # | b | b++ | b* | b*++ | b0 | b0++ | Seq. | y | y++ | y* | y*++ | y0 | y0++ | # |
|---|---|---|---|---|---|---|---|---|---|---|---|---|---|---|
| 1 | 123.1478 | 62.0776 | | | | | A | | | | | | | 26 |
| 2 | 270.1832 | 135.5953 | | | | | M | 2817.3142 | 1409.1607 | 2800.2876 | 1400.6475 | 2799.3036 | 1400.1554 | 25 |
| 3 | 384.2262 | 192.6167 | 367.1996 | 184.1034 | | | N | 2670.2788 | 1335.6430 | 2653.2522 | 1327.1298 | 2652.2682 | 1326.6377 | 24 |
| 4 | 485.2738 | 243.1406 | 468.2473 | 234.6273 | 467.2633 | 234.1353 | T | 2556.2359 | 1278.6216 | 2539.2093 | 1270.1083 | 2538.2253 | 1269.6163 | 23 |
| 5 | 542.2953 | 271.6513 | 525.2688 | 263.1380 | 524.2847 | 262.6460 | G | 2455.1882 | 1228.0977 | 2438.1616 | 1219.5845 | 2437.1776 | 1219.0924 | 22 |
| 6 | 599.3168 | 300.1620 | 582.2902 | 291.6487 | 581.3062 | 291.1567 | G | 2398.1667 | 1199.5870 | 2381.1402 | 1191.0737 | 2380.1561 | 1190.5817 | 21 |
| 7 | **746.3852** | 373.6962 | 729.3586 | 365.1830 | 728.3746 | 364.6909 | F | 2341.1452 | 1171.0763 | 2324.1187 | 1162.5630 | 2323.1347 | 1162.0710 | 20 |
| 8 | 861.4121 | 431.2097 | 844.3856 | 422.6964 | 843.4016 | 422.2044 | D | 2194.0768 | 1097.5421 | 2177.0503 | 1089.0288 | 2176.0663 | 1088.5368 | 19 |
| 9 | 948.4441 | ***474.7257*** | 931.4176 | 466.2124 | 930.4336 | 465.7204 | S | 2079.0499 | 1040.0286 | 2062.0233 | 1031.5153 | 2061.0393 | 1031.0233 | 18 |
| 10 | 1079.4846 | 540.2460 | 1062.4581 | 531.7327 | 1061.4741 | 531.2407 | M | 1992.0179 | 996.5126 | 1974.9913 | 987.9993 | 1974.0073 | 987.5073 | 17 |
| 11 | 1207.5432 | ***604.2752*** | 1190.5167 | 595.7620 | 1189.5326 | 595.2700 | Q | 1860.9774 | 930.9923 | 1843.9508 | 922.4791 | 1842.9668 | 921.9870 | 16 |
| 12 | 1363.6443 | 682.3258 | 1346.6178 | 673.8125 | 1345.6338 | 673.3205 | R | 1732.9188 | 866.9630 | 1715.8923 | 858.4498 | 1714.9082 | 857.9578 | 15 |
| 13 | **1491.7029** | ***746.3551*** | 1474.6764 | 737.8418 | 1473.6923 | 737.3498 | Q | 1576.8177 | 788.9125 | 1559.7911 | 780.3992 | 1558.8071 | 779.9072 | 14 |
| 14 | **1604.7870** | ***802.8971*** | 1587.7604 | 794.3838 | 1586.7764 | 793.8918 | L | 1448.7591 | 724.8832 | 1431.7326 | 716.3699 | 1430.7485 | 715.8779 | 13 |
| 15 | 1733.8296 | 867.4184 | 1716.8030 | 858.9051 | 1715.8190 | 858.4131 | E | 1335.6750 | 668.3412 | 1318.6485 | 659.8279 | 1317.6645 | 659.3359 | 12 |
| 16 | 1804.8667 | 902.9370 | 1787.8401 | 894.4237 | 1786.8561 | 893.9317 | A | 1206.6325 | 603.8199 | 1189.6059 | 595.3066 | 1188.6219 | 594.8146 | 11 |
| 17 | 1933.9093 | 967.4583 | 1916.8827 | 958.9450 | 1915.8987 | 958.4530 | E | 1135.5953 | 568.3013 | 1118.5688 | 559.7880 | 1117.5848 | 559.2960 | 10 |
| 18 | 2046.9933 | 1024.0003 | 2029.9668 | 1015.4870 | 2028.9828 | 1014.9950 | L | 1006.5527 | 503.7800 | 989.5262 | 495.2667 | 988.5422 | 494.7747 | 9 |
| 19 | 2203.0944 | 1102.0509 | 2186.0679 | 1093.5376 | 2185.0839 | 1093.0456 | R | 893.4687 | 447.2380 | 876.4421 | 438.7247 | 875.4581 | 438.2327 | 8 |
| 20 | 2290.1265 | 1145.5669 | 2273.0999 | 1137.0536 | 2272.1159 | 1136.5616 | S | 737.3676 | 369.1874 | 720.3410 | 360.6742 | 719.3570 | 360.1821 | 7 |
| 21 | 2403.2105 | 1202.1089 | 2386.1840 | 1193.5956 | 2385.2000 | 1193.1036 | L | 650.3355 | 325.6714 | 633.3090 | 317.1581 | 632.3250 | 316.6661 | 6 |
| 22 | 2490.2426 | 1245.6249 | 2473.2160 | 1237.1116 | 2472.2320 | 1236.6196 | S | 537.2515 | 269.1294 | 520.2249 | 260.6161 | 519.2409 | 260.1241 | 5 |
| 23 | 2577.2746 | 1289.1409 | 2560.2480 | 1280.6277 | 2559.2640 | 1280.1356 | S | 450.2195 | 225.6134 | 433.1929 | 217.1001 | 432.2089 | 216.6081 | 4 |
| 24 | 2706.3172 | ***1353.6622*** | 2689.2906 | 1345.1490 | 2688.3066 | 1344.6569 | E | 363.1874 | 182.0974 | 346.1609 | 173.5841 | 345.1769 | 173.0921 | 3 |
| 25 | 2793.3492 | ***1397.1782*** | 2776.3227 | 1388.6650 | 2775.3386 | 1388.1730 | S | 234.1448 | 117.5761 | 217.1183 | 109.0628 | 216.1343 | 108.5708 | 2 |
| 26 | | | | | | | K | 147.1128 | 74.0600 | 130.0863 | 65.5468 | | | 1 |

MS/MS Fragmentation of **AMNTGGFDSMQRQLEAELRSLSSESK**
Found in **P48563** in **Uniprot_yeastS288C**, Protein MON2 OS=Saccharomyces cerevisiae (strain ATCC 204508 / S288c) OX=559292 GN=MON2 PE=1 SV=1

Match to Query 16213: 2938.433859 from(980.485229,3+) intensity(915230.3125) rtinseconds(6465.8409) index(16496)
Title: 20100530_methyl300mM_1.18350.18350.3
Data file D:\Panyue\Proteomic dataset\20100530_methyl300mM_1.RAW

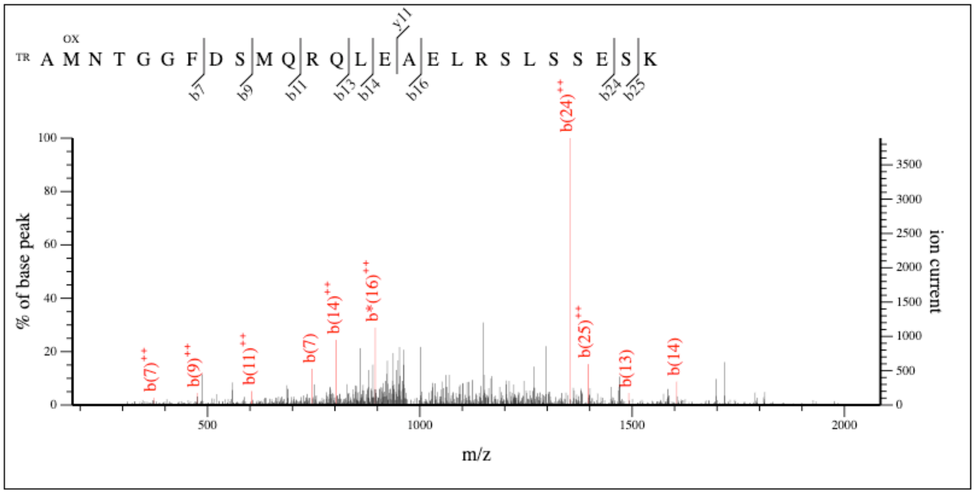


```
Monoisotopic mass of neutral peptide Mr(calc): 2938.4474
Fixed modifications: Carbamidomethyl (C) (apply to specified residues or termini only)
Variable modifications:
N-term : Trimethyl:2H(9) (Protein N-term)
M2     : Oxidation (M), with neutral losses 0.0000(shown in table), 63.9983
Ions Score: 30  Expect: 0.01
Matches : 12/432 fragment ions using 18 most intense peaks   (help)
```

| # | b | b++ | b* | b*++ | b0 | b0++ | Seq. | y | y++ | y* | y*++ | y0 | y0++ | # |
|---|---|---|---|---|---|---|---|---|---|---|---|---|---|---|
| 1 | 123.1478 | 62.0776 | | | | | A | | | | | | | 26 |
| 2 | 270.1832 | 135.5953 | | | | | M | 2817.3142 | 1409.1607 | 2800.2876 | 1400.6475 | 2799.3036 | 1400.1554 | 25 |
| 3 | 384.2262 | 192.6167 | 367.1996 | 184.1034 | | | N | 2670.2788 | 1335.6430 | 2653.2522 | 1327.1298 | 2652.2682 | 1326.6377 | 24 |
| 4 | 485.2738 | 243.1406 | 468.2473 | 234.6273 | 467.2633 | 234.1353 | T | 2556.2359 | 1278.6216 | 2539.2093 | 1270.1083 | 2538.2253 | 1269.6163 | 23 |
| 5 | 542.2953 | 271.6513 | 525.2688 | 263.1380 | 524.2847 | 262.6460 | G | 2455.1882 | 1228.0977 | 2438.1616 | 1219.5845 | 2437.1776 | 1219.0924 | 22 |
| 6 | 599.3168 | 300.1620 | 582.2902 | 291.6487 | 581.3062 | 291.1567 | G | 2398.1667 | 1199.5870 | 2381.1402 | 1191.0737 | 2380.1561 | 1190.5817 | 21 |
| 7 | 746.3852 | 373.6962 | 729.3586 | 365.1830 | 728.3746 | 364.6909 | F | 2341.1452 | 1171.0763 | 2324.1187 | 1162.5630 | 2323.1347 | 1162.0710 | 20 |
| 8 | 861.4121 | 431.2097 | 844.3856 | 422.6964 | 843.4016 | 422.2044 | D | 2194.0768 | 1097.5421 | 2177.0503 | 1089.0288 | 2176.0663 | 1088.5368 | 19 |
| 9 | 948.4441 | 474.7257 | 931.4176 | 466.2124 | 930.4336 | 465.7204 | S | 2079.0499 | 1040.0286 | 2062.0233 | 1031.5153 | 2061.0393 | 1031.0233 | 18 |
| 10 | 1079.4846 | 540.2460 | 1062.4581 | 531.7327 | 1061.4741 | 531.2407 | M | 1992.0179 | 996.5126 | 1974.9913 | 987.9993 | 1974.0073 | 987.5073 | 17 |
| 11 | 1207.5432 | 604.2752 | 1190.5167 | 595.7620 | 1189.5326 | 595.2700 | Q | 1860.9774 | 930.9923 | 1843.9508 | 922.4791 | 1842.9668 | 921.9870 | 16 |
| 12 | 1363.6443 | 682.3258 | 1346.6178 | 673.8125 | 1345.6338 | 673.3205 | R | 1732.9188 | 866.9630 | 1715.8923 | 858.4498 | 1714.9082 | 857.9578 | 15 |
| 13 | 1491.7029 | 746.3551 | 1474.6764 | 737.8418 | 1473.6923 | 737.3498 | Q | 1576.8177 | 788.9125 | 1559.7911 | 780.3992 | 1558.8071 | 779.9072 | 14 |
| 14 | 1604.7870 | 802.8971 | 1587.7604 | 794.3838 | 1586.7764 | 793.8918 | L | 1448.7591 | 724.8832 | 1431.7326 | 716.3699 | 1430.7485 | 715.8779 | 13 |
| 15 | 1733.8296 | 867.4184 | 1716.8030 | 858.9051 | 1715.8190 | 858.4131 | E | 1335.6750 | 668.3412 | 1318.6485 | 659.8279 | 1317.6645 | 659.3359 | 12 |
| 16 | 1804.8667 | 902.9370 | 1787.8401 | 894.4237 | 1786.8561 | 893.9317 | A | 1206.6325 | 603.8199 | 1189.6059 | 595.3066 | 1188.6219 | 594.8146 | 11 |
| 17 | 1933.9093 | 967.4583 | 1916.8827 | 958.9450 | 1915.8987 | 958.4530 | E | 1135.5953 | 568.3013 | 1118.5688 | 559.7880 | 1117.5848 | 559.2960 | 10 |
| 18 | 2046.9933 | 1024.0003 | 2029.9668 | 1015.4870 | 2028.9828 | 1014.9950 | L | 1006.5527 | 503.7800 | 989.5262 | 495.2667 | 988.5422 | 494.7747 | 9 |
| 19 | 2203.0944 | 1102.0509 | 2186.0679 | 1093.5376 | 2185.0839 | 1093.0456 | R | 893.4687 | 447.2380 | 876.4421 | 438.7247 | 875.4581 | 438.2327 | 8 |
| 20 | 2290.1265 | 1145.5669 | 2273.0999 | 1137.0536 | 2272.1159 | 1136.5616 | S | 737.3676 | 369.1874 | 720.3410 | 360.6742 | 719.3570 | 360.1821 | 7 |
| 21 | 2403.2105 | 1202.1089 | 2386.1840 | 1193.5956 | 2385.2000 | 1193.1036 | L | 650.3355 | 325.6714 | 633.3090 | 317.1581 | 632.3250 | 316.6661 | 6 |
| 22 | 2490.2426 | 1245.6249 | 2473.2160 | 1237.1116 | 2472.2320 | 1236.6196 | S | 537.2515 | 269.1294 | 520.2249 | 260.6161 | 519.2409 | 260.1241 | 5 |
| 23 | 2577.2746 | 1289.1409 | 2560.2480 | 1280.6277 | 2559.2640 | 1280.1356 | S | 450.2195 | 225.6134 | 433.1929 | 217.1001 | 432.2089 | 216.6081 | 4 |
| 24 | 2706.3172 | 1353.6622 | 2689.2906 | 1345.1490 | 2688.3066 | 1344.6569 | E | 363.1874 | 182.0974 | 346.1609 | 173.5841 | 345.1769 | 173.0921 | 3 |
| 25 | 2793.3492 | 1397.1782 | 2776.3227 | 1388.6650 | 2775.3386 | 1388.1730 | S | 234.1448 | 117.5761 | 217.1183 | 109.0628 | 216.1343 | 108.5708 | 2 |
| 26 | | | | | | | K | 147.1128 | 74.0600 | 130.0863 | 65.5468 | | | 1 |

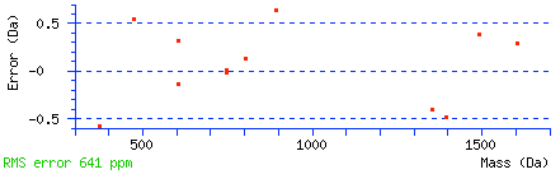


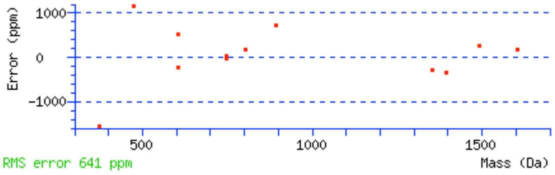


### 30. Pdc6 (2 peptides in 300mM fraction 1)

MS/MS Fragmentation of **SEITLGK**
Found in **P26263** in **Uniprot_yeastS288C**, Pyruvate decarboxylase isozyme 3 OS=Saccharomyces cerevisiae (strain ATCC 204508 / S288c) OX=559292 GN=PDC6 PE=1 SV=3

Match to Query 2027: 780.447924 from(781.455200,1+) intensity(1382002.5000) rtinseconds(965.5585) index(2410)
Title: 20100530_methyl300mM_1.2681.2681.1
Data file D:\Panyue\Proteomic dataset\20100530_methyl300mM_1.RAW

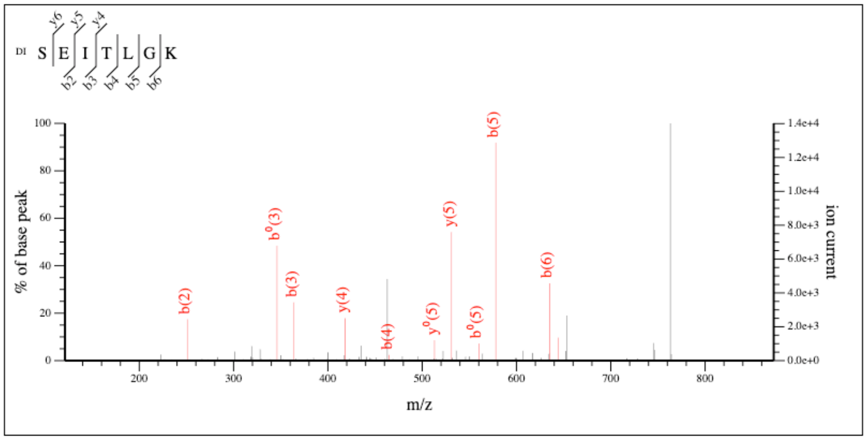


**Monoisotopic mass of neutral peptide Mr(calc):** 780.4864
**Fixed modifications:** Carbamidomethyl (C) (apply to specified residues or termini only)
**Variable modifications:**
**N-term :** Dimethyl:2H(6) (Protein N-term)
**Ions Score:** 34 **Expect:** 0.0046
**Matches :** 11/27 fragment ions using 24 most intense peaks (help)

| # | b | $b^0$ | Seq. | y | y* | $y^0$ | # |
|---|---|---|---|---|---|---|---|
| 1 | 122.1083 | 104.0977 | S | | | | 7 |
| 2 | *251.1509* | 233.1403 | E | 660.3927 | 643.3661 | 642.3821 | 6 |
| 3 | *364.2349* | 346.2244 | I | **531.3501** | 514.3235 | 513.3395 | 5 |
| 4 | *465.2826* | 447.2720 | T | **418.2660** | 401.2395 | 400.2554 | 4 |
| 5 | *578.3667* | 560.3561 | L | 317.2183 | 300.1918 | | 3 |
| 6 | *635.3881* | 617.3776 | G | 204.1343 | 187.1077 | | 2 |
| 7 | | | K | 147.1128 | 130.0863 | | 1 |

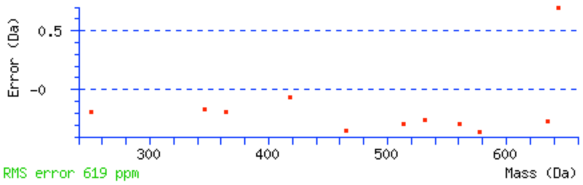


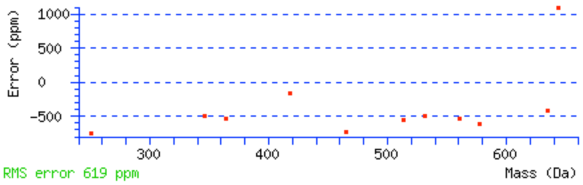

MS/MS Fragmentation of **SEITLGK**
Found in **P26263** in **Uniprot_yeastS288C**, Pyruvate decarboxylase isozyme 3 OS=Saccharomyces cerevisiae (strain ATCC 204508 / S288c) OX=559292 GN=PDC6 PE=1 SV=3

Match to Query 2028: 780.448901 from(781.456177,1+) intensity(579563.8125) rtinseconds(962.3724) index(2401)
Title: 20100530_methyl300mM_1.2671.2671.1
Data file D:\Panyue\Proteomic dataset\20100530_methyl300mM_1.RAW

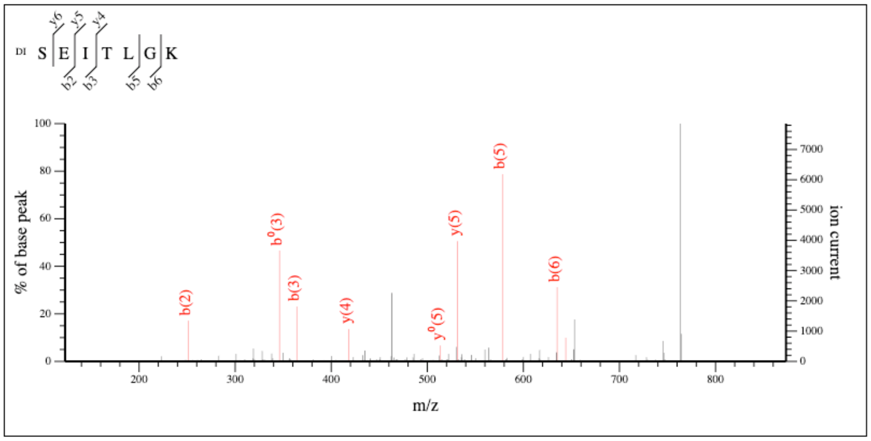


**Monoisotopic mass of neutral peptide Mr(calc):** 780.4864
**Fixed modifications:** Carbamidomethyl (C) (apply to specified residues or termini only)
**Variable modifications:**
**N-term :** Dimethyl:2H(6) (Protein N-term)
**Ions Score:** 29 **Expect:** 0.017
**Matches :** 9/27 fragment ions using 18 most intense peaks (help)

| # | b | $b^0$ | Seq. | y | y* | $y^0$ | # |
|---|---|---|---|---|---|---|---|
| 1 | 122.1083 | 104.0977 | S | | | | 7 |
| 2 | 251.1509 | 233.1403 | E | 660.3927 | 643.3661 | 642.3821 | 6 |
| 3 | 364.2349 | 346.2244 | I | 531.3501 | 514.3235 | 513.3395 | 5 |
| 4 | 465.2826 | 447.2720 | T | 418.2660 | 401.2395 | 400.2554 | 4 |
| 5 | 578.3667 | 560.3561 | L | 317.2183 | 300.1918 | | 3 |
| 6 | 635.3881 | 617.3776 | G | 204.1343 | 187.1077 | | 2 |
| 7 | | | K | 147.1128 | 130.0863 | | 1 |

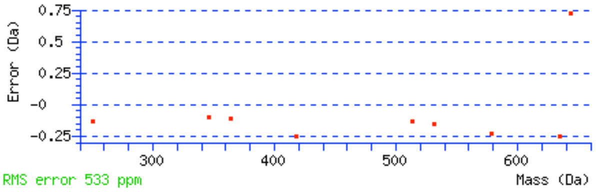


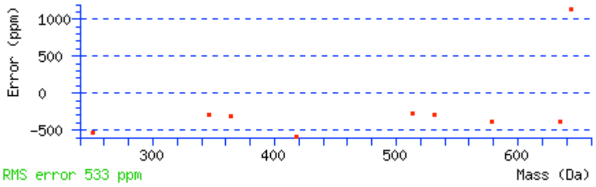


31. Pdc5 (2 peptides in 300mM fraction 1)

MS/MS Fragmentation of **SEITLGK**
Found in **P16467** in **Uniprot_yeastS288C**, Pyruvate decarboxylase isozyme 2 OS=Saccharomyces cerevisiae (strain ATCC 204508 / S288c) OX=559292 GN=PDC5 PE=1 SV=4

Match to Query 2027: 780.447924 from(781.455200,1+) intensity(1382002.5000) rtinseconds(965.5585) index(2410)
Title: 20100530_methyl300mM_1.2681.2681.1
Data file D:\Panyue\Proteomic dataset\20100530_methyl300mM_1.RAW

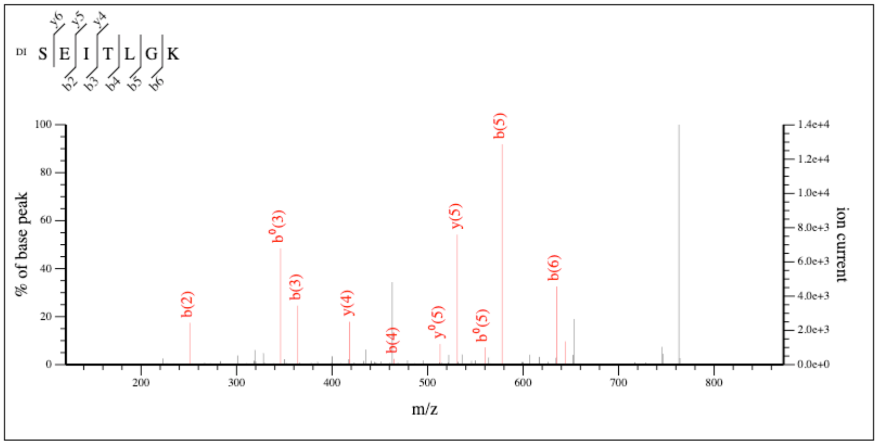


**Monoisotopic mass of neutral peptide Mr(calc):** 780.4864
**Fixed modifications:** Carbamidomethyl (C) (apply to specified residues or termini only)
**Variable modifications:**
**N-term :** Dimethyl:2H(6) (Protein N-term)
**Ions Score:** 34 **Expect:** 0.0046
**Matches :** 11/27 fragment ions using 24 most intense peaks (help)

| # | b | $b^0$ | Seq. | y | y* | $y^0$ | # |
|---|---|---|---|---|---|---|---|
| 1 | 122.1083 | 104.0977 | S | | | | 7 |
| 2 | ***251.1509*** | 233.1403 | E | 660.3927 | 643.3661 | 642.3821 | 6 |
| 3 | ***364.2349*** | 346.2244 | I | **531.3501** | 514.3235 | 513.3395 | 5 |
| 4 | ***465.2826*** | 447.2720 | T | **418.2660** | 401.2395 | 400.2554 | 4 |
| 5 | ***578.3667*** | 560.3561 | L | 317.2183 | 300.1918 | | 3 |
| 6 | ***635.3881*** | 617.3776 | G | 204.1343 | 187.1077 | | 2 |
| 7 | | | K | 147.1128 | 130.0863 | | 1 |

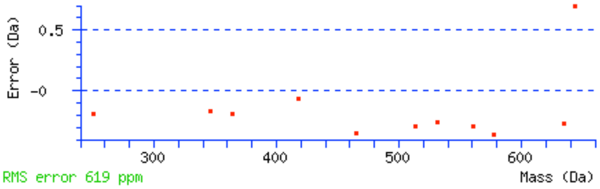


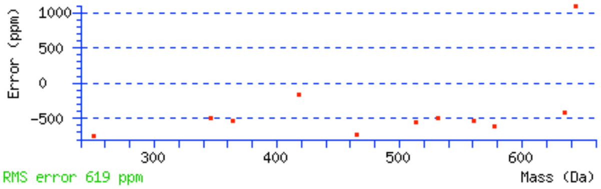

MS/MS Fragmentation of SEITLGK
Found in P16467 in Uniprot_yeastS288C, Pyruvate decarboxylase isozyme 2 OS=Saccharomyces cerevisiae (strain ATCC 204508 / S288c) OX=559292 GN=PDC5 PE=1 SV=4

Match to Query 2028: 780.448901 from(781.456177,1+) intensity(579563.8125) rtinseconds(962.3724) index(2401)
Title: 20100530_methyl300mM_1.2671.2671.1
Data file D:\Panyue\Proteomic dataset\20100530_methyl300mM_1.RAW

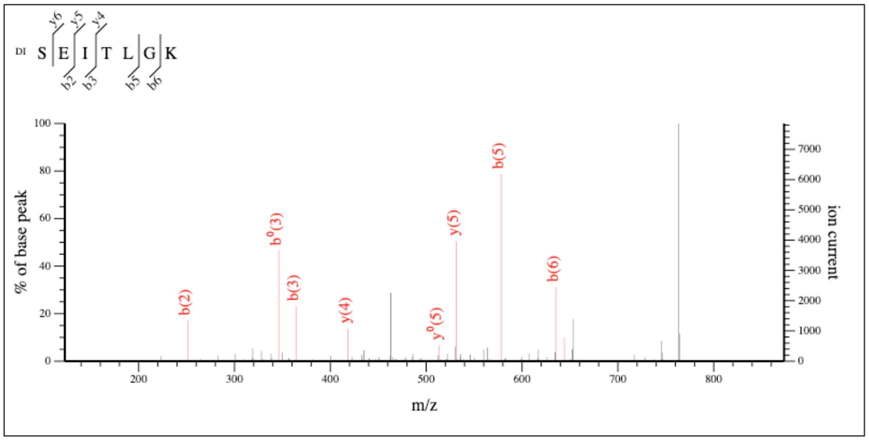


**Monoisotopic mass of neutral peptide Mr(calc):** 780.4864
**Fixed modifications:** Carbamidomethyl (C) (apply to specified residues or termini only)
**Variable modifications:**
**N-term :** Dimethyl:2H(6) (Protein N-term)
**Ions Score:** 29 **Expect:** 0.017
**Matches :** 9/27 fragment ions using 18 most intense peaks (help)

| # | b | b$^0$ | Seq. | y | y* | y$^0$ | # |
|---|---|---|---|---|---|---|---|
| 1 | 122.1083 | 104.0977 | S | | | | 7 |
| 2 | *251.1509* | 233.1403 | E | 660.3927 | 643.3661 | 642.3821 | 6 |
| 3 | *364.2349* | 346.2244 | I | **531.3501** | 514.3235 | 513.3395 | 5 |
| 4 | 465.2826 | 447.2720 | T | **418.2660** | 401.2395 | 400.2554 | 4 |
| 5 | *578.3667* | 560.3561 | L | 317.2183 | 300.1918 | | 3 |
| 6 | *635.3881* | 617.3776 | G | 204.1343 | 187.1077 | | 2 |
| 7 | | | K | 147.1128 | 130.0863 | | 1 |

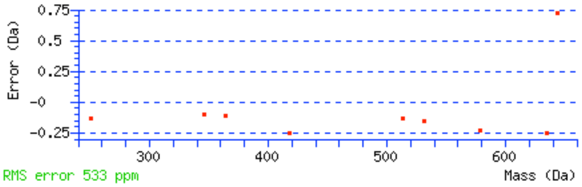


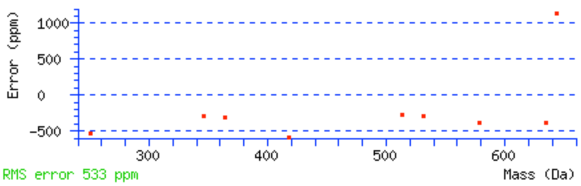


32. Pdc1 (2peptides in 300mM fraction 1)

MS/MS Fragmentation of **SEITLGK**
Found in **P06169** in **Uniprot_yeastS288C**, Pyruvate decarboxylase isozyme 1 OS=Saccharomyces cerevisiae (strain ATCC 204508 / S288c) OX=559292 GN=PDC1 PE=1 SV=7

Match to Query 2027: 780.447924 from(781.455200,1+) intensity(1382002.5000) rtinseconds(965.5585) index(2410)
Title: 20100530_methyl300mM_1.2681.2681.1
Data file D:\Panyue\Proteomic dataset\20100530_methyl300mM_1.RAW

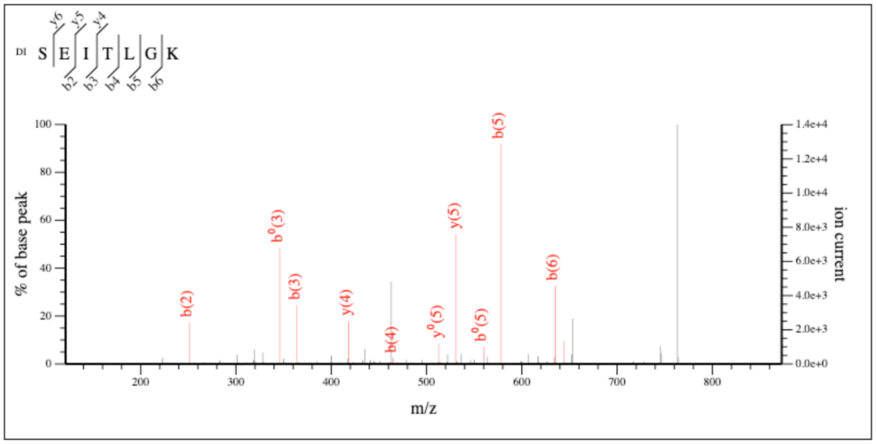

**Monoisotopic mass of neutral peptide Mr(calc):** 780.4864
**Fixed modifications:** Carbamidomethyl (C) (apply to specified residues or termini only)
**Variable modifications:**
**N-term :** Dimethyl:2H(6) (Protein N-term)
**Ions Score:** 34 **Expect:** 0.0046
**Matches :** 11/27 fragment ions using 24 most intense peaks (help)

| # | b | $b^0$ | Seq. | y | y* | $y^0$ | # |
|---|---|---|---|---|---|---|---|
| 1 | 122.1083 | 104.0977 | S | | | | 7 |
| 2 | ***251.1509*** | 233.1403 | E | 660.3927 | 643.3661 | 642.3821 | 6 |
| 3 | ***364.2349*** | 346.2244 | I | **531.3501** | 514.3235 | 513.3395 | 5 |
| 4 | ***465.2826*** | 447.2720 | T | **418.2660** | 401.2395 | 400.2554 | 4 |
| 5 | ***578.3667*** | 560.3561 | L | 317.2183 | 300.1918 | | 3 |
| 6 | ***635.3881*** | 617.3776 | G | 204.1343 | 187.1077 | | 2 |
| 7 | | | K | 147.1128 | 130.0863 | | 1 |

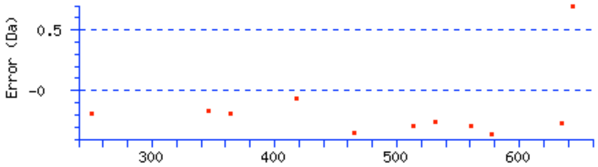

RMS error 619 ppm

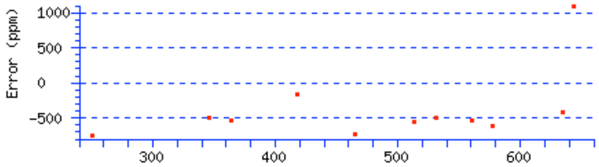

RMS error 619 ppm

MS/MS Fragmentation of SEITLGK
Found in P06169 in Uniprot_yeastS288C, Pyruvate decarboxylase isozyme 1 OS=Saccharomyces cerevisiae (strain ATCC 204508 / S288c) OX=559292 GN=PDC1 PE=1 SV=7

Match to Query 2028: 780.448901 from(781.456177,1+) intensity(579563.8125) rtinseconds(962.3724) index(2401)
Title: 20100530_methyl300mM_1.2671.2671.1
Data file D:\Panyue\Proteomic dataset\20100530_methyl300mM_1.RAW

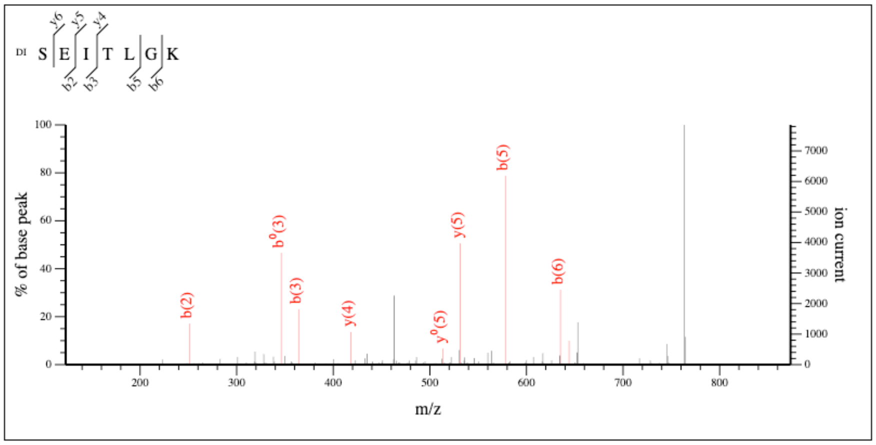


**Monoisotopic mass of neutral peptide Mr(calc):** 780.4864
**Fixed modifications:** Carbamidomethyl (C) (apply to specified residues or termini only)
**Variable modifications:**
**N-term :** Dimethyl:2H(6) (Protein N-term)
**Ions Score:** 29 **Expect:** 0.017
**Matches :** 9/27 fragment ions using 18 most intense peaks (help)

| # | b | $b^0$ | Seq. | y | y* | $y^0$ | # |
|---|---|---|---|---|---|---|---|
| 1 | 122.1083 | 104.0977 | S | | | | 7 |
| 2 | ***251.1509*** | 233.1403 | E | 660.3927 | 643.3661 | 642.3821 | 6 |
| 3 | ***364.2349*** | 346.2244 | I | **531.3501** | 514.3235 | 513.3395 | 5 |
| 4 | 465.2826 | 447.2720 | T | **418.2660** | 401.2395 | 400.2554 | 4 |
| 5 | ***578.3667*** | 560.3561 | L | 317.2183 | 300.1918 | | 3 |
| 6 | ***635.3881*** | 617.3776 | G | 204.1343 | 187.1077 | | 2 |
| 7 | | | K | 147.1128 | 130.0863 | | 1 |

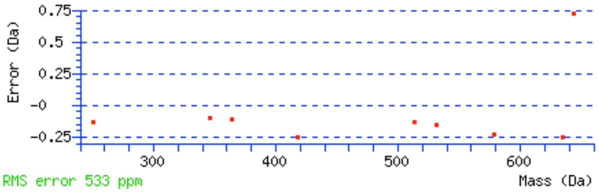


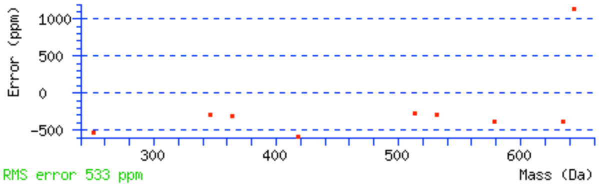


33. Sec3 (1 peptide in 300mM fraction 1)

MS/MS Fragmentation of **RSSKSPFK**
Found in **P33332** in **Uniprot_yeastS288C**, Exocyst complex component SEC3 OS=Saccharomyces cerevisiae (strain ATCC 204508 / S288c) OX=559292 GN=SEC3 PE=1 SV=1

Match to Query 4135: 986.585448 from(494.300000,2+) intensity(476399.1250) rtinseconds(1108.6913) index(2810)
Title: 20100530_methyl300mM_1.3126.3126.
Data file D:\Panyue\Proteomic dataset\20100530_methyl300mM_1.RAW

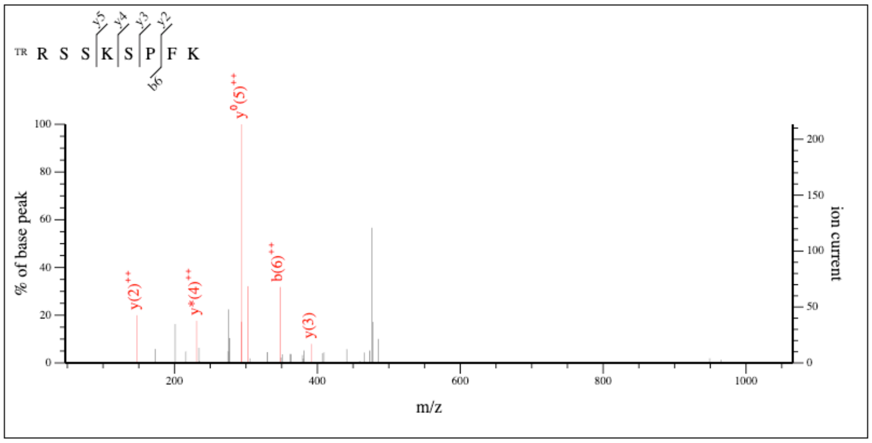


**Monoisotopic mass of neutral peptide Mr(calc):** 986.6223
**Fixed modifications:** Carbamidomethyl (C) (apply to specified residues or termini only)
**Variable modifications:**
**N-term :** Trimethyl:2H(9) (Protein N-term)
**Ions Score:** 27 **Expect:** 0.05
**Matches :** 11/76 fragment ions using 10 most intense peaks (help)

| # | b | b++ | b* | b*++ | b0 | b0++ | Seq. | y | y++ | y* | y*++ | y0 | y0++ | # |
|---|---|---|---|---|---|---|---|---|---|---|---|---|---|---|
| 1 | 208.2118 | 104.6096 | 191.1853 | 96.0963 | | | R | | | | | | | 8 |
| 2 | 295.2439 | 148.1256 | 278.2173 | 139.6123 | 277.2333 | 139.1203 | S | 780.4250 | 390.7162 | 763.3985 | 382.2029 | 762.4145 | 381.7109 | 7 |
| 3 | 382.2759 | 191.6416 | 365.2493 | 183.1283 | 364.2653 | 182.6363 | S | 693.3930 | 347.2001 | 676.3665 | 338.6869 | 675.3824 | 338.1949 | 6 |
| 4 | 510.3708 | 255.6891 | 493.3443 | 247.1758 | 492.3603 | 246.6838 | K | 606.3610 | 303.6841 | 589.3344 | 295.1709 | 588.3504 | 294.6788 | 5 |
| 5 | 597.4029 | 299.2051 | 580.3763 | 290.6918 | 579.3923 | 290.1998 | S | 478.2660 | 239.6366 | 461.2395 | 231.1234 | 460.2554 | 230.6314 | 4 |
| 6 | 694.4556 | 347.7315 | 677.4291 | 339.2182 | 676.4451 | 338.7262 | P | 391.2340 | 196.1206 | 374.2074 | 187.6074 | | | 3 |
| 7 | 841.5241 | 421.2657 | 824.4975 | 412.7524 | 823.5135 | 412.2604 | F | 294.1812 | 147.5942 | 277.1547 | 139.0810 | | | 2 |
| 8 | | | | | | | K | 147.1128 | 74.0600 | 130.0863 | 65.5468 | | | 1 |

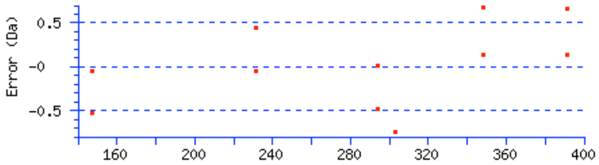


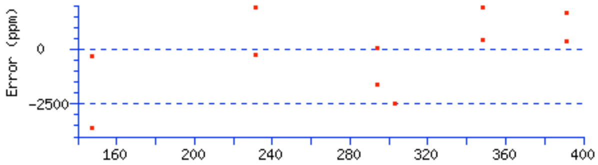


34. Rpn5 (1 peptide in 300mM fraction 1)

MS/MS Fragmentation of **SRDAPIK**
Found in **Q12250** in **Uniprot_yeastS288C**, 26S proteasome regulatory subunit RPN5 OS=Saccharomyces cerevisiae (strain ATCC 204508 / S288c) OX=559292 GN=RPN5 PE=1 SV=3

Match to Query 2184: 802.442724 from(803.450000,1+) intensity(170525.0156) rtinseconds(1658.3588) index(4312)
Title: 20100530_methyl300mM_1.4797.4797.1
Data file D:\Panyue\Proteomic dataset\20100530_methyl300mM_1.RAW

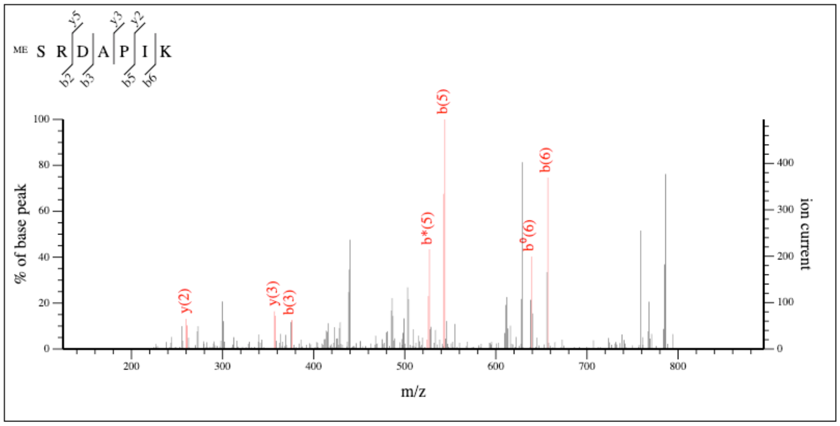


**Monoisotopic mass of neutral peptide Mr(calc):** 802.4740
**Fixed modifications:** Carbamidomethyl (C) (apply to specified residues or termini only)
**Variable modifications:**
**N-term :** Methyl:2H(3) (Protein N-term)
**Ions Score:** 22 **Expect:** 0.14
**Matches :** 13/31 fragment ions using 24 most intense peaks (help)

| # | b | b* | $b^0$ | Seq. | y | y* | $y^0$ | # |
|---|---|---|---|---|---|---|---|---|
| 1 | 105.0738 | | 87.0632 | S | | | | 7 |
| 2 | ***261.1749*** | 244.1483 | 243.1643 | R | 699.4148 | 682.3883 | 681.4042 | 6 |
| 3 | ***376.2018*** | 359.1753 | 358.1913 | D | ***543.3137*** | 526.2871 | 525.3031 | 5 |
| 4 | 447.2390 | 430.2124 | 429.2284 | A | 428.2867 | 411.2602 | | 4 |
| 5 | ***544.2917*** | 527.2652 | 526.2812 | P | ***357.2496*** | 340.2231 | | 3 |
| 6 | ***657.3758*** | 640.3492 | 639.3652 | I | ***260.1969*** | 243.1703 | | 2 |
| 7 | | | | K | 147.1128 | 130.0863 | | 1 |

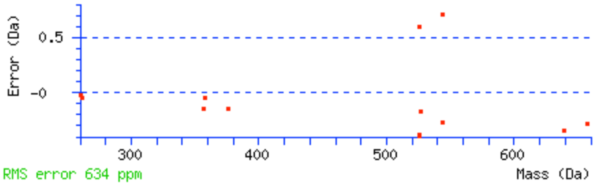


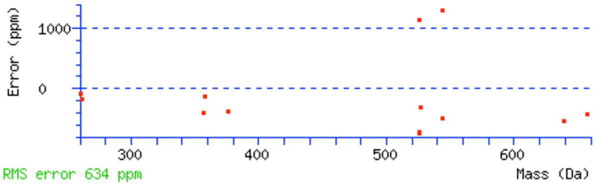


35. Nmd4 (1 peptide in 300mM fraction 1)

MS/MS Fragmentation of **TQYNFIIDASAFEKGLGNIKR**
Found in **Q12129** in **Uniprot_yeastS288C**, Nonsense-mediated decay protein 4 OS=Saccharomyces cerevisiae (strain ATCC 204508 / S288c) OX=559292 GN=NMD4 PE=1 SV=1

Match to Query 8775: 2401.358172 from(801.460000,3+) intensity(400408.9375) rtinseconds(1878.2484) index(4903)
Title: 20100530_methyl300mM_1.5455.5455.
Data file D:\Panyue\Proteomic dataset\20100530_methyl300mM_1.RAW

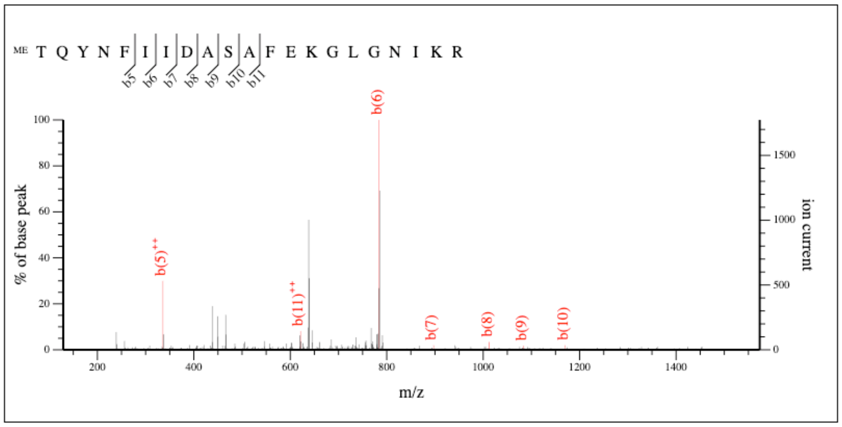


```
Monoisotopic mass of neutral peptide Mr(calc): 2401.2833
Fixed modifications: Carbamidomethyl (C) (apply to specified residues or termini only)
Variable modifications:
N-term : Methyl:2H(3) (Protein N-term)
Ions Score: 20  Expect: 0.027
Matches : 7/222 fragment ions using 13 most intense peaks   (help)
```

| # | b | b++ | b* | b*++ | b0 | b0++ | Seq. | y | y++ | y* | y*++ | y0 | y0++ | # |
|---|---|---|---|---|---|---|---|---|---|---|---|---|---|---|
| 1 | 119.0894 | 60.0484 | | | 101.0789 | 51.0431 | T | | | | | | | 21 |
| 2 | 247.1480 | 124.0776 | 230.1215 | 115.5644 | 229.1374 | 115.0724 | Q | 2284.2084 | 1142.6079 | 2267.1819 | 1134.0946 | 2266.1979 | 1133.6026 | 20 |
| 3 | 410.2113 | 205.6093 | 393.1848 | 197.0960 | 392.2008 | 196.6040 | Y | 2156.1499 | 1078.5786 | 2139.1233 | 1070.0653 | 2138.1393 | 1069.5733 | 19 |
| 4 | 524.2543 | 262.6308 | 507.2277 | 254.1175 | 506.2437 | 253.6255 | N | 1993.0865 | 997.0469 | 1976.0600 | 988.5336 | 1975.0760 | 988.0416 | 18 |
| 5 | 671.3227 | **336.1650** | 654.2961 | 327.6517 | 653.3121 | 327.1597 | F | 1879.0436 | 940.0254 | 1862.0171 | 931.5122 | 1861.0330 | 931.0202 | 17 |
| 6 | ***784.4067*** | 392.7070 | 767.3802 | 384.1937 | 766.3962 | 383.7017 | I | 1731.9752 | 866.4912 | 1714.9486 | 857.9780 | 1713.9646 | 857.4860 | 16 |
| 7 | ***897.4908*** | 449.2490 | 880.4643 | 440.7358 | 879.4802 | 440.2438 | I | 1618.8911 | 809.9492 | 1601.8646 | 801.4359 | 1600.8806 | 800.9439 | 15 |
| 8 | ***1012.5178*** | 506.7625 | 995.4912 | 498.2492 | 994.5072 | 497.7572 | D | 1505.8071 | 753.4072 | 1488.7805 | 744.8939 | 1487.7965 | 744.4019 | 14 |
| 9 | ***1083.5549*** | 542.2811 | 1066.5283 | 533.7678 | 1065.5443 | 533.2758 | A | 1390.7801 | 695.8937 | 1373.7536 | 687.3804 | 1372.7696 | 686.8884 | 13 |
| 10 | ***1170.5869*** | 585.7971 | 1153.5603 | 577.2838 | 1152.5763 | 576.7918 | S | 1319.7430 | 660.3751 | 1302.7165 | 651.8619 | 1301.7324 | 651.3699 | 12 |
| 11 | 1241.6240 | **621.3156** | 1224.5975 | 612.8024 | 1223.6134 | 612.3104 | A | 1232.7110 | 616.8591 | 1215.6844 | 608.3459 | 1214.7004 | 607.8538 | 11 |
| 12 | 1388.6924 | 694.8499 | 1371.6659 | 686.3366 | 1370.6819 | 685.8446 | F | 1161.6739 | 581.3406 | 1144.6473 | 572.8273 | 1143.6633 | 572.3353 | 10 |
| 13 | 1517.7350 | 759.3711 | 1500.7085 | 750.8579 | 1499.7245 | 750.3659 | E | 1014.6055 | 507.8064 | 997.5789 | 499.2931 | 996.5949 | 498.8011 | 9 |
| 14 | 1645.8300 | 823.4186 | 1628.8034 | 814.9054 | 1627.8194 | 814.4133 | K | 885.5629 | 443.2851 | 868.5363 | 434.7718 | | | 8 |
| 15 | 1702.8514 | 851.9294 | 1685.8249 | 843.4161 | 1684.8409 | 842.9241 | G | 757.4679 | 379.2376 | 740.4413 | 370.7243 | | | 7 |
| 16 | 1815.9355 | 908.4714 | 1798.9090 | 899.9581 | 1797.9249 | 899.4661 | L | 700.4464 | 350.7269 | 683.4199 | 342.2136 | | | 6 |
| 17 | 1872.9570 | 936.9821 | 1855.9304 | 928.4688 | 1854.9464 | 927.9768 | G | 587.3624 | 294.1848 | 570.3358 | 285.6715 | | | 5 |
| 18 | 1986.9999 | 994.0036 | 1969.9734 | 985.4903 | 1968.9893 | 984.9983 | N | 530.3409 | 265.6741 | 513.3144 | 257.1608 | | | 4 |
| 19 | 2100.0840 | 1050.5456 | 2083.0574 | 1042.0323 | 2082.0734 | 1041.5403 | I | 416.2980 | 208.6526 | 399.2714 | 200.1394 | | | 3 |
| 20 | 2228.1789 | 1114.5931 | 2211.1524 | 1106.0798 | 2210.1684 | 1105.5878 | K | 303.2139 | 152.1106 | 286.1874 | 143.5973 | | | 2 |
| 21 | | | | | | | R | 175.1190 | 88.0631 | 158.0924 | 79.5498 | | | 1 |

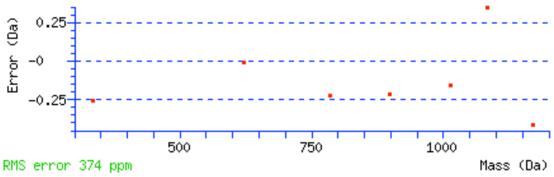


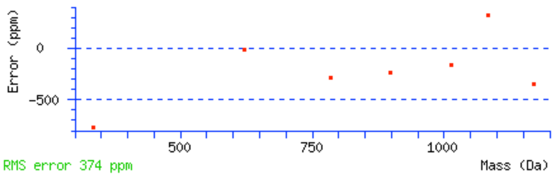


36. Eft1 (2 peptides in 300mM fraction 1)

MS/MS Fragmentation of VAFTVDQMRSLMDK
Found in P32324 in Uniprot_yeastS288C, Elongation factor 2 OS=Saccharomyces cerevisiae (strain ATCC 204508 / S288c) OX=559292 GN=EFT1 PE=1 SV=1

Match to Query 9324: 1706.872656 from(854.443604,2+) intensity(1093425.6250) rtinseconds(2570.8239) index(6784)
Title: 20100530_methyl300mM_1.7547.7547.2
Data file D:\Panyue\Proteomic dataset\20100530_methyl300mM_1.RAW

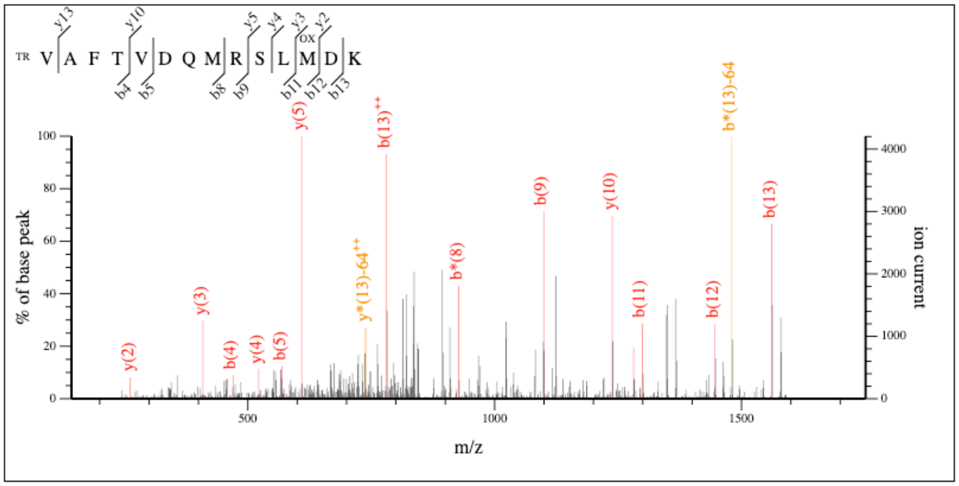


**Monoisotopic mass of neutral peptide Mr(calc):** 1706.9046
**Fixed modifications:** Carbamidomethyl (C) (apply to specified residues or termini only)
**Variable modifications:**
**N-term :** Trimethyl:2H(9) (Protein N-term)
**M12** : Oxidation (M), with neutral losses 0.0000(shown in table), 63.9983
**Ions Score:** 28 **Expect:** 0.026
**Matches :** 19/214 fragment ions using 29 most intense peaks (help)

| # | b | $b^{++}$ | b* | $b^{*++}$ | $b^0$ | $b^{0++}$ | Seq. | y | $y^{++}$ | y* | $y^{*++}$ | $y^0$ | $y^{0++}$ | # |
|---|---|---|---|---|---|---|---|---|---|---|---|---|---|---|
| 1 | 151.1791 | 76.0932 | | | | | V | | | | | | | 14 |
| 2 | 222.2162 | 111.6118 | | | | | A | 1557.7400 | 779.3736 | 1540.7134 | 770.8604 | 1539.7294 | 770.3683 | 13 |
| 3 | 369.2847 | 185.1460 | | | | | F | 1486.7029 | 743.8551 | 1469.6763 | 735.3418 | 1468.6923 | 734.8498 | 12 |
| 4 | ***470.3323*** | 235.6698 | | | 452.3218 | 226.6645 | T | 1339.6344 | 670.3209 | 1322.6079 | 661.8076 | 1321.6239 | 661.3156 | 11 |
| 5 | ***569.4008*** | 285.2040 | | | 551.3902 | 276.1987 | V | ***1238.5868*** | 619.7970 | 1221.5602 | 611.2837 | 1220.5762 | 610.7917 | 10 |
| 6 | 684.4277 | 342.7175 | | | 666.4171 | 333.7122 | D | 1139.5184 | 570.2628 | 1122.4918 | 561.7495 | 1121.5078 | 561.2575 | 9 |
| 7 | 812.4863 | 406.7468 | 795.4597 | 398.2335 | 794.4757 | 397.7415 | Q | 1024.4914 | 512.7493 | 1007.4649 | 504.2361 | 1006.4808 | 503.7441 | 8 |
| 8 | 943.5268 | 472.2670 | 926.5002 | 463.7537 | 925.5162 | 463.2617 | M | 896.4328 | 448.7201 | 879.4063 | 440.2068 | 878.4223 | 439.7148 | 7 |
| 9 | ***1099.6279*** | 550.3176 | 1082.6013 | 541.8043 | 1081.6173 | 541.3123 | R | 765.3924 | 383.1998 | 748.3658 | 374.6865 | 747.3818 | 374.1945 | 6 |
| 10 | 1186.6599 | 593.8336 | 1169.6333 | 585.3203 | 1168.6493 | 584.8283 | S | ***609.2912*** | 305.1493 | 592.2647 | 296.6360 | 591.2807 | 296.1440 | 5 |
| 11 | ***1299.7440*** | 650.3756 | 1282.7174 | 641.8623 | 1281.7334 | 641.3703 | L | ***522.2592*** | 261.6332 | 505.2327 | 253.1200 | 504.2486 | 252.6280 | 4 |
| 12 | ***1446.7794*** | 723.8933 | 1429.7528 | 715.3800 | 1428.7688 | 714.8880 | M | ***409.1751*** | 205.0912 | 392.1486 | 196.5779 | 391.1646 | 196.0859 | 3 |
| 13 | ***1561.8063*** | 781.4068 | 1544.7798 | 772.8935 | 1543.7957 | 772.4015 | D | ***262.1397*** | 131.5735 | 245.1132 | 123.0602 | 244.1292 | 122.5682 | 2 |
| 14 | | | | | | | K | 147.1128 | 74.0600 | 130.0863 | 65.5468 | | | 1 |

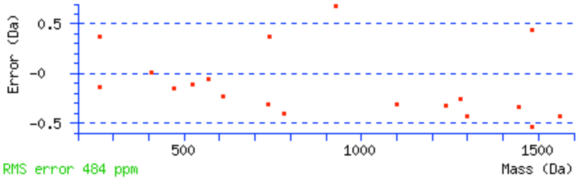


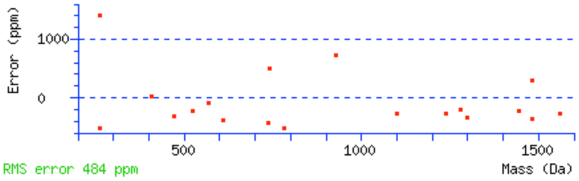

MS/MS Fragmentation of **VAFTVDQMRSLMDK**
Found in **P32324** in **Uniprot_yeastS288C**, Elongation factor 2 OS=Saccharomyces cerevisiae (strain ATCC 204508 / S288c) OX=559292 GN=EFT1 PE=1 SV=1

Match to Query 9325: 1706.873510 from(854.444031,2+) intensity(1613264.0000) rtinseconds(2573.1806) index(6790)
Title: 20100530_methyl300mM_1.7554.7554.2
Data file D:\Panyue\Proteomic dataset\20100530_methyl300mM_1.RAW

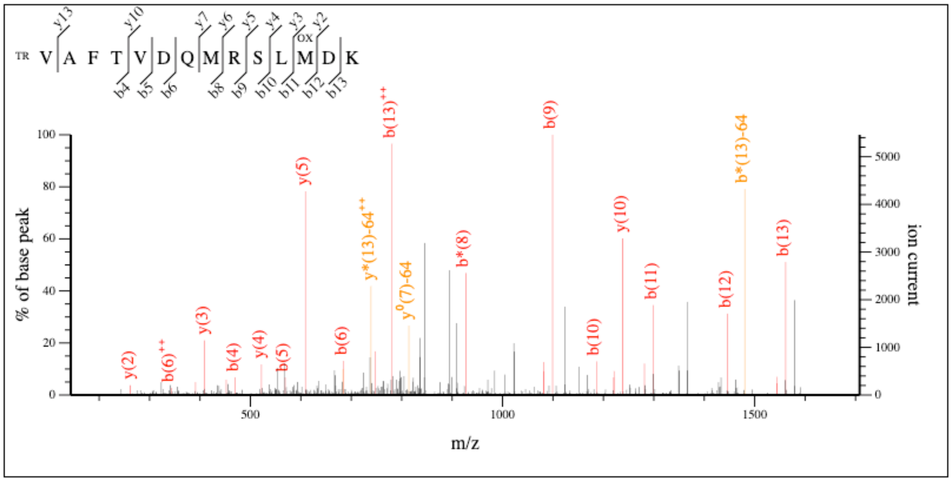


**Monoisotopic mass of neutral peptide Mr(calc):** 1706.9046
**Fixed modifications:** Carbamidomethyl (C) (apply to specified residues or termini only)
**Variable modifications:**
**N-term :** Trimethyl:2H(9) (Protein N-term)
**M12** : Oxidation (M), with neutral losses 0.0000(shown in table), 63.9983
**Ions Score:** 32 **Expect:** 0.011
**Matches :** 39/214 fragment ions using 56 most intense peaks (help)

| # | b | $b^{++}$ | $b^{*}$ | $b^{*++}$ | $b^{0}$ | $b^{0++}$ | Seq. | y | $y^{++}$ | $y^{*}$ | $y^{*++}$ | $y^{0}$ | $y^{0++}$ | # |
|---|---|---|---|---|---|---|---|---|---|---|---|---|---|---|
| 1 | 151.1791 | 76.0932 | | | | | V | | | | | | | 14 |
| 2 | 222.2162 | 111.6118 | | | | | A | 1557.7400 | 779.3736 | 1540.7134 | 770.8604 | 1539.7294 | 770.3683 | 13 |
| 3 | 369.2847 | 185.1460 | | | | | F | 1486.7029 | 743.8551 | 1469.6763 | 735.3418 | 1468.6923 | 734.8498 | 12 |
| 4 | ***470.3323*** | 235.6698 | | | 452.3218 | 226.6645 | T | 1339.6344 | 670.3209 | 1322.6079 | 661.8076 | 1321.6239 | 661.3156 | 11 |
| 5 | ***569.4008*** | 285.2040 | | | 551.3902 | 276.1987 | V | **1238.5868** | 619.7970 | 1221.5602 | 611.2837 | 1220.5762 | 610.7917 | 10 |
| 6 | ***684.4277*** | 342.7175 | | | 666.4171 | 333.7122 | D | 1139.5184 | 570.2628 | 1122.4918 | 561.7495 | 1121.5078 | 561.2575 | 9 |
| 7 | 812.4863 | 406.7468 | 795.4597 | 398.2335 | 794.4757 | 397.7415 | Q | 1024.4914 | 512.7493 | 1007.4649 | 504.2361 | 1006.4808 | 503.7441 | 8 |
| 8 | 943.5268 | 472.2670 | 926.5002 | 463.7537 | 925.5162 | 463.2617 | M | 896.4328 | 448.7201 | 879.4063 | 440.2068 | 878.4223 | 439.7148 | 7 |
| 9 | ***1099.6279*** | 550.3176 | 1082.6013 | 541.8043 | 1081.6173 | 541.3123 | R | 765.3924 | 383.1998 | 748.3658 | 374.6865 | 747.3818 | 374.1945 | 6 |
| 10 | ***1186.6599*** | 593.8336 | 1169.6333 | 585.3203 | 1168.6493 | 584.8283 | S | **609.2912** | 305.1493 | 592.2647 | 296.6360 | 591.2807 | 296.1440 | 5 |
| 11 | ***1299.7440*** | 650.3756 | 1282.7174 | 641.8623 | 1281.7334 | 641.3703 | L | **522.2592** | 261.6332 | 505.2327 | 253.1200 | 504.2486 | 252.6280 | 4 |
| 12 | ***1446.7794*** | 723.8933 | 1429.7528 | 715.3800 | 1428.7688 | 714.8880 | M | **409.1751** | 205.0912 | 392.1486 | 196.5779 | 391.1646 | 196.0859 | 3 |
| 13 | ***1561.8063*** | 781.4068 | 1544.7798 | 772.8935 | 1543.7957 | 772.4015 | D | **262.1397** | 131.5735 | 245.1132 | 123.0602 | 244.1292 | 122.5682 | 2 |
| 14 | | | | | | | K | 147.1128 | 74.0600 | 130.0863 | 65.5468 | | | 1 |

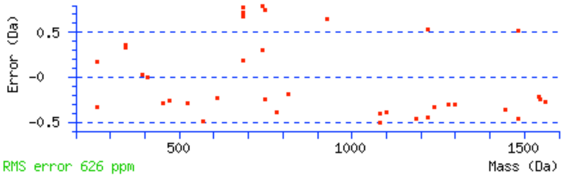


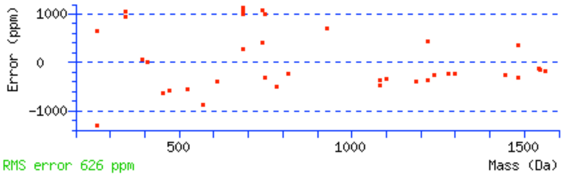


37. YJL133C-A (1 peptide in 300mM fraction 2)

MS/MS Fragmentation of **IAQSTRLAAAVSSSAASAGVSRIAASAMASTIFK**
Found in **Q3E7A3** in **Uniprot_yeastS288C**, Uncharacterized protein YJL133C-A OS=Saccharomyces cerevisiae (strain ATCC 204508 / S288c) OX=559292 GN=YJL133C-A PE=1 SV=1

Match to Query 17288: 3318.820016 from(830.712280,4+) intensity(1376235.5000) rtinseconds(4110.2645) index(10766)
Title: 20100530_methyl300mM_2.11977.11977.4
Data file D:\Panyue\Proteomic dataset\20100530_methyl300mM_2.RAW

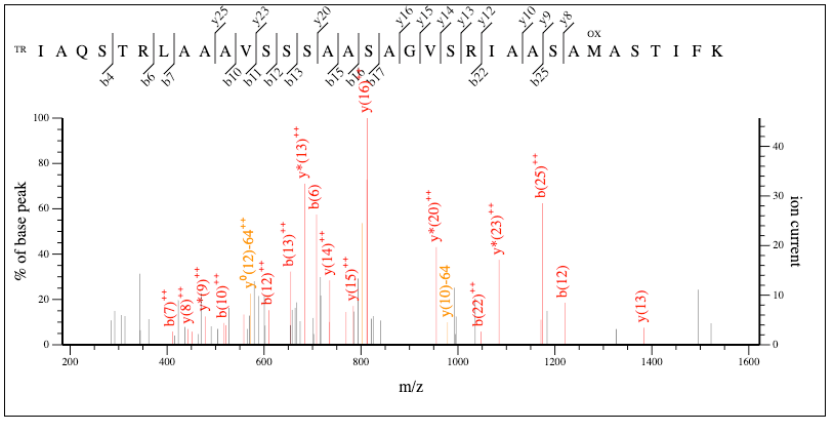


**Monoisotopic mass of neutral peptide Mr(calc):** 3318.8279
**Fixed modifications:** Carbamidomethyl (C) (apply to specified residues or termini only)
**Variable modifications:**
**N-term :** Trimethyl:2H(9) (Protein N-term)
**M28 :** Oxidation (M), with neutral losses 0.0000(shown in table), 63.9983
**Ions Score:** 21 **Expect:** 0.036
**Matches :** 32/578 fragment ions using 58 most intense peaks (help)

| # | b | b++ | b* | b*++ | b0 | b0++ | Seq. | y | y++ | y* | y*++ | y0 | y0++ | # |
|---|---|---|---|---|---|---|---|---|---|---|---|---|---|---|
| 1 | 165.1948 | 83.1010 | | | | | I | | | | | | | 34 |
| 2 | 236.2319 | 118.6196 | | | | | A | 3155.6477 | 1578.3275 | 3138.6212 | 1569.8142 | 3137.6372 | 1569.3222 | 33 |
| 3 | 364.2905 | 182.6489 | 347.2639 | 174.1356 | | | Q | 3084.6106 | 1542.8090 | 3067.5841 | 1534.2957 | 3066.6001 | 1533.8037 | 32 |
| 4 | **451.3225** | 226.1649 | 434.2960 | 217.6516 | 433.3119 | 217.1596 | S | 2956.5520 | 1478.7797 | 2939.5255 | 1470.2664 | 2938.5415 | 1469.7744 | 31 |
| 5 | 552.3702 | 276.6887 | 535.3436 | 268.1755 | 534.3596 | 267.6834 | T | 2869.5200 | 1435.2636 | 2852.4935 | 1426.7504 | 2851.5095 | 1426.2584 | 30 |
| 6 | **708.4713** | 354.7393 | 691.4447 | 346.2260 | 690.4607 | 345.7340 | R | 2768.4723 | 1384.7398 | 2751.4458 | 1376.2265 | 2750.4618 | 1375.7345 | 29 |
| 7 | 821.5554 | ***411.2813*** | 804.5288 | 402.7680 | 803.5448 | 402.2760 | L | 2612.3712 | 1306.6893 | 2595.3447 | 1298.1760 | 2594.3607 | 1297.6840 | 28 |
| 8 | 892.5925 | 446.7999 | 875.5659 | 438.2866 | 874.5819 | 437.7946 | A | 2499.2872 | 1250.1472 | 2482.2606 | 1241.6339 | 2481.2766 | 1241.1419 | 27 |
| 9 | 963.6296 | 482.3184 | 946.6030 | 473.8052 | 945.6190 | 473.3131 | A | 2428.2501 | 1214.6287 | 2411.2235 | 1206.1154 | 2410.2395 | 1205.6234 | 26 |
| 10 | 1034.6667 | ***517.8370*** | 1017.6401 | 509.3237 | 1016.6561 | 508.8317 | A | 2357.2129 | 1179.1101 | 2340.1864 | 1170.5968 | 2339.2024 | 1170.1048 | 25 |
| 11 | 1133.7351 | 567.3712 | 1116.7086 | 558.8579 | 1115.7245 | 558.3659 | V | 2286.1758 | 1143.5915 | 2269.1493 | 1135.0783 | 2268.1653 | 1134.5863 | 24 |
| 12 | **1220.7671** | ***610.8872*** | 1203.7406 | 602.3739 | 1202.7566 | 601.8819 | S | 2187.1074 | 1094.0573 | 2170.0809 | 1085.5441 | 2169.0968 | 1085.0521 | 23 |
| 13 | 1307.7992 | ***654.4032*** | 1290.7726 | 645.8899 | 1289.7886 | 645.3979 | S | 2100.0754 | 1050.5413 | 2083.0488 | 1042.0281 | 2082.0648 | 1041.5360 | 22 |
| 14 | 1394.8312 | 697.9192 | 1377.8046 | 689.4060 | 1376.8206 | 688.9140 | S | 2013.0434 | 1007.0253 | 1996.0168 | 998.5120 | 1995.0328 | 998.0200 | 21 |
| 15 | 1465.8683 | ***733.4378*** | 1448.8418 | 724.9245 | 1447.8577 | 724.4325 | A | 1926.0113 | 963.5093 | 1908.9848 | 954.9960 | 1908.0008 | 954.5040 | 20 |
| 16 | 1536.9054 | ***768.9563*** | 1519.8789 | 760.4431 | 1518.8949 | 759.9511 | A | 1854.9742 | 927.9907 | 1837.9477 | 919.4775 | 1836.9636 | 918.9855 | 19 |
| 17 | 1623.9375 | ***812.4724*** | 1606.9109 | 803.9591 | 1605.9269 | 803.4671 | S | 1783.9371 | 892.4722 | 1766.9105 | 883.9589 | 1765.9265 | 883.4669 | 18 |
| 18 | 1694.9746 | 847.9909 | 1677.9480 | 839.4776 | 1676.9640 | 838.9856 | A | 1696.9051 | 848.9562 | 1679.8785 | 840.4429 | 1678.8945 | 839.9509 | 17 |
| 19 | 1751.9960 | 876.5017 | 1734.9695 | 867.9884 | 1733.9855 | 867.4964 | G | 1625.8680 | ***813.4376*** | 1608.8414 | 804.9243 | 1607.8574 | 804.4323 | 16 |
| 20 | 1851.0644 | 926.0359 | 1834.0379 | 917.5226 | 1833.0539 | 917.0306 | V | 1568.8465 | ***784.9269*** | 1551.8199 | 776.4136 | 1550.8359 | 775.9216 | 15 |
| 21 | 1938.0965 | 969.5519 | 1921.0699 | 961.0386 | 1920.0859 | 960.5466 | S | 1469.7781 | ***735.3927*** | 1452.7515 | 726.8794 | 1451.7675 | 726.3874 | 14 |
| 22 | 2094.1976 | ***1047.6024*** | 2077.1710 | 1039.0892 | 2076.1870 | 1038.5971 | R | 1382.7460 | 691.8767 | 1365.7195 | 683.3634 | 1364.7355 | 682.8714 | 13 |
| 23 | 2207.2816 | 1104.1445 | 2190.2551 | 1095.6312 | 2189.2711 | 1095.1392 | I | 1226.6449 | 613.8261 | 1209.6184 | 605.3128 | 1208.6344 | 604.8208 | 12 |
| 24 | 2278.3188 | 1139.6630 | 2261.2922 | 1131.1497 | 2260.3082 | 1130.6577 | A | 1113.5609 | 557.2841 | 1096.5343 | 548.7708 | 1095.5503 | 548.2788 | 11 |
| 25 | 2349.3559 | ***1175.1816*** | 2332.3293 | 1166.6683 | 2331.3453 | 1166.1763 | A | 1042.5238 | ***521.7655*** | 1025.4972 | 513.2522 | 1024.5132 | 512.7602 | 10 |
| 26 | 2436.3879 | 1218.6976 | 2419.3614 | 1210.1843 | 2418.3773 | 1209.6923 | S | 971.4866 | 486.2470 | 954.4601 | 477.7337 | 953.4761 | 477.2417 | 9 |
| 27 | 2507.4250 | 1254.2161 | 2490.3985 | 1245.7029 | 2489.4145 | 1245.2109 | A | 884.4546 | ***442.7309*** | 867.4281 | 434.2177 | 866.4441 | 433.7257 | 8 |

| | | | | | | | | | | | | | | |
|---|---|---|---|---|---|---|---|---|---|---|---|---|---|---|
| 28 | 2654.4604 | 1327.7338 | 2637.4339 | 1319.2206 | 2636.4499 | 1318.7286 | M | 813.4175 | 407.2124 | 796.3910 | 398.6991 | 795.4069 | 398.2071 | 7 |
| 29 | 2725.4975 | 1363.2524 | 2708.4710 | 1354.7391 | 2707.4870 | 1354.2471 | A | 666.3821 | 333.6947 | 649.3556 | 325.1814 | 648.3715 | 324.6894 | 6 |
| 30 | 2812.5296 | 1406.7684 | 2795.5030 | 1398.2551 | 2794.5190 | 1397.7631 | S | 595.3450 | 298.1761 | 578.3184 | 289.6629 | 577.3344 | 289.1708 | 5 |
| 31 | 2913.5772 | 1457.2923 | 2896.5507 | 1448.7790 | 2895.5667 | 1448.2870 | T | 508.3130 | 254.6601 | 491.2864 | 246.1468 | 490.3024 | 245.6548 | 4 |
| 32 | 3026.6613 | 1513.8343 | 3009.6348 | 1505.3210 | 3008.6507 | 1504.8290 | I | 407.2653 | 204.1363 | 390.2387 | 195.6230 | | | 3 |
| 33 | 3173.7297 | 1587.3685 | 3156.7032 | 1578.8552 | 3155.7192 | 1578.3632 | F | 294.1812 | 147.5942 | 277.1547 | 139.0810 | | | 2 |
| 34 | | | | | | | K | 147.1128 | 74.0600 | 130.0863 | 65.5468 | | | 1 |

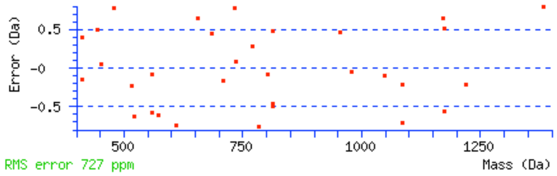


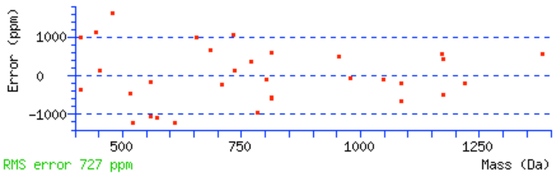


### 38. PDC6 (2 peptides in 300mM fraction2)

MS/MS Fragmentation of SEITLGK
Found in P26263 in Uniprot_yeastS288C, Pyruvate decarboxylase isozyme 3 OS=Saccharomyces cerevisiae (strain ATCC 204508 / S288c) OX=559292 GN=PDC6 PE=1 SV=3

Match to Query 2328: 780.448962 from(781.456238,1+) intensity(548515.6250) rtinseconds(1050.5666) index(2619)
Title: 20100530_methyl300mM_2.2915.2915.1
Data file D:\Panyue\Proteomic dataset\20100530_methyl300mM_2.RAW

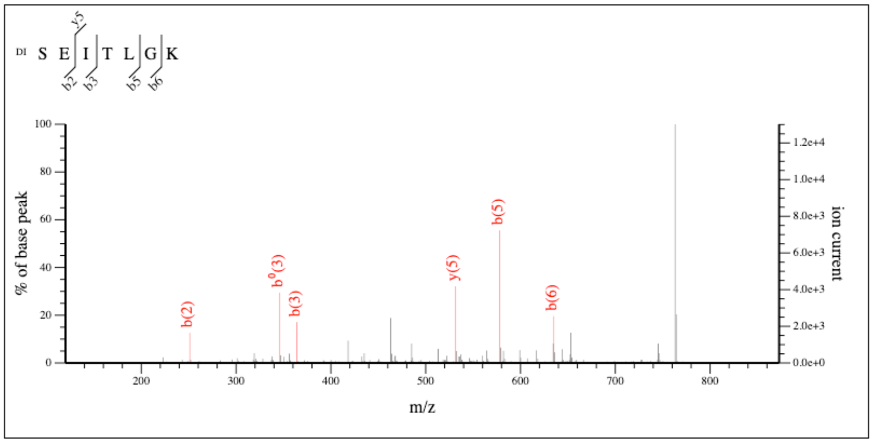


**Monoisotopic mass of neutral peptide Mr(calc):** 780.4864
**Fixed modifications:** Carbamidomethyl (C) (apply to specified residues or termini only)
**Variable modifications:**
**N-term :** Dimethyl:2H(6) (Protein N-term)
**Ions Score:** 26 **Expect:** 0.03
**Matches :** 6/27 fragment ions using 12 most intense peaks (help)

| # | b | $b^0$ | Seq. | y | y* | $y^0$ | # |
|---|---|---|---|---|---|---|---|
| 1 | 122.1083 | 104.0977 | S | | | | 7 |
| 2 | ***251.1509*** | 233.1403 | E | 660.3927 | 643.3661 | 642.3821 | 6 |
| 3 | ***364.2349*** | 346.2244 | I | 531.3501 | 514.3235 | 513.3395 | 5 |
| 4 | 465.2826 | 447.2720 | T | 418.2660 | 401.2395 | 400.2554 | 4 |
| 5 | ***578.3667*** | 560.3561 | L | 317.2183 | 300.1918 | | 3 |
| 6 | ***635.3881*** | 617.3776 | G | 204.1343 | 187.1077 | | 2 |
| 7 | | | K | 147.1128 | 130.0863 | | 1 |

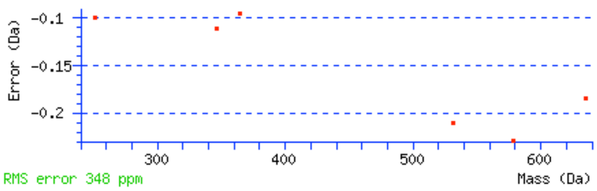


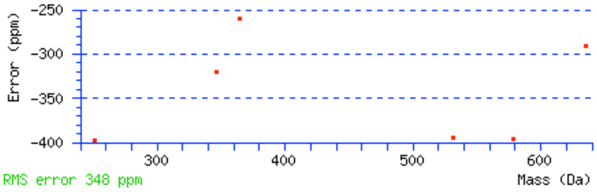

MS/MS Fragmentation of **SEITLGK**
Found in **P26263** in **Uniprot_yeastS288C**, Pyruvate decarboxylase isozyme 3 OS=Saccharomyces cerevisiae (strain ATCC 204508 / S288c) OX=559292 GN=PDC6 PE=1 SV=3

Match to Query 2329: 780.449450 from(781.456726,1+) intensity(1119942.0000) rtinseconds(1053.19) index(2626)
Title: 20100530_methyl300mM_2.2923.2923.1
Data file D:\Panyue\Proteomic dataset\20100530_methyl300mM_2.RAW

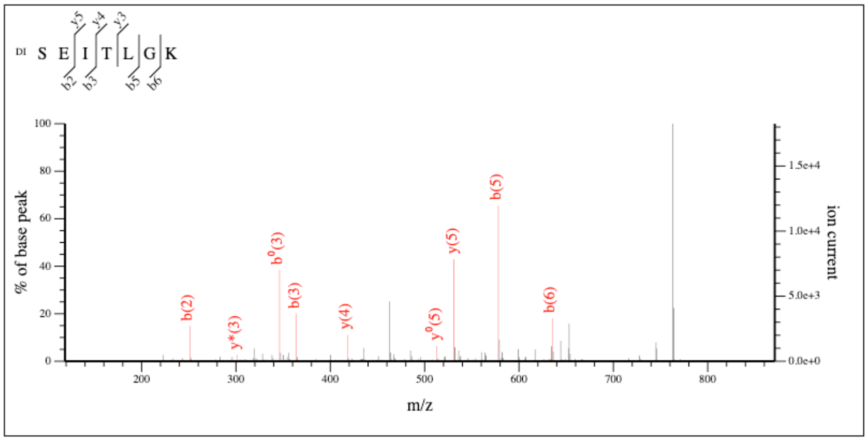


**Monoisotopic mass of neutral peptide Mr(calc):** 780.4864
**Fixed modifications:** Carbamidomethyl (C) (apply to specified residues or termini only)
**Variable modifications:**
**N-term :** Dimethyl:2H(6) (Protein N-term)
**Ions Score:** 27 **Expect:** 0.027
**Matches :** 9/27 fragment ions using 18 most intense peaks (help)

| # | b | $b^0$ | Seq. | y | y* | $y^0$ | # |
|---|---|---|---|---|---|---|---|
| 1 | 122.1083 | 104.0977 | S | | | | 7 |
| 2 | ***251.1509*** | 233.1403 | E | 660.3927 | 643.3661 | 642.3821 | 6 |
| 3 | ***364.2349*** | 346.2244 | I | **531.3501** | 514.3235 | 513.3395 | 5 |
| 4 | 465.2826 | 447.2720 | T | **418.2660** | 401.2395 | 400.2554 | 4 |
| 5 | ***578.3667*** | 560.3561 | L | 317.2183 | 300.1918 | | 3 |
| 6 | ***635.3881*** | 617.3776 | G | 204.1343 | 187.1077 | | 2 |
| 7 | | | K | 147.1128 | 130.0863 | | 1 |

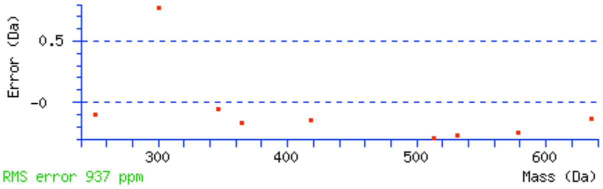


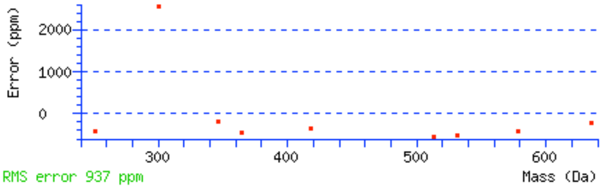


39. PDC5 (2 peptides in 300mM fraction2)

MS/MS Fragmentation of **SEITLGK**
Found in **P16467** in **Uniprot_yeastS288C**, Pyruvate decarboxylase isozyme 2 OS=Saccharomyces cerevisiae (strain ATCC 204508 / S288c) OX=559292 GN=PDC5 PE=1 SV=4

Match to Query 2328: 780.448962 from(781.456238,1+) intensity(548515.6250) rtinseconds(1050.5666) index(2619)
Title: 20100530_methyl300mM_2.2915.2915.1
Data file D:\Panyue\Proteomic dataset\20100530_methyl300mM_2.RAW

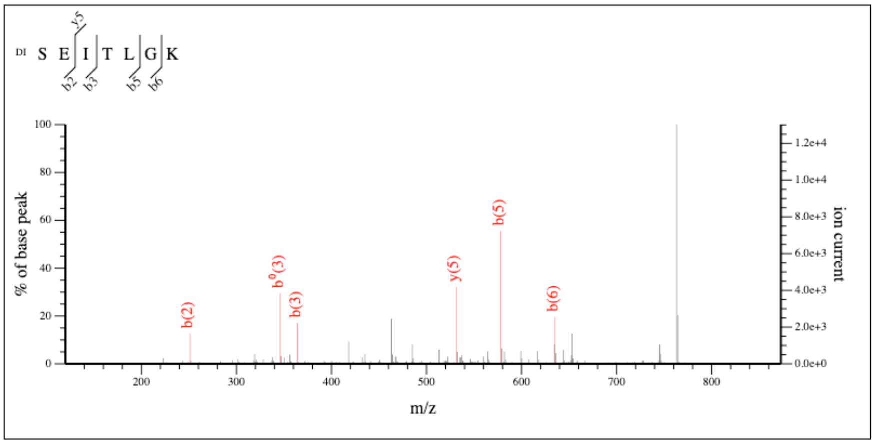

**Monoisotopic mass of neutral peptide Mr(calc):** 780.4864
**Fixed modifications:** Carbamidomethyl (C) (apply to specified residues or termini only)
**Variable modifications:**
**N-term :** Dimethyl:2H(6) (Protein N-term)
**Ions Score:** 26 **Expect:** 0.03
**Matches :** 6/27 fragment ions using 12 most intense peaks (help)

| # | b | $b^0$ | Seq. | y | y* | $y^0$ | # |
|---|---|---|---|---|---|---|---|
| 1 | 122.1083 | 104.0977 | S | | | | 7 |
| 2 | ***251.1509*** | 233.1403 | E | 660.3927 | 643.3661 | 642.3821 | 6 |
| 3 | ***364.2349*** | 346.2244 | I | 531.3501 | 514.3235 | 513.3395 | 5 |
| 4 | 465.2826 | 447.2720 | T | 418.2660 | 401.2395 | 400.2554 | 4 |
| 5 | ***578.3667*** | 560.3561 | L | 317.2183 | 300.1918 | | 3 |
| 6 | ***635.3881*** | 617.3776 | G | 204.1343 | 187.1077 | | 2 |
| 7 | | | K | 147.1128 | 130.0863 | | 1 |

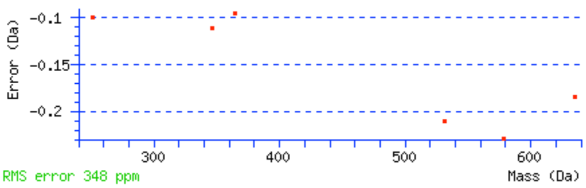

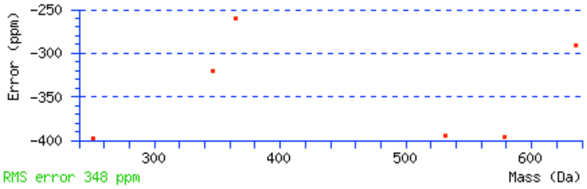

MS/MS Fragmentation of **SEITLGK**
Found in **P16467** in **Uniprot_yeastS288C**, Pyruvate decarboxylase isozyme 2 OS=Saccharomyces cerevisiae (strain ATCC 204508 / S288c) OX=559292 GN=PDC5 PE=1 SV=4

Match to Query 2329: 780.449450 from(781.456726,1+) intensity(1119942.0000) rtinseconds(1053.19) index(2626)
Title: 20100530_methyl300mM_2.2923.2923.1
Data file D:\Panyue\Proteomic dataset\20100530_methyl300mM_2.RAW

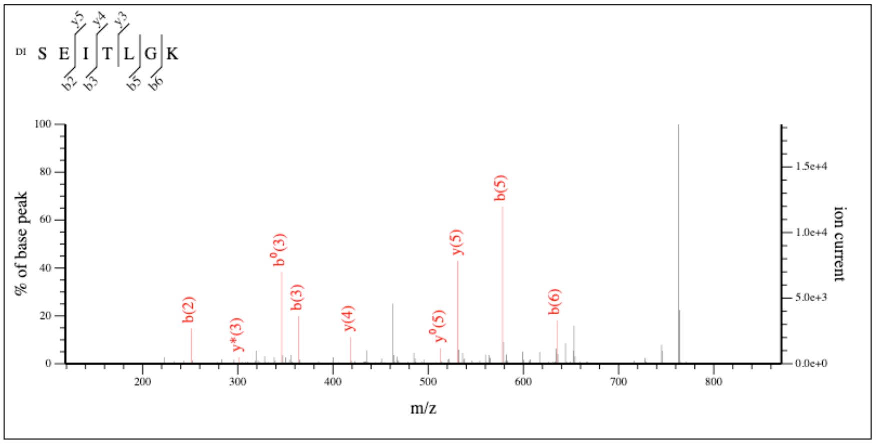

**Monoisotopic mass of neutral peptide Mr(calc):** 780.4864
**Fixed modifications:** Carbamidomethyl (C) (apply to specified residues or termini only)
**Variable modifications:**
**N-term :** Dimethyl:2H(6) (Protein N-term)
**Ions Score:** 27 **Expect:** 0.027
**Matches :** 9/27 fragment ions using 18 most intense peaks (help)

| # | b | b0 | Seq. | y | y* | y0 | # |
|---|---|---|---|---|---|---|---|
| 1 | 122.1083 | 104.0977 | S | | | | 7 |
| 2 | *251.1509* | 233.1403 | E | 660.3927 | 643.3661 | 642.3821 | 6 |
| 3 | *364.2349* | 346.2244 | I | **531.3501** | 514.3235 | 513.3395 | 5 |
| 4 | 465.2826 | 447.2720 | T | **418.2660** | 401.2395 | 400.2554 | 4 |
| 5 | *578.3667* | 560.3561 | L | 317.2183 | 300.1918 | | 3 |
| 6 | *635.3881* | 617.3776 | G | 204.1343 | 187.1077 | | 2 |
| 7 | | | K | 147.1128 | 130.0863 | | 1 |

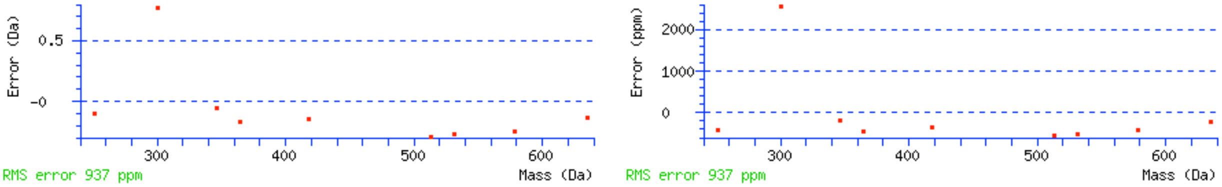

40. PDC1 (2 peptides in 300mM fraction 2)

MS/MS Fragmentation of **SEITLGK**
Found in **P06169** in **Uniprot_yeastS288C**, Pyruvate decarboxylase isozyme 1 OS=Saccharomyces cerevisiae (strain ATCC 204508 / S288c) OX=559292 GN=PDC1 PE=1 SV=7

Match to Query 2328: 780.448962 from(781.456238,1+) intensity(548515.6250) rtinseconds(1050.5666) index(2619)
Title: 20100530_methyl300mM_2.2915.2915.1
Data file D:\Panyue\Proteomic dataset\20100530_methyl300mM_2.RAW

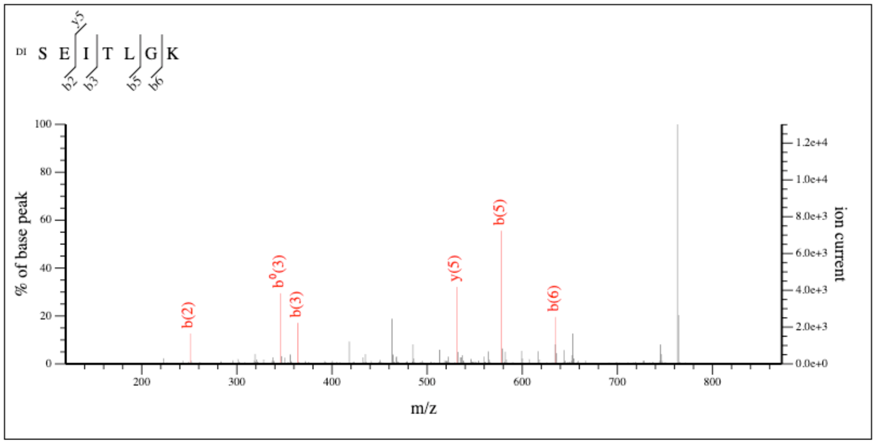


**Monoisotopic mass of neutral peptide Mr(calc):** 780.4864
**Fixed modifications:** Carbamidomethyl (C) (apply to specified residues or termini only)
**Variable modifications:**
**N-term :** Dimethyl:2H(6) (Protein N-term)
**Ions Score:** 26 **Expect:** 0.03
**Matches :** 6/27 fragment ions using 12 most intense peaks (help)

| # | b | $b^0$ | Seq. | y | y* | $y^0$ | # |
|---|---|---|---|---|---|---|---|
| 1 | 122.1083 | 104.0977 | S | | | | 7 |
| 2 | ***251.1509*** | 233.1403 | E | 660.3927 | 643.3661 | 642.3821 | 6 |
| 3 | ***364.2349*** | 346.2244 | I | 531.3501 | 514.3235 | 513.3395 | 5 |
| 4 | 465.2826 | 447.2720 | T | 418.2660 | 401.2395 | 400.2554 | 4 |
| 5 | ***578.3667*** | 560.3561 | L | 317.2183 | 300.1918 | | 3 |
| 6 | ***635.3881*** | 617.3776 | G | 204.1343 | 187.1077 | | 2 |
| 7 | | | K | 147.1128 | 130.0863 | | 1 |

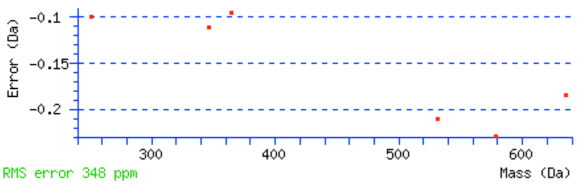


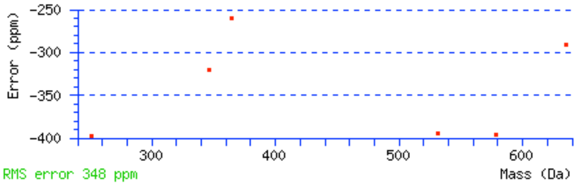

MS/MS Fragmentation of **SEITLGK**
Found in **P06169** in **Uniprot_yeastS288C**, Pyruvate decarboxylase isozyme 1 OS=Saccharomyces cerevisiae (strain ATCC 204508 / S288c) OX=559292 GN=PDC1 PE=1 SV=7

Match to Query 2329: 780.449450 from(781.456726,1+) intensity(1119942.0000) rtinseconds(1053.19) index(2626)
Title: 20100530_methyl300mM_2.2923.2923.1
Data file D:\Panyue\Proteomic dataset\20100530_methyl300mM_2.RAW

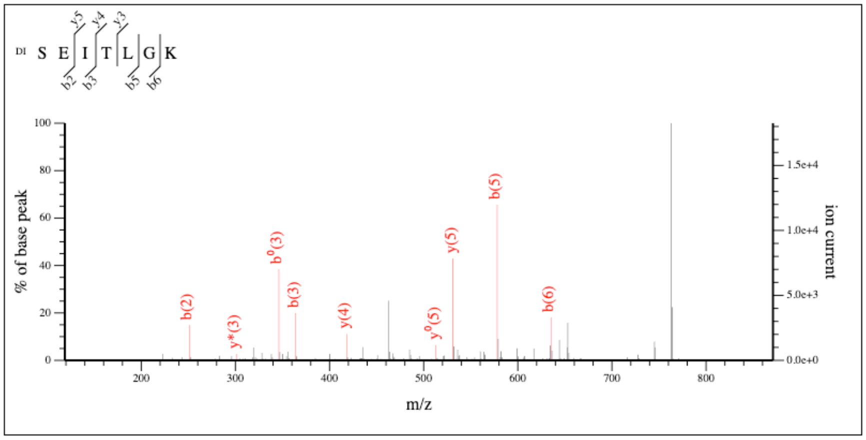


**Monoisotopic mass of neutral peptide Mr(calc):** 780.4864
**Fixed modifications:** Carbamidomethyl (C) (apply to specified residues or termini only)
**Variable modifications:**
**N-term :** Dimethyl:2H(6) (Protein N-term)
**Ions Score:** 27 **Expect:** 0.027
**Matches :** 9/27 fragment ions using 18 most intense peaks (help)

| # | b | $b^0$ | Seq. | y | y* | $y^0$ | # |
|---|---|---|---|---|---|---|---|
| 1 | 122.1083 | 104.0977 | S | | | | 7 |
| 2 | ***251.1509*** | 233.1403 | E | 660.3927 | 643.3661 | 642.3821 | 6 |
| 3 | ***364.2349*** | 346.2244 | I | **531.3501** | 514.3235 | 513.3395 | 5 |
| 4 | 465.2826 | 447.2720 | T | **418.2660** | 401.2395 | 400.2554 | 4 |
| 5 | ***578.3667*** | 560.3561 | L | 317.2183 | 300.1918 | | 3 |
| 6 | ***635.3881*** | 617.3776 | G | 204.1343 | 187.1077 | | 2 |
| 7 | | | K | 147.1128 | 130.0863 | | 1 |

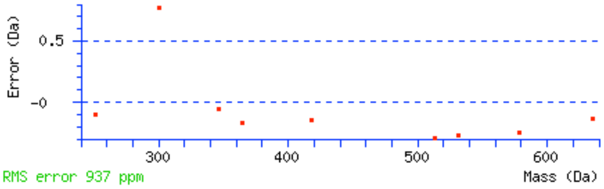


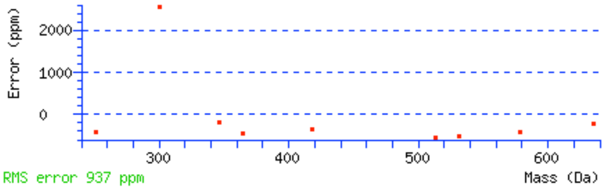


41. Nop14 (2 peptides in 300mM fraction 2)

MS/MS Fragmentation of **AGSQLKNLK**
Found in **Q99207** in **Uniprot_yeastS288C**, Nucleolar complex protein 14 OS=Saccharomyces cerevisiae (strain ATCC 204508 / S288c) OX=559292 GN=NOP14 PE=1 SV=1

Match to Query 4597: 1008.669238 from(1009.676514,1+) intensity(512111.0000) rtinseconds(1692.3747) index(4351)
Title: 20100530_methyl300mM_2.4842.4842.1
Data file D:\Panyue\Proteomic dataset\20100530_methyl300mM_2.RAW

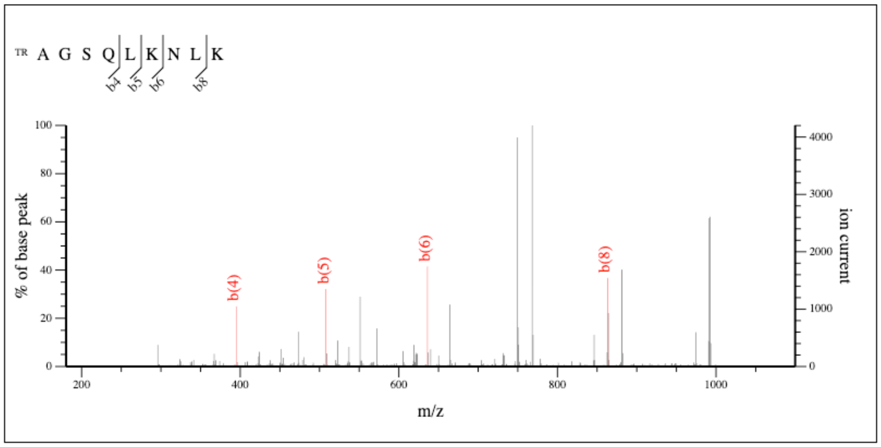


**Monoisotopic mass of neutral peptide Mr(calc):** 1008.6641
**Fixed modifications:** Carbamidomethyl (C) (apply to specified residues or termini only)
**Variable modifications:**
**N-term :** Trimethyl:2H(9) (Protein N-term)
**Ions Score:** 23 **Expect:** 0.011
**Matches :** 4/37 fragment ions using 8 most intense peaks (help)

| # | b | b* | $b^0$ | Seq. | y | y* | $y^0$ | # |
|---|---|---|---|---|---|---|---|---|
| 1 | 123.1478 | | | A | | | | 9 |
| 2 | 180.1693 | | | G | 887.5309 | 870.5043 | 869.5203 | 8 |
| 3 | 267.2013 | | 249.1908 | S | 830.5094 | 813.4829 | 812.4989 | 7 |
| 4 | ***395.2599*** | 378.2334 | 377.2493 | Q | 743.4774 | 726.4509 | | 6 |
| 5 | ***508.3440*** | 491.3174 | 490.3334 | L | 615.4188 | 598.3923 | | 5 |
| 6 | ***636.4389*** | 619.4124 | 618.4284 | K | 502.3348 | 485.3082 | | 4 |
| 7 | 750.4819 | 733.4553 | 732.4713 | N | 374.2398 | 357.2132 | | 3 |
| 8 | ***863.5659*** | 846.5394 | 845.5554 | L | 260.1969 | 243.1703 | | 2 |
| 9 | | | | K | 147.1128 | 130.0863 | | 1 |

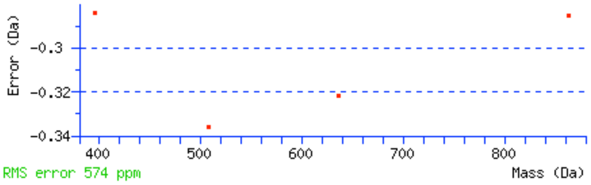


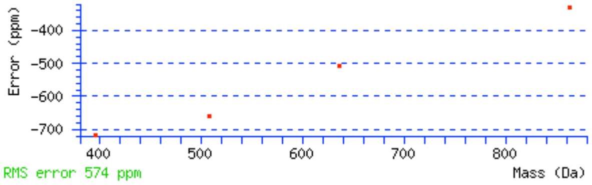

MS/MS Fragmentation of **AGSQLKNLK**
Found in **Q99207** in **Uniprot_yeastS288C**, Nucleolar complex protein 14 OS=Saccharomyces cerevisiae (strain ATCC 204508 / S288c) OX=559292 GN=NOP14 PE=1 SV=1

Match to Query 4602: 1008.670397 from(1009.677673,1+) intensity(1075322.1250) rtinseconds(1695.2647) index(4359)
Title: 20100530_methyl300mM_2.4851.4851.1
Data file D:\Panyue\Proteomic dataset\20100530_methyl300mM_2.RAW

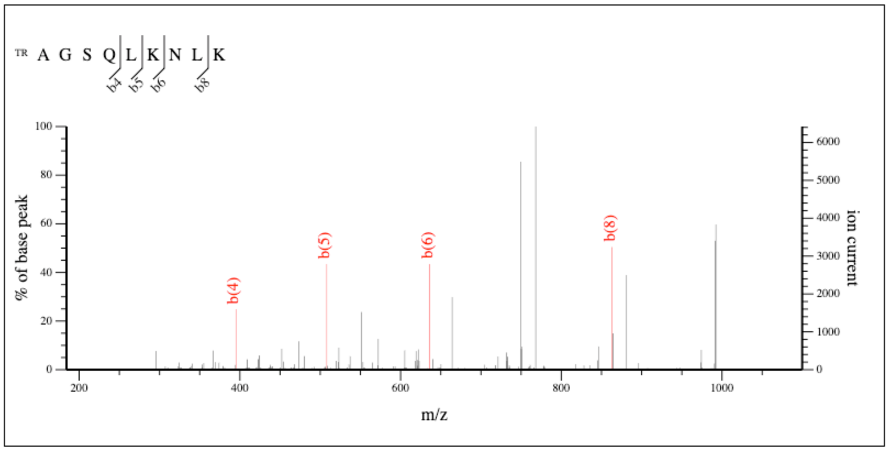


**Monoisotopic mass of neutral peptide Mr(calc):** 1008.6641
**Fixed modifications:** Carbamidomethyl (C) (apply to specified residues or termini only)
**Variable modifications:**
**N-term :** Trimethyl:2H(9) (Protein N-term)
**Ions Score:** 27 **Expect:** 0.0039
**Matches :** 4/37 fragment ions using 8 most intense peaks (help)

| # | b | b* | b$^0$ | Seq. | y | y* | y$^0$ | # |
|---|---|---|---|---|---|---|---|---|
| 1 | 123.1478 | | | A | | | | 9 |
| 2 | 180.1693 | | | G | 887.5309 | 870.5043 | 869.5203 | 8 |
| 3 | 267.2013 | | 249.1908 | S | 830.5094 | 813.4829 | 812.4989 | 7 |
| 4 | ***395.2599*** | 378.2334 | 377.2493 | Q | 743.4774 | 726.4509 | | 6 |
| 5 | ***508.3440*** | 491.3174 | 490.3334 | L | 615.4188 | 598.3923 | | 5 |
| 6 | ***636.4389*** | 619.4124 | 618.4284 | K | 502.3348 | 485.3082 | | 4 |
| 7 | 750.4819 | 733.4553 | 732.4713 | N | 374.2398 | 357.2132 | | 3 |
| 8 | ***863.5659*** | 846.5394 | 845.5554 | L | 260.1969 | 243.1703 | | 2 |
| 9 | | | | K | 147.1128 | 130.0863 | | 1 |

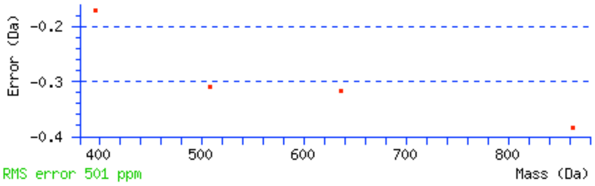


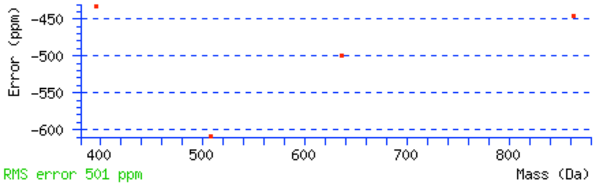


42. Eft1 (4 peptides in 300mM fraction2)

MS/MS Fragmentation of **VAFTVDQMRSLMDK**
Found in **P32324** in **Uniprot_yeastS288C**, Elongation factor 2 OS=Saccharomyces cerevisiae (strain ATCC 204508 / S288c) OX=559292 GN=EFT1 PE=1 SV=1

Match to Query 9596: 1706.868992 from(854.441772,2+) intensity(497797.5625) rtinseconds(2662.8691) index(6988)
Title: 20100530_methyl300mM_2.7775.7775.2
Data file D:\Panyue\Proteomic dataset\20100530_methyl300mM_2.RAW

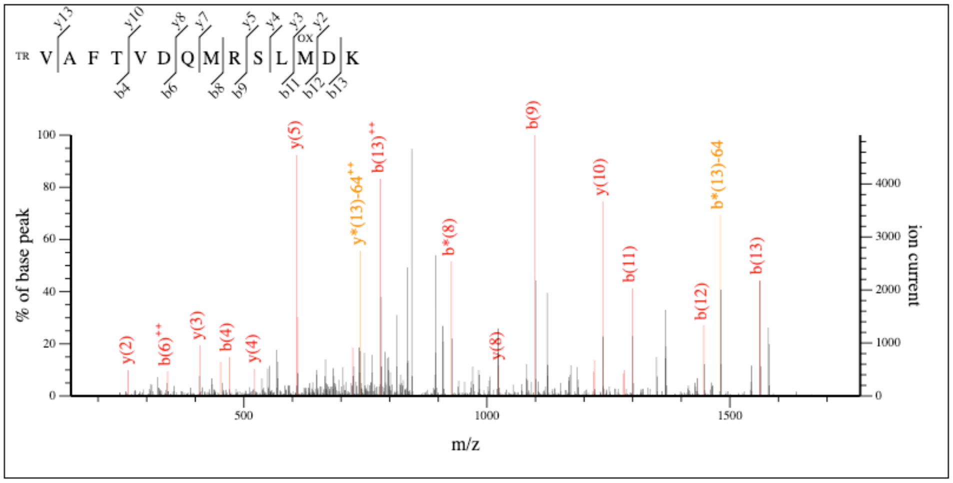


**Monoisotopic mass of neutral peptide Mr(calc):** 1706.9046
**Fixed modifications:** Carbamidomethyl (C) (apply to specified residues or termini only)
**Variable modifications:**
**N-term** : Trimethyl:2H(9) (Protein N-term)
**M12** : Oxidation (M), with neutral losses 0.0000(shown in table), 63.9983
**Ions Score:** 28 **Expect:** 0.024
**Matches** : 28/214 fragment ions using 43 most intense peaks (help)

| # | b | b++ | b* | b*++ | b0 | b0++ | Seq. | y | y++ | y* | y*++ | y0 | y0++ | # |
|---|---|---|---|---|---|---|---|---|---|---|---|---|---|---|
| 1 | 151.1791 | 76.0932 | | | | | V | | | | | | | 14 |
| 2 | 222.2162 | 111.6118 | | | | | A | 1557.7400 | 779.3736 | 1540.7134 | 770.8604 | 1539.7294 | 770.3683 | 13 |
| 3 | 369.2847 | 185.1460 | | | | | F | 1486.7029 | 743.8551 | 1469.6763 | 735.3418 | 1468.6923 | 734.8498 | 12 |
| 4 | ***470.3323*** | 235.6698 | | | 452.3218 | 226.6645 | T | 1339.6344 | 670.3209 | 1322.6079 | 661.8076 | 1321.6239 | 661.3156 | 11 |
| 5 | 569.4008 | 285.2040 | | | 551.3902 | 276.1987 | V | ***1238.5868*** | 619.7970 | 1221.5602 | 611.2837 | 1220.5762 | 610.7917 | 10 |
| 6 | 684.4277 | **342.7175** | | | 666.4171 | 333.7122 | D | 1139.5184 | 570.2628 | 1122.4918 | 561.7495 | 1121.5078 | 561.2575 | 9 |
| 7 | 812.4863 | 406.7468 | 795.4597 | 398.2335 | 794.4757 | 397.7415 | Q | ***1024.4914*** | 512.7493 | 1007.4649 | 504.2361 | 1006.4808 | 503.7441 | 8 |
| 8 | 943.5268 | 472.2670 | 926.5002 | 463.7537 | 925.5162 | 463.2617 | M | 896.4328 | 448.7201 | 879.4063 | 440.2068 | 878.4223 | 439.7148 | 7 |
| 9 | ***1099.6279*** | 550.3176 | 1082.6013 | 541.8043 | 1081.6173 | 541.3123 | R | 765.3924 | 383.1998 | 748.3658 | 374.6865 | 747.3818 | 374.1945 | 6 |
| 10 | 1186.6599 | 593.8336 | 1169.6333 | 585.3203 | 1168.6493 | 584.8283 | S | ***609.2912*** | 305.1493 | 592.2647 | 296.6360 | 591.2807 | 296.1440 | 5 |
| 11 | ***1299.7440*** | 650.3756 | 1282.7174 | 641.8623 | 1281.7334 | 641.3703 | L | ***522.2592*** | 261.6332 | 505.2327 | 253.1200 | 504.2486 | 252.6280 | 4 |
| 12 | ***1446.7794*** | **723.8933** | 1429.7528 | 715.3800 | 1428.7688 | 714.8880 | M | ***409.1751*** | 205.0912 | 392.1486 | 196.5779 | 391.1646 | 196.0859 | 3 |
| 13 | ***1561.8063*** | **781.4068** | 1544.7798 | 772.8935 | 1543.7957 | 772.4015 | D | ***262.1397*** | 131.5735 | 245.1132 | 123.0602 | 244.1292 | 122.5682 | 2 |
| 14 | | | | | | | K | 147.1128 | 74.0600 | 130.0863 | 65.5468 | | | 1 |

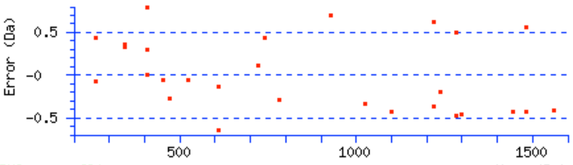


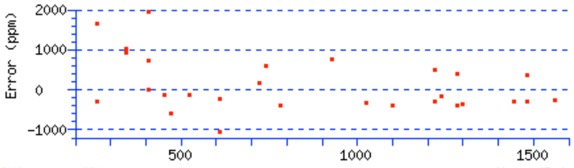

MS/MS Fragmentation of **VAFTVDQMRSLMDK**
Found in **P32324** in **Uniprot_yeastS288C**, Elongation factor 2 OS=Saccharomyces cerevisiae (strain ATCC 204508 / S288c) OX=559292 GN=EFT1 PE=1 SV=1

Match to Query 9597: 1706.871312 from(854.442932,2+) intensity(2050476.7500) rtinseconds(2664.2804) index(6991)
Title: 20100530_methyl300mM_2.7779.7779.2
Data file D:\Panyue\Proteomic dataset\20100530_methyl300mM_2.RAW

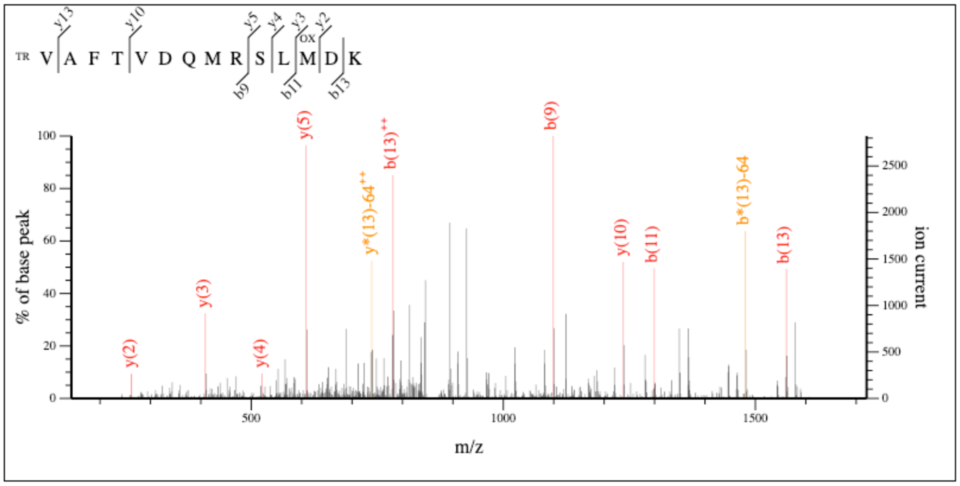


**Monoisotopic mass of neutral peptide Mr(calc):** 1706.9046
**Fixed modifications:** Carbamidomethyl (C) (apply to specified residues or termini only)
**Variable modifications:**
**N-term :** Trimethyl:2H(9) (Protein N-term)
**M12** : Oxidation (M), with neutral losses 0.0000(shown in table), 63.9983
**Ions Score:** 23 **Expect:** 0.075
**Matches :** 14/214 fragment ions using 14 most intense peaks (help)

| # | b | b$^{++}$ | b* | b*$^{++}$ | b$^{0}$ | b$^{0++}$ | Seq. | y | y$^{++}$ | y* | y*$^{++}$ | y$^{0}$ | y$^{0++}$ | # |
|---|---|---|---|---|---|---|---|---|---|---|---|---|---|---|
| 1 | 151.1791 | 76.0932 | | | | | V | | | | | | | 14 |
| 2 | 222.2162 | 111.6118 | | | | | A | 1557.7400 | 779.3736 | 1540.7134 | 770.8604 | 1539.7294 | 770.3683 | 13 |
| 3 | 369.2847 | 185.1460 | | | | | F | 1486.7029 | 743.8551 | 1469.6763 | 735.3418 | 1468.6923 | 734.8498 | 12 |
| 4 | 470.3323 | 235.6698 | | | 452.3218 | 226.6645 | T | 1339.6344 | 670.3209 | 1322.6079 | 661.8076 | 1321.6239 | 661.3156 | 11 |
| 5 | 569.4008 | 285.2040 | | | 551.3902 | 276.1987 | V | ***1238.5868*** | 619.7970 | 1221.5602 | 611.2837 | 1220.5762 | 610.7917 | 10 |
| 6 | 684.4277 | 342.7175 | | | 666.4171 | 333.7122 | D | 1139.5184 | 570.2628 | 1122.4918 | 561.7495 | 1121.5078 | 561.2575 | 9 |
| 7 | 812.4863 | 406.7468 | 795.4597 | 398.2335 | 794.4757 | 397.7415 | Q | 1024.4914 | 512.7493 | 1007.4649 | 504.2361 | 1006.4808 | 503.7441 | 8 |
| 8 | 943.5268 | 472.2670 | 926.5002 | 463.7537 | 925.5162 | 463.2617 | M | 896.4328 | 448.7201 | 879.4063 | 440.2068 | 878.4223 | 439.7148 | 7 |
| 9 | **1099.6279** | 550.3176 | 1082.6013 | 541.8043 | 1081.6173 | 541.3123 | R | 765.3924 | 383.1998 | 748.3658 | 374.6865 | 747.3818 | 374.1945 | 6 |
| 10 | 1186.6599 | 593.8336 | 1169.6333 | 585.3203 | 1168.6493 | 584.8283 | S | ***609.2912*** | 305.1493 | 592.2647 | 296.6360 | 591.2807 | 296.1440 | 5 |
| 11 | **1299.7440** | 650.3756 | 1282.7174 | 641.8623 | 1281.7334 | 641.3703 | L | ***522.2592*** | 261.6332 | 505.2327 | 253.1200 | 504.2486 | 252.6280 | 4 |
| 12 | 1446.7794 | 723.8933 | 1429.7528 | 715.3800 | 1428.7688 | 714.8880 | M | ***409.1751*** | 205.0912 | 392.1486 | 196.5779 | 391.1646 | 196.0859 | 3 |
| 13 | **1561.8063** | 781.4068 | 1544.7798 | 772.8935 | 1543.7957 | 772.4015 | D | ***262.1397*** | 131.5735 | 245.1132 | 123.0602 | 244.1292 | 122.5682 | 2 |
| 14 | | | | | | | K | 147.1128 | 74.0600 | 130.0863 | 65.5468 | | | 1 |

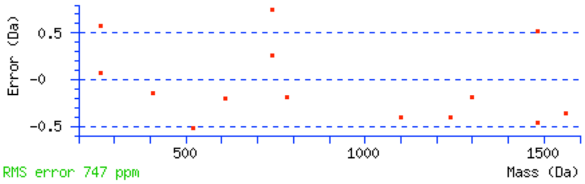


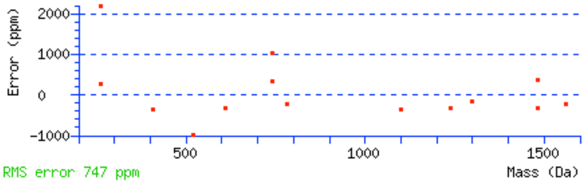

MS/MS Fragmentation of **VAFTVDQMRSLMDK**
Found in **P32324** in **Uniprot_yeastS288C**, Elongation factor 2 OS=Saccharomyces cerevisiae (strain ATCC 204508 / S288c) OX=559292 GN=EFT1 PE=1 SV=1

Match to Query 9598: 1706.872532 from(854.443542,2+) intensity(497797.5625) rtinseconds(2663.396) index(6990)
Title: 20100530_methyl300mM_2.7777.7777.2
Data file D:\Panyue\Proteomic dataset\20100530_methyl300mM_2.RAW

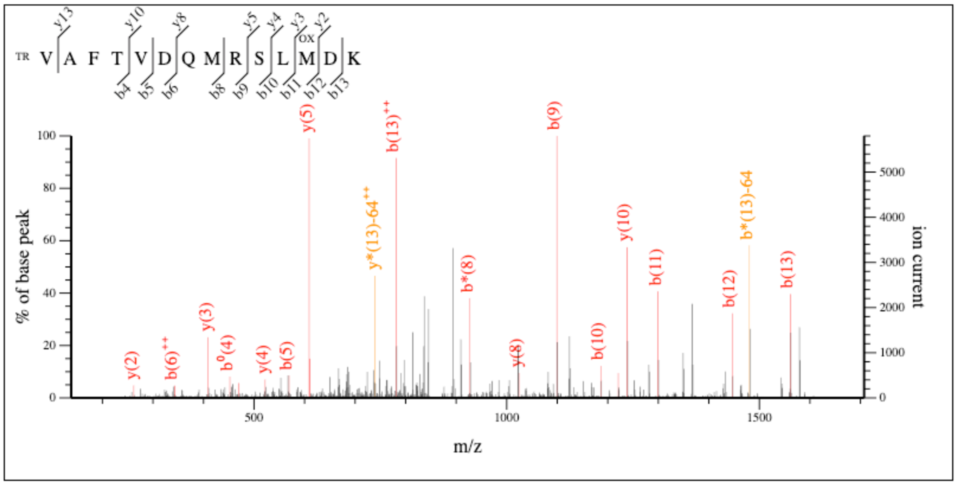


**Monoisotopic mass of neutral peptide Mr(calc):** 1706.9046
**Fixed modifications:** Carbamidomethyl (C) (apply to specified residues or termini only)
**Variable modifications:**
**N-term :** Trimethyl:2H(9) (Protein N-term)
**M12** : Oxidation (M), with neutral losses 0.0000(shown in table), 63.9983
**Ions Score:** 32 **Expect:** 0.011
**Matches :** 25/214 fragment ions using 42 most intense peaks (help)

| # | b | $b^{++}$ | b* | $b^{*++}$ | $b^{0}$ | $b^{0++}$ | Seq. | y | $y^{++}$ | y* | $y^{*++}$ | $y^{0}$ | $y^{0++}$ | # |
|---|---|---|---|---|---|---|---|---|---|---|---|---|---|---|
| 1 | 151.1791 | 76.0932 | | | | | V | | | | | | | 14 |
| 2 | 222.2162 | 111.6118 | | | | | A | 1557.7400 | 779.3736 | 1540.7134 | 770.8604 | 1539.7294 | 770.3683 | 13 |
| 3 | 369.2847 | 185.1460 | | | | | F | 1486.7029 | 743.8551 | 1469.6763 | 735.3418 | 1468.6923 | 734.8498 | 12 |
| 4 | ***470.3323*** | 235.6698 | | | 452.3218 | 226.6645 | T | 1339.6344 | 670.3209 | 1322.6079 | 661.8076 | 1321.6239 | 661.3156 | 11 |
| 5 | ***569.4008*** | 285.2040 | | | 551.3902 | 276.1987 | V | ***1238.5868*** | 619.7970 | 1221.5602 | 611.2837 | 1220.5762 | 610.7917 | 10 |
| 6 | 684.4277 | 342.7175 | | | 666.4171 | 333.7122 | D | 1139.5184 | 570.2628 | 1122.4918 | 561.7495 | 1121.5078 | 561.2575 | 9 |
| 7 | 812.4863 | 406.7468 | 795.4597 | 398.2335 | 794.4757 | 397.7415 | Q | ***1024.4914*** | 512.7493 | 1007.4649 | 504.2361 | 1006.4808 | 503.7441 | 8 |
| 8 | 943.5268 | 472.2670 | 926.5002 | 463.7537 | 925.5162 | 463.2617 | M | 896.4328 | 448.7201 | 879.4063 | 440.2068 | 878.4223 | 439.7148 | 7 |
| 9 | ***1099.6279*** | 550.3176 | 1082.6013 | 541.8043 | 1081.6173 | 541.3123 | R | 765.3924 | 383.1998 | 748.3658 | 374.6865 | 747.3818 | 374.1945 | 6 |
| 10 | ***1186.6599*** | 593.8336 | 1169.6333 | 585.3203 | 1168.6493 | 584.8283 | S | ***609.2912*** | 305.1493 | 592.2647 | 296.6360 | 591.2807 | 296.1440 | 5 |
| 11 | ***1299.7440*** | 650.3756 | 1282.7174 | 641.8623 | 1281.7334 | 641.3703 | L | ***522.2592*** | 261.6332 | 505.2327 | 253.1200 | 504.2486 | 252.6280 | 4 |
| 12 | ***1446.7794*** | 723.8933 | 1429.7528 | 715.3800 | 1428.7688 | 714.8880 | M | ***409.1751*** | 205.0912 | 392.1486 | 196.5779 | 391.1646 | 196.0859 | 3 |
| 13 | ***1561.8063*** | 781.4068 | 1544.7798 | 772.8935 | 1543.7957 | 772.4015 | D | ***262.1397*** | 131.5735 | 245.1132 | 123.0602 | 244.1292 | 122.5682 | 2 |
| 14 | | | | | | | K | 147.1128 | 74.0600 | 130.0863 | 65.5468 | | | 1 |

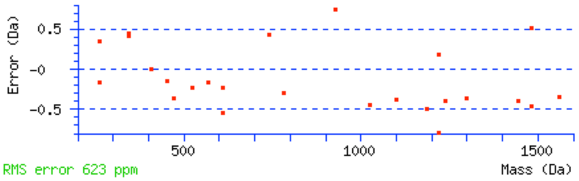


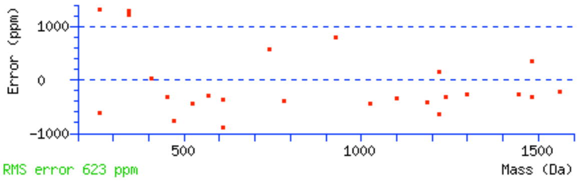

MS/MS Fragmentation of **VAFTVDQMRSLMDK**
Found in **P32324** in **Uniprot_yeastS288C**, Elongation factor 2 OS=Saccharomyces cerevisiae (strain ATCC 204508 / S288c) OX=559292 GN=EFT1 PE=1 SV=1

Match to Query 9599: 1706.872778 from(854.443665,2+) intensity(2025475.2500) rtinseconds(2664.5713) index(6992)
Title: 20100530_methyl300mM_2.7780.7780.2
Data file D:\Panyue\Proteomic dataset\20100530_methyl300mM_2.RAW

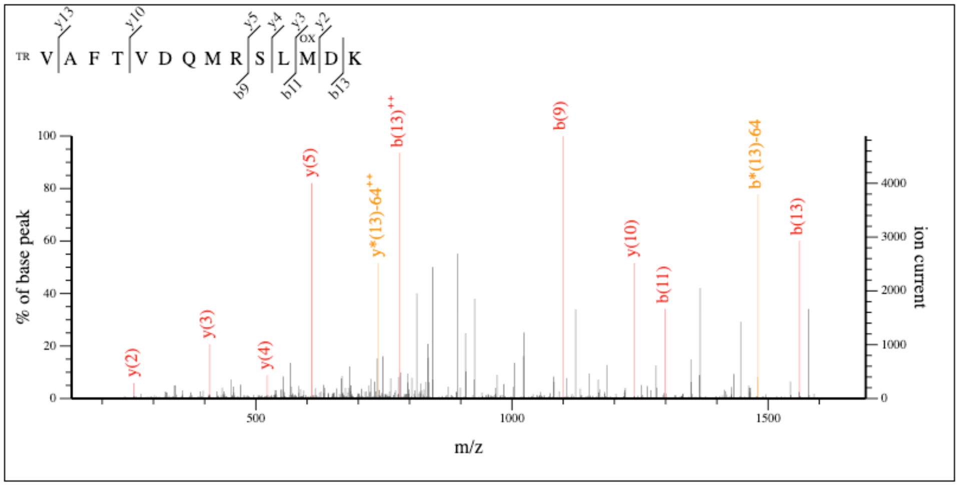


**Monoisotopic mass of neutral peptide Mr(calc):** 1706.9046
**Fixed modifications:** Carbamidomethyl (C) (apply to specified residues or termini only)
**Variable modifications:**
**N-term :** Trimethyl:2H(9) (Protein N-term)
**M12 :** Oxidation (M), with neutral losses 0.0000(shown in table), 63.9983
**Ions Score:** 24 **Expect:** 0.061
**Matches :** 13/214 fragment ions using 14 most intense peaks (help)

| # | b | b++ | b* | b*++ | b0 | b0++ | Seq. | y | y++ | y* | y*++ | y0 | y0++ | # |
|---|---|---|---|---|---|---|---|---|---|---|---|---|---|---|
| 1 | 151.1791 | 76.0932 | | | | | V | | | | | | | 14 |
| 2 | 222.2162 | 111.6118 | | | | | A | 1557.7400 | 779.3736 | 1540.7134 | 770.8604 | 1539.7294 | 770.3683 | 13 |
| 3 | 369.2847 | 185.1460 | | | | | F | 1486.7029 | 743.8551 | 1469.6763 | 735.3418 | 1468.6923 | 734.8498 | 12 |
| 4 | 470.3323 | 235.6698 | | | 452.3218 | 226.6645 | T | 1339.6344 | 670.3209 | 1322.6079 | 661.8076 | 1321.6239 | 661.3156 | 11 |
| 5 | 569.4008 | 285.2040 | | | 551.3902 | 276.1987 | V | ***1238.5868*** | 619.7970 | 1221.5602 | 611.2837 | 1220.5762 | 610.7917 | 10 |
| 6 | 684.4277 | 342.7175 | | | 666.4171 | 333.7122 | D | 1139.5184 | 570.2628 | 1122.4918 | 561.7495 | 1121.5078 | 561.2575 | 9 |
| 7 | 812.4863 | 406.7468 | 795.4597 | 398.2335 | 794.4757 | 397.7415 | Q | 1024.4914 | 512.7493 | 1007.4649 | 504.2361 | 1006.4808 | 503.7441 | 8 |
| 8 | 943.5268 | 472.2670 | 926.5002 | 463.7537 | 925.5162 | 463.2617 | M | 896.4328 | 448.7201 | 879.4063 | 440.2068 | 878.4223 | 439.7148 | 7 |
| 9 | **1099.6279** | 550.3176 | 1082.6013 | 541.8043 | 1081.6173 | 541.3123 | R | 765.3924 | 383.1998 | 748.3658 | 374.6865 | 747.3818 | 374.1945 | 6 |
| 10 | 1186.6599 | 593.8336 | 1169.6333 | 585.3203 | 1168.6493 | 584.8283 | S | ***609.2912*** | 305.1493 | 592.2647 | 296.6360 | 591.2807 | 296.1440 | 5 |
| 11 | **1299.7440** | 650.3756 | 1282.7174 | 641.8623 | 1281.7334 | 641.3703 | L | ***522.2592*** | 261.6332 | 505.2327 | 253.1200 | 504.2486 | 252.6280 | 4 |
| 12 | 1446.7794 | 723.8933 | 1429.7528 | 715.3800 | 1428.7688 | 714.8880 | M | ***409.1751*** | 205.0912 | 392.1486 | 196.5779 | 391.1646 | 196.0859 | 3 |
| 13 | **1561.8063** | 781.4068 | 1544.7798 | 772.8935 | 1543.7957 | 772.4015 | D | ***262.1397*** | 131.5735 | 245.1132 | 123.0602 | 244.1292 | 122.5682 | 2 |
| 14 | | | | | | | K | 147.1128 | 74.0600 | 130.0863 | 65.5468 | | | 1 |

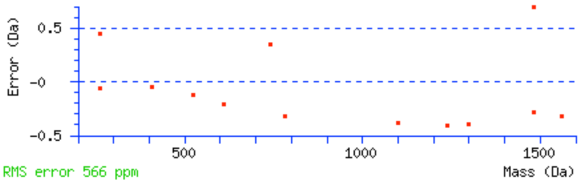


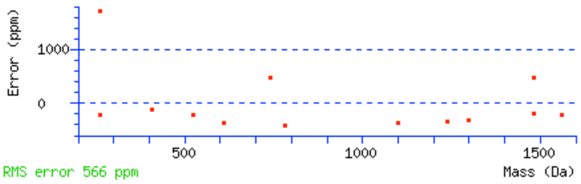


43. Asc1 (1 peptide in 300mM fraction 2)

MS/MS Fragmentation of **ASNEVLVLR**
Found in **P38011** in **Uniprot_yeastS288C**, Guanine nucleotide-binding protein subunit beta-like protein OS=Saccharomyces cerevisiae (strain ATCC 204508 / S288c) OX=559292 GN=ASC1 PE=1 SV=4

Match to Query 5002: 1050.681689 from(1051.688965,1+) intensity(761395.4375) rtinseconds(3370.3069) index(8883)
Title: 20100530_methyl300mM_2.9883.9883.1
Data file D:\Panyue\Proteomic dataset\20100530_methyl300mM_2.RAW

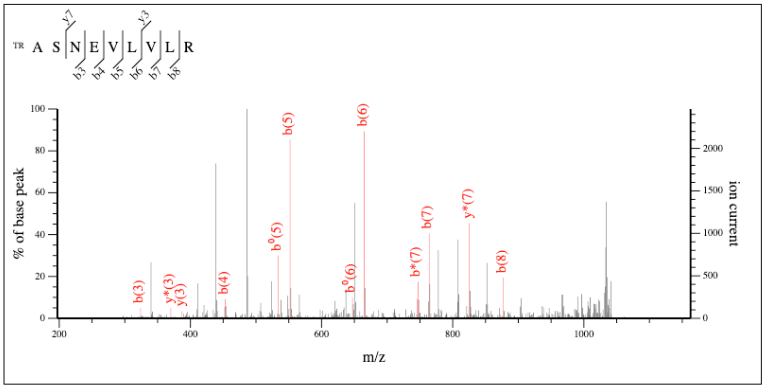

**Monoisotopic mass of neutral peptide Mr(calc):** 1050.6747
**Fixed modifications:** Carbamidomethyl (C) (apply to specified residues or termini only)
**Variable modifications:**
**N-term :** Trimethyl:2H(9) (Protein N-term)
**Ions Score:** 20 **Expect:** 0.022
**Matches :** 13/40 fragment ions using 40 most intense peaks (help)

| # | b | b* | $b^0$ | Seq. | y | y* | $y^0$ | # |
|---|---|---|---|---|---|---|---|---|
| 1 | 123.1478 | | | A | | | | 9 |
| 2 | 210.1799 | | 192.1693 | S | 929.5415 | 912.5149 | 911.5309 | 8 |
| 3 | ***324.2228*** | 307.1962 | 306.2122 | N | 842.5094 | 825.4829 | 824.4989 | 7 |
| 4 | ***453.2654*** | 436.2388 | 435.2548 | E | 728.4665 | 711.4400 | 710.4559 | 6 |
| 5 | ***552.3338*** | 535.3072 | 534.3232 | V | 599.4239 | 582.3974 | | 5 |
| 6 | ***665.4179*** | 648.3913 | 647.4073 | L | 500.3555 | 483.3289 | | 4 |
| 7 | ***764.4863*** | 747.4597 | 746.4757 | V | 387.2714 | 370.2449 | | 3 |
| 8 | ***877.5703*** | 860.5438 | 859.5598 | L | 288.2030 | 271.1765 | | 2 |
| 9 | | | | R | 175.1190 | 158.0924 | | 1 |

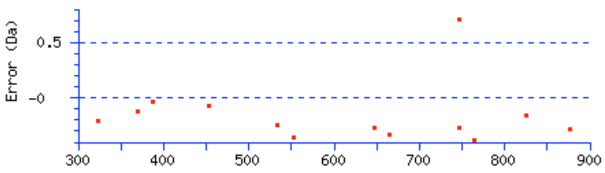


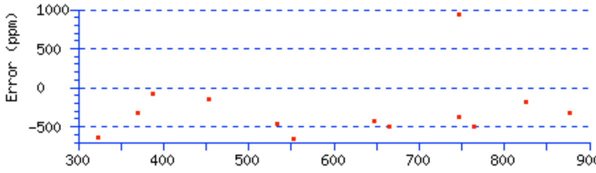


44. Mon2 (6 peptides in 300mM fraction 3)

MS/MS Fragmentation of **AMNTGGFDSMQRQLEAELRSLSSESK**
Found in **P48563** in **Uniprot_yeastS288C**, Protein MON2 OS=Saccharomyces cerevisiae (strain ATCC 204508 / S288c) OX=559292 GN=MON2 PE=1 SV=1

Match to Query 16224: 2938.437339 from(980.486389,3+) intensity(359560.2500) rtinseconds(5899.3213) index(15021)
Title: 20100530_methyl300mM_3.16710.16710.3
Data file D:\Panyue\Proteomic dataset\20100530_methyl300mM_3.RAW

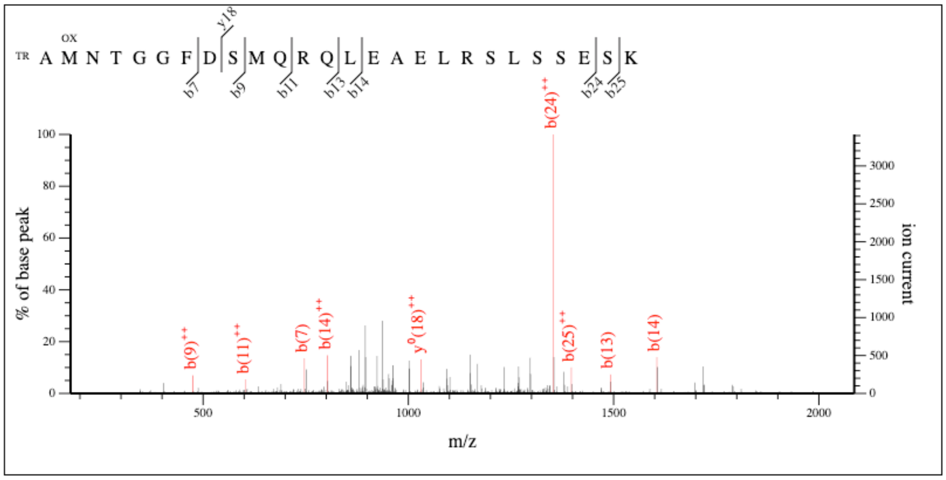


**Monoisotopic mass of neutral peptide Mr(calc):** 2938.4474
**Fixed modifications:** Carbamidomethyl (C) (apply to specified residues or termini only)
**Variable modifications:**
**N-term :** Trimethyl:2H(9) (Protein N-term)
**M2** : Oxidation (M), with neutral losses 0.0000(shown in table), 63.9983
**Ions Score:** 20 **Expect:** 0.1
**Matches :** 11/432 fragment ions using 18 most intense peaks (help)

| # | b | $b^{++}$ | b* | $b^{*++}$ | $b^0$ | $b^{0++}$ | Seq. | y | $y^{++}$ | y* | $y^{*++}$ | $y^0$ | $y^{0++}$ | # |
|---|---|---|---|---|---|---|---|---|---|---|---|---|---|---|
| 1 | 123.1478 | 62.0776 | | | | | A | | | | | | | 26 |
| 2 | 270.1832 | 135.5953 | | | | | M | 2817.3142 | 1409.1607 | 2800.2876 | 1400.6475 | 2799.3036 | 1400.1554 | 25 |
| 3 | 384.2262 | 192.6167 | 367.1996 | 184.1034 | | | N | 2670.2788 | 1335.6430 | 2653.2522 | 1327.1298 | 2652.2682 | 1326.6377 | 24 |
| 4 | 485.2738 | 243.1406 | 468.2473 | 234.6273 | 467.2633 | 234.1353 | T | 2556.2359 | 1278.6216 | 2539.2093 | 1270.1083 | 2538.2253 | 1269.6163 | 23 |
| 5 | 542.2953 | 271.6513 | 525.2688 | 263.1380 | 524.2847 | 262.6460 | G | 2455.1882 | 1228.0977 | 2438.1616 | 1219.5845 | 2437.1776 | 1219.0924 | 22 |
| 6 | 599.3168 | 300.1620 | 582.2902 | 291.6487 | 581.3062 | 291.1567 | G | 2398.1667 | 1199.5870 | 2381.1402 | 1191.0737 | 2380.1561 | 1190.5817 | 21 |
| 7 | **746.3852** | 373.6962 | 729.3586 | 365.1830 | 728.3746 | 364.6909 | F | 2341.1452 | 1171.0763 | 2324.1187 | 1162.5630 | 2323.1347 | 1162.0710 | 20 |
| 8 | 861.4121 | 431.2097 | 844.3856 | 422.6964 | 843.4016 | 422.2044 | D | 2194.0768 | 1097.5421 | 2177.0503 | 1089.0288 | 2176.0663 | 1088.5368 | 19 |
| 9 | 948.4441 | ***474.7257*** | 931.4176 | 466.2124 | 930.4336 | 465.7204 | S | 2079.0499 | 1040.0286 | 2062.0233 | 1031.5153 | 2061.0393 | 1031.0233 | 18 |
| 10 | 1079.4846 | 540.2460 | 1062.4581 | 531.7327 | 1061.4741 | 531.2407 | M | 1992.0179 | 996.5126 | 1974.9913 | 987.9993 | 1974.0073 | 987.5073 | 17 |
| 11 | 1207.5432 | ***604.2752*** | 1190.5167 | 595.7620 | 1189.5326 | 595.2700 | Q | 1860.9774 | 930.9923 | 1843.9508 | 922.4791 | 1842.9668 | 921.9870 | 16 |
| 12 | 1363.6443 | 682.3258 | 1346.6178 | 673.8125 | 1345.6338 | 673.3205 | R | 1732.9188 | 866.9630 | 1715.8923 | 858.4498 | 1714.9082 | 857.9578 | 15 |
| 13 | **1491.7029** | ***746.3551*** | 1474.6764 | 737.8418 | 1473.6923 | 737.3498 | Q | 1576.8177 | 788.9125 | 1559.7911 | 780.3992 | 1558.8071 | 779.9072 | 14 |
| 14 | **1604.7870** | ***802.8971*** | 1587.7604 | 794.3838 | 1586.7764 | 793.8918 | L | 1448.7591 | 724.8832 | 1431.7326 | 716.3699 | 1430.7485 | 715.8779 | 13 |
| 15 | 1733.8296 | 867.4184 | 1716.8030 | 858.9051 | 1715.8190 | 858.4131 | E | 1335.6750 | 668.3412 | 1318.6485 | 659.8279 | 1317.6645 | 659.3359 | 12 |
| 16 | 1804.8667 | 902.9370 | 1787.8401 | 894.4237 | 1786.8561 | 893.9317 | A | 1206.6325 | 603.8199 | 1189.6059 | 595.3066 | 1188.6219 | 594.8146 | 11 |
| 17 | 1933.9093 | 967.4583 | 1916.8827 | 958.9450 | 1915.8987 | 958.4530 | E | 1135.5953 | 568.3013 | 1118.5688 | 559.7880 | 1117.5848 | 559.2960 | 10 |
| 18 | 2046.9933 | 1024.0003 | 2029.9668 | 1015.4870 | 2028.9828 | 1014.9950 | L | 1006.5527 | 503.7800 | 989.5262 | 495.2667 | 988.5422 | 494.7747 | 9 |
| 19 | 2203.0944 | 1102.0509 | 2186.0679 | 1093.5376 | 2185.0839 | 1093.0456 | R | 893.4687 | 447.2380 | 876.4421 | 438.7247 | 875.4581 | 438.2327 | 8 |
| 20 | 2290.1265 | 1145.5669 | 2273.0999 | 1137.0536 | 2272.1159 | 1136.5616 | S | 737.3676 | 369.1874 | 720.3410 | 360.6742 | 719.3570 | 360.1821 | 7 |
| 21 | 2403.2105 | 1202.1089 | 2386.1840 | 1193.5956 | 2385.2000 | 1193.1036 | L | 650.3355 | 325.6714 | 633.3090 | 317.1581 | 632.3250 | 316.6661 | 6 |
| 22 | 2490.2426 | 1245.6249 | 2473.2160 | 1237.1116 | 2472.2320 | 1236.6196 | S | 537.2515 | 269.1294 | 520.2249 | 260.6161 | 519.2409 | 260.1241 | 5 |
| 23 | 2577.2746 | 1289.1409 | 2560.2480 | 1280.6277 | 2559.2640 | 1280.1356 | S | 450.2195 | 225.6134 | 433.1929 | 217.1001 | 432.2089 | 216.6081 | 4 |
| 24 | 2706.3172 | ***1353.6622*** | 2689.2906 | 1345.1490 | 2688.3066 | 1344.6569 | E | 363.1874 | 182.0974 | 346.1609 | 173.5841 | 345.1769 | 173.0921 | 3 |
| 25 | 2793.3492 | ***1397.1782*** | 2776.3227 | 1388.6650 | 2775.3386 | 1388.1730 | S | 234.1448 | 117.5761 | 217.1183 | 109.0628 | 216.1343 | 108.5708 | 2 |
| 26 | | | | | | | K | 147.1128 | 74.0600 | 130.0863 | 65.5468 | | | 1 |

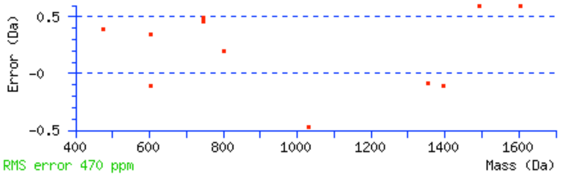

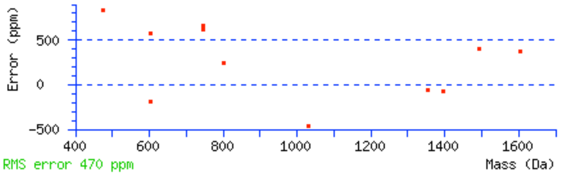

MS/MS Fragmentation of **AMNTGGFDSMQRQLEAELRSLSSESK**
Found in **P48563** in **Uniprot_yeastS288C**, Protein MON2 OS=Saccharomyces cerevisiae (strain ATCC 204508 / S288c) OX=559292 GN=MON2 PE=1 SV=1

Match to Query 16228: 2938.437705 from(980.486511,3+) intensity(1156262.5000) rtinseconds(6342.7228) index(16077)
Title: 20100530_methyl300mM_3.17884.17884.3
Data file D:\Panyue\Proteomic dataset\20100530_methyl300mM_3.RAW

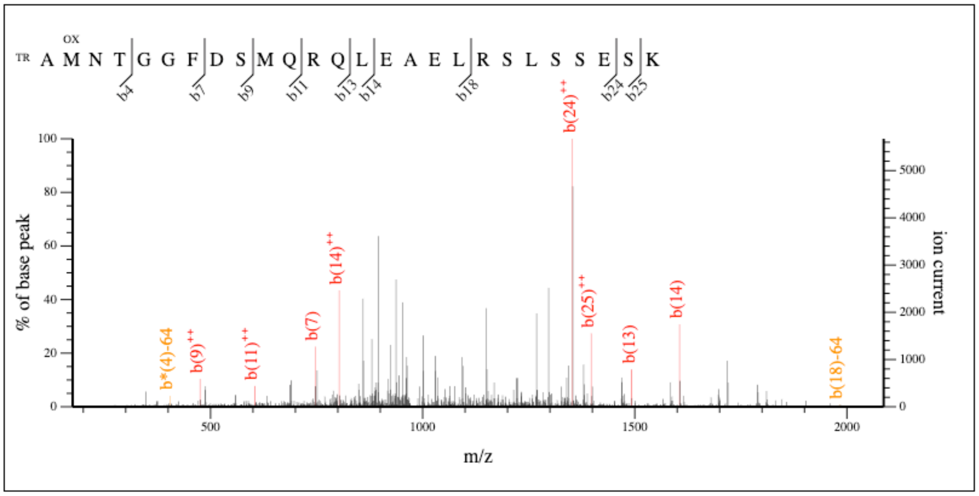

**Monoisotopic mass of neutral peptide Mr(calc):** 2938.4474
**Fixed modifications:** Carbamidomethyl (C) (apply to specified residues or termini only)
**Variable modifications:**
**N-term :** Trimethyl:2H(9) (Protein N-term)
**M2** : Oxidation (M), with neutral losses 0.0000(shown in table), 63.9983
**Ions Score:** 21 **Expect:** 0.093
**Matches :** 12/432 fragment ions using 18 most intense peaks (help)

| # | b | $b^{++}$ | $b^*$ | $b^{*++}$ | $b^0$ | $b^{0++}$ | Seq. | y | $y^{++}$ | $y^*$ | $y^{*++}$ | $y^0$ | $y^{0++}$ | # |
|---|---|---|---|---|---|---|---|---|---|---|---|---|---|---|
| 1 | 123.1478 | 62.0776 | | | | | A | | | | | | | 26 |
| 2 | 270.1832 | 135.5953 | | | | | M | 2817.3142 | 1409.1607 | 2800.2876 | 1400.6475 | 2799.3036 | 1400.1554 | 25 |
| 3 | 384.2262 | 192.6167 | 367.1996 | 184.1034 | | | N | 2670.2788 | 1335.6430 | 2653.2522 | 1327.1298 | 2652.2682 | 1326.6377 | 24 |
| 4 | 485.2738 | 243.1406 | 468.2473 | 234.6273 | 467.2633 | 234.1353 | T | 2556.2359 | 1278.6216 | 2539.2093 | 1270.1083 | 2538.2253 | 1269.6163 | 23 |
| 5 | 542.2953 | 271.6513 | 525.2688 | 263.1380 | 524.2847 | 262.6460 | G | 2455.1882 | 1228.0977 | 2438.1616 | 1219.5845 | 2437.1776 | 1219.0924 | 22 |
| 6 | 599.3168 | 300.1620 | 582.2902 | 291.6487 | 581.3062 | 291.1567 | G | 2398.1667 | 1199.5870 | 2381.1402 | 1191.0737 | 2380.1561 | 1190.5817 | 21 |
| 7 | **746.3852** | 373.6962 | 729.3586 | 365.1830 | 728.3746 | 364.6909 | F | 2341.1452 | 1171.0763 | 2324.1187 | 1162.5630 | 2323.1347 | 1162.0710 | 20 |
| 8 | 861.4121 | 431.2097 | 844.3856 | 422.6964 | 843.4016 | 422.2044 | D | 2194.0768 | 1097.5421 | 2177.0503 | 1089.0288 | 2176.0663 | 1088.5368 | 19 |
| 9 | 948.4441 | ***474.7257*** | 931.4176 | 466.2124 | 930.4336 | 465.7204 | S | 2079.0499 | 1040.0286 | 2062.0233 | 1031.5153 | 2061.0393 | 1031.0233 | 18 |
| 10 | 1079.4846 | 540.2460 | 1062.4581 | 531.7327 | 1061.4741 | 531.2407 | M | 1992.0179 | 996.5126 | 1974.9913 | 987.9993 | 1974.0073 | 987.5073 | 17 |
| 11 | 1207.5432 | ***604.2752*** | 1190.5167 | 595.7620 | 1189.5326 | 595.2700 | Q | 1860.9774 | 930.9923 | 1843.9508 | 922.4791 | 1842.9668 | 921.9870 | 16 |
| 12 | 1363.6443 | 682.3258 | 1346.6178 | 673.8125 | 1345.6338 | 673.3205 | R | 1732.9188 | 866.9630 | 1715.8923 | 858.4498 | 1714.9082 | 857.9578 | 15 |
| 13 | **1491.7029** | ***746.3551*** | 1474.6764 | 737.8418 | 1473.6923 | 737.3498 | Q | 1576.8177 | 788.9125 | 1559.7911 | 780.3992 | 1558.8071 | 779.9072 | 14 |
| 14 | **1604.7870** | ***802.8971*** | 1587.7604 | 794.3838 | 1586.7764 | 793.8918 | L | 1448.7591 | 724.8832 | 1431.7326 | 716.3699 | 1430.7485 | 715.8779 | 13 |
| 15 | 1733.8296 | 867.4184 | 1716.8030 | 858.9051 | 1715.8190 | 858.4131 | E | 1335.6750 | 668.3412 | 1318.6485 | 659.8279 | 1317.6645 | 659.3359 | 12 |
| 16 | 1804.8667 | 902.9370 | 1787.8401 | 894.4237 | 1786.8561 | 893.9317 | A | 1206.6325 | 603.8199 | 1189.6059 | 595.3066 | 1188.6219 | 594.8146 | 11 |
| 17 | 1933.9093 | 967.4583 | 1916.8827 | 958.9450 | 1915.8987 | 958.4530 | E | 1135.5953 | 568.3013 | 1118.5688 | 559.7880 | 1117.5848 | 559.2960 | 10 |
| 18 | 2046.9933 | 1024.0003 | 2029.9668 | 1015.4870 | 2028.9828 | 1014.9950 | L | 1006.5527 | 503.7800 | 989.5262 | 495.2667 | 988.5422 | 494.7747 | 9 |
| 19 | 2203.0944 | 1102.0509 | 2186.0679 | 1093.5376 | 2185.0839 | 1093.0456 | R | 893.4687 | 447.2380 | 876.4421 | 438.7247 | 875.4581 | 438.2327 | 8 |
| 20 | 2290.1265 | 1145.5669 | 2273.0999 | 1137.0536 | 2272.1159 | 1136.5616 | S | 737.3676 | 369.1874 | 720.3410 | 360.6742 | 719.3570 | 360.1821 | 7 |
| 21 | 2403.2105 | 1202.1089 | 2386.1840 | 1193.5956 | 2385.2000 | 1193.1036 | L | 650.3355 | 325.6714 | 633.3090 | 317.1581 | 632.3250 | 316.6661 | 6 |
| 22 | 2490.2426 | 1245.6249 | 2473.2160 | 1237.1116 | 2472.2320 | 1236.6196 | S | 537.2515 | 269.1294 | 520.2249 | 260.6161 | 519.2409 | 260.1241 | 5 |
| 23 | 2577.2746 | 1289.1409 | 2560.2480 | 1280.6277 | 2559.2640 | 1280.1356 | S | 450.2195 | 225.6134 | 433.1929 | 217.1001 | 432.2089 | 216.6081 | 4 |
| 24 | 2706.3172 | ***1353.6622*** | 2689.2906 | 1345.1490 | 2688.3066 | 1344.6569 | E | 363.1874 | 182.0974 | 346.1609 | 173.5841 | 345.1769 | 173.0921 | 3 |
| 25 | 2793.3492 | ***1397.1782*** | 2776.3227 | 1388.6650 | 2775.3386 | 1388.1730 | S | 234.1448 | 117.5761 | 217.1183 | 109.0628 | 216.1343 | 108.5708 | 2 |
| 26 | | | | | | | K | 147.1128 | 74.0600 | 130.0863 | 65.5468 | | | 1 |

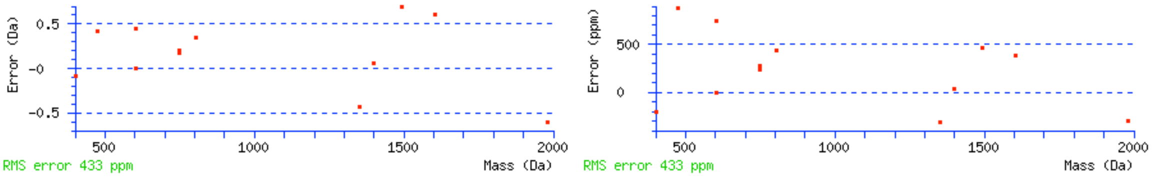


MS/MS Fragmentation of **AMNTGGFDSMQRQLEAELRSLSSESK**
Found in **P48563** in **Uniprot_yeastS288C**, Protein MON2 OS=Saccharomyces cerevisiae (strain ATCC 204508 / S288c) OX=559292 GN=MON2 PE=1 SV=1

Match to Query 16235: 2938.440453 from(980.487427,3+) intensity(1415059.2500) rtinseconds(6510.8986) index(16498)
Title: 20100530_methyl300mM_3.18352.18352.3
Data file D:\Panyue\Proteomic dataset\20100530_methyl300mM_3.RAW

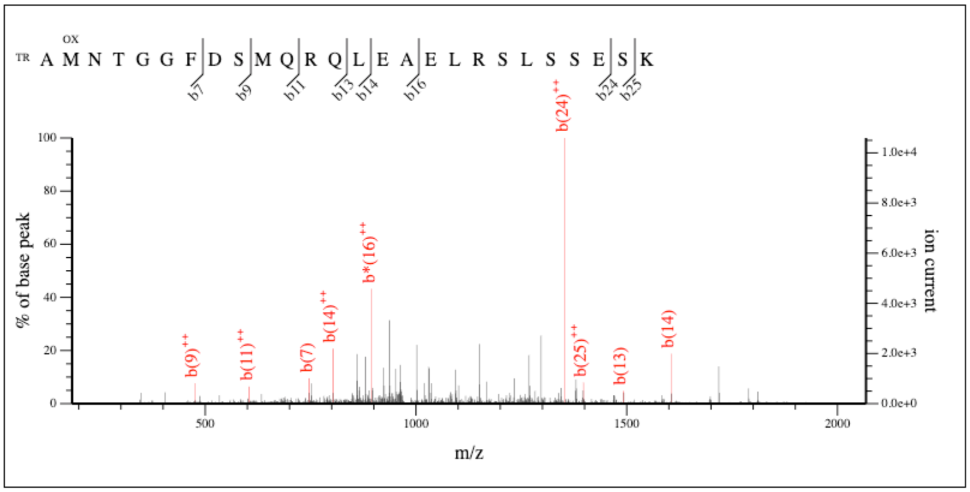

**Monoisotopic mass of neutral peptide Mr(calc):** 2938.4474
**Fixed modifications:** Carbamidomethyl (C) (apply to specified residues or termini only)
**Variable modifications:**
**N-term :** Trimethyl:2H(9) (Protein N-term)
**M2** : Oxidation (M), with neutral losses 0.0000(shown in table), 63.9983
**Ions Score:** 24 **Expect:** 0.045
**Matches :** 11/432 fragment ions using 17 most intense peaks (help)

| # | b | b++ | b* | b*++ | b0 | b0++ | Seq. | y | y++ | y* | y*++ | y0 | y0++ | # |
|---|---|---|---|---|---|---|---|---|---|---|---|---|---|---|
| 1 | 123.1478 | 62.0776 | | | | | A | | | | | | | 26 |
| 2 | 270.1832 | 135.5953 | | | | | M | 2817.3142 | 1409.1607 | 2800.2876 | 1400.6475 | 2799.3036 | 1400.1554 | 25 |
| 3 | 384.2262 | 192.6167 | 367.1996 | 184.1034 | | | N | 2670.2788 | 1335.6430 | 2653.2522 | 1327.1298 | 2652.2682 | 1326.6377 | 24 |
| 4 | 485.2738 | 243.1406 | 468.2473 | 234.6273 | 467.2633 | 234.1353 | T | 2556.2359 | 1278.6216 | 2539.2093 | 1270.1083 | 2538.2253 | 1269.6163 | 23 |
| 5 | 542.2953 | 271.6513 | 525.2688 | 263.1380 | 524.2847 | 262.6460 | G | 2455.1882 | 1228.0977 | 2438.1616 | 1219.5845 | 2437.1776 | 1219.0924 | 22 |
| 6 | 599.3168 | 300.1620 | 582.2902 | 291.6487 | 581.3062 | 291.1567 | G | 2398.1667 | 1199.5870 | 2381.1402 | 1191.0737 | 2380.1561 | 1190.5817 | 21 |
| 7 | **746.3852** | 373.6962 | 729.3586 | 365.1830 | 728.3746 | 364.6909 | F | 2341.1452 | 1171.0763 | 2324.1187 | 1162.5630 | 2323.1347 | 1162.0710 | 20 |
| 8 | 861.4121 | 431.2097 | 844.3856 | 422.6964 | 843.4016 | 422.2044 | D | 2194.0768 | 1097.5421 | 2177.0503 | 1089.0288 | 2176.0663 | 1088.5368 | 19 |
| 9 | 948.4441 | ***474.7257*** | 931.4176 | 466.2124 | 930.4336 | 465.7204 | S | 2079.0499 | 1040.0286 | 2062.0233 | 1031.5153 | 2061.0393 | 1031.0233 | 18 |
| 10 | 1079.4846 | 540.2460 | 1062.4581 | 531.7327 | 1061.4741 | 531.2407 | M | 1992.0179 | 996.5126 | 1974.9913 | 987.9993 | 1974.0073 | 987.5073 | 17 |
| 11 | 1207.5432 | ***604.2752*** | 1190.5167 | 595.7620 | 1189.5326 | 595.2700 | Q | 1860.9774 | 930.9923 | 1843.9508 | 922.4791 | 1842.9668 | 921.9870 | 16 |
| 12 | 1363.6443 | 682.3258 | 1346.6178 | 673.8125 | 1345.6338 | 673.3205 | R | 1732.9188 | 866.9630 | 1715.8923 | 858.4498 | 1714.9082 | 857.9578 | 15 |
| 13 | **1491.7029** | ***746.3551*** | 1474.6764 | 737.8418 | 1473.6923 | 737.3498 | Q | 1576.8177 | 788.9125 | 1559.7911 | 780.3992 | 1558.8071 | 779.9072 | 14 |
| 14 | **1604.7870** | ***802.8971*** | 1587.7604 | 794.3838 | 1586.7764 | 793.8918 | L | 1448.7591 | 724.8832 | 1431.7326 | 716.3699 | 1430.7485 | 715.8779 | 13 |
| 15 | 1733.8296 | 867.4184 | 1716.8030 | 858.9051 | 1715.8190 | 858.4131 | E | 1335.6750 | 668.3412 | 1318.6485 | 659.8279 | 1317.6645 | 659.3359 | 12 |
| 16 | 1804.8667 | 902.9370 | 1787.8401 | 894.4237 | 1786.8561 | 893.9317 | A | 1206.6325 | 603.8199 | 1189.6059 | 595.3066 | 1188.6219 | 594.8146 | 11 |
| 17 | 1933.9093 | 967.4583 | 1916.8827 | 958.9450 | 1915.8987 | 958.4530 | E | 1135.5953 | 568.3013 | 1118.5688 | 559.7880 | 1117.5848 | 559.2960 | 10 |
| 18 | 2046.9933 | 1024.0003 | 2029.9668 | 1015.4870 | 2028.9828 | 1014.9950 | L | 1006.5527 | 503.7800 | 989.5262 | 495.2667 | 988.5422 | 494.7747 | 9 |
| 19 | 2203.0944 | 1102.0509 | 2186.0679 | 1093.5376 | 2185.0839 | 1093.0456 | R | 893.4687 | 447.2380 | 876.4421 | 438.7247 | 875.4581 | 438.2327 | 8 |
| 20 | 2290.1265 | 1145.5669 | 2273.0999 | 1137.0536 | 2272.1159 | 1136.5616 | S | 737.3676 | 369.1874 | 720.3410 | 360.6742 | 719.3570 | 360.1821 | 7 |
| 21 | 2403.2105 | 1202.1089 | 2386.1840 | 1193.5956 | 2385.2000 | 1193.1036 | L | 650.3355 | 325.6714 | 633.3090 | 317.1581 | 632.3250 | 316.6661 | 6 |
| 22 | 2490.2426 | 1245.6249 | 2473.2160 | 1237.1116 | 2472.2320 | 1236.6196 | S | 537.2515 | 269.1294 | 520.2249 | 260.6161 | 519.2409 | 260.1241 | 5 |
| 23 | 2577.2746 | 1289.1409 | 2560.2480 | 1280.6277 | 2559.2640 | 1280.1356 | S | 450.2195 | 225.6134 | 433.1929 | 217.1001 | 432.2089 | 216.6081 | 4 |
| 24 | 2706.3172 | ***1353.6622*** | 2689.2906 | 1345.1490 | 2688.3066 | 1344.6569 | E | 363.1874 | 182.0974 | 346.1609 | 173.5841 | 345.1769 | 173.0921 | 3 |
| 25 | 2793.3492 | ***1397.1782*** | 2776.3227 | 1388.6650 | 2775.3386 | 1388.1730 | S | 234.1448 | 117.5761 | 217.1183 | 109.0628 | 216.1343 | 108.5708 | 2 |
| 26 | | | | | | | K | 147.1128 | 74.0600 | 130.0863 | 65.5468 | | | 1 |

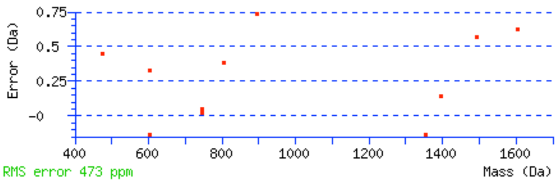


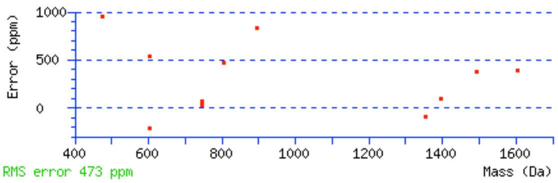

MS/MS Fragmentation of **AMNTGGFDSMQRQLEAELRSLSSESK**
Found in **P48563** in **Uniprot_yeastS288C**, Protein MON2 OS=Saccharomyces cerevisiae (strain ATCC 204508 / S288c) OX=559292 GN=MON2 PE=1 SV=1

Match to Query 16264: 2938.443747 from(980.488525,3+) intensity(359560.2500) rtinseconds(5900.0622) index(15023)
Title: 20100530_methyl300mM_3.16712.16712.3
Data file D:\Panyue\Proteomic dataset\20100530_methyl300mM_3.RAW

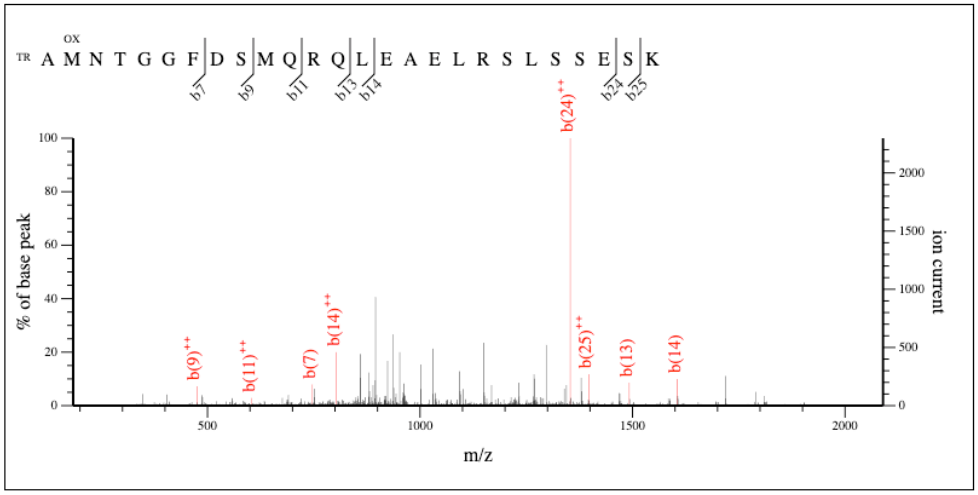


**Monoisotopic mass of neutral peptide Mr(calc):** 2938.4474
**Fixed modifications:** Carbamidomethyl (C) (apply to specified residues or termini only)
**Variable modifications:**
**N-term :** Trimethyl:2H(9) (Protein N-term)
**M2** : Oxidation (M), with neutral losses 0.0000(shown in table), 63.9983
**Ions Score:** 20 **Expect:** 0.12
**Matches :** 10/432 fragment ions using 18 most intense peaks (help)

| # | b | b++ | b* | b*++ | b0 | b0++ | Seq. | y | y++ | y* | y*++ | y0 | y0++ | # |
|---|---|---|---|---|---|---|---|---|---|---|---|---|---|---|
| 1 | 123.1478 | 62.0776 | | | | | A | | | | | | | 26 |
| 2 | 270.1832 | 135.5953 | | | | | M | 2817.3142 | 1409.1607 | 2800.2876 | 1400.6475 | 2799.3036 | 1400.1554 | 25 |
| 3 | 384.2262 | 192.6167 | 367.1996 | 184.1034 | | | N | 2670.2788 | 1335.6430 | 2653.2522 | 1327.1298 | 2652.2682 | 1326.6377 | 24 |
| 4 | 485.2738 | 243.1406 | 468.2473 | 234.6273 | 467.2633 | 234.1353 | T | 2556.2359 | 1278.6216 | 2539.2093 | 1270.1083 | 2538.2253 | 1269.6163 | 23 |
| 5 | 542.2953 | 271.6513 | 525.2688 | 263.1380 | 524.2847 | 262.6460 | G | 2455.1882 | 1228.0977 | 2438.1616 | 1219.5845 | 2437.1776 | 1219.0924 | 22 |
| 6 | 599.3168 | 300.1620 | 582.2902 | 291.6487 | 581.3062 | 291.1567 | G | 2398.1667 | 1199.5870 | 2381.1402 | 1191.0737 | 2380.1561 | 1190.5817 | 21 |
| 7 | **746.3852** | 373.6962 | 729.3586 | 365.1830 | 728.3746 | 364.6909 | F | 2341.1452 | 1171.0763 | 2324.1187 | 1162.5630 | 2323.1347 | 1162.0710 | 20 |
| 8 | 861.4121 | 431.2097 | 844.3856 | 422.6964 | 843.4016 | 422.2044 | D | 2194.0768 | 1097.5421 | 2177.0503 | 1089.0288 | 2176.0663 | 1088.5368 | 19 |
| 9 | 948.4441 | ***474.7257*** | 931.4176 | 466.2124 | 930.4336 | 465.7204 | S | 2079.0499 | 1040.0286 | 2062.0233 | 1031.5153 | 2061.0393 | 1031.0233 | 18 |
| 10 | 1079.4846 | 540.2460 | 1062.4581 | 531.7327 | 1061.4741 | 531.2407 | M | 1992.0179 | 996.5126 | 1974.9913 | 987.9993 | 1974.0073 | 987.5073 | 17 |
| 11 | 1207.5432 | ***604.2752*** | 1190.5167 | 595.7620 | 1189.5326 | 595.2700 | Q | 1860.9774 | 930.9923 | 1843.9508 | 922.4791 | 1842.9668 | 921.9870 | 16 |
| 12 | 1363.6443 | 682.3258 | 1346.6178 | 673.8125 | 1345.6338 | 673.3205 | R | 1732.9188 | 866.9630 | 1715.8923 | 858.4498 | 1714.9082 | 857.9578 | 15 |
| 13 | **1491.7029** | ***746.3551*** | 1474.6764 | 737.8418 | 1473.6923 | 737.3498 | Q | 1576.8177 | 788.9125 | 1559.7911 | 780.3992 | 1558.8071 | 779.9072 | 14 |
| 14 | **1604.7870** | ***802.8971*** | 1587.7604 | 794.3838 | 1586.7764 | 793.8918 | L | 1448.7591 | 724.8832 | 1431.7326 | 716.3699 | 1430.7485 | 715.8779 | 13 |
| 15 | 1733.8296 | 867.4184 | 1716.8030 | 858.9051 | 1715.8190 | 858.4131 | E | 1335.6750 | 668.3412 | 1318.6485 | 659.8279 | 1317.6645 | 659.3359 | 12 |
| 16 | 1804.8667 | 902.9370 | 1787.8401 | 894.4237 | 1786.8561 | 893.9317 | A | 1206.6325 | 603.8199 | 1189.6059 | 595.3066 | 1188.6219 | 594.8146 | 11 |
| 17 | 1933.9093 | 967.4583 | 1916.8827 | 958.9450 | 1915.8987 | 958.4530 | E | 1135.5953 | 568.3013 | 1118.5688 | 559.7880 | 1117.5848 | 559.2960 | 10 |
| 18 | 2046.9933 | 1024.0003 | 2029.9668 | 1015.4870 | 2028.9828 | 1014.9950 | L | 1006.5527 | 503.7800 | 989.5262 | 495.2667 | 988.5422 | 494.7747 | 9 |
| 19 | 2203.0944 | 1102.0509 | 2186.0679 | 1093.5376 | 2185.0839 | 1093.0456 | R | 893.4687 | 447.2380 | 876.4421 | 438.7247 | 875.4581 | 438.2327 | 8 |
| 20 | 2290.1265 | 1145.5669 | 2273.0999 | 1137.0536 | 2272.1159 | 1136.5616 | S | 737.3676 | 369.1874 | 720.3410 | 360.6742 | 719.3570 | 360.1821 | 7 |
| 21 | 2403.2105 | 1202.1089 | 2386.1840 | 1193.5956 | 2385.2000 | 1193.1036 | L | 650.3355 | 325.6714 | 633.3090 | 317.1581 | 632.3250 | 316.6661 | 6 |
| 22 | 2490.2426 | 1245.6249 | 2473.2160 | 1237.1116 | 2472.2320 | 1236.6196 | S | 537.2515 | 269.1294 | 520.2249 | 260.6161 | 519.2409 | 260.1241 | 5 |
| 23 | 2577.2746 | 1289.1409 | 2560.2480 | 1280.6277 | 2559.2640 | 1280.1356 | S | 450.2195 | 225.6134 | 433.1929 | 217.1001 | 432.2089 | 216.6081 | 4 |
| 24 | 2706.3172 | ***1353.6622*** | 2689.2906 | 1345.1490 | 2688.3066 | 1344.6569 | E | 363.1874 | 182.0974 | 346.1609 | 173.5841 | 345.1769 | 173.0921 | 3 |
| 25 | 2793.3492 | ***1397.1782*** | 2776.3227 | 1388.6650 | 2775.3386 | 1388.1730 | S | 234.1448 | 117.5761 | 217.1183 | 109.0628 | 216.1343 | 108.5708 | 2 |
| 26 | | | | | | | K | 147.1128 | 74.0600 | 130.0863 | 65.5468 | | | 1 |

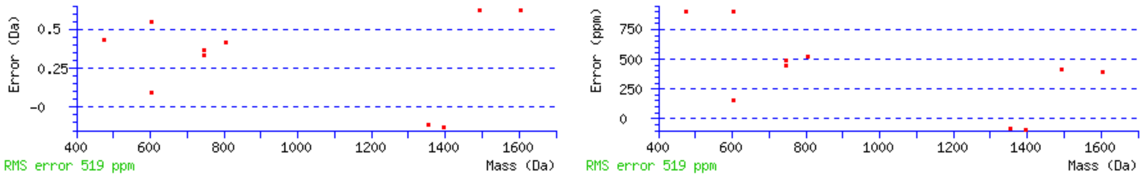

MS/MS Fragmentation of **AMNTGGFDSMQRQLEAELRSLSSESK**
Found in **P48563** in **Uniprot_yeastS288C**, Protein MON2 OS=Saccharomyces cerevisiae (strain ATCC 204508 / S288c) OX=559292 GN=MON2 PE=1 SV=1

Match to Query 16272: 2938.445397 from(980.489075,3+) intensity(180881.5000) rtinseconds(5352.1475) index(13734)
Title: 20100530_methyl300mM_3.15279.15279.3
Data file D:\Panyue\Proteomic dataset\20100530_methyl300mM_3.RAW

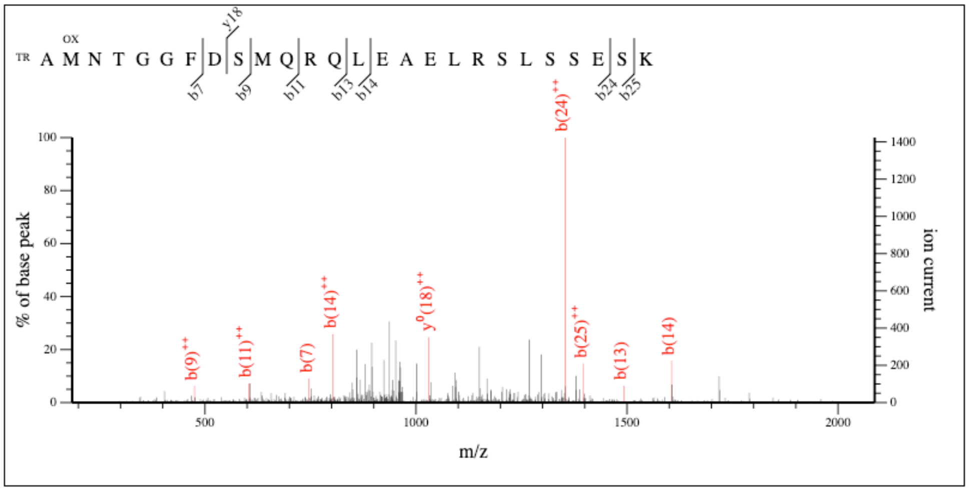

**Monoisotopic mass of neutral peptide Mr(calc):** 2938.4474
**Fixed modifications:** Carbamidomethyl (C) (apply to specified residues or termini only)
**Variable modifications:**
**N-term** : Trimethyl:2H(9) (Protein N-term)
**M2** : Oxidation (M), with neutral losses 0.0000(shown in table), 63.9983
**Ions Score:** 20 **Expect:** 0.11
**Matches** : 11/432 fragment ions using 18 most intense peaks (help)

| # | b | b++ | b* | b*++ | b0 | b0++ | Seq. | y | y++ | y* | y*++ | y0 | y0++ | # |
|---|---|---|---|---|---|---|---|---|---|---|---|---|---|---|
| 1 | 123.1478 | 62.0776 | | | | | A | | | | | | | 26 |
| 2 | 270.1832 | 135.5953 | | | | | M | 2817.3142 | 1409.1607 | 2800.2876 | 1400.6475 | 2799.3036 | 1400.1554 | 25 |
| 3 | 384.2262 | 192.6167 | 367.1996 | 184.1034 | | | N | 2670.2788 | 1335.6430 | 2653.2522 | 1327.1298 | 2652.2682 | 1326.6377 | 24 |
| 4 | 485.2738 | 243.1406 | 468.2473 | 234.6273 | 467.2633 | 234.1353 | T | 2556.2359 | 1278.6216 | 2539.2093 | 1270.1083 | 2538.2253 | 1269.6163 | 23 |
| 5 | 542.2953 | 271.6513 | 525.2688 | 263.1380 | 524.2847 | 262.6460 | G | 2455.1882 | 1228.0977 | 2438.1616 | 1219.5845 | 2437.1776 | 1219.0924 | 22 |
| 6 | 599.3168 | 300.1620 | 582.2902 | 291.6487 | 581.3062 | 291.1567 | G | 2398.1667 | 1199.5870 | 2381.1402 | 1191.0737 | 2380.1561 | 1190.5817 | 21 |
| 7 | **746.3852** | 373.6962 | 729.3586 | 365.1830 | 728.3746 | 364.6909 | F | 2341.1452 | 1171.0763 | 2324.1187 | 1162.5630 | 2323.1347 | 1162.0710 | 20 |
| 8 | 861.4121 | 431.2097 | 844.3856 | 422.6964 | 843.4016 | 422.2044 | D | 2194.0768 | 1097.5421 | 2177.0503 | 1089.0288 | 2176.0663 | 1088.5368 | 19 |
| 9 | 948.4441 | ***474.7257*** | 931.4176 | 466.2124 | 930.4336 | 465.7204 | S | 2079.0499 | 1040.0286 | 2062.0233 | 1031.5153 | 2061.0393 | 1031.0233 | 18 |
| 10 | 1079.4846 | 540.2460 | 1062.4581 | 531.7327 | 1061.4741 | 531.2407 | M | 1992.0179 | 996.5126 | 1974.9913 | 987.9993 | 1974.0073 | 987.5073 | 17 |
| 11 | 1207.5432 | ***604.2752*** | 1190.5167 | 595.7620 | 1189.5326 | 595.2700 | Q | 1860.9774 | 930.9923 | 1843.9508 | 922.4791 | 1842.9668 | 921.9870 | 16 |
| 12 | 1363.6443 | 682.3258 | 1346.6178 | 673.8125 | 1345.6338 | 673.3205 | R | 1732.9188 | 866.9630 | 1715.8923 | 858.4498 | 1714.9082 | 857.9578 | 15 |
| 13 | **1491.7029** | ***746.3551*** | 1474.6764 | 737.8418 | 1473.6923 | 737.3498 | Q | 1576.8177 | 788.9125 | 1559.7911 | 780.3992 | 1558.8071 | 779.9072 | 14 |
| 14 | **1604.7870** | ***802.8971*** | 1587.7604 | 794.3838 | 1586.7764 | 793.8918 | L | 1448.7591 | 724.8832 | 1431.7326 | 716.3699 | 1430.7485 | 715.8779 | 13 |
| 15 | 1733.8296 | 867.4184 | 1716.8030 | 858.9051 | 1715.8190 | 858.4131 | E | 1335.6750 | 668.3412 | 1318.6485 | 659.8279 | 1317.6645 | 659.3359 | 12 |
| 16 | 1804.8667 | 902.9370 | 1787.8401 | 894.4237 | 1786.8561 | 893.9317 | A | 1206.6325 | 603.8199 | 1189.6059 | 595.3066 | 1188.6219 | 594.8146 | 11 |
| 17 | 1933.9093 | 967.4583 | 1916.8827 | 958.9450 | 1915.8987 | 958.4530 | E | 1135.5953 | 568.3013 | 1118.5688 | 559.7880 | 1117.5848 | 559.2960 | 10 |
| 18 | 2046.9933 | 1024.0003 | 2029.9668 | 1015.4870 | 2028.9828 | 1014.9950 | L | 1006.5527 | 503.7800 | 989.5262 | 495.2667 | 988.5422 | 494.7747 | 9 |
| 19 | 2203.0944 | 1102.0509 | 2186.0679 | 1093.5376 | 2185.0839 | 1093.0456 | R | 893.4687 | 447.2380 | 876.4421 | 438.7247 | 875.4581 | 438.2327 | 8 |
| 20 | 2290.1265 | 1145.5669 | 2273.0999 | 1137.0536 | 2272.1159 | 1136.5616 | S | 737.3676 | 369.1874 | 720.3410 | 360.6742 | 719.3570 | 360.1821 | 7 |
| 21 | 2403.2105 | 1202.1089 | 2386.1840 | 1193.5956 | 2385.2000 | 1193.1036 | L | 650.3355 | 325.6714 | 633.3090 | 317.1581 | 632.3250 | 316.6661 | 6 |
| 22 | 2490.2426 | 1245.6249 | 2473.2160 | 1237.1116 | 2472.2320 | 1236.6196 | S | 537.2515 | 269.1294 | 520.2249 | 260.6161 | 519.2409 | 260.1241 | 5 |
| 23 | 2577.2746 | 1289.1409 | 2560.2480 | 1280.6277 | 2559.2640 | 1280.1356 | S | 450.2195 | 225.6134 | 433.1929 | 217.1001 | 432.2089 | 216.6081 | 4 |
| 24 | 2706.3172 | ***1353.6622*** | 2689.2906 | 1345.1490 | 2688.3066 | 1344.6569 | E | 363.1874 | 182.0974 | 346.1609 | 173.5841 | 345.1769 | 173.0921 | 3 |
| 25 | 2793.3492 | ***1397.1782*** | 2776.3227 | 1388.6650 | 2775.3386 | 1388.1730 | S | 234.1448 | 117.5761 | 217.1183 | 109.0628 | 216.1343 | 108.5708 | 2 |
| 26 | | | | | | | K | 147.1128 | 74.0600 | 130.0863 | 65.5468 | | | 1 |

MS/MS Fragmentation of **AMNTGGFDSMQRQLEAELRSLSSESK**
Found in **P48563** in **Uniprot_yeastS288C**, Protein MON2 OS=Saccharomyces cerevisiae (strain ATCC 204508 / S288c) OX=559292 GN=MON2 PE=1 SV=1

Match to Query 16429: 2954.433066 from(985.818298,3+) intensity(85402.8281) rtinseconds(5946.845) index(15134)
Title: 20100530_methyl300mM_3.16835.16835.3
Data file D:\Panyue\Proteomic dataset\20100530_methyl300mM_3.RAW

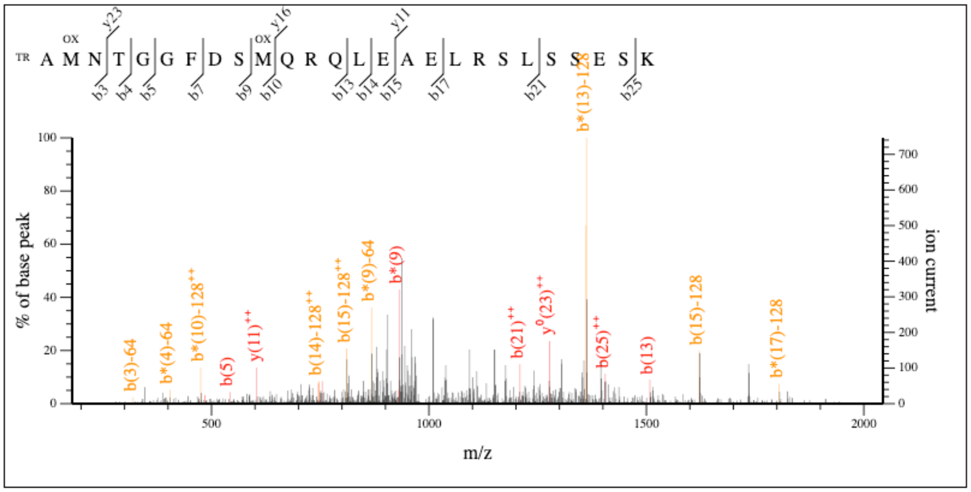


**Monoisotopic mass of neutral peptide Mr(calc):** 2954.4423
**Fixed modifications:** Carbamidomethyl (C) (apply to specified residues or termini only)
**Variable modifications:**
**N-term** : Trimethyl:2H(9) (Protein N-term)
**M2** : Oxidation (M), with neutral losses 0.0000(shown in table), 63.9983
**M10** : Oxidation (M), with neutral losses 0.0000(shown in table), 63.9983
**Ions Score:** 21 **Expect:** 0.021
**Matches** : 30/480 fragment ions using 34 most intense peaks (help)

| # | b | b++ | b* | b*++ | b0 | b0++ | Seq. | y | y++ | y* | y*++ | y0 | y0++ | # |
|---|---|---|---|---|---|---|---|---|---|---|---|---|---|---|
| 1 | 123.1478 | 62.0776 | | | | | A | | | | | | | 26 |
| 2 | 270.1832 | 135.5953 | | | | | M | 2833.3091 | 1417.1582 | 2816.2825 | 1408.6449 | 2815.2985 | 1408.1529 | 25 |
| 3 | 384.2262 | 192.6167 | 367.1996 | 184.1034 | | | N | 2686.2737 | 1343.6405 | 2669.2471 | 1335.1272 | 2668.2631 | 1334.6352 | 24 |
| 4 | 485.2738 | 243.1406 | 468.2473 | 234.6273 | 467.2633 | 234.1353 | T | 2572.2308 | 1286.6190 | 2555.2042 | 1278.1057 | 2554.2202 | 1277.6137 | 23 |
| 5 | 542.2953 | 271.6513 | 525.2688 | 263.1380 | 524.2847 | 262.6460 | G | 2471.1831 | 1236.0952 | 2454.1565 | 1227.5819 | 2453.1725 | 1227.0899 | 22 |
| 6 | 599.3168 | 300.1620 | 582.2902 | 291.6487 | 581.3062 | 291.1567 | G | 2414.1616 | 1207.5845 | 2397.1351 | 1199.0712 | 2396.1511 | 1198.5792 | 21 |
| 7 | 746.3852 | 373.6962 | 729.3586 | 365.1830 | 728.3746 | 364.6909 | F | 2357.1402 | 1179.0737 | 2340.1136 | 1170.5604 | 2339.1296 | 1170.0684 | 20 |
| 8 | 861.4121 | 431.2097 | 844.3856 | 422.6964 | 843.4016 | 422.2044 | D | 2210.0717 | 1105.5395 | 2193.0452 | 1097.0262 | 2192.0612 | 1096.5342 | 19 |
| 9 | 948.4441 | 474.7257 | 931.4176 | 466.2124 | 930.4336 | 465.7204 | S | 2095.0448 | 1048.0260 | 2078.0183 | 1039.5128 | 2077.0342 | 1039.0208 | 18 |
| 10 | 1095.4795 | 548.2434 | 1078.4530 | 539.7301 | 1077.4690 | 539.2381 | M | 2008.0128 | 1004.5100 | 1990.9862 | 995.9968 | 1990.0022 | 995.5047 | 17 |
| 11 | 1223.5381 | 612.2727 | 1206.5116 | 603.7594 | 1205.5276 | 603.2674 | Q | 1860.9774 | 930.9923 | 1843.9508 | 922.4791 | 1842.9668 | 921.9870 | 16 |
| 12 | 1379.6392 | 690.3233 | 1362.6127 | 681.8100 | 1361.6287 | 681.3180 | R | 1732.9188 | 866.9630 | 1715.8923 | 858.4498 | 1714.9082 | 857.9578 | 15 |
| 13 | 1507.6978 | 754.3525 | 1490.6713 | 745.8393 | 1489.6873 | 745.3473 | Q | 1576.8177 | 788.9125 | 1559.7911 | 780.3992 | 1558.8071 | 779.9072 | 14 |
| 14 | 1620.7819 | 810.8946 | 1603.7553 | 802.3813 | 1602.7713 | 801.8893 | L | 1448.7591 | 724.8832 | 1431.7326 | 716.3699 | 1430.7485 | 715.8779 | 13 |
| 15 | 1749.8245 | 875.4159 | 1732.7979 | 866.9026 | 1731.8139 | 866.4106 | E | 1335.6750 | 668.3412 | 1318.6485 | 659.8279 | 1317.6645 | 659.3359 | 12 |
| 16 | 1820.8616 | 910.9344 | 1803.8350 | 902.4212 | 1802.8510 | 901.9291 | A | 1206.6325 | 603.8199 | 1189.6059 | 595.3066 | 1188.6219 | 594.8146 | 11 |
| 17 | 1949.9042 | 975.4557 | 1932.8776 | 966.9425 | 1931.8936 | 966.4504 | E | 1135.5953 | 568.3013 | 1118.5688 | 559.7880 | 1117.5848 | 559.2960 | 10 |
| 18 | 2062.9882 | 1031.9978 | 2045.9617 | 1023.4845 | 2044.9777 | 1022.9925 | L | 1006.5527 | 503.7800 | 989.5262 | 495.2667 | 988.5422 | 494.7747 | 9 |
| 19 | 2219.0894 | 1110.0483 | 2202.0628 | 1101.5350 | 2201.0788 | 1101.0430 | R | 893.4687 | 447.2380 | 876.4421 | 438.7247 | 875.4581 | 438.2327 | 8 |
| 20 | 2306.1214 | 1153.5643 | 2289.0948 | 1145.0511 | 2288.1108 | 1144.5590 | S | 737.3676 | 369.1874 | 720.3410 | 360.6742 | 719.3570 | 360.1821 | 7 |
| 21 | 2419.2054 | 1210.1064 | 2402.1789 | 1201.5931 | 2401.1949 | 1201.1011 | L | 650.3355 | 325.6714 | 633.3090 | 317.1581 | 632.3250 | 316.6661 | 6 |
| 22 | 2506.2375 | 1253.6224 | 2489.2109 | 1245.1091 | 2488.2269 | 1244.6171 | S | 537.2515 | 269.1294 | 520.2249 | 260.6161 | 519.2409 | 260.1241 | 5 |
| 23 | 2593.2695 | 1297.1384 | 2576.2430 | 1288.6251 | 2575.2589 | 1288.1331 | S | 450.2195 | 225.6134 | 433.1929 | 217.1001 | 432.2089 | 216.6081 | 4 |
| 24 | 2722.3121 | 1361.6597 | 2705.2855 | 1353.1464 | 2704.3015 | 1352.6544 | E | 363.1874 | 182.0974 | 346.1609 | 173.5841 | 345.1769 | 173.0921 | 3 |
| 25 | 2809.3441 | 1405.1757 | 2792.3176 | 1396.6624 | 2791.3336 | 1396.1704 | S | 234.1448 | 117.5761 | 217.1183 | 109.0628 | 216.1343 | 108.5708 | 2 |
| 26 | | | | | | | K | 147.1128 | 74.0600 | 130.0863 | 65.5468 | | | 1 |

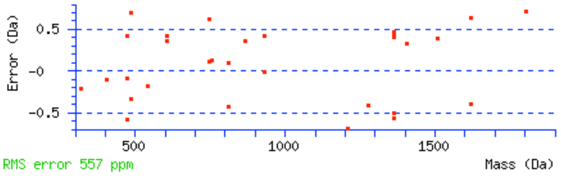


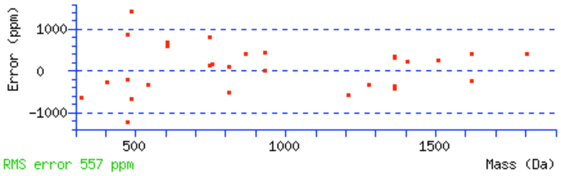


### 45. Nop14 (2 peptides in 300mM fraction 3)

MS/MS Fragmentation of **AGSQLKNLK**
Found in **Q99207** in **Uniprot_yeastS288C**, Nucleolar complex protein 14 OS=Saccharomyces cerevisiae (strain ATCC 204508 / S288c) OX=559292 GN=NOP14 PE=1 SV=1

Match to Query 4522: 1008.669970 from(1009.677246,1+) intensity(578424.0625) rtinseconds(1693.4415) index(4359)
Title: 20100530_methyl300mM_3.4850.4850.1
Data file D:\Panyue\Proteomic dataset\20100530_methyl300mM_3.RAW

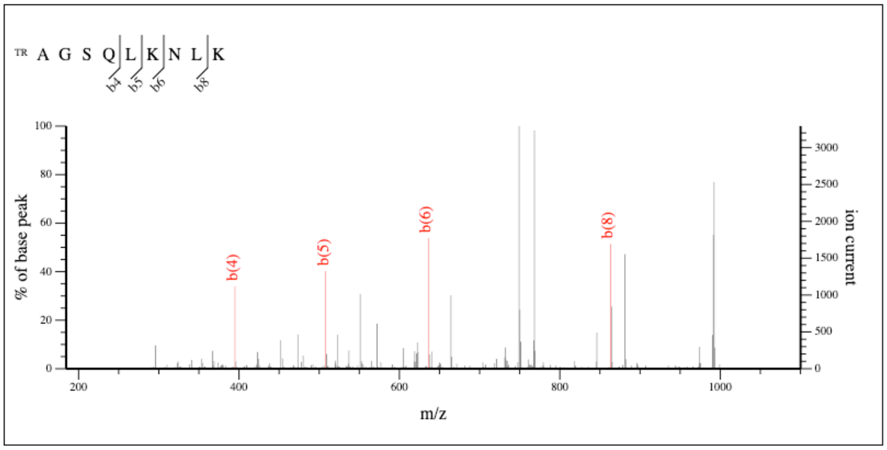

**Monoisotopic mass of neutral peptide Mr(calc):** 1008.6641
**Fixed modifications:** Carbamidomethyl (C) (apply to specified residues or termini only)
**Variable modifications:**
**N-term :** Trimethyl:2H(9) (Protein N-term)
**Ions Score:** 27 **Expect:** 0.0041
**Matches :** 4/37 fragment ions using 8 most intense peaks (help)

| # | b | b* | b$^0$ | Seq. | y | y* | y$^0$ | # |
|---|---|---|---|---|---|---|---|---|
| 1 | 123.1478 | | | A | | | | 9 |
| 2 | 180.1693 | | | G | 887.5309 | 870.5043 | 869.5203 | 8 |
| 3 | 267.2013 | | 249.1908 | S | 830.5094 | 813.4829 | 812.4989 | 7 |
| 4 | ***395.2599*** | 378.2334 | 377.2493 | Q | 743.4774 | 726.4509 | | 6 |
| 5 | ***508.3440*** | 491.3174 | 490.3334 | L | 615.4188 | 598.3923 | | 5 |
| 6 | ***636.4389*** | 619.4124 | 618.4284 | K | 502.3348 | 485.3082 | | 4 |
| 7 | 750.4819 | 733.4553 | 732.4713 | N | 374.2398 | 357.2132 | | 3 |
| 8 | ***863.5659*** | 846.5394 | 845.5554 | L | 260.1969 | 243.1703 | | 2 |
| 9 | | | | K | 147.1128 | 130.0863 | | 1 |

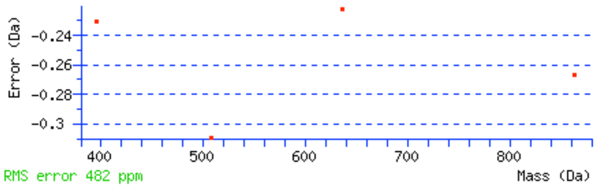


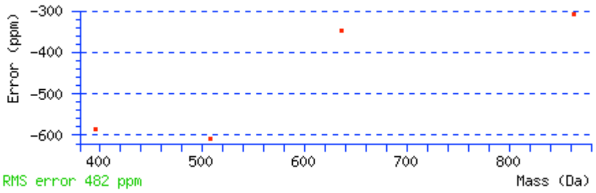


MS/MS Fragmentation of **AGSQLKNLK**
Found in **Q99207** in **Uniprot_yeastS288C**, Nucleolar complex protein 14 OS=Saccharomyces cerevisiae (strain ATCC 204508 / S288c) OX=559292 GN=NOP14 PE=1 SV=1

Match to Query 4523: 1008.670031 from(1009.677307,1+) intensity(218454.9219) rtinseconds(1691.8806) index(4355)
Title: 20100530_methyl300mM_3.4845.4845.1
Data file D:\Panyue\Proteomic dataset\20100530_methyl300mM_3.RAW

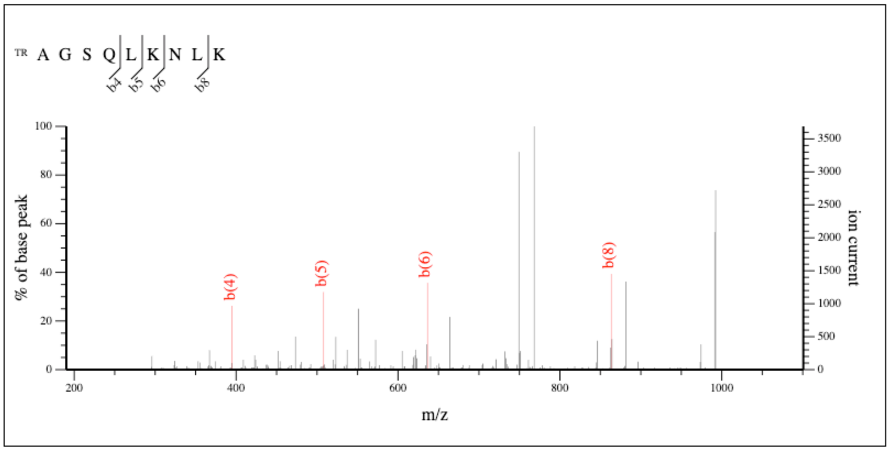

**Monoisotopic mass of neutral peptide Mr(calc):** 1008.6641
**Fixed modifications:** Carbamidomethyl (C) (apply to specified residues or termini only)
**Variable modifications:**
**N-term :** Trimethyl:2H(9) (Protein N-term)
**Ions Score:** 27 **Expect:** 0.0041
**Matches :** 4/37 fragment ions using 8 most intense peaks (help)

| # | b | b* | b0 | Seq. | y | y* | y0 | # |
|---|---|---|---|---|---|---|---|---|
| 1 | 123.1478 | | | A | | | | 9 |
| 2 | 180.1693 | | | G | 887.5309 | 870.5043 | 869.5203 | 8 |
| 3 | 267.2013 | | 249.1908 | S | 830.5094 | 813.4829 | 812.4989 | 7 |
| 4 | ***395.2599*** | 378.2334 | 377.2493 | Q | 743.4774 | 726.4509 | | 6 |
| 5 | ***508.3440*** | 491.3174 | 490.3334 | L | 615.4188 | 598.3923 | | 5 |
| 6 | ***636.4389*** | 619.4124 | 618.4284 | K | 502.3348 | 485.3082 | | 4 |
| 7 | 750.4819 | 733.4553 | 732.4713 | N | 374.2398 | 357.2132 | | 3 |
| 8 | ***863.5659*** | 846.5394 | 845.5554 | L | 260.1969 | 243.1703 | | 2 |
| 9 | | | | K | 147.1128 | 130.0863 | | 1 |

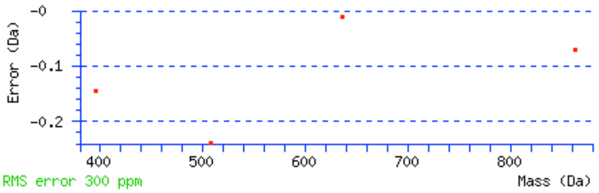


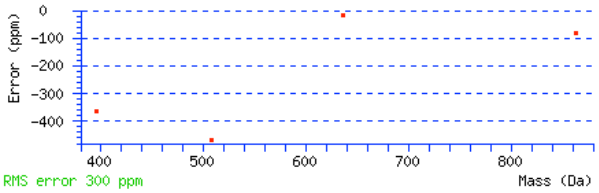


### 46. Eft1 (3 peptides in 300mM fraction 3)

MS/MS Fragmentation of **VAFTVDQMRSLMDK**
Found in **P32324** in **Uniprot_yeastS288C**, Elongation factor 2 OS=Saccharomyces cerevisiae (strain ATCC 204508 / S288c) OX=559292 GN=EFT1 PE=1 SV=1

Match to Query 9478: 1706.871190 from(854.442871,2+) intensity(892155.6875) rtinseconds(2674.9329) index(7022)
Title: 20100530_methyl300mM_3.7813.7813.2
Data file D:\Panyue\Proteomic dataset\20100530_methyl300mM_3.RAW

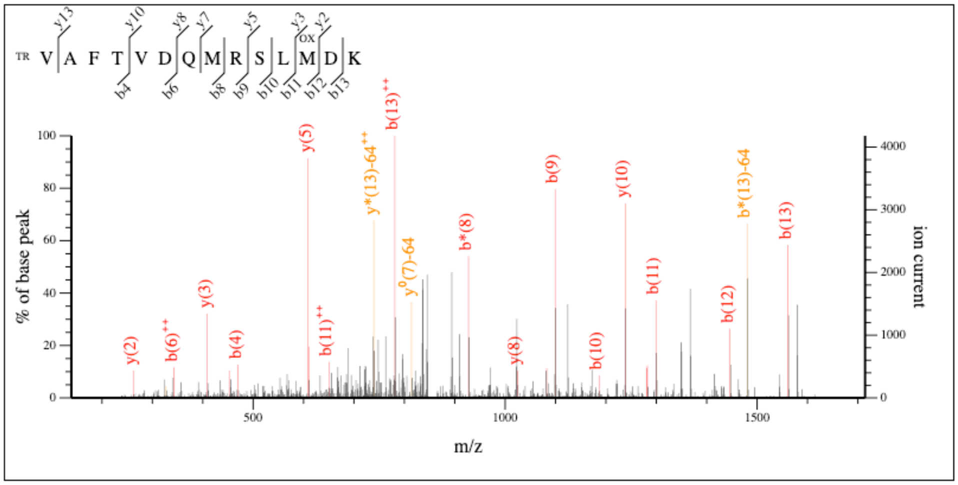

**Monoisotopic mass of neutral peptide Mr(calc):** 1706.9046
**Fixed modifications:** Carbamidomethyl (C) (apply to specified residues or termini only)
**Variable modifications:**
**N-term :** Trimethyl:2H(9) (Protein N-term)
**M12** : Oxidation (M), with neutral losses 0.0000(shown in table), 63.9983
**Ions Score:** 21 **Expect:** 0.12
**Matches :** 27/214 fragment ions using 56 most intense peaks (help)

| # | b | $b^{++}$ | b* | $b^{*++}$ | $b^0$ | $b^{0++}$ | Seq. | y | $y^{++}$ | y* | $y^{*++}$ | $y^0$ | $y^{0++}$ | # |
|---|---|---|---|---|---|---|---|---|---|---|---|---|---|---|
| 1 | 151.1791 | 76.0932 | | | | | V | | | | | | | 14 |
| 2 | 222.2162 | 111.6118 | | | | | A | 1557.7400 | 779.3736 | 1540.7134 | 770.8604 | 1539.7294 | 770.3683 | 13 |
| 3 | 369.2847 | 185.1460 | | | | | F | 1486.7029 | 743.8551 | 1469.6763 | 735.3418 | 1468.6923 | 734.8498 | 12 |
| 4 | ***470.3323*** | 235.6698 | | | 452.3218 | 226.6645 | T | 1339.6344 | 670.3209 | 1322.6079 | 661.8076 | 1321.6239 | 661.3156 | 11 |
| 5 | 569.4008 | 285.2040 | | | 551.3902 | 276.1987 | V | ***1238.5868*** | 619.7970 | 1221.5602 | 611.2837 | 1220.5762 | 610.7917 | 10 |
| 6 | 684.4277 | **342.7175** | | | 666.4171 | 333.7122 | D | 1139.5184 | 570.2628 | 1122.4918 | 561.7495 | 1121.5078 | 561.2575 | 9 |
| 7 | 812.4863 | 406.7468 | 795.4597 | 398.2335 | 794.4757 | 397.7415 | Q | ***1024.4914*** | 512.7493 | 1007.4649 | 504.2361 | 1006.4808 | 503.7441 | 8 |
| 8 | 943.5268 | 472.2670 | 926.5002 | 463.7537 | 925.5162 | 463.2617 | M | 896.4328 | 448.7201 | 879.4063 | 440.2068 | 878.4223 | 439.7148 | 7 |
| 9 | ***1099.6279*** | 550.3176 | 1082.6013 | 541.8043 | 1081.6173 | 541.3123 | R | 765.3924 | 383.1998 | 748.3658 | 374.6865 | 747.3818 | 374.1945 | 6 |
| 10 | ***1186.6599*** | 593.8336 | 1169.6333 | 585.3203 | 1168.6493 | 584.8283 | S | ***609.2912*** | 305.1493 | 592.2647 | 296.6360 | 591.2807 | 296.1440 | 5 |
| 11 | ***1299.7440*** | **650.3756** | 1282.7174 | 641.8623 | 1281.7334 | 641.3703 | L | 522.2592 | 261.6332 | 505.2327 | 253.1200 | 504.2486 | 252.6280 | 4 |
| 12 | ***1446.7794*** | 723.8933 | 1429.7528 | 715.3800 | 1428.7688 | 714.8880 | M | ***409.1751*** | 205.0912 | 392.1486 | 196.5779 | 391.1646 | 196.0859 | 3 |
| 13 | ***1561.8063*** | **781.4068** | 1544.7798 | 772.8935 | 1543.7957 | 772.4015 | D | ***262.1397*** | 131.5735 | 245.1132 | 123.0602 | 244.1292 | 122.5682 | 2 |
| 14 | | | | | | | K | 147.1128 | 74.0600 | 130.0863 | 65.5468 | | | 1 |

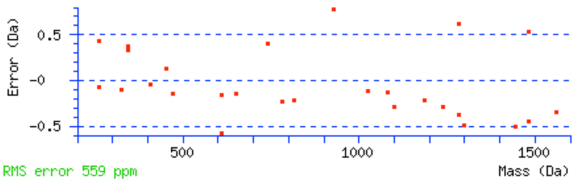


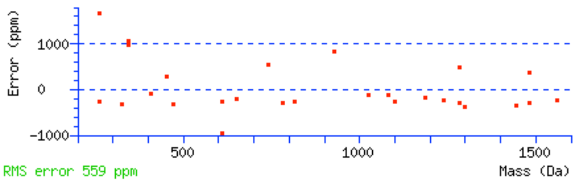


MS/MS Fragmentation of **VAFTVDQMRSLMDK**
Found in **P32324** in **Uniprot_yeastS288C**, Elongation factor 2 OS=Saccharomyces cerevisiae (strain ATCC 204508 / S288c) OX=559292 GN=EFT1 PE=1 SV=1

Match to Query 9479: 1706.871556 from(854.443054,2+) intensity(2018150.8750) rtinseconds(2676.8971) index(7027)
Title: 20100530_methyl300mM_3.7819.7819.2
Data file D:\Panyue\Proteomic dataset\20100530_methyl300mM_3.RAW

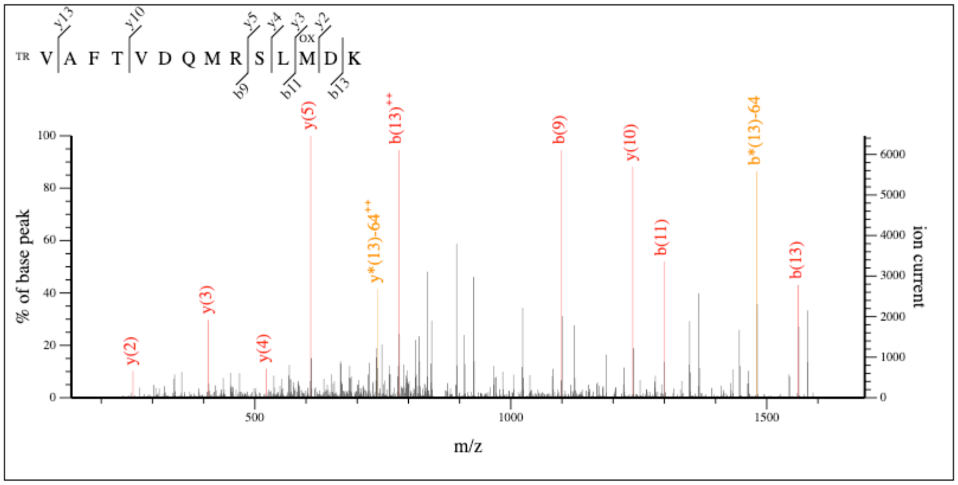

**Monoisotopic mass of neutral peptide Mr(calc):** 1706.9046
**Fixed modifications:** Carbamidomethyl (C) (apply to specified residues or termini only)
**Variable modifications:**
**N-term :** Trimethyl:2H(9) (Protein N-term)
**M12** : Oxidation (M), with neutral losses 0.0000(shown in table), 63.9983
**Ions Score:** 24 **Expect:** 0.071
**Matches :** 13/214 fragment ions using 14 most intense peaks (help)

| # | b | b++ | b* | b*++ | b0 | b0++ | Seq. | y | y++ | y* | y*++ | y0 | y0++ | # |
|---|---|---|---|---|---|---|---|---|---|---|---|---|---|---|
| 1 | 151.1791 | 76.0932 | | | | | V | | | | | | | 14 |
| 2 | 222.2162 | 111.6118 | | | | | A | 1557.7400 | 779.3736 | 1540.7134 | 770.8604 | 1539.7294 | 770.3683 | 13 |
| 3 | 369.2847 | 185.1460 | | | | | F | 1486.7029 | 743.8551 | 1469.6763 | 735.3418 | 1468.6923 | 734.8498 | 12 |
| 4 | 470.3323 | 235.6698 | | | 452.3218 | 226.6645 | T | 1339.6344 | 670.3209 | 1322.6079 | 661.8076 | 1321.6239 | 661.3156 | 11 |
| 5 | 569.4008 | 285.2040 | | | 551.3902 | 276.1987 | V | *1238.5868* | 619.7970 | 1221.5602 | 611.2837 | 1220.5762 | 610.7917 | 10 |
| 6 | 684.4277 | 342.7175 | | | 666.4171 | 333.7122 | D | 1139.5184 | 570.2628 | 1122.4918 | 561.7495 | 1121.5078 | 561.2575 | 9 |
| 7 | 812.4863 | 406.7468 | 795.4597 | 398.2335 | 794.4757 | 397.7415 | Q | 1024.4914 | 512.7493 | 1007.4649 | 504.2361 | 1006.4808 | 503.7441 | 8 |
| 8 | 943.5268 | 472.2670 | 926.5002 | 463.7537 | 925.5162 | 463.2617 | M | 896.4328 | 448.7201 | 879.4063 | 440.2068 | 878.4223 | 439.7148 | 7 |
| 9 | **1099.6279** | 550.3176 | 1082.6013 | 541.8043 | 1081.6173 | 541.3123 | R | 765.3924 | 383.1998 | 748.3658 | 374.6865 | 747.3818 | 374.1945 | 6 |
| 10 | 1186.6599 | 593.8336 | 1169.6333 | 585.3203 | 1168.6493 | 584.8283 | S | *609.2912* | 305.1493 | 592.2647 | 296.6360 | 591.2807 | 296.1440 | 5 |
| 11 | **1299.7440** | 650.3756 | 1282.7174 | 641.8623 | 1281.7334 | 641.3703 | L | *522.2592* | 261.6332 | 505.2327 | 253.1200 | 504.2486 | 252.6280 | 4 |
| 12 | 1446.7794 | 723.8933 | 1429.7528 | 715.3800 | 1428.7688 | 714.8880 | M | *409.1751* | 205.0912 | 392.1486 | 196.5779 | 391.1646 | 196.0859 | 3 |
| 13 | **1561.8063** | 781.4068 | 1544.7798 | 772.8935 | 1543.7957 | 772.4015 | D | *262.1397* | 131.5735 | 245.1132 | 123.0602 | 244.1292 | 122.5682 | 2 |
| 14 | | | | | | | K | 147.1128 | 74.0600 | 130.0863 | 65.5468 | | | 1 |

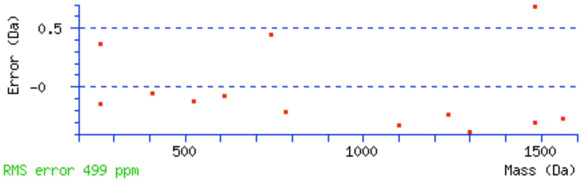


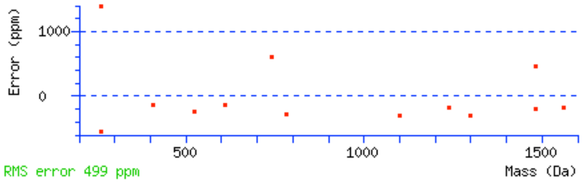


MS/MS Fragmentation of **VAFTVDQMRSLMDK**
Found in **P32324** in **Uniprot_yeastS288C**, Elongation factor 2 OS=Saccharomyces cerevisiae (strain ATCC 204508 / S288c) OX=559292 GN=EFT1 PE=1 SV=1

Match to Query 9480: 1706.873144 from(854.443848,2+) intensity(892155.6875) rtinseconds(2675.4829) index(7024)
Title: 20100530_methyl1300mM_3.7815.7815.2
Data file D:\Panyue\Proteomic dataset\20100530_methyl1300mM_3.RAW

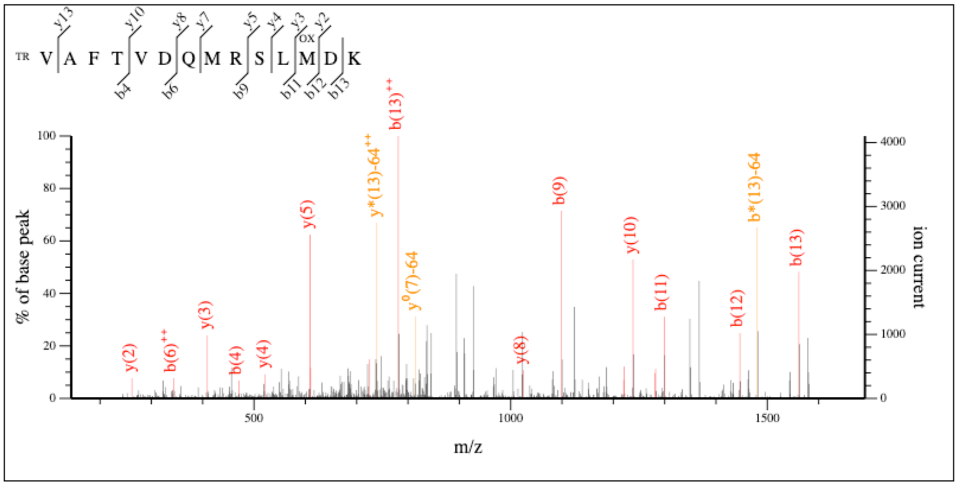

**Monoisotopic mass of neutral peptide Mr(calc):** 1706.9046
**Fixed modifications:** Carbamidomethyl (C) (apply to specified residues or termini only)
**Variable modifications:**
**N-term :** Trimethyl:2H(9) (Protein N-term)
**M12** : Oxidation (M), with neutral losses 0.0000(shown in table), 63.9983
**Ions Score:** 25 **Expect:** 0.047
**Matches :** 25/214 fragment ions using 42 most intense peaks (help)

| # | b | b++ | b* | b*++ | b0 | b0++ | Seq. | y | y++ | y* | y*++ | y0 | y0++ | # |
|---|---|---|---|---|---|---|---|---|---|---|---|---|---|---|
| 1 | 151.1791 | 76.0932 | | | | | V | | | | | | | 14 |
| 2 | 222.2162 | 111.6118 | | | | | A | 1557.7400 | 779.3736 | 1540.7134 | 770.8604 | 1539.7294 | 770.3683 | 13 |
| 3 | 369.2847 | 185.1460 | | | | | F | 1486.7029 | 743.8551 | 1469.6763 | 735.3418 | 1468.6923 | 734.8498 | 12 |
| 4 | ***470.3323*** | 235.6698 | | | 452.3218 | 226.6645 | T | 1339.6344 | 670.3209 | 1322.6079 | 661.8076 | 1321.6239 | 661.3156 | 11 |
| 5 | 569.4008 | 285.2040 | | | 551.3902 | 276.1987 | V | ***1238.5868*** | 619.7970 | 1221.5602 | 611.2837 | 1220.5762 | 610.7917 | 10 |
| 6 | 684.4277 | **342.7175** | | | 666.4171 | 333.7122 | D | 1139.5184 | 570.2628 | 1122.4918 | 561.7495 | 1121.5078 | 561.2575 | 9 |
| 7 | 812.4863 | 406.7468 | 795.4597 | 398.2335 | 794.4757 | 397.7415 | Q | ***1024.4914*** | 512.7493 | 1007.4649 | 504.2361 | 1006.4808 | 503.7441 | 8 |
| 8 | 943.5268 | 472.2670 | 926.5002 | 463.7537 | 925.5162 | 463.2617 | M | 896.4328 | 448.7201 | 879.4063 | 440.2068 | 878.4223 | 439.7148 | 7 |
| 9 | ***1099.6279*** | 550.3176 | 1082.6013 | 541.8043 | 1081.6173 | 541.3123 | R | 765.3924 | 383.1998 | 748.3658 | 374.6865 | 747.3818 | 374.1945 | 6 |
| 10 | 1186.6599 | 593.8336 | 1169.6333 | 585.3203 | 1168.6493 | 584.8283 | S | ***609.2912*** | 305.1493 | 592.2647 | 296.6360 | 591.2807 | 296.1440 | 5 |
| 11 | ***1299.7440*** | 650.3756 | 1282.7174 | 641.8623 | 1281.7334 | 641.3703 | L | ***522.2592*** | 261.6332 | 505.2327 | 253.1200 | 504.2486 | 252.6280 | 4 |
| 12 | ***1446.7794*** | **723.8933** | 1429.7528 | 715.3800 | 1428.7688 | 714.8880 | M | ***409.1751*** | 205.0912 | 392.1486 | 196.5779 | 391.1646 | 196.0859 | 3 |
| 13 | ***1561.8063*** | **781.4068** | 1544.7798 | 772.8935 | 1543.7957 | 772.4015 | D | ***262.1397*** | 131.5735 | 245.1132 | 123.0602 | 244.1292 | 122.5682 | 2 |
| 14 | | | | | | | K | 147.1128 | 74.0600 | 130.0863 | 65.5468 | | | 1 |

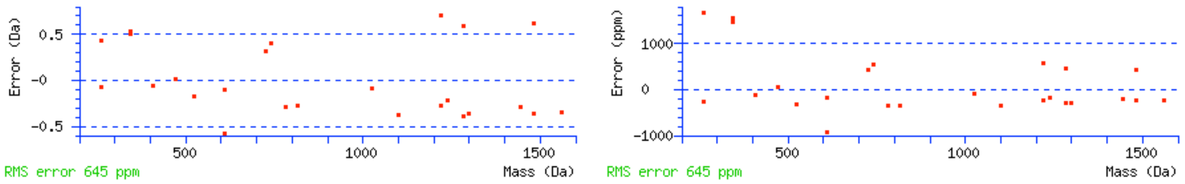


### 47. Rfa1 (1 peptide in 350mM fraction 1)

MS/MS Fragmentation of **SSVQLSR**
Found in **P22336** in **Uniprot_yeastS288C**, Replication factor A protein 1 OS=Saccharomyces cerevisiae (strain ATCC 204508 / S288c) OX=559292 GN=RFA1 PE=1 SV=1

Match to Query 2478: 826.525448 from(414.270000,2+) intensity(13124.7412) rtinseconds(7382.8074) index(18698)
Title: 20100530_methyl350mM_1.20799.20799.
Data file D:\Panyue\Proteomic dataset\20100530_methyl350mM_1.RAW

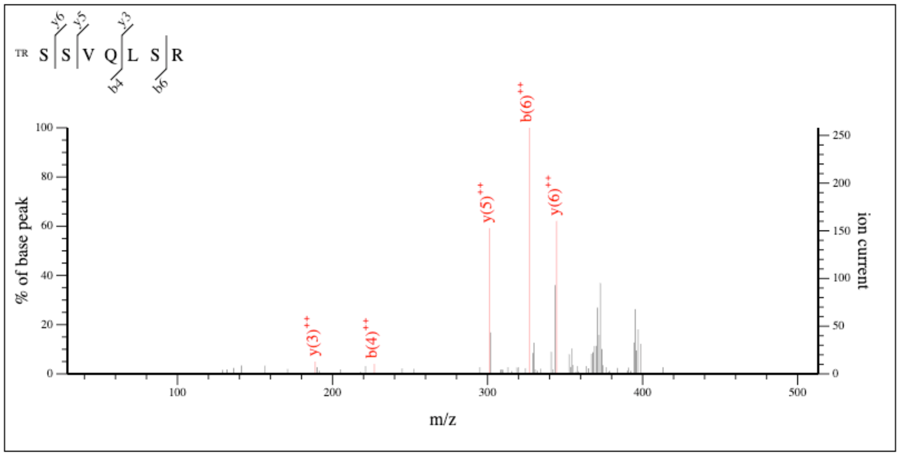

**Monoisotopic mass of neutral peptide Mr(calc):** 826.5222
**Fixed modifications:** Carbamidomethyl (C) (apply to specified residues or termini only)
**Variable modifications:**
**N-term :** Trimethyl:2H(9) (Protein N-term)
**Ions Score:** 25 **Expect:** 0.034
**Matches :** 5/64 fragment ions using 6 most intense peaks (help)

| # | b | b++ | b* | b*++ | b0 | b0++ | Seq. | y | y++ | y* | y*++ | y0 | y0++ | # |
|---|---|---|---|---|---|---|---|---|---|---|---|---|---|---|
| 1 | 139.1427 | 70.0750 | | | 121.1322 | 61.0697 | S | | | | | | | 7 |
| 2 | 226.1748 | 113.5910 | | | 208.1642 | 104.5857 | S | 689.3941 | ***345.2007*** | 672.3675 | 336.6874 | 671.3835 | 336.1954 | 6 |
| 3 | 325.2432 | 163.1252 | | | 307.2326 | 154.1199 | V | 602.3620 | ***301.6847*** | 585.3355 | 293.1714 | 584.3515 | 292.6794 | 5 |
| 4 | 453.3018 | ***227.1545*** | 436.2752 | 218.6412 | 435.2912 | 218.1492 | Q | 503.2936 | 252.1504 | 486.2671 | 243.6372 | 485.2831 | 243.1452 | 4 |
| 5 | 566.3858 | 283.6966 | 549.3593 | 275.1833 | 548.3753 | 274.6913 | L | 375.2350 | ***188.1212*** | 358.2085 | 179.6079 | 357.2245 | 179.1159 | 3 |
| 6 | 653.4179 | ***327.2126*** | 636.3913 | 318.6993 | 635.4073 | 318.2073 | S | 262.1510 | 131.5791 | 245.1244 | 123.0659 | 244.1404 | 122.5738 | 2 |
| 7 | | | | | | | R | 175.1190 | 88.0631 | 158.0924 | 79.5498 | | | 1 |

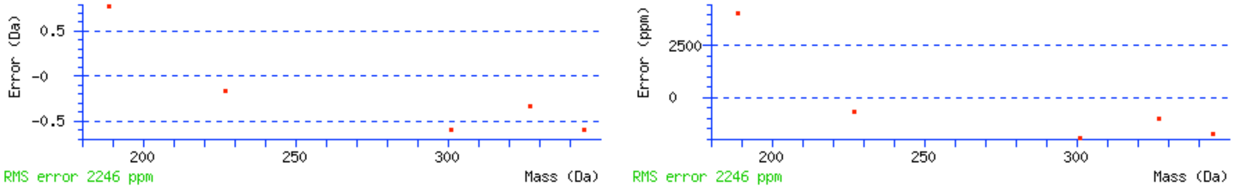


### 48. Pdc6 (2 peptides in 350mM fraction 1)

MS/MS Fragmentation of **SEITLGK**
Found in **P26263** in **Uniprot_yeastS288C**, Pyruvate decarboxylase isozyme 3 OS=Saccharomyces cerevisiae (strain ATCC 204508 / S288c) OX=559292 GN=PDC6 PE=1 SV=3

Match to Query 2066: 780.448718 from(781.455994,1+) intensity(675320.9375) rtinseconds(986.8442) index(2468)
Title: 20100530_methyl350mM_1.2745.2745.1
Data file D:\Panyue\Proteomic dataset\20100530_methyl350mM_1.RAW

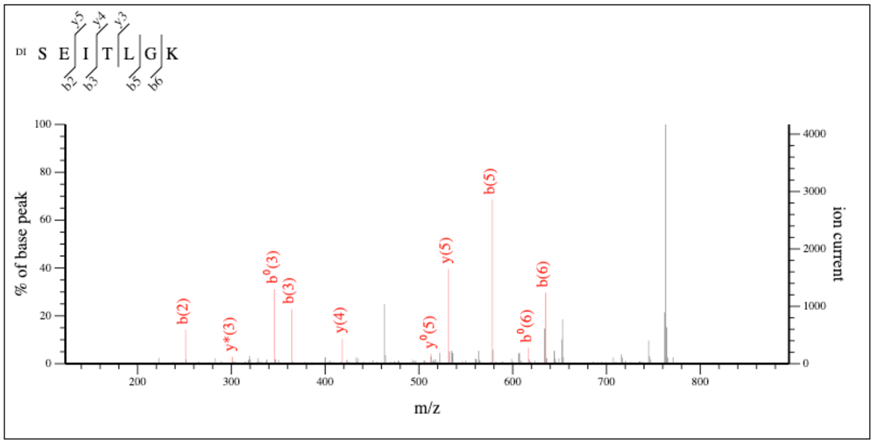

**Monoisotopic mass of neutral peptide Mr(calc):** 780.4864
**Fixed modifications:** Carbamidomethyl (C) (apply to specified residues or termini only)
**Variable modifications:**
**N-term :** Dimethyl:2H(6) (Protein N-term)
**Ions Score:** 26 **Expect:** 0.029
**Matches :** 10/27 fragment ions using 18 most intense peaks (help)

| # | b | $b^0$ | Seq. | y | y* | $y^0$ | # |
|---|---|---|---|---|---|---|---|
| 1 | 122.1083 | 104.0977 | S | | | | 7 |
| 2 | ***251.1509*** | 233.1403 | E | 660.3927 | 643.3661 | 642.3821 | 6 |
| 3 | ***364.2349*** | 346.2244 | I | **531.3501** | 514.3235 | 513.3395 | 5 |
| 4 | 465.2826 | 447.2720 | T | **418.2660** | 401.2395 | 400.2554 | 4 |
| 5 | ***578.3667*** | 560.3561 | L | 317.2183 | 300.1918 | | 3 |
| 6 | ***635.3881*** | 617.3776 | G | 204.1343 | 187.1077 | | 2 |
| 7 | | | K | 147.1128 | 130.0863 | | 1 |

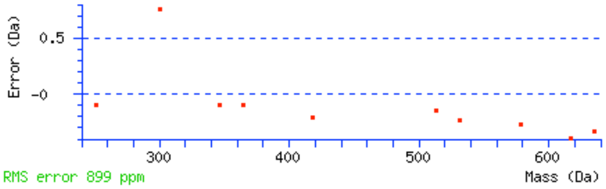


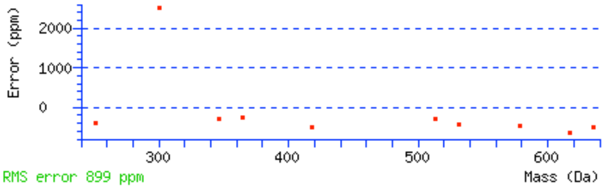


MS/MS Fragmentation of **SEITLGK**
Found in **P26263** in **Uniprot_yeastS288C**, Pyruvate decarboxylase isozyme 3 OS=Saccharomyces cerevisiae (strain ATCC 204508 / S288c) OX=559292 GN=PDC6 PE=1 SV=3

Match to Query 2065: 780.448596 from(781.455872,1+) intensity(1431024.3750) rtinseconds(989.4157) index(2475)
Title: 20100530_methyl350mM_1.2753.2753.1
Data file D:\Panyue\Proteomic dataset\20100530_methyl350mM_1.RAW

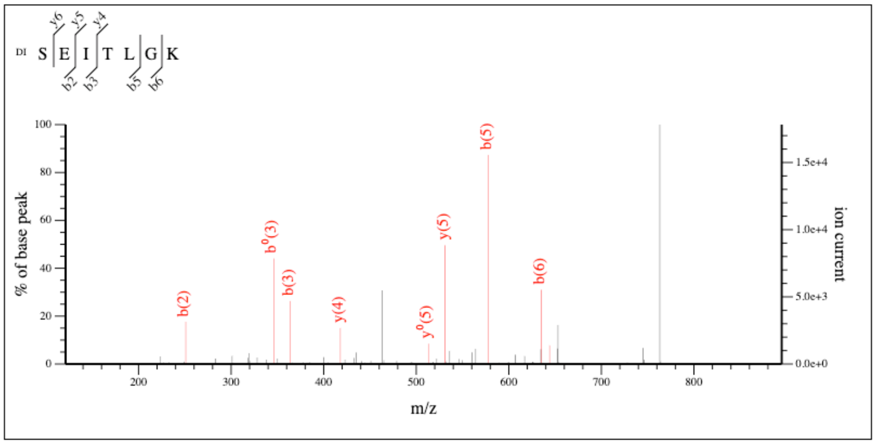

**Monoisotopic mass of neutral peptide Mr(calc):** 780.4864
**Fixed modifications:** Carbamidomethyl (C) (apply to specified residues or termini only)
**Variable modifications:**
**N-term :** Dimethyl:2H(6) (Protein N-term)
**Ions Score:** 27 **Expect:** 0.024
**Matches :** 9/27 fragment ions using 18 most intense peaks (help)

| # | b | b⁰ | Seq. | y | y* | y⁰ | # |
|---|---|---|---|---|---|---|---|
| 1 | 122.1083 | 104.0977 | S | | | | 7 |
| 2 | 251.1509 | 233.1403 | E | 660.3927 | 643.3661 | 642.3821 | 6 |
| 3 | 364.2349 | 346.2244 | I | 531.3501 | 514.3235 | 513.3395 | 5 |
| 4 | 465.2826 | 447.2720 | T | 418.2660 | 401.2395 | 400.2554 | 4 |
| 5 | 578.3667 | 560.3561 | L | 317.2183 | 300.1918 | | 3 |
| 6 | 635.3881 | 617.3776 | G | 204.1343 | 187.1077 | | 2 |
| 7 | | | K | 147.1128 | 130.0863 | | 1 |

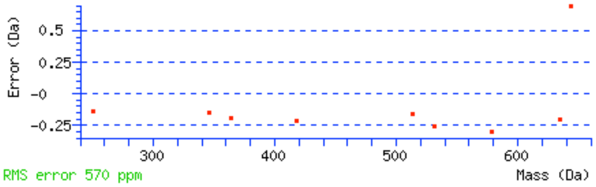


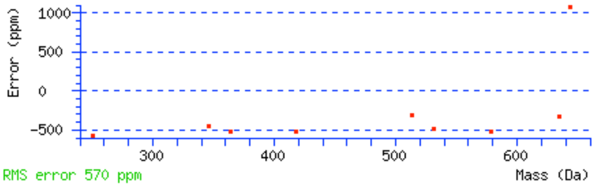


### 49. Pdc5 (2 peptides in 350mM fraction 1)

MS/MS Fragmentation of SEITLGK
Found in P16467 in Uniprot_yeastS288C, Pyruvate decarboxylase isozyme 2 OS=Saccharomyces cerevisiae (strain ATCC 204508 / S288c) OX=559292 GN=PDC5 PE=1 SV=4

Match to Query 2065: 780.448596 from(781.455872,1+) intensity(1431024.3750) rtinseconds(989.4157) index(2475)
Title: 20100530_methyl350mM_1.2753.2753.1
Data file D:\Panyue\Proteomic dataset\20100530_methyl350mM_1.RAW

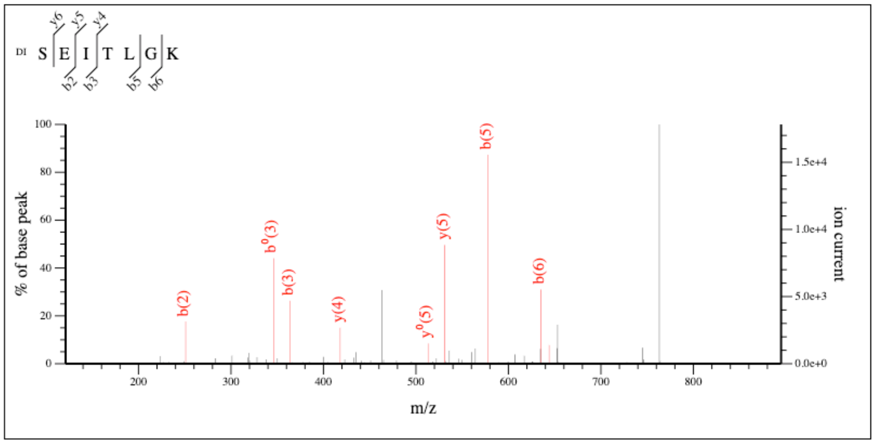

**Monoisotopic mass of neutral peptide Mr(calc):** 780.4864
**Fixed modifications:** Carbamidomethyl (C) (apply to specified residues or termini only)
**Variable modifications:**
**N-term :** Dimethyl:2H(6) (Protein N-term)
**Ions Score:** 27 **Expect:** 0.024
**Matches :** 9/27 fragment ions using 18 most intense peaks (help)

| # | b | $b^0$ | Seq. | y | y* | $y^0$ | # |
|---|---|---|---|---|---|---|---|
| 1 | 122.1083 | 104.0977 | S | | | | 7 |
| 2 | 251.1509 | 233.1403 | E | 660.3927 | 643.3661 | 642.3821 | 6 |
| 3 | 364.2349 | 346.2244 | I | 531.3501 | 514.3235 | 513.3395 | 5 |
| 4 | 465.2826 | 447.2720 | T | 418.2660 | 401.2395 | 400.2554 | 4 |
| 5 | 578.3667 | 560.3561 | L | 317.2183 | 300.1918 | | 3 |
| 6 | 635.3881 | 617.3776 | G | 204.1343 | 187.1077 | | 2 |
| 7 | | | K | 147.1128 | 130.0863 | | 1 |

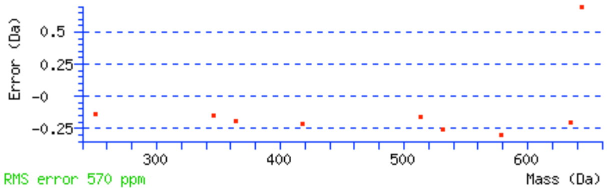


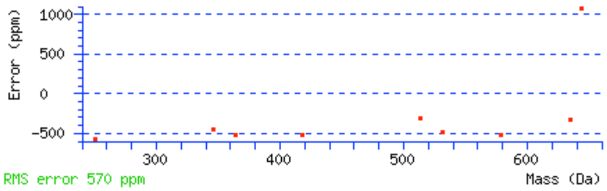


MS/MS Fragmentation of **SEITLGK**
Found in **P16467** in **Uniprot_yeastS288C**, Pyruvate decarboxylase isozyme 2 OS=Saccharomyces cerevisiae (strain ATCC 204508 / S288c) OX=559292 GN=PDC5 PE=1 SV=4

Match to Query 2066: 780.448718 from(781.455994,1+) intensity(675320.9375) rtinseconds(986.8442) index(2468)
Title: 20100530_methyl350mM_1.2745.2745.1
Data file D:\Panyue\Proteomic dataset\20100530_methyl350mM_1.RAW

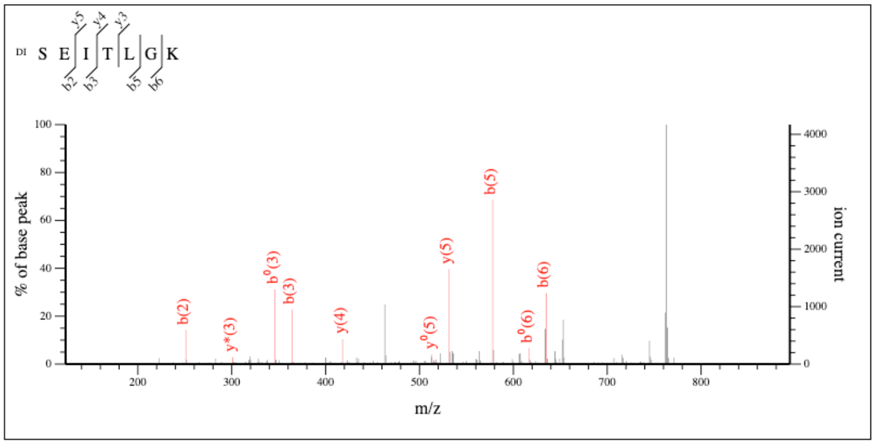

**Monoisotopic mass of neutral peptide Mr(calc):** 780.4864
**Fixed modifications:** Carbamidomethyl (C) (apply to specified residues or termini only)
**Variable modifications:**
**N-term :** Dimethyl:2H(6) (Protein N-term)
**Ions Score:** 26 **Expect:** 0.029
**Matches :** 10/27 fragment ions using 18 most intense peaks (help)

| # | b | $b^0$ | Seq. | y | y* | $y^0$ | # |
|---|---|---|---|---|---|---|---|
| 1 | 122.1083 | 104.0977 | S | | | | 7 |
| 2 | ***251.1509*** | 233.1403 | E | 660.3927 | 643.3661 | 642.3821 | 6 |
| 3 | ***364.2349*** | 346.2244 | I | **531.3501** | 514.3235 | 513.3395 | 5 |
| 4 | 465.2826 | 447.2720 | T | **418.2660** | 401.2395 | 400.2554 | 4 |
| 5 | ***578.3667*** | 560.3561 | L | 317.2183 | 300.1918 | | 3 |
| 6 | ***635.3881*** | 617.3776 | G | 204.1343 | 187.1077 | | 2 |
| 7 | | | K | 147.1128 | 130.0863 | | 1 |

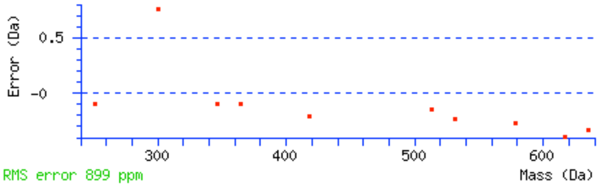


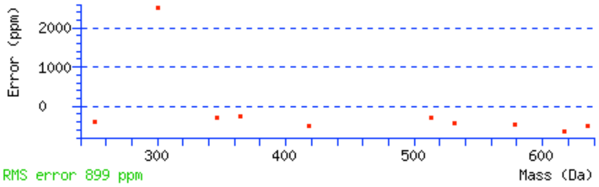


### 50. Pdc1 (2peptides in 350mM fraction1)

MS/MS Fragmentation of **SEITLGK**
Found in **P06169** in **Uniprot_yeastS288C**, Pyruvate decarboxylase isozyme 1 OS=Saccharomyces cerevisiae (strain ATCC 204508 / S288c) OX=559292 GN=PDC1 PE=1 SV=7

Match to Query 2065: 780.448596 from(781.455872,1+) intensity(1431024.3750) rtinseconds(989.4157) index(2475)
Title: 20100530_methyl350mM_1.2753.2753.1
Data file D:\Panyue\Proteomic dataset\20100530_methyl350mM_1.RAW

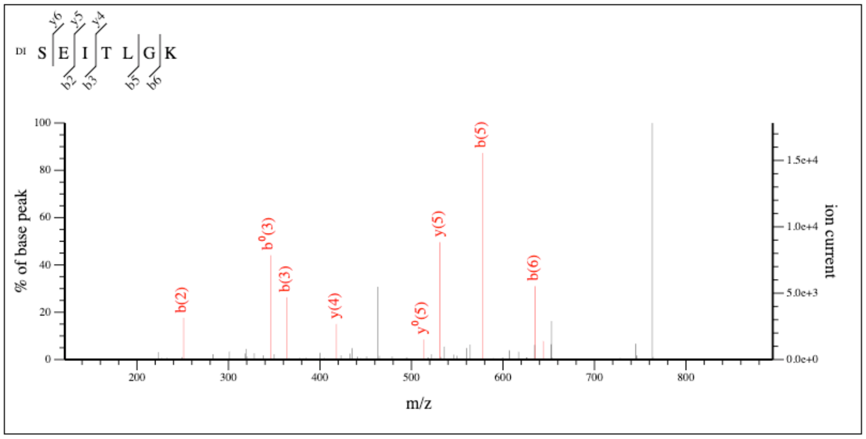

**Monoisotopic mass of neutral peptide Mr(calc):** 780.4864
**Fixed modifications:** Carbamidomethyl (C) (apply to specified residues or termini only)
**Variable modifications:**
**N-term :** Dimethyl:2H(6) (Protein N-term)
**Ions Score:** 27 **Expect:** 0.024
**Matches :** 9/27 fragment ions using 18 most intense peaks (help)

| # | b | $b^0$ | Seq. | y | y* | $y^0$ | # |
|---|---|---|---|---|---|---|---|
| 1 | 122.1083 | 104.0977 | S | | | | 7 |
| 2 | 251.1509 | 233.1403 | E | 660.3927 | 643.3661 | 642.3821 | 6 |
| 3 | 364.2349 | 346.2244 | I | 531.3501 | 514.3235 | 513.3395 | 5 |
| 4 | 465.2826 | 447.2720 | T | 418.2660 | 401.2395 | 400.2554 | 4 |
| 5 | 578.3667 | 560.3561 | L | 317.2183 | 300.1918 | | 3 |
| 6 | 635.3881 | 617.3776 | G | 204.1343 | 187.1077 | | 2 |
| 7 | | | K | 147.1128 | 130.0863 | | 1 |

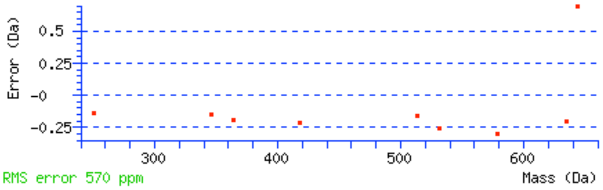


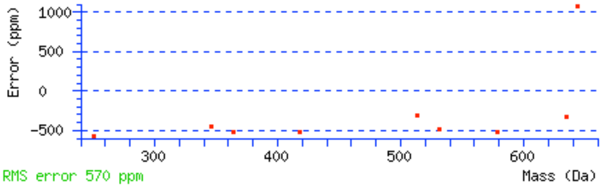


MS/MS Fragmentation of **SEITLGK**
Found in **P06169** in **Uniprot_yeastS288C**, Pyruvate decarboxylase isozyme 1 OS=Saccharomyces cerevisiae (strain ATCC 204508 / S288c) OX=559292 GN=PDC1 PE=1 SV=7

Match to Query 2066: 780.448718 from(781.455994,1+) intensity(675320.9375) rtinseconds(986.8442) index(2468)
Title: 20100530_methyl350mM_1.2745.2745.1
Data file D:\Panyue\Proteomic dataset\20100530_methyl350mM_1.RAW

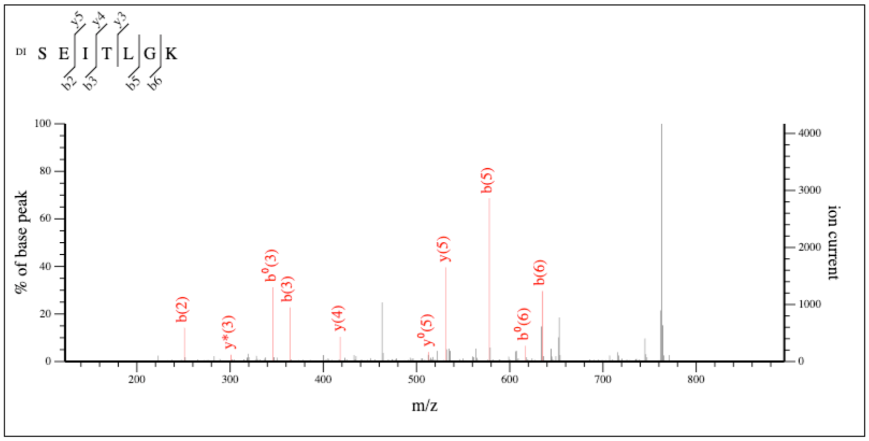

**Monoisotopic mass of neutral peptide Mr(calc):** 780.4864
**Fixed modifications:** Carbamidomethyl (C) (apply to specified residues or termini only)
**Variable modifications:**
**N-term :** Dimethyl:2H(6) (Protein N-term)
**Ions Score:** 26 **Expect:** 0.029
**Matches :** 10/27 fragment ions using 18 most intense peaks (help)

| # | b | $b^0$ | Seq. | y | y* | $y^0$ | # |
|---|---|---|---|---|---|---|---|
| 1 | 122.1083 | 104.0977 | S | | | | 7 |
| 2 | ***251.1509*** | 233.1403 | E | 660.3927 | 643.3661 | 642.3821 | 6 |
| 3 | ***364.2349*** | 346.2244 | I | **531.3501** | 514.3235 | 513.3395 | 5 |
| 4 | 465.2826 | 447.2720 | T | **418.2660** | 401.2395 | 400.2554 | 4 |
| 5 | ***578.3667*** | 560.3561 | L | 317.2183 | 300.1918 | | 3 |
| 6 | ***635.3881*** | 617.3776 | G | 204.1343 | 187.1077 | | 2 |
| 7 | | | K | 147.1128 | 130.0863 | | 1 |

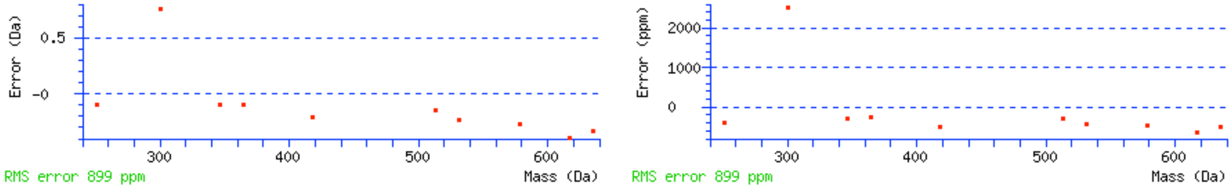


### 51. Atg41 (1 peptide in 350mM fraction1)

MS/MS Fragmentation of **SSVESSPISR**
Found in **Q12048** in **Uniprot_yeastS288C**, Autophagy-related protein 41 OS=Saccharomyces cerevisiae (strain ATCC 204508 / S288c) OX=559292 GN=ATG41 PE=1 SV=1

Match to Query 5164: 1081.561376 from(541.787964,2+) intensity(221117.6719) rtinseconds(1682.8938) index(4355)
Title: 20100530_methyl350mM_1.4844.4844.2
Data file D:\Panyue\Proteomic dataset\20100530_methyl350mM_1.RAW

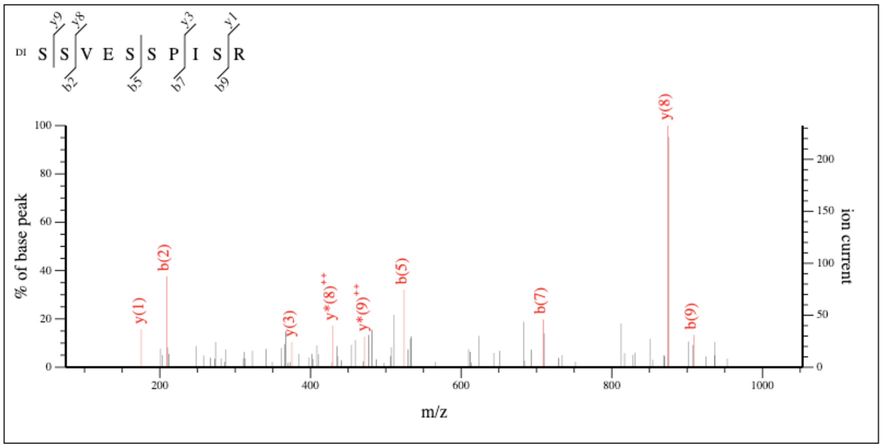

**Monoisotopic mass of neutral peptide Mr(calc):** 1081.5886
**Fixed modifications:** Carbamidomethyl (C) (apply to specified residues or termini only)
**Variable modifications:**
**N-term :** Dimethyl:2H(6) (Protein N-term)
**Ions Score:** 23 **Expect:** 0.019
**Matches :** 12/88 fragment ions using 16 most intense peaks (help)

| # | b | b++ | b0 | b0++ | Seq. | y | y++ | y* | y*++ | y0 | y0++ | # |
|---|---|---|---|---|---|---|---|---|---|---|---|---|
| 1 | 122.1083 | 61.5578 | 104.0977 | 52.5525 | S | | | | | | | 10 |
| 2 | 209.1403 | 105.0738 | 191.1297 | 96.0685 | S | 961.4949 | 481.2511 | 944.4684 | 472.7378 | 943.4843 | 472.2458 | 9 |
| 3 | 308.2087 | 154.6080 | 290.1981 | 145.6027 | V | 874.4629 | 437.7351 | 857.4363 | 429.2218 | 856.4523 | 428.7298 | 8 |
| 4 | 437.2513 | 219.1293 | 419.2407 | 210.1240 | E | 775.3945 | 388.2009 | 758.3679 | 379.6876 | 757.3839 | 379.1956 | 7 |
| 5 | 524.2833 | 262.6453 | 506.2728 | 253.6400 | S | 646.3519 | 323.6796 | 629.3253 | 315.1663 | 628.3413 | 314.6743 | 6 |
| 6 | 611.3154 | 306.1613 | 593.3048 | 297.1560 | S | 559.3198 | 280.1636 | 542.2933 | 271.6503 | 541.3093 | 271.1583 | 5 |
| 7 | 708.3681 | 354.6877 | 690.3576 | 345.6824 | P | 472.2878 | 236.6475 | 455.2613 | 228.1343 | 454.2772 | 227.6423 | 4 |
| 8 | 821.4522 | 411.2297 | 803.4416 | 402.2244 | I | 375.2350 | 188.1212 | 358.2085 | 179.6079 | 357.2245 | 179.1159 | 3 |
| 9 | 908.4842 | 454.7457 | 890.4736 | 445.7405 | S | 262.1510 | 131.5791 | 245.1244 | 123.0659 | 244.1404 | 122.5738 | 2 |
| 10 | | | | | R | 175.1190 | 88.0631 | 158.0924 | 79.5498 | | | 1 |

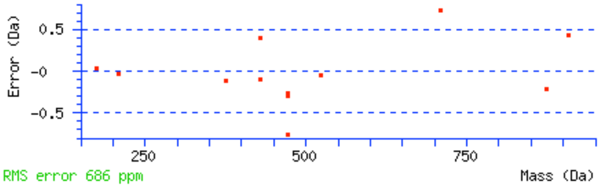


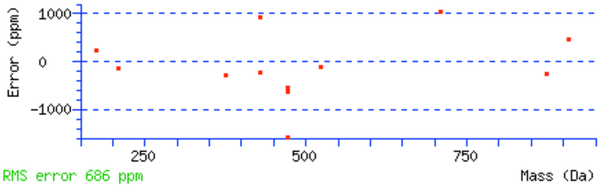


### 52. Eft1 (4 peptides in 350mM fraction1)

MS/MS Fragmentation of **VAFTVDQMRSLMDK**
Found in **P32324** in **Uniprot_yeastS288C**, Elongation factor 2 OS=Saccharomyces cerevisiae (strain ATCC 204508 / S288c) OX=559292 GN=EFT1 PE=1 SV=1

Match to Query 9305: 1706.869360 from(854.441956,2+) intensity(801496.0000) rtinseconds(2605.9476) index(6852)
Title: 20100530_methyl350mM_1.7622.7622.2
Data file D:\Panyue\Proteomic dataset\20100530_methyl350mM_1.RAW

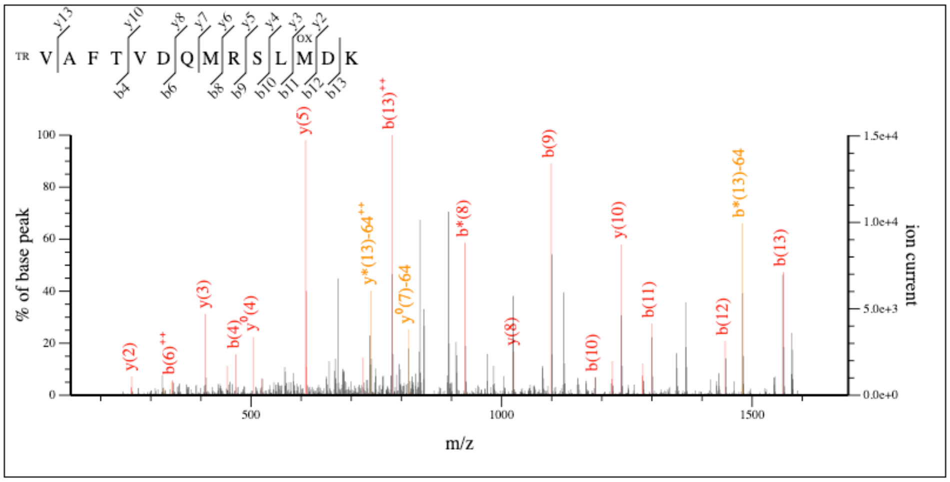

```
Monoisotopic mass of neutral peptide Mr(calc): 1706.9046
Fixed modifications: Carbamidomethyl (C) (apply to specified residues or termini only)
Variable modifications:
N-term : Trimethyl:2H(9) (Protein N-term)
M12    : Oxidation (M), with neutral losses 0.0000(shown in table), 63.9983
Ions Score: 25  Expect: 0.057
Matches : 34/214 fragment ions using 56 most intense peaks   (help)
```

| # | b | $b^{++}$ | $b^{*}$ | $b^{*++}$ | $b^{0}$ | $b^{0++}$ | Seq. | y | $y^{++}$ | $y^{*}$ | $y^{*++}$ | $y^{0}$ | $y^{0++}$ | # |
|---|---|---|---|---|---|---|---|---|---|---|---|---|---|---|
| 1 | 151.1791 | 76.0932 | | | | | V | | | | | | | 14 |
| 2 | 222.2162 | 111.6118 | | | | | A | 1557.7400 | 779.3736 | 1540.7134 | 770.8604 | 1539.7294 | 770.3683 | 13 |
| 3 | 369.2847 | 185.1460 | | | | | F | 1486.7029 | 743.8551 | 1469.6763 | 735.3418 | 1468.6923 | 734.8498 | 12 |
| 4 | ***470.3323*** | 235.6698 | | | 452.3218 | 226.6645 | T | 1339.6344 | 670.3209 | 1322.6079 | 661.8076 | 1321.6239 | 661.3156 | 11 |
| 5 | 569.4008 | 285.2040 | | | 551.3902 | 276.1987 | V | ***1238.5868*** | 619.7970 | 1221.5602 | 611.2837 | 1220.5762 | 610.7917 | 10 |
| 6 | 684.4277 | **342.7175** | | | 666.4171 | 333.7122 | D | 1139.5184 | 570.2628 | 1122.4918 | 561.7495 | 1121.5078 | 561.2575 | 9 |
| 7 | 812.4863 | 406.7468 | 795.4597 | 398.2335 | 794.4757 | 397.7415 | Q | ***1024.4914*** | 512.7493 | 1007.4649 | 504.2361 | 1006.4808 | 503.7441 | 8 |
| 8 | 943.5268 | 472.2670 | 926.5002 | 463.7537 | 925.5162 | 463.2617 | M | 896.4328 | 448.7201 | 879.4063 | 440.2068 | 878.4223 | 439.7148 | 7 |
| 9 | ***1099.6279*** | 550.3176 | 1082.6013 | 541.8043 | 1081.6173 | 541.3123 | R | 765.3924 | 383.1998 | 748.3658 | 374.6865 | 747.3818 | 374.1945 | 6 |
| 10 | ***1186.6599*** | 593.8336 | 1169.6333 | 585.3203 | 1168.6493 | 584.8283 | S | ***609.2912*** | 305.1493 | 592.2647 | 296.6360 | 591.2807 | 296.1440 | 5 |
| 11 | ***1299.7440*** | 650.3756 | 1282.7174 | 641.8623 | 1281.7334 | 641.3703 | L | ***522.2592*** | 261.6332 | 505.2327 | 253.1200 | 504.2486 | 252.6280 | 4 |
| 12 | ***1446.7794*** | **723.8933** | 1429.7528 | 715.3800 | 1428.7688 | 714.8880 | M | ***409.1751*** | 205.0912 | 392.1486 | 196.5779 | 391.1646 | 196.0859 | 3 |
| 13 | ***1561.8063*** | **781.4068** | 1544.7798 | 772.8935 | 1543.7957 | 772.4015 | D | ***262.1397*** | 131.5735 | 245.1132 | 123.0602 | 244.1292 | 122.5682 | 2 |
| 14 | | | | | | | K | 147.1128 | 74.0600 | 130.0863 | 65.5468 | | | 1 |

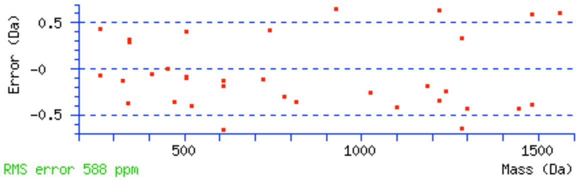


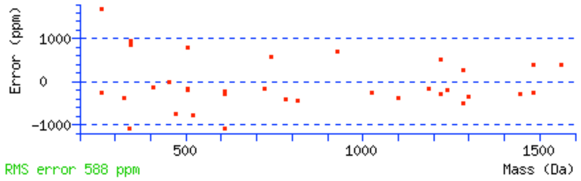


MS/MS Fragmentation of **VAFTVDQMRSLMDK**
Found in **P32324** in **Uniprot_yeastS288C**, Elongation factor 2 OS=Saccharomyces cerevisiae (strain ATCC 204508 / S288c) OX=559292 GN=EFT1 PE=1 SV=1

Match to Query 9306: 1706.869848 from(854.442200,2+) intensity(906835.1875) rtinseconds(2602.1931) index(6841)
Title: 20100530_methyl350mM_1.7610.7610.2
Data file D:\Panyue\Proteomic dataset\20100530_methyl350mM_1.RAW

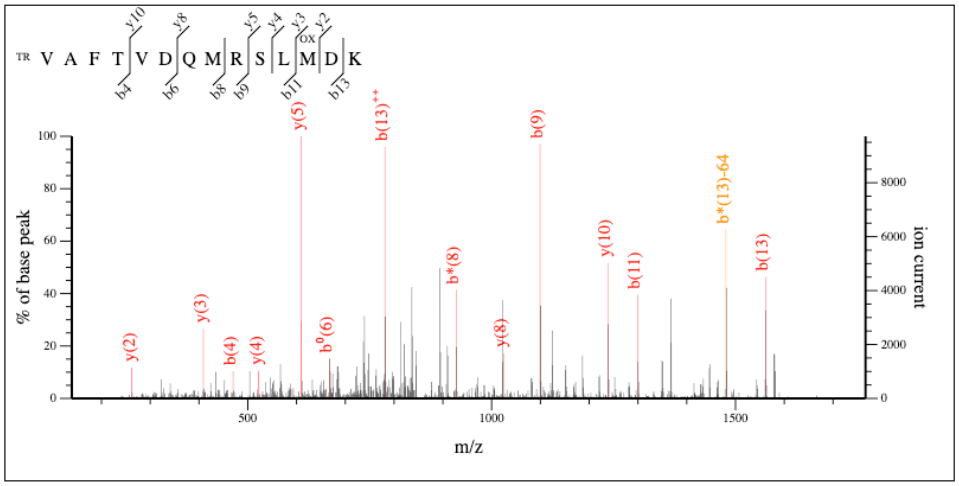

```
Monoisotopic mass of neutral peptide Mr(calc): 1706.9046
Fixed modifications: Carbamidomethyl (C) (apply to specified residues or termini only)
Variable modifications:
N-term : Trimethyl:2H(9) (Protein N-term)
M12    : Oxidation (M), with neutral losses 0.0000(shown in table), 63.9983
Ions Score: 30  Expect: 0.017
Matches : 17/214 fragment ions using 29 most intense peaks   (help)
```

| # | b | b++ | b* | b*++ | b0 | b0++ | Seq. | y | y++ | y* | y*++ | y0 | y0++ | # |
|---|---|---|---|---|---|---|---|---|---|---|---|---|---|---|
| 1 | 151.1791 | 76.0932 | | | | | V | | | | | | | 14 |
| 2 | 222.2162 | 111.6118 | | | | | A | 1557.7400 | 779.3736 | 1540.7134 | 770.8604 | 1539.7294 | 770.3683 | 13 |
| 3 | 369.2847 | 185.1460 | | | | | F | 1486.7029 | 743.8551 | 1469.6763 | 735.3418 | 1468.6923 | 734.8498 | 12 |
| 4 | **470.3323** | 235.6698 | | | 452.3218 | 226.6645 | T | 1339.6344 | 670.3209 | 1322.6079 | 661.8076 | 1321.6239 | 661.3156 | 11 |
| 5 | 569.4008 | 285.2040 | | | 551.3902 | 276.1987 | V | ***1238.5868*** | 619.7970 | 1221.5602 | 611.2837 | 1220.5762 | 610.7917 | 10 |
| 6 | 684.4277 | 342.7175 | | | 666.4171 | 333.7122 | D | 1139.5184 | 570.2628 | 1122.4918 | 561.7495 | 1121.5078 | 561.2575 | 9 |
| 7 | 812.4863 | 406.7468 | 795.4597 | 398.2335 | 794.4757 | 397.7415 | Q | ***1024.4914*** | 512.7493 | 1007.4649 | 504.2361 | 1006.4808 | 503.7441 | 8 |
| 8 | 943.5268 | 472.2670 | 926.5002 | 463.7537 | 925.5162 | 463.2617 | M | 896.4328 | 448.7201 | 879.4063 | 440.2068 | 878.4223 | 439.7148 | 7 |
| 9 | **1099.6279** | 550.3176 | 1082.6013 | 541.8043 | 1081.6173 | 541.3123 | R | 765.3924 | 383.1998 | 748.3658 | 374.6865 | 747.3818 | 374.1945 | 6 |
| 10 | 1186.6599 | 593.8336 | 1169.6333 | 585.3203 | 1168.6493 | 584.8283 | S | ***609.2912*** | 305.1493 | 592.2647 | 296.6360 | 591.2807 | 296.1440 | 5 |
| 11 | **1299.7440** | 650.3756 | 1282.7174 | 641.8623 | 1281.7334 | 641.3703 | L | ***522.2592*** | 261.6332 | 505.2327 | 253.1200 | 504.2486 | 252.6280 | 4 |
| 12 | 1446.7794 | 723.8933 | 1429.7528 | 715.3800 | 1428.7688 | 714.8880 | M | ***409.1751*** | 205.0912 | 392.1486 | 196.5779 | 391.1646 | 196.0859 | 3 |
| 13 | **1561.8063** | 781.4068 | 1544.7798 | 772.8935 | 1543.7957 | 772.4015 | D | ***262.1397*** | 131.5735 | 245.1132 | 123.0602 | 244.1292 | 122.5682 | 2 |
| 14 | | | | | | | K | 147.1128 | 74.0600 | 130.0863 | 65.5468 | | | 1 |

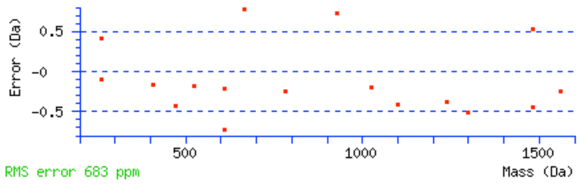


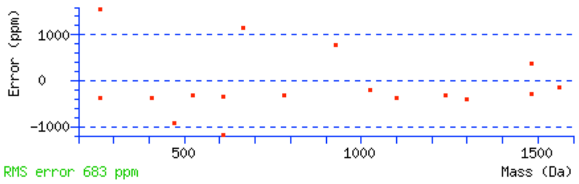


MS/MS Fragmentation of **VAFTVDQMRSLMDK**
Found in **P32324** in **Uniprot_yeastS288C**, Elongation factor 2 OS=Saccharomyces cerevisiae (strain ATCC 204508 / S288c) OX=559292 GN=EFT1 PE=1 SV=1

Match to Query 9307: 1706.872044 from(854.443298,2+) intensity(755776.0625) rtinseconds(2600.1285) index(6836)
Title: 20100530_methyl350mM_1.7604.7604.2
Data file D:\Panyue\Proteomic dataset\20100530_methyl350mM_1.RAW

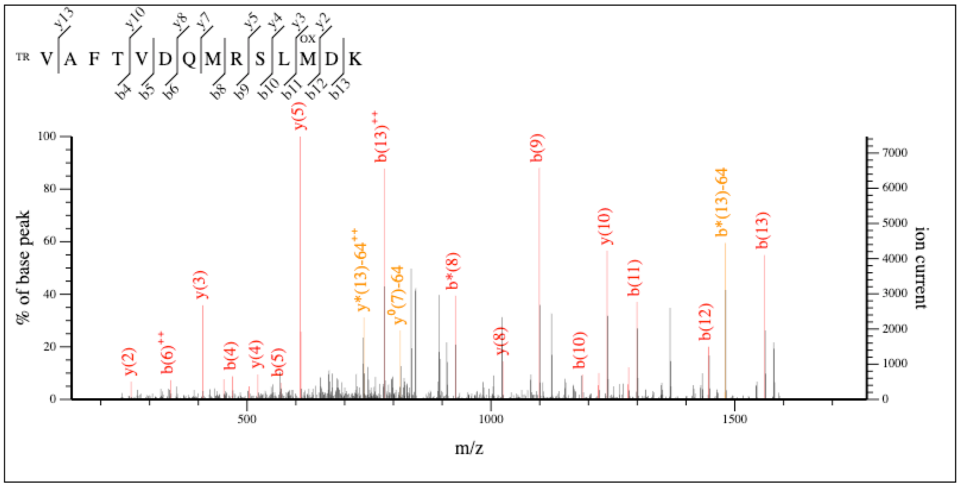

**Monoisotopic mass of neutral peptide Mr(calc):** 1706.9046
**Fixed modifications:** Carbamidomethyl (C) (apply to specified residues or termini only)
**Variable modifications:**
**N-term :** Trimethyl:2H(9) (Protein N-term)
**M12** : Oxidation (M), with neutral losses 0.0000(shown in table), 63.9983
**Ions Score:** 24 **Expect:** 0.058
**Matches :** 32/214 fragment ions using 57 most intense peaks (help)

| # | b | $b^{++}$ | b* | $b^{*++}$ | $b^0$ | $b^{0++}$ | Seq. | y | $y^{++}$ | y* | $y^{*++}$ | $y^0$ | $y^{0++}$ | # |
|---|---|---|---|---|---|---|---|---|---|---|---|---|---|---|
| 1 | 151.1791 | 76.0932 | | | | | V | | | | | | | 14 |
| 2 | 222.2162 | 111.6118 | | | | | A | 1557.7400 | 779.3736 | 1540.7134 | 770.8604 | 1539.7294 | 770.3683 | 13 |
| 3 | 369.2847 | 185.1460 | | | | | F | 1486.7029 | 743.8551 | 1469.6763 | 735.3418 | 1468.6923 | 734.8498 | 12 |
| 4 | ***470.3323*** | 235.6698 | | | 452.3218 | 226.6645 | T | 1339.6344 | 670.3209 | 1322.6079 | 661.8076 | 1321.6239 | 661.3156 | 11 |
| 5 | ***569.4008*** | 285.2040 | | | 551.3902 | 276.1987 | V | ***1238.5868*** | 619.7970 | 1221.5602 | 611.2837 | 1220.5762 | 610.7917 | 10 |
| 6 | 684.4277 | 342.7175 | | | 666.4171 | 333.7122 | D | 1139.5184 | 570.2628 | 1122.4918 | 561.7495 | 1121.5078 | 561.2575 | 9 |
| 7 | 812.4863 | 406.7468 | 795.4597 | 398.2335 | 794.4757 | 397.7415 | Q | ***1024.4914*** | 512.7493 | 1007.4649 | 504.2361 | 1006.4808 | 503.7441 | 8 |
| 8 | 943.5268 | 472.2670 | 926.5002 | 463.7537 | 925.5162 | 463.2617 | M | 896.4328 | 448.7201 | 879.4063 | 440.2068 | 878.4223 | 439.7148 | 7 |
| 9 | ***1099.6279*** | 550.3176 | 1082.6013 | 541.8043 | 1081.6173 | 541.3123 | R | 765.3924 | 383.1998 | 748.3658 | 374.6865 | 747.3818 | 374.1945 | 6 |
| 10 | ***1186.6599*** | 593.8336 | 1169.6333 | 585.3203 | 1168.6493 | 584.8283 | S | ***609.2912*** | 305.1493 | 592.2647 | 296.6360 | 591.2807 | 296.1440 | 5 |
| 11 | ***1299.7440*** | 650.3756 | 1282.7174 | 641.8623 | 1281.7334 | 641.3703 | L | ***522.2592*** | 261.6332 | 505.2327 | 253.1200 | 504.2486 | 252.6280 | 4 |
| 12 | ***1446.7794*** | 723.8933 | 1429.7528 | 715.3800 | 1428.7688 | 714.8880 | M | ***409.1751*** | 205.0912 | 392.1486 | 196.5779 | 391.1646 | 196.0859 | 3 |
| 13 | ***1561.8063*** | 781.4068 | 1544.7798 | 772.8935 | 1543.7957 | 772.4015 | D | ***262.1397*** | 131.5735 | 245.1132 | 123.0602 | 244.1292 | 122.5682 | 2 |
| 14 | | | | | | | K | 147.1128 | 74.0600 | 130.0863 | 65.5468 | | | 1 |

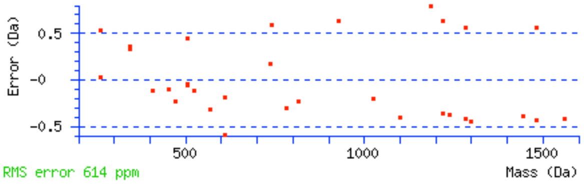


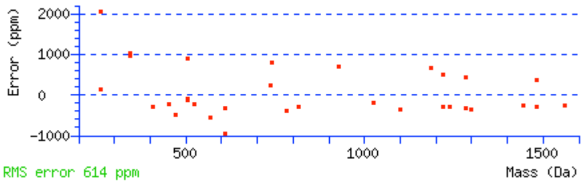


MS/MS Fragmentation of **VAFTVDQMRSLMDK**
Found in **P32324** in **Uniprot_yeastS288C**, Elongation factor 2 OS=Saccharomyces cerevisiae (strain ATCC 204508 / S288c) OX=559292 GN=EFT1 PE=1 SV=1

Match to Query 9308: 1706.872656 from(854.443604,2+) intensity(801496.0000) rtinseconds(2606.783) index(6855)
Title: 20100530_methyl350mM_1.7625.7625.2
Data file D:\Panyue\Proteomic dataset\20100530_methyl350mM_1.RAW

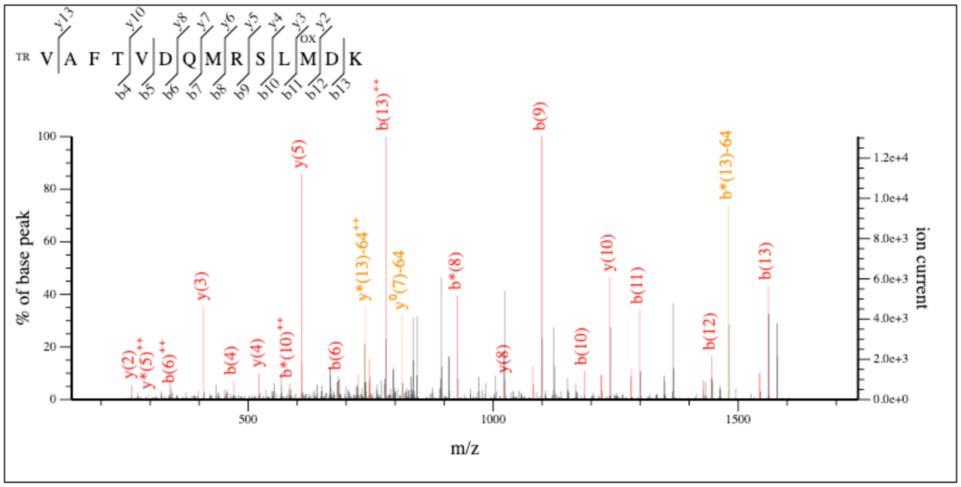

```
Monoisotopic mass of neutral peptide Mr(calc): 1706.9046
Fixed modifications: Carbamidomethyl (C) (apply to specified residues or termini only)
Variable modifications:
N-term : Trimethyl:2H(9) (Protein N-term)
M12    : Oxidation (M), with neutral losses 0.0000(shown in table), 63.9983
Ions Score: 27  Expect: 0.032
Matches : 47/214 fragment ions using 85 most intense peaks   (help)
```

| # | b | b++ | b* | b*++ | b0 | b0++ | Seq. | y | y++ | y* | y*++ | y0 | y0++ | # |
|---|---|---|---|---|---|---|---|---|---|---|---|---|---|---|
| 1 | 151.1791 | 76.0932 | | | | | V | | | | | | | 14 |
| 2 | 222.2162 | 111.6118 | | | | | A | 1557.7400 | 779.3736 | 1540.7134 | 770.8604 | 1539.7294 | 770.3683 | 13 |
| 3 | 369.2847 | 185.1460 | | | | | F | 1486.7029 | 743.8551 | 1469.6763 | 735.3418 | 1468.6923 | 734.8498 | 12 |
| 4 | ***470.3323*** | 235.6698 | | | 452.3218 | 226.6645 | T | 1339.6344 | 670.3209 | 1322.6079 | 661.8076 | 1321.6239 | 661.3156 | 11 |
| 5 | ***569.4008*** | 285.2040 | | | 551.3902 | 276.1987 | V | **1238.5868** | 619.7970 | 1221.5602 | 611.2837 | 1220.5762 | 610.7917 | 10 |
| 6 | ***684.4277*** | 342.7175 | | | 666.4171 | 333.7122 | D | 1139.5184 | 570.2628 | 1122.4918 | 561.7495 | 1121.5078 | 561.2575 | 9 |
| 7 | 812.4863 | 406.7468 | 795.4597 | 398.2335 | 794.4757 | 397.7415 | Q | **1024.4914** | 512.7493 | 1007.4649 | 504.2361 | 1006.4808 | 503.7441 | 8 |
| 8 | 943.5268 | 472.2670 | 926.5002 | 463.7537 | 925.5162 | 463.2617 | M | 896.4328 | 448.7201 | 879.4063 | 440.2068 | 878.4223 | 439.7148 | 7 |
| 9 | ***1099.6279*** | 550.3176 | 1082.6013 | 541.8043 | 1081.6173 | 541.3123 | R | 765.3924 | 383.1998 | 748.3658 | 374.6865 | 747.3818 | 374.1945 | 6 |
| 10 | ***1186.6599*** | 593.8336 | 1169.6333 | 585.3203 | 1168.6493 | 584.8283 | S | **609.2912** | 305.1493 | 592.2647 | 296.6360 | 591.2807 | 296.1440 | 5 |
| 11 | ***1299.7440*** | 650.3756 | 1282.7174 | 641.8623 | 1281.7334 | 641.3703 | L | **522.2592** | 261.6332 | 505.2327 | 253.1200 | 504.2486 | 252.6280 | 4 |
| 12 | ***1446.7794*** | 723.8933 | 1429.7528 | 715.3800 | 1428.7688 | 714.8880 | M | **409.1751** | 205.0912 | 392.1486 | 196.5779 | 391.1646 | 196.0859 | 3 |
| 13 | ***1561.8063*** | 781.4068 | 1544.7798 | 772.8935 | 1543.7957 | 772.4015 | D | **262.1397** | 131.5735 | 245.1132 | 123.0602 | 244.1292 | 122.5682 | 2 |
| 14 | | | | | | | K | 147.1128 | 74.0600 | 130.0863 | 65.5468 | | | 1 |

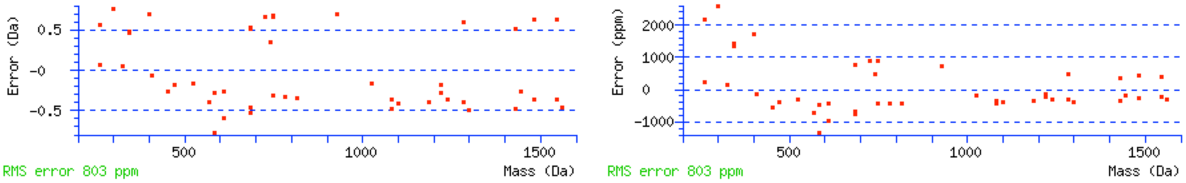


### 53. Pdc6 (2 peptides in 350mM fraction 2)

MS/MS Fragmentation of **SEITLGK**
Found in **P26263** in **Uniprot_yeastS288C**, Pyruvate decarboxylase isozyme 3 OS=Saccharomyces cerevisiae (strain ATCC 204508 / S288c) OX=559292 GN=PDC6 PE=1 SV=3

Match to Query 2158: 780.448412 from(781.455688,1+) intensity(339328.8750) rtinseconds(1035.2799) index(2578)
Title: 20100530_methyl350mM_2.2868.2868.1
Data file D:\Panyue\Proteomic dataset\20100530_methyl350mM_2.RAW

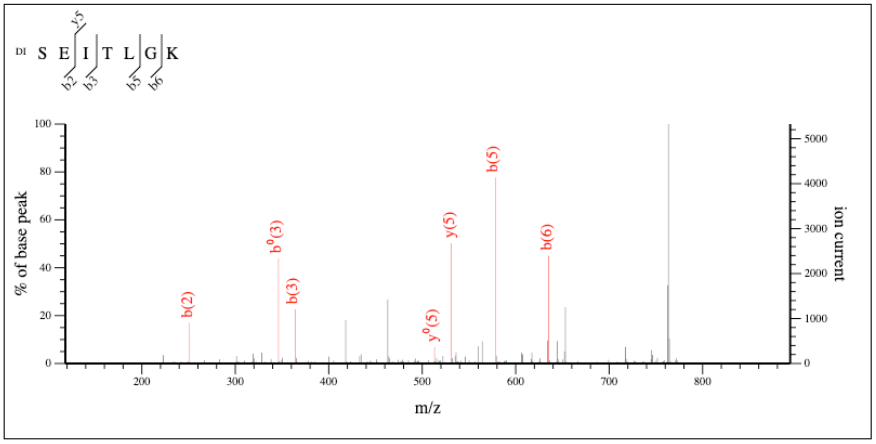

**Monoisotopic mass of neutral peptide Mr(calc):** 780.4864
**Fixed modifications:** Carbamidomethyl (C) (apply to specified residues or termini only)
**Variable modifications:**
**N-term :** Dimethyl:2H(6) (Protein N-term)
**Ions Score:** 26 **Expect:** 0.029
**Matches :** 7/27 fragment ions using 12 most intense peaks (help)

| # | b | $b^0$ | Seq. | y | y* | $y^0$ | # |
|---|---|---|---|---|---|---|---|
| 1 | 122.1083 | 104.0977 | S | | | | 7 |
| 2 | ***251.1509*** | 233.1403 | E | 660.3927 | 643.3661 | 642.3821 | 6 |
| 3 | ***364.2349*** | 346.2244 | I | 531.3501 | 514.3235 | 513.3395 | 5 |
| 4 | 465.2826 | 447.2720 | T | 418.2660 | 401.2395 | 400.2554 | 4 |
| 5 | ***578.3667*** | 560.3561 | L | 317.2183 | 300.1918 | | 3 |
| 6 | ***635.3881*** | 617.3776 | G | 204.1343 | 187.1077 | | 2 |
| 7 | | | K | 147.1128 | 130.0863 | | 1 |

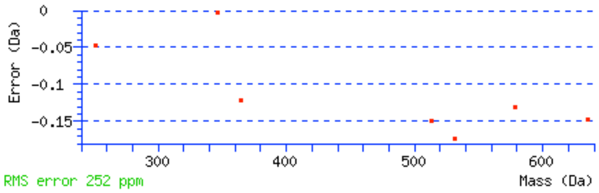


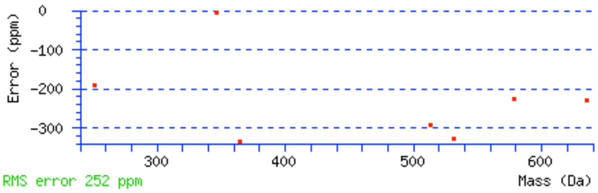


MS/MS Fragmentation of **SEITLGK**
Found in **P26263** in **Uniprot_yeastS288C**, Pyruvate decarboxylase isozyme 3 OS=Saccharomyces cerevisiae (strain ATCC 204508 / S288c) OX=559292 GN=PDC6 PE=1 SV=3

Match to Query 2159: 780.449145 from(781.456421,1+) intensity(967923.8125) rtinseconds(1038.388) index(2586)
Title: 20100530_methyl350mM_2.2877.2877.1
Data file D:\Panyue\Proteomic dataset\20100530_methyl350mM_2.RAW

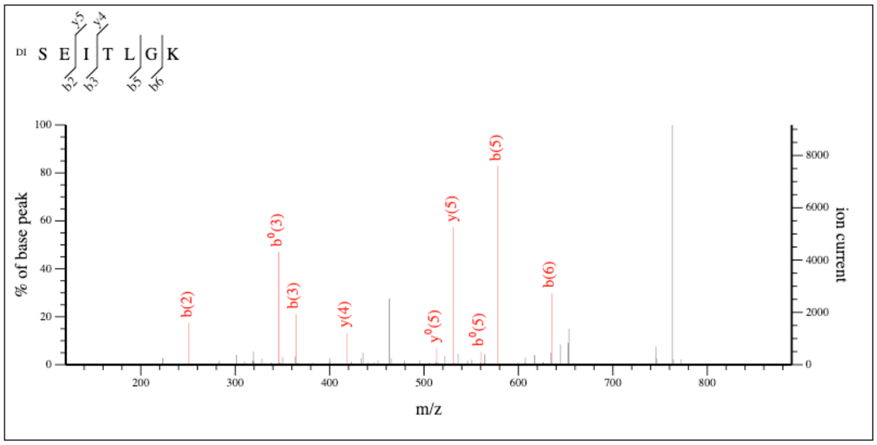

**Monoisotopic mass of neutral peptide Mr(calc):** 780.4864
**Fixed modifications:** Carbamidomethyl (C) (apply to specified residues or termini only)
**Variable modifications:**
**N-term :** Dimethyl:2H(6) (Protein N-term)
**Ions Score:** 27 **Expect:** 0.025
**Matches :** 9/27 fragment ions using 18 most intense peaks (help)

| # | b | $b^0$ | Seq. | y | y* | $y^0$ | # |
|---|---|---|---|---|---|---|---|
| 1 | 122.1083 | 104.0977 | S | | | | 7 |
| 2 | ***251.1509*** | 233.1403 | E | 660.3927 | 643.3661 | 642.3821 | 6 |
| 3 | ***364.2349*** | 346.2244 | I | **531.3501** | 514.3235 | 513.3395 | 5 |
| 4 | 465.2826 | 447.2720 | T | **418.2660** | 401.2395 | 400.2554 | 4 |
| 5 | ***578.3667*** | 560.3561 | L | 317.2183 | 300.1918 | | 3 |
| 6 | ***635.3881*** | 617.3776 | G | 204.1343 | 187.1077 | | 2 |
| 7 | | | K | 147.1128 | 130.0863 | | 1 |

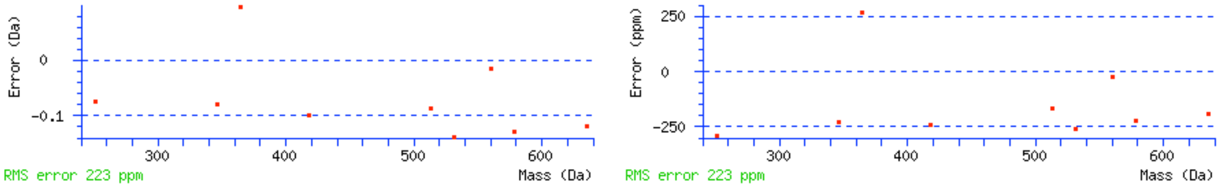


### 54. Pdc5 (2 peptides in 350mM fraction 2)

MS/MS Fragmentation of **SEITLGK**
Found in **P16467** in **Uniprot_yeastS288C**, Pyruvate decarboxylase isozyme 2 OS=Saccharomyces cerevisiae (strain ATCC 204508 / S288c) OX=559292 GN=PDC5 PE=1 SV=4

Match to Query 2158: 780.448412 from(781.455688,1+) intensity(339328.8750) rtinseconds(1035.2799) index(2578)
Title: 20100530_methyl350mM_2.2868.2868.1
Data file D:\Panyue\Proteomic dataset\20100530_methyl350mM_2.RAW

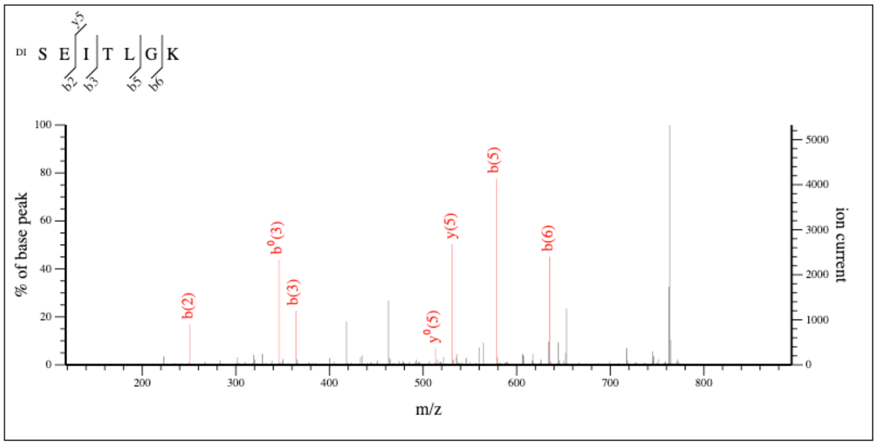

**Monoisotopic mass of neutral peptide Mr(calc):** 780.4864
**Fixed modifications:** Carbamidomethyl (C) (apply to specified residues or termini only)
**Variable modifications:**
**N-term :** Dimethyl:2H(6) (Protein N-term)
**Ions Score:** 26 **Expect:** 0.029
**Matches :** 7/27 fragment ions using 12 most intense peaks (help)

| # | b | b⁰ | Seq. | y | y* | y⁰ | # |
|---|---|---|---|---|---|---|---|
| 1 | 122.1083 | 104.0977 | S | | | | 7 |
| 2 | ***251.1509*** | 233.1403 | E | 660.3927 | 643.3661 | 642.3821 | 6 |
| 3 | ***364.2349*** | 346.2244 | I | 531.3501 | 514.3235 | 513.3395 | 5 |
| 4 | 465.2826 | 447.2720 | T | 418.2660 | 401.2395 | 400.2554 | 4 |
| 5 | ***578.3667*** | 560.3561 | L | 317.2183 | 300.1918 | | 3 |
| 6 | ***635.3881*** | 617.3776 | G | 204.1343 | 187.1077 | | 2 |
| 7 | | | K | 147.1128 | 130.0863 | | 1 |

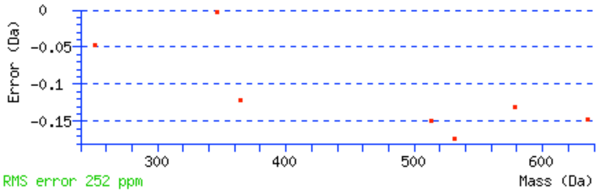


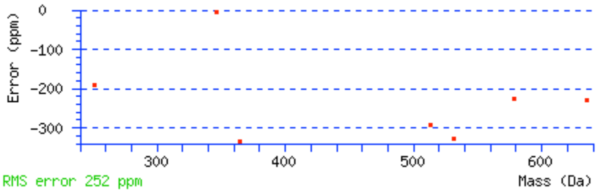


MS/MS Fragmentation of **SEITLGK**
Found in **P16467** in **Uniprot_yeastS288C**, Pyruvate decarboxylase isozyme 2 OS=Saccharomyces cerevisiae (strain ATCC 204508 / S288c) OX=559292 GN=PDC5 PE=1 SV=4

Match to Query 2159: 780.449145 from(781.456421,1+) intensity(967923.8125) rtinseconds(1038.388) index(2586)
Title: 20100530_methyl350mM_2.2877.2877.1
Data file D:\Panyue\Proteomic dataset\20100530_methyl350mM_2.RAW

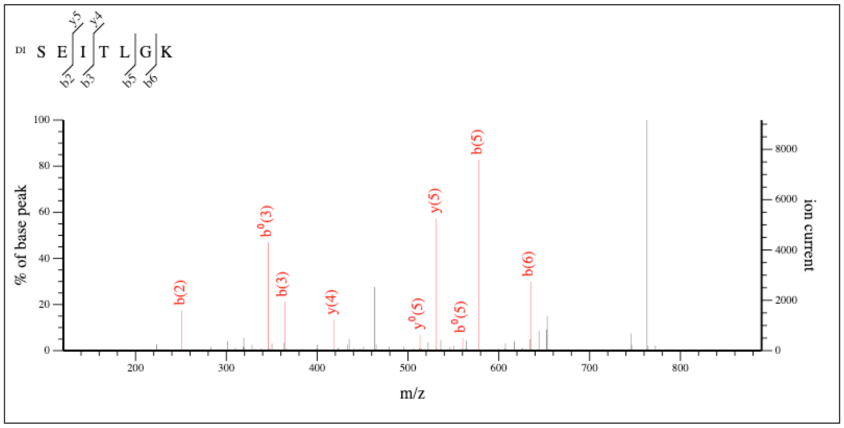

**Monoisotopic mass of neutral peptide Mr(calc):** 780.4864
**Fixed modifications:** Carbamidomethyl (C) (apply to specified residues or termini only)
**Variable modifications:**
**N-term :** Dimethyl:2H(6) (Protein N-term)
**Ions Score:** 27 **Expect:** 0.025
**Matches :** 9/27 fragment ions using 18 most intense peaks (help)

| # | b | $b^0$ | Seq. | y | $y^*$ | $y^0$ | # |
|---|---|---|---|---|---|---|---|
| 1 | 122.1083 | 104.0977 | S | | | | 7 |
| 2 | ***251.1509*** | 233.1403 | E | 660.3927 | 643.3661 | 642.3821 | 6 |
| 3 | ***364.2349*** | 346.2244 | I | **531.3501** | 514.3235 | 513.3395 | 5 |
| 4 | 465.2826 | 447.2720 | T | **418.2660** | 401.2395 | 400.2554 | 4 |
| 5 | ***578.3667*** | 560.3561 | L | 317.2183 | 300.1918 | | 3 |
| 6 | ***635.3881*** | 617.3776 | G | 204.1343 | 187.1077 | | 2 |
| 7 | | | K | 147.1128 | 130.0863 | | 1 |

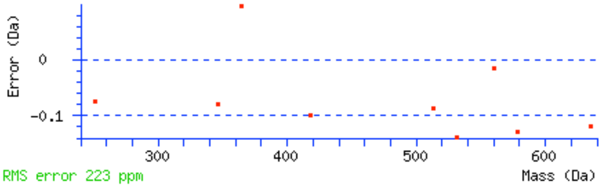


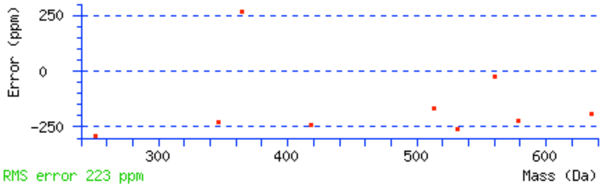


### 55. Pdc1 (2 peptides in 350mM fraction 2)

MS/MS Fragmentation of **SEITLGK**
Found in **P06169** in **Uniprot_yeastS288C**, Pyruvate decarboxylase isozyme 1 OS=Saccharomyces cerevisiae (strain ATCC 204508 / S288c) OX=559292 GN=PDC1 PE=1 SV=7

Match to Query 2158: 780.448412 from(781.455688,1+) intensity(339328.8750) rtinseconds(1035.2799) index(2578)
Title: 20100530_methyl350mM_2.2868.2868.1
Data file D:\Panyue\Proteomic dataset\20100530_methyl350mM_2.RAW

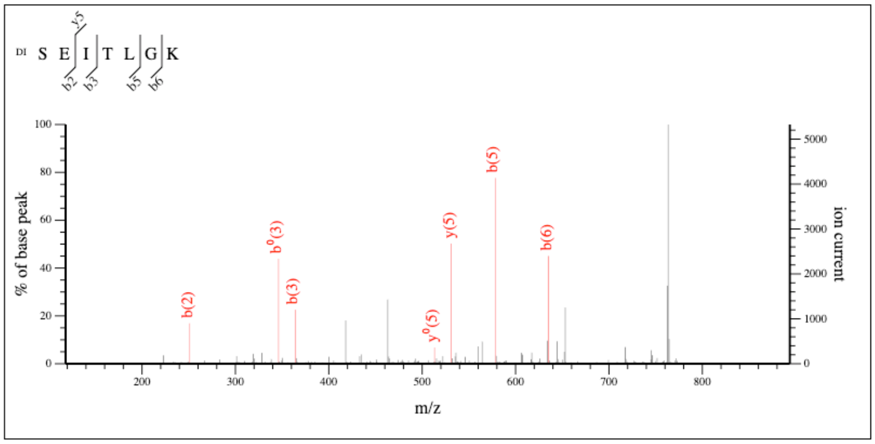

**Monoisotopic mass of neutral peptide Mr(calc):** 780.4864
**Fixed modifications:** Carbamidomethyl (C) (apply to specified residues or termini only)
**Variable modifications:**
**N-term :** Dimethyl:2H(6) (Protein N-term)
**Ions Score:** 26 **Expect:** 0.029
**Matches :** 7/27 fragment ions using 12 most intense peaks (help)

| # | b | $b^0$ | Seq. | y | y* | $y^0$ | # |
|---|---|---|---|---|---|---|---|
| 1 | 122.1083 | 104.0977 | S | | | | 7 |
| 2 | *251.1509* | 233.1403 | E | 660.3927 | 643.3661 | 642.3821 | 6 |
| 3 | *364.2349* | 346.2244 | I | 531.3501 | 514.3235 | 513.3395 | 5 |
| 4 | 465.2826 | 447.2720 | T | 418.2660 | 401.2395 | 400.2554 | 4 |
| 5 | *578.3667* | 560.3561 | L | 317.2183 | 300.1918 | | 3 |
| 6 | *635.3881* | 617.3776 | G | 204.1343 | 187.1077 | | 2 |
| 7 | | | K | 147.1128 | 130.0863 | | 1 |

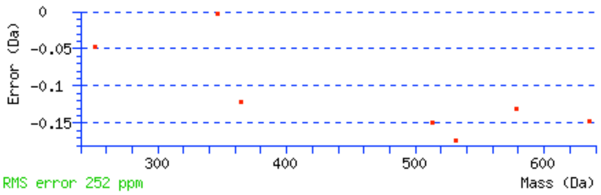


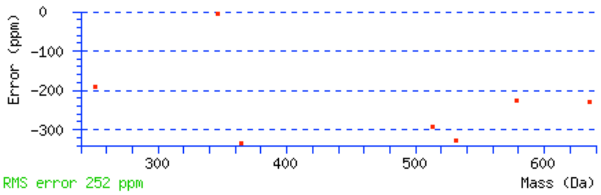


MS/MS Fragmentation of **SEITLGK**
Found in **P06169** in **Uniprot_yeastS288C**, Pyruvate decarboxylase isozyme 1 OS=Saccharomyces cerevisiae (strain ATCC 204508 / S288c) OX=559292 GN=PDC1 PE=1 SV=7

Match to Query 2159: 780.449145 from(781.456421,1+) intensity(967923.8125) rtinseconds(1038.388) index(2586)
Title: 20100530_methyl350mM_2.2877.2877.1
Data file D:\Panyue\Proteomic dataset\20100530_methyl350mM_2.RAW

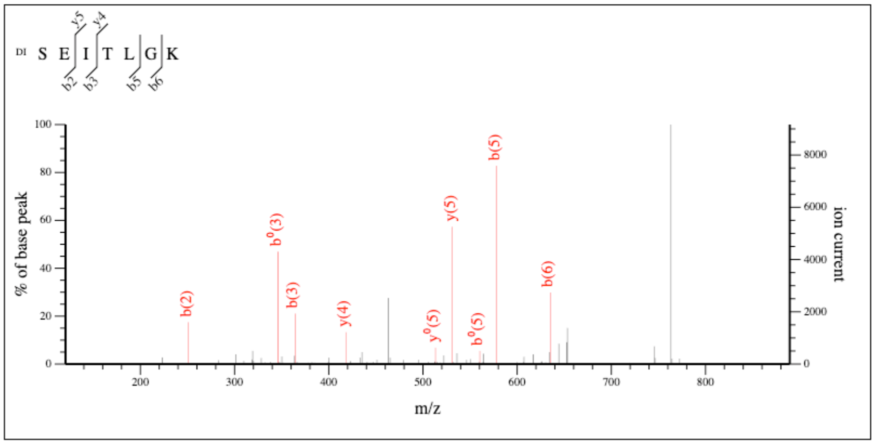

**Monoisotopic mass of neutral peptide Mr(calc):** 780.4864
**Fixed modifications:** Carbamidomethyl (C) (apply to specified residues or termini only)
**Variable modifications:**
**N-term :** Dimethyl:2H(6) (Protein N-term)
**Ions Score:** 27 **Expect:** 0.025
**Matches :** 9/27 fragment ions using 18 most intense peaks (help)

| # | b | b$^0$ | Seq. | y | y* | y$^0$ | # |
|---|---|---|---|---|---|---|---|
| 1 | 122.1083 | 104.0977 | S | | | | 7 |
| 2 | 251.1509 | 233.1403 | E | 660.3927 | 643.3661 | 642.3821 | 6 |
| 3 | 364.2349 | 346.2244 | I | 531.3501 | 514.3235 | 513.3395 | 5 |
| 4 | 465.2826 | 447.2720 | T | 418.2660 | 401.2395 | 400.2554 | 4 |
| 5 | 578.3667 | 560.3561 | L | 317.2183 | 300.1918 | | 3 |
| 6 | 635.3881 | 617.3776 | G | 204.1343 | 187.1077 | | 2 |
| 7 | | | K | 147.1128 | 130.0863 | | 1 |

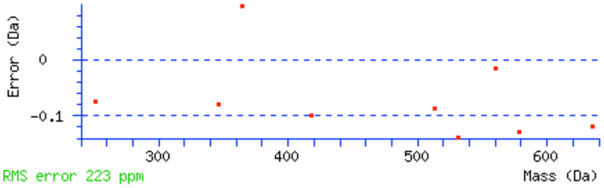


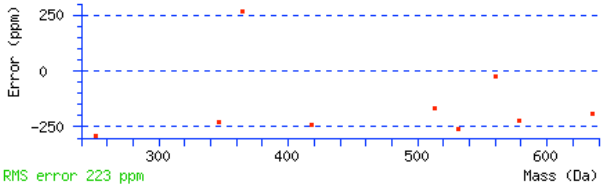


### 56. Ssc1 (1 peptide in 350mM fraction 2)

MS/MS Fragmentation of LAAKNILNR
Found in P0CS90 in Uniprot_yeastS288C, Heat shock protein SSC1, mitochondrial OS=Saccharomyces cerevisiae (strain ATCC 204508 / S288c) OX=559292 GN=SSC1 PE=1 SV=1

Match to Query 4722: 1028.602270 from(515.308411,2+) intensity(711733.1250) rtinseconds(1247.4135) index(3137)
Title: 20100530_methyl350mM_2.3491.3491.2
Data file D:\Panyue\Proteomic dataset\20100530_methyl350mM_2.RAW

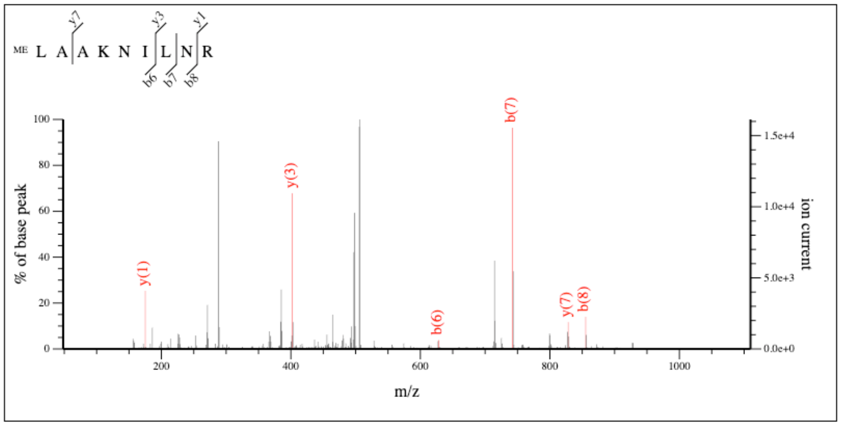

**Monoisotopic mass of neutral peptide Mr(calc):** 1028.6534
**Fixed modifications:** Carbamidomethyl (C) (apply to specified residues or termini only)
**Variable modifications:**
**N-term :** Methyl:2H(3) (Protein N-term)
**Ions Score:** 20 **Expect:** 0.15
**Matches :** 6/58 fragment ions using 9 most intense peaks (help)

| # | b | b++ | b* | b*++ | Seq. | y | y++ | y* | y*++ | # |
|---|---|---|---|---|---|---|---|---|---|---|
| 1 | 131.1258 | 66.0665 | | | L | | | | | 9 |
| 2 | 202.1629 | 101.5851 | | | A | 899.5421 | 450.2747 | 882.5156 | 441.7614 | 8 |
| 3 | 273.2000 | 137.1037 | | | A | ***828.5050*** | 414.7561 | 811.4785 | 406.2429 | 7 |
| 4 | 401.2950 | 201.1511 | 384.2685 | 192.6379 | K | 757.4679 | 379.2376 | 740.4413 | 370.7243 | 6 |
| 5 | 515.3379 | 258.1726 | 498.3114 | 249.6593 | N | 629.3729 | 315.1901 | 612.3464 | 306.6768 | 5 |
| 6 | ***628.4220*** | 314.7146 | 611.3955 | 306.2014 | I | 515.3300 | 258.1686 | 498.3035 | 249.6554 | 4 |
| 7 | ***741.5061*** | 371.2567 | 724.4795 | 362.7434 | L | ***402.2459*** | 201.6266 | 385.2194 | 193.1133 | 3 |
| 8 | ***855.5490*** | 428.2781 | 838.5224 | 419.7649 | N | 289.1619 | 145.0846 | 272.1353 | 136.5713 | 2 |
| 9 | | | | | R | ***175.1190*** | 88.0631 | 158.0924 | 79.5498 | 1 |

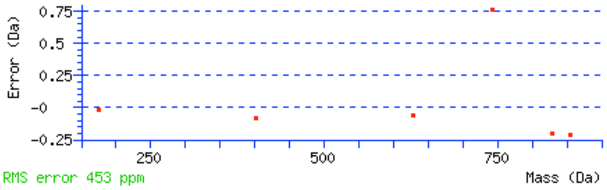


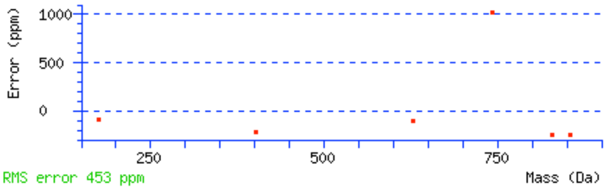


### 57. Nop14 (1 peptide in 350mM fraction 2)

MS/MS Fragmentation of **AGSQLKNLK**
Found in **Q99207** in **Uniprot_yeastS288C**, Nucleolar complex protein 14 OS=Saccharomyces cerevisiae (strain ATCC 204508 / S288c) OX=559292 GN=NOP14 PE=1 SV=1

Match to Query 4529: 1008.670519 from(1009.677795,1+) intensity(272300.1562) rtinseconds(1700.9615) index(4366)
Title: 20100530_methyl350mM_2.4858.4858.1
Data file D:\Panyue\Proteomic dataset\20100530_methyl350mM_2.RAW

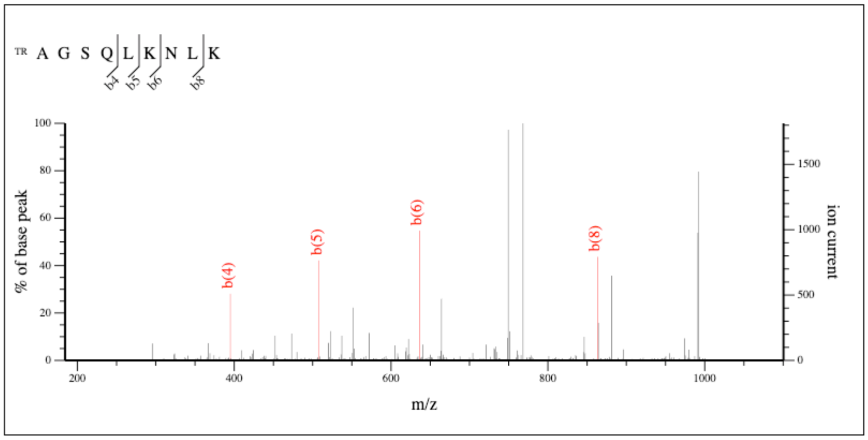

**Monoisotopic mass of neutral peptide Mr(calc):** 1008.6641
**Fixed modifications:** Carbamidomethyl (C) (apply to specified residues or termini only)
**Variable modifications:**
**N-term :** Trimethyl:2H(9) (Protein N-term)
**Ions Score:** 27 **Expect:** 0.0037
**Matches :** 4/37 fragment ions using 8 most intense peaks (help)

| # | b | b* | b⁰ | Seq. | y | y* | y⁰ | # |
|---|---|---|---|---|---|---|---|---|
| 1 | 123.1478 | | | A | | | | 9 |
| 2 | 180.1693 | | | G | 887.5309 | 870.5043 | 869.5203 | 8 |
| 3 | 267.2013 | | 249.1908 | S | 830.5094 | 813.4829 | 812.4989 | 7 |
| 4 | ***395.2599*** | 378.2334 | 377.2493 | Q | 743.4774 | 726.4509 | | 6 |
| 5 | ***508.3440*** | 491.3174 | 490.3334 | L | 615.4188 | 598.3923 | | 5 |
| 6 | ***636.4389*** | 619.4124 | 618.4284 | K | 502.3348 | 485.3082 | | 4 |
| 7 | 750.4819 | 733.4553 | 732.4713 | N | 374.2398 | 357.2132 | | 3 |
| 8 | ***863.5659*** | 846.5394 | 845.5554 | L | 260.1969 | 243.1703 | | 2 |
| 9 | | | | K | 147.1128 | 130.0863 | | 1 |

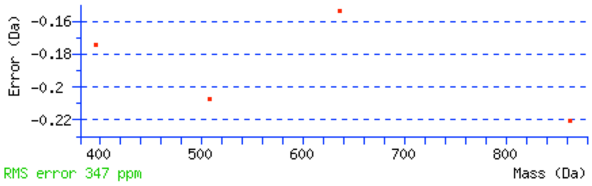


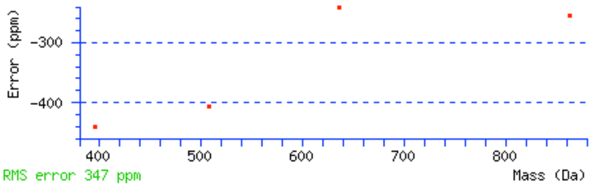


### 58. Eft1 (1 peptide in 350mM fraction 2)

MS/MS Fragmentation of **VAFTVDQMRSLMDK**
Found in **P32324** in **Uniprot_yeastS288C**, Elongation factor 2 OS=Saccharomyces cerevisiae (strain ATCC 204508 / S288c) OX=559292 GN=EFT1 PE=1 SV=1

Match to Query 9524: 1706.871312 from(854.442932,2+) intensity(1077469.7500) rtinseconds(2679.2239) index(6991)
Title: 20100530_methyl350mM_2.7778.7778.2
Data file D:\Panyue\Proteomic dataset\20100530_methyl350mM_2.RAW

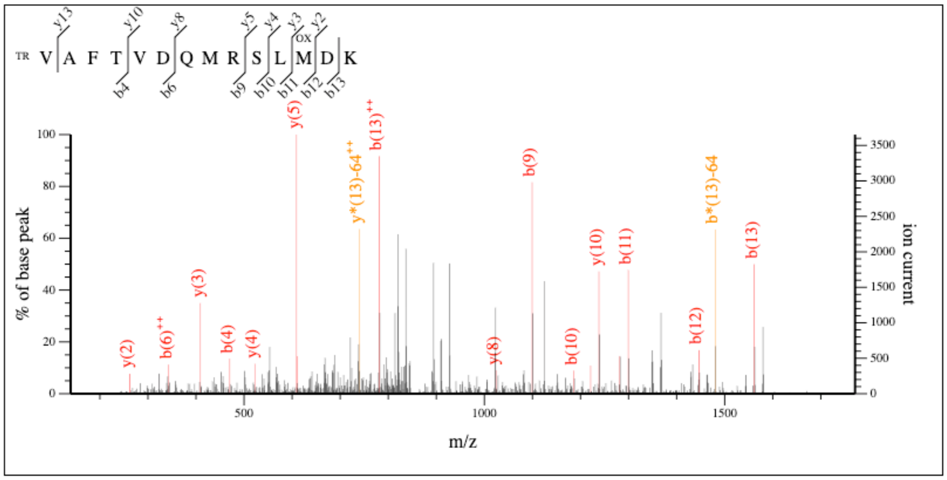

**Monoisotopic mass of neutral peptide Mr(calc):** 1706.9046
**Fixed modifications:** Carbamidomethyl (C) (apply to specified residues or termini only)
**Variable modifications:**
**N-term :** Trimethyl:2H(9) (Protein N-term)
**M12** : Oxidation (M), with neutral losses 0.0000(shown in table), 63.9983
**Ions Score:** 22 **Expect:** 0.11
**Matches :** 24/214 fragment ions using 43 most intense peaks (help)

| # | b | b++ | b* | b*++ | b0 | b0++ | Seq. | y | y++ | y* | y*++ | y0 | y0++ | # |
|---|---|---|---|---|---|---|---|---|---|---|---|---|---|---|
| 1 | 151.1791 | 76.0932 | | | | | V | | | | | | | 14 |
| 2 | 222.2162 | 111.6118 | | | | | A | 1557.7400 | 779.3736 | 1540.7134 | 770.8604 | 1539.7294 | 770.3683 | 13 |
| 3 | 369.2847 | 185.1460 | | | | | F | 1486.7029 | 743.8551 | 1469.6763 | 735.3418 | 1468.6923 | 734.8498 | 12 |
| 4 | 470.3323 | 235.6698 | | | 452.3218 | 226.6645 | T | 1339.6344 | 670.3209 | 1322.6079 | 661.8076 | 1321.6239 | 661.3156 | 11 |
| 5 | 569.4008 | 285.2040 | | | 551.3902 | 276.1987 | V | 1238.5868 | 619.7970 | 1221.5602 | 611.2837 | 1220.5762 | 610.7917 | 10 |
| 6 | 684.4277 | 342.7175 | | | 666.4171 | 333.7122 | D | 1139.5184 | 570.2628 | 1122.4918 | 561.7495 | 1121.5078 | 561.2575 | 9 |
| 7 | 812.4863 | 406.7468 | 795.4597 | 398.2335 | 794.4757 | 397.7415 | Q | 1024.4914 | 512.7493 | 1007.4649 | 504.2361 | 1006.4808 | 503.7441 | 8 |
| 8 | 943.5268 | 472.2670 | 926.5002 | 463.7537 | 925.5162 | 463.2617 | M | 896.4328 | 448.7201 | 879.4063 | 440.2068 | 878.4223 | 439.7148 | 7 |
| 9 | 1099.6279 | 550.3176 | 1082.6013 | 541.8043 | 1081.6173 | 541.3123 | R | 765.3924 | 383.1998 | 748.3658 | 374.6865 | 747.3818 | 374.1945 | 6 |
| 10 | 1186.6599 | 593.8336 | 1169.6333 | 585.3203 | 1168.6493 | 584.8283 | S | 609.2912 | 305.1493 | 592.2647 | 296.6360 | 591.2807 | 296.1440 | 5 |
| 11 | 1299.7440 | 650.3756 | 1282.7174 | 641.8623 | 1281.7334 | 641.3703 | L | 522.2592 | 261.6332 | 505.2327 | 253.1200 | 504.2486 | 252.6280 | 4 |
| 12 | 1446.7794 | 723.8933 | 1429.7528 | 715.3800 | 1428.7688 | 714.8880 | M | 409.1751 | 205.0912 | 392.1486 | 196.5779 | 391.1646 | 196.0859 | 3 |
| 13 | 1561.8063 | 781.4068 | 1544.7798 | 772.8935 | 1543.7957 | 772.4015 | D | 262.1397 | 131.5735 | 245.1132 | 123.0602 | 244.1292 | 122.5682 | 2 |
| 14 | | | | | | | K | 147.1128 | 74.0600 | 130.0863 | 65.5468 | | | 1 |

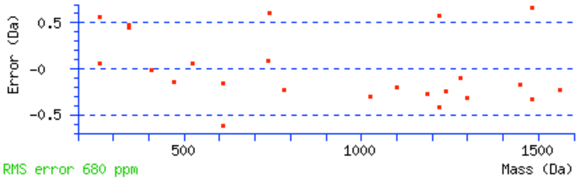


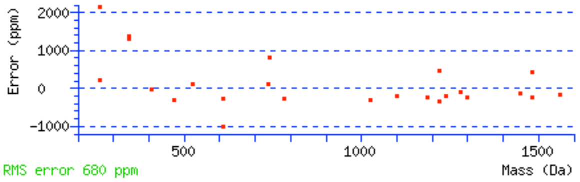


### 59. Pdc6 (2 peptides in 350mM fraction 3)

MS/MS Fragmentation of **SEITLGK**
Found in **P26263** in **Uniprot_yeastS288C**, Pyruvate decarboxylase isozyme 3 OS=Saccharomyces cerevisiae (strain ATCC 204508 / S288c) OX=559292 GN=PDC6 PE=1 SV=3

Match to Query 2082: 780.449145 from(781.456421,1+) intensity(1526053.3750) rtinseconds(1029.7648) index(2567)
Title: 20100530_methyl350mM_3.2856.2856.1
Data file D:\Panyue\Proteomic dataset\20100530_methyl350mM_3.RAW

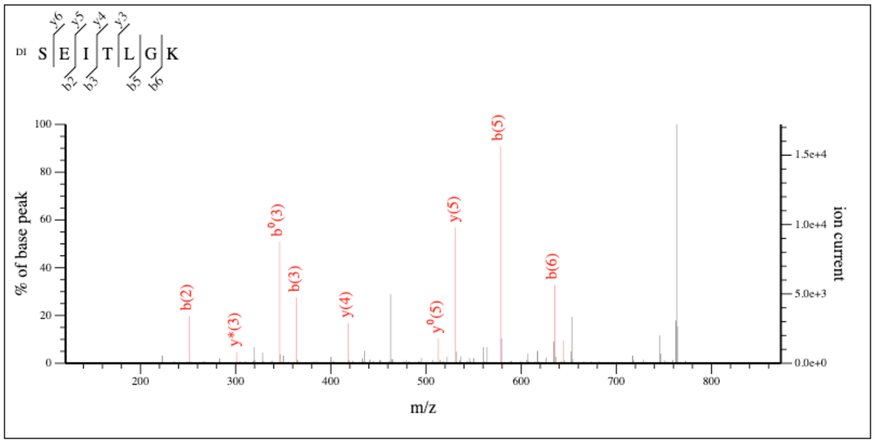

**Monoisotopic mass of neutral peptide Mr(calc):** 780.4864
**Fixed modifications:** Carbamidomethyl (C) (apply to specified residues or termini only)
**Variable modifications:**
**N-term :** Dimethyl:2H(6) (Protein N-term)
**Ions Score:** 27 **Expect:** 0.026
**Matches :** 10/27 fragment ions using 18 most intense peaks (help)

| # | b | $b^0$ | Seq. | y | y* | $y^0$ | # |
|---|---|---|---|---|---|---|---|
| 1 | 122.1083 | 104.0977 | S | | | | 7 |
| 2 | ***251.1509*** | 233.1403 | E | 660.3927 | 643.3661 | 642.3821 | 6 |
| 3 | ***364.2349*** | 346.2244 | I | **531.3501** | 514.3235 | 513.3395 | 5 |
| 4 | 465.2826 | 447.2720 | T | **418.2660** | 401.2395 | 400.2554 | 4 |
| 5 | ***578.3667*** | 560.3561 | L | 317.2183 | 300.1918 | | 3 |
| 6 | ***635.3881*** | 617.3776 | G | 204.1343 | 187.1077 | | 2 |
| 7 | | | K | 147.1128 | 130.0863 | | 1 |

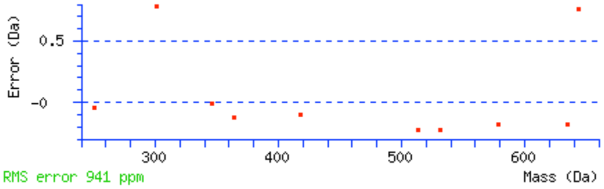


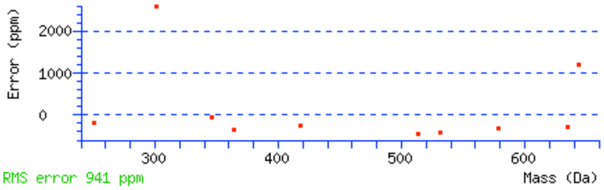


MS/MS Fragmentation of **SEITLGK**
Found in **P26263** in **Uniprot_yeastS288C**, Pyruvate decarboxylase isozyme 3 OS=Saccharomyces cerevisiae (strain ATCC 204508 / S288c) OX=559292 GN=PDC6 PE=1 SV=3

Match to Query 2083: 780.449328 from(781.456604,1+) intensity(615472.8750) rtinseconds(1027.341) index(2561)
Title: 20100530_methyl350mM_3.2849.2849.1
Data file D:\Panyue\Proteomic dataset\20100530_methyl350mM_3.RAW

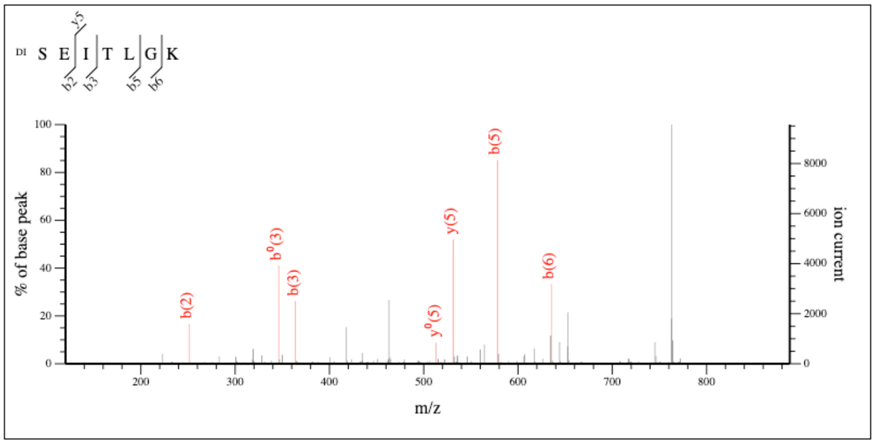

**Monoisotopic mass of neutral peptide Mr(calc):** 780.4864
**Fixed modifications:** Carbamidomethyl (C) (apply to specified residues or termini only)
**Variable modifications:**
**N-term :** Dimethyl:2H(6) (Protein N-term)
**Ions Score:** 24 **Expect:** 0.05
**Matches :** 7/27 fragment ions using 12 most intense peaks (help)

| # | b | b$^0$ | Seq. | y | y* | y$^0$ | # |
|---|---|---|---|---|---|---|---|
| 1 | 122.1083 | 104.0977 | S | | | | 7 |
| 2 | ***251.1509*** | 233.1403 | E | 660.3927 | 643.3661 | 642.3821 | 6 |
| 3 | ***364.2349*** | 346.2244 | I | 531.3501 | 514.3235 | 513.3395 | 5 |
| 4 | 465.2826 | 447.2720 | T | 418.2660 | 401.2395 | 400.2554 | 4 |
| 5 | ***578.3667*** | 560.3561 | L | 317.2183 | 300.1918 | | 3 |
| 6 | ***635.3881*** | 617.3776 | G | 204.1343 | 187.1077 | | 2 |
| 7 | | | K | 147.1128 | 130.0863 | | 1 |

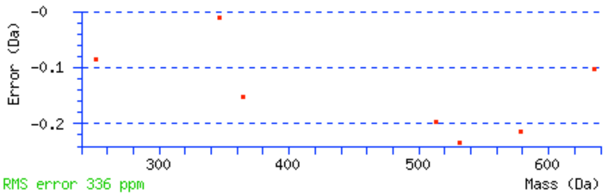


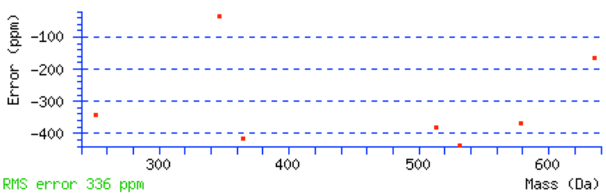


### 60. Pdc5 (2 peptides in 350mM fraction 3)

MS/MS Fragmentation of **SEITLGK**
Found in **P16467** in **Uniprot_yeastS288C**, Pyruvate decarboxylase isozyme 2 OS=Saccharomyces cerevisiae (strain ATCC 204508 / S288c) OX=559292 GN=PDC5 PE=1 SV=4

Match to Query 2082: 780.449145 from(781.456421,1+) intensity(1526053.3750) rtinseconds(1029.7648) index(2567)
Title: 20100530_methyl350mM_3.2856.2856.1
Data file D:\Panyue\Proteomic dataset\20100530_methyl350mM_3.RAW

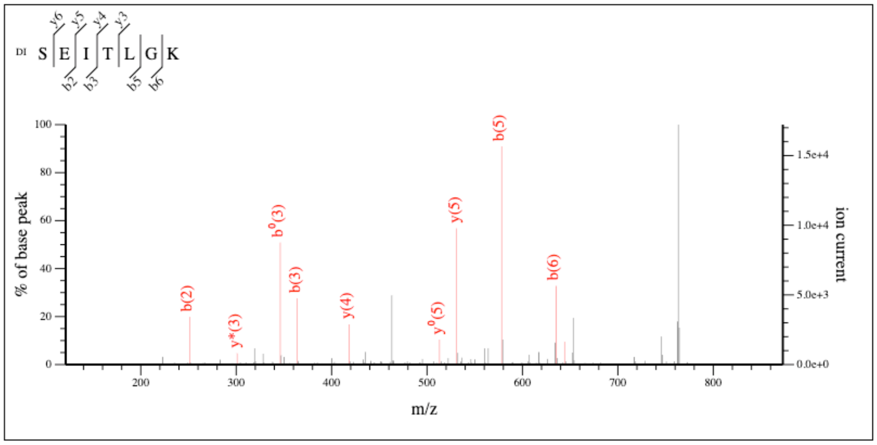

**Monoisotopic mass of neutral peptide Mr(calc):** 780.4864
**Fixed modifications:** Carbamidomethyl (C) (apply to specified residues or termini only)
**Variable modifications:**
**N-term :** Dimethyl:2H(6) (Protein N-term)
**Ions Score:** 27 **Expect:** 0.026
**Matches :** 10/27 fragment ions using 18 most intense peaks (help)

| # | b | $b^0$ | Seq. | y | y* | $y^0$ | # |
|---|---|---|---|---|---|---|---|
| 1 | 122.1083 | 104.0977 | S | | | | 7 |
| 2 | 251.1509 | 233.1403 | E | 660.3927 | 643.3661 | 642.3821 | 6 |
| 3 | 364.2349 | 346.2244 | I | 531.3501 | 514.3235 | 513.3395 | 5 |
| 4 | 465.2826 | 447.2720 | T | 418.2660 | 401.2395 | 400.2554 | 4 |
| 5 | 578.3667 | 560.3561 | L | 317.2183 | 300.1918 | | 3 |
| 6 | 635.3881 | 617.3776 | G | 204.1343 | 187.1077 | | 2 |
| 7 | | | K | 147.1128 | 130.0863 | | 1 |

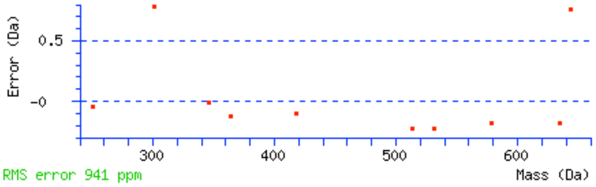


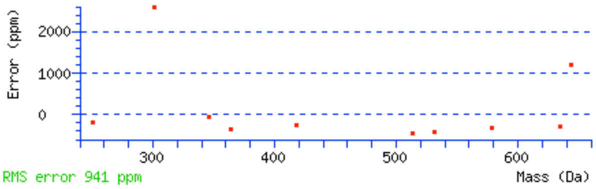


MS/MS Fragmentation of **SEITLGK**
Found in **P16467** in **Uniprot_yeastS288C**, Pyruvate decarboxylase isozyme 2 OS=Saccharomyces cerevisiae (strain ATCC 204508 / S288c) OX=559292 GN=PDC5 PE=1 SV=4

Match to Query 2083: 780.449328 from(781.456604,1+) intensity(615472.8750) rtinseconds(1027.341) index(2561)
Title: 20100530_methyl350mM_3.2849.2849.1
Data file D:\Panyue\Proteomic dataset\20100530_methyl350mM_3.RAW

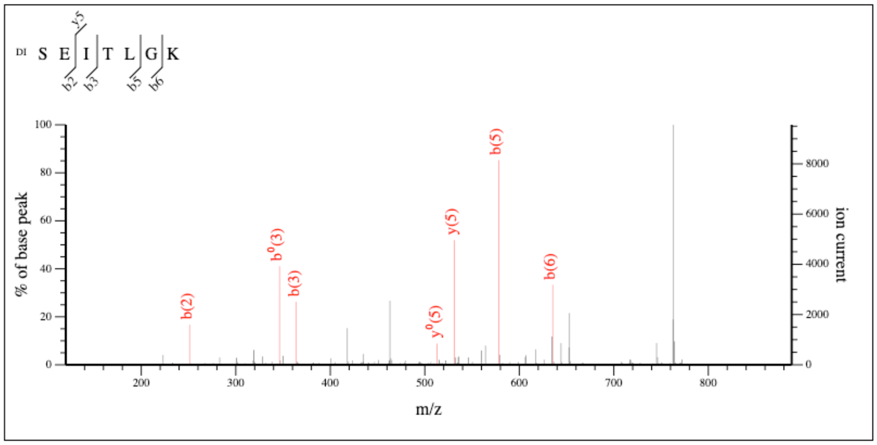

**Monoisotopic mass of neutral peptide Mr(calc):** 780.4864
**Fixed modifications:** Carbamidomethyl (C) (apply to specified residues or termini only)
**Variable modifications:**
**N-term :** Dimethyl:2H(6) (Protein N-term)
**Ions Score:** 24 **Expect:** 0.05
**Matches :** 7/27 fragment ions using 12 most intense peaks (help)

| # | b | b0 | Seq. | y | y* | y0 | # |
|---|---|---|---|---|---|---|---|
| 1 | 122.1083 | 104.0977 | S | | | | 7 |
| 2 | *251.1509* | 233.1403 | E | 660.3927 | 643.3661 | 642.3821 | 6 |
| 3 | *364.2349* | 346.2244 | I | 531.3501 | 514.3235 | 513.3395 | 5 |
| 4 | 465.2826 | 447.2720 | T | 418.2660 | 401.2395 | 400.2554 | 4 |
| 5 | *578.3667* | 560.3561 | L | 317.2183 | 300.1918 | | 3 |
| 6 | *635.3881* | 617.3776 | G | 204.1343 | 187.1077 | | 2 |
| 7 | | | K | 147.1128 | 130.0863 | | 1 |

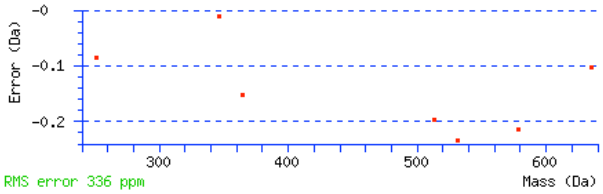


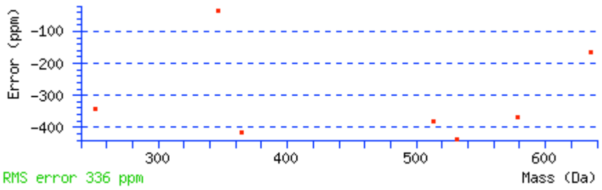


### 61. Pdc1 (2 peptides in 350mM fraction 3)

MS/MS Fragmentation of **SEITLGK**
Found in **P06169** in **Uniprot_yeastS288C**, Pyruvate decarboxylase isozyme 1 OS=Saccharomyces cerevisiae (strain ATCC 204508 / S288c) OX=559292 GN=PDC1 PE=1 SV=7

Match to Query 2082: 780.449145 from(781.456421,1+) intensity(1526053.3750) rtinseconds(1029.7648) index(2567)
Title: 20100530_methyl350mM_3.2856.2856.1
Data file D:\Panyue\Proteomic dataset\20100530_methyl350mM_3.RAW

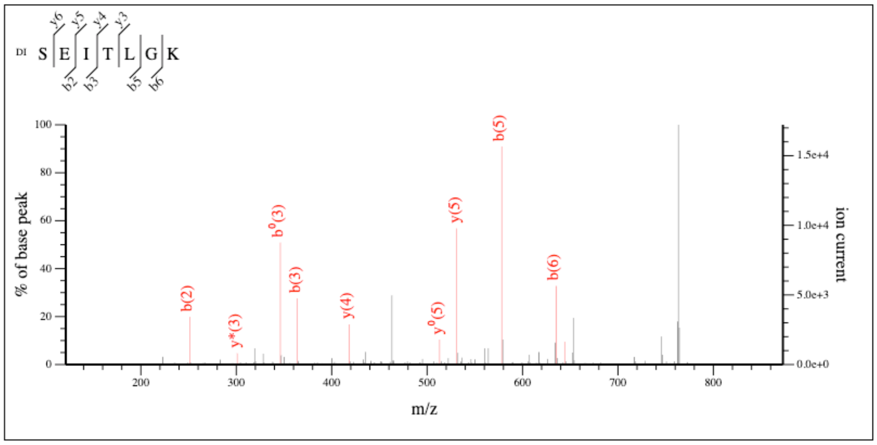

**Monoisotopic mass of neutral peptide Mr(calc):** 780.4864
**Fixed modifications:** Carbamidomethyl (C) (apply to specified residues or termini only)
**Variable modifications:**
**N-term :** Dimethyl:2H(6) (Protein N-term)
**Ions Score:** 27 **Expect:** 0.026
**Matches :** 10/27 fragment ions using 18 most intense peaks (help)

| # | b | $b^0$ | Seq. | y | y* | $y^0$ | # |
|---|---|---|---|---|---|---|---|
| 1 | 122.1083 | 104.0977 | S | | | | 7 |
| 2 | ***251.1509*** | 233.1403 | E | 660.3927 | 643.3661 | 642.3821 | 6 |
| 3 | ***364.2349*** | 346.2244 | I | **531.3501** | 514.3235 | 513.3395 | 5 |
| 4 | 465.2826 | 447.2720 | T | **418.2660** | 401.2395 | 400.2554 | 4 |
| 5 | ***578.3667*** | 560.3561 | L | 317.2183 | 300.1918 | | 3 |
| 6 | ***635.3881*** | 617.3776 | G | 204.1343 | 187.1077 | | 2 |
| 7 | | | K | 147.1128 | 130.0863 | | 1 |

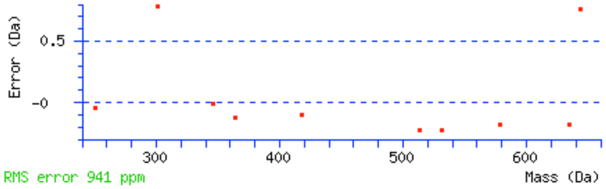


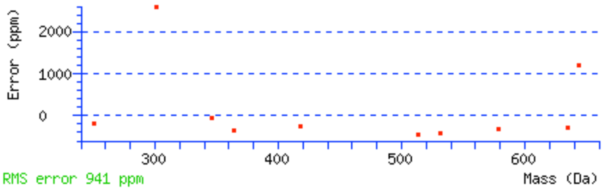


MS/MS Fragmentation of **SEITLGK**
Found in **P06169** in **Uniprot_yeastS288C**, Pyruvate decarboxylase isozyme 1 OS=Saccharomyces cerevisiae (strain ATCC 204508 / S288c) OX=559292 GN=PDC1 PE=1 SV=7

Match to Query 2083: 780.449328 from(781.456604,1+) intensity(615472.8750) rtinseconds(1027.341) index(2561)
Title: 20100530_methyl350mM_3.2849.2849.1
Data file D:\Panyue\Proteomic dataset\20100530_methyl350mM_3.RAW

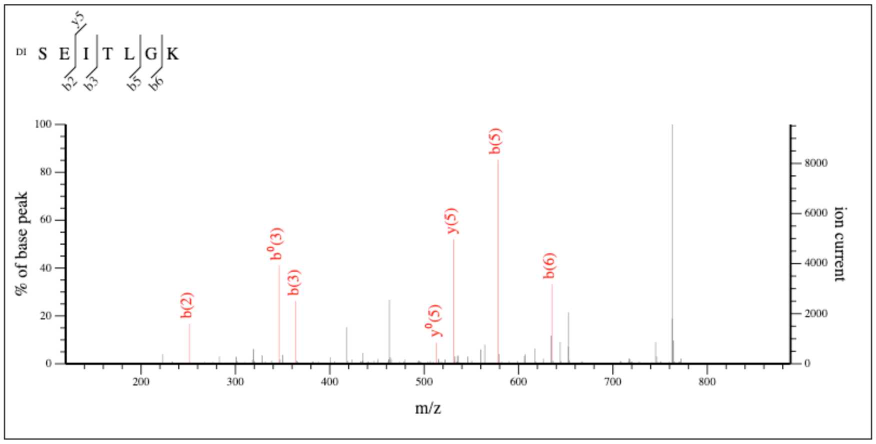

**Monoisotopic mass of neutral peptide Mr(calc):** 780.4864
**Fixed modifications:** Carbamidomethyl (C) (apply to specified residues or termini only)
**Variable modifications:**
**N-term :** Dimethyl:2H(6) (Protein N-term)
**Ions Score:** 24 **Expect:** 0.05
**Matches :** 7/27 fragment ions using 12 most intense peaks (help)

| # | b | b$^0$ | Seq. | y | y* | y$^0$ | # |
|---|---|---|---|---|---|---|---|
| 1 | 122.1083 | 104.0977 | S | | | | 7 |
| 2 | 251.1509 | 233.1403 | E | 660.3927 | 643.3661 | 642.3821 | 6 |
| 3 | 364.2349 | 346.2244 | I | 531.3501 | 514.3235 | 513.3395 | 5 |
| 4 | 465.2826 | 447.2720 | T | 418.2660 | 401.2395 | 400.2554 | 4 |
| 5 | 578.3667 | 560.3561 | L | 317.2183 | 300.1918 | | 3 |
| 6 | 635.3881 | 617.3776 | G | 204.1343 | 187.1077 | | 2 |
| 7 | | | K | 147.1128 | 130.0863 | | 1 |

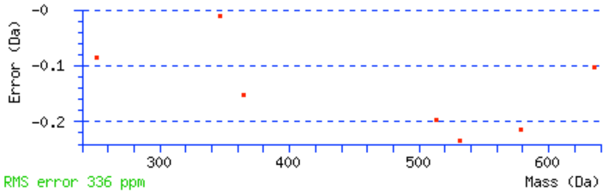


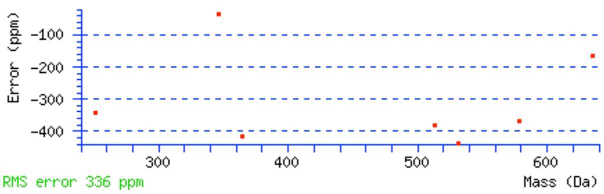


### 62. Ted1 (1 peptide in 350mM fraction 3)

MS/MS Fragmentation of **LRCAVK**
Found in **P40533** in **Uniprot_yeastS288C**, Protein TED1 OS=Saccharomyces cerevisiae (strain ATCC 204508 / S288c) OX=559292 GN=TED1 PE=1 SV=1

Match to Query 2078: 779.468737 from(780.476013,1+) intensity(142841.1406) rtinseconds(1083.7816) index(2715)
Title: 20100530_methyl350mM_3.3020.3020.1
Data file D:\Panyue\Proteomic dataset\20100530_methyl350mM_3.RAW

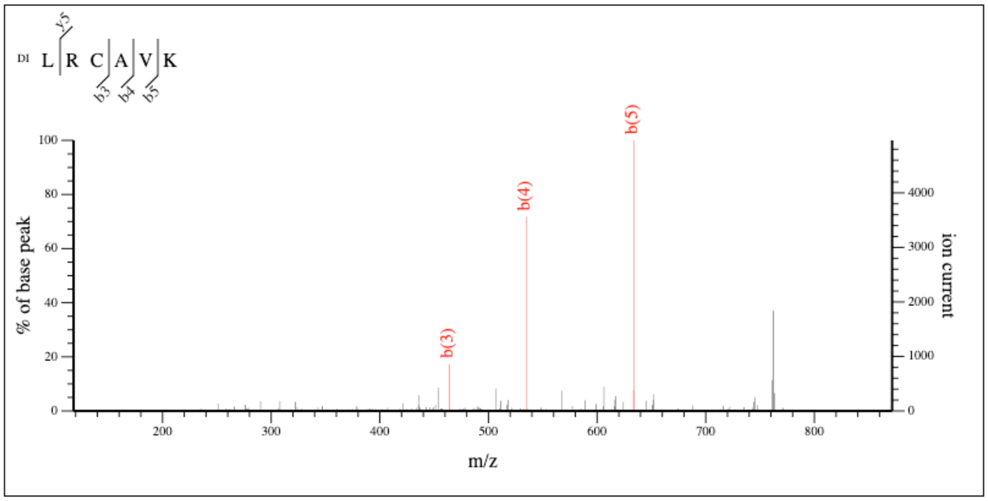

```
Monoisotopic mass of neutral peptide Mr(calc): 779.4958
Fixed modifications: Carbamidomethyl (C) (apply to specified residues or termini only)
Variable modifications:
N-term : Dimethyl:2H(6) (Protein N-term)
Ions Score: 21  Expect: 0.059
Matches : 4/19 fragment ions using 6 most intense peaks   (help)
```

| # | b | b* | Seq. | y | y* | # |
|---|---|---|---|---|---|---|
| 1 | 148.1603 | | L | | | 6 |
| 2 | 304.2614 | 287.2349 | R | 633.3501 | 616.3235 | 5 |
| 3 | 464.2921 | 447.2655 | C | 477.2490 | 460.2224 | 4 |
| 4 | 535.3292 | 518.3026 | A | 317.2183 | 300.1918 | 3 |
| 5 | 634.3976 | 617.3710 | V | 246.1812 | 229.1547 | 2 |
| 6 | | | K | 147.1128 | 130.0863 | 1 |

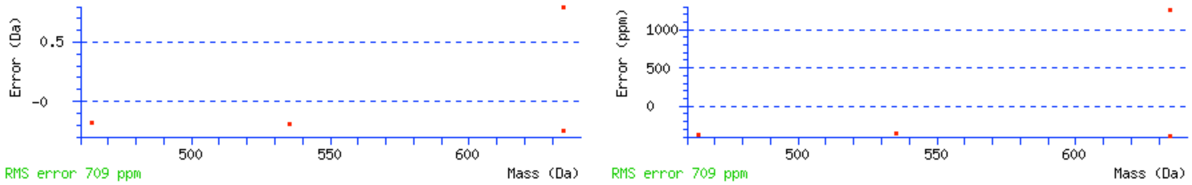


### 63. Eft1 (4 peptides in 350mM fraction 3)

MS/MS Fragmentation of **VAFTVDQMRSLMDK**
Found in **P32324** in **Uniprot_yeastS288C**, Elongation factor 2 OS=Saccharomyces cerevisiae (strain ATCC 204508 / S288c) OX=559292 GN=EFT1 PE=1 SV=1

Match to Query 9391: 1706.871678 from(854.443115,2+) intensity(1530816.5000) rtinseconds(2617.2481) index(6873)
Title: 20100530_methyl350mM_3.7645.7645.2
Data file D:\Panyue\Proteomic dataset\20100530_methyl350mM_3.RAW

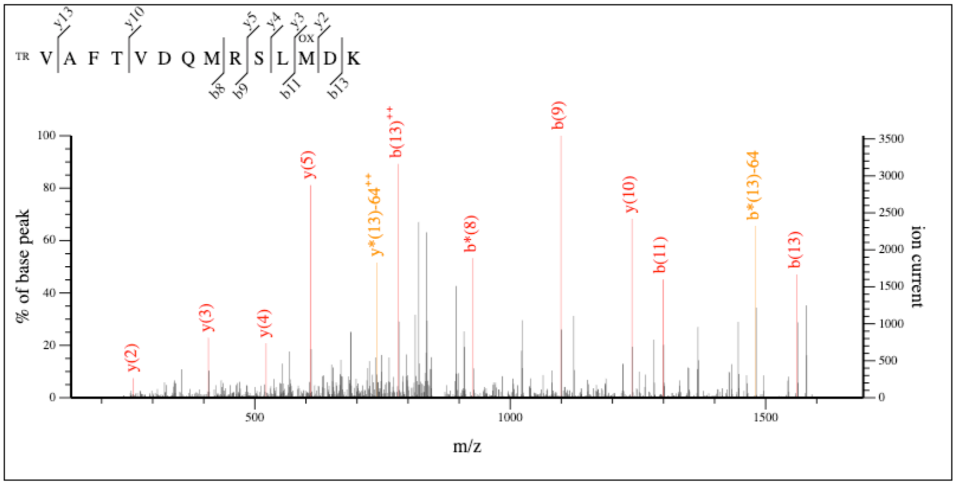

**Monoisotopic mass of neutral peptide Mr(calc):** 1706.9046
**Fixed modifications:** Carbamidomethyl (C) (apply to specified residues or termini only)
**Variable modifications:**
**N-term :** Trimethyl:2H(9) (Protein N-term)
**M12 :** Oxidation (M), with neutral losses 0.0000(shown in table), 63.9983
**Ions Score:** 26 **Expect:** 0.043
**Matches :** 14/214 fragment ions using 14 most intense peaks (help)

| # | b | b++ | b* | b*++ | b0 | b0++ | Seq. | y | y++ | y* | y*++ | y0 | y0++ | # |
|---|---|---|---|---|---|---|---|---|---|---|---|---|---|---|
| 1 | 151.1791 | 76.0932 | | | | | V | | | | | | | 14 |
| 2 | 222.2162 | 111.6118 | | | | | A | 1557.7400 | 779.3736 | 1540.7134 | 770.8604 | 1539.7294 | 770.3683 | 13 |
| 3 | 369.2847 | 185.1460 | | | | | F | 1486.7029 | 743.8551 | 1469.6763 | 735.3418 | 1468.6923 | 734.8498 | 12 |
| 4 | 470.3323 | 235.6698 | | | 452.3218 | 226.6645 | T | 1339.6344 | 670.3209 | 1322.6079 | 661.8076 | 1321.6239 | 661.3156 | 11 |
| 5 | 569.4008 | 285.2040 | | | 551.3902 | 276.1987 | V | 1238.5868 | 619.7970 | 1221.5602 | 611.2837 | 1220.5762 | 610.7917 | 10 |
| 6 | 684.4277 | 342.7175 | | | 666.4171 | 333.7122 | D | 1139.5184 | 570.2628 | 1122.4918 | 561.7495 | 1121.5078 | 561.2575 | 9 |
| 7 | 812.4863 | 406.7468 | 795.4597 | 398.2335 | 794.4757 | 397.7415 | Q | 1024.4914 | 512.7493 | 1007.4649 | 504.2361 | 1006.4808 | 503.7441 | 8 |
| 8 | 943.5268 | 472.2670 | 926.5002 | 463.7537 | 925.5162 | 463.2617 | M | 896.4328 | 448.7201 | 879.4063 | 440.2068 | 878.4223 | 439.7148 | 7 |
| 9 | 1099.6279 | 550.3176 | 1082.6013 | 541.8043 | 1081.6173 | 541.3123 | R | 765.3924 | 383.1998 | 748.3658 | 374.6865 | 747.3818 | 374.1945 | 6 |
| 10 | 1186.6599 | 593.8336 | 1169.6333 | 585.3203 | 1168.6493 | 584.8283 | S | 609.2912 | 305.1493 | 592.2647 | 296.6360 | 591.2807 | 296.1440 | 5 |
| 11 | 1299.7440 | 650.3756 | 1282.7174 | 641.8623 | 1281.7334 | 641.3703 | L | 522.2592 | 261.6332 | 505.2327 | 253.1200 | 504.2486 | 252.6280 | 4 |
| 12 | 1446.7794 | 723.8933 | 1429.7528 | 715.3800 | 1428.7688 | 714.8880 | M | 409.1751 | 205.0912 | 392.1486 | 196.5779 | 391.1646 | 196.0859 | 3 |
| 13 | 1561.8063 | 781.4068 | 1544.7798 | 772.8935 | 1543.7957 | 772.4015 | D | 262.1397 | 131.5735 | 245.1132 | 123.0602 | 244.1292 | 122.5682 | 2 |
| 14 | | | | | | | K | 147.1128 | 74.0600 | 130.0863 | 65.5468 | | | 1 |

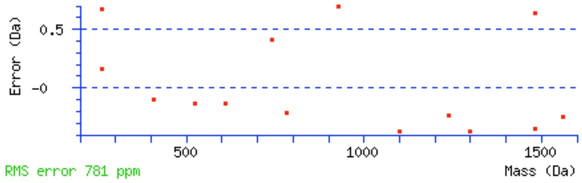


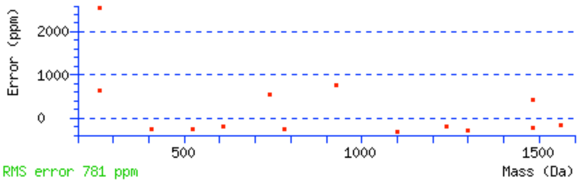


MS/MS Fragmentation of **VAFTVDQMRSLMDK**
Found in **P32324** in **Uniprot_yeastS288C**, Elongation factor 2 OS=Saccharomyces cerevisiae (strain ATCC 204508 / S288c) OX=559292 GN=EFT1 PE=1 SV=1

Match to Query 9392: 1706.871800 from(854.443176,2+) intensity(1791171.5000) rtinseconds(2619.8269) index(6880)
Title: 20100530_methyl350mM_3.7653.7653.2
Data file D:\Panyue\Proteomic dataset\20100530_methyl350mM_3.RAW

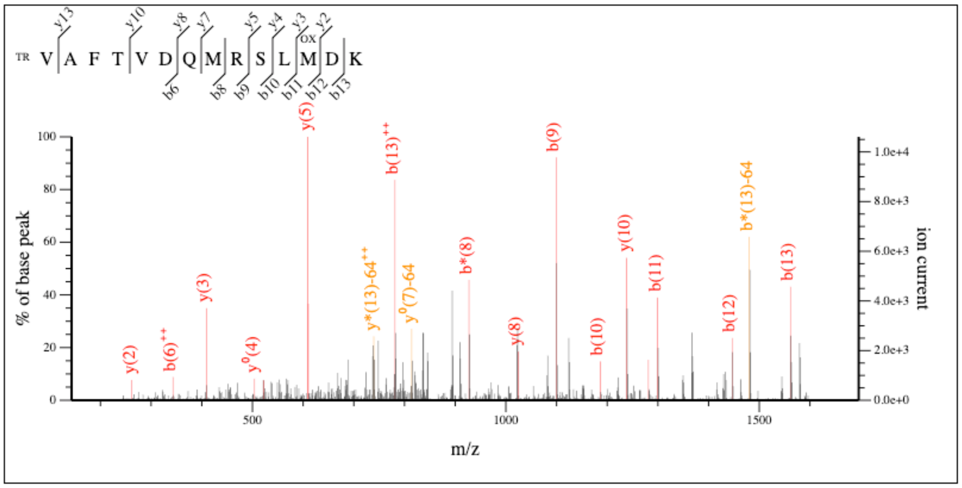

**Monoisotopic mass of neutral peptide Mr(calc):** 1706.9046
**Fixed modifications:** Carbamidomethyl (C) (apply to specified residues or termini only)
**Variable modifications:**
**N-term :** Trimethyl:2H(9) (Protein N-term)
**M12** : Oxidation (M), with neutral losses 0.0000(shown in table), 63.9983
**Ions Score:** 23 **Expect:** 0.083
**Matches :** 26/214 fragment ions using 42 most intense peaks (help)

| # | b | b++ | b* | b*++ | b0 | b0++ | Seq. | y | y++ | y* | y*++ | y0 | y0++ | # |
|---|---|---|---|---|---|---|---|---|---|---|---|---|---|---|
| 1 | 151.1791 | 76.0932 | | | | | V | | | | | | | 14 |
| 2 | 222.2162 | 111.6118 | | | | | A | 1557.7400 | 779.3736 | 1540.7134 | 770.8604 | 1539.7294 | 770.3683 | 13 |
| 3 | 369.2847 | 185.1460 | | | | | F | 1486.7029 | 743.8551 | 1469.6763 | 735.3418 | 1468.6923 | 734.8498 | 12 |
| 4 | 470.3323 | 235.6698 | | | 452.3218 | 226.6645 | T | 1339.6344 | 670.3209 | 1322.6079 | 661.8076 | 1321.6239 | 661.3156 | 11 |
| 5 | 569.4008 | 285.2040 | | | 551.3902 | 276.1987 | V | ***1238.5868*** | 619.7970 | 1221.5602 | 611.2837 | 1220.5762 | 610.7917 | 10 |
| 6 | 684.4277 | 342.7175 | | | 666.4171 | 333.7122 | D | 1139.5184 | 570.2628 | 1122.4918 | 561.7495 | 1121.5078 | 561.2575 | 9 |
| 7 | 812.4863 | 406.7468 | 795.4597 | 398.2335 | 794.4757 | 397.7415 | Q | ***1024.4914*** | 512.7493 | 1007.4649 | 504.2361 | 1006.4808 | 503.7441 | 8 |
| 8 | 943.5268 | 472.2670 | 926.5002 | 463.7537 | 925.5162 | 463.2617 | M | 896.4328 | 448.7201 | 879.4063 | 440.2068 | 878.4223 | 439.7148 | 7 |
| 9 | ***1099.6279*** | 550.3176 | 1082.6013 | 541.8043 | 1081.6173 | 541.3123 | R | 765.3924 | 383.1998 | 748.3658 | 374.6865 | 747.3818 | 374.1945 | 6 |
| 10 | ***1186.6599*** | 593.8336 | 1169.6333 | 585.3203 | 1168.6493 | 584.8283 | S | ***609.2912*** | 305.1493 | 592.2647 | 296.6360 | 591.2807 | 296.1440 | 5 |
| 11 | ***1299.7440*** | 650.3756 | 1282.7174 | 641.8623 | 1281.7334 | 641.3703 | L | ***522.2592*** | 261.6332 | 505.2327 | 253.1200 | 504.2486 | 252.6280 | 4 |
| 12 | ***1446.7794*** | 723.8933 | 1429.7528 | 715.3800 | 1428.7688 | 714.8880 | M | ***409.1751*** | 205.0912 | 392.1486 | 196.5779 | 391.1646 | 196.0859 | 3 |
| 13 | ***1561.8063*** | 781.4068 | 1544.7798 | 772.8935 | 1543.7957 | 772.4015 | D | ***262.1397*** | 131.5735 | 245.1132 | 123.0602 | 244.1292 | 122.5682 | 2 |
| 14 | | | | | | | K | 147.1128 | 74.0600 | 130.0863 | 65.5468 | | | 1 |

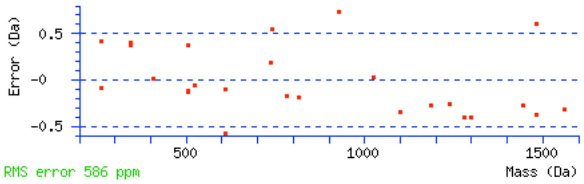


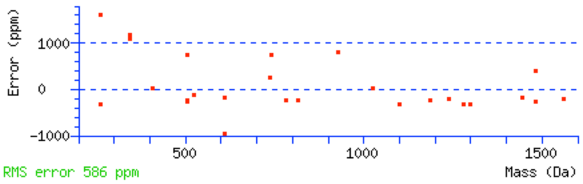


MS/MS Fragmentation of **VAFTVDQMRSLMDK**
Found in **P32324** in **Uniprot_yeastS288C**, Elongation factor 2 OS=Saccharomyces cerevisiae (strain ATCC 204508 / S288c) OX=559292 GN=EFT1 PE=1 SV=1

Match to Query 9393: 1706.872778 from(854.443665,2+) intensity(1791171.5000) rtinseconds(2620.0961) index(6881)
Title: 20100530_methyl350mM_3.7654.7654.2
Data file D:\Panyue\Proteomic dataset\20100530_methyl350mM_3.RAW

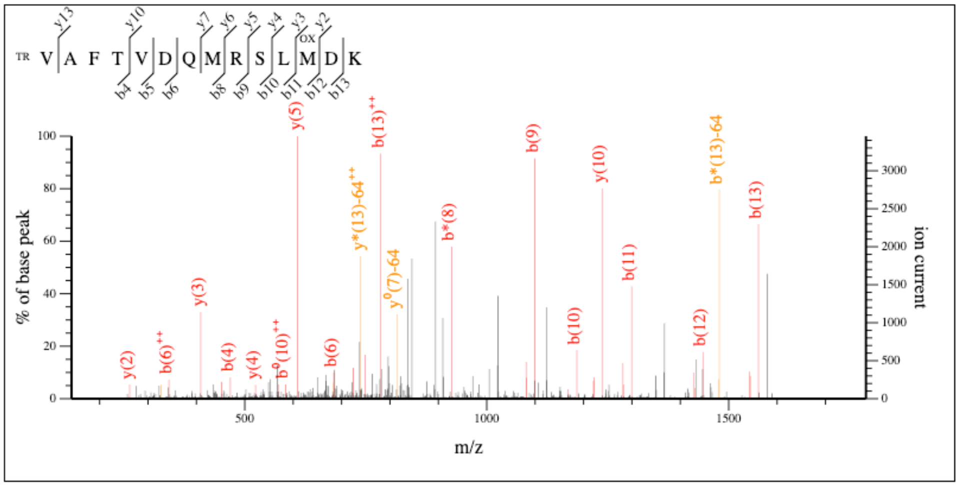

**Monoisotopic mass of neutral peptide Mr(calc):** 1706.9046
**Fixed modifications:** Carbamidomethyl (C) (apply to specified residues or termini only)
**Variable modifications:**
**N-term :** Trimethyl:2H(9) (Protein N-term)
**M12** : Oxidation (M), with neutral losses 0.0000(shown in table), 63.9983
**Ions Score:** 25 **Expect:** 0.051
**Matches :** 41/214 fragment ions using 85 most intense peaks (help)

| # | b | $b^{++}$ | b* | $b^{*++}$ | $b^0$ | $b^{0++}$ | Seq. | y | $y^{++}$ | y* | $y^{*++}$ | $y^0$ | $y^{0++}$ | # |
|---|---|---|---|---|---|---|---|---|---|---|---|---|---|---|
| 1 | 151.1791 | 76.0932 | | | | | V | | | | | | | 14 |
| 2 | 222.2162 | 111.6118 | | | | | A | 1557.7400 | 779.3736 | 1540.7134 | 770.8604 | 1539.7294 | 770.3683 | 13 |
| 3 | 369.2847 | 185.1460 | | | | | F | 1486.7029 | 743.8551 | 1469.6763 | 735.3418 | 1468.6923 | 734.8498 | 12 |
| 4 | ***470.3323*** | 235.6698 | | | 452.3218 | 226.6645 | T | 1339.6344 | 670.3209 | 1322.6079 | 661.8076 | 1321.6239 | 661.3156 | 11 |
| 5 | ***569.4008*** | 285.2040 | | | 551.3902 | 276.1987 | V | **1238.5868** | 619.7970 | 1221.5602 | 611.2837 | 1220.5762 | 610.7917 | 10 |
| 6 | ***684.4277*** | 342.7175 | | | 666.4171 | 333.7122 | D | 1139.5184 | 570.2628 | 1122.4918 | 561.7495 | 1121.5078 | 561.2575 | 9 |
| 7 | 812.4863 | 406.7468 | 795.4597 | 398.2335 | 794.4757 | 397.7415 | Q | 1024.4914 | 512.7493 | 1007.4649 | 504.2361 | 1006.4808 | 503.7441 | 8 |
| 8 | 943.5268 | 472.2670 | 926.5002 | 463.7537 | 925.5162 | 463.2617 | M | 896.4328 | 448.7201 | 879.4063 | 440.2068 | 878.4223 | 439.7148 | 7 |
| 9 | ***1099.6279*** | 550.3176 | 1082.6013 | 541.8043 | 1081.6173 | 541.3123 | R | 765.3924 | 383.1998 | 748.3658 | 374.6865 | 747.3818 | 374.1945 | 6 |
| 10 | ***1186.6599*** | 593.8336 | 1169.6333 | 585.3203 | 1168.6493 | 584.8283 | S | **609.2912** | 305.1493 | 592.2647 | 296.6360 | 591.2807 | 296.1440 | 5 |
| 11 | ***1299.7440*** | 650.3756 | 1282.7174 | 641.8623 | 1281.7334 | 641.3703 | L | **522.2592** | 261.6332 | 505.2327 | 253.1200 | 504.2486 | 252.6280 | 4 |
| 12 | ***1446.7794*** | 723.8933 | 1429.7528 | 715.3800 | 1428.7688 | 714.8880 | M | **409.1751** | 205.0912 | 392.1486 | 196.5779 | 391.1646 | 196.0859 | 3 |
| 13 | ***1561.8063*** | 781.4068 | 1544.7798 | 772.8935 | 1543.7957 | 772.4015 | D | **262.1397** | 131.5735 | 245.1132 | 123.0602 | 244.1292 | 122.5682 | 2 |
| 14 | | | | | | | K | 147.1128 | 74.0600 | 130.0863 | 65.5468 | | | 1 |

MS/MS Fragmentation of **VAFTVDQMRSLMDK**
Found in **P32324** in **Uniprot_yeastS288C**, Elongation factor 2 OS=Saccharomyces cerevisiae (strain ATCC 204508 / S288c) OX=559292 GN=EFT1 PE=1 SV=1

Match to Query 9394: 1706.874852 from(854.444702,2+) intensity(2586890.0000) rtinseconds(2623.0387) index(6889)
Title: 20100530_methyl350mM_3.7663.7663.2
Data file D:\Panyue\Proteomic dataset\20100530_methyl350mM_3.RAW

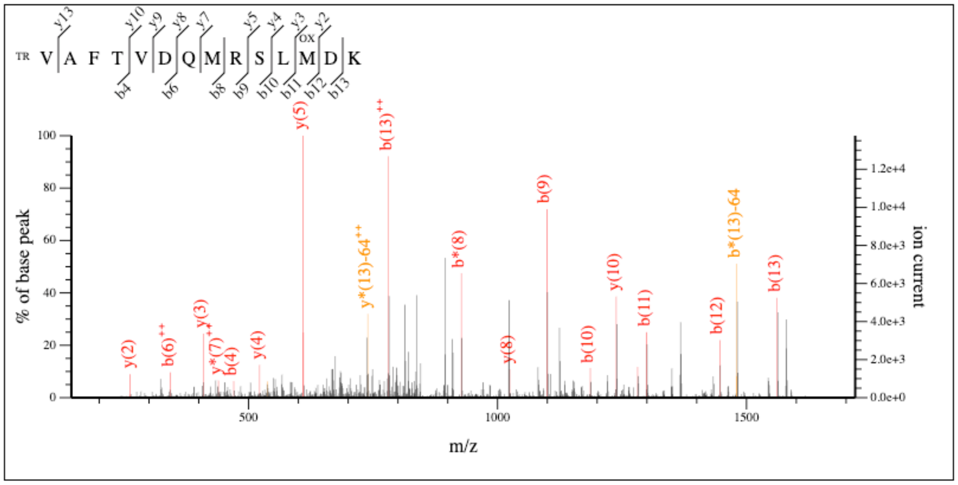

```
Monoisotopic mass of neutral peptide Mr(calc): 1706.9046
Fixed modifications: Carbamidomethyl (C) (apply to specified residues or termini only)
Variable modifications:
N-term : Trimethyl:2H(9) (Protein N-term)
M12    : Oxidation (M), with neutral losses 0.0000(shown in table), 63.9983
Ions Score: 24  Expect: 0.068
Matches : 25/214 fragment ions using 42 most intense peaks   (help)
```

| # | b | b++ | b* | b*++ | b0 | b0++ | Seq. | y | y++ | y* | y*++ | y0 | y0++ | # |
|---|---|---|---|---|---|---|---|---|---|---|---|---|---|---|
| 1 | 151.1791 | 76.0932 | | | | | V | | | | | | | 14 |
| 2 | 222.2162 | 111.6118 | | | | | A | 1557.7400 | 779.3736 | 1540.7134 | 770.8604 | 1539.7294 | 770.3683 | 13 |
| 3 | 369.2847 | 185.1460 | | | | | F | 1486.7029 | 743.8551 | 1469.6763 | 735.3418 | 1468.6923 | 734.8498 | 12 |
| 4 | 470.3323 | 235.6698 | | | 452.3218 | 226.6645 | T | 1339.6344 | 670.3209 | 1322.6079 | 661.8076 | 1321.6239 | 661.3156 | 11 |
| 5 | 569.4008 | 285.2040 | | | 551.3902 | 276.1987 | V | 1238.5868 | 619.7970 | 1221.5602 | 611.2837 | 1220.5762 | 610.7917 | 10 |
| 6 | 684.4277 | 342.7175 | | | 666.4171 | 333.7122 | D | 1139.5184 | 570.2628 | 1122.4918 | 561.7495 | 1121.5078 | 561.2575 | 9 |
| 7 | 812.4863 | 406.7468 | 795.4597 | 398.2335 | 794.4757 | 397.7415 | Q | 1024.4914 | 512.7493 | 1007.4649 | 504.2361 | 1006.4808 | 503.7441 | 8 |
| 8 | 943.5268 | 472.2670 | 926.5002 | 463.7537 | 925.5162 | 463.2617 | M | 896.4328 | 448.7201 | 879.4063 | 440.2068 | 878.4223 | 439.7148 | 7 |
| 9 | 1099.6279 | 550.3176 | 1082.6013 | 541.8043 | 1081.6173 | 541.3123 | R | 765.3924 | 383.1998 | 748.3658 | 374.6865 | 747.3818 | 374.1945 | 6 |
| 10 | 1186.6599 | 593.8336 | 1169.6333 | 585.3203 | 1168.6493 | 584.8283 | S | 609.2912 | 305.1493 | 592.2647 | 296.6360 | 591.2807 | 296.1440 | 5 |
| 11 | 1299.7440 | 650.3756 | 1282.7174 | 641.8623 | 1281.7334 | 641.3703 | L | 522.2592 | 261.6332 | 505.2327 | 253.1200 | 504.2486 | 252.6280 | 4 |
| 12 | 1446.7794 | 723.8933 | 1429.7528 | 715.3800 | 1428.7688 | 714.8880 | M | 409.1751 | 205.0912 | 392.1486 | 196.5779 | 391.1646 | 196.0859 | 3 |
| 13 | 1561.8063 | 781.4068 | 1544.7798 | 772.8935 | 1543.7957 | 772.4015 | D | 262.1397 | 131.5735 | 245.1132 | 123.0602 | 244.1292 | 122.5682 | 2 |
| 14 | | | | | | | K | 147.1128 | 74.0600 | 130.0863 | 65.5468 | | | 1 |

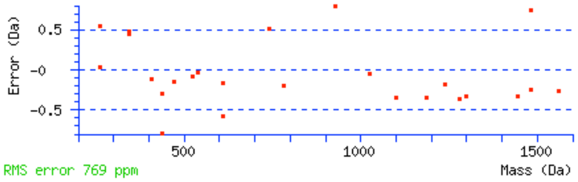


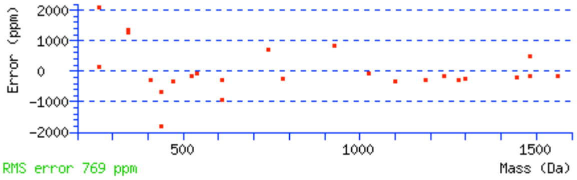


### 64. Dma2 (1 peptide in 400mM fraction 1)

MS/MS Fragmentation of **YTPIPANTPAPTAPTSSMTSNSSSASNANTTSSSGINPRNR**
Found in **P53924** in **Uniprot_yeastS288C**, E3 ubiquitin-protein ligase DMA2 OS=Saccharomyces cerevisiae (strain ATCC 204508 / S288c) OX=559292 GN=DMA2 PE=1 SV=1

Match to Query 11756: 4124.010896 from(1032.010000,4+) intensity(322014.1562) rtinseconds(4430.0867) index(11423)
Title: 20100530_methyl400mM_1.12706.12706.
Data file D:\Panyue\Proteomic dataset\20100530_methyl400mM_1.RAW

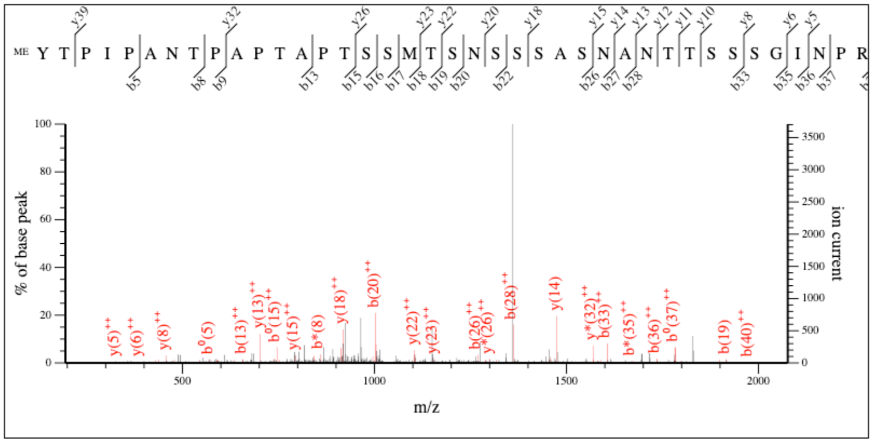


```
Monoisotopic mass of neutral peptide Mr(calc): 4123.9544
Fixed modifications: Carbamidomethyl (C) (apply to specified residues or termini only)
Variable modifications:
N-term : Methyl:2H(3) (Protein N-term)
Ions Score: 22  Expect: 0.028
Matches : 59/452 fragment ions using 153 most intense peaks   (help)
```

| # | b | b++ | b* | b*++ | b0 | b0++ | Seq. | y | y++ | y* | y*++ | y0 | y0++ | # |
|---|---|---|---|---|---|---|---|---|---|---|---|---|---|---|
| 1 | 181.1051 | 91.0562 | | | | | Y | | | | | | | 41 |
| 2 | 282.1528 | 141.5800 | | | 264.1422 | 132.5747 | T | 3944.8639 | 1972.9356 | 3927.8373 | 1964.4223 | 3926.8533 | 1963.9303 | 40 |
| 3 | 379.2055 | 190.1064 | | | 361.1950 | 181.1011 | P | 3843.8162 | 1922.4117 | 3826.7897 | 1913.8985 | 3825.8056 | 1913.4065 | 39 |
| 4 | 492.2896 | 246.6484 | | | 474.2790 | 237.6432 | I | 3746.7634 | 1873.8854 | 3729.7369 | 1865.3721 | 3728.7529 | 1864.8801 | 38 |
| 5 | 589.3424 | 295.1748 | | | 571.3318 | 286.1695 | P | 3633.6794 | 1817.3433 | 3616.6528 | 1808.8301 | 3615.6688 | 1808.3380 | 37 |
| 6 | 660.3795 | 330.6934 | | | 642.3689 | 321.6881 | A | 3536.6266 | 1768.8169 | 3519.6001 | 1760.3037 | 3518.6161 | 1759.8117 | 36 |
| 7 | 774.4224 | 387.7148 | 757.3958 | 379.2016 | 756.4118 | 378.7096 | N | 3465.5895 | 1733.2984 | 3448.5630 | 1724.7851 | 3447.5789 | 1724.2931 | 35 |
| 8 | 875.4701 | 438.2387 | 858.4435 | 429.7254 | 857.4595 | 429.2334 | T | 3351.5466 | 1676.2769 | 3334.5200 | 1667.7637 | 3333.5360 | 1667.2716 | 34 |
| 9 | 972.5228 | 486.7651 | 955.4963 | 478.2518 | 954.5123 | 477.7598 | P | 3250.4989 | 1625.7531 | 3233.4723 | 1617.2398 | 3232.4883 | 1616.7478 | 33 |
| 10 | 1043.5600 | 522.2836 | 1026.5334 | 513.7703 | 1025.5494 | 513.2783 | A | 3153.4461 | 1577.2267 | 3136.4196 | 1568.7134 | 3135.4356 | 1568.2214 | 32 |
| 11 | 1140.6127 | 570.8100 | 1123.5862 | 562.2967 | 1122.6022 | 561.8047 | P | 3082.4090 | 1541.7081 | 3065.3825 | 1533.1949 | 3064.3985 | 1532.7029 | 31 |
| 12 | 1241.6604 | 621.3338 | 1224.6338 | 612.8206 | 1223.6498 | 612.3286 | T | 2985.3563 | 1493.1818 | 2968.3297 | 1484.6685 | 2967.3457 | 1484.1765 | 30 |
| 13 | 1312.6975 | 656.8524 | 1295.6710 | 648.3391 | 1294.6869 | 647.8471 | A | 2884.3086 | 1442.6579 | 2867.2820 | 1434.1447 | 2866.2980 | 1433.6526 | 29 |
| 14 | 1409.7503 | 705.3788 | 1392.7237 | 696.8655 | 1391.7397 | 696.3735 | P | 2813.2715 | 1407.1394 | 2796.2449 | 1398.6261 | 2795.2609 | 1398.1341 | 28 |
| 15 | 1510.7980 | 755.9026 | 1493.7714 | 747.3893 | 1492.7874 | 746.8973 | T | 2716.2187 | 1358.6130 | 2699.1922 | 1350.0997 | 2698.2081 | 1349.6077 | 27 |
| 16 | 1597.8300 | 799.4186 | 1580.8034 | 790.9054 | 1579.8194 | 790.4133 | S | 2615.1710 | 1308.0891 | 2598.1445 | 1299.5759 | 2597.1605 | 1299.0839 | 26 |
| 17 | 1684.8620 | 842.9346 | 1667.8355 | 834.4214 | 1666.8514 | 833.9294 | S | 2528.1390 | 1264.5731 | 2511.1124 | 1256.0599 | 2510.1284 | 1255.5679 | 25 |
| 18 | 1815.9025 | 908.4549 | 1798.8759 | 899.9416 | 1797.8919 | 899.4496 | M | 2441.1070 | 1221.0571 | 2424.0804 | 1212.5438 | 2423.0964 | 1212.0518 | 24 |
| 19 | 1916.9502 | 958.9787 | 1899.9236 | 950.4655 | 1898.9396 | 949.9734 | T | 2310.0665 | 1155.5369 | 2293.0399 | 1147.0236 | 2292.0559 | 1146.5316 | 23 |
| 20 | 2003.9822 | 1002.4947 | 1986.9557 | 993.9815 | 1985.9716 | 993.4895 | S | 2209.0188 | 1105.0130 | 2191.9923 | 1096.4998 | 2191.0082 | 1096.0078 | 22 |
| 21 | 2118.0251 | 1059.5162 | 2100.9986 | 1051.0029 | 2100.0146 | 1050.5109 | N | 2121.9868 | 1061.4970 | 2104.9602 | 1052.9837 | 2103.9762 | 1052.4917 | 21 |
| 22 | 2205.0572 | 1103.0322 | 2188.0306 | 1094.5189 | 2187.0466 | 1094.0269 | S | 2007.9438 | 1004.4756 | 1990.9173 | 995.9623 | 1989.9333 | 995.4703 | 20 |
| 23 | 2292.0892 | 1146.5482 | 2275.0626 | 1138.0350 | 2274.0786 | 1137.5429 | S | 1920.9118 | 960.9595 | 1903.8853 | 952.4463 | 1902.9013 | 951.9543 | 19 |
| 24 | 2379.1212 | 1190.0642 | 2362.0947 | 1181.5510 | 2361.1106 | 1181.0590 | S | 1833.8798 | 917.4435 | 1816.8532 | 908.9303 | 1815.8692 | 908.4383 | 18 |
| 25 | 2450.1583 | 1225.5828 | 2433.1318 | 1217.0695 | 2432.1478 | 1216.5775 | A | 1746.8478 | 873.9275 | 1729.8212 | 865.4142 | 1728.8372 | 864.9222 | 17 |
| 26 | 2537.1904 | 1269.0988 | 2520.1638 | 1260.5855 | 2519.1798 | 1260.0935 | S | 1675.8106 | 838.4090 | 1658.7841 | 829.8957 | 1657.8001 | 829.4037 | 16 |
| 27 | 2651.2333 | 1326.1203 | 2634.2067 | 1317.6070 | 2633.2227 | 1317.1150 | N | 1588.7786 | 794.8929 | 1571.7521 | 786.3797 | 1570.7681 | 785.8877 | 15 |
| 28 | 2722.2704 | 1361.6388 | 2705.2438 | 1353.1256 | 2704.2598 | 1352.6336 | A | 1474.7357 | 737.8715 | 1457.7091 | 729.3582 | 1456.7251 | 728.8662 | 14 |
| 29 | 2836.3133 | 1418.6603 | 2819.2868 | 1410.1470 | 2818.3028 | 1409.6550 | N | 1403.6986 | 702.3529 | 1386.6720 | 693.8397 | 1385.6880 | 693.3476 | 13 |
| 30 | 2937.3610 | 1469.1841 | 2920.3345 | 1460.6709 | 2919.3504 | 1460.1789 | T | 1289.6557 | 645.3315 | 1272.6291 | 636.8182 | 1271.6451 | 636.3262 | 12 |
| 31 | 3038.4087 | 1519.7080 | 3021.3821 | 1511.1947 | 3020.3981 | 1510.7027 | T | 1188.6080 | 594.8076 | 1171.5814 | 586.2943 | 1170.5974 | 585.8023 | 11 |
| 32 | 3125.4407 | 1563.2240 | 3108.4142 | 1554.7107 | 3107.4301 | 1554.2187 | S | 1087.5603 | 544.2838 | 1070.5337 | 535.7705 | 1069.5497 | 535.2785 | 10 |
| 33 | 3212.4727 | 1606.7400 | 3195.4462 | 1598.2267 | 3194.4622 | 1597.7347 | S | 1000.5283 | 500.7678 | 983.5017 | 492.2545 | 982.5177 | 491.7625 | 9 |
| 34 | 3299.5048 | 1650.2560 | 3282.4782 | 1641.7427 | 3281.4942 | 1641.2507 | S | 913.4962 | 457.2518 | 896.4697 | 448.7385 | 895.4857 | 448.2465 | 8 |
| 35 | 3356.5262 | 1678.7668 | 3339.4997 | 1670.2535 | 3338.5157 | 1669.7615 | G | 826.4642 | 413.7357 | 809.4377 | 405.2225 | | | 7 |
| 36 | 3469.6103 | 1735.3088 | 3452.5837 | 1726.7955 | 3451.5997 | 1726.3035 | I | 769.4427 | 385.2250 | 752.4162 | 376.7117 | | | 6 |
| 37 | 3583.6532 | 1792.3302 | 3566.6267 | 1783.8170 | 3565.6427 | 1783.3250 | N | 656.3587 | 328.6830 | 639.3321 | 320.1697 | | | 5 |
| 38 | 3680.7060 | 1840.8566 | 3663.6794 | 1832.3434 | 3662.6954 | 1831.8513 | P | 542.3158 | 271.6615 | 525.2892 | 263.1482 | | | 4 |
| 39 | 3836.8071 | 1918.9072 | 3819.7805 | 1910.3939 | 3818.7965 | 1909.9019 | R | 445.2630 | 223.1351 | 428.2364 | 214.6219 | | | 3 |
| 40 | 3950.8500 | 1975.9286 | 3933.8235 | 1967.4154 | 3932.8395 | 1966.9234 | N | 289.1619 | 145.0846 | 272.1353 | 136.5713 | | | 2 |
| 41 | | | | | | | R | 175.1190 | 88.0631 | 158.0924 | 79.5498 | | | 1 |

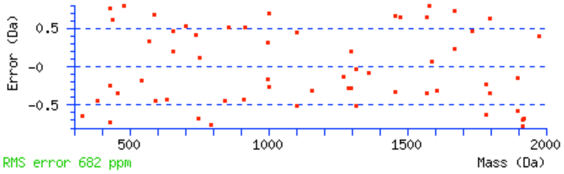


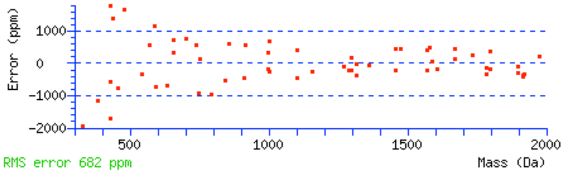


65. Mon2 (4 peptides in 400mM fraction 1)

MS/MS Fragmentation of **AMNTGGFDSMQRQLEAELRSLSSESK**
Found in **P48563** in **Uniprot_yeastS288C**, Protein MON2 OS=Saccharomyces cerevisiae (strain ATCC 204508 / S288c) OX=559292 GN=MON2 PE=1 SV=1

Match to Query 16233: 2938.457847 from(980.493225,3+) intensity(147633.9688) rtinseconds(5335.513) index(13591)
Title: 20100530_methyl400mM_1.15117.15117.3
Data file D:\Panyue\Proteomic dataset\20100530_methyl400mM_1.RAW

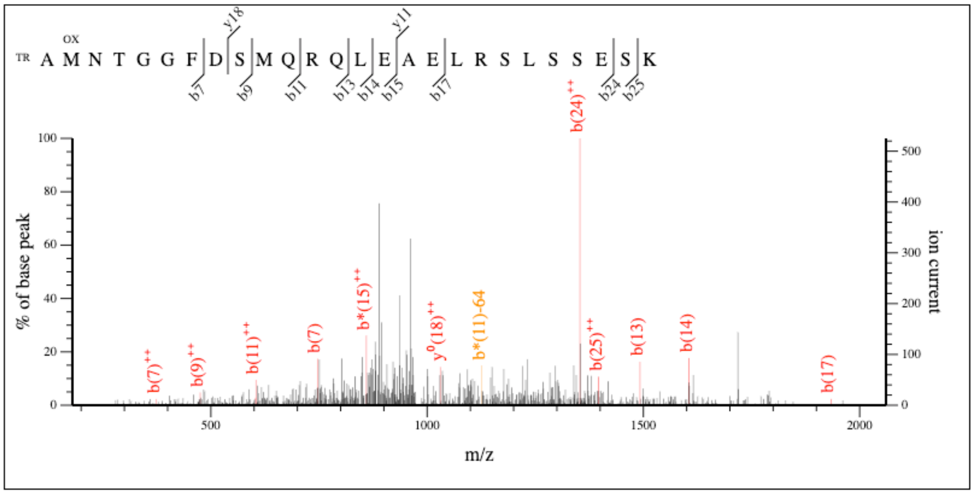


**Monoisotopic mass of neutral peptide Mr(calc):** 2938.4474
**Fixed modifications:** Carbamidomethyl (C) (apply to specified residues or termini only)
**Variable modifications:**
**N-term** : Trimethyl:2H(9) (Protein N-term)
**M2** : Oxidation (M), with neutral losses 0.0000(shown in table), 63.9983
**Ions Score:** 24 **Expect:** 0.048
**Matches** : 14/432 fragment ions using 17 most intense peaks (help)

| # | b | b++ | b* | b*++ | b0 | b0++ | Seq. | y | y++ | y* | y*++ | y0 | y0++ | # |
|---|---|---|---|---|---|---|---|---|---|---|---|---|---|---|
| 1 | 123.1478 | 62.0776 | | | | | A | | | | | | | 26 |
| 2 | 270.1832 | 135.5953 | | | | | M | 2817.3142 | 1409.1607 | 2800.2876 | 1400.6475 | 2799.3036 | 1400.1554 | 25 |
| 3 | 384.2262 | 192.6167 | 367.1996 | 184.1034 | | | N | 2670.2788 | 1335.6430 | 2653.2522 | 1327.1298 | 2652.2682 | 1326.6377 | 24 |
| 4 | 485.2738 | 243.1406 | 468.2473 | 234.6273 | 467.2633 | 234.1353 | T | 2556.2359 | 1278.6216 | 2539.2093 | 1270.1083 | 2538.2253 | 1269.6163 | 23 |
| 5 | 542.2953 | 271.6513 | 525.2688 | 263.1380 | 524.2847 | 262.6460 | G | 2455.1882 | 1228.0977 | 2438.1616 | 1219.5845 | 2437.1776 | 1219.0924 | 22 |
| 6 | 599.3168 | 300.1620 | 582.2902 | 291.6487 | 581.3062 | 291.1567 | G | 2398.1667 | 1199.5870 | 2381.1402 | 1191.0737 | 2380.1561 | 1190.5817 | 21 |
| 7 | ***746.3852*** | ***373.6962*** | 729.3586 | 365.1830 | 728.3746 | 364.6909 | F | 2341.1452 | 1171.0763 | 2324.1187 | 1162.5630 | 2323.1347 | 1162.0710 | 20 |
| 8 | 861.4121 | 431.2097 | 844.3856 | 422.6964 | 843.4016 | 422.2044 | D | 2194.0768 | 1097.5421 | 2177.0503 | 1089.0288 | 2176.0663 | 1088.5368 | 19 |
| 9 | 948.4441 | ***474.7257*** | 931.4176 | 466.2124 | 930.4336 | 465.7204 | S | 2079.0499 | 1040.0286 | 2062.0233 | 1031.5153 | 2061.0393 | 1031.0233 | 18 |
| 10 | 1079.4846 | 540.2460 | 1062.4581 | 531.7327 | 1061.4741 | 531.2407 | M | 1992.0179 | 996.5126 | 1974.9913 | 987.9993 | 1974.0073 | 987.5073 | 17 |
| 11 | 1207.5432 | ***604.2752*** | 1190.5167 | 595.7620 | 1189.5326 | 595.2700 | Q | 1860.9774 | 930.9923 | 1843.9508 | 922.4791 | 1842.9668 | 921.9870 | 16 |
| 12 | 1363.6443 | 682.3258 | 1346.6178 | 673.8125 | 1345.6338 | 673.3205 | R | 1732.9188 | 866.9630 | 1715.8923 | 858.4498 | 1714.9082 | 857.9578 | 15 |
| 13 | ***1491.7029*** | ***746.3551*** | 1474.6764 | 737.8418 | 1473.6923 | 737.3498 | Q | 1576.8177 | 788.9125 | 1559.7911 | 780.3992 | 1558.8071 | 779.9072 | 14 |
| 14 | ***1604.7870*** | 802.8971 | 1587.7604 | 794.3838 | 1586.7764 | 793.8918 | L | 1448.7591 | 724.8832 | 1431.7326 | 716.3699 | 1430.7485 | 715.8779 | 13 |
| 15 | 1733.8296 | 867.4184 | 1716.8030 | 858.9051 | 1715.8190 | 858.4131 | E | 1335.6750 | 668.3412 | 1318.6485 | 659.8279 | 1317.6645 | 659.3359 | 12 |
| 16 | 1804.8667 | 902.9370 | 1787.8401 | 894.4237 | 1786.8561 | 893.9317 | A | 1206.6325 | 603.8199 | 1189.6059 | 595.3066 | 1188.6219 | 594.8146 | 11 |
| 17 | ***1933.9093*** | 967.4583 | 1916.8827 | 958.9450 | 1915.8987 | 958.4530 | E | 1135.5953 | 568.3013 | 1118.5688 | 559.7880 | 1117.5848 | 559.2960 | 10 |
| 18 | 2046.9933 | 1024.0003 | 2029.9668 | 1015.4870 | 2028.9828 | 1014.9950 | L | 1006.5527 | 503.7800 | 989.5262 | 495.2667 | 988.5422 | 494.7747 | 9 |
| 19 | 2203.0944 | 1102.0509 | 2186.0679 | 1093.5376 | 2185.0839 | 1093.0456 | R | 893.4687 | 447.2380 | 876.4421 | 438.7247 | 875.4581 | 438.2327 | 8 |
| 20 | 2290.1265 | 1145.5669 | 2273.0999 | 1137.0536 | 2272.1159 | 1136.5616 | S | 737.3676 | 369.1874 | 720.3410 | 360.6742 | 719.3570 | 360.1821 | 7 |
| 21 | 2403.2105 | 1202.1089 | 2386.1840 | 1193.5956 | 2385.2000 | 1193.1036 | L | 650.3355 | 325.6714 | 633.3090 | 317.1581 | 632.3250 | 316.6661 | 6 |
| 22 | 2490.2426 | 1245.6249 | 2473.2160 | 1237.1116 | 2472.2320 | 1236.6196 | S | 537.2515 | 269.1294 | 520.2249 | 260.6161 | 519.2409 | 260.1241 | 5 |
| 23 | 2577.2746 | 1289.1409 | 2560.2480 | 1280.6277 | 2559.2640 | 1280.1356 | S | 450.2195 | 225.6134 | 433.1929 | 217.1001 | 432.2089 | 216.6081 | 4 |
| 24 | 2706.3172 | ***1353.6622*** | 2689.2906 | 1345.1490 | 2688.3066 | 1344.6569 | E | 363.1874 | 182.0974 | 346.1609 | 173.5841 | 345.1769 | 173.0921 | 3 |
| 25 | 2793.3492 | ***1397.1782*** | 2776.3227 | 1388.6650 | 2775.3386 | 1388.1730 | S | 234.1448 | 117.5761 | 217.1183 | 109.0628 | 216.1343 | 108.5708 | 2 |
| 26 | | | | | | | K | 147.1128 | 74.0600 | 130.0863 | 65.5468 | | | 1 |

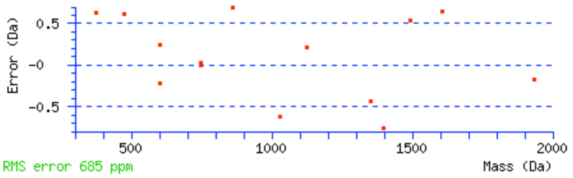


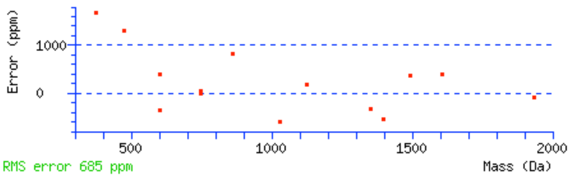


MS/MS Fragmentation of **AMNTGGFDSMQRQLEAELRSLSSESK**
Found in **P48563** in **Uniprot_yeastS288C**, Protein MON2 OS=Saccharomyces cerevisiae (strain ATCC 204508 / S288c) OX=559292 GN=MON2 PE=1 SV=1

Match to Query 16162: 2938.438254 from(980.486694,3+) intensity(637150.1875) rtinseconds(6766.0396) index(17031)
Title: 20100530_methyl400mM_1.18944.18944.3
Data file D:\Panyue\Proteomic dataset\20100530_methyl400mM_1.RAW

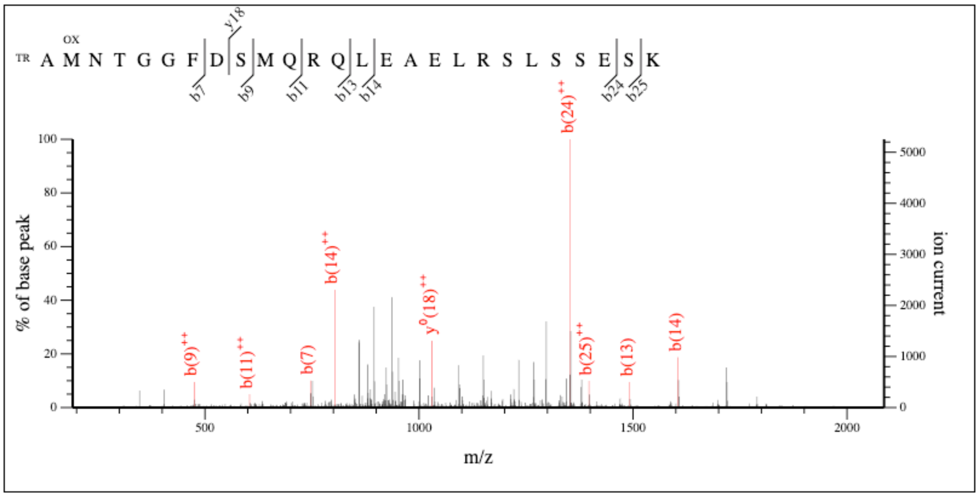


**Monoisotopic mass of neutral peptide Mr(calc):** 2938.4474
**Fixed modifications:** Carbamidomethyl (C) (apply to specified residues or termini only)
**Variable modifications:**
**N-term :** Trimethyl:2H(9) (Protein N-term)
**M2** : Oxidation (M), with neutral losses 0.0000(shown in table), 63.9983
**Ions Score:** 21 **Expect:** 0.088
**Matches :** 11/432 fragment ions using 17 most intense peaks (help)

| # | b | b++ | b* | b*++ | b0 | b0++ | Seq. | y | y++ | y* | y*++ | y0 | y0++ | # |
|---|---|---|---|---|---|---|---|---|---|---|---|---|---|---|
| 1 | 123.1478 | 62.0776 | | | | | A | | | | | | | 26 |
| 2 | 270.1832 | 135.5953 | | | | | M | 2817.3142 | 1409.1607 | 2800.2876 | 1400.6475 | 2799.3036 | 1400.1554 | 25 |
| 3 | 384.2262 | 192.6167 | 367.1996 | 184.1034 | | | N | 2670.2788 | 1335.6430 | 2653.2522 | 1327.1298 | 2652.2682 | 1326.6377 | 24 |
| 4 | 485.2738 | 243.1406 | 468.2473 | 234.6273 | 467.2633 | 234.1353 | T | 2556.2359 | 1278.6216 | 2539.2093 | 1270.1083 | 2538.2253 | 1269.6163 | 23 |
| 5 | 542.2953 | 271.6513 | 525.2688 | 263.1380 | 524.2847 | 262.6460 | G | 2455.1882 | 1228.0977 | 2438.1616 | 1219.5845 | 2437.1776 | 1219.0924 | 22 |
| 6 | 599.3168 | 300.1620 | 582.2902 | 291.6487 | 581.3062 | 291.1567 | G | 2398.1667 | 1199.5870 | 2381.1402 | 1191.0737 | 2380.1561 | 1190.5817 | 21 |
| 7 | **746.3852** | 373.6962 | 729.3586 | 365.1830 | 728.3746 | 364.6909 | F | 2341.1452 | 1171.0763 | 2324.1187 | 1162.5630 | 2323.1347 | 1162.0710 | 20 |
| 8 | 861.4121 | 431.2097 | 844.3856 | 422.6964 | 843.4016 | 422.2044 | D | 2194.0768 | 1097.5421 | 2177.0503 | 1089.0288 | 2176.0663 | 1088.5368 | 19 |
| 9 | 948.4441 | ***474.7257*** | 931.4176 | 466.2124 | 930.4336 | 465.7204 | S | 2079.0499 | 1040.0286 | 2062.0233 | 1031.5153 | 2061.0393 | 1031.0233 | 18 |
| 10 | 1079.4846 | 540.2460 | 1062.4581 | 531.7327 | 1061.4741 | 531.2407 | M | 1992.0179 | 996.5126 | 1974.9913 | 987.9993 | 1974.0073 | 987.5073 | 17 |
| 11 | 1207.5432 | ***604.2752*** | 1190.5167 | 595.7620 | 1189.5326 | 595.2700 | Q | 1860.9774 | 930.9923 | 1843.9508 | 922.4791 | 1842.9668 | 921.9870 | 16 |
| 12 | 1363.6443 | 682.3258 | 1346.6178 | 673.8125 | 1345.6338 | 673.3205 | R | 1732.9188 | 866.9630 | 1715.8923 | 858.4498 | 1714.9082 | 857.9578 | 15 |
| 13 | **1491.7029** | ***746.3551*** | 1474.6764 | 737.8418 | 1473.6923 | 737.3498 | Q | 1576.8177 | 788.9125 | 1559.7911 | 780.3992 | 1558.8071 | 779.9072 | 14 |
| 14 | **1604.7870** | ***802.8971*** | 1587.7604 | 794.3838 | 1586.7764 | 793.8918 | L | 1448.7591 | 724.8832 | 1431.7326 | 716.3699 | 1430.7485 | 715.8779 | 13 |
| 15 | 1733.8296 | 867.4184 | 1716.8030 | 858.9051 | 1715.8190 | 858.4131 | E | 1335.6750 | 668.3412 | 1318.6485 | 659.8279 | 1317.6645 | 659.3359 | 12 |
| 16 | 1804.8667 | 902.9370 | 1787.8401 | 894.4237 | 1786.8561 | 893.9317 | A | 1206.6325 | 603.8199 | 1189.6059 | 595.3066 | 1188.6219 | 594.8146 | 11 |
| 17 | 1933.9093 | 967.4583 | 1916.8827 | 958.9450 | 1915.8987 | 958.4530 | E | 1135.5953 | 568.3013 | 1118.5688 | 559.7880 | 1117.5848 | 559.2960 | 10 |
| 18 | 2046.9933 | 1024.0003 | 2029.9668 | 1015.4870 | 2028.9828 | 1014.9950 | L | 1006.5527 | 503.7800 | 989.5262 | 495.2667 | 988.5422 | 494.7747 | 9 |
| 19 | 2203.0944 | 1102.0509 | 2186.0679 | 1093.5376 | 2185.0839 | 1093.0456 | R | 893.4687 | 447.2380 | 876.4421 | 438.7247 | 875.4581 | 438.2327 | 8 |
| 20 | 2290.1265 | 1145.5669 | 2273.0999 | 1137.0536 | 2272.1159 | 1136.5616 | S | 737.3676 | 369.1874 | 720.3410 | 360.6742 | 719.3570 | 360.1821 | 7 |
| 21 | 2403.2105 | 1202.1089 | 2386.1840 | 1193.5956 | 2385.2000 | 1193.1036 | L | 650.3355 | 325.6714 | 633.3090 | 317.1581 | 632.3250 | 316.6661 | 6 |
| 22 | 2490.2426 | 1245.6249 | 2473.2160 | 1237.1116 | 2472.2320 | 1236.6196 | S | 537.2515 | 269.1294 | 520.2249 | 260.6161 | 519.2409 | 260.1241 | 5 |
| 23 | 2577.2746 | 1289.1409 | 2560.2480 | 1280.6277 | 2559.2640 | 1280.1356 | S | 450.2195 | 225.6134 | 433.1929 | 217.1001 | 432.2089 | 216.6081 | 4 |
| 24 | 2706.3172 | ***1353.6622*** | 2689.2906 | 1345.1490 | 2688.3066 | 1344.6569 | E | 363.1874 | 182.0974 | 346.1609 | 173.5841 | 345.1769 | 173.0921 | 3 |
| 25 | 2793.3492 | ***1397.1782*** | 2776.3227 | 1388.6650 | 2775.3386 | 1388.1730 | S | 234.1448 | 117.5761 | 217.1183 | 109.0628 | 216.1343 | 108.5708 | 2 |
| 26 | | | | | | | K | 147.1128 | 74.0600 | 130.0863 | 65.5468 | | | 1 |

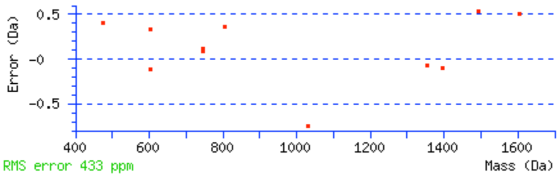


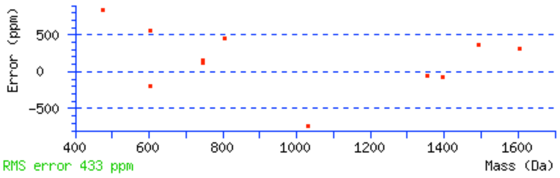

MS/MS Fragmentation of **AMNTGGFDSMQRQLEAELRSLSSESK**
Found in **P48563** in **Uniprot_yeastS288C**, Protein MON2 OS=Saccharomyces cerevisiae (strain ATCC 204508 / S288c) OX=559292 GN=MON2 PE=1 SV=1

Match to Query 16177: 2938.442466 from(980.488098,3+) intensity(154237.6406) rtinseconds(5697.0321) index(14436)
Title: 20100530_methyl400mM_1.16057.16057.3
Data file D:\Panyue\Proteomic dataset\20100530_methyl400mM_1.RAW

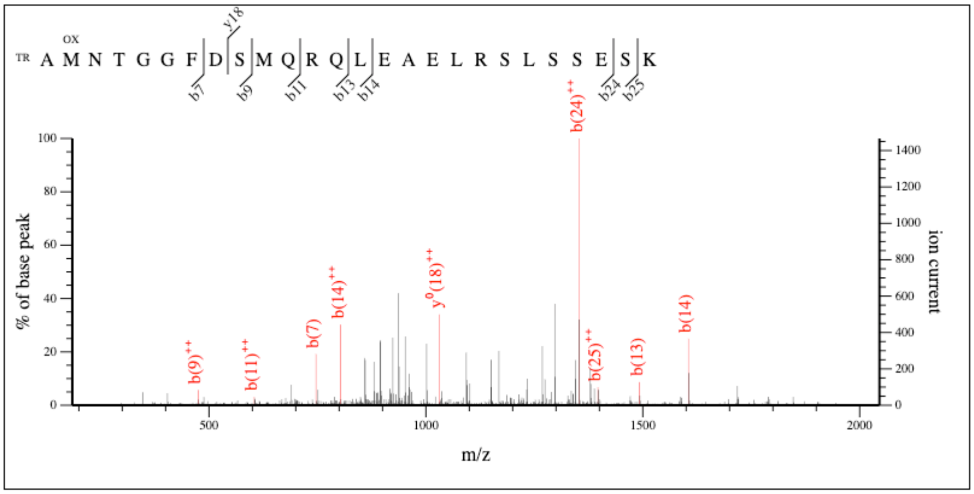


**Monoisotopic mass of neutral peptide Mr(calc):** 2938.4474
**Fixed modifications:** Carbamidomethyl (C) (apply to specified residues or termini only)
**Variable modifications:**
**N-term :** Trimethyl:2H(9) (Protein N-term)
**M2** : Oxidation (M), with neutral losses 0.0000(shown in table), 63.9983
**Ions Score:** 21 **Expect:** 0.094
**Matches :** 11/432 fragment ions using 17 most intense peaks (help)

| # | b | b++ | b* | b*++ | b0 | b0++ | Seq. | y | y++ | y* | y*++ | y0 | y0++ | # |
|---|---|---|---|---|---|---|---|---|---|---|---|---|---|---|
| 1 | 123.1478 | 62.0776 | | | | | A | | | | | | | 26 |
| 2 | 270.1832 | 135.5953 | | | | | M | 2817.3142 | 1409.1607 | 2800.2876 | 1400.6475 | 2799.3036 | 1400.1554 | 25 |
| 3 | 384.2262 | 192.6167 | 367.1996 | 184.1034 | | | N | 2670.2788 | 1335.6430 | 2653.2522 | 1327.1298 | 2652.2682 | 1326.6377 | 24 |
| 4 | 485.2738 | 243.1406 | 468.2473 | 234.6273 | 467.2633 | 234.1353 | T | 2556.2359 | 1278.6216 | 2539.2093 | 1270.1083 | 2538.2253 | 1269.6163 | 23 |
| 5 | 542.2953 | 271.6513 | 525.2688 | 263.1380 | 524.2847 | 262.6460 | G | 2455.1882 | 1228.0977 | 2438.1616 | 1219.5845 | 2437.1776 | 1219.0924 | 22 |
| 6 | 599.3168 | 300.1620 | 582.2902 | 291.6487 | 581.3062 | 291.1567 | G | 2398.1667 | 1199.5870 | 2381.1402 | 1191.0737 | 2380.1561 | 1190.5817 | 21 |
| 7 | **746.3852** | 373.6962 | 729.3586 | 365.1830 | 728.3746 | 364.6909 | F | 2341.1452 | 1171.0763 | 2324.1187 | 1162.5630 | 2323.1347 | 1162.0710 | 20 |
| 8 | 861.4121 | 431.2097 | 844.3856 | 422.6964 | 843.4016 | 422.2044 | D | 2194.0768 | 1097.5421 | 2177.0503 | 1089.0288 | 2176.0663 | 1088.5368 | 19 |
| 9 | 948.4441 | ***474.7257*** | 931.4176 | 466.2124 | 930.4336 | 465.7204 | S | 2079.0499 | 1040.0286 | 2062.0233 | 1031.5153 | 2061.0393 | 1031.0233 | 18 |
| 10 | 1079.4846 | 540.2460 | 1062.4581 | 531.7327 | 1061.4741 | 531.2407 | M | 1992.0179 | 996.5126 | 1974.9913 | 987.9993 | 1974.0073 | 987.5073 | 17 |
| 11 | 1207.5432 | ***604.2752*** | 1190.5167 | 595.7620 | 1189.5326 | 595.2700 | Q | 1860.9774 | 930.9923 | 1843.9508 | 922.4791 | 1842.9668 | 921.9870 | 16 |
| 12 | 1363.6443 | 682.3258 | 1346.6178 | 673.8125 | 1345.6338 | 673.3205 | R | 1732.9188 | 866.9630 | 1715.8923 | 858.4498 | 1714.9082 | 857.9578 | 15 |
| 13 | **1491.7029** | ***746.3551*** | 1474.6764 | 737.8418 | 1473.6923 | 737.3498 | Q | 1576.8177 | 788.9125 | 1559.7911 | 780.3992 | 1558.8071 | 779.9072 | 14 |
| 14 | **1604.7870** | ***802.8971*** | 1587.7604 | 794.3838 | 1586.7764 | 793.8918 | L | 1448.7591 | 724.8832 | 1431.7326 | 716.3699 | 1430.7485 | 715.8779 | 13 |
| 15 | 1733.8296 | 867.4184 | 1716.8030 | 858.9051 | 1715.8190 | 858.4131 | E | 1335.6750 | 668.3412 | 1318.6485 | 659.8279 | 1317.6645 | 659.3359 | 12 |
| 16 | 1804.8667 | 902.9370 | 1787.8401 | 894.4237 | 1786.8561 | 893.9317 | A | 1206.6325 | 603.8199 | 1189.6059 | 595.3066 | 1188.6219 | 594.8146 | 11 |
| 17 | 1933.9093 | 967.4583 | 1916.8827 | 958.9450 | 1915.8987 | 958.4530 | E | 1135.5953 | 568.3013 | 1118.5688 | 559.7880 | 1117.5848 | 559.2960 | 10 |
| 18 | 2046.9933 | 1024.0003 | 2029.9668 | 1015.4870 | 2028.9828 | 1014.9950 | L | 1006.5527 | 503.7800 | 989.5262 | 495.2667 | 988.5422 | 494.7747 | 9 |
| 19 | 2203.0944 | 1102.0509 | 2186.0679 | 1093.5376 | 2185.0839 | 1093.0456 | R | 893.4687 | 447.2380 | 876.4421 | 438.7247 | 875.4581 | 438.2327 | 8 |
| 20 | 2290.1265 | 1145.5669 | 2273.0999 | 1137.0536 | 2272.1159 | 1136.5616 | S | 737.3676 | 369.1874 | 720.3410 | 360.6742 | 719.3570 | 360.1821 | 7 |
| 21 | 2403.2105 | 1202.1089 | 2386.1840 | 1193.5956 | 2385.2000 | 1193.1036 | L | 650.3355 | 325.6714 | 633.3090 | 317.1581 | 632.3250 | 316.6661 | 6 |
| 22 | 2490.2426 | 1245.6249 | 2473.2160 | 1237.1116 | 2472.2320 | 1236.6196 | S | 537.2515 | 269.1294 | 520.2249 | 260.6161 | 519.2409 | 260.1241 | 5 |
| 23 | 2577.2746 | 1289.1409 | 2560.2480 | 1280.6277 | 2559.2640 | 1280.1356 | S | 450.2195 | 225.6134 | 433.1929 | 217.1001 | 432.2089 | 216.6081 | 4 |
| 24 | 2706.3172 | ***1353.6622*** | 2689.2906 | 1345.1490 | 2688.3066 | 1344.6569 | E | 363.1874 | 182.0974 | 346.1609 | 173.5841 | 345.1769 | 173.0921 | 3 |
| 25 | 2793.3492 | ***1397.1782*** | 2776.3227 | 1388.6650 | 2775.3386 | 1388.1730 | S | 234.1448 | 117.5761 | 217.1183 | 109.0628 | 216.1343 | 108.5708 | 2 |
| 26 | | | | | | | K | 147.1128 | 74.0600 | 130.0863 | 65.5468 | | | 1 |

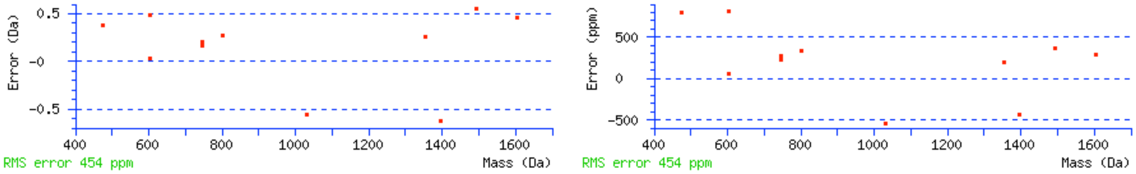


MS/MS Fragmentation of **AMNTGGFDSMQRQLEAELRSLSSESK**
Found in **P48563** in **Uniprot_yeastS288C**, Protein MON2 OS=Saccharomyces cerevisiae (strain ATCC 204508 / S288c) OX=559292 GN=MON2 PE=1 SV=1

Match to Query 16188: 2938.444113 from(980.488647,3+) intensity(179135.7812) rtinseconds(5701.282) index(14446)
Title: 20100530_methyl400mM_1.16068.16068.3
Data file D:\Panyue\Proteomic dataset\20100530_methyl400mM_1.RAW

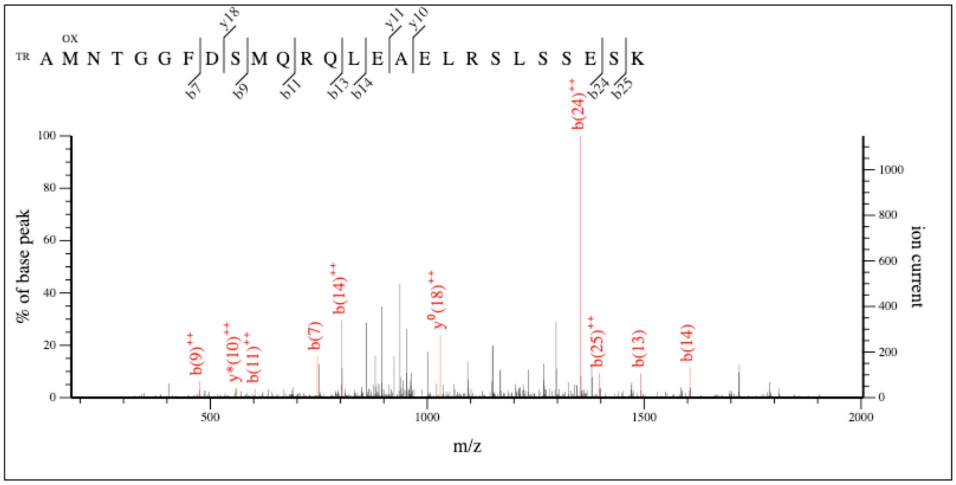


**Monoisotopic mass of neutral peptide Mr(calc):** 2938.4474
**Fixed modifications:** Carbamidomethyl (C) (apply to specified residues or termini only)
**Variable modifications:**
**N-term :** Trimethyl:2H(9) (Protein N-term)
**M2** : Oxidation (M), with neutral losses 0.0000(shown in table), 63.9983
**Ions Score:** 21 **Expect:** 0.094
**Matches :** 13/432 fragment ions using 17 most intense peaks (help)

| # | b | $b^{++}$ | $b^{*}$ | $b^{*++}$ | $b^{0}$ | $b^{0++}$ | Seq. | y | $y^{++}$ | $y^{*}$ | $y^{*++}$ | $y^{0}$ | $y^{0++}$ | # |
|---|---|---|---|---|---|---|---|---|---|---|---|---|---|---|
| 1 | 123.1478 | 62.0776 | | | | | A | | | | | | | 26 |
| 2 | 270.1832 | 135.5953 | | | | | M | 2817.3142 | 1409.1607 | 2800.2876 | 1400.6475 | 2799.3036 | 1400.1554 | 25 |
| 3 | 384.2262 | 192.6167 | 367.1996 | 184.1034 | | | N | 2670.2788 | 1335.6430 | 2653.2522 | 1327.1298 | 2652.2682 | 1326.6377 | 24 |
| 4 | 485.2738 | 243.1406 | 468.2473 | 234.6273 | 467.2633 | 234.1353 | T | 2556.2359 | 1278.6216 | 2539.2093 | 1270.1083 | 2538.2253 | 1269.6163 | 23 |
| 5 | 542.2953 | 271.6513 | 525.2688 | 263.1380 | 524.2847 | 262.6460 | G | 2455.1882 | 1228.0977 | 2438.1616 | 1219.5845 | 2437.1776 | 1219.0924 | 22 |
| 6 | 599.3168 | 300.1620 | 582.2902 | 291.6487 | 581.3062 | 291.1567 | G | 2398.1667 | 1199.5870 | 2381.1402 | 1191.0737 | 2380.1561 | 1190.5817 | 21 |
| 7 | **746.3852** | 373.6962 | 729.3586 | 365.1830 | 728.3746 | 364.6909 | F | 2341.1452 | 1171.0763 | 2324.1187 | 1162.5630 | 2323.1347 | 1162.0710 | 20 |
| 8 | 861.4121 | 431.2097 | 844.3856 | 422.6964 | 843.4016 | 422.2044 | D | 2194.0768 | 1097.5421 | 2177.0503 | 1089.0288 | 2176.0663 | 1088.5368 | 19 |
| 9 | 948.4441 | ***474.7257*** | 931.4176 | 466.2124 | 930.4336 | 465.7204 | S | 2079.0499 | 1040.0286 | 2062.0233 | 1031.5153 | 2061.0393 | 1031.0233 | 18 |
| 10 | 1079.4846 | 540.2460 | 1062.4581 | 531.7327 | 1061.4741 | 531.2407 | M | 1992.0179 | 996.5126 | 1974.9913 | 987.9993 | 1974.0073 | 987.5073 | 17 |
| 11 | 1207.5432 | ***604.2752*** | 1190.5167 | 595.7620 | 1189.5326 | 595.2700 | Q | 1860.9774 | 930.9923 | 1843.9508 | 922.4791 | 1842.9668 | 921.9870 | 16 |
| 12 | 1363.6443 | 682.3258 | 1346.6178 | 673.8125 | 1345.6338 | 673.3205 | R | 1732.9188 | 866.9630 | 1715.8923 | 858.4498 | 1714.9082 | 857.9578 | 15 |
| 13 | **1491.7029** | ***746.3551*** | 1474.6764 | 737.8418 | 1473.6923 | 737.3498 | Q | 1576.8177 | 788.9125 | 1559.7911 | 780.3992 | 1558.8071 | 779.9072 | 14 |
| 14 | **1604.7870** | ***802.8971*** | 1587.7604 | 794.3838 | 1586.7764 | 793.8918 | L | 1448.7591 | 724.8832 | 1431.7326 | 716.3699 | 1430.7485 | 715.8779 | 13 |
| 15 | 1733.8296 | 867.4184 | 1716.8030 | 858.9051 | 1715.8190 | 858.4131 | E | 1335.6750 | 668.3412 | 1318.6485 | 659.8279 | 1317.6645 | 659.3359 | 12 |
| 16 | 1804.8667 | 902.9370 | 1787.8401 | 894.4237 | 1786.8561 | 893.9317 | A | 1206.6325 | 603.8199 | 1189.6059 | 595.3066 | 1188.6219 | 594.8146 | 11 |
| 17 | 1933.9093 | 967.4583 | 1916.8827 | 958.9450 | 1915.8987 | 958.4530 | E | 1135.5953 | 568.3013 | 1118.5688 | 559.7880 | 1117.5848 | 559.2960 | 10 |
| 18 | 2046.9933 | 1024.0003 | 2029.9668 | 1015.4870 | 2028.9828 | 1014.9950 | L | 1006.5527 | 503.7800 | 989.5262 | 495.2667 | 988.5422 | 494.7747 | 9 |
| 19 | 2203.0944 | 1102.0509 | 2186.0679 | 1093.5376 | 2185.0839 | 1093.0456 | R | 893.4687 | 447.2380 | 876.4421 | 438.7247 | 875.4581 | 438.2327 | 8 |
| 20 | 2290.1265 | 1145.5669 | 2273.0999 | 1137.0536 | 2272.1159 | 1136.5616 | S | 737.3676 | 369.1874 | 720.3410 | 360.6742 | 719.3570 | 360.1821 | 7 |
| 21 | 2403.2105 | 1202.1089 | 2386.1840 | 1193.5956 | 2385.2000 | 1193.1036 | L | 650.3355 | 325.6714 | 633.3090 | 317.1581 | 632.3250 | 316.6661 | 6 |
| 22 | 2490.2426 | 1245.6249 | 2473.2160 | 1237.1116 | 2472.2320 | 1236.6196 | S | 537.2515 | 269.1294 | 520.2249 | 260.6161 | 519.2409 | 260.1241 | 5 |
| 23 | 2577.2746 | 1289.1409 | 2560.2480 | 1280.6277 | 2559.2640 | 1280.1356 | S | 450.2195 | 225.6134 | 433.1929 | 217.1001 | 432.2089 | 216.6081 | 4 |
| 24 | 2706.3172 | ***1353.6622*** | 2689.2906 | 1345.1490 | 2688.3066 | 1344.6569 | E | 363.1874 | 182.0974 | 346.1609 | 173.5841 | 345.1769 | 173.0921 | 3 |
| 25 | 2793.3492 | ***1397.1782*** | 2776.3227 | 1388.6650 | 2775.3386 | 1388.1730 | S | 234.1448 | 117.5761 | 217.1183 | 109.0628 | 216.1343 | 108.5708 | 2 |
| 26 | | | | | | | K | 147.1128 | 74.0600 | 130.0863 | 65.5468 | | | 1 |

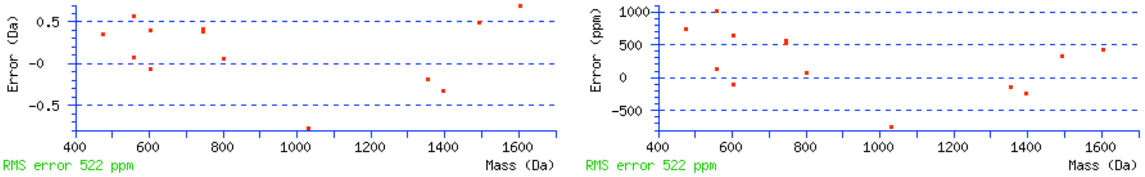


### 66. Pdc6 (3 peptides in 400mM fraction 1)

MS/MS Fragmentation of **SEITLGK**
Found in **P26263** in **Uniprot_yeastS288C**, Pyruvate decarboxylase isozyme 3 OS=Saccharomyces cerevisiae (strain ATCC 204508 / S288c) OX=559292 GN=PDC6 PE=1 SV=3

Match to Query 2214: 780.449389 from(781.456665,1+) intensity(13546679.0000) rtinseconds(1048.8241) index(2597)
Title: 20100530_methyl400mM_1.2890.2890.1
Data file D:\Panyue\Proteomic dataset\20100530_methyl400mM_1.RAW

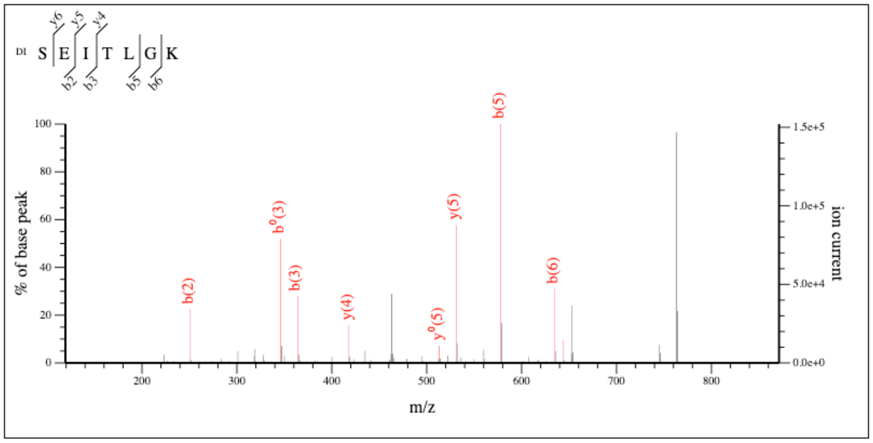

**Monoisotopic mass of neutral peptide Mr(calc):** 780.4864
**Fixed modifications:** Carbamidomethyl (C) (apply to specified residues or termini only)
**Variable modifications:**
**N-term :** Dimethyl:2H(6) (Protein N-term)
**Ions Score:** 27 **Expect:** 0.025
**Matches :** 9/27 fragment ions using 18 most intense peaks (help)

| # | b | $b^0$ | Seq. | y | y* | $y^0$ | # |
|---|---|---|---|---|---|---|---|
| 1 | 122.1083 | 104.0977 | S | | | | 7 |
| 2 | ***251.1509*** | 233.1403 | E | 660.3927 | 643.3661 | 642.3821 | 6 |
| 3 | ***364.2349*** | 346.2244 | I | **531.3501** | 514.3235 | 513.3395 | 5 |
| 4 | 465.2826 | 447.2720 | T | **418.2660** | 401.2395 | 400.2554 | 4 |
| 5 | ***578.3667*** | 560.3561 | L | 317.2183 | 300.1918 | | 3 |
| 6 | ***635.3881*** | 617.3776 | G | 204.1343 | 187.1077 | | 2 |
| 7 | | | K | 147.1128 | 130.0863 | | 1 |

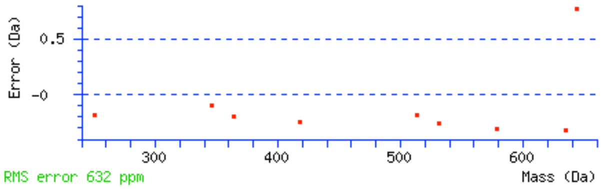


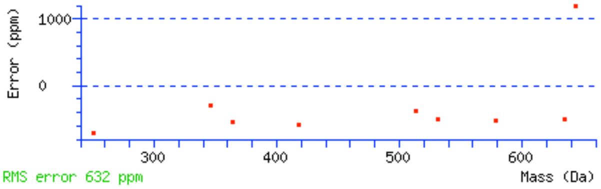


MS/MS Fragmentation of **SEITLGK**
Found in **P26263** in **Uniprot_yeastS288C**, Pyruvate decarboxylase isozyme 3 OS=Saccharomyces cerevisiae (strain ATCC 204508 / S288c) OX=559292 GN=PDC6 PE=1 SV=3

Match to Query 2215: 780.449755 from(781.457031,1+) intensity(16567665.0000) rtinseconds(1045.5965) index(2589)
Title: 20100530_methyl400mM_1.2881.2881.1
Data file D:\Panyue\Proteomic dataset\20100530_methyl400mM_1.RAW

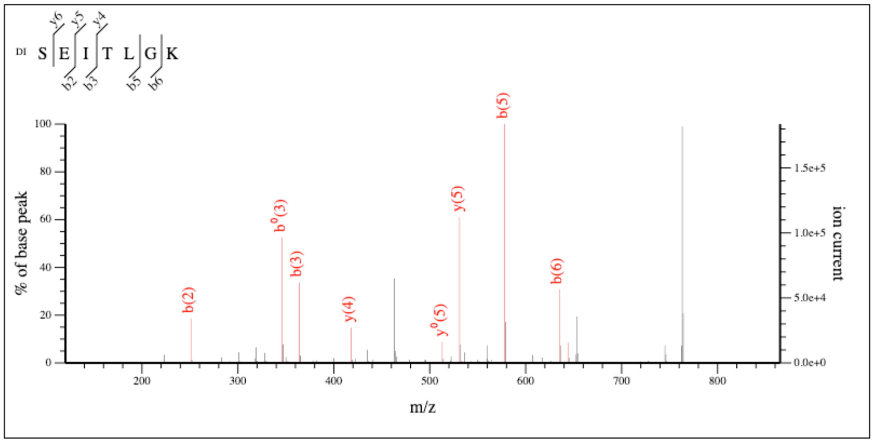

**Monoisotopic mass of neutral peptide Mr(calc):** 780.4864
**Fixed modifications:** Carbamidomethyl (C) (apply to specified residues or termini only)
**Variable modifications:**
**N-term :** Dimethyl:2H(6) (Protein N-term)
**Ions Score:** 27 **Expect:** 0.026
**Matches :** 9/27 fragment ions using 18 most intense peaks (help)

| # | b | $b^0$ | Seq. | y | y* | $y^0$ | # |
|---|---|---|---|---|---|---|---|
| 1 | 122.1083 | 104.0977 | S | | | | 7 |
| 2 | 251.1509 | 233.1403 | E | 660.3927 | 643.3661 | 642.3821 | 6 |
| 3 | 364.2349 | 346.2244 | I | 531.3501 | 514.3235 | 513.3395 | 5 |
| 4 | 465.2826 | 447.2720 | T | 418.2660 | 401.2395 | 400.2554 | 4 |
| 5 | 578.3667 | 560.3561 | L | 317.2183 | 300.1918 | | 3 |
| 6 | 635.3881 | 617.3776 | G | 204.1343 | 187.1077 | | 2 |
| 7 | | | K | 147.1128 | 130.0863 | | 1 |

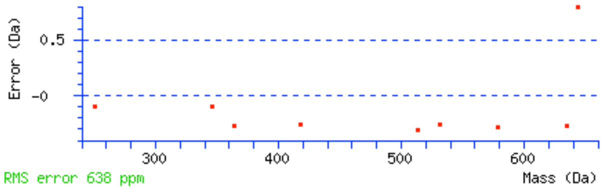


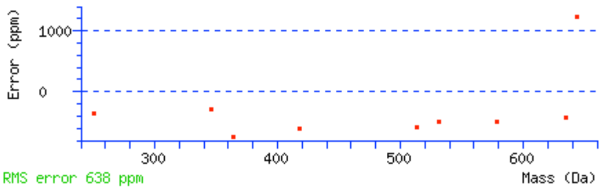


MS/MS Fragmentation of **SEITLGK**
Found in **P26263** in **Uniprot_yeastS288C**, Pyruvate decarboxylase isozyme 3 OS=Saccharomyces cerevisiae (strain ATCC 204508 / S288c) OX=559292 GN=PDC6 PE=1 SV=3

Match to Query 2217: 780.452724 from(781.460000,1+) intensity(65215.5391) rtinseconds(983.4678) index(2427)
Title: 20100530_methyl400mM_1.2700.2700.1
Data file D:\Panyue\Proteomic dataset\20100530_methyl400mM_1.RAW

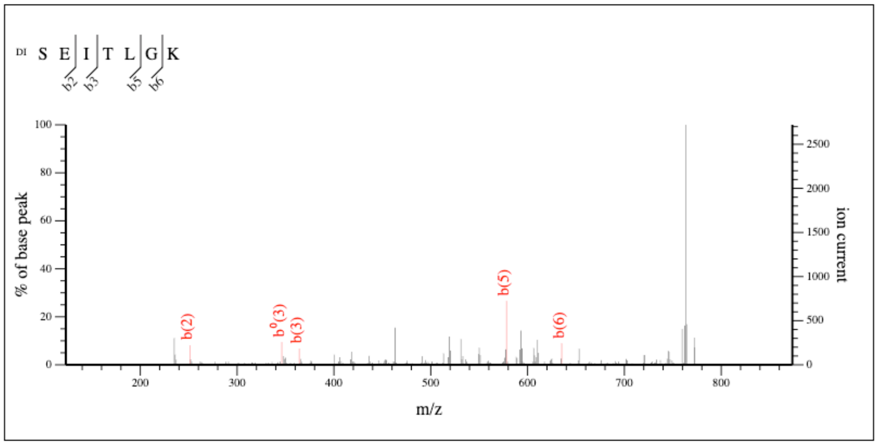

**Monoisotopic mass of neutral peptide Mr(calc):** 780.4864
**Fixed modifications:** Carbamidomethyl (C) (apply to specified residues or termini only)
**Variable modifications:**
**N-term :** Dimethyl:2H(6) (Protein N-term)
**Ions Score:** 24 **Expect:** 0.042
**Matches :** 5/27 fragment ions using 12 most intense peaks (help)

| # | b | $b^0$ | Seq. | y | y* | $y^0$ | # |
|---|---|---|---|---|---|---|---|
| 1 | 122.1083 | 104.0977 | S | | | | 7 |
| 2 | *251.1509* | 233.1403 | E | 660.3927 | 643.3661 | 642.3821 | 6 |
| 3 | *364.2349* | 346.2244 | I | 531.3501 | 514.3235 | 513.3395 | 5 |
| 4 | 465.2826 | 447.2720 | T | 418.2660 | 401.2395 | 400.2554 | 4 |
| 5 | *578.3667* | 560.3561 | L | 317.2183 | 300.1918 | | 3 |
| 6 | *635.3881* | 617.3776 | G | 204.1343 | 187.1077 | | 2 |
| 7 | | | K | 147.1128 | 130.0863 | | 1 |

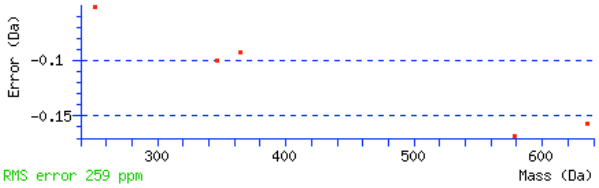


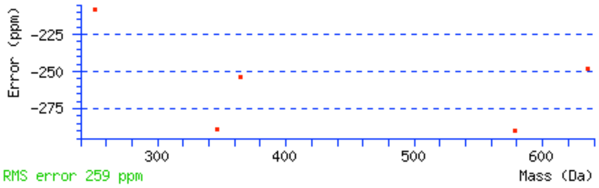


### 67. Pdc5 (3 peptides in 400mM fraction 1)

MS/MS Fragmentation of **SEITLGK**
Found in **P16467** in **Uniprot_yeastS288C**, Pyruvate decarboxylase isozyme 2 OS=Saccharomyces cerevisiae (strain ATCC 204508 / S288c) OX=559292 GN=PDC5 PE=1 SV=4

Match to Query 2214: 780.449389 from(781.456665,1+) intensity(13546679.0000) rtinseconds(1048.8241) index(2597)
Title: 20100530_methyl400mM_1.2890.2890.1
Data file D:\Panyue\Proteomic dataset\20100530_methyl400mM_1.RAW

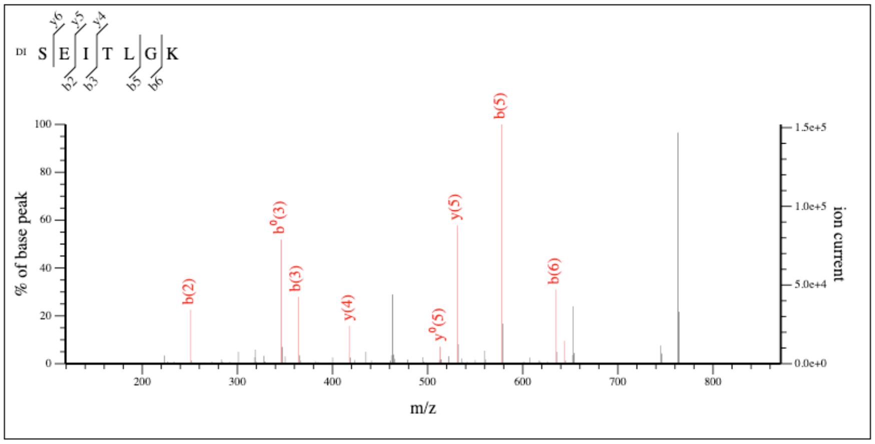

**Monoisotopic mass of neutral peptide Mr(calc):** 780.4864
**Fixed modifications:** Carbamidomethyl (C) (apply to specified residues or termini only)
**Variable modifications:**
**N-term :** Dimethyl:2H(6) (Protein N-term)
**Ions Score:** 27 **Expect:** 0.025
**Matches :** 9/27 fragment ions using 18 most intense peaks (help)

| # | b | b⁰ | Seq. | y | y* | y⁰ | # |
|---|---|---|---|---|---|---|---|
| 1 | 122.1083 | 104.0977 | S | | | | 7 |
| 2 | ***251.1509*** | 233.1403 | E | 660.3927 | 643.3661 | 642.3821 | 6 |
| 3 | ***364.2349*** | 346.2244 | I | **531.3501** | 514.3235 | 513.3395 | 5 |
| 4 | 465.2826 | 447.2720 | T | **418.2660** | 401.2395 | 400.2554 | 4 |
| 5 | ***578.3667*** | 560.3561 | L | 317.2183 | 300.1918 | | 3 |
| 6 | ***635.3881*** | 617.3776 | G | 204.1343 | 187.1077 | | 2 |
| 7 | | | K | 147.1128 | 130.0863 | | 1 |

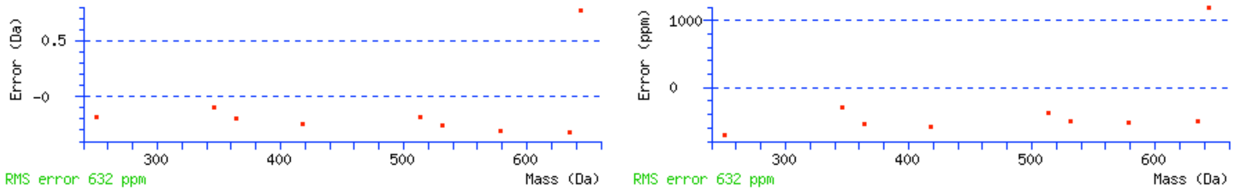


MS/MS Fragmentation of **SEITLGK**
Found in **P16467** in **Uniprot_yeastS288C**, Pyruvate decarboxylase isozyme 2 OS=Saccharomyces cerevisiae (strain ATCC 204508 / S288c) OX=559292 GN=PDC5 PE=1 SV=4

Match to Query 2215: 780.449755 from(781.457031,1+) intensity(16567665.0000) rtinseconds(1045.5965) index(2589)
Title: 20100530_methyl400mM_1.2881.2881.1
Data file D:\Panyue\Proteomic dataset\20100530_methyl400mM_1.RAW

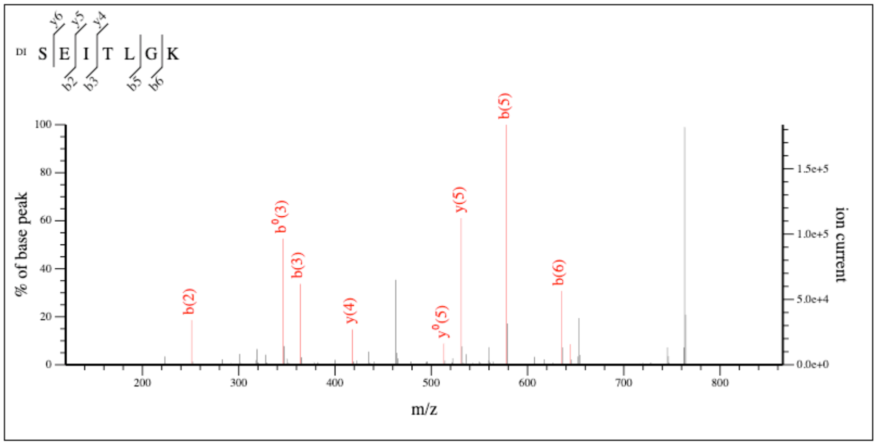

**Monoisotopic mass of neutral peptide Mr(calc):** 780.4864
**Fixed modifications:** Carbamidomethyl (C) (apply to specified residues or termini only)
**Variable modifications:**
**N-term :** Dimethyl:2H(6) (Protein N-term)
**Ions Score:** 27 **Expect:** 0.026
**Matches :** 9/27 fragment ions using 18 most intense peaks (help)

| # | b | $b^0$ | Seq. | y | y* | $y^0$ | # |
|---|---|---|---|---|---|---|---|
| 1 | 122.1083 | 104.0977 | S | | | | 7 |
| 2 | ***251.1509*** | 233.1403 | E | 660.3927 | 643.3661 | 642.3821 | 6 |
| 3 | ***364.2349*** | 346.2244 | I | **531.3501** | 514.3235 | 513.3395 | 5 |
| 4 | 465.2826 | 447.2720 | T | **418.2660** | 401.2395 | 400.2554 | 4 |
| 5 | ***578.3667*** | 560.3561 | L | 317.2183 | 300.1918 | | 3 |
| 6 | ***635.3881*** | 617.3776 | G | 204.1343 | 187.1077 | | 2 |
| 7 | | | K | 147.1128 | 130.0863 | | 1 |

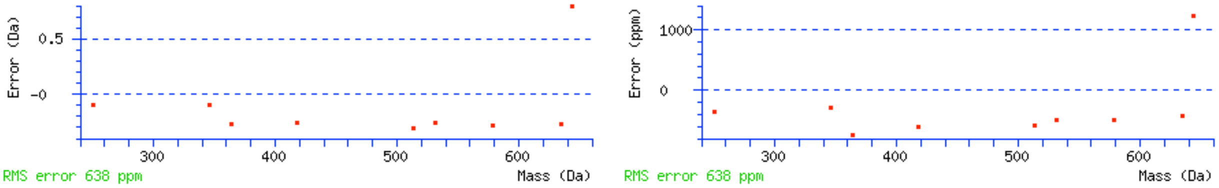


MS/MS Fragmentation of **SEITLGK**
Found in **P16467** in **Uniprot_yeastS288C**, Pyruvate decarboxylase isozyme 2 OS=Saccharomyces cerevisiae (strain ATCC 204508 / S288c) OX=559292 GN=PDC5 PE=1 SV=4

Match to Query 2217: 780.452724 from(781.460000,1+) intensity(65215.5391) rtinseconds(983.4678) index(2427)
Title: 20100530_methyl400mM_1.2700.2700.1
Data file D:\Panyue\Proteomic dataset\20100530_methyl400mM_1.RAW

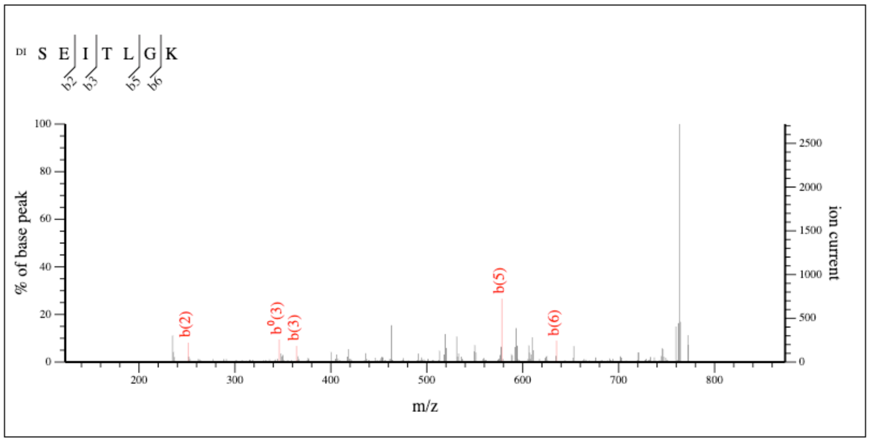

**Monoisotopic mass of neutral peptide Mr(calc):** 780.4864
**Fixed modifications:** Carbamidomethyl (C) (apply to specified residues or termini only)
**Variable modifications:**
**N-term :** Dimethyl:2H(6) (Protein N-term)
**Ions Score:** 24 **Expect:** 0.042
**Matches :** 5/27 fragment ions using 12 most intense peaks (help)

| # | b | b⁰ | Seq. | y | y* | y⁰ | # |
|---|---|---|---|---|---|---|---|
| 1 | 122.1083 | 104.0977 | S | | | | 7 |
| 2 | ***251.1509*** | 233.1403 | E | 660.3927 | 643.3661 | 642.3821 | 6 |
| 3 | ***364.2349*** | 346.2244 | I | 531.3501 | 514.3235 | 513.3395 | 5 |
| 4 | 465.2826 | 447.2720 | T | 418.2660 | 401.2395 | 400.2554 | 4 |
| 5 | ***578.3667*** | 560.3561 | L | 317.2183 | 300.1918 | | 3 |
| 6 | ***635.3881*** | 617.3776 | G | 204.1343 | 187.1077 | | 2 |
| 7 | | | K | 147.1128 | 130.0863 | | 1 |

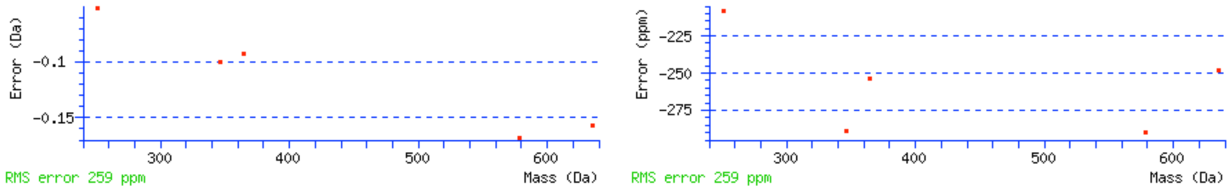


### 68. Pdc1 (3 peptides in 400mM fraction 1)

MS/MS Fragmentation of **SEITLGK**
Found in **P06169** in **Uniprot_yeastS288C**, Pyruvate decarboxylase isozyme 1 OS=Saccharomyces cerevisiae (strain ATCC 204508 / S288c) OX=559292 GN=PDC1 PE=1 SV=7

Match to Query 2214: 780.449389 from(781.456665,1+) intensity(13546679.0000) rtinseconds(1048.8241) index(2597)
Title: 20100530_methyl400mM_1.2890.2890.1
Data file D:\Panyue\Proteomic dataset\20100530_methyl400mM_1.RAW

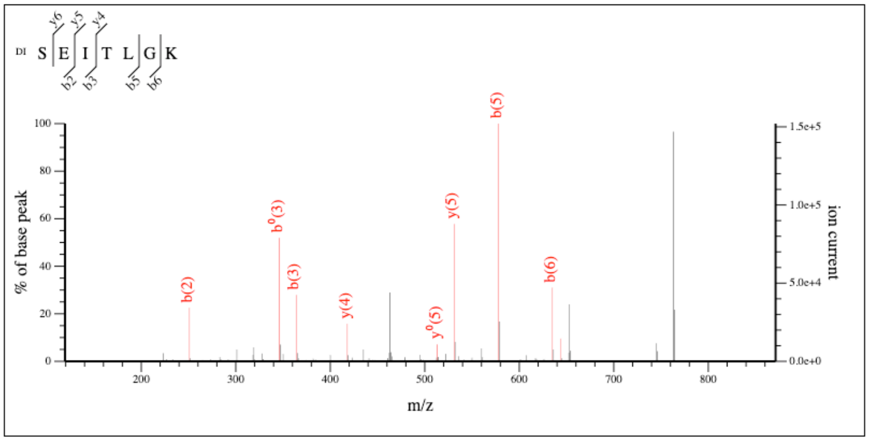

**Monoisotopic mass of neutral peptide Mr(calc):** 780.4864
**Fixed modifications:** Carbamidomethyl (C) (apply to specified residues or termini only)
**Variable modifications:**
**N-term :** Dimethyl:2H(6) (Protein N-term)
**Ions Score:** 27 **Expect:** 0.025
**Matches :** 9/27 fragment ions using 18 most intense peaks (help)

| # | b | $b^0$ | Seq. | y | y* | $y^0$ | # |
|---|---|---|---|---|---|---|---|
| 1 | 122.1083 | 104.0977 | S | | | | 7 |
| 2 | ***251.1509*** | 233.1403 | E | 660.3927 | 643.3661 | 642.3821 | 6 |
| 3 | ***364.2349*** | 346.2244 | I | **531.3501** | 514.3235 | 513.3395 | 5 |
| 4 | 465.2826 | 447.2720 | T | **418.2660** | 401.2395 | 400.2554 | 4 |
| 5 | ***578.3667*** | 560.3561 | L | 317.2183 | 300.1918 | | 3 |
| 6 | ***635.3881*** | 617.3776 | G | 204.1343 | 187.1077 | | 2 |
| 7 | | | K | 147.1128 | 130.0863 | | 1 |

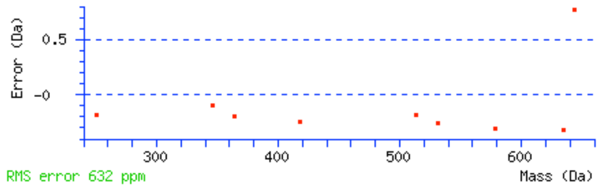


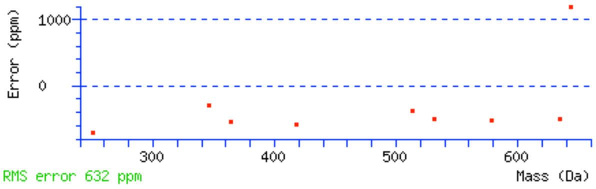


MS/MS Fragmentation of **SEITLGK**
Found in **P06169** in **Uniprot_yeastS288C**, Pyruvate decarboxylase isozyme 1 OS=Saccharomyces cerevisiae (strain ATCC 204508 / S288c) OX=559292 GN=PDC1 PE=1 SV=7

Match to Query 2215: 780.449755 from(781.457031,1+) intensity(16567665.0000) rtinseconds(1045.5965) index(2589)
Title: 20100530_methyl400mM_1.2881.2881.1
Data file D:\Panyue\Proteomic dataset\20100530_methyl400mM_1.RAW

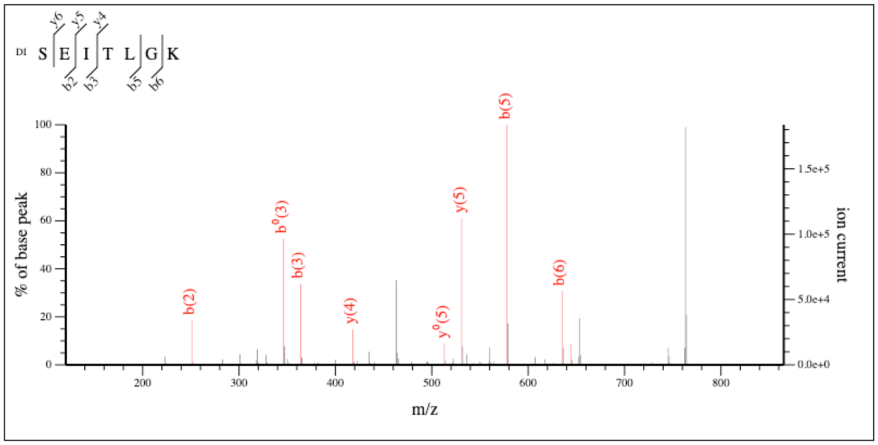

**Monoisotopic mass of neutral peptide Mr(calc):** 780.4864
**Fixed modifications:** Carbamidomethyl (C) (apply to specified residues or termini only)
**Variable modifications:**
**N-term :** Dimethyl:2H(6) (Protein N-term)
**Ions Score:** 27 **Expect:** 0.026
**Matches :** 9/27 fragment ions using 18 most intense peaks (help)

| # | b | $b^0$ | Seq. | y | y* | $y^0$ | # |
|---|---|---|---|---|---|---|---|
| 1 | 122.1083 | 104.0977 | S | | | | 7 |
| 2 | ***251.1509*** | 233.1403 | E | 660.3927 | 643.3661 | 642.3821 | 6 |
| 3 | ***364.2349*** | 346.2244 | I | **531.3501** | 514.3235 | 513.3395 | 5 |
| 4 | 465.2826 | 447.2720 | T | **418.2660** | 401.2395 | 400.2554 | 4 |
| 5 | ***578.3667*** | 560.3561 | L | 317.2183 | 300.1918 | | 3 |
| 6 | ***635.3881*** | 617.3776 | G | 204.1343 | 187.1077 | | 2 |
| 7 | | | K | 147.1128 | 130.0863 | | 1 |

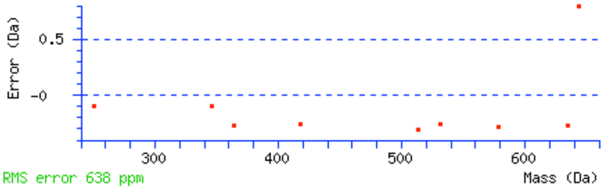


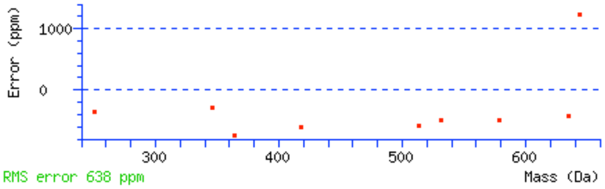


MS/MS Fragmentation of **SEITLGK**
Found in **P06169** in **Uniprot_yeastS288C**, Pyruvate decarboxylase isozyme 1 OS=Saccharomyces cerevisiae (strain ATCC 204508 / S288c) OX=559292 GN=PDC1 PE=1 SV=7

Match to Query 2217: 780.452724 from(781.460000,1+) intensity(65215.5391) rtinseconds(983.4678) index(2427)
Title: 20100530_methyl400mM_1.2700.2700.1
Data file D:\Panyue\Proteomic dataset\20100530_methyl400mM_1.RAW

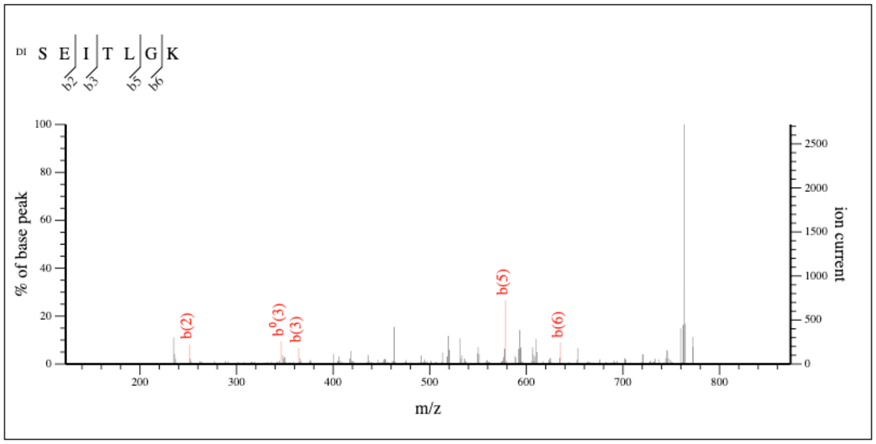

**Monoisotopic mass of neutral peptide Mr(calc):** 780.4864
**Fixed modifications:** Carbamidomethyl (C) (apply to specified residues or termini only)
**Variable modifications:**
**N-term :** Dimethyl:2H(6) (Protein N-term)
**Ions Score:** 24 **Expect:** 0.042
**Matches :** 5/27 fragment ions using 12 most intense peaks (help)

| # | b | $b^0$ | Seq. | y | y* | $y^0$ | # |
|---|---|---|---|---|---|---|---|
| 1 | 122.1083 | 104.0977 | S | | | | 7 |
| 2 | ***251.1509*** | 233.1403 | E | 660.3927 | 643.3661 | 642.3821 | 6 |
| 3 | ***364.2349*** | 346.2244 | I | 531.3501 | 514.3235 | 513.3395 | 5 |
| 4 | 465.2826 | 447.2720 | T | 418.2660 | 401.2395 | 400.2554 | 4 |
| 5 | ***578.3667*** | 560.3561 | L | 317.2183 | 300.1918 | | 3 |
| 6 | ***635.3881*** | 617.3776 | G | 204.1343 | 187.1077 | | 2 |
| 7 | | | K | 147.1128 | 130.0863 | | 1 |

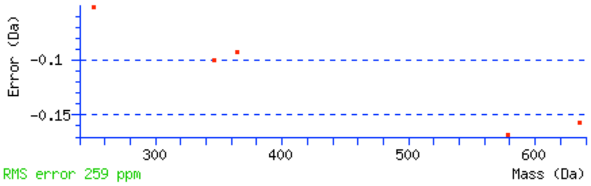


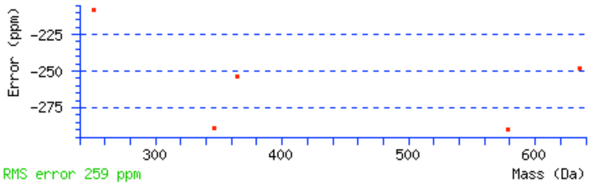


### 69. Fzo1 (2 peptides in 400mM fraction 1)

MS/MS Fragmentation of **SEGKQQFK**
Found in **P38297** in **Uniprot_yeastS288C**, Mitofusin FZO1 OS=Saccharomyces cerevisiae (strain ATCC 204508 / S288c) OX=559292 GN=FZO1 PE=1 SV=1

Match to Query 4361: 984.539220 from(493.276886,2+) intensity(382754.2500) rtinseconds(1624.5608) index(4125)
Title: 20100530_methyl400mM_1.4590.4590.2
Data file D:\Panyue\Proteomic dataset\20100530_methyl400mM_1.RAW

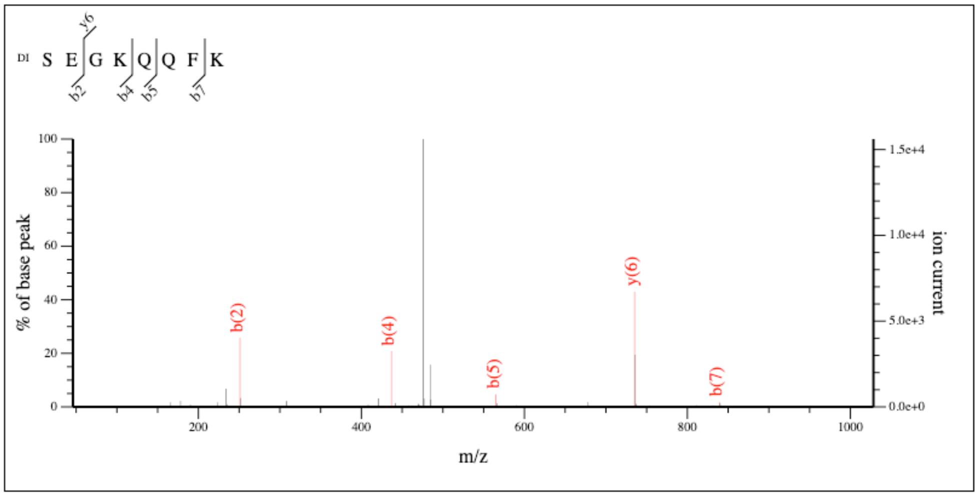

**Monoisotopic mass of neutral peptide Mr(calc):** 984.5511
**Fixed modifications:** Carbamidomethyl (C) (apply to specified residues or termini only)
**Variable modifications:**
**N-term :** Dimethyl:2H(6) (Protein N-term)
**Ions Score:** 29 **Expect:** 0.02
**Matches :** 5/66 fragment ions using 8 most intense peaks (help)

| # | b | b++ | b* | b*++ | b0 | b0++ | Seq. | y | y++ | y* | y*++ | y0 | y0++ | # |
|---|---|---|---|---|---|---|---|---|---|---|---|---|---|---|
| 1 | 122.1083 | 61.5578 | | | 104.0977 | 52.5525 | S | | | | | | | 8 |
| 2 | ***251.1509*** | 126.0791 | | | 233.1403 | 117.0738 | E | 864.4574 | 432.7323 | 847.4308 | 424.2191 | 846.4468 | 423.7271 | 7 |
| 3 | 308.1723 | 154.5898 | | | 290.1618 | 145.5845 | G | 735.4148 | 368.2110 | 718.3883 | 359.6978 | | | 6 |
| 4 | ***436.2673*** | 218.6373 | 419.2407 | 210.1240 | 418.2567 | 209.6320 | K | 678.3933 | 339.7003 | 661.3668 | 331.1870 | | | 5 |
| 5 | ***564.3259*** | 282.6666 | 547.2993 | 274.1533 | 546.3153 | 273.6613 | Q | 550.2984 | 275.6528 | 533.2718 | 267.1396 | | | 4 |
| 6 | 692.3844 | 346.6959 | 675.3579 | 338.1826 | 674.3739 | 337.6906 | Q | 422.2398 | 211.6235 | 405.2132 | 203.1103 | | | 3 |
| 7 | ***839.4529*** | 420.2301 | 822.4263 | 411.7168 | 821.4423 | 411.2248 | F | 294.1812 | 147.5942 | 277.1547 | 139.0810 | | | 2 |
| 8 | | | | | | | K | 147.1128 | 74.0600 | 130.0863 | 65.5468 | | | 1 |

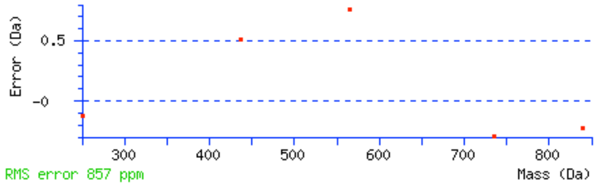


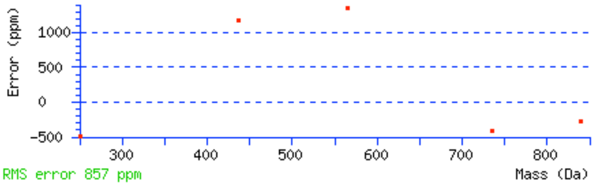


MS/MS Fragmentation of **SEGKQQFK**
Found in **P38297** in **Uniprot_yeastS288C**, Mitofusin FZO1 OS=Saccharomyces cerevisiae (strain ATCC 204508 / S288c) OX=559292 GN=FZO1 PE=1 SV=1

Match to Query 4362: 984.539526 from(493.277039,2+) intensity(585241.8125) rtinseconds(1626.2064) index(4129)
Title: 20100530_methyl400mM_1.4595.4595.2
Data file D:\Panyue\Proteomic dataset\20100530_methyl400mM_1.RAW

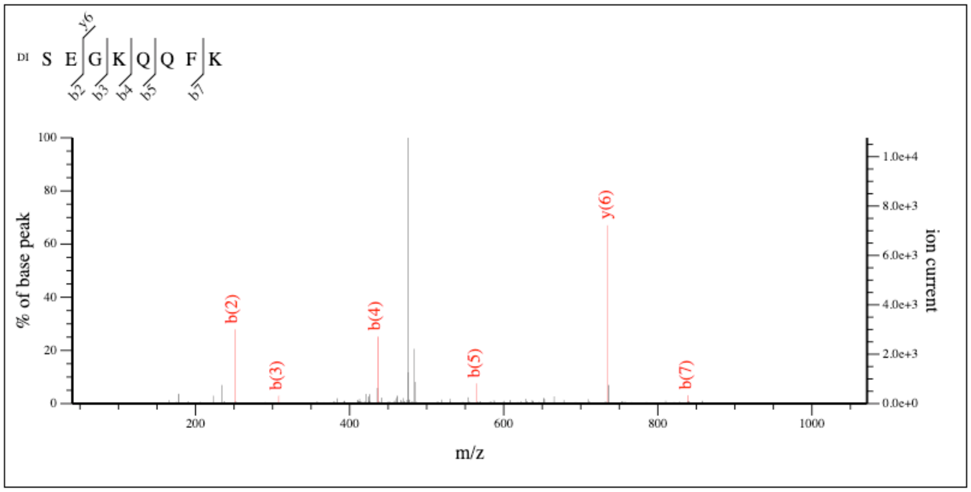

**Monoisotopic mass of neutral peptide Mr(calc):** 984.5511
**Fixed modifications:** Carbamidomethyl (C) (apply to specified residues or termini only)
**Variable modifications:**
**N-term :** Dimethyl:2H(6) (Protein N-term)
**Ions Score:** 28 **Expect:** 0.026
**Matches :** 6/66 fragment ions using 17 most intense peaks (help)

| # | b | b++ | b* | b*++ | b0 | b0++ | Seq. | y | y++ | y* | y*++ | y0 | y0++ | # |
|---|---|---|---|---|---|---|---|---|---|---|---|---|---|---|
| 1 | 122.1083 | 61.5578 | | | 104.0977 | 52.5525 | S | | | | | | | 8 |
| 2 | ***251.1509*** | 126.0791 | | | 233.1403 | 117.0738 | E | 864.4574 | 432.7323 | 847.4308 | 424.2191 | 846.4468 | 423.7271 | 7 |
| 3 | ***308.1723*** | 154.5898 | | | 290.1618 | 145.5845 | G | 735.4148 | 368.2110 | 718.3883 | 359.6978 | | | 6 |
| 4 | ***436.2673*** | 218.6373 | 419.2407 | 210.1240 | 418.2567 | 209.6320 | K | 678.3933 | 339.7003 | 661.3668 | 331.1870 | | | 5 |
| 5 | ***564.3259*** | 282.6666 | 547.2993 | 274.1533 | 546.3153 | 273.6613 | Q | 550.2984 | 275.6528 | 533.2718 | 267.1396 | | | 4 |
| 6 | 692.3844 | 346.6959 | 675.3579 | 338.1826 | 674.3739 | 337.6906 | Q | 422.2398 | 211.6235 | 405.2132 | 203.1103 | | | 3 |
| 7 | ***839.4529*** | 420.2301 | 822.4263 | 411.7168 | 821.4423 | 411.2248 | F | 294.1812 | 147.5942 | 277.1547 | 139.0810 | | | 2 |
| 8 | | | | | | | K | 147.1128 | 74.0600 | 130.0863 | 65.5468 | | | 1 |

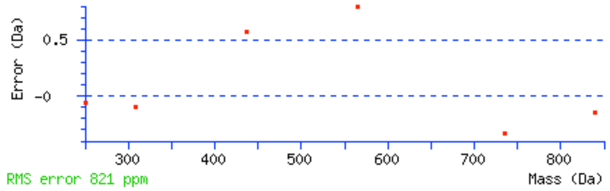


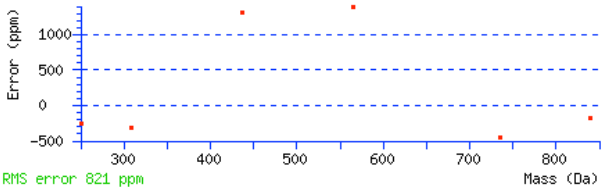


### 70. Eft1 (1 peptide in 400mM fraction 1)

MS/MS Fragmentation of **VAFTVDQMRSLMDK**
Found in **P32324** in **Uniprot_yeastS288C**, Elongation factor 2 OS=Saccharomyces cerevisiae (strain ATCC 204508 / S288c) OX=559292 GN=EFT1 PE=1 SV=1

Match to Query 9353: 1706.865448 from(854.440000,2+) intensity(1247188.1250) rtinseconds(2686.387) index(6972)
Title: 20100530_methyl400mM_1.7756.7756.2
Data file D:\Panyue\Proteomic dataset\20100530_methyl400mM_1.RAW

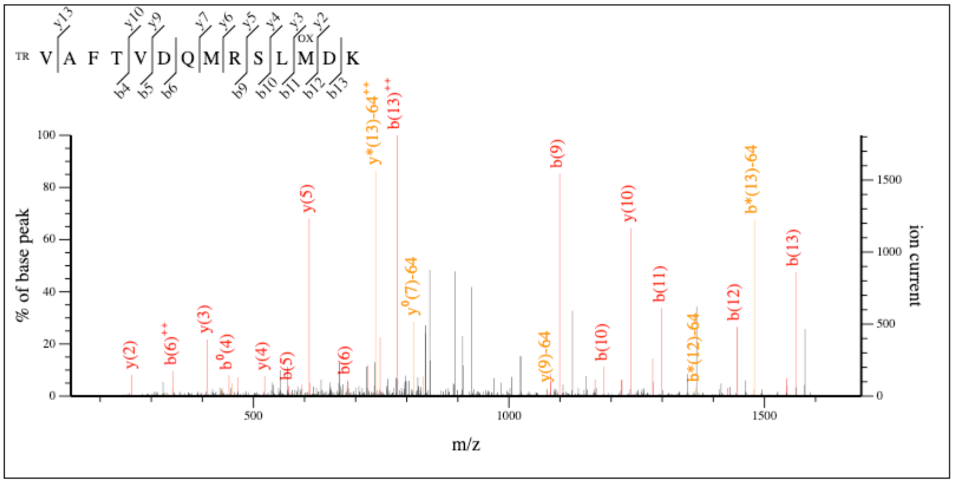

**Monoisotopic mass of neutral peptide Mr(calc):** 1706.9046
**Fixed modifications:** Carbamidomethyl (C) (apply to specified residues or termini only)
**Variable modifications:**
**N-term :** Trimethyl:2H(9) (Protein N-term)
**M12 :** Oxidation (M), with neutral losses 0.0000(shown in table), 63.9983
**Ions Score:** 25 **Expect:** 0.049
**Matches :** 52/214 fragment ions using 98 most intense peaks (help)

| # | b | b++ | b* | b*++ | b0 | b0++ | Seq. | y | y++ | y* | y*++ | y0 | y0++ | # |
|---|---|---|---|---|---|---|---|---|---|---|---|---|---|---|
| 1 | 151.1791 | 76.0932 | | | | | V | | | | | | | 14 |
| 2 | 222.2162 | 111.6118 | | | | | A | 1557.7400 | 779.3736 | 1540.7134 | 770.8604 | 1539.7294 | 770.3683 | 13 |
| 3 | 369.2847 | 185.1460 | | | | | F | 1486.7029 | 743.8551 | 1469.6763 | 735.3418 | 1468.6923 | 734.8498 | 12 |
| 4 | ***470.3323*** | 235.6698 | | | 452.3218 | 226.6645 | T | 1339.6344 | 670.3209 | 1322.6079 | 661.8076 | 1321.6239 | 661.3156 | 11 |
| 5 | ***569.4008*** | 285.2040 | | | 551.3902 | 276.1987 | V | **1238.5868** | 619.7970 | 1221.5602 | 611.2837 | 1220.5762 | 610.7917 | 10 |
| 6 | ***684.4277*** | **342.7175** | | | 666.4171 | 333.7122 | D | 1139.5184 | 570.2628 | 1122.4918 | 561.7495 | 1121.5078 | 561.2575 | 9 |
| 7 | 812.4863 | 406.7468 | 795.4597 | 398.2335 | 794.4757 | 397.7415 | Q | 1024.4914 | 512.7493 | 1007.4649 | 504.2361 | 1006.4808 | 503.7441 | 8 |
| 8 | 943.5268 | 472.2670 | 926.5002 | 463.7537 | 925.5162 | 463.2617 | M | 896.4328 | 448.7201 | 879.4063 | 440.2068 | 878.4223 | 439.7148 | 7 |
| 9 | ***1099.6279*** | 550.3176 | 1082.6013 | 541.8043 | 1081.6173 | 541.3123 | R | 765.3924 | 383.1998 | 748.3658 | 374.6865 | 747.3818 | 374.1945 | 6 |
| 10 | ***1186.6599*** | **593.8336** | 1169.6333 | 585.3203 | 1168.6493 | 584.8283 | S | **609.2912** | 305.1493 | 592.2647 | 296.6360 | 591.2807 | 296.1440 | 5 |
| 11 | ***1299.7440*** | 650.3756 | 1282.7174 | 641.8623 | 1281.7334 | 641.3703 | L | **522.2592** | 261.6332 | 505.2327 | 253.1200 | 504.2486 | 252.6280 | 4 |
| 12 | ***1446.7794*** | **723.8933** | 1429.7528 | 715.3800 | 1428.7688 | 714.8880 | M | **409.1751** | 205.0912 | 392.1486 | 196.5779 | 391.1646 | 196.0859 | 3 |
| 13 | ***1561.8063*** | **781.4068** | 1544.7798 | 772.8935 | 1543.7957 | 772.4015 | D | **262.1397** | 131.5735 | 245.1132 | 123.0602 | 244.1292 | 122.5682 | 2 |
| 14 | | | | | | | K | 147.1128 | 74.0600 | 130.0863 | 65.5468 | | | 1 |

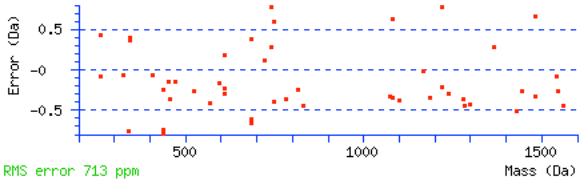


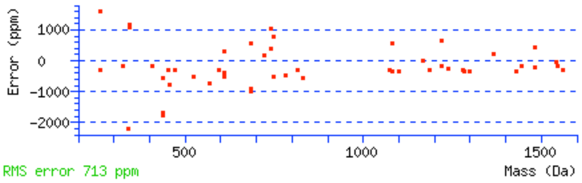


### 71. Pdc6 (2 peptides in 400mM fraction 2)

MS/MS Fragmentation of **SEITLGK**
Found in **P26263** in **Uniprot_yeastS288C**, Pyruvate decarboxylase isozyme 3 OS=Saccharomyces cerevisiae (strain ATCC 204508 / S288c) OX=559292 GN=PDC6 PE=1 SV=3

Match to Query 2320: 780.449877 from(781.457153,1+) intensity(6784650.5000) rtinseconds(1067.5029) index(2650)
Title: 20100530_methyl400mM_2.2950.2950.1
Data file D:\Panyue\Proteomic dataset\20100530_methyl400mM_2.RAW

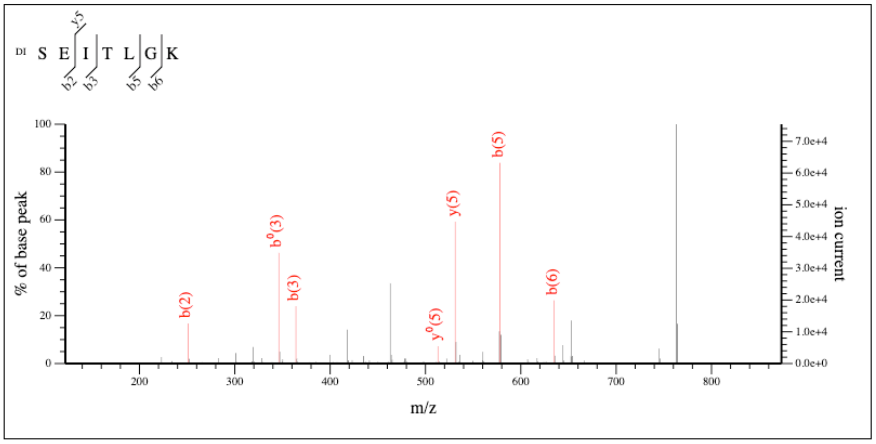

**Monoisotopic mass of neutral peptide Mr(calc):** 780.4864
**Fixed modifications:** Carbamidomethyl (C) (apply to specified residues or termini only)
**Variable modifications:**
**N-term :** Dimethyl:2H(6) (Protein N-term)
**Ions Score:** 27 **Expect:** 0.028
**Matches :** 7/27 fragment ions using 12 most intense peaks (help)

| # | b | $b^0$ | Seq. | y | y* | $y^0$ | # |
|---|---|---|---|---|---|---|---|
| 1 | 122.1083 | 104.0977 | S | | | | 7 |
| 2 | ***251.1509*** | 233.1403 | E | 660.3927 | 643.3661 | 642.3821 | 6 |
| 3 | ***364.2349*** | 346.2244 | I | 531.3501 | 514.3235 | 513.3395 | 5 |
| 4 | 465.2826 | 447.2720 | T | 418.2660 | 401.2395 | 400.2554 | 4 |
| 5 | ***578.3667*** | 560.3561 | L | 317.2183 | 300.1918 | | 3 |
| 6 | ***635.3881*** | 617.3776 | G | 204.1343 | 187.1077 | | 2 |
| 7 | | | K | 147.1128 | 130.0863 | | 1 |

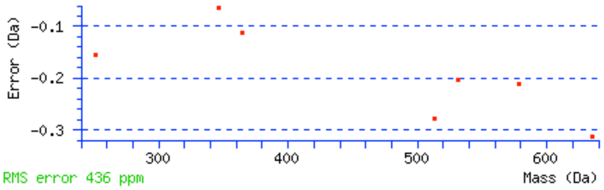


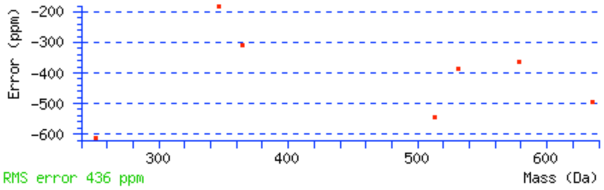


MS/MS Fragmentation of **SEITLGK**
Found in **P26263** in **Uniprot_yeastS288C**, Pyruvate decarboxylase isozyme 3 OS=Saccharomyces cerevisiae (strain ATCC 204508 / S288c) OX=559292 GN=PDC6 PE=1 SV=3

Match to Query 2321: 780.450244 from(781.457520,1+) intensity(10227573.0000) rtinseconds(1064.2357) index(2642)
Title: 20100530_methyl400mM_2.2941.2941.1
Data file D:\Panyue\Proteomic dataset\20100530_methyl400mM_2.RAW

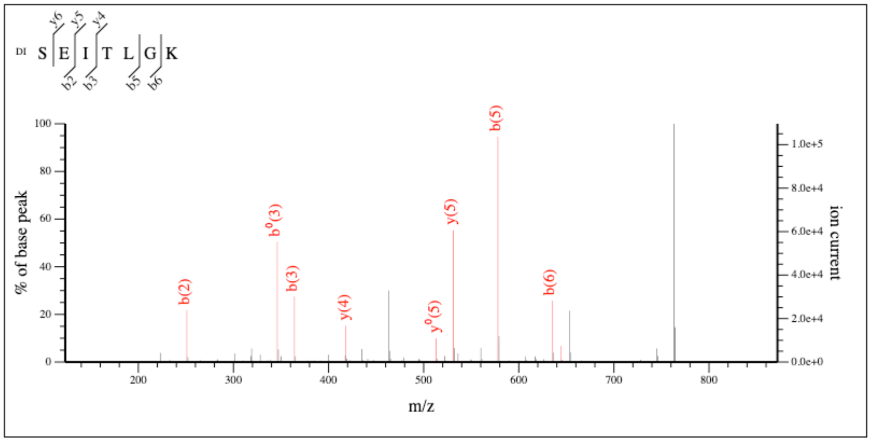

**Monoisotopic mass of neutral peptide Mr(calc):** 780.4864
**Fixed modifications:** Carbamidomethyl (C) (apply to specified residues or termini only)
**Variable modifications:**
**N-term :** Dimethyl:2H(6) (Protein N-term)
**Ions Score:** 25 **Expect:** 0.037
**Matches :** 9/27 fragment ions using 18 most intense peaks (help)

| # | b | $b^0$ | Seq. | y | y* | $y^0$ | # |
|---|---|---|---|---|---|---|---|
| 1 | 122.1083 | 104.0977 | S | | | | 7 |
| 2 | 251.1509 | 233.1403 | E | 660.3927 | 643.3661 | 642.3821 | 6 |
| 3 | 364.2349 | 346.2244 | I | 531.3501 | 514.3235 | 513.3395 | 5 |
| 4 | 465.2826 | 447.2720 | T | 418.2660 | 401.2395 | 400.2554 | 4 |
| 5 | 578.3667 | 560.3561 | L | 317.2183 | 300.1918 | | 3 |
| 6 | 635.3881 | 617.3776 | G | 204.1343 | 187.1077 | | 2 |
| 7 | | | K | 147.1128 | 130.0863 | | 1 |

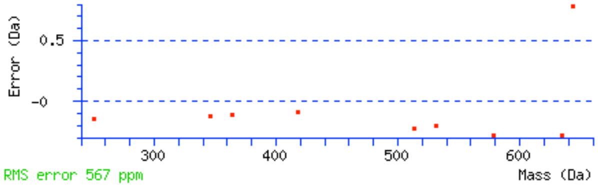


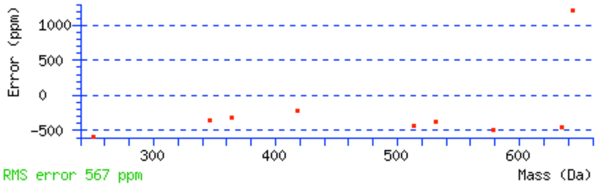


### 72. Pdc5 (2 peptides in 400mM fraction 2)

MS/MS Fragmentation of **SEITLGK**
Found in **P16467** in **Uniprot_yeastS288C**, Pyruvate decarboxylase isozyme 2 OS=Saccharomyces cerevisiae (strain ATCC 204508 / S288c) OX=559292 GN=PDC5 PE=1 SV=4

Match to Query 2320: 780.449877 from(781.457153,1+) intensity(6784650.5000) rtinseconds(1067.5029) index(2650)
Title: 20100530_methyl400mM_2.2950.2950.1
Data file D:\Panyue\Proteomic dataset\20100530_methyl400mM_2.RAW

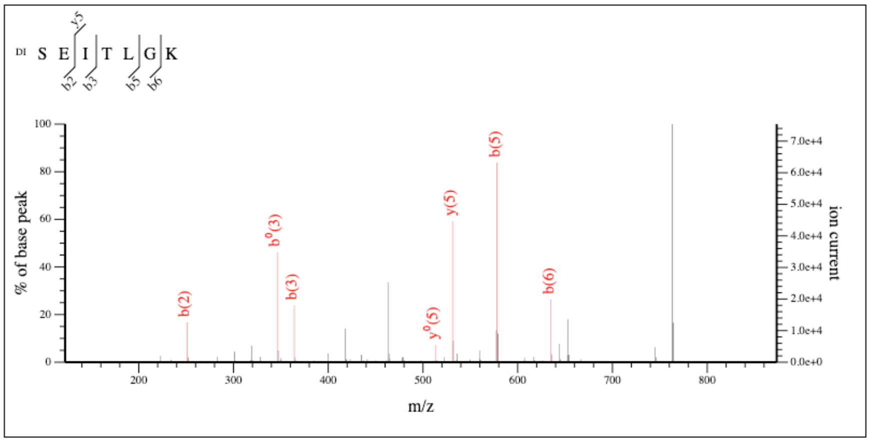

**Monoisotopic mass of neutral peptide Mr(calc):** 780.4864
**Fixed modifications:** Carbamidomethyl (C) (apply to specified residues or termini only)
**Variable modifications:**
**N-term :** Dimethyl:2H(6) (Protein N-term)
**Ions Score:** 27 **Expect:** 0.028
**Matches :** 7/27 fragment ions using 12 most intense peaks (help)

| # | b | $b^0$ | Seq. | y | y* | $y^0$ | # |
|---|---|---|---|---|---|---|---|
| 1 | 122.1083 | 104.0977 | S | | | | 7 |
| 2 | ***251.1509*** | 233.1403 | E | 660.3927 | 643.3661 | 642.3821 | 6 |
| 3 | ***364.2349*** | 346.2244 | I | 531.3501 | 514.3235 | 513.3395 | 5 |
| 4 | 465.2826 | 447.2720 | T | 418.2660 | 401.2395 | 400.2554 | 4 |
| 5 | ***578.3667*** | 560.3561 | L | 317.2183 | 300.1918 | | 3 |
| 6 | ***635.3881*** | 617.3776 | G | 204.1343 | 187.1077 | | 2 |
| 7 | | | K | 147.1128 | 130.0863 | | 1 |

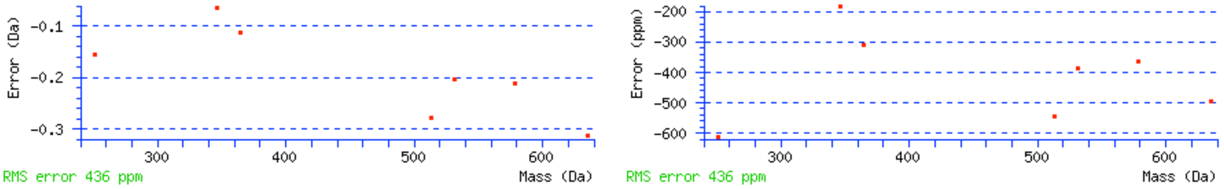


MS/MS Fragmentation of **SEITLGK**
Found in **P16467** in **Uniprot_yeastS288C**, Pyruvate decarboxylase isozyme 2 OS=Saccharomyces cerevisiae (strain ATCC 204508 / S288c) OX=559292 GN=PDC5 PE=1 SV=4

Match to Query 2321: 780.450244 from(781.457520,1+) intensity(10227573.0000) rtinseconds(1064.2357) index(2642)
Title: 20100530_methyl400mM_2.2941.2941.1
Data file D:\Panyue\Proteomic dataset\20100530_methyl400mM_2.RAW

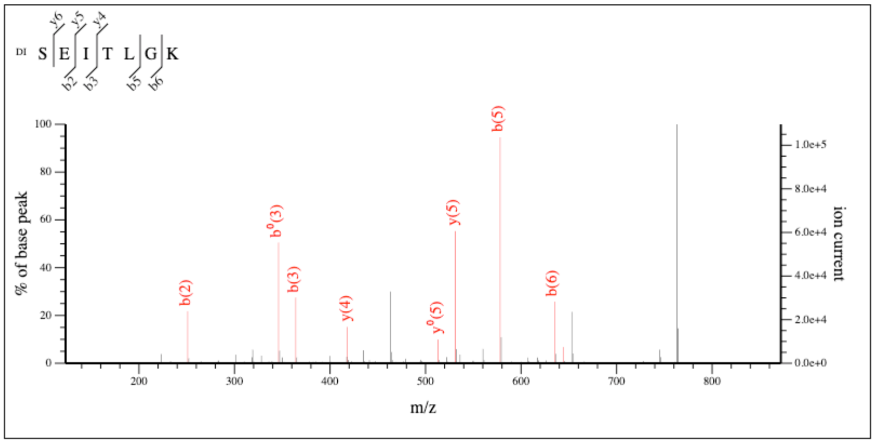

**Monoisotopic mass of neutral peptide Mr(calc):** 780.4864
**Fixed modifications:** Carbamidomethyl (C) (apply to specified residues or termini only)
**Variable modifications:**
**N-term :** Dimethyl:2H(6) (Protein N-term)
**Ions Score:** 25 **Expect:** 0.037
**Matches :** 9/27 fragment ions using 18 most intense peaks (help)

| # | b | b0 | Seq. | y | y* | y0 | # |
|---|---|---|---|---|---|---|---|
| 1 | 122.1083 | 104.0977 | S | | | | 7 |
| 2 | ***251.1509*** | 233.1403 | E | 660.3927 | 643.3661 | 642.3821 | 6 |
| 3 | ***364.2349*** | 346.2244 | I | **531.3501** | 514.3235 | 513.3395 | 5 |
| 4 | 465.2826 | 447.2720 | T | **418.2660** | 401.2395 | 400.2554 | 4 |
| 5 | ***578.3667*** | 560.3561 | L | 317.2183 | 300.1918 | | 3 |
| 6 | ***635.3881*** | 617.3776 | G | 204.1343 | 187.1077 | | 2 |
| 7 | | | K | 147.1128 | 130.0863 | | 1 |

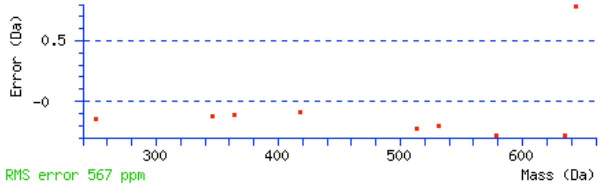


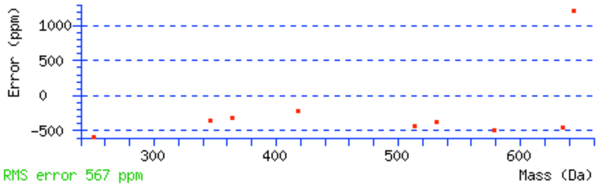


### 73. Pdc1 (2 peptides in 400mM fraction 2)

MS/MS Fragmentation of **SEITLGK**
Found in **P06169** in **Uniprot_yeastS288C**, Pyruvate decarboxylase isozyme 1 OS=Saccharomyces cerevisiae (strain ATCC 204508 / S288c) OX=559292 GN=PDC1 PE=1 SV=7

Match to Query 2321: 780.450244 from(781.457520,1+) intensity(10227573.0000) rtinseconds(1064.2357) index(2642)
Title: 20100530_methyl400mM_2.2941.2941.1
Data file D:\Panyue\Proteomic dataset\20100530_methyl400mM_2.RAW

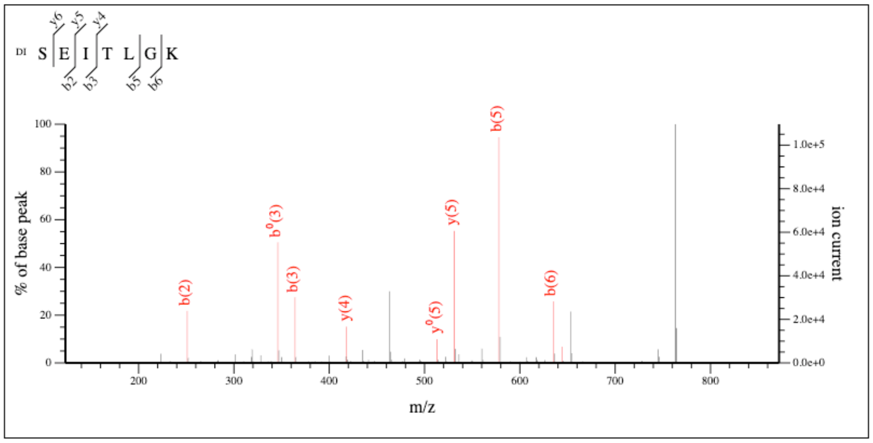

**Monoisotopic mass of neutral peptide Mr(calc):** 780.4864
**Fixed modifications:** Carbamidomethyl (C) (apply to specified residues or termini only)
**Variable modifications:**
**N-term :** Dimethyl:2H(6) (Protein N-term)
**Ions Score:** 25 **Expect:** 0.037
**Matches :** 9/27 fragment ions using 18 most intense peaks (help)

| # | b | $b^0$ | Seq. | y | y* | $y^0$ | # |
|---|---|---|---|---|---|---|---|
| 1 | 122.1083 | 104.0977 | S | | | | 7 |
| 2 | ***251.1509*** | 233.1403 | E | 660.3927 | 643.3661 | 642.3821 | 6 |
| 3 | ***364.2349*** | 346.2244 | I | **531.3501** | 514.3235 | 513.3395 | 5 |
| 4 | 465.2826 | 447.2720 | T | **418.2660** | 401.2395 | 400.2554 | 4 |
| 5 | ***578.3667*** | 560.3561 | L | 317.2183 | 300.1918 | | 3 |
| 6 | ***635.3881*** | 617.3776 | G | 204.1343 | 187.1077 | | 2 |
| 7 | | | K | 147.1128 | 130.0863 | | 1 |

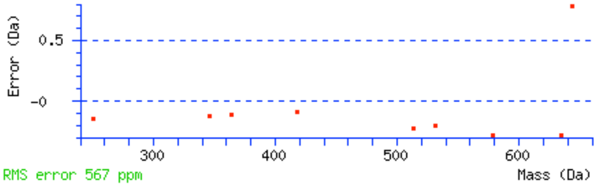


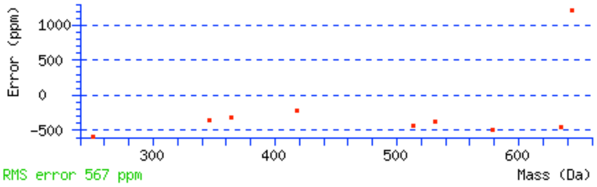


MS/MS Fragmentation of **SEITLGK**
Found in **P06169** in **Uniprot_yeastS288C**, Pyruvate decarboxylase isozyme 1 OS=Saccharomyces cerevisiae (strain ATCC 204508 / S288c) OX=559292 GN=PDC1 PE=1 SV=7

Match to Query 2320: 780.449877 from(781.457153,1+) intensity(6784650.5000) rtinseconds(1067.5029) index(2650)
Title: 20100530_methyl400mM_2.2950.2950.1
Data file D:\Panyue\Proteomic dataset\20100530_methyl400mM_2.RAW

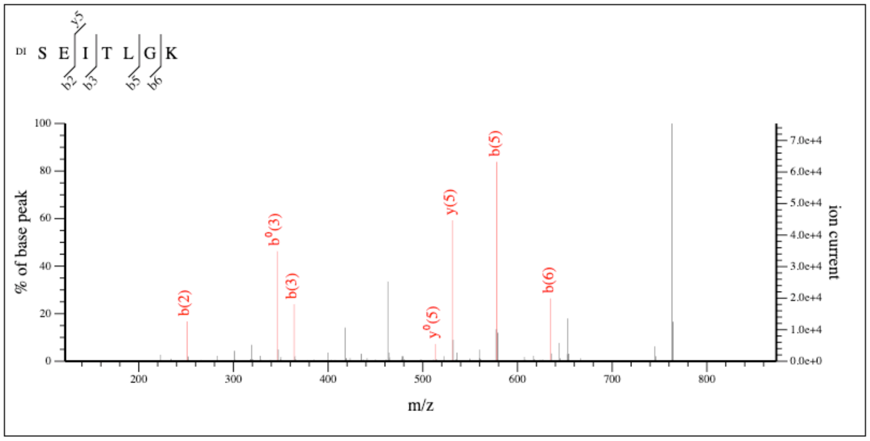

**Monoisotopic mass of neutral peptide Mr(calc):** 780.4864
**Fixed modifications:** Carbamidomethyl (C) (apply to specified residues or termini only)
**Variable modifications:**
**N-term :** Dimethyl:2H(6) (Protein N-term)
**Ions Score:** 27 **Expect:** 0.028
**Matches :** 7/27 fragment ions using 12 most intense peaks (help)

| # | b | $b^0$ | Seq. | y | y* | $y^0$ | # |
|---|---|---|---|---|---|---|---|
| 1 | 122.1083 | 104.0977 | S | | | | 7 |
| 2 | ***251.1509*** | 233.1403 | E | 660.3927 | 643.3661 | 642.3821 | 6 |
| 3 | ***364.2349*** | 346.2244 | I | 531.3501 | 514.3235 | 513.3395 | 5 |
| 4 | 465.2826 | 447.2720 | T | 418.2660 | 401.2395 | 400.2554 | 4 |
| 5 | ***578.3667*** | 560.3561 | L | 317.2183 | 300.1918 | | 3 |
| 6 | ***635.3881*** | 617.3776 | G | 204.1343 | 187.1077 | | 2 |
| 7 | | | K | 147.1128 | 130.0863 | | 1 |

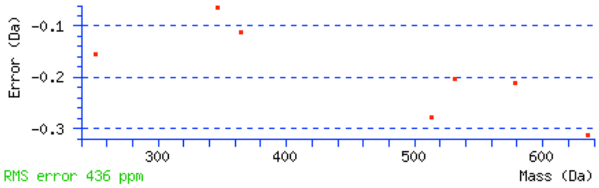


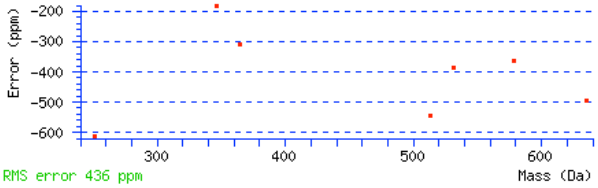


### 74. Eaf6 (1 peptide in 400mM fraction2)

MS/MS Fragmentation of **TDELKSYEALK**
Found in **P47128** in **Uniprot_yeastS288C**, Chromatin modification-related protein EAF6 OS=Saccharomyces cerevisiae (strain ATCC 204508 / S288c) OX=559292 GN=EAF6 PE=1 SV=1

Match to Query 6963: 1312.662206 from(657.338379,2+) intensity(345217.1875) rtinseconds(1183.9777) index(2955)
Title: 20100530_methyl400mM_2.3289.3289.2
Data file D:\Panyue\Proteomic dataset\20100530_methyl400mM_2.RAW

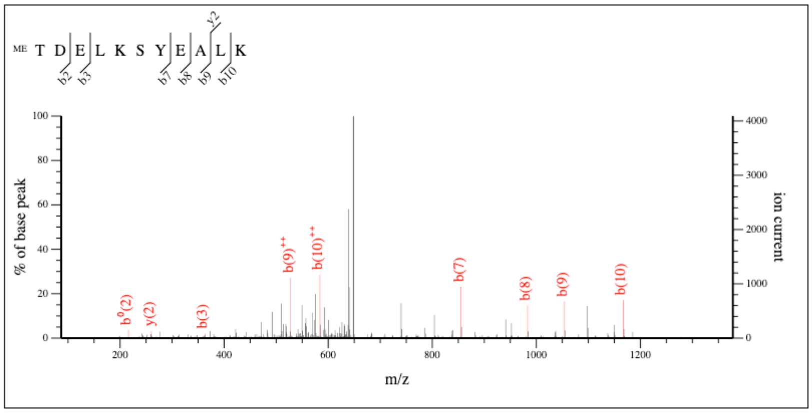

**Monoisotopic mass of neutral peptide Mr(calc):** 1312.6953
**Fixed modifications:** Carbamidomethyl (C) (apply to specified residues or termini only)
**Variable modifications:**
**N-term :** Methyl:2H(3) (Protein N-term)
**Ions Score:** 25 **Expect:** 0.047
**Matches :** 9/106 fragment ions using 22 most intense peaks (help)

| # | b | b++ | b* | b*++ | b0 | b0++ | Seq. | y | y++ | y* | y*++ | y0 | y0++ | # |
|---|---|---|---|---|---|---|---|---|---|---|---|---|---|---|
| 1 | 119.0894 | 60.0484 | | | 101.0789 | 51.0431 | T | | | | | | | 11 |
| 2 | 234.1164 | 117.5618 | | | 216.1058 | 108.5565 | D | 1195.6205 | 598.3139 | 1178.5939 | 589.8006 | 1177.6099 | 589.3086 | 10 |
| 3 | ***363.1590*** | 182.0831 | | | 345.1484 | 173.0778 | E | 1080.5936 | 540.8004 | 1063.5670 | 532.2871 | 1062.5830 | 531.7951 | 9 |
| 4 | 476.2430 | 238.6252 | | | 458.2325 | 229.6199 | L | 951.5510 | 476.2791 | 934.5244 | 467.7658 | 933.5404 | 467.2738 | 8 |
| 5 | 604.3380 | 302.6726 | 587.3114 | 294.1594 | 586.3274 | 293.6674 | K | 838.4669 | 419.7371 | 821.4403 | 411.2238 | 820.4563 | 410.7318 | 7 |
| 6 | 691.3700 | 346.1887 | 674.3435 | 337.6754 | 673.3595 | 337.1834 | S | 710.3719 | 355.6896 | 693.3454 | 347.1763 | 692.3614 | 346.6843 | 6 |
| 7 | ***854.4334*** | 427.7203 | 837.4068 | 419.2070 | 836.4228 | 418.7150 | Y | 623.3399 | 312.1736 | 606.3134 | 303.6603 | 605.3293 | 303.1683 | 5 |
| 8 | ***983.4759*** | 492.2416 | 966.4494 | 483.7283 | 965.4654 | 483.2363 | E | 460.2766 | 230.6419 | 443.2500 | 222.1287 | 442.2660 | 221.6366 | 4 |
| 9 | ***1054.5131*** | **527.7602** | 1037.4865 | 519.2469 | 1036.5025 | 518.7549 | A | 331.2340 | 166.1206 | 314.2074 | 157.6074 | | | 3 |
| 10 | ***1167.5971*** | **584.3022** | 1150.5706 | 575.7889 | 1149.5866 | 575.2969 | L | 260.1969 | 130.6021 | 243.1703 | 122.0888 | | | 2 |
| 11 | | | | | | | K | 147.1128 | 74.0600 | 130.0863 | 65.5468 | | | 1 |

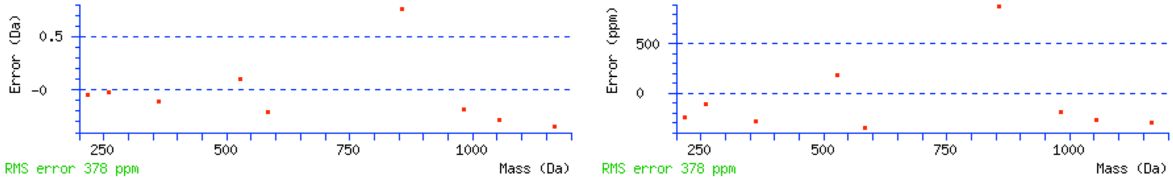


### 75. YLR460C (1 peptide in 400mM fraction 2)

MS/MS Fragmentation of **QVAIPETMK**
Found in **P54007** in **Uniprot_yeastS288C**, Uncharacterized protein YLR460C OS=Saccharomyces cerevisiae (strain ATCC 204508 / S288C) OX=559292 GN=YLR460C PE=1 SV=1

Match to Query 5326: 1082.585448 from(542.300000,2+) intensity(388884.5000) rtinseconds(1428.4894) index(3593)
Title: 20100530_methyl400mM_2.4000.4000.
Data file D:\Panyue\Proteomic dataset\20100530_methyl400mM_2.RAW

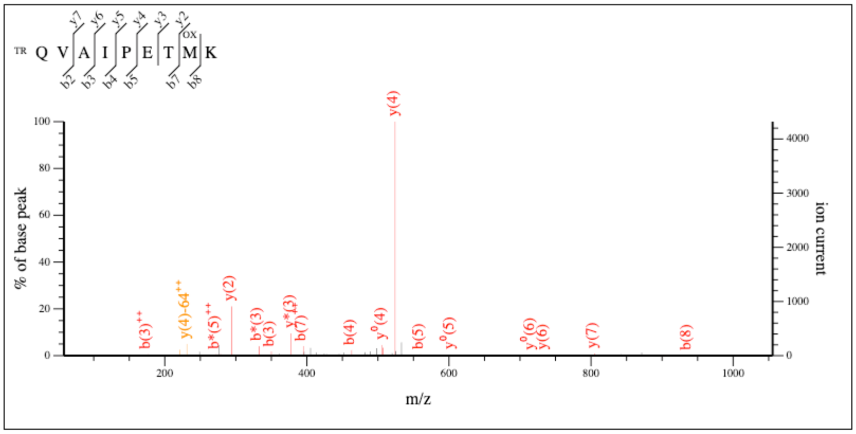

**Monoisotopic mass of neutral peptide Mr(calc):** 1082.6356
**Fixed modifications:** Carbamidomethyl (C) (apply to specified residues or termini only)
**Variable modifications:**
**N-term :** Trimethyl:2H(9) (Protein N-term)
**M8** : Oxidation (M), with neutral losses 0.0000(shown in table), 63.9983
**Ions Score:** 25 **Expect:** 0.039
**Matches :** 26/128 fragment ions using 55 most intense peaks (help)

| # | b | $b^{++}$ | b* | $b^{*++}$ | $b^{0}$ | $b^{0++}$ | Seq. | y | $y^{++}$ | y* | $y^{*++}$ | $y^{0}$ | $y^{0++}$ | # |
|---|---|---|---|---|---|---|---|---|---|---|---|---|---|---|
| 1 | 180.1693 | 90.5883 | 163.1427 | 82.0750 | | | Q | | | | | | | 9 |
| 2 | **279.2377** | 140.1225 | 262.2112 | 131.6092 | | | V | 904.4808 | 452.7441 | 887.4543 | 444.2308 | 886.4703 | 443.7388 | 8 |
| 3 | **350.2748** | **175.6410** | 333.2483 | 167.1278 | | | A | **805.4124** | 403.2098 | 788.3859 | 394.6966 | 787.4019 | 394.2046 | 7 |
| 4 | **463.3589** | **232.1831** | 446.3323 | 223.6698 | | | I | **734.3753** | 367.6913 | 717.3488 | 359.1780 | 716.3647 | 358.6860 | 6 |
| 5 | **560.4117** | 280.7095 | 543.3851 | 272.1962 | | | P | 621.2912 | 311.1493 | 604.2647 | 302.6360 | 603.2807 | 302.1440 | 5 |
| 6 | 689.4542 | 345.2308 | 672.4277 | 336.7175 | 671.4437 | 336.2255 | E | **524.2385** | 262.6229 | 507.2119 | 254.1096 | 506.2279 | 253.6176 | 4 |
| 7 | 790.5019 | **395.7546** | 773.4754 | 387.2413 | 772.4914 | 386.7493 | T | **395.1959** | 198.1016 | 378.1693 | 189.5883 | 377.1853 | 189.0963 | 3 |
| 8 | **937.5373** | 469.2723 | 920.5108 | 460.7590 | 919.5268 | 460.2670 | M | **294.1482** | 147.5777 | 277.1217 | 139.0645 | | | 2 |
| 9 | | | | | | | K | 147.1128 | 74.0600 | 130.0863 | 65.5468 | | | 1 |

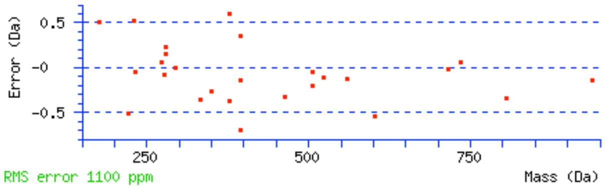


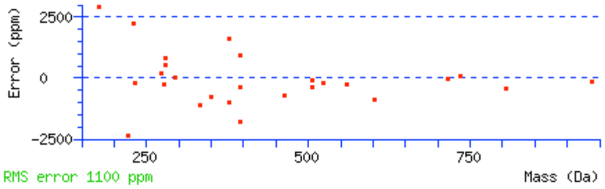


### 76. Fzo1 (1 peptide in 400mM fraction 2)

MS/MS Fragmentation of **SEGKQQFK**
Found in **P38297** in **Uniprot_yeastS288C**, Mitofusin FZO1 OS=Saccharomyces cerevisiae (strain ATCC 204508 / S288c) OX=559292 GN=FZO1 PE=1 SV=1

Match to Query 4425: 984.539158 from(493.276855,2+) intensity(626786.1875) rtinseconds(1629.841) index(4126)
Title: 20100530_methyl400mM_2.4593.4593.2
Data file D:\Panyue\Proteomic dataset\20100530_methyl400mM_2.RAW

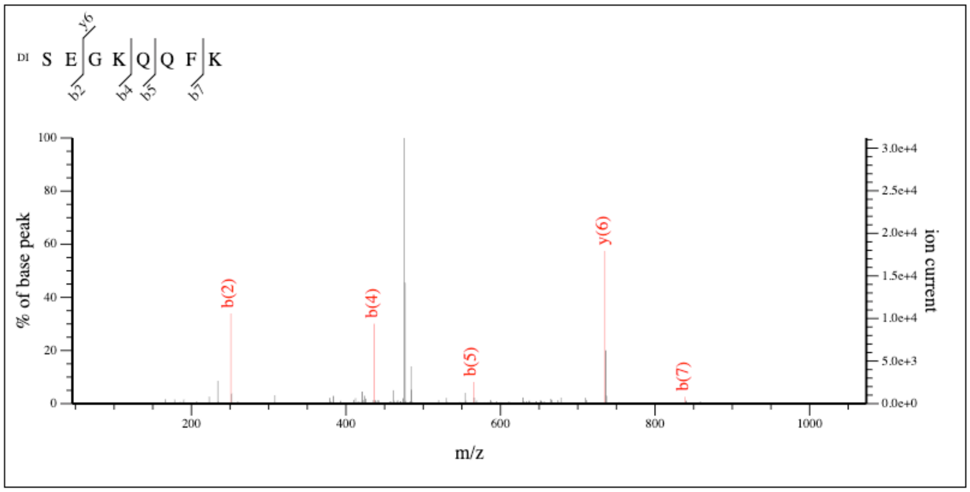

**Monoisotopic mass of neutral peptide Mr(calc):** 984.5511
**Fixed modifications:** Carbamidomethyl (C) (apply to specified residues or termini only)
**Variable modifications:**
**N-term :** Dimethyl:2H(6) (Protein N-term)
**Ions Score:** 28 **Expect:** 0.022
**Matches :** 5/66 fragment ions using 9 most intense peaks (help)

| # | b | b++ | b* | b*++ | b0 | b0++ | Seq. | y | y++ | y* | y*++ | y0 | y0++ | # |
|---|---|---|---|---|---|---|---|---|---|---|---|---|---|---|
| 1 | 122.1083 | 61.5578 | | | 104.0977 | 52.5525 | S | | | | | | | 8 |
| 2 | *251.1509* | 126.0791 | | | 233.1403 | 117.0738 | E | 864.4574 | 432.7323 | 847.4308 | 424.2191 | 846.4468 | 423.7271 | 7 |
| 3 | 308.1723 | 154.5898 | | | 290.1618 | 145.5845 | G | 735.4148 | 368.2110 | 718.3883 | 359.6978 | | | 6 |
| 4 | *436.2673* | 218.6373 | 419.2407 | 210.1240 | 418.2567 | 209.6320 | K | 678.3933 | 339.7003 | 661.3668 | 331.1870 | | | 5 |
| 5 | *564.3259* | 282.6666 | 547.2993 | 274.1533 | 546.3153 | 273.6613 | Q | 550.2984 | 275.6528 | 533.2718 | 267.1396 | | | 4 |
| 6 | 692.3844 | 346.6959 | 675.3579 | 338.1826 | 674.3739 | 337.6906 | Q | 422.2398 | 211.6235 | 405.2132 | 203.1103 | | | 3 |
| 7 | *839.4529* | 420.2301 | 822.4263 | 411.7168 | 821.4423 | 411.2248 | F | 294.1812 | 147.5942 | 277.1547 | 139.0810 | | | 2 |
| 8 | | | | | | | K | 147.1128 | 74.0600 | 130.0863 | 65.5468 | | | 1 |

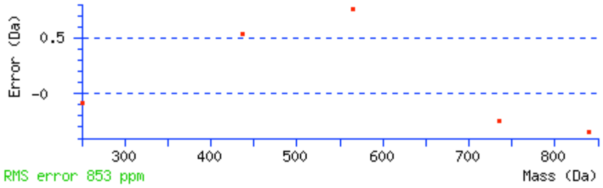


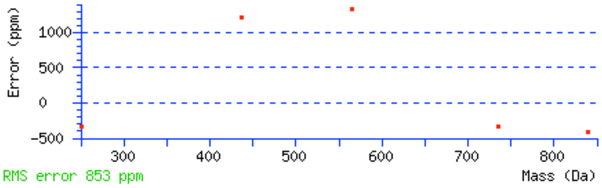


### 77. Stp22 (1 peptide in 400mM fraction 2)

MS/MS Fragmentation of **SANGKISVPEAVVNWLFK**
Found in **P25604** in **Uniprot_yeastS288C**, Suppressor protein STP22 of temperature-sensitive alpha-factor receptor and arginine permease OS=Saccharomyces cerevisiae (strain ATCC 204508 / S288c) OX=559292 GN=STP22 PE=1 SV=3

Match to Query 11259: 1975.045448 from(988.530000,2+) intensity(425332.9375) rtinseconds(3357.3891) index(8694)
Title: 20100530_methyl400mM_2.9673.9673.
Data file D:\Panyue\Proteomic dataset\20100530_methyl400mM_2.RAW

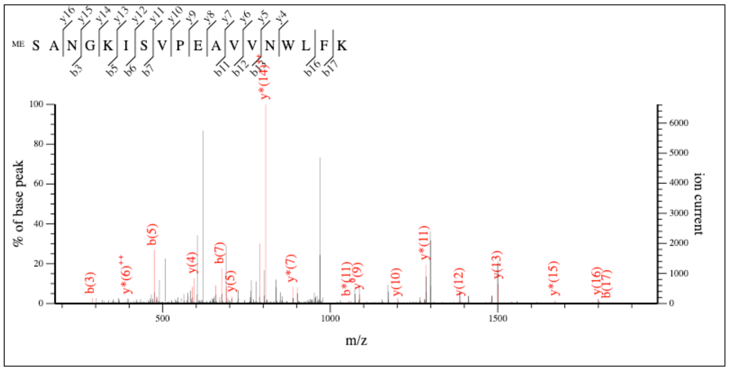

**Monoisotopic mass of neutral peptide Mr(calc):** 1975.0970
**Fixed modifications:** Carbamidomethyl (C) (apply to specified residues or termini only)
**Variable modifications:**
**N-term :** Methyl:2H(3) (Protein N-term)
**Ions Score:** 20 **Expect:** 0.048
**Matches :** 36/184 fragment ions using 96 most intense peaks (help)

| # | b | $b^{++}$ | b* | $b^{*++}$ | $b^{0}$ | $b^{0++}$ | Seq. | y | $y^{++}$ | y* | $y^{*++}$ | $y^{0}$ | $y^{0++}$ | # |
|---|---|---|---|---|---|---|---|---|---|---|---|---|---|---|
| 1 | 105.0738 | 53.0405 | | | 87.0632 | 44.0352 | S | | | | | | | 18 |
| 2 | 176.1109 | 88.5591 | | | 158.1003 | 79.5538 | A | 1872.0378 | 936.5225 | 1855.0112 | 928.0093 | 1854.0272 | 927.5173 | 17 |
| 3 | 290.1538 | 145.5806 | 273.1273 | 137.0673 | 272.1433 | 136.5753 | N | 1801.0007 | 901.0040 | 1783.9741 | 892.4907 | 1782.9901 | 891.9987 | 16 |
| 4 | 347.1753 | 174.0913 | 330.1487 | 165.5780 | 329.1647 | 165.0860 | G | 1686.9578 | 843.9825 | 1669.9312 | 835.4692 | 1668.9472 | 834.9772 | 15 |
| 5 | 475.2703 | 238.1388 | 458.2437 | 229.6255 | 457.2597 | 229.1335 | K | 1629.9363 | 815.4718 | 1612.9097 | 806.9585 | 1611.9257 | 806.4665 | 14 |
| 6 | 588.3543 | 294.6808 | 571.3278 | 286.1675 | 570.3438 | 285.6755 | I | 1501.8413 | 751.4243 | 1484.8148 | 742.9110 | 1483.8308 | 742.4190 | 13 |
| 7 | 675.3863 | 338.1968 | 658.3598 | 329.6835 | 657.3758 | 329.1915 | S | 1388.7573 | 694.8823 | 1371.7307 | 686.3690 | 1370.7467 | 685.8770 | 12 |
| 8 | 774.4548 | 387.7310 | 757.4282 | 379.2177 | 756.4442 | 378.7257 | V | 1301.7252 | 651.3663 | 1284.6987 | 642.8530 | 1283.7147 | 642.3610 | 11 |
| 9 | 871.5075 | 436.2574 | 854.4810 | 427.7441 | 853.4970 | 427.2521 | P | 1202.6568 | 601.8320 | 1185.6303 | 593.3188 | 1184.6463 | 592.8268 | 10 |
| 10 | 1000.5501 | 500.7787 | 983.5236 | 492.2654 | 982.5395 | 491.7734 | E | 1105.6041 | 553.3057 | 1088.5775 | 544.7924 | 1087.5935 | 544.3004 | 9 |
| 11 | 1071.5872 | 536.2973 | 1054.5607 | 527.7840 | 1053.5767 | 527.2920 | A | 976.5615 | 488.7844 | 959.5349 | 480.2711 | | | 8 |
| 12 | 1170.6556 | 585.8315 | 1153.6291 | 577.3182 | 1152.6451 | 576.8262 | V | 905.5244 | 453.2658 | 888.4978 | 444.7525 | | | 7 |
| 13 | 1269.7241 | 635.3657 | 1252.6975 | 626.8524 | 1251.7135 | 626.3604 | V | 806.4559 | 403.7316 | 789.4294 | 395.2183 | | | 6 |
| 14 | 1383.7670 | 692.3871 | 1366.7404 | 683.8739 | 1365.7564 | 683.3818 | N | 707.3875 | 354.1974 | 690.3610 | 345.6841 | | | 5 |
| 15 | 1569.8463 | 785.4268 | 1552.8197 | 776.9135 | 1551.8357 | 776.4215 | W | 593.3446 | 297.1759 | 576.3180 | 288.6627 | | | 4 |
| 16 | 1682.9304 | 841.9688 | 1665.9038 | 833.4555 | 1664.9198 | 832.9635 | L | 407.2653 | 204.1363 | 390.2387 | 195.6230 | | | 3 |
| 17 | 1829.9988 | 915.5030 | 1812.9722 | 906.9898 | 1811.9882 | 906.4977 | F | 294.1812 | 147.5942 | 277.1547 | 139.0810 | | | 2 |
| 18 | | | | | | | K | 147.1128 | 74.0600 | 130.0863 | 65.5468 | | | 1 |

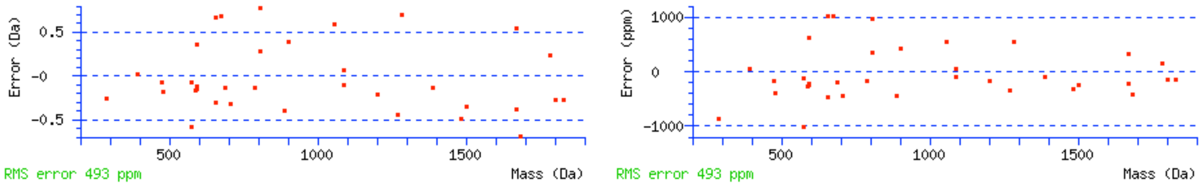


### 78. Pdc6 (2 peptides in 400mM fraction 3)

MS/MS Fragmentation of **SEITLGK**
Found in **P26263** in **Uniprot_yeastS288C**, Pyruvate decarboxylase isozyme 3 OS=Saccharomyces cerevisiae (strain ATCC 204508 / S288c) OX=559292 GN=PDC6 PE=1 SV=3

Match to Query 2053: 780.448840 from(781.456116,1+) intensity(12796055.0000) rtinseconds(1047.2379) index(2599)
Title: 20100530_methyl400mM_3.2892.2892.1
Data file D:\Panyue\Proteomic dataset\20100530_methyl400mM_3.RAW

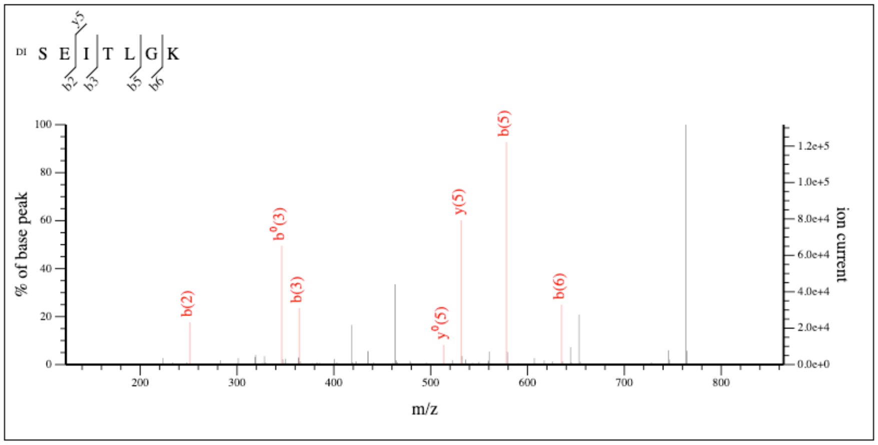

**Monoisotopic mass of neutral peptide Mr(calc):** 780.4864
**Fixed modifications:** Carbamidomethyl (C) (apply to specified residues or termini only)
**Variable modifications:**
**N-term :** Dimethyl:2H(6) (Protein N-term)
**Ions Score:** 27 **Expect:** 0.026
**Matches :** 7/27 fragment ions using 12 most intense peaks (help)

| # | b | b⁰ | Seq. | y | y* | y⁰ | # |
|---|---|---|---|---|---|---|---|
| 1 | 122.1083 | 104.0977 | S | | | | 7 |
| 2 | 251.1509 | 233.1403 | E | 660.3927 | 643.3661 | 642.3821 | 6 |
| 3 | 364.2349 | 346.2244 | I | 531.3501 | 514.3235 | 513.3395 | 5 |
| 4 | 465.2826 | 447.2720 | T | 418.2660 | 401.2395 | 400.2554 | 4 |
| 5 | 578.3667 | 560.3561 | L | 317.2183 | 300.1918 | | 3 |
| 6 | 635.3881 | 617.3776 | G | 204.1343 | 187.1077 | | 2 |
| 7 | | | K | 147.1128 | 130.0863 | | 1 |

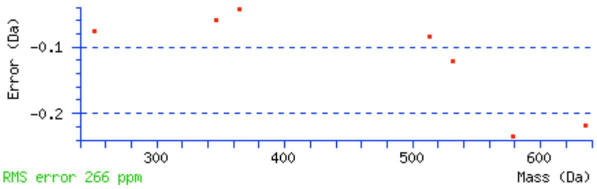


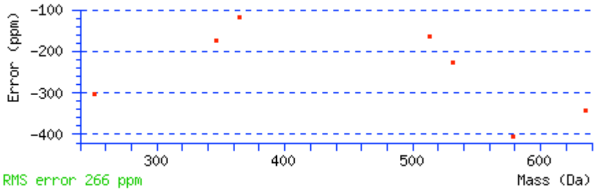


MS/MS Fragmentation of **SEITLGK**
Found in **P26263** in **Uniprot_yeastS288C**, Pyruvate decarboxylase isozyme 3 OS=Saccharomyces cerevisiae (strain ATCC 204508 / S288c) OX=559292 GN=PDC6 PE=1 SV=3

Match to Query 2055: 780.449999 from(781.457275,1+) intensity(8012099.0000) rtinseconds(1050.4695) index(2608)
Title: 20100530_methyl400mM_3.2902.2902.1
Data file D:\Panyue\Proteomic dataset\20100530_methyl400mM_3.RAW

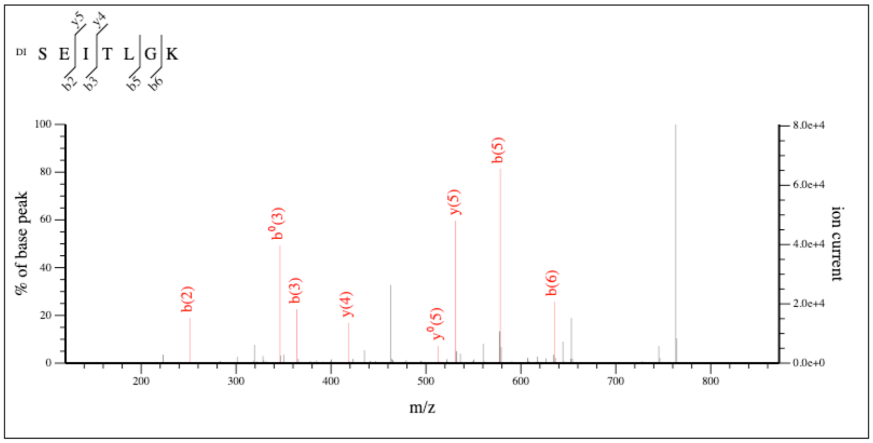

**Monoisotopic mass of neutral peptide Mr(calc):** 780.4864
**Fixed modifications:** Carbamidomethyl (C) (apply to specified residues or termini only)
**Variable modifications:**
**N-term :** Dimethyl:2H(6) (Protein N-term)
**Ions Score:** 25 **Expect:** 0.036
**Matches :** 8/27 fragment ions using 18 most intense peaks (help)

| # | b | $b^0$ | Seq. | y | y* | $y^0$ | # |
|---|---|---|---|---|---|---|---|
| 1 | 122.1083 | 104.0977 | S | | | | 7 |
| 2 | *251.1509* | 233.1403 | E | 660.3927 | 643.3661 | 642.3821 | 6 |
| 3 | *364.2349* | 346.2244 | I | **531.3501** | 514.3235 | 513.3395 | 5 |
| 4 | 465.2826 | 447.2720 | T | **418.2660** | 401.2395 | 400.2554 | 4 |
| 5 | *578.3667* | 560.3561 | L | 317.2183 | 300.1918 | | 3 |
| 6 | *635.3881* | 617.3776 | G | 204.1343 | 187.1077 | | 2 |
| 7 | | | K | 147.1128 | 130.0863 | | 1 |

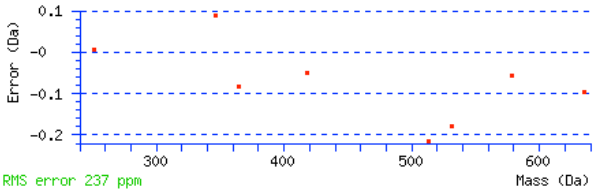


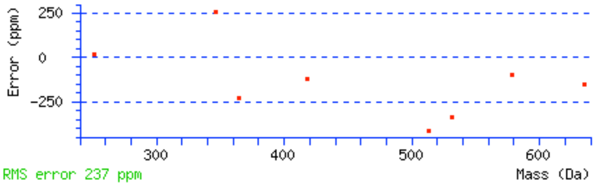


### 79. Pdc5 (2 peptides in 400mM fraction 3)

MS/MS Fragmentation of SEITLGK
Found in P16467 in Uniprot_yeastS288C, Pyruvate decarboxylase isozyme 2 OS=Saccharomyces cerevisiae (strain ATCC 204508 / S288c) OX=559292 GN=PDC5 PE=1 SV=4

Match to Query 2055: 780.449999 from(781.457275,1+) intensity(8012099.0000) rtinseconds(1050.4695) index(2608)
Title: 20100530_methyl400mM_3.2902.2902.1
Data file D:\Panyue\Proteomic dataset\20100530_methyl400mM_3.RAW

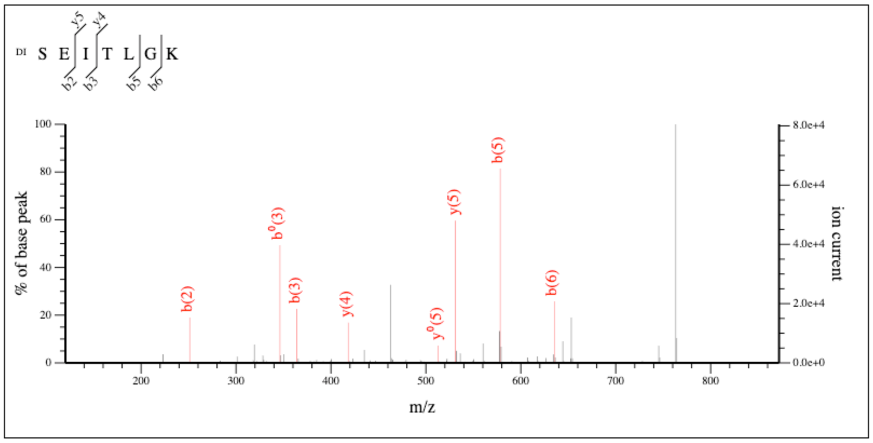

**Monoisotopic mass of neutral peptide Mr(calc):** 780.4864
**Fixed modifications:** Carbamidomethyl (C) (apply to specified residues or termini only)
**Variable modifications:**
**N-term :** Dimethyl:2H(6) (Protein N-term)
**Ions Score:** 25 **Expect:** 0.036
**Matches :** 8/27 fragment ions using 18 most intense peaks (help)

| # | b | $b^0$ | Seq. | y | y* | $y^0$ | # |
|---|---|---|---|---|---|---|---|
| 1 | 122.1083 | 104.0977 | S | | | | 7 |
| 2 | 251.1509 | 233.1403 | E | 660.3927 | 643.3661 | 642.3821 | 6 |
| 3 | 364.2349 | 346.2244 | I | 531.3501 | 514.3235 | 513.3395 | 5 |
| 4 | 465.2826 | 447.2720 | T | 418.2660 | 401.2395 | 400.2554 | 4 |
| 5 | 578.3667 | 560.3561 | L | 317.2183 | 300.1918 | | 3 |
| 6 | 635.3881 | 617.3776 | G | 204.1343 | 187.1077 | | 2 |
| 7 | | | K | 147.1128 | 130.0863 | | 1 |

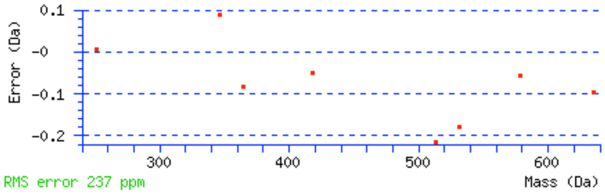


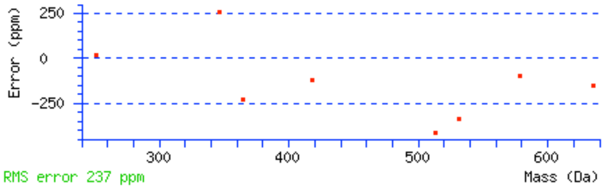


MS/MS Fragmentation of SEITLGK
Found in P16467 in Uniprot_yeastS288C, Pyruvate decarboxylase isozyme 2 OS=Saccharomyces cerevisiae (strain ATCC 204508 / S288c) OX=559292 GN=PDC5 PE=1 SV=4

Match to Query 2053: 780.448840 from(781.456116,1+) intensity(12796055.0000) rtinseconds(1047.2379) index(2599)
Title: 20100530_methyl400mM_3.2892.2892.1
Data file D:\Panyue\Proteomic dataset\20100530_methyl400mM_3.RAW

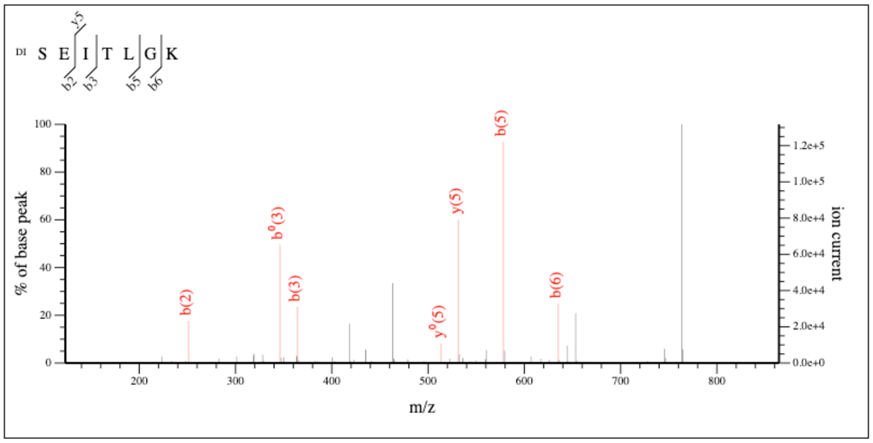

**Monoisotopic mass of neutral peptide Mr(calc):** 780.4864
**Fixed modifications:** Carbamidomethyl (C) (apply to specified residues or termini only)
**Variable modifications:**
**N-term :** Dimethyl:2H(6) (Protein N-term)
**Ions Score:** 27 **Expect:** 0.026
**Matches :** 7/27 fragment ions using 12 most intense peaks (help)

| # | b | b$^0$ | Seq. | y | y* | y$^0$ | # |
|---|---|---|---|---|---|---|---|
| 1 | 122.1083 | 104.0977 | S | | | | 7 |
| 2 | 251.1509 | 233.1403 | E | 660.3927 | 643.3661 | 642.3821 | 6 |
| 3 | 364.2349 | 346.2244 | I | 531.3501 | 514.3235 | 513.3395 | 5 |
| 4 | 465.2826 | 447.2720 | T | 418.2660 | 401.2395 | 400.2554 | 4 |
| 5 | 578.3667 | 560.3561 | L | 317.2183 | 300.1918 | | 3 |
| 6 | 635.3881 | 617.3776 | G | 204.1343 | 187.1077 | | 2 |
| 7 | | | K | 147.1128 | 130.0863 | | 1 |

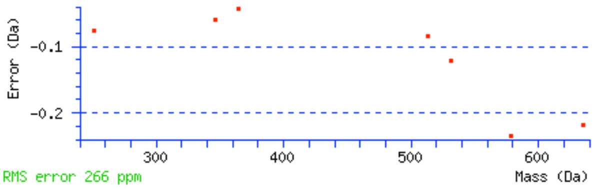


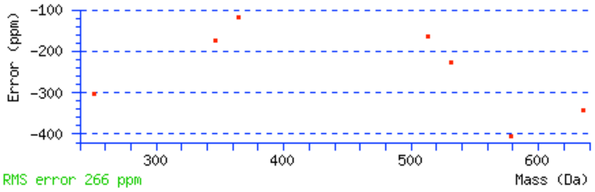


### 80. GRS1 (1 peptide in 400mM fraction 3)

MS/MS Fragmentation of **SVEDIKK**
Found in **P38088-2** in **Uniprot_yeastS288C**, Isoform Cytoplasmic of Glycine--tRNA ligase 1, mitochondrial OS=Saccharomyces cerevisiae (strain ATCC 204508 / S288c) OX=559292 GN=GRS1

Match to Query 2578: 834.459874 from(418.237213,2+) intensity(280349.9688) rtinseconds(1123.5963) index(2808)
Title: 20100530_methyl400mM_3.3124.3124.2
Data file D:\Panyue\Proteomic dataset\20100530_methyl400mM_3.RAW

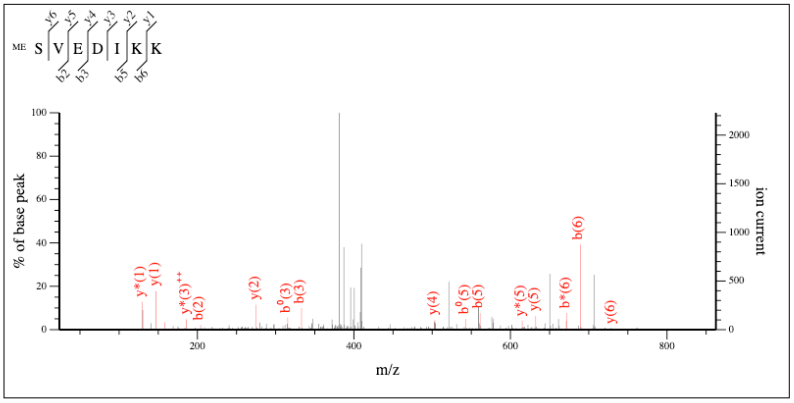

**Monoisotopic mass of neutral peptide Mr(calc):** 834.4890
**Fixed modifications:** Carbamidomethyl (C) (apply to specified residues or termini only)
**Variable modifications:**
**N-term :** Methyl:2H(3) (Protein N-term)
**Ions Score:** 20 **Expect:** 0.18
**Matches :** 20/56 fragment ions using 48 most intense peaks (help)

| # | b | b++ | b* | b*++ | b0 | b0++ | Seq. | y | y++ | y* | y*++ | y0 | y0++ | # |
|---|---|---|---|---|---|---|---|---|---|---|---|---|---|---|
| 1 | 105.0738 | 53.0405 | | | 87.0632 | 44.0352 | S | | | | | | | 7 |
| 2 | 204.1422 | 102.5747 | | | 186.1316 | 93.5695 | V | 731.4298 | 366.2185 | 714.4032 | 357.7053 | 713.4192 | 357.2132 | 6 |
| 3 | 333.1848 | 167.0960 | | | 315.1742 | 158.0908 | E | 632.3614 | 316.6843 | 615.3348 | 308.1710 | 614.3508 | 307.6790 | 5 |
| 4 | 448.2117 | 224.6095 | | | 430.2012 | 215.6042 | D | 503.3188 | 252.1630 | 486.2922 | 243.6498 | 485.3082 | 243.1577 | 4 |
| 5 | 561.2958 | 281.1515 | | | 543.2852 | 272.1463 | I | 388.2918 | 194.6496 | 371.2653 | 186.1363 | | | 3 |
| 6 | 689.3908 | 345.1990 | 672.3642 | 336.6857 | 671.3802 | 336.1937 | K | 275.2078 | 138.1075 | 258.1812 | 129.5942 | | | 2 |
| 7 | | | | | | | K | 147.1128 | 74.0600 | 130.0863 | 65.5468 | | | 1 |

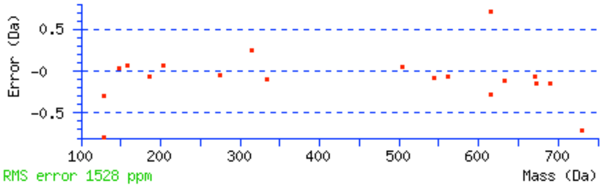


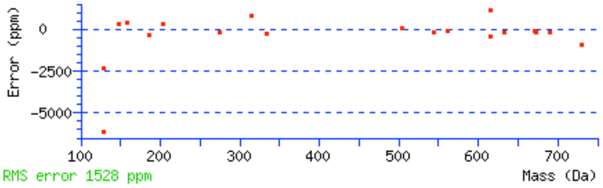


### 81. Eft1 (1 peptide in 400mM fraction 3)

MS/MS Fragmentation of **VAFTVDQMRSLMDK**
Found in **P32324** in **Uniprot_yeastS288C**, Elongation factor 2 OS=Saccharomyces cerevisiae (strain ATCC 204508 / S288c) OX=559292 GN=EFT1 PE=1 SV=1

Match to Query 9380: 1706.865448 from(854.440000,2+) intensity(1266505.0000) rtinseconds(2687.4876) index(6973)
Title: 20100530_methyl400mM_3.7757.7757.
Data file D:\Panyue\Proteomic dataset\20100530_methyl400mM_3.RAW

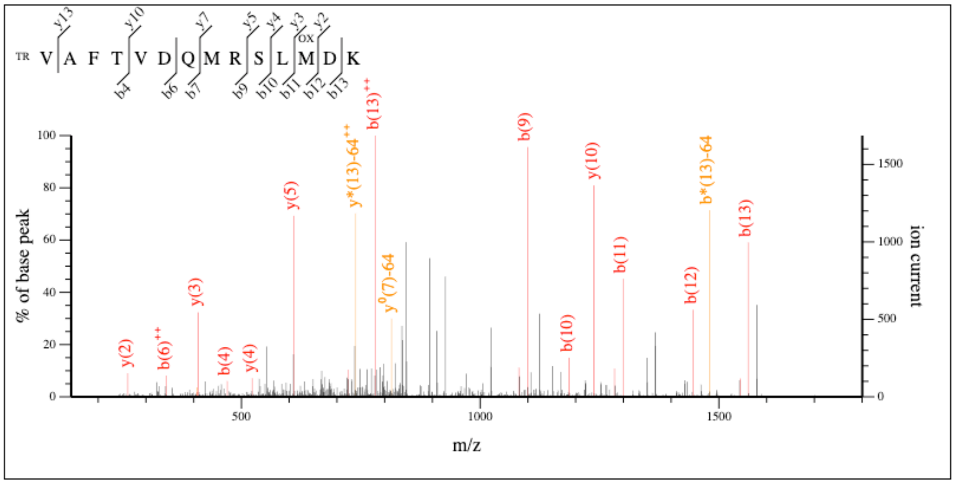

```
Monoisotopic mass of neutral peptide Mr(calc): 1706.9046
Fixed modifications: Carbamidomethyl (C) (apply to specified residues or termini only)
Variable modifications:
N-term : Trimethyl:2H(9) (Protein N-term)
M12    : Oxidation (M), with neutral losses 0.0000(shown in table), 63.9983
Ions Score: 25  Expect: 0.13
Matches : 25/214 fragment ions using 43 most intense peaks   (help)
```

| # | b | b++ | b* | b*++ | b0 | b0++ | Seq. | y | y++ | y* | y*++ | y0 | y0++ | # |
|---|---|---|---|---|---|---|---|---|---|---|---|---|---|---|
| 1 | 151.1791 | 76.0932 | | | | | V | | | | | | | 14 |
| 2 | 222.2162 | 111.6118 | | | | | A | 1557.7400 | 779.3736 | 1540.7134 | 770.8604 | 1539.7294 | 770.3683 | 13 |
| 3 | 369.2847 | 185.1460 | | | | | F | 1486.7029 | 743.8551 | 1469.6763 | 735.3418 | 1468.6923 | 734.8498 | 12 |
| 4 | ***470.3323*** | 235.6698 | | | 452.3218 | 226.6645 | T | 1339.6344 | 670.3209 | 1322.6079 | 661.8076 | 1321.6239 | 661.3156 | 11 |
| 5 | 569.4008 | 285.2040 | | | 551.3902 | 276.1987 | V | **1238.5868** | 619.7970 | 1221.5602 | 611.2837 | 1220.5762 | 610.7917 | 10 |
| 6 | 684.4277 | ***342.7175*** | | | 666.4171 | 333.7122 | D | 1139.5184 | 570.2628 | 1122.4918 | 561.7495 | 1121.5078 | 561.2575 | 9 |
| 7 | 812.4863 | ***406.7468*** | 795.4597 | 398.2335 | 794.4757 | 397.7415 | Q | 1024.4914 | 512.7493 | 1007.4649 | 504.2361 | 1006.4808 | 503.7441 | 8 |
| 8 | 943.5268 | 472.2670 | 926.5002 | 463.7537 | 925.5162 | 463.2617 | M | 896.4328 | 448.7201 | 879.4063 | 440.2068 | 878.4223 | 439.7148 | 7 |
| 9 | ***1099.6279*** | 550.3176 | 1082.6013 | 541.8043 | 1081.6173 | 541.3123 | R | 765.3924 | 383.1998 | 748.3658 | 374.6865 | 747.3818 | 374.1945 | 6 |
| 10 | ***1186.6599*** | 593.8336 | 1169.6333 | 585.3203 | 1168.6493 | 584.8283 | S | **609.2912** | 305.1493 | 592.2647 | 296.6360 | 591.2807 | 296.1440 | 5 |
| 11 | ***1299.7440*** | 650.3756 | 1282.7174 | 641.8623 | 1281.7334 | 641.3703 | L | **522.2592** | 261.6332 | 505.2327 | 253.1200 | 504.2486 | 252.6280 | 4 |
| 12 | ***1446.7794*** | ***723.8933*** | 1429.7528 | 715.3800 | 1428.7688 | 714.8880 | M | **409.1751** | 205.0912 | 392.1486 | 196.5779 | 391.1646 | 196.0859 | 3 |
| 13 | ***1561.8063*** | ***781.4068*** | 1544.7798 | 772.8935 | 1543.7957 | 772.4015 | D | **262.1397** | 131.5735 | 245.1132 | 123.0602 | 244.1292 | 122.5682 | 2 |
| 14 | | | | | | | K | 147.1128 | 74.0600 | 130.0863 | 65.5468 | | | 1 |

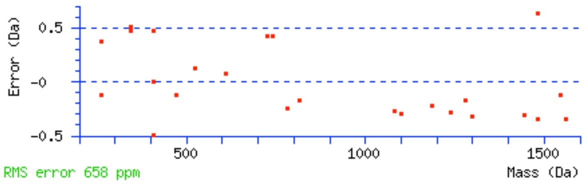


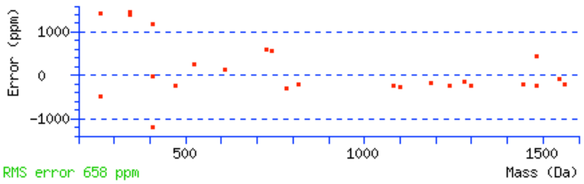


### 82. Pdc5 (2 peptides in 450mM fraction 1)

MS/MS Fragmentation of **SEITLGK**
Found in **P16467** in **Uniprot_yeastS288C**, Pyruvate decarboxylase isozyme 2 OS=Saccharomyces cerevisiae (strain ATCC 204508 / S288c) OX=559292 GN=PDC5 PE=1 SV=4

Match to Query 2255: 780.449694 from(781.456970,1+) intensity(14809349.0000) rtinseconds(1062.9977) index(2620)
Title: 20100530_methyl450mM_1.2917.2917.1
Data file D:\Panyue\Proteomic dataset\20100530_methyl450mM_1.RAW

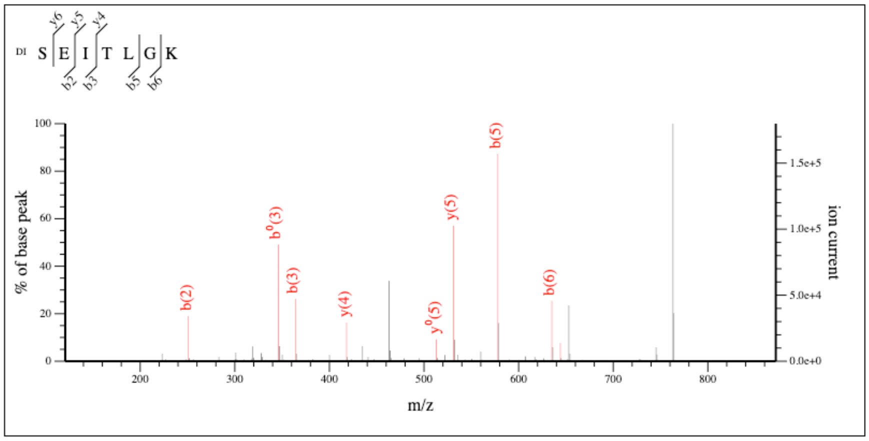

**Monoisotopic mass of neutral peptide Mr(calc):** 780.4864
**Fixed modifications:** Carbamidomethyl (C) (apply to specified residues or termini only)
**Variable modifications:**
**N-term :** Dimethyl:2H(6) (Protein N-term)
**Ions Score:** 29 **Expect:** 0.017
**Matches :** 9/27 fragment ions using 18 most intense peaks (help)

| # | b | $b^0$ | Seq. | y | y* | $y^0$ | # |
|---|---|---|---|---|---|---|---|
| 1 | 122.1083 | 104.0977 | S | | | | 7 |
| 2 | *251.1509* | 233.1403 | E | 660.3927 | 643.3661 | 642.3821 | 6 |
| 3 | *364.2349* | 346.2244 | I | **531.3501** | 514.3235 | 513.3395 | 5 |
| 4 | 465.2826 | 447.2720 | T | **418.2660** | 401.2395 | 400.2554 | 4 |
| 5 | *578.3667* | 560.3561 | L | 317.2183 | 300.1918 | | 3 |
| 6 | *635.3881* | 617.3776 | G | 204.1343 | 187.1077 | | 2 |
| 7 | | | K | 147.1128 | 130.0863 | | 1 |

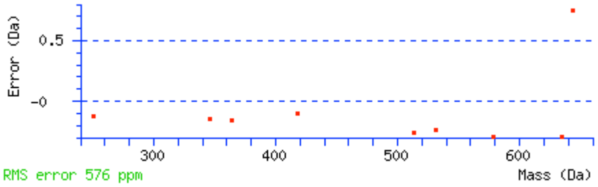


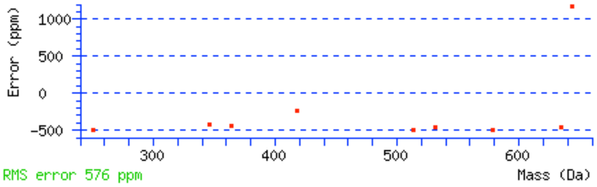


MS/MS Fragmentation of **SEITLGK**
Found in **P16467** in **Uniprot_yeastS288C**, Pyruvate decarboxylase isozyme 2 OS=Saccharomyces cerevisiae (strain ATCC 204508 / S288c) OX=559292 GN=PDC5 PE=1 SV=4

Match to Query 2257: 780.449999 from(781.457275,1+) intensity(15218069.0000) rtinseconds(1059.8959) index(2612)
Title: 20100530_methyl450mM_1.2908.2908.1
Data file D:\Panyue\Proteomic dataset\20100530_methyl450mM_1.RAW

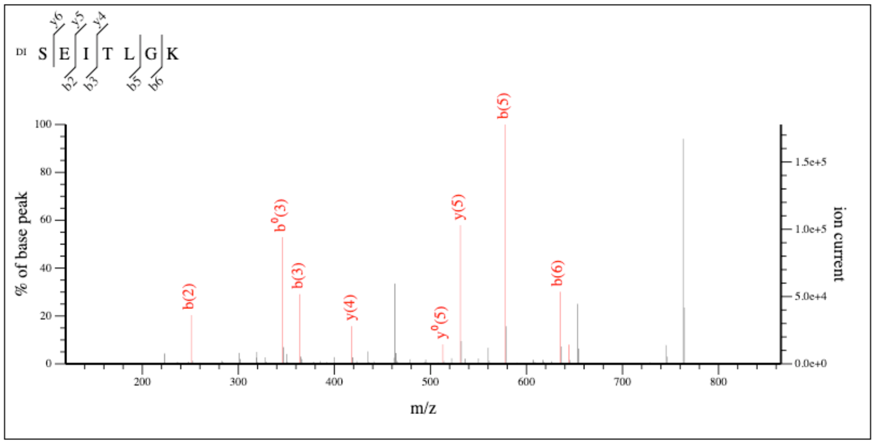

**Monoisotopic mass of neutral peptide Mr(calc):** 780.4864
**Fixed modifications:** Carbamidomethyl (C) (apply to specified residues or termini only)
**Variable modifications:**
**N-term :** Dimethyl:2H(6) (Protein N-term)
**Ions Score:** 27 **Expect:** 0.026
**Matches :** 9/27 fragment ions using 18 most intense peaks (help)

| # | b | $b^0$ | Seq. | y | y* | $y^0$ | # |
|---|---|---|---|---|---|---|---|
| 1 | 122.1083 | 104.0977 | S | | | | 7 |
| 2 | 251.1509 | 233.1403 | E | 660.3927 | 643.3661 | 642.3821 | 6 |
| 3 | 364.2349 | 346.2244 | I | 531.3501 | 514.3235 | 513.3395 | 5 |
| 4 | 465.2826 | 447.2720 | T | 418.2660 | 401.2395 | 400.2554 | 4 |
| 5 | 578.3667 | 560.3561 | L | 317.2183 | 300.1918 | | 3 |
| 6 | 635.3881 | 617.3776 | G | 204.1343 | 187.1077 | | 2 |
| 7 | | | K | 147.1128 | 130.0863 | | 1 |

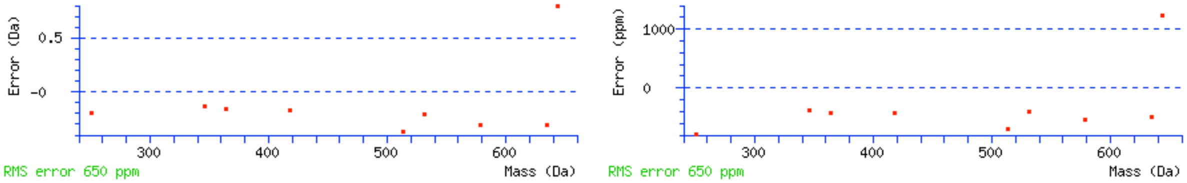


### 83. Pdc1 (2 peptides in 450mM fraction 1)

MS/MS Fragmentation of **SEITLGK**
Found in **P06169** in **Uniprot_yeastS288C**, Pyruvate decarboxylase isozyme 1 OS=Saccharomyces cerevisiae (strain ATCC 204508 / S288c) OX=559292 GN=PDC1 PE=1 SV=7

Match to Query 2255: 780.449694 from(781.456970,1+) intensity(14809349.0000) rtinseconds(1062.9977) index(2620)
Title: 20100530_methyl450mM_1.2917.2917.1
Data file D:\Panyue\Proteomic dataset\20100530_methyl450mM_1.RAW

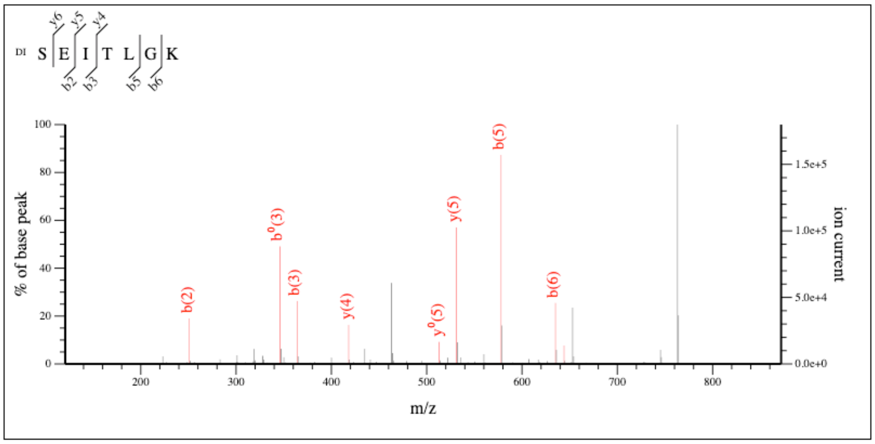

**Monoisotopic mass of neutral peptide Mr(calc):** 780.4864
**Fixed modifications:** Carbamidomethyl (C) (apply to specified residues or termini only)
**Variable modifications:**
**N-term :** Dimethyl:2H(6) (Protein N-term)
**Ions Score:** 29 **Expect:** 0.017
**Matches :** 9/27 fragment ions using 18 most intense peaks (help)

| # | b | $b^0$ | Seq. | y | y* | $y^0$ | # |
|---|---|---|---|---|---|---|---|
| 1 | 122.1083 | 104.0977 | S | | | | 7 |
| 2 | *251.1509* | 233.1403 | E | 660.3927 | 643.3661 | 642.3821 | 6 |
| 3 | *364.2349* | 346.2244 | I | **531.3501** | 514.3235 | 513.3395 | 5 |
| 4 | 465.2826 | 447.2720 | T | **418.2660** | 401.2395 | 400.2554 | 4 |
| 5 | *578.3667* | 560.3561 | L | 317.2183 | 300.1918 | | 3 |
| 6 | *635.3881* | 617.3776 | G | 204.1343 | 187.1077 | | 2 |
| 7 | | | K | 147.1128 | 130.0863 | | 1 |

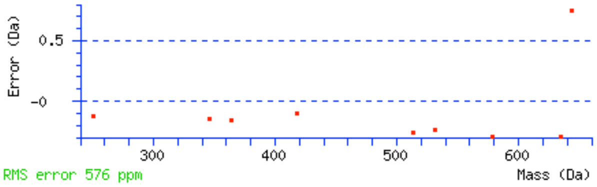


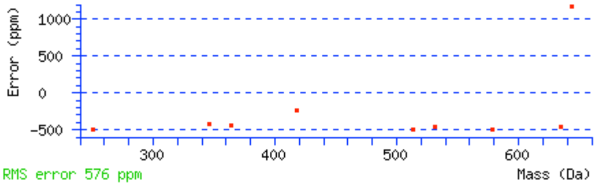


MS/MS Fragmentation of **SEITLGK**
Found in **P06169** in **Uniprot_yeastS288C**, Pyruvate decarboxylase isozyme 1 OS=Saccharomyces cerevisiae (strain ATCC 204508 / S288c) OX=559292 GN=PDC1 PE=1 SV=7

Match to Query 2257: 780.449999 from(781.457275,1+) intensity(15218069.0000) rtinseconds(1059.8959) index(2612)
Title: 20100530_methyl450mM_1.2908.2908.1
Data file D:\Panyue\Proteomic dataset\20100530_methyl450mM_1.RAW

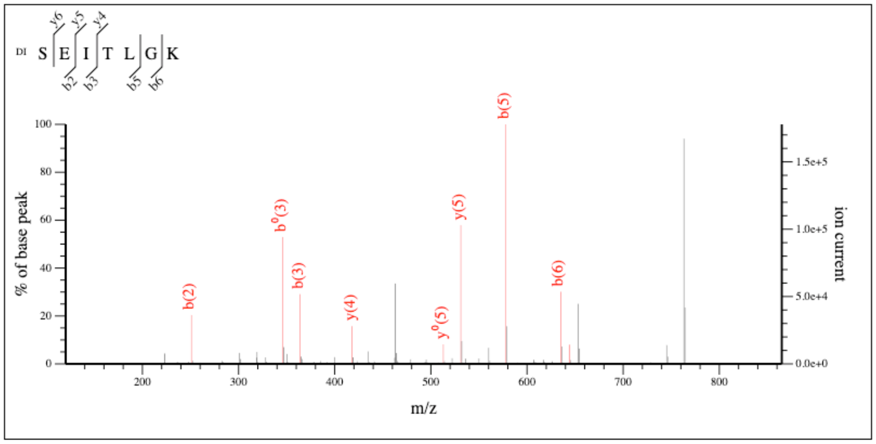

**Monoisotopic mass of neutral peptide Mr(calc):** 780.4864
**Fixed modifications:** Carbamidomethyl (C) (apply to specified residues or termini only)
**Variable modifications:**
**N-term :** Dimethyl:2H(6) (Protein N-term)
**Ions Score:** 27 **Expect:** 0.026
**Matches :** 9/27 fragment ions using 18 most intense peaks (help)

| # | b | $b^0$ | Seq. | y | y* | $y^0$ | # |
|---|---|---|---|---|---|---|---|
| 1 | 122.1083 | 104.0977 | S | | | | 7 |
| 2 | ***251.1509*** | 233.1403 | E | 660.3927 | 643.3661 | 642.3821 | 6 |
| 3 | ***364.2349*** | 346.2244 | I | **531.3501** | 514.3235 | 513.3395 | 5 |
| 4 | 465.2826 | 447.2720 | T | **418.2660** | 401.2395 | 400.2554 | 4 |
| 5 | ***578.3667*** | 560.3561 | L | 317.2183 | 300.1918 | | 3 |
| 6 | ***635.3881*** | 617.3776 | G | 204.1343 | 187.1077 | | 2 |
| 7 | | | K | 147.1128 | 130.0863 | | 1 |

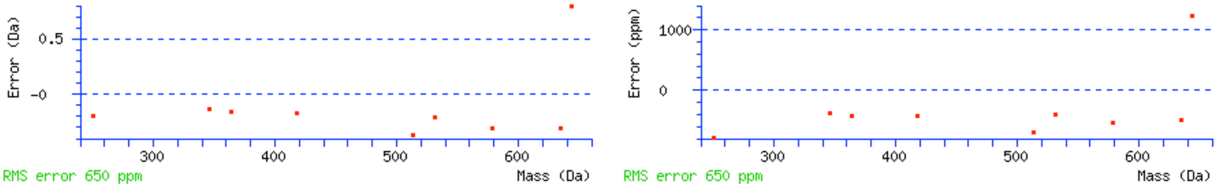


### 84. Fzo1 (2 peptides in 450mM fraction 1)

MS/MS Fragmentation of **SEGKQQFK**
Found in **P38297** in **Uniprot_yeastS288C**, Mitofusin FZO1 OS=Saccharomyces cerevisiae (strain ATCC 204508 / S288c) OX=559292 GN=FZO1 PE=1 SV=1

Match to Query 4284: 984.538670 from(493.276611,2+) intensity(298876.4688) rtinseconds(1646.6198) index(4145)
Title: 20100530_methyl450mM_1.4613.4613.2
Data file D:\Panyue\Proteomic dataset\20100530_methyl450mM_1.RAW

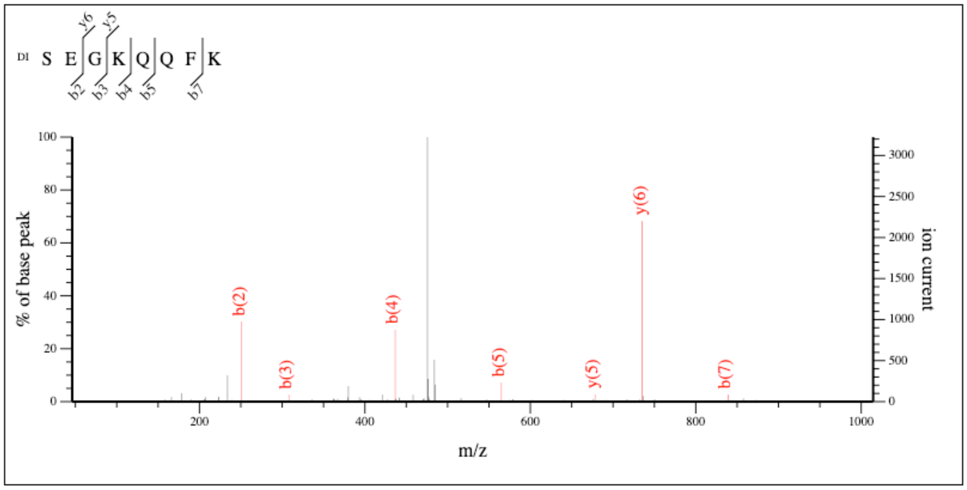

**Monoisotopic mass of neutral peptide Mr(calc):** 984.5511
**Fixed modifications:** Carbamidomethyl (C) (apply to specified residues or termini only)
**Variable modifications:**
**N-term :** Dimethyl:2H(6) (Protein N-term)
**Ions Score:** 31 **Expect:** 0.011
**Matches :** 7/66 fragment ions using 16 most intense peaks (help)

| # | b | b++ | b* | b*++ | b0 | b0++ | Seq. | y | y++ | y* | y*++ | y0 | y0++ | # |
|---|---|---|---|---|---|---|---|---|---|---|---|---|---|---|
| 1 | 122.1083 | 61.5578 | | | 104.0977 | 52.5525 | S | | | | | | | 8 |
| 2 | ***251.1509*** | 126.0791 | | | 233.1403 | 117.0738 | E | 864.4574 | 432.7323 | 847.4308 | 424.2191 | 846.4468 | 423.7271 | 7 |
| 3 | ***308.1723*** | 154.5898 | | | 290.1618 | 145.5845 | G | **735.4148** | 368.2110 | 718.3883 | 359.6978 | | | 6 |
| 4 | ***436.2673*** | 218.6373 | 419.2407 | 210.1240 | 418.2567 | 209.6320 | K | **678.3933** | 339.7003 | 661.3668 | 331.1870 | | | 5 |
| 5 | ***564.3259*** | 282.6666 | 547.2993 | 274.1533 | 546.3153 | 273.6613 | Q | 550.2984 | 275.6528 | 533.2718 | 267.1396 | | | 4 |
| 6 | 692.3844 | 346.6959 | 675.3579 | 338.1826 | 674.3739 | 337.6906 | Q | 422.2398 | 211.6235 | 405.2132 | 203.1103 | | | 3 |
| 7 | ***839.4529*** | 420.2301 | 822.4263 | 411.7168 | 821.4423 | 411.2248 | F | 294.1812 | 147.5942 | 277.1547 | 139.0810 | | | 2 |
| 8 | | | | | | | K | 147.1128 | 74.0600 | 130.0863 | 65.5468 | | | 1 |

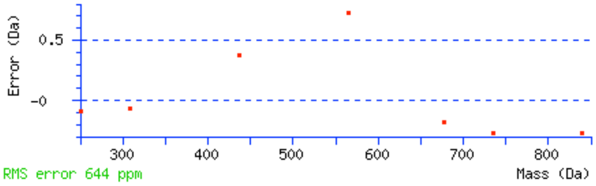


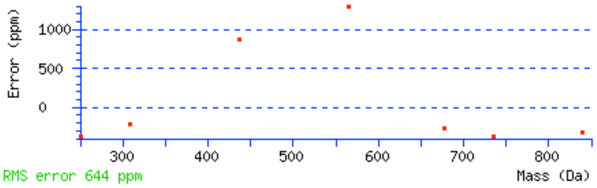


MS/MS Fragmentation of **SEGKQQFK**
Found in **P38297** in **Uniprot_yeastS288C**, Mitofusin FZO1 OS=Saccharomyces cerevisiae (strain ATCC 204508 / S288c) OX=559292 GN=FZO1 PE=1 SV=1

Match to Query 4285: 984.540087 from(985.547363,1+) intensity(180540.5469) rtinseconds(1656.771) index(4172)
Title: 20100530_methyl450mM_1.4643.4643.1
Data file D:\Panyue\Proteomic dataset\20100530_methyl450mM_1.RAW

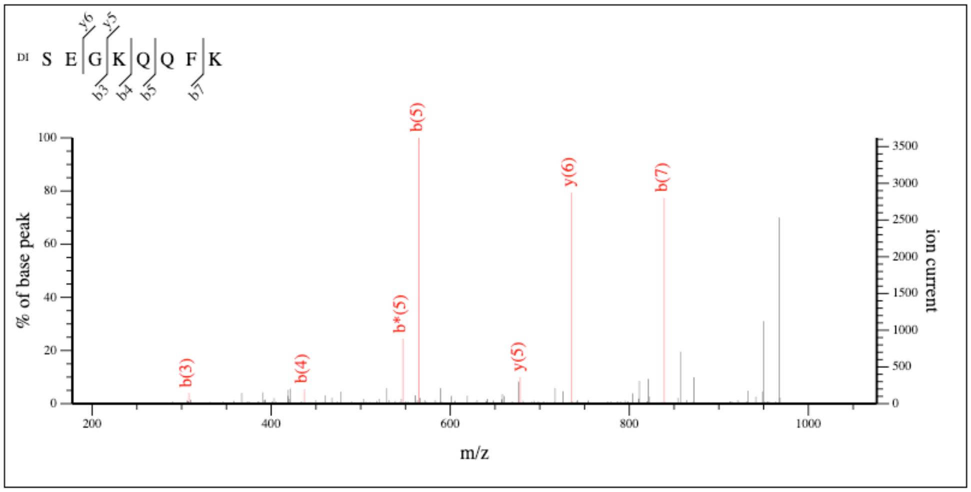

**Monoisotopic mass of neutral peptide Mr(calc):** 984.5511
**Fixed modifications:** Carbamidomethyl (C) (apply to specified residues or termini only)
**Variable modifications:**
**N-term :** Dimethyl:2H(6) (Protein N-term)
**Ions Score:** 26 **Expect:** 0.034
**Matches :** 8/33 fragment ions using 14 most intense peaks (help)

| # | b | b* | $b^0$ | Seq. | y | y* | $y^0$ | # |
|---|---|---|---|---|---|---|---|---|
| 1 | 122.1083 | | 104.0977 | S | | | | 8 |
| 2 | 251.1509 | | 233.1403 | E | 864.4574 | 847.4308 | 846.4468 | 7 |
| 3 | 308.1723 | | 290.1618 | G | 735.4148 | 718.3883 | | 6 |
| 4 | 436.2673 | 419.2407 | 418.2567 | K | 678.3933 | 661.3668 | | 5 |
| 5 | 564.3259 | 547.2993 | 546.3153 | Q | 550.2984 | 533.2718 | | 4 |
| 6 | 692.3844 | 675.3579 | 674.3739 | Q | 422.2398 | 405.2132 | | 3 |
| 7 | 839.4529 | 822.4263 | 821.4423 | F | 294.1812 | 277.1547 | | 2 |
| 8 | | | | K | 147.1128 | 130.0863 | | 1 |

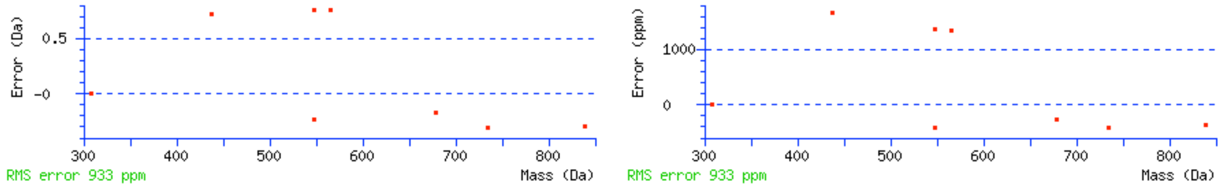


### 85. Osh2 (1 peptide in 450mM fraction 1)

MS/MS Fragmentation of **SREDLSIAEDLNQVSK**
Found in **Q12451** in **Uniprot_yeastS288C**, Oxysterol-binding protein homolog 2 OS=Saccharomyces cerevisiae (strain ATCC 204508 / S288c) OX=559292 GN=OSH2 PE=1 SV=1

Match to Query 10205: 1836.920872 from(919.467712,2+) intensity(342045.0625) rtinseconds(2430.2606) index(6208)
Title: 20100530_methyl450mM_1.6908.6908.2
Data file D:\Panyue\Proteomic dataset\20100530_methyl450mM_1.RAW

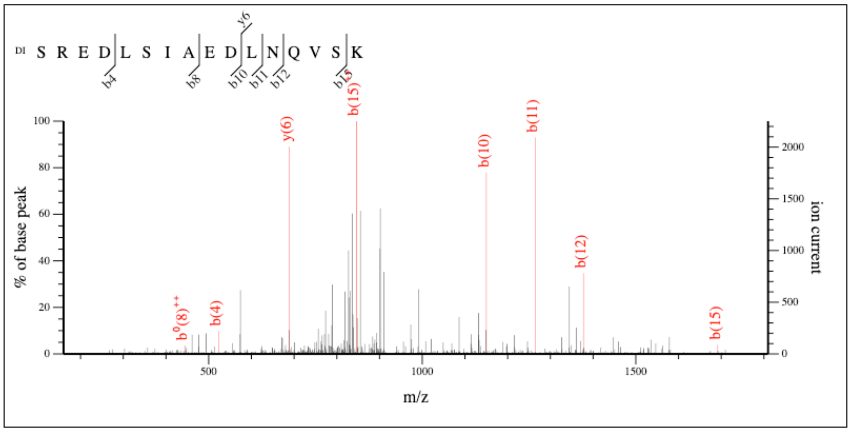

**Monoisotopic mass of neutral peptide Mr(calc):** 1836.9699
**Fixed modifications:** Carbamidomethyl (C) (apply to specified residues or termini only)
**Variable modifications:**
**N-term :** Dimethyl:2H(6) (Protein N-term)
**Ions Score:** 21 **Expect:** 0.12
**Matches :** 9/176 fragment ions using 15 most intense peaks (help)

| # | b | b++ | b* | b*++ | b0 | b0++ | Seq. | y | y++ | y* | y*++ | y0 | y0++ | # |
|---|---|---|---|---|---|---|---|---|---|---|---|---|---|---|
| 1 | 122.1083 | 61.5578 | | | 104.0977 | 52.5525 | S | | | | | | | 16 |
| 2 | 278.2094 | 139.6083 | 261.1828 | 131.0951 | 260.1988 | 130.6030 | R | 1716.8763 | 858.9418 | 1699.8497 | 850.4285 | 1698.8657 | 849.9365 | 15 |
| 3 | 407.2520 | 204.1296 | 390.2254 | 195.6163 | 389.2414 | 195.1243 | E | 1560.7752 | 780.8912 | 1543.7486 | 772.3779 | 1542.7646 | 771.8859 | 14 |
| 4 | 522.2789 | 261.6431 | 505.2524 | 253.1298 | 504.2683 | 252.6378 | D | 1431.7326 | 716.3699 | 1414.7060 | 707.8566 | 1413.7220 | 707.3646 | 13 |
| 5 | 635.3630 | 318.1851 | 618.3364 | 309.6719 | 617.3524 | 309.1798 | L | 1316.7056 | 658.8564 | 1299.6791 | 650.3432 | 1298.6951 | 649.8512 | 12 |
| 6 | 722.3950 | 361.7011 | 705.3685 | 353.1879 | 704.3844 | 352.6959 | S | 1203.6216 | 602.3144 | 1186.5950 | 593.8011 | 1185.6110 | 593.3091 | 11 |
| 7 | 835.4791 | 418.2432 | 818.4525 | 409.7299 | 817.4685 | 409.2379 | I | 1116.5895 | 558.7984 | 1099.5630 | 550.2851 | 1098.5790 | 549.7931 | 10 |
| 8 | 906.5162 | 453.7617 | 889.4896 | 445.2485 | 888.5056 | 444.7564 | A | 1003.5055 | 502.2564 | 986.4789 | 493.7431 | 985.4949 | 493.2511 | 9 |
| 9 | 1035.5588 | 518.2830 | 1018.5322 | 509.7698 | 1017.5482 | 509.2777 | E | 932.4684 | 466.7378 | 915.4418 | 458.2245 | 914.4578 | 457.7325 | 8 |
| 10 | 1150.5857 | 575.7965 | 1133.5592 | 567.2832 | 1132.5752 | 566.7912 | D | 803.4258 | 402.2165 | 786.3992 | 393.7032 | 785.4152 | 393.2112 | 7 |
| 11 | 1263.6698 | 632.3385 | 1246.6432 | 623.8253 | 1245.6592 | 623.3332 | L | 688.3988 | 344.7030 | 671.3723 | 336.1898 | 670.3882 | 335.6978 | 6 |
| 12 | 1377.7127 | 689.3600 | 1360.6862 | 680.8467 | 1359.7021 | 680.3547 | N | 575.3148 | 288.1610 | 558.2882 | 279.6477 | 557.3042 | 279.1557 | 5 |
| 13 | 1505.7713 | 753.3893 | 1488.7447 | 744.8760 | 1487.7607 | 744.3840 | Q | 461.2718 | 231.1396 | 444.2453 | 222.6263 | 443.2613 | 222.1343 | 4 |
| 14 | 1604.8397 | 802.9235 | 1587.8132 | 794.4102 | 1586.8291 | 793.9182 | V | 333.2132 | 167.1103 | 316.1867 | 158.5970 | 315.2027 | 158.1050 | 3 |
| 15 | 1691.8717 | 846.4395 | 1674.8452 | 837.9262 | 1673.8612 | 837.4342 | S | 234.1448 | 117.5761 | 217.1183 | 109.0628 | 216.1343 | 108.5708 | 2 |
| 16 | | | | | | | K | 147.1128 | 74.0600 | 130.0863 | 65.5468 | | | 1 |

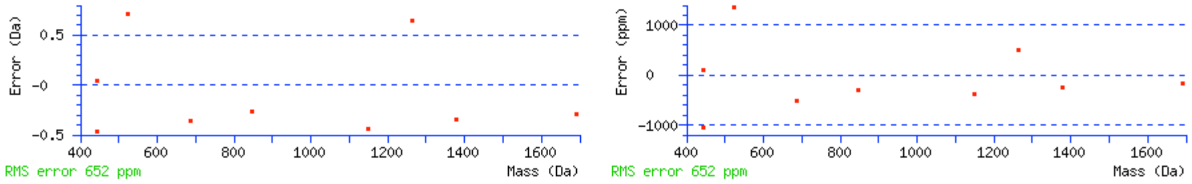


### 86. YNL247W (1 peptide in 450mM fraction 1)

MS/MS Fragmentation of **NIFIKALRR**
Found in **P53852** in **Uniprot_yeastS288C**, Cysteine--tRNA ligase OS=Saccharomyces cerevisiae (strain ATCC 204508 / S288c) OX=559292 GN=YNL247W PE=1 SV=1

Match to Query 5760: 1163.745448 from(582.880000,2+) intensity(3519.8865) rtinseconds(305.7753) index(751)
Title: 20100530_methyl450mM_1.837.837.
Data file D:\Panyue\Proteomic dataset\20100530_methyl450mM_1.RAW

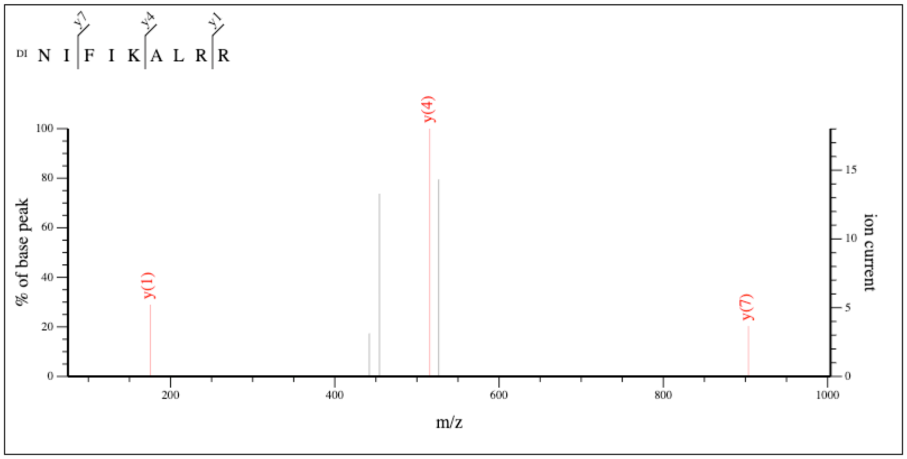

**Monoisotopic mass of neutral peptide Mr(calc):** 1163.7773
**Fixed modifications:** Carbamidomethyl (C) (apply to specified residues or termini only)
**Variable modifications:**
**N-term :** Dimethyl:2H(6) (Protein N-term)
**Ions Score:** 21 **Expect:** 0.027
**Matches :** 3/64 fragment ions using 4 most intense peaks (help)

| # | b | b++ | b* | b*++ | Seq. | y | y++ | y* | y*++ | # |
|---|---|---|---|---|---|---|---|---|---|---|
| 1 | 149.1192 | 75.0632 | 132.0926 | 66.5499 | N | | | | | 9 |
| 2 | 262.2032 | 131.6053 | 245.1767 | 123.0920 | I | 1016.6727 | 508.8400 | 999.6462 | 500.3267 | 8 |
| 3 | 409.2716 | 205.1395 | 392.2451 | 196.6262 | F | *903.5887* | 452.2980 | 886.5621 | 443.7847 | 7 |
| 4 | 522.3557 | 261.6815 | 505.3292 | 253.1682 | I | 756.5203 | 378.7638 | 739.4937 | 370.2505 | 6 |
| 5 | 650.4507 | 325.7290 | 633.4241 | 317.2157 | K | 643.4362 | 322.2217 | 626.4097 | 313.7085 | 5 |
| 6 | 721.4878 | 361.2475 | 704.4612 | 352.7343 | A | *515.3412* | 258.1743 | 498.3147 | 249.6610 | 4 |
| 7 | 834.5718 | 417.7896 | 817.5453 | 409.2763 | L | 444.3041 | 222.6557 | 427.2776 | 214.1424 | 3 |
| 8 | 990.6730 | 495.8401 | 973.6464 | 487.3268 | R | 331.2201 | 166.1137 | 314.1935 | 157.6004 | 2 |
| 9 | | | | | R | *175.1190* | 88.0631 | 158.0924 | 79.5498 | 1 |

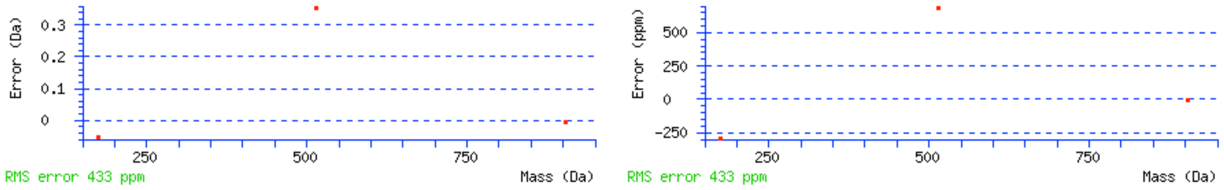


### 87. Rps2 (1 peptide in 450mM fraction 2)

MS/MS Fragmentation of **SAPEAQQQKR**
Found in **P25443** in **Uniprot_yeastS288C**, 40S ribosomal protein S2 OS=Saccharomyces cerevisiae (strain ATCC 204508 / S288c) OX=559292 GN=RPS2 PE=1 SV=3

Match to Query 5729: 1158.605448 from(580.310000,2+) intensity(2055.4792) rtinseconds(520.6008) index(1260)
Title: 20100530_methyl450mM_2.1404.1404.
Data file D:\Panyue\Proteomic dataset\20100530_methyl450mM_2.RAW

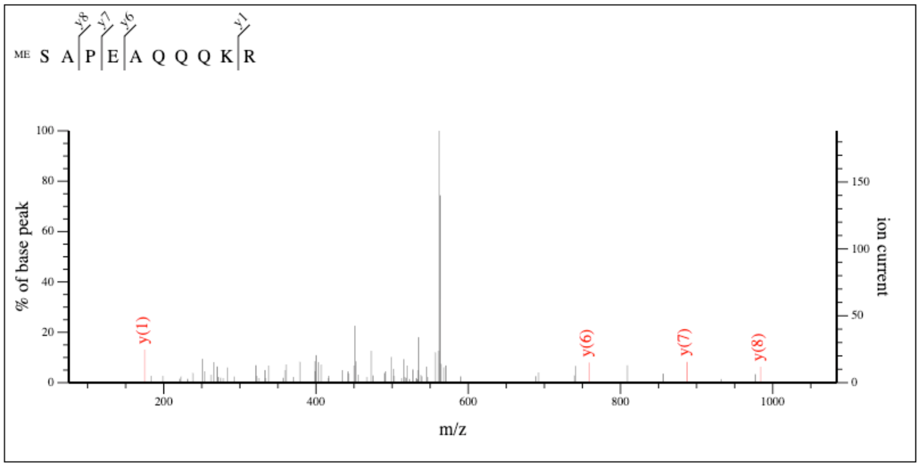

**Monoisotopic mass of neutral peptide Mr(calc):** 1158.6184
**Fixed modifications:** Carbamidomethyl (C) (apply to specified residues or termini only)
**Variable modifications:**
**N-term :** Methyl:2H(3) (Protein N-term)
**Ions Score:** 21 **Expect:** 0.16
**Matches :** 4/86 fragment ions using 9 most intense peaks (help)

| # | b | b++ | b* | b*++ | b0 | b0++ | Seq. | y | y++ | y* | y*++ | y0 | y0++ | # |
|---|---|---|---|---|---|---|---|---|---|---|---|---|---|---|
| 1 | 105.0738 | 53.0405 | | | 87.0632 | 44.0352 | S | | | | | | | 10 |
| 2 | 176.1109 | 88.5591 | | | 158.1003 | 79.5538 | A | 1055.5592 | 528.2833 | 1038.5327 | 519.7700 | 1037.5487 | 519.2780 | 9 |
| 3 | 273.1637 | 137.0855 | | | 255.1531 | 128.0802 | P | ***984.5221*** | 492.7647 | 967.4956 | 484.2514 | 966.5116 | 483.7594 | 8 |
| 4 | 402.2063 | 201.6068 | | | 384.1957 | 192.6015 | E | ***887.4694*** | 444.2383 | 870.4428 | 435.7250 | 869.4588 | 435.2330 | 7 |
| 5 | 473.2434 | 237.1253 | | | 455.2328 | 228.1200 | A | ***758.4268*** | 379.7170 | 741.4002 | 371.2037 | | | 6 |
| 6 | 601.3019 | 301.1546 | 584.2754 | 292.6413 | 583.2914 | 292.1493 | Q | 687.3896 | 344.1985 | 670.3631 | 335.6852 | | | 5 |
| 7 | 729.3605 | 365.1839 | 712.3340 | 356.6706 | 711.3500 | 356.1786 | Q | 559.3311 | 280.1692 | 542.3045 | 271.6559 | | | 4 |
| 8 | 857.4191 | 429.2132 | 840.3926 | 420.6999 | 839.4085 | 420.2079 | Q | 431.2725 | 216.1399 | 414.2459 | 207.6266 | | | 3 |
| 9 | 985.5141 | 493.2607 | 968.4875 | 484.7474 | 967.5035 | 484.2554 | K | 303.2139 | 152.1106 | 286.1874 | 143.5973 | | | 2 |
| 10 | | | | | | | R | ***175.1190*** | 88.0631 | 158.0924 | 79.5498 | | | 1 |

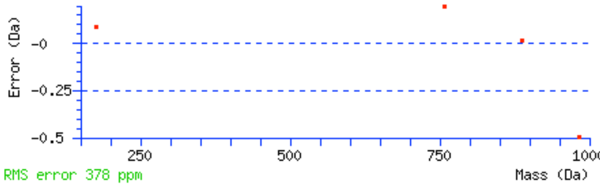


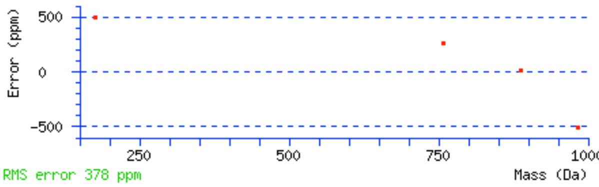


### 88. YLR171W (1 peptide in 450mM fraction 2)

MS/MS Fragmentation of **SSSQTLPK**
Found in **O13562** in **Uniprot_yeastS288C**, Putative uncharacterized protein YLR171W OS=Saccharomyces cerevisiae (strain ATCC 204508 / S288c) OX=559292 GN=YLR171W PE=5 SV=1

Match to Query 3541: 897.565526 from(449.790039,2+) intensity(39602.1641) rtinseconds(838.2299) index(2031)
Title: 20100530_methyl450mM_2.2262.2262.2
Data file D:\Panyue\Proteomic dataset\20100530_methyl450mM_2.RAW

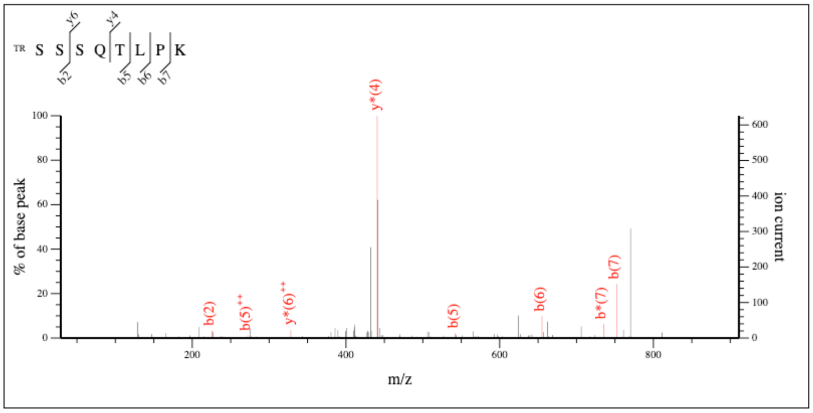

Label all possible matches ○ Label matches used for scoring ◉

**Monoisotopic mass of neutral peptide Mr(calc):** 897.5481
**Fixed modifications:** Carbamidomethyl (C) (apply to specified residues or termini only)
**Variable modifications:**
**N-term :** Trimethyl:2H(9) (Protein N-term)
**Ions Score:** 27 **Expect:** 0.012
**Matches :** 13/72 fragment ions using 21 most intense peaks (help)

| # | b | $b^{++}$ | b* | $b^{*++}$ | $b^0$ | $b^{0++}$ | Seq. | y | $y^{++}$ | y* | $y^{*++}$ | $y^0$ | $y^{0++}$ | # |
|---|---|---|---|---|---|---|---|---|---|---|---|---|---|---|
| 1 | 139.1427 | 70.0750 | | | 121.1322 | 61.0697 | S | | | | | | | 8 |
| 2 | ***226.1748*** | 113.5910 | | | 208.1642 | 104.5857 | S | 760.4199 | 380.7136 | 743.3934 | 372.2003 | 742.4094 | 371.7083 | 7 |
| 3 | 313.2068 | 157.1070 | | | 295.1962 | 148.1018 | S | 673.3879 | 337.1976 | 656.3614 | 328.6843 | 655.3774 | 328.1923 | 6 |
| 4 | ***441.2654*** | 221.1363 | 424.2388 | 212.6231 | 423.2548 | 212.1310 | Q | 586.3559 | 293.6816 | 569.3293 | 285.1683 | 568.3453 | 284.6763 | 5 |
| 5 | ***542.3131*** | **271.6602** | 525.2865 | 263.1469 | 524.3025 | 262.6549 | T | 458.2973 | 229.6523 | 441.2708 | 221.1390 | 440.2867 | 220.6470 | 4 |
| 6 | ***655.3971*** | **328.2022** | 638.3706 | 319.6889 | 637.3866 | 319.1969 | L | 357.2496 | 179.1285 | 340.2231 | 170.6152 | | | 3 |
| 7 | ***752.4499*** | 376.7286 | 735.4233 | 368.2153 | 734.4393 | 367.7233 | P | 244.1656 | 122.5864 | 227.1390 | 114.0731 | | | 2 |
| 8 | | | | | | | K | 147.1128 | 74.0600 | 130.0863 | 65.5468 | | | 1 |

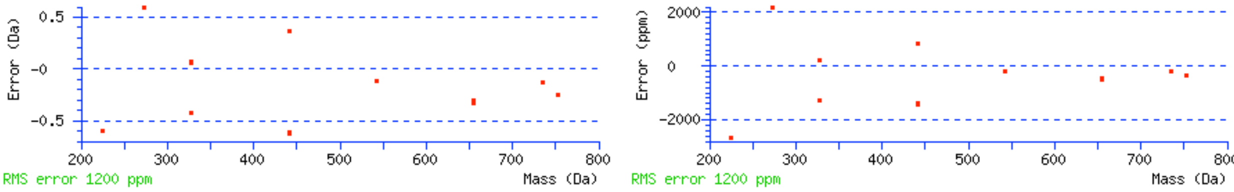


### 89. Pdc5 (2 peptides in 450mM fraction 2)

MS/MS Fragmentation of **SEITLGK**
Found in **P16467** in **Uniprot_yeastS288C**, Pyruvate decarboxylase isozyme 2 OS=Saccharomyces cerevisiae (strain ATCC 204508 / S288c) OX=559292 GN=PDC5 PE=1 SV=4

Match to Query 2306: 780.449999 from(781.457275,1+) intensity(12283579.0000) rtinseconds(1056.5849) index(2589)
Title: 20100530_methyl450mM_2.2883.2883.1
Data file D:\Panyue\Proteomic dataset\20100530_methyl450mM_2.RAW

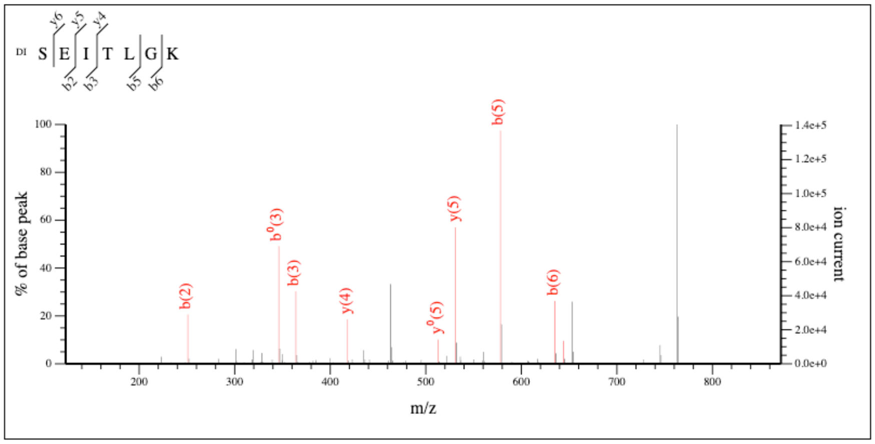

**Monoisotopic mass of neutral peptide Mr(calc):** 780.4864
**Fixed modifications:** Carbamidomethyl (C) (apply to specified residues or termini only)
**Variable modifications:**
**N-term :** Dimethyl:2H(6) (Protein N-term)
**Ions Score:** 27 **Expect:** 0.026
**Matches :** 9/27 fragment ions using 18 most intense peaks (help)

| # | b | $b^0$ | Seq. | y | y* | $y^0$ | # |
|---|---|---|---|---|---|---|---|
| 1 | 122.1083 | 104.0977 | S | | | | 7 |
| 2 | ***251.1509*** | 233.1403 | E | 660.3927 | 643.3661 | 642.3821 | 6 |
| 3 | ***364.2349*** | 346.2244 | I | **531.3501** | 514.3235 | 513.3395 | 5 |
| 4 | 465.2826 | 447.2720 | T | **418.2660** | 401.2395 | 400.2554 | 4 |
| 5 | ***578.3667*** | 560.3561 | L | 317.2183 | 300.1918 | | 3 |
| 6 | ***635.3881*** | 617.3776 | G | 204.1343 | 187.1077 | | 2 |
| 7 | | | K | 147.1128 | 130.0863 | | 1 |

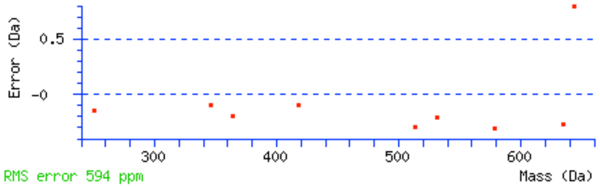


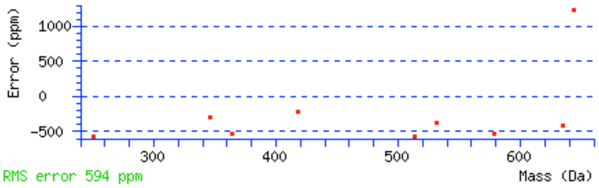


MS/MS Fragmentation of **SEITLGK**
Found in **P16467** in **Uniprot_yeastS288C**, Pyruvate decarboxylase isozyme 2 OS=Saccharomyces cerevisiae (strain ATCC 204508 / S288c) OX=559292 GN=PDC5 PE=1 SV=4

Match to Query 2305: 780.449938 from(781.457214,1+) intensity(11503263.0000) rtinseconds(1059.8771) index(2598)
Title: 20100530_methyl450mM_2.2893.2893.1
Data file D:\Panyue\Proteomic dataset\20100530_methyl450mM_2.RAW

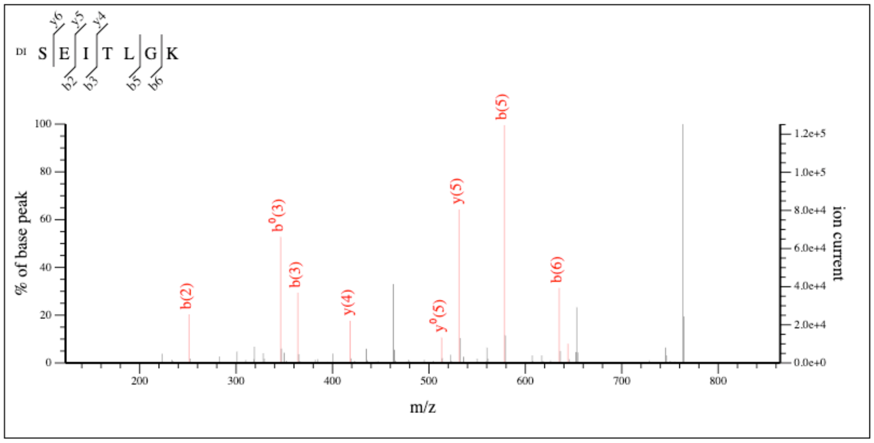

**Monoisotopic mass of neutral peptide Mr(calc):** 780.4864
**Fixed modifications:** Carbamidomethyl (C) (apply to specified residues or termini only)
**Variable modifications:**
**N-term :** Dimethyl:2H(6) (Protein N-term)
**Ions Score:** 27 **Expect:** 0.025
**Matches :** 9/27 fragment ions using 18 most intense peaks (help)

| # | b | b0 | Seq. | y | y* | y0 | # |
|---|---|---|---|---|---|---|---|
| 1 | 122.1083 | 104.0977 | S | | | | 7 |
| 2 | 251.1509 | 233.1403 | E | 660.3927 | 643.3661 | 642.3821 | 6 |
| 3 | 364.2349 | 346.2244 | I | 531.3501 | 514.3235 | 513.3395 | 5 |
| 4 | 465.2826 | 447.2720 | T | 418.2660 | 401.2395 | 400.2554 | 4 |
| 5 | 578.3667 | 560.3561 | L | 317.2183 | 300.1918 | | 3 |
| 6 | 635.3881 | 617.3776 | G | 204.1343 | 187.1077 | | 2 |
| 7 | | | K | 147.1128 | 130.0863 | | 1 |

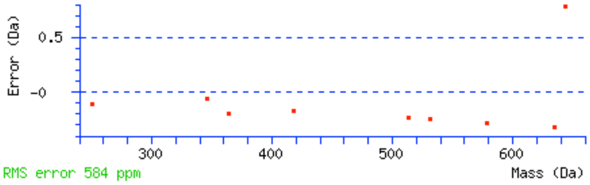


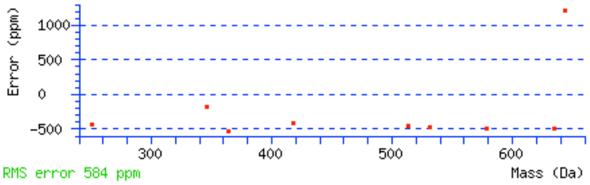


### 90. Pdc1 (2 peptides in 450mM fraction 2)

MS/MS Fragmentation of **SEITLGK**
Found in **P06169** in **Uniprot_yeastS288C**, Pyruvate decarboxylase isozyme 1 OS=Saccharomyces cerevisiae (strain ATCC 204508 / S288c) OX=559292 GN=PDC1 PE=1 SV=7

Match to Query 2306: 780.449999 from(781.457275,1+) intensity(12283579.0000) rtinseconds(1056.5849) index(2589)
Title: 20100530_methyl450mM_2.2883.2883.1
Data file D:\Panyue\Proteomic dataset\20100530_methyl450mM_2.RAW

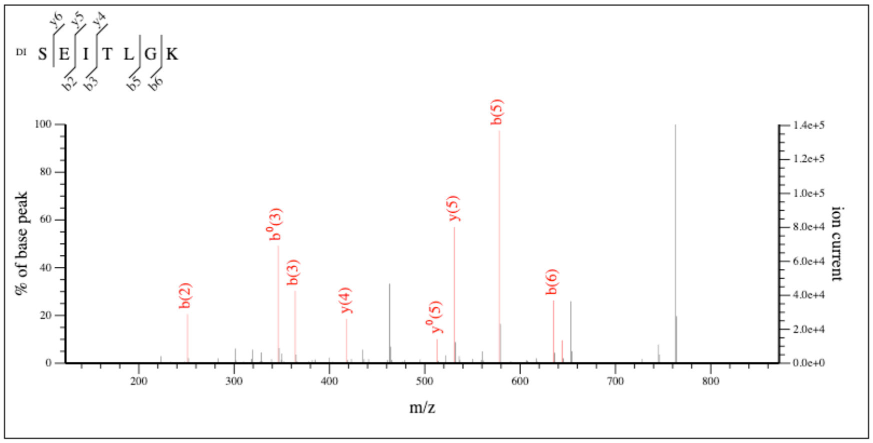

**Monoisotopic mass of neutral peptide Mr(calc):** 780.4864
**Fixed modifications:** Carbamidomethyl (C) (apply to specified residues or termini only)
**Variable modifications:**
**N-term :** Dimethyl:2H(6) (Protein N-term)
**Ions Score:** 27 **Expect:** 0.026
**Matches :** 9/27 fragment ions using 18 most intense peaks (help)

| # | b | b⁰ | Seq. | y | y* | y⁰ | # |
|---|---|---|---|---|---|---|---|
| 1 | 122.1083 | 104.0977 | S | | | | 7 |
| 2 | ***251.1509*** | 233.1403 | E | 660.3927 | 643.3661 | 642.3821 | 6 |
| 3 | ***364.2349*** | 346.2244 | I | **531.3501** | 514.3235 | 513.3395 | 5 |
| 4 | 465.2826 | 447.2720 | T | **418.2660** | 401.2395 | 400.2554 | 4 |
| 5 | ***578.3667*** | 560.3561 | L | 317.2183 | 300.1918 | | 3 |
| 6 | ***635.3881*** | 617.3776 | G | 204.1343 | 187.1077 | | 2 |
| 7 | | | K | 147.1128 | 130.0863 | | 1 |

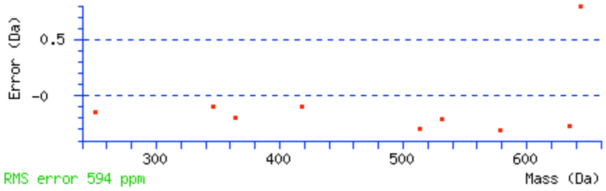


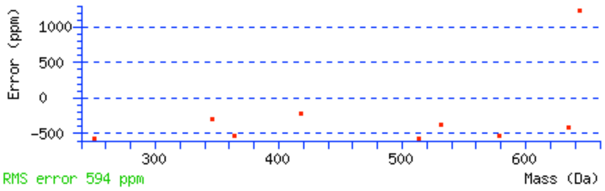


MS/MS Fragmentation of **SEITLGK**
Found in **P06169** in **Uniprot_yeastS288C**, Pyruvate decarboxylase isozyme 1 OS=Saccharomyces cerevisiae (strain ATCC 204508 / S288c) OX=559292 GN=PDC1 PE=1 SV=7

Match to Query 2305: 780.449938 from(781.457214,1+) intensity(11503263.0000) rtinseconds(1059.8771) index(2598)
Title: 20100530_methyl450mM_2.2893.2893.1
Data file D:\Panyue\Proteomic dataset\20100530_methyl450mM_2.RAW

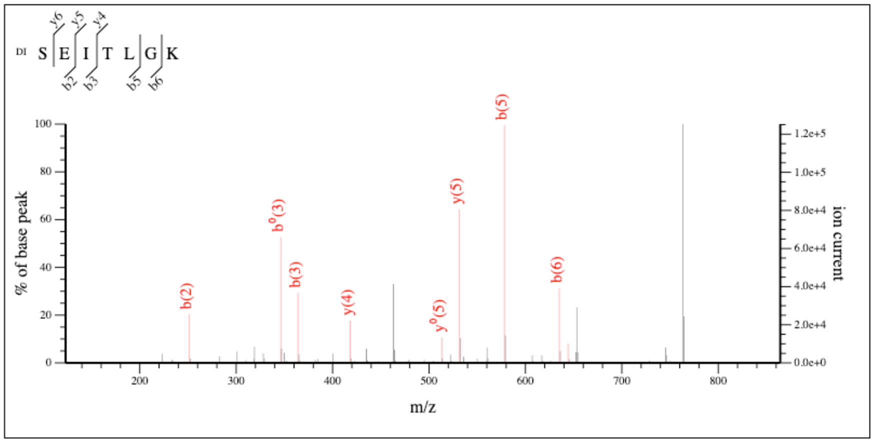

**Monoisotopic mass of neutral peptide Mr(calc):** 780.4864
**Fixed modifications:** Carbamidomethyl (C) (apply to specified residues or termini only)
**Variable modifications:**
**N-term :** Dimethyl:2H(6) (Protein N-term)
**Ions Score:** 27 **Expect:** 0.025
**Matches :** 9/27 fragment ions using 18 most intense peaks (help)

| # | b | $b^0$ | Seq. | y | y* | $y^0$ | # |
|---|---|---|---|---|---|---|---|
| 1 | 122.1083 | 104.0977 | S | | | | 7 |
| 2 | 251.1509 | 233.1403 | E | 660.3927 | 643.3661 | 642.3821 | 6 |
| 3 | 364.2349 | 346.2244 | I | 531.3501 | 514.3235 | 513.3395 | 5 |
| 4 | 465.2826 | 447.2720 | T | 418.2660 | 401.2395 | 400.2554 | 4 |
| 5 | 578.3667 | 560.3561 | L | 317.2183 | 300.1918 | | 3 |
| 6 | 635.3881 | 617.3776 | G | 204.1343 | 187.1077 | | 2 |
| 7 | | | K | 147.1128 | 130.0863 | | 1 |

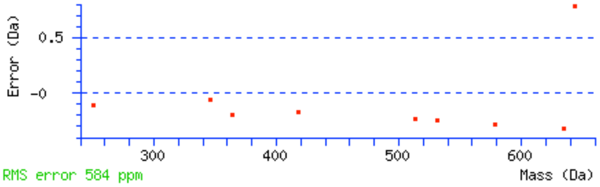


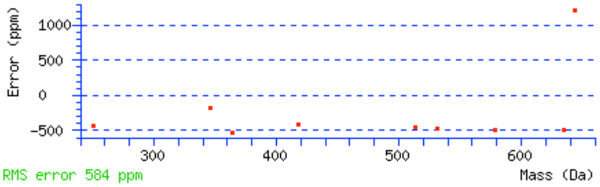


91. Fzo1 (2 peptides in 450mM fraction 2)

MS/MS Fragmentation of **SEGKQQFK**
Found in **P38297** in **Uniprot_yeastS288C**, Mitofusin FZO1 OS=Saccharomyces cerevisiae (strain ATCC 204508 / S288c) OX=559292 GN=FZO1 PE=1 SV=1

Match to Query 4317: 984.538976 from(493.276764,2+) intensity(881398.3750) rtinseconds(1642.1013) index(4108)
Title: 20100530_methyl450mM_2.4574.4574.2
Data file D:\Panyue\Proteomic dataset\20100530_methyl450mM_2.RAW

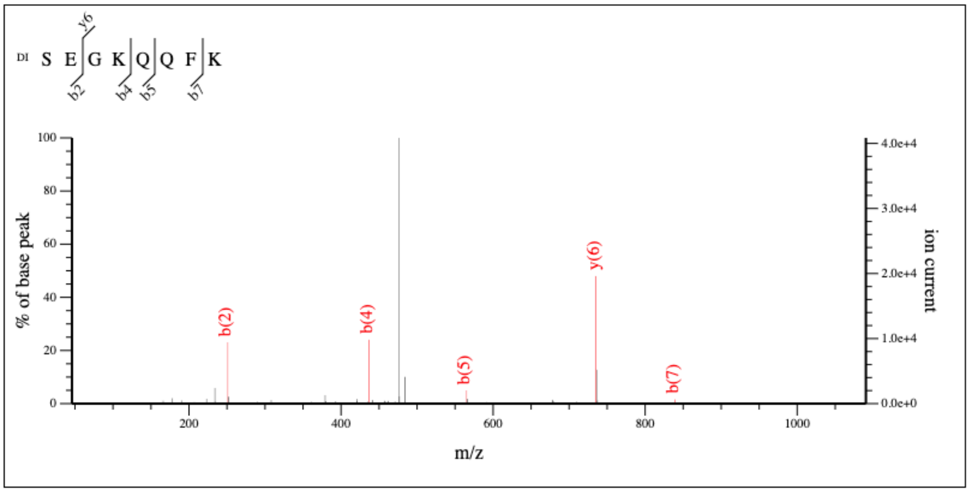

**Monoisotopic mass of neutral peptide Mr(calc):** 984.5511
**Fixed modifications:** Carbamidomethyl (C) (apply to specified residues or termini only)
**Variable modifications:**
**N-term :** Dimethyl:2H(6) (Protein N-term)
**Ions Score:** 27 **Expect:** 0.032
**Matches :** 5/66 fragment ions using 9 most intense peaks (help)

| # | b | b++ | b* | b*++ | b0 | b0++ | Seq. | y | y++ | y* | y*++ | y0 | y0++ | # |
|---|---|---|---|---|---|---|---|---|---|---|---|---|---|---|
| 1 | 122.1083 | 61.5578 | | | 104.0977 | 52.5525 | S | | | | | | | 8 |
| 2 | ***251.1509*** | 126.0791 | | | 233.1403 | 117.0738 | E | 864.4574 | 432.7323 | 847.4308 | 424.2191 | 846.4468 | 423.7271 | 7 |
| 3 | 308.1723 | 154.5898 | | | 290.1618 | 145.5845 | G | 735.4148 | 368.2110 | 718.3883 | 359.6978 | | | 6 |
| 4 | ***436.2673*** | 218.6373 | 419.2407 | 210.1240 | 418.2567 | 209.6320 | K | 678.3933 | 339.7003 | 661.3668 | 331.1870 | | | 5 |
| 5 | ***564.3259*** | 282.6666 | 547.2993 | 274.1533 | 546.3153 | 273.6613 | Q | 550.2984 | 275.6528 | 533.2718 | 267.1396 | | | 4 |
| 6 | 692.3844 | 346.6959 | 675.3579 | 338.1826 | 674.3739 | 337.6906 | Q | 422.2398 | 211.6235 | 405.2132 | 203.1103 | | | 3 |
| 7 | ***839.4529*** | 420.2301 | 822.4263 | 411.7168 | 821.4423 | 411.2248 | F | 294.1812 | 147.5942 | 277.1547 | 139.0810 | | | 2 |
| 8 | | | | | | | K | 147.1128 | 74.0600 | 130.0863 | 65.5468 | | | 1 |

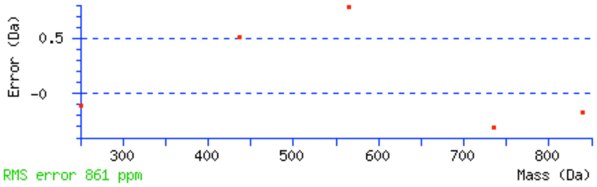


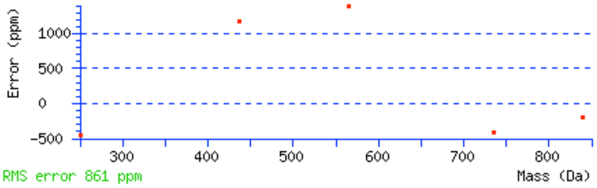


MS/MS Fragmentation of **SEGKQQFK**
Found in **P38297** in **Uniprot_yeastS288C**, Mitofusin FZO1 OS=Saccharomyces cerevisiae (strain ATCC 204508 / S288c) OX=559292 GN=FZO1 PE=1 SV=1

Match to Query 4319: 984.540209 from(985.547485,1+) intensity(146440.6406) rtinseconds(1651.4468) index(4134)
Title: 20100530_methyl450mM_2.4602.4602.1
Data file D:\Panyue\Proteomic dataset\20100530_methyl450mM_2.RAW

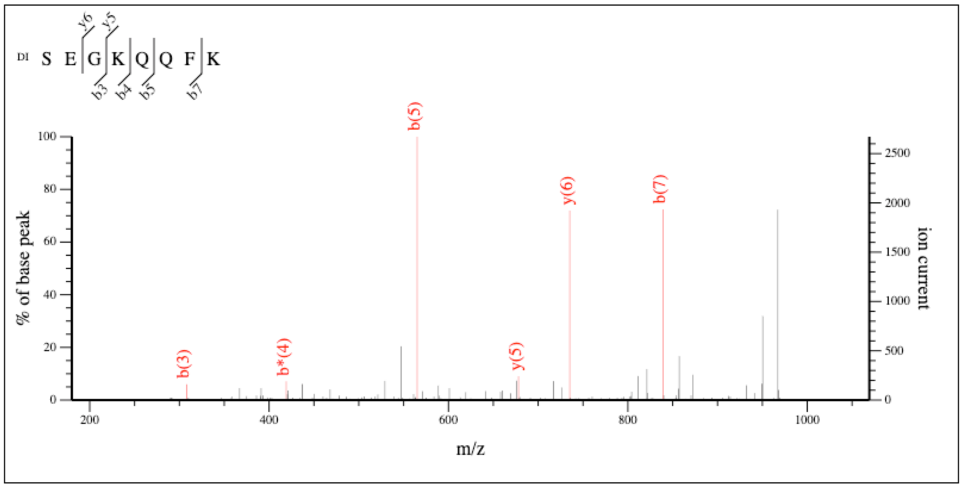

**Monoisotopic mass of neutral peptide Mr(calc):** 984.5511
**Fixed modifications:** Carbamidomethyl (C) (apply to specified residues or termini only)
**Variable modifications:**
**N-term :** Dimethyl:2H(6) (Protein N-term)
**Ions Score:** 30 **Expect:** 0.016
**Matches :** 6/33 fragment ions using 7 most intense peaks (help)

| # | b | b* | $b^0$ | Seq. | y | y* | $y^0$ | # |
|---|---|---|---|---|---|---|---|---|
| 1 | 122.1083 | | 104.0977 | S | | | | 8 |
| 2 | 251.1509 | | 233.1403 | E | 864.4574 | 847.4308 | 846.4468 | 7 |
| 3 | ***308.1723*** | | 290.1618 | G | ***735.4148*** | 718.3883 | | 6 |
| 4 | 436.2673 | 419.2407 | 418.2567 | K | ***678.3933*** | 661.3668 | | 5 |
| 5 | ***564.3259*** | 547.2993 | 546.3153 | Q | 550.2984 | 533.2718 | | 4 |
| 6 | 692.3844 | 675.3579 | 674.3739 | Q | 422.2398 | 405.2132 | | 3 |
| 7 | ***839.4529*** | 822.4263 | 821.4423 | F | 294.1812 | 277.1547 | | 2 |
| 8 | | | | K | 147.1128 | 130.0863 | | 1 |

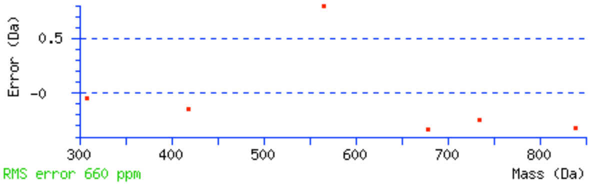


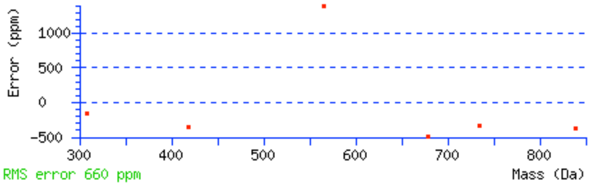


### 92. Atg23 (2 peptides in 450mM fraction2)

MS/MS Fragmentation of **ELNQVLEK**
Found in **Q06671** in **Uniprot_yeastS288C**, Autophagy-related protein 23 OS=Saccharomyces cerevisiae (strain ATCC 204508 / S288c) OX=559292 GN=ATG23 PE=1 SV=1

Match to Query 4332: 988.571020 from(495.292786,2+) intensity(306389.5000) rtinseconds(2012.664) index(5076)
Title: 20100530_methyl450mM_2.5650.5650.2
Data file D:\Panyue\Proteomic dataset\20100530_methyl450mM_2.RAW

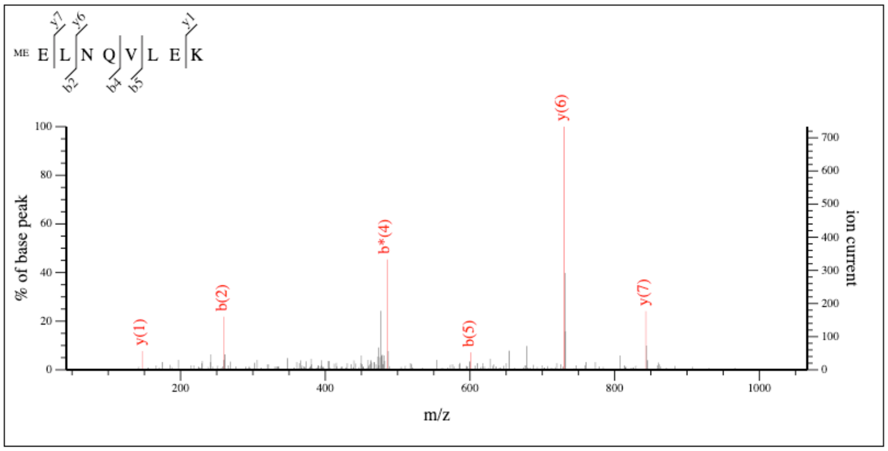

**Monoisotopic mass of neutral peptide Mr(calc):** 988.5632
**Fixed modifications:** Carbamidomethyl (C) (apply to specified residues or termini only)
**Variable modifications:**
**N-term :** Methyl:2H(3) (Protein N-term)
**Ions Score:** 27 **Expect:** 0.033
**Matches :** 8/78 fragment ions using 9 most intense peaks (help)

| # | b | b++ | b* | b*++ | b0 | b0++ | Seq. | y | y++ | y* | y*++ | y0 | y0++ | # |
|---|---|---|---|---|---|---|---|---|---|---|---|---|---|---|
| 1 | ***147.0843*** | 74.0458 | | | 129.0738 | 65.0405 | E | | | | | | | 8 |
| 2 | ***260.1684*** | 130.5878 | | | 242.1578 | 121.5826 | L | **843.4934** | 422.2504 | 826.4669 | 413.7371 | 825.4829 | 413.2451 | 7 |
| 3 | 374.2113 | 187.6093 | 357.1848 | 179.0960 | 356.2008 | 178.6040 | N | **730.4094** | 365.7083 | 713.3828 | 357.1951 | 712.3988 | 356.7030 | 6 |
| 4 | 502.2699 | 251.6386 | 485.2434 | 243.1253 | 484.2594 | 242.6333 | Q | 616.3665 | 308.6869 | 599.3399 | 300.1736 | 598.3559 | 299.6816 | 5 |
| 5 | ***601.3383*** | 301.1728 | 584.3118 | 292.6595 | 583.3278 | 292.1675 | V | 488.3079 | 244.6576 | 471.2813 | 236.1443 | 470.2973 | 235.6523 | 4 |
| 6 | 714.4224 | 357.7148 | 697.3958 | 349.2016 | 696.4118 | 348.7096 | L | 389.2395 | 195.1234 | 372.2129 | 186.6101 | 371.2289 | 186.1181 | 3 |
| 7 | ***843.4650*** | 422.2361 | 826.4384 | 413.7229 | 825.4544 | 413.2309 | E | 276.1554 | 138.5813 | 259.1288 | 130.0681 | 258.1448 | 129.5761 | 2 |
| 8 | | | | | | | K | **147.1128** | 74.0600 | 130.0863 | 65.5468 | | | 1 |

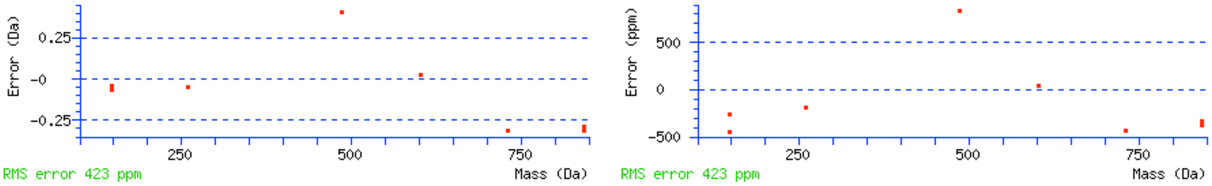


MS/MS Fragmentation of **ELNQVLEK**
Found in **Q06671** in **Uniprot_yeastS288C**, Autophagy-related protein 23 OS=Saccharomyces cerevisiae (strain ATCC 204508 / S288c) OX=559292 GN=ATG23 PE=1 SV=1

Match to Query 4331: 988.570286 from(495.292419,2+) intensity(405435.7812) rtinseconds(2014.9292) index(5081)
Title: 20100530_methyl450mM_2.5656.5656.2
Data file D:\Panyue\Proteomic dataset\20100530_methyl450mM_2.RAW

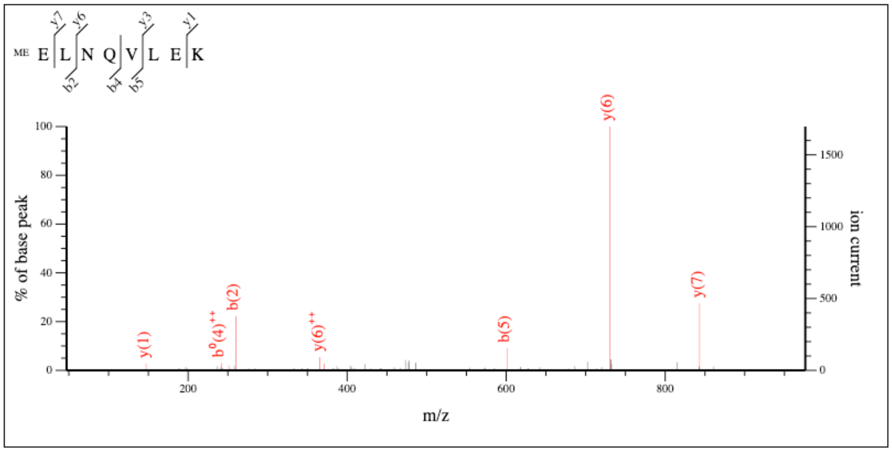

**Monoisotopic mass of neutral peptide Mr(calc):** 988.5632
**Fixed modifications:** Carbamidomethyl (C) (apply to specified residues or termini only)
**Variable modifications:**
**N-term :** Methyl:2H(3) (Protein N-term)
**Ions Score:** 22 **Expect:** 0.096
**Matches :** 12/78 fragment ions using 16 most intense peaks (help)

| # | b | b++ | b* | b*++ | b0 | b0++ | Seq. | y | y++ | y* | y*++ | y0 | y0++ | # |
|---|---|---|---|---|---|---|---|---|---|---|---|---|---|---|
| 1 | ***147.0843*** | 74.0458 | | | 129.0738 | 65.0405 | E | | | | | | | 8 |
| 2 | ***260.1684*** | 130.5878 | | | 242.1578 | 121.5826 | L | **843.4934** | 422.2504 | 826.4669 | 413.7371 | 825.4829 | 413.2451 | 7 |
| 3 | 374.2113 | 187.6093 | 357.1848 | 179.0960 | 356.2008 | 178.6040 | N | **730.4094** | 365.7083 | 713.3828 | 357.1951 | 712.3988 | 356.7030 | 6 |
| 4 | 502.2699 | 251.6386 | 485.2434 | 243.1253 | 484.2594 | 242.6333 | Q | 616.3665 | 308.6869 | 599.3399 | 300.1736 | 598.3559 | 299.6816 | 5 |
| 5 | ***601.3383*** | 301.1728 | 584.3118 | 292.6595 | 583.3278 | 292.1675 | V | 488.3079 | 244.6576 | 471.2813 | 236.1443 | 470.2973 | 235.6523 | 4 |
| 6 | 714.4224 | 357.7148 | 697.3958 | 349.2016 | 696.4118 | 348.7096 | L | 389.2395 | 195.1234 | 372.2129 | 186.6101 | 371.2289 | 186.1181 | 3 |
| 7 | ***843.4650*** | 422.2361 | 826.4384 | 413.7229 | 825.4544 | 413.2309 | E | 276.1554 | 138.5813 | 259.1288 | 130.0681 | 258.1448 | 129.5761 | 2 |
| 8 | | | | | | | K | **147.1128** | 74.0600 | 130.0863 | 65.5468 | | | 1 |

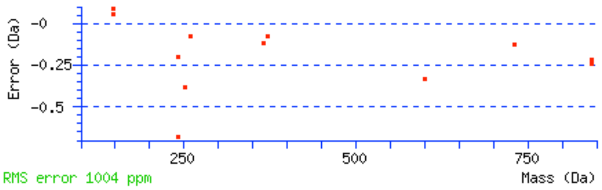


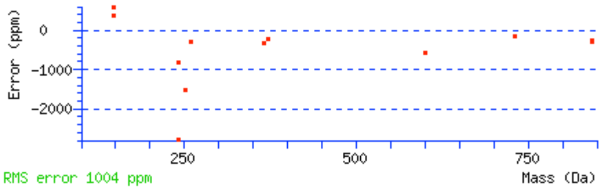


### 93. Ahp1 (1 peptide in 450mM fraction 2)

MS/MS Fragmentation of **SDLVNK**
Found in **P38013** in **Uniprot_yeastS288C**, Peroxiredoxin AHP1 OS=Saccharomyces cerevisiae (strain ATCC 204508 / S288c) OX=559292 GN=AHP1 PE=1 SV=4

Match to Query 1889: 691.405993 from(692.413269,1+) intensity(341576.5000) rtinseconds(2583.5414) index(6566)
Title: 20100530_methyl450mM_2.7307.7307.1
Data file D:\Panyue\Proteomic dataset\20100530_methyl450mM_2.RAW

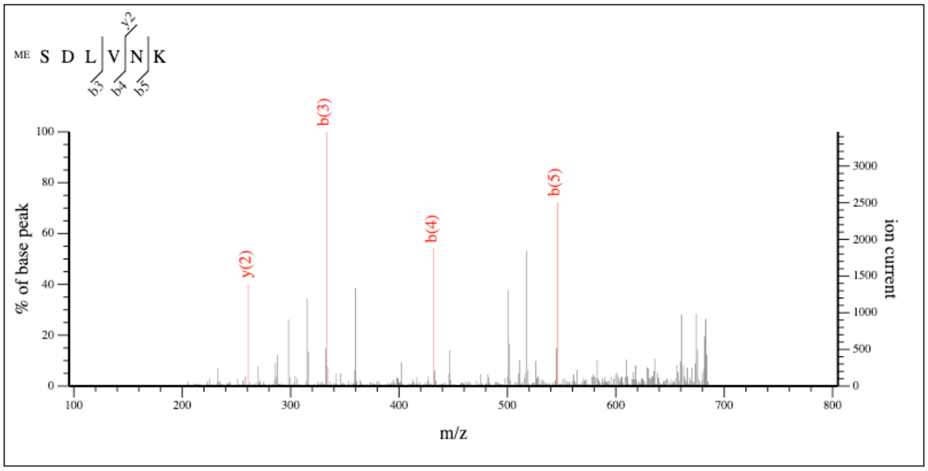

**Monoisotopic mass of neutral peptide Mr(calc):** 691.3944
**Fixed modifications:** Carbamidomethyl (C) (apply to specified residues or termini only)
**Variable modifications:**
**N-term :** Methyl:2H(3) (Protein N-term)
**Ions Score:** 22 **Expect:** 0.079
**Matches :** 4/22 fragment ions using 5 most intense peaks (help)

| # | b | b* | b0 | Seq. | y | y* | y0 | # |
|---|---|---|---|---|---|---|---|---|
| 1 | 105.0738 | | 87.0632 | S | | | | 6 |
| 2 | 220.1007 | | 202.0902 | D | 588.3352 | 571.3086 | 570.3246 | 5 |
| 3 | ***333.1848*** | | 315.1742 | L | 473.3082 | 456.2817 | | 4 |
| 4 | ***432.2532*** | | 414.2426 | V | 360.2241 | 343.1976 | | 3 |
| 5 | ***546.2961*** | 529.2696 | 528.2856 | N | 261.1557 | 244.1292 | | 2 |
| 6 | | | | K | 147.1128 | 130.0863 | | 1 |

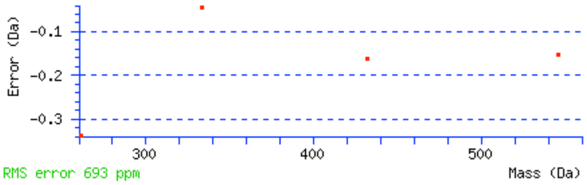


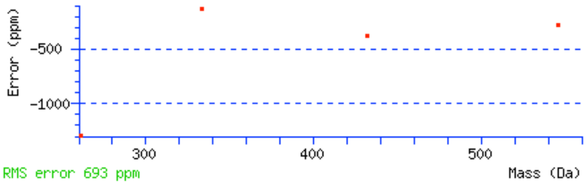


### 94. Pdc5 (2 peptides in 450mM fraction 3)

MS/MS Fragmentation of **SEITLGK**
Found in **P16467** in **Uniprot_yeastS288C**, Pyruvate decarboxylase isozyme 2 OS=Saccharomyces cerevisiae (strain ATCC 204508 / S288c) OX=559292 GN=PDC5 PE=1 SV=4

Match to Query 2323: 780.450549 from(781.457825,1+) intensity(12241962.0000) rtinseconds(1055.6165) index(2554)
Title: 20100530_methyl450mM_3.2845.2845.1
Data file D:\Panyue\Proteomic dataset\20100530_methyl450mM_3.RAW

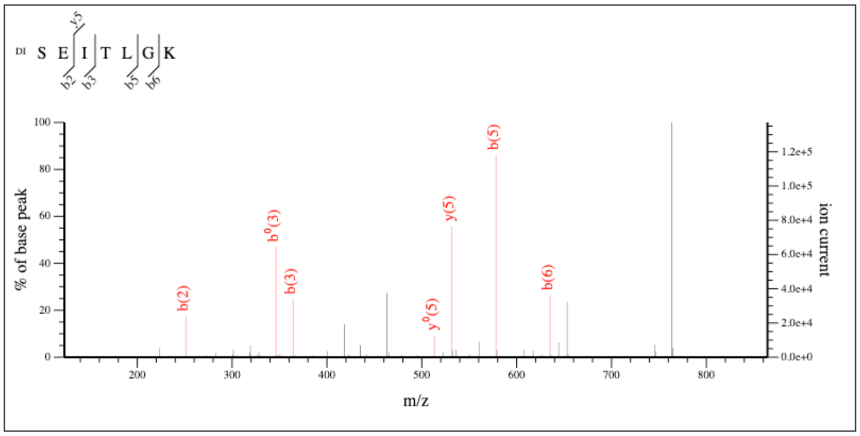

**Monoisotopic mass of neutral peptide Mr(calc):** 780.4864
**Fixed modifications:** Carbamidomethyl (C) (apply to specified residues or termini only)
**Variable modifications:**
**N-term :** Dimethyl:2H(6) (Protein N-term)
**Ions Score:** 27 **Expect:** 0.026
**Matches :** 7/27 fragment ions using 12 most intense peaks (help)

| # | b | b⁰ | Seq. | y | y* | y⁰ | # |
|---|---|---|---|---|---|---|---|
| 1 | 122.1083 | 104.0977 | S | | | | 7 |
| 2 | ***251.1509*** | 233.1403 | E | 660.3927 | 643.3661 | 642.3821 | 6 |
| 3 | ***364.2349*** | 346.2244 | I | 531.3501 | 514.3235 | 513.3395 | 5 |
| 4 | 465.2826 | 447.2720 | T | 418.2660 | 401.2395 | 400.2554 | 4 |
| 5 | ***578.3667*** | 560.3561 | L | 317.2183 | 300.1918 | | 3 |
| 6 | ***635.3881*** | 617.3776 | G | 204.1343 | 187.1077 | | 2 |
| 7 | | | K | 147.1128 | 130.0863 | | 1 |

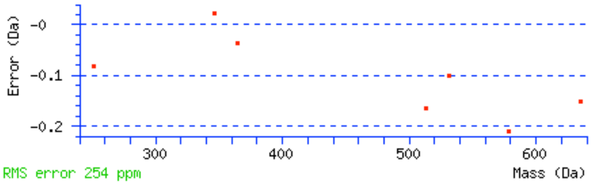


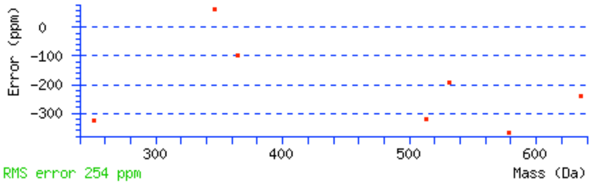


MS/MS Fragmentation of **SEITLGK**
Found in **P16467** in **Uniprot_yeastS288C**, Pyruvate decarboxylase isozyme 2 OS=Saccharomyces cerevisiae (strain ATCC 204508 / S288c) OX=559292 GN=PDC5 PE=1 SV=4

Match to Query 2321: 780.449755 from(781.457031,1+) intensity(12691800.0000) rtinseconds(1052.3514) index(2545)
Title: 20100530_methyl450mM_3.2835.2835.1
Data file D:\Panyue\Proteomic dataset\20100530_methyl450mM_3.RAW

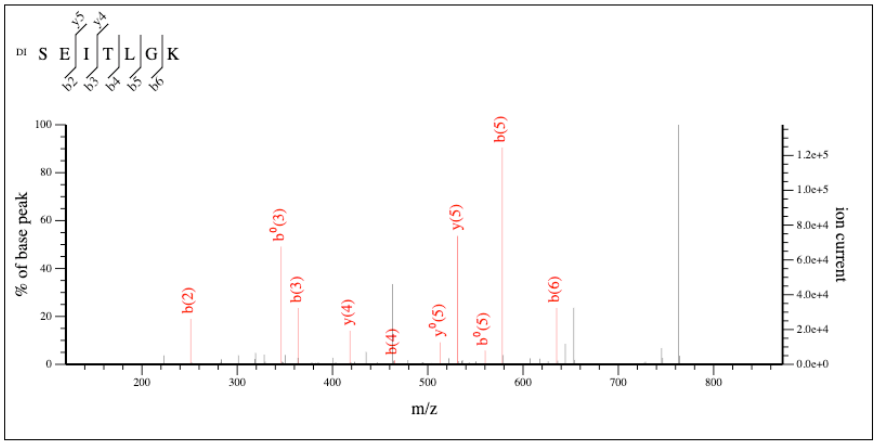

**Monoisotopic mass of neutral peptide Mr(calc):** 780.4864
**Fixed modifications:** Carbamidomethyl (C) (apply to specified residues or termini only)
**Variable modifications:**
**N-term :** Dimethyl:2H(6) (Protein N-term)
**Ions Score:** 31 **Expect:** 0.0094
**Matches :** 10/27 fragment ions using 24 most intense peaks (help)

| # | b | $b^0$ | Seq. | y | y* | $y^0$ | # |
|---|---|---|---|---|---|---|---|
| 1 | 122.1083 | 104.0977 | S | | | | 7 |
| 2 | 251.1509 | 233.1403 | E | 660.3927 | 643.3661 | 642.3821 | 6 |
| 3 | 364.2349 | 346.2244 | I | 531.3501 | 514.3235 | 513.3395 | 5 |
| 4 | 465.2826 | 447.2720 | T | 418.2660 | 401.2395 | 400.2554 | 4 |
| 5 | 578.3667 | 560.3561 | L | 317.2183 | 300.1918 | | 3 |
| 6 | 635.3881 | 617.3776 | G | 204.1343 | 187.1077 | | 2 |
| 7 | | | K | 147.1128 | 130.0863 | | 1 |

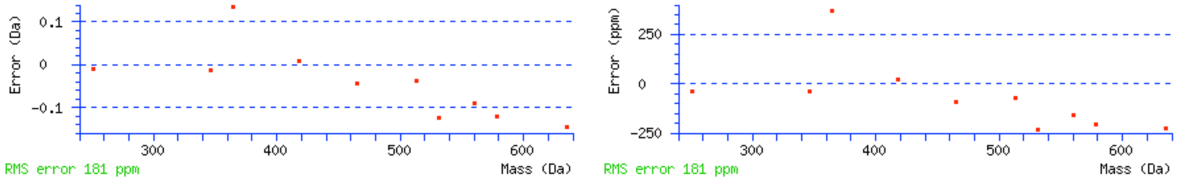


### 95. Pdc 1 (2 peptides in 450mM fraction 3)

MS/MS Fragmentation of SEITLGK
Found in P06169 in Uniprot_yeastS288C, Pyruvate decarboxylase isozyme 1 OS=Saccharomyces cerevisiae (strain ATCC 204508 / S288c) OX=559292 GN=PDC1 PE=1 SV=7

Match to Query 2323: 780.450549 from(781.457825,1+) intensity(12241962.0000) rtinseconds(1055.6165) index(2554)
Title: 20100530_methyl450mM_3.2845.2845.1
Data file D:\Panyue\Proteomic dataset\20100530_methyl450mM_3.RAW

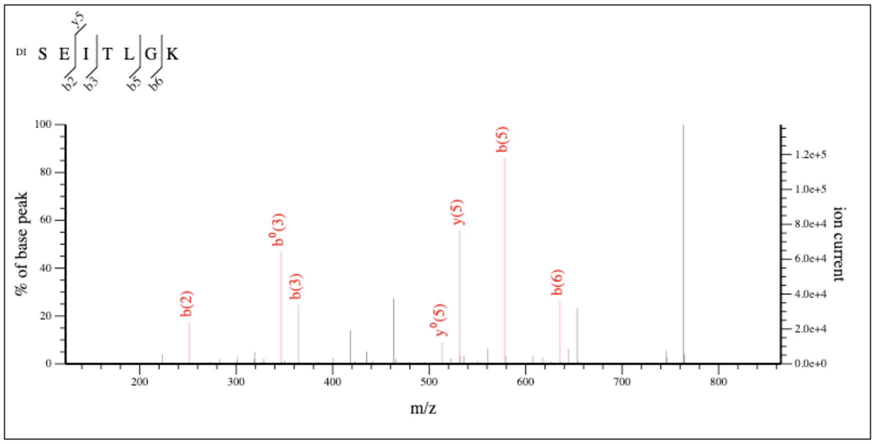

**Monoisotopic mass of neutral peptide Mr(calc):** 780.4864
**Fixed modifications:** Carbamidomethyl (C) (apply to specified residues or termini only)
**Variable modifications:**
**N-term :** Dimethyl:2H(6) (Protein N-term)
**Ions Score:** 27 **Expect:** 0.026
**Matches :** 7/27 fragment ions using 12 most intense peaks (help)

| # | b | b$^0$ | Seq. | y | y* | y$^0$ | # |
|---|---|---|---|---|---|---|---|
| 1 | 122.1083 | 104.0977 | S | | | | 7 |
| 2 | *251.1509* | 233.1403 | E | 660.3927 | 643.3661 | 642.3821 | 6 |
| 3 | *364.2349* | 346.2244 | I | 531.3501 | 514.3235 | 513.3395 | 5 |
| 4 | 465.2826 | 447.2720 | T | 418.2660 | 401.2395 | 400.2554 | 4 |
| 5 | *578.3667* | 560.3561 | L | 317.2183 | 300.1918 | | 3 |
| 6 | *635.3881* | 617.3776 | G | 204.1343 | 187.1077 | | 2 |
| 7 | | | K | 147.1128 | 130.0863 | | 1 |

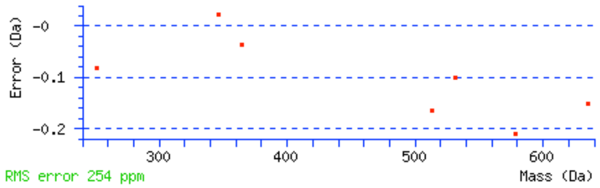


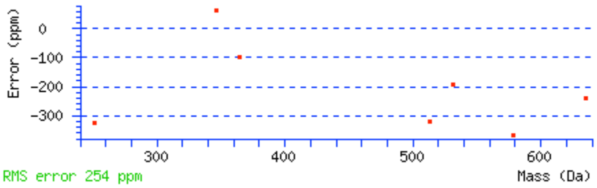


MS/MS Fragmentation of SEITLGK
Found in P06169 in Uniprot_yeastS288C, Pyruvate decarboxylase isozyme 1 OS=Saccharomyces cerevisiae (strain ATCC 204508 / S288c) OX=559292 GN=PDC1 PE=1 SV=7

Match to Query 2321: 780.449755 from(781.457031,1+) intensity(12691800.0000) rtinseconds(1052.3514) index(2545)
Title: 20100530_methyl450mM_3.2835.2835.1
Data file D:\Panyue\Proteomic dataset\20100530_methyl450mM_3.RAW

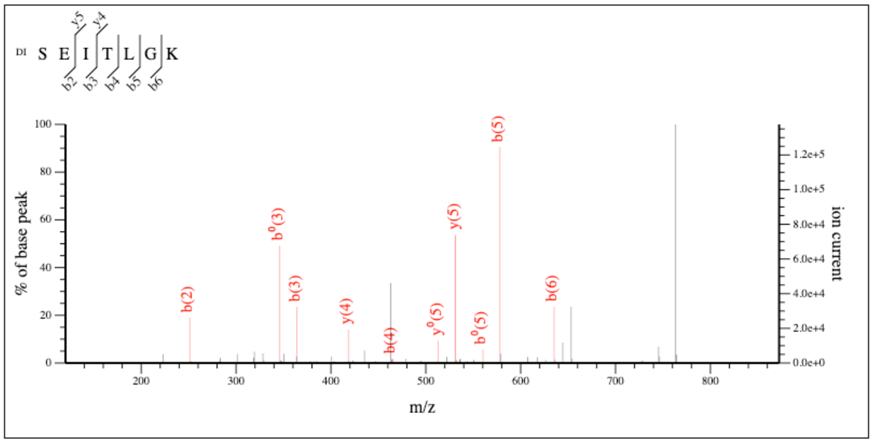

**Monoisotopic mass of neutral peptide Mr(calc):** 780.4864
**Fixed modifications:** Carbamidomethyl (C) (apply to specified residues or termini only)
**Variable modifications:**
**N-term :** Dimethyl:2H(6) (Protein N-term)
**Ions Score:** 31 **Expect:** 0.0094
**Matches :** 10/27 fragment ions using 24 most intense peaks (help)

| # | b | $b^0$ | Seq. | y | y* | $y^0$ | # |
|---|---|---|---|---|---|---|---|
| 1 | 122.1083 | 104.0977 | S | | | | 7 |
| 2 | 251.1509 | 233.1403 | E | 660.3927 | 643.3661 | 642.3821 | 6 |
| 3 | 364.2349 | 346.2244 | I | 531.3501 | 514.3235 | 513.3395 | 5 |
| 4 | 465.2826 | 447.2720 | T | 418.2660 | 401.2395 | 400.2554 | 4 |
| 5 | 578.3667 | 560.3561 | L | 317.2183 | 300.1918 | | 3 |
| 6 | 635.3881 | 617.3776 | G | 204.1343 | 187.1077 | | 2 |
| 7 | | | K | 147.1128 | 130.0863 | | 1 |

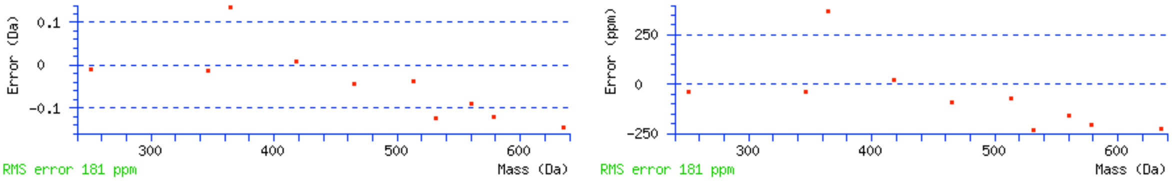


### 96. Psa1 (1 peptide in 450mM fraction 3)

MS/MS Fragmentation of **KGLILVGGYGTR**
Found in **P41940** in **Uniprot_yeastS288C**, Mannose-1-phosphate guanyltransferase OS=Saccharomyces cerevisiae (strain ATCC 204508 / S288c) OX=559292 GN=PSA1 PE=1 SV=2

Match to Query 2929: 1266.758172 from(423.260000,3+) intensity(1157.6622) rtinseconds(140.0563) index(341)
Title: 20100530_methyl450mM_3.381.381.
Data file D:\Panyue\Proteomic dataset\20100530_methyl450mM_3.RAW

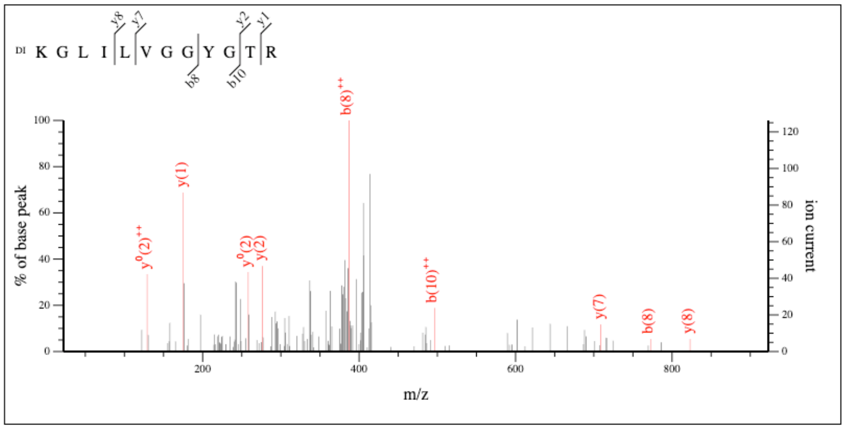

**Monoisotopic mass of neutral peptide Mr(calc):** 1266.7931
**Fixed modifications:** Carbamidomethyl (C) (apply to specified residues or termini only)
**Variable modifications:**
**N-term :** Dimethyl:2H(6) (Protein N-term)
**Ions Score:** 25 **Expect:** 0.088
**Matches :** 10/110 fragment ions using 15 most intense peaks (help)

| # | b | b++ | b* | b*++ | b0 | b0++ | Seq. | y | y++ | y* | y*++ | y0 | y0++ | # |
|---|---|---|---|---|---|---|---|---|---|---|---|---|---|---|
| 1 | 163.1712 | 82.0892 | 146.1447 | 73.5760 | | | K | | | | | | | 12 |
| 2 | 220.1927 | 110.6000 | 203.1661 | 102.0867 | | | G | 1105.6364 | 553.3218 | 1088.6099 | 544.8086 | 1087.6259 | 544.3166 | 11 |
| 3 | 333.2767 | 167.1420 | 316.2502 | 158.6287 | | | L | 1048.6150 | 524.8111 | 1031.5884 | 516.2978 | 1030.6044 | 515.8058 | 10 |
| 4 | 446.3608 | 223.6840 | 429.3342 | 215.1708 | | | I | 935.5309 | 468.2691 | 918.5043 | 459.7558 | 917.5203 | 459.2638 | 9 |
| 5 | 559.4449 | 280.2261 | 542.4183 | 271.7128 | | | L | ***822.4468*** | 411.7271 | 805.4203 | 403.2138 | 804.4363 | 402.7218 | 8 |
| 6 | 658.5133 | 329.7603 | 641.4867 | 321.2470 | | | V | ***709.3628*** | 355.1850 | 692.3362 | 346.6717 | 691.3522 | 346.1797 | 7 |
| 7 | 715.5347 | 358.2710 | 698.5082 | 349.7577 | | | G | 610.2944 | 305.6508 | 593.2678 | 297.1375 | 592.2838 | 296.6455 | 6 |
| 8 | 772.5562 | **386.7817** | 755.5296 | 378.2685 | | | G | 553.2729 | 277.1401 | 536.2463 | 268.6268 | 535.2623 | 268.1348 | 5 |
| 9 | 935.6195 | 468.3134 | 918.5930 | 459.8001 | | | Y | ***496.2514*** | 248.6293 | 479.2249 | 240.1161 | 478.2409 | 239.6241 | 4 |
| 10 | 992.6410 | **496.8241** | 975.6144 | 488.3109 | | | G | 333.1881 | 167.0977 | 316.1615 | 158.5844 | 315.1775 | 158.0924 | 3 |
| 11 | 1093.6887 | 547.3480 | 1076.6621 | 538.8347 | 1075.6781 | 538.3427 | T | ***276.1666*** | 138.5870 | 259.1401 | 130.0737 | 258.1561 | 129.5817 | 2 |
| 12 | | | | | | | R | ***175.1190*** | 88.0631 | 158.0924 | 79.5498 | | | 1 |

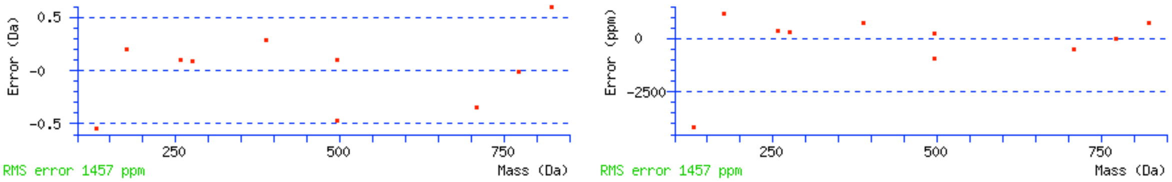


### 97. Fzo1 (1 peptide in 450mM fraction 3)

MS/MS Fragmentation of **SEGKQQFK**
Found in **P38297** in **Uniprot_yeastS288C**, Mitofusin FZO1 OS=Saccharomyces cerevisiae (strain ATCC 204508 / S288c) OX=559292 GN=FZO1 PE=1 SV=1

Match to Query 4281: 984.539586 from(493.277069,2+) intensity(399240.3125) rtinseconds(1653.1527) index(4102)
Title: 20100530_methyl450mM_3.4567.4567.2
Data file D:\Panyue\Proteomic dataset\20100530_methyl450mM_3.RAW

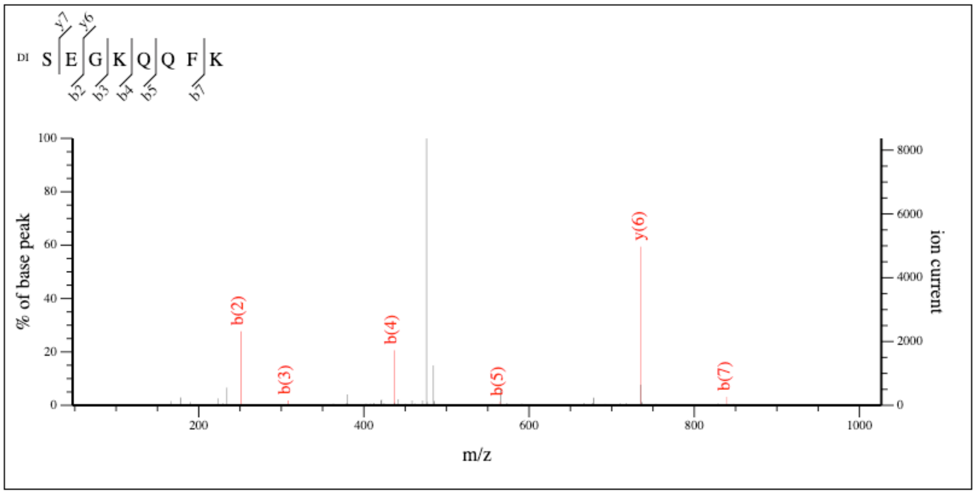

**Monoisotopic mass of neutral peptide Mr(calc):** 984.5511
**Fixed modifications:** Carbamidomethyl (C) (apply to specified residues or termini only)
**Variable modifications:**
**N-term :** Dimethyl:2H(6) (Protein N-term)
**Ions Score:** 27  **Expect:** 0.029
**Matches :** 7/66 fragment ions using 16 most intense peaks  (help)

| # | b | b++ | b* | b*++ | b0 | b0++ | Seq. | y | y++ | y* | y*++ | y0 | y0++ | # |
|---|---|---|---|---|---|---|---|---|---|---|---|---|---|---|
| 1 | 122.1083 | 61.5578 | | | 104.0977 | 52.5525 | S | | | | | | | 8 |
| 2 | 251.1509 | 126.0791 | | | 233.1403 | 117.0738 | E | 864.4574 | 432.7323 | 847.4308 | 424.2191 | 846.4468 | 423.7271 | 7 |
| 3 | 308.1723 | 154.5898 | | | 290.1618 | 145.5845 | G | 735.4148 | 368.2110 | 718.3883 | 359.6978 | | | 6 |
| 4 | 436.2673 | 218.6373 | 419.2407 | 210.1240 | 418.2567 | 209.6320 | K | 678.3933 | 339.7003 | 661.3668 | 331.1870 | | | 5 |
| 5 | 564.3259 | 282.6666 | 547.2993 | 274.1533 | 546.3153 | 273.6613 | Q | 550.2984 | 275.6528 | 533.2718 | 267.1396 | | | 4 |
| 6 | 692.3844 | 346.6959 | 675.3579 | 338.1826 | 674.3739 | 337.6906 | Q | 422.2398 | 211.6235 | 405.2132 | 203.1103 | | | 3 |
| 7 | 839.4529 | 420.2301 | 822.4263 | 411.7168 | 821.4423 | 411.2248 | F | 294.1812 | 147.5942 | 277.1547 | 139.0810 | | | 2 |
| 8 | | | | | | | K | 147.1128 | 74.0600 | 130.0863 | 65.5468 | | | 1 |

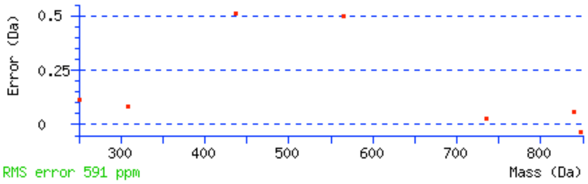


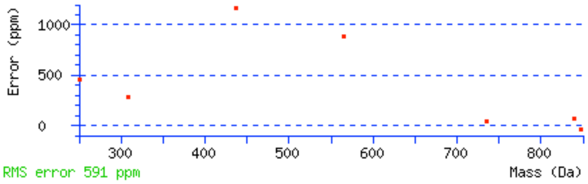


### 98. Hpc2 (1 peptide in 500mM fraction 2)

MS/MS Fragmentation of **DQKAIVLDNSKSGSK**
Found in **Q01448** in **Uniprot_yeastS288C**, Histone promoter control protein 2 OS=Saccharomyces cerevisiae (strain ATCC 204508 / S288c) OX=559292 GN=HPC2 PE=1 SV=2

Match to Query 8657: 1605.809422 from(803.911987,2+) intensity(193736.0156) rtinseconds(3732.6176) index(9410)
Title: 20100530_methyl500mM_2.10472.10472.2
Data file D:\Panyue\Proteomic dataset\20100530_methyl500mM_2.RAW

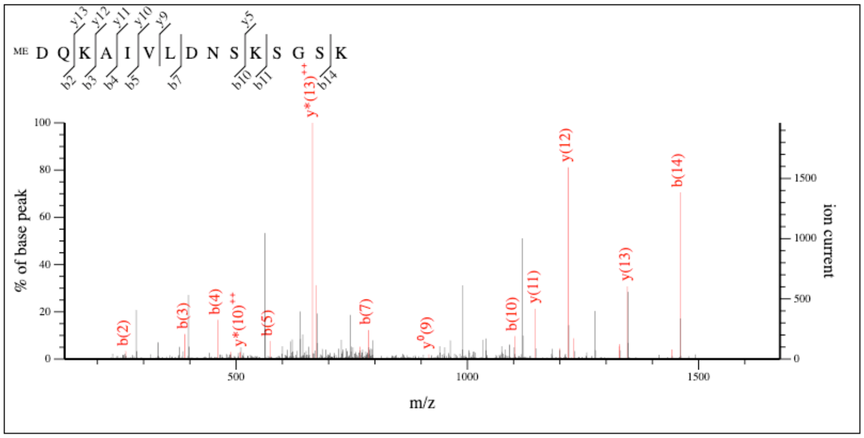

```
Monoisotopic mass of neutral peptide Mr(calc): 1605.8765
Fixed modifications: Carbamidomethyl (C) (apply to specified residues or termini only)
Variable modifications:
N-term : Methyl:2H(3) (Protein N-term)
Ions Score: 31  Expect: 0.012
Matches : 24/164 fragment ions using 55 most intense peaks   (help)
```

| # | b | b++ | b* | b*++ | b0 | b0++ | Seq. | y | y++ | y* | y*++ | y0 | y0++ | # |
|---|---|---|---|---|---|---|---|---|---|---|---|---|---|---|
| 1 | 133.0687 | 67.0380 | | | 115.0581 | 58.0327 | D | | | | | | | 15 |
| 2 | 261.1273 | 131.0673 | 244.1007 | 122.5540 | 243.1167 | 122.0620 | Q | 1474.8224 | 737.9148 | 1457.7958 | 729.4016 | 1456.8118 | 728.9095 | 14 |
| 3 | 389.2222 | 195.1148 | 372.1957 | 186.6015 | 371.2117 | 186.1095 | K | 1346.7638 | 673.8855 | 1329.7373 | 665.3723 | 1328.7532 | 664.8803 | 13 |
| 4 | 460.2594 | 230.6333 | 443.2328 | 222.1200 | 442.2488 | 221.6280 | A | 1218.6688 | 609.8381 | 1201.6423 | 601.3248 | 1200.6583 | 600.8328 | 12 |
| 5 | 573.3434 | 287.1753 | 556.3169 | 278.6621 | 555.3329 | 278.1701 | I | 1147.6317 | 574.3195 | 1130.6052 | 565.8062 | 1129.6212 | 565.3142 | 11 |
| 6 | 672.4118 | 336.7096 | 655.3853 | 328.1963 | 654.4013 | 327.7043 | V | 1034.5477 | 517.7775 | 1017.5211 | 509.2642 | 1016.5371 | 508.7722 | 10 |
| 7 | 785.4959 | 393.2516 | 768.4693 | 384.7383 | 767.4853 | 384.2463 | L | 935.4792 | 468.2433 | 918.4527 | 459.7300 | 917.4687 | 459.2380 | 9 |
| 8 | 900.5228 | 450.7651 | 883.4963 | 442.2518 | 882.5123 | 441.7598 | D | 822.3952 | 411.7012 | 805.3686 | 403.1880 | 804.3846 | 402.6959 | 8 |
| 9 | 1014.5658 | 507.7865 | 997.5392 | 499.2732 | 996.5552 | 498.7812 | N | 707.3682 | 354.1878 | 690.3417 | 345.6745 | 689.3577 | 345.1825 | 7 |
| 10 | 1101.5978 | 551.3025 | 1084.5712 | 542.7893 | 1083.5872 | 542.2973 | S | 593.3253 | 297.1663 | 576.2988 | 288.6530 | 575.3148 | 288.1610 | 6 |
| 11 | 1229.6928 | 615.3500 | 1212.6662 | 606.8367 | 1211.6822 | 606.3447 | K | 506.2933 | 253.6503 | 489.2667 | 245.1370 | 488.2827 | 244.6450 | 5 |
| 12 | 1316.7248 | 658.8660 | 1299.6982 | 650.3528 | 1298.7142 | 649.8607 | S | 378.1983 | 189.6028 | 361.1718 | 181.0895 | 360.1878 | 180.5975 | 4 |
| 13 | 1373.7462 | 687.3768 | 1356.7197 | 678.8635 | 1355.7357 | 678.3715 | G | 291.1663 | 146.0868 | 274.1397 | 137.5735 | 273.1557 | 137.0815 | 3 |
| 14 | 1460.7783 | 730.8928 | 1443.7517 | 722.3795 | 1442.7677 | 721.8875 | S | 234.1448 | 117.5761 | 217.1183 | 109.0628 | 216.1343 | 108.5708 | 2 |
| 15 | | | | | | | K | 147.1128 | 74.0600 | 130.0863 | 65.5468 | | | 1 |

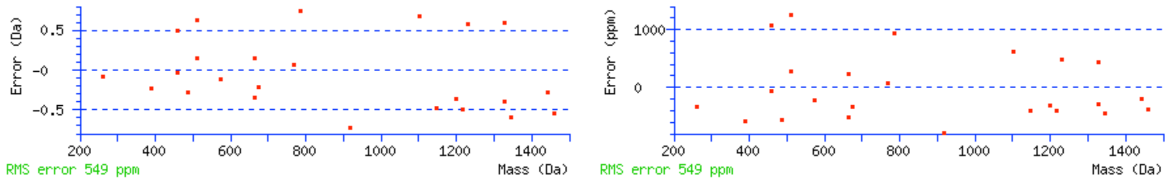


### 99. Eft1 (1 peptide in 500mM fraction 2)

MS/MS Fragmentation of **VAFTVDQMRSLMDK**
Found in **P32324** in **Uniprot_yeastS288C**, Elongation factor 2 OS=Saccharomyces cerevisiae (strain ATCC 204508 / S288c) OX=559292 GN=EFT1 PE=1 SV=1

Match to Query 9168: 1706.871190 from(854.442871,2+) intensity(351496.2500) rtinseconds(2759.1235) index(6914)
Title: 20100530_methyl500mM_2.7696.7696.2
Data file D:\Panyue\Proteomic dataset\20100530_methyl500mM_2.RAW

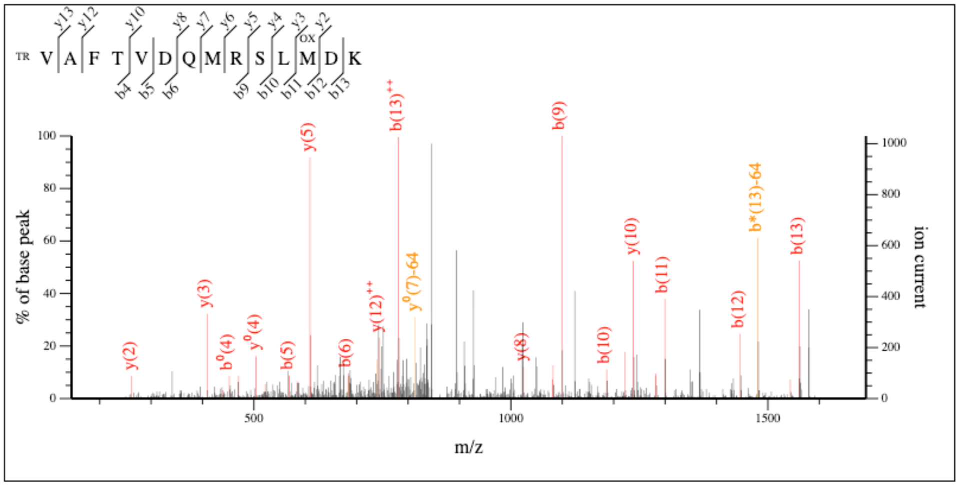

**Monoisotopic mass of neutral peptide Mr(calc):** 1706.9046
**Fixed modifications:** Carbamidomethyl (C) (apply to specified residues or termini only)
**Variable modifications:**
**N-term :** Trimethyl:2H(9) (Protein N-term)
**M12 :** Oxidation (M), with neutral losses 0.0000(shown in table), 63.9983
**Ions Score:** 22 **Expect:** 0.11
**Matches :** 39/214 fragment ions using 98 most intense peaks (help)

| # | b | b++ | b* | b*++ | b0 | b0++ | Seq. | y | y++ | y* | y*++ | y0 | y0++ | # |
|---|---|---|---|---|---|---|---|---|---|---|---|---|---|---|
| 1 | 151.1791 | 76.0932 | | | | | V | | | | | | | 14 |
| 2 | 222.2162 | 111.6118 | | | | | A | 1557.7400 | 779.3736 | 1540.7134 | 770.8604 | 1539.7294 | 770.3683 | 13 |
| 3 | 369.2847 | 185.1460 | | | | | F | 1486.7029 | 743.8551 | 1469.6763 | 735.3418 | 1468.6923 | 734.8498 | 12 |
| 4 | 470.3323 | 235.6698 | | | 452.3218 | 226.6645 | T | 1339.6344 | 670.3209 | 1322.6079 | 661.8076 | 1321.6239 | 661.3156 | 11 |
| 5 | 569.4008 | 285.2040 | | | 551.3902 | 276.1987 | V | 1238.5868 | 619.7970 | 1221.5602 | 611.2837 | 1220.5762 | 610.7917 | 10 |
| 6 | 684.4277 | 342.7175 | | | 666.4171 | 333.7122 | D | 1139.5184 | 570.2628 | 1122.4918 | 561.7495 | 1121.5078 | 561.2575 | 9 |
| 7 | 812.4863 | 406.7468 | 795.4597 | 398.2335 | 794.4757 | 397.7415 | Q | 1024.4914 | 512.7493 | 1007.4649 | 504.2361 | 1006.4808 | 503.7441 | 8 |
| 8 | 943.5268 | 472.2670 | 926.5002 | 463.7537 | 925.5162 | 463.2617 | M | 896.4328 | 448.7201 | 879.4063 | 440.2068 | 878.4223 | 439.7148 | 7 |
| 9 | 1099.6279 | 550.3176 | 1082.6013 | 541.8043 | 1081.6173 | 541.3123 | R | 765.3924 | 383.1998 | 748.3658 | 374.6865 | 747.3818 | 374.1945 | 6 |
| 10 | 1186.6599 | 593.8336 | 1169.6333 | 585.3203 | 1168.6493 | 584.8283 | S | 609.2912 | 305.1493 | 592.2647 | 296.6360 | 591.2807 | 296.1440 | 5 |
| 11 | 1299.7440 | 650.3756 | 1282.7174 | 641.8623 | 1281.7334 | 641.3703 | L | 522.2592 | 261.6332 | 505.2327 | 253.1200 | 504.2486 | 252.6280 | 4 |
| 12 | 1446.7794 | 723.8933 | 1429.7528 | 715.3800 | 1428.7688 | 714.8880 | M | 409.1751 | 205.0912 | 392.1486 | 196.5779 | 391.1646 | 196.0859 | 3 |
| 13 | 1561.8063 | 781.4068 | 1544.7798 | 772.8935 | 1543.7957 | 772.4015 | D | 262.1397 | 131.5735 | 245.1132 | 123.0602 | 244.1292 | 122.5682 | 2 |
| 14 | | | | | | | K | 147.1128 | 74.0600 | 130.0863 | 65.5468 | | | 1 |

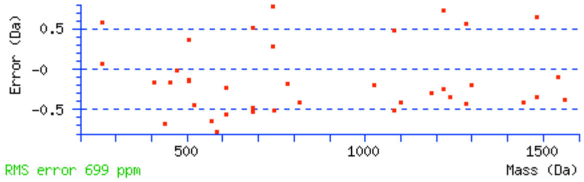


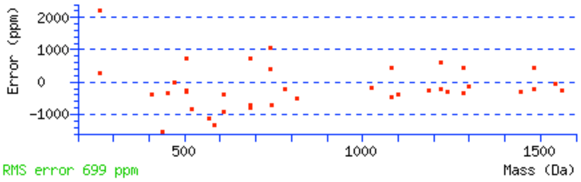


### 100. MRX11 (1 peptide in 500mM fraction 2)

MS/MS Fragmentation of **TVMNLFFRPCQLQMGSGPLELMLKR**
Found in **Q03079** in **Uniprot_yeastS288C**, MIOREX complex component 11 OS=Saccharomyces cerevisiae (strain ATCC 204508 / S288c) OX=559292 GN=MRX11 PE=1 SV=1

Match to Query 16072: 2998.424157 from(1000.481995,3+) intensity(203715.4062) rtinseconds(2851.7978) index(7147)
Title: 20100530_methyl500mM_2.7955.7955.3
Data file D:\Panyue\Proteomic dataset\20100530_methyl500mM_2.RAW

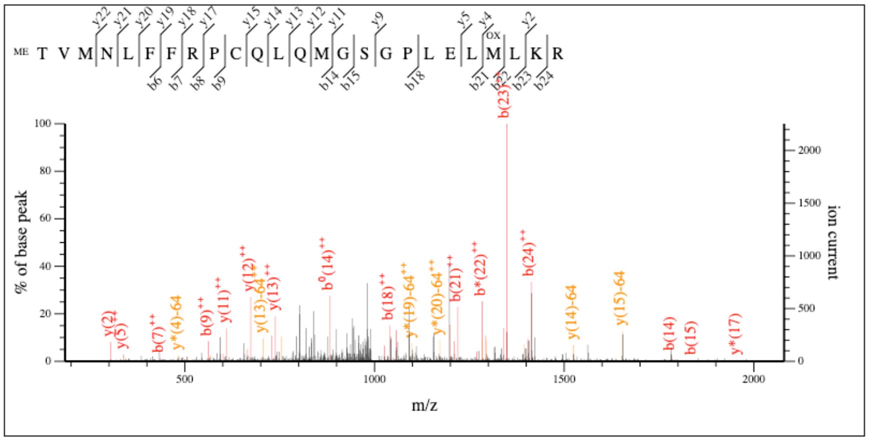


**Monoisotopic mass of neutral peptide Mr(calc):** 2998.5456
**Fixed modifications:** Carbamidomethyl (C) (apply to specified residues or termini only)
**Variable modifications:**
**N-term :** Methyl:2H(3) (Protein N-term)
**M22 :** Oxidation (M), with neutral losses 0.0000(shown in table), 63.9983
**Ions Score:** 20 **Expect:** 0.021
**Matches :** 64/412 fragment ions using 119 most intense peaks (help)

| # | b | b++ | b* | b*++ | b0 | b0++ | Seq. | y | y++ | y* | y*++ | y0 | y0++ | # |
|---|---|---|---|---|---|---|---|---|---|---|---|---|---|---|
| 1 | 119.0894 | 60.0484 | | | 101.0789 | 51.0431 | T | | | | | | | 25 |
| 2 | 218.1578 | 109.5826 | | | 200.1473 | 100.5773 | V | 2881.4708 | 1441.2390 | 2864.4442 | 1432.7257 | 2863.4602 | 1432.2337 | 24 |
| 3 | 349.1983 | 175.1028 | | | 331.1878 | 166.0975 | M | 2782.4024 | 1391.7048 | 2765.3758 | 1383.1915 | 2764.3918 | 1382.6995 | 23 |
| 4 | 463.2413 | 232.1243 | 446.2147 | 223.6110 | 445.2307 | 223.1190 | N | 2651.3619 | 1326.1846 | 2634.3353 | 1317.6713 | 2633.3513 | 1317.1793 | 22 |
| 5 | 576.3253 | 288.6663 | 559.2988 | 280.1530 | 558.3148 | 279.6610 | L | 2537.3189 | **1269.1631** | 2520.2924 | 1260.6498 | 2519.3084 | 1260.1578 | 21 |
| 6 | 723.3937 | 362.2005 | 706.3672 | 353.6872 | 705.3832 | 353.1952 | F | 2424.2349 | 1212.6211 | 2407.2083 | 1204.1078 | 2406.2243 | 1203.6158 | 20 |
| 7 | 870.4622 | ***435.7347*** | 853.4356 | 427.2214 | 852.4516 | 426.7294 | F | 2277.1665 | 1139.0869 | 2260.1399 | 1130.5736 | 2259.1559 | 1130.0816 | 19 |
| 8 | ***1026.5633*** | 513.7853 | 1009.5367 | 505.2720 | 1008.5527 | 504.7800 | R | 2130.0980 | 1065.5527 | 2113.0715 | 1057.0394 | 2112.0875 | 1056.5474 | 18 |
| 9 | 1123.6160 | ***562.3117*** | 1106.5895 | 553.7984 | 1105.6055 | 553.3064 | P | 1973.9969 | 987.5021 | 1956.9704 | 978.9888 | 1955.9864 | 978.4968 | 17 |
| 10 | ***1283.6467*** | 642.3270 | 1266.6201 | 633.8137 | 1265.6361 | 633.3217 | C | 1876.9442 | 938.9757 | 1859.9176 | 930.4625 | 1858.9336 | 929.9704 | 16 |
| 11 | 1411.7053 | ***706.3563*** | 1394.6787 | 697.8430 | 1393.6947 | 697.3510 | Q | 1716.9135 | 858.9604 | 1699.8870 | 850.4471 | 1698.9030 | 849.9551 | 15 |
| 12 | ***1524.7893*** | 762.8983 | 1507.7628 | 754.3850 | 1506.7788 | 753.8930 | L | 1588.8549 | 794.9311 | 1571.8284 | 786.4178 | 1570.8444 | 785.9258 | 14 |
| 13 | ***1652.8479*** | 826.9276 | 1635.8213 | 818.4143 | 1634.8373 | 817.9223 | Q | 1475.7709 | **738.3891** | 1458.7443 | 729.8758 | 1457.7603 | 729.3838 | 13 |
| 14 | ***1783.8884*** | 892.4478 | 1766.8618 | 883.9346 | 1765.8778 | 883.4425 | M | 1347.7123 | **674.3598** | 1330.6858 | 665.8465 | 1329.7017 | 665.3545 | 12 |
| 15 | ***1840.9098*** | 920.9586 | 1823.8833 | 912.4453 | 1822.8993 | 911.9533 | G | 1216.6718 | **608.8395** | 1199.6453 | 600.3263 | 1198.6613 | 599.8343 | 11 |
| 16 | 1927.9419 | 964.4746 | 1910.9153 | 955.9613 | 1909.9313 | 955.4693 | S | 1159.6504 | 580.3288 | 1142.6238 | 571.8155 | 1141.6398 | 571.3235 | 10 |
| 17 | 1984.9633 | 992.9853 | 1967.9368 | 984.4720 | 1966.9528 | 983.9800 | G | 1072.6183 | 536.8128 | 1055.5918 | 528.2995 | 1054.6078 | 527.8075 | 9 |
| 18 | 2082.0161 | ***1041.5117*** | 2064.9896 | 1032.9984 | 2064.0055 | 1032.5064 | P | 1015.5969 | 508.3021 | 998.5703 | 499.7888 | 997.5863 | 499.2968 | 8 |
| 19 | 2195.1002 | ***1098.0537*** | 2178.0736 | 1089.5404 | 2177.0896 | 1089.0484 | L | 918.5441 | 459.7757 | 901.5176 | 451.2624 | 900.5335 | 450.7704 | 7 |
| 20 | 2324.1428 | 1162.5750 | 2307.1162 | 1154.0617 | 2306.1322 | 1153.5697 | E | 805.4600 | 403.2337 | 788.4335 | 394.7204 | 787.4495 | 394.2284 | 6 |
| 21 | 2437.2268 | ***1219.1170*** | 2420.2003 | 1210.6038 | 2419.2163 | 1210.1118 | L | 676.4174 | **338.7124** | 659.3909 | 330.1991 | | | 5 |
| 22 | 2584.2622 | ***1292.6347*** | 2567.2357 | 1284.1215 | 2566.2517 | 1283.6295 | M | 563.3334 | 282.1703 | 546.3068 | 273.6571 | | | 4 |
| 23 | 2697.3463 | ***1349.1768*** | 2680.3197 | 1340.6635 | 2679.3357 | 1340.1715 | L | 416.2980 | 208.6526 | 399.2714 | 200.1394 | | | 3 |
| 24 | 2825.4412 | ***1413.2243*** | 2808.4147 | 1404.7110 | 2807.4307 | 1404.2190 | K | 303.2139 | 152.1106 | 286.1874 | 143.5973 | | | 2 |
| 25 | | | | | | | R | 175.1190 | 88.0631 | 158.0924 | 79.5498 | | | 1 |

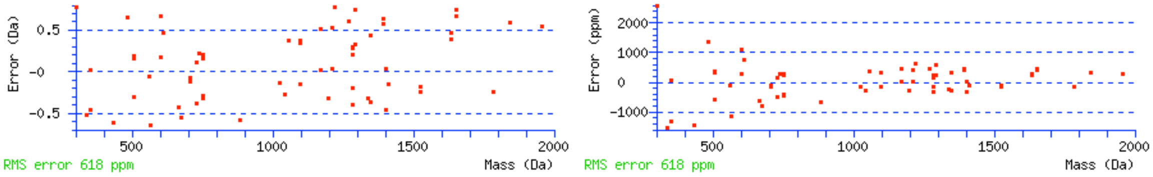


101. NUS1 (1 peptide in 600mM fraction 1)

MS/MS Fragmentation of **PTMIK**
Found in **Q12063** in **Uniprot_yeastS288C**, Dehydrodolichyl diphosphate synthase complex subunit NUS1 OS=Saccharomyces cerevisiae (strain ATCC 204508 / S288c) OX=559292 GN=NUS1 PE=1 SV=1

Match to Query 1806: 638.375720 from(639.382996,1+) intensity(3783.9541) rtinseconds(597.4514) index(1436)
Title: 20100530_methyl600mM_1.1601.1601.1
Data file D:\Panyue\Proteomic dataset\20100530_methyl600mM_1.RAW

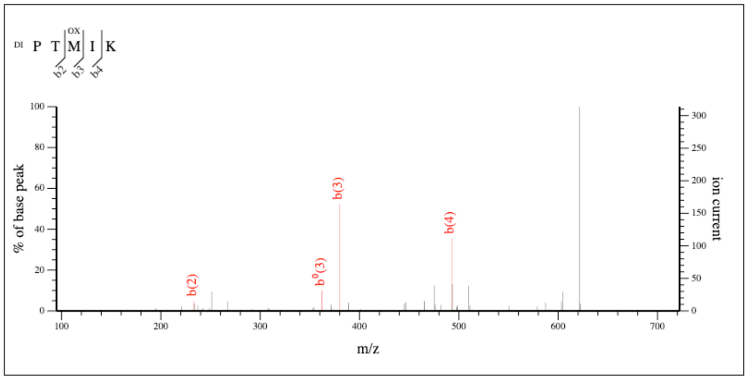

```
Monoisotopic mass of neutral peptide Mr(calc): 638.3944
Fixed modifications: Carbamidomethyl (C) (apply to specified residues or termini only)
Variable modifications:
N-term : Dimethyl:2H(6) (Protein N-term)
M3     : Oxidation (M), with neutral losses 0.0000(shown in table), 63.9983
Ions Score: 22  Expect: 0.039
Matches : 4/25 fragment ions using 10 most intense peaks   (help)
```

| # | b | b⁰ | Seq. | y | y* | y⁰ | # |
|---|---|---|---|---|---|---|---|
| 1 | 132.1290 | | P | | | | 5 |
| 2 | 233.1767 | 215.1661 | T | 508.2799 | 491.2534 | 490.2694 | 4 |
| 3 | 380.2121 | 362.2015 | M | 407.2323 | 390.2057 | | 3 |
| 4 | 493.2961 | 475.2856 | I | 260.1969 | 243.1703 | | 2 |
| 5 | | | K | 147.1128 | 130.0863 | | 1 |

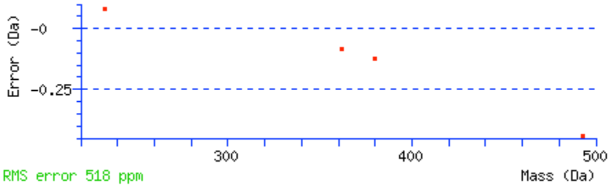


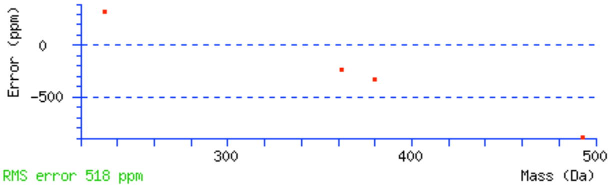


### 102. DED81 (1 peptide in 600mM fraction 1)

MS/MS Fragmentation of SSLYIK
Found in P38707 in Uniprot_yeastS288C, Asparagine--tRNA ligase, cytoplasmic OS=Saccharomyces cerevisiae (strain ATCC 204508 / S288c) OX=559292 GN=DED81 PE=1 SV=1

Match to Query 2178: 743.434619 from(744.441895,1+) intensity(158141.6875) rtinseconds(1292.3655) index(3134)
Title: 20100530_methyl600mM_1.3492.3492.1
Data file D:\Panyue\Proteomic dataset\20100530_methyl600mM_1.RAW

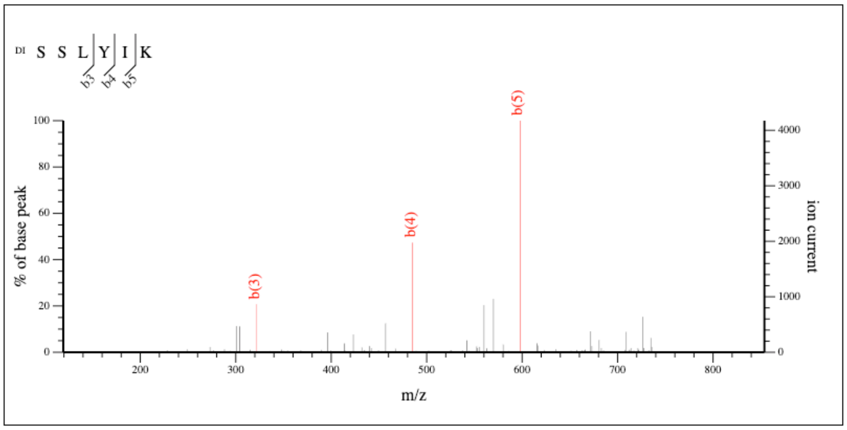

**Monoisotopic mass of neutral peptide Mr(calc):** 743.4700
**Fixed modifications:** Carbamidomethyl (C) (apply to specified residues or termini only)
**Variable modifications:**
**N-term :** Dimethyl:2H(6) (Protein N-term)
**Ions Score:** 21 **Expect:** 0.094
**Matches :** 3/21 fragment ions using 6 most intense peaks (help)

| # | b | $b^0$ | Seq. | y | y* | $y^0$ | # |
|---|---|---|---|---|---|---|---|
| 1 | 122.1083 | 104.0977 | S | | | | 6 |
| 2 | 209.1403 | 191.1297 | S | 623.3763 | 606.3497 | 605.3657 | 5 |
| 3 | *322.2244* | 304.2138 | L | 536.3443 | 519.3177 | | 4 |
| 4 | *485.2877* | 467.2771 | Y | 423.2602 | 406.2336 | | 3 |
| 5 | *598.3718* | 580.3612 | I | 260.1969 | 243.1703 | | 2 |
| 6 | | | K | 147.1128 | 130.0863 | | 1 |

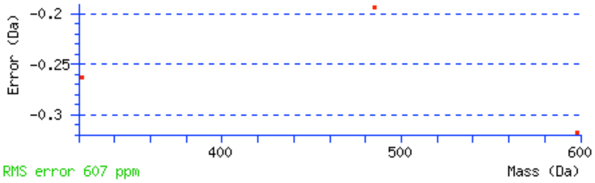


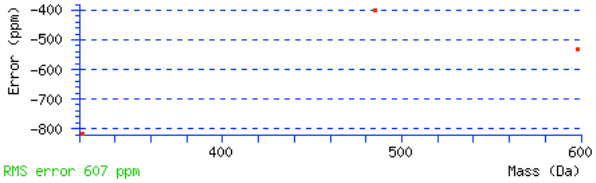


### 103. Apl4 (2 peptides in 600mM fraction 1)

MS/MS Fragmentation of **GSSLRSFIK**
Found in **Q12028** in **Uniprot_yeastS288C**, AP-1 complex subunit gamma-1 OS=Saccharomyces cerevisiae (strain ATCC 204508 / S288c) OX=559292 GN=APL4 PE=1 SV=1

Match to Query 4469: 1010.554247 from(1011.561523,1+) intensity(213313.2500) rtinseconds(2584.2685) index(6422)
Title: 20100530_methyl600mM_1.7150.7150.1
Data file D:\Panyue\Proteomic dataset\20100530_methyl600mM_1.RAW

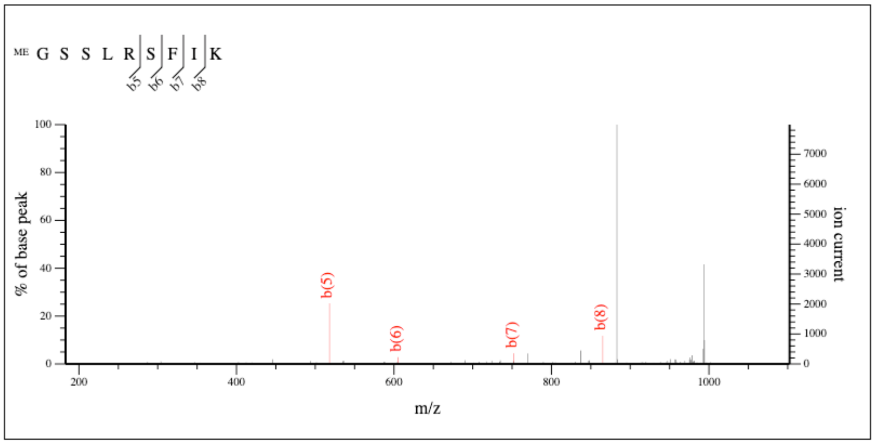

**Monoisotopic mass of neutral peptide Mr(calc):** 1010.5952
**Fixed modifications:** Carbamidomethyl (C) (apply to specified residues or termini only)
**Variable modifications:**
**N-term :** Methyl:2H(3) (Protein N-term)
**Ions Score:** 24 **Expect:** 0.043
**Matches :** 4/40 fragment ions using 8 most intense peaks (help)

| # | b | b* | $b^0$ | Seq. | y | y* | $y^0$ | # |
|---|---|---|---|---|---|---|---|---|
| 1 | 75.0632 | | | G | | | | 9 |
| 2 | 162.0952 | | 144.0847 | S | 937.5465 | 920.5200 | 919.5360 | 8 |
| 3 | 249.1273 | | 231.1167 | S | 850.5145 | 833.4880 | 832.5039 | 7 |
| 4 | 362.2113 | | 344.2008 | L | 763.4825 | 746.4559 | 745.4719 | 6 |
| 5 | 518.3125 | 501.2859 | 500.3019 | R | 650.3984 | 633.3719 | 632.3879 | 5 |
| 6 | 605.3445 | 588.3179 | 587.3339 | S | 494.2973 | 477.2708 | 476.2867 | 4 |
| 7 | 752.4129 | 735.3863 | 734.4023 | F | 407.2653 | 390.2387 | | 3 |
| 8 | 865.4970 | 848.4704 | 847.4864 | I | 260.1969 | 243.1703 | | 2 |
| 9 | | | | K | 147.1128 | 130.0863 | | 1 |

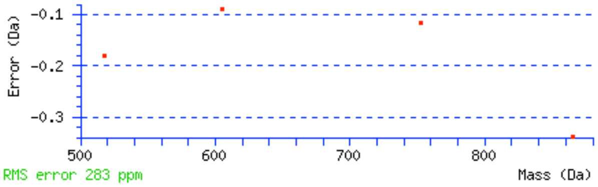


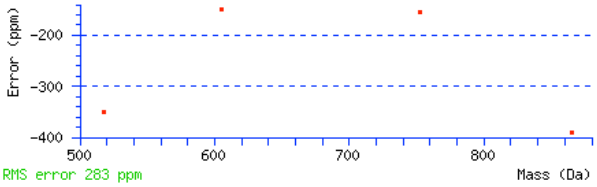


MS/MS Fragmentation of GSSLRSFIK
Found in Q12028 in Uniprot_yeastS288C, AP-1 complex subunit gamma-1 OS=Saccharomyces cerevisiae (strain ATCC 204508 / S288c) OX=559292 GN=APL4 PE=1 SV=1

Match to Query 4468: 1010.553868 from(506.284210,2+) intensity(504566.5312) rtinseconds(2569.6078) index(6384)
Title: 20100530_methyl600mM_1.7108.7108.2
Data file D:\Panyue\Proteomic dataset\20100530_methyl600mM_1.RAW

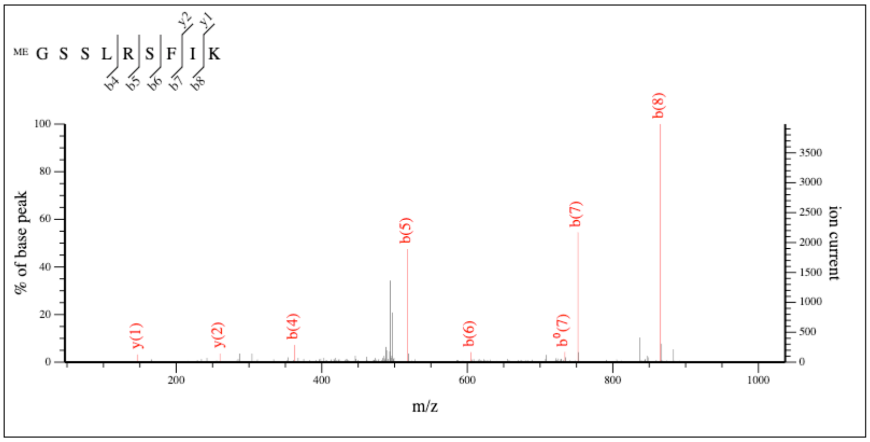

**Monoisotopic mass of neutral peptide Mr(calc):** 1010.5952
**Fixed modifications:** Carbamidomethyl (C) (apply to specified residues or termini only)
**Variable modifications:**
**N-term :** Methyl:2H(3) (Protein N-term)
**Ions Score:** 50 **Expect:** 0.00011
**Matches :** 9/80 fragment ions using 8 most intense peaks (help)

| # | b | $b^{++}$ | b* | $b^{*++}$ | $b^0$ | $b^{0++}$ | Seq. | y | $y^{++}$ | y* | $y^{*++}$ | $y^0$ | $y^{0++}$ | # |
|---|---|---|---|---|---|---|---|---|---|---|---|---|---|---|
| 1 | 75.0632 | 38.0352 | | | | | G | | | | | | | 9 |
| 2 | 162.0952 | 81.5513 | | | 144.0847 | 72.5460 | S | 937.5465 | 469.2769 | 920.5200 | 460.7636 | 919.5360 | 460.2716 | 8 |
| 3 | 249.1273 | 125.0673 | | | 231.1167 | 116.0620 | S | 850.5145 | 425.7609 | 833.4880 | 417.2476 | 832.5039 | 416.7556 | 7 |
| 4 | ***362.2113*** | 181.6093 | | | 344.2008 | 172.6040 | L | 763.4825 | 382.2449 | 746.4559 | 373.7316 | 745.4719 | 373.2396 | 6 |
| 5 | ***518.3125*** | 259.6599 | 501.2859 | 251.1466 | 500.3019 | 250.6546 | R | 650.3984 | 325.7028 | 633.3719 | 317.1896 | 632.3879 | 316.6976 | 5 |
| 6 | ***605.3445*** | 303.1759 | 588.3179 | 294.6626 | 587.3339 | 294.1706 | S | 494.2973 | 247.6523 | 477.2708 | 239.1390 | 476.2867 | 238.6470 | 4 |
| 7 | ***752.4129*** | 376.7101 | 735.3863 | 368.1968 | 734.4023 | 367.7048 | F | 407.2653 | 204.1363 | 390.2387 | 195.6230 | | | 3 |
| 8 | ***865.4970*** | 433.2521 | 848.4704 | 424.7388 | 847.4864 | 424.2468 | I | **260.1969** | 130.6021 | 243.1703 | 122.0888 | | | 2 |
| 9 | | | | | | | K | **147.1128** | 74.0600 | 130.0863 | 65.5468 | | | 1 |

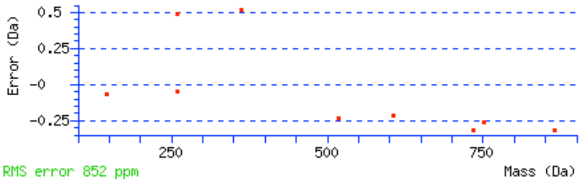


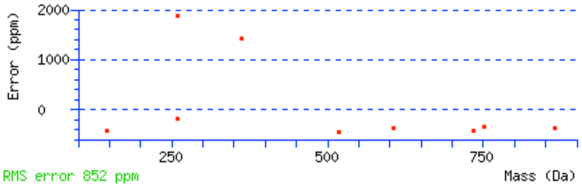


### 104. Seg2 (1 peptide in 600mM fraction 1)

MS/MS Fragmentation of **IQESSPDALAAAAAIGNALSYNGR**
Found in **P34250** in **Uniprot_yeastS288C**, Eisosome protein SEG2 OS=Saccharomyces cerevisiae (strain ATCC 204508 / S288c) OX=559292 GN=SEG2 PE=1 SV=1

Match to Query 13399: 2410.268172 from(804.430000,3+) intensity(287449.1562) rtinseconds(3261.5166) index(8149)
Title: 20100530_methyl600mM_1.9072.9072.3
Data file D:\Panyue\Proteomic dataset\20100530_methyl600mM_1.RAW

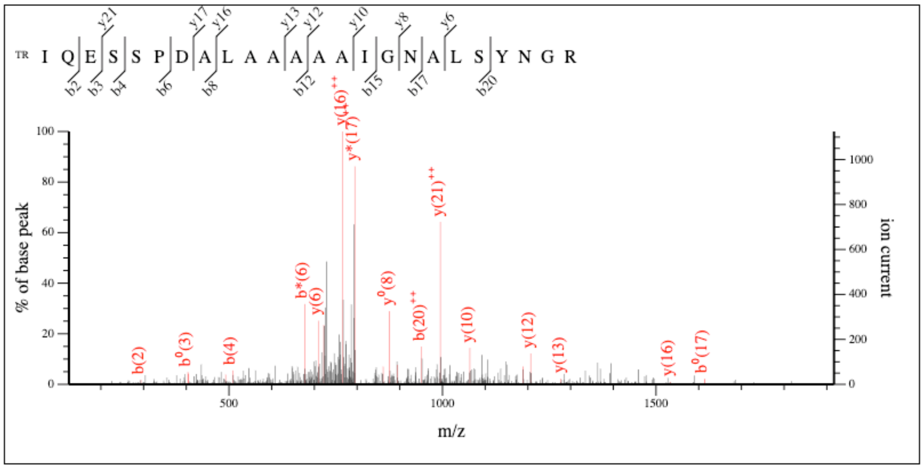


**Monoisotopic mass of neutral peptide Mr(calc):** 2410.2802
**Fixed modifications:** Carbamidomethyl (C) (apply to specified residues or termini only)
**Variable modifications:**
**N-term :** Trimethyl:2H(9) (Protein N-term)
**Ions Score:** 21 **Expect:** 0.01
**Matches :** 30/262 fragment ions using 60 most intense peaks (help)

| # | b | b++ | b* | b*++ | b0 | b0++ | Seq. | y | y++ | y* | y*++ | y0 | y0++ | # |
|---|---|---|---|---|---|---|---|---|---|---|---|---|---|---|
| 1 | 165.1948 | 83.1010 | | | | | I | | | | | | | 24 |
| 2 | **293.2534** | 147.1303 | 276.2268 | 138.6170 | | | Q | 2247.1000 | 1124.0536 | 2230.0735 | 1115.5404 | 2229.0894 | 1115.0484 | 23 |
| 3 | 422.2960 | 211.6516 | 405.2694 | 203.1383 | 404.2854 | 202.6463 | E | 2119.0414 | 1060.0244 | 2102.0149 | 1051.5111 | 2101.0309 | 1051.0191 | 22 |
| 4 | **509.3280** | 255.1676 | 492.3014 | 246.6544 | 491.3174 | 246.1623 | S | 1989.9988 | **995.5031** | 1972.9723 | 986.9898 | 1971.9883 | 986.4978 | 21 |
| 5 | 596.3600 | 298.6836 | 579.3335 | 290.1704 | 578.3494 | 289.6784 | S | 1902.9668 | **951.9870** | 1885.9403 | 943.4738 | 1884.9563 | 942.9818 | 20 |
| 6 | 693.4128 | 347.2100 | 676.3862 | 338.6967 | 675.4022 | 338.2047 | P | 1815.9348 | 908.4710 | 1798.9082 | 899.9578 | 1797.9242 | 899.4657 | 19 |
| 7 | 808.4397 | 404.7235 | 791.4132 | 396.2102 | 790.4291 | 395.7182 | D | 1718.8820 | 859.9446 | 1701.8555 | 851.4314 | 1700.8715 | 850.9394 | 18 |
| 8 | 879.4768 | 440.2421 | 862.4503 | 431.7288 | 861.4663 | 431.2368 | A | 1603.8551 | 802.4312 | 1586.8285 | 793.9179 | 1585.8445 | 793.4259 | 17 |
| 9 | 992.5609 | 496.7841 | 975.5343 | 488.2708 | 974.5503 | 487.7788 | L | ***1532.8180*** | **766.9126** | 1515.7914 | 758.3993 | 1514.8074 | 757.9073 | 16 |
| 10 | **1063.5980** | 532.3026 | 1046.5715 | 523.7894 | 1045.5874 | 523.2974 | A | 1419.7339 | 710.3706 | 1402.7074 | 701.8573 | 1401.7233 | 701.3653 | 15 |
| 11 | 1134.6351 | 567.8212 | 1117.6086 | 559.3079 | 1116.6246 | 558.8159 | A | 1348.6968 | 674.8520 | 1331.6702 | 666.3388 | 1330.6862 | 665.8467 | 14 |
| 12 | 1205.6722 | 603.3398 | 1188.6457 | 594.8265 | 1187.6617 | 594.3345 | A | ***1277.6597*** | 639.3335 | 1260.6331 | 630.8202 | 1259.6491 | 630.3282 | 13 |
| 13 | 1276.7093 | 638.8583 | 1259.6828 | 630.3450 | 1258.6988 | 629.8530 | A | ***1206.6226*** | 603.8149 | 1189.5960 | 595.3016 | 1188.6120 | 594.8096 | 12 |
| 14 | 1347.7465 | 674.3769 | 1330.7199 | 665.8636 | 1329.7359 | 665.3716 | A | 1135.5854 | 568.2964 | 1118.5589 | 559.7831 | 1117.5749 | 559.2911 | 11 |
| 15 | 1460.8305 | 730.9189 | 1443.8040 | 722.4056 | 1442.8200 | 721.9136 | I | ***1064.5483*** | 532.7778 | 1047.5218 | 524.2645 | 1046.5378 | 523.7725 | 10 |
| 16 | 1517.8520 | 759.4296 | 1500.8254 | 750.9164 | 1499.8414 | 750.4244 | G | ***951.4643*** | 476.2358 | 934.4377 | 467.7225 | 933.4537 | 467.2305 | 9 |
| 17 | 1631.8949 | 816.4511 | 1614.8684 | 807.9378 | 1613.8844 | 807.4458 | N | ***894.4428*** | 447.7250 | 877.4163 | 439.2118 | 876.4322 | 438.7198 | 8 |
| 18 | 1702.9320 | 851.9697 | 1685.9055 | 843.4564 | 1684.9215 | 842.9644 | A | 780.3999 | 390.7036 | 763.3733 | 382.1903 | 762.3893 | 381.6983 | 7 |
| 19 | 1816.0161 | 908.5117 | 1798.9895 | 899.9984 | 1798.0055 | 899.5064 | L | ***709.3628*** | 355.1850 | 692.3362 | 346.6717 | 691.3522 | 346.1797 | 6 |
| 20 | 1903.0481 | 952.0277 | 1886.0216 | 943.5144 | 1885.0376 | 943.0224 | S | 596.2787 | 298.6430 | 579.2522 | 290.1297 | 578.2681 | 289.6377 | 5 |
| 21 | 2066.1115 | 1033.5594 | 2049.0849 | 1025.0461 | 2048.1009 | 1024.5541 | Y | ***509.2467*** | 255.1270 | 492.2201 | 246.6137 | | | 4 |
| 22 | 2180.1544 | 1090.5808 | 2163.1278 | 1082.0676 | 2162.1438 | 1081.5755 | N | 346.1833 | 173.5953 | 329.1568 | 165.0820 | | | 3 |
| 23 | 2237.1758 | 1119.0916 | 2220.1493 | 1110.5783 | 2219.1653 | 1110.0863 | G | 232.1404 | 116.5738 | 215.1139 | 108.0606 | | | 2 |
| 24 | | | | | | | R | 175.1190 | 88.0631 | 158.0924 | 79.5498 | | | 1 |

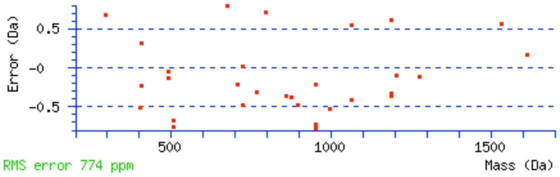


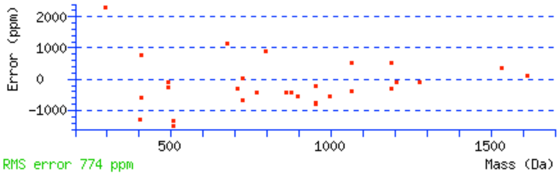


### 105. Ded81 (2 peptides in 600mM fraction 2)

MS/MS Fragmentation of **SSLYIK**
Found in **P38707** in **Uniprot_yeastS288C**, Asparagine--tRNA ligase, cytoplasmic OS=Saccharomyces cerevisiae (strain ATCC 204508 / S288c) OX=559292 GN=DED81 PE=1 SV=1

Match to Query 2205: 743.433337 from(744.440613,1+) intensity(166316.6250) rtinseconds(1282.3702) index(3118)
Title: 20100530_methyl600mM_2.3473.3473.1
Data file D:\Panyue\Proteomic dataset\20100530_methyl600mM_2.RAW

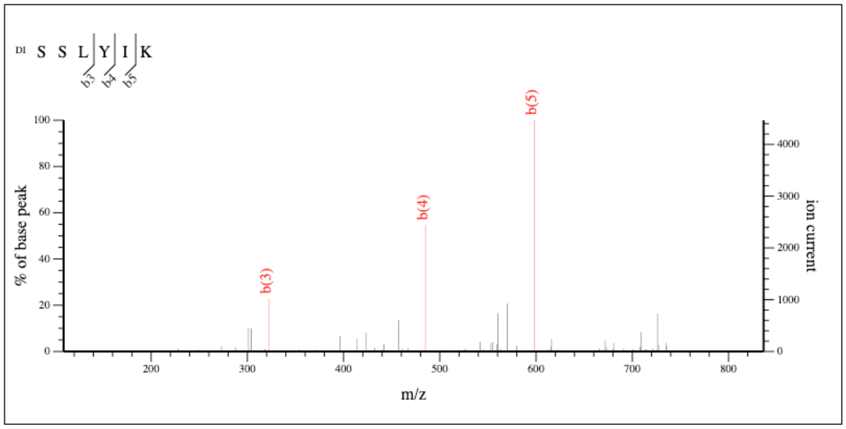

**Monoisotopic mass of neutral peptide Mr(calc):** 743.4700
**Fixed modifications:** Carbamidomethyl (C) (apply to specified residues or termini only)
**Variable modifications:**
**N-term :** Dimethyl:2H(6) (Protein N-term)
**Ions Score:** 21 **Expect:** 0.096
**Matches :** 3/21 fragment ions using 6 most intense peaks (help)

| # | b | $b^0$ | Seq. | y | y* | $y^0$ | # |
|---|---|---|---|---|---|---|---|
| 1 | 122.1083 | 104.0977 | S | | | | 6 |
| 2 | 209.1403 | 191.1297 | S | 623.3763 | 606.3497 | 605.3657 | 5 |
| 3 | 322.2244 | 304.2138 | L | 536.3443 | 519.3177 | | 4 |
| 4 | 485.2877 | 467.2771 | Y | 423.2602 | 406.2336 | | 3 |
| 5 | 598.3718 | 580.3612 | I | 260.1969 | 243.1703 | | 2 |
| 6 | | | K | 147.1128 | 130.0863 | | 1 |

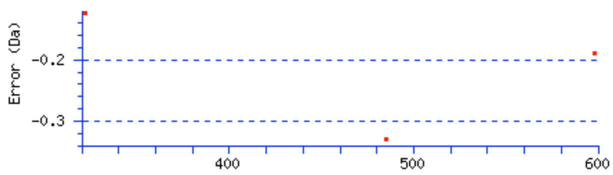

RMS error 486 ppm

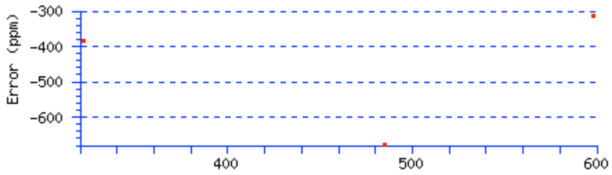

RMS error 486 ppm

MS/MS Fragmentation of **SSLYIK**
Found in **P38707** in **Uniprot_yeastS288C**, Asparagine--tRNA ligase, cytoplasmic OS=Saccharomyces cerevisiae (strain ATCC 204508 / S288c) OX=559292 GN=DED81 PE=1 SV=1

Match to Query 2204: 743.433154 from(744.440430,1+) intensity(211844.5156) rtinseconds(1284.1834) index(3122)
Title: 20100530_methyl600mM_2.3478.3478.1
Data file D:\Panyue\Proteomic dataset\20100530_methyl600mM_2.RAW

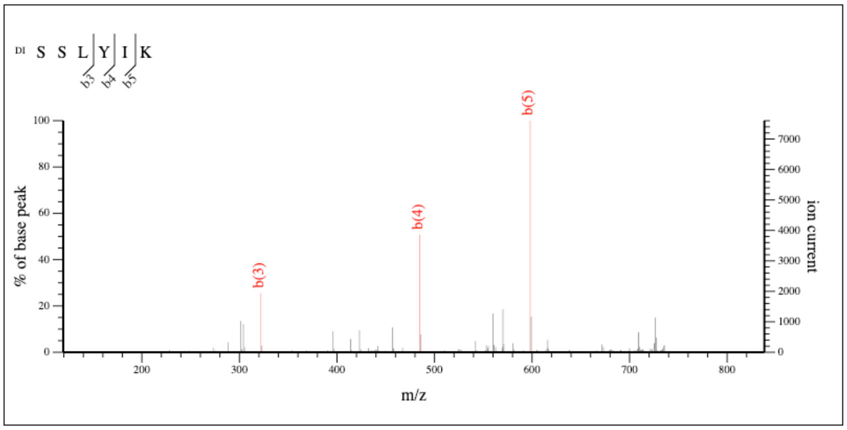

**Monoisotopic mass of neutral peptide Mr(calc):** 743.4700
**Fixed modifications:** Carbamidomethyl (C) (apply to specified residues or termini only)
**Variable modifications:**
**N-term :** Dimethyl:2H(6) (Protein N-term)
**Ions Score:** 24 **Expect:** 0.065
**Matches :** 3/21 fragment ions using 6 most intense peaks (help)

| # | b | $b^0$ | Seq. | y | y* | $y^0$ | # |
|---|---|---|---|---|---|---|---|
| 1 | 122.1083 | 104.0977 | S | | | | 6 |
| 2 | 209.1403 | 191.1297 | S | 623.3763 | 606.3497 | 605.3657 | 5 |
| 3 | *322.2244* | 304.2138 | L | 536.3443 | 519.3177 | | 4 |
| 4 | *485.2877* | 467.2771 | Y | 423.2602 | 406.2336 | | 3 |
| 5 | *598.3718* | 580.3612 | I | 260.1969 | 243.1703 | | 2 |
| 6 | | | K | 147.1128 | 130.0863 | | 1 |

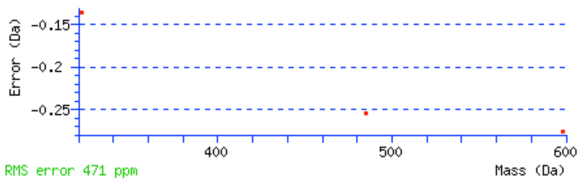


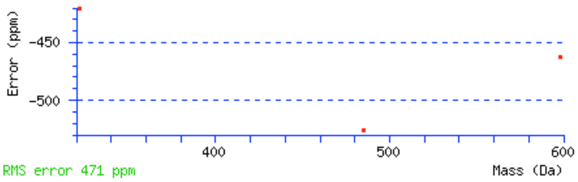


### 106. VMA1 (1 peptide in 600mM fraction 2)

MS/MS Fragmentation of **AGAIENAR**
Found in **P17255** in **Uniprot_yeastS288C**, V-type proton ATPase catalytic subunit A OS=Saccharomyces cerevisiae (strain ATCC 204508 / S288c) OX=559292 GN=VMA1 PE=1 SV=3

Match to Query 2671: 817.415440 from(409.714996,2+) intensity(147836.2031) rtinseconds(2065.3296) index(5117)
Title: 20100530_methyl600mM_2.5697.5697.2
Data file D:\Panyue\Proteomic dataset\20100530_methyl600mM_2.RAW

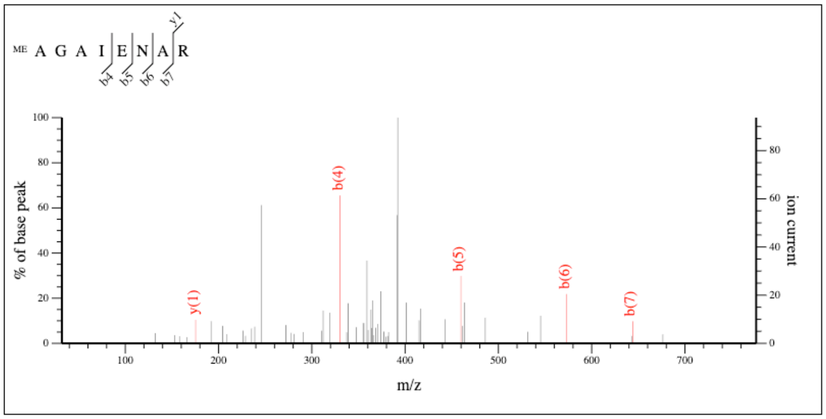

**Monoisotopic mass of neutral peptide Mr(calc):** 817.4485
**Fixed modifications:** Carbamidomethyl (C) (apply to specified residues or termini only)
**Variable modifications:**
**N-term :** Methyl:2H(3) (Protein N-term)
**Ions Score:** 33 **Expect:** 0.01
**Matches :** 5/60 fragment ions using 6 most intense peaks (help)

| # | b | b++ | b* | b*++ | b0 | b0++ | Seq. | y | y++ | y* | y*++ | y0 | y0++ | # |
|---|---|---|---|---|---|---|---|---|---|---|---|---|---|---|
| 1 | 89.0789 | 45.0431 | | | | | A | | | | | | | 8 |
| 2 | 146.1003 | 73.5538 | | | | | G | 730.3842 | 365.6958 | 713.3577 | 357.1825 | 712.3737 | 356.6905 | 7 |
| 3 | 217.1374 | 109.0724 | | | | | A | 673.3628 | 337.1850 | 656.3362 | 328.6717 | 655.3522 | 328.1797 | 6 |
| 4 | ***330.2215*** | 165.6144 | | | | | I | 602.3257 | 301.6665 | 585.2991 | 293.1532 | 584.3151 | 292.6612 | 5 |
| 5 | ***459.2641*** | 230.1357 | | | 441.2535 | 221.1304 | E | 489.2416 | 245.1244 | 472.2150 | 236.6112 | 471.2310 | 236.1191 | 4 |
| 6 | ***573.3070*** | 287.1572 | 556.2805 | 278.6439 | 555.2965 | 278.1519 | N | 360.1990 | 180.6031 | 343.1724 | 172.0899 | | | 3 |
| 7 | ***644.3441*** | 322.6757 | 627.3176 | 314.1624 | 626.3336 | 313.6704 | A | 246.1561 | 123.5817 | 229.1295 | 115.0684 | | | 2 |
| 8 | | | | | | | R | 175.1190 | 88.0631 | 158.0924 | 79.5498 | | | 1 |

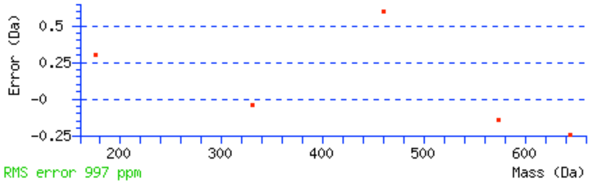


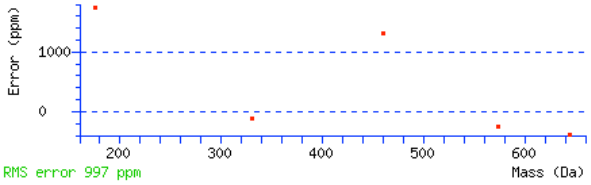


### 107. Seg2 (1 peptide in 600mM fraction 2)

MS/MS Fragmentation of **IQESSPDALAAAAAIGNALSYNGR**
Found in **P34250** in **Uniprot_yeastS288C**, Eisosome protein SEG2 OS=Saccharomyces cerevisiae (strain ATCC 204508 / S288c) OX=559292 GN=SEG2 PE=1 SV=1

Match to Query 13371: 2410.268172 from(804.430000,3+) intensity(167101.1250) rtinseconds(3267.4064) index(8180)
Title: 20100530_methyl600mM_2.9104.9104.3
Data file D:\Panyue\Proteomic dataset\20100530_methyl600mM_2.RAW

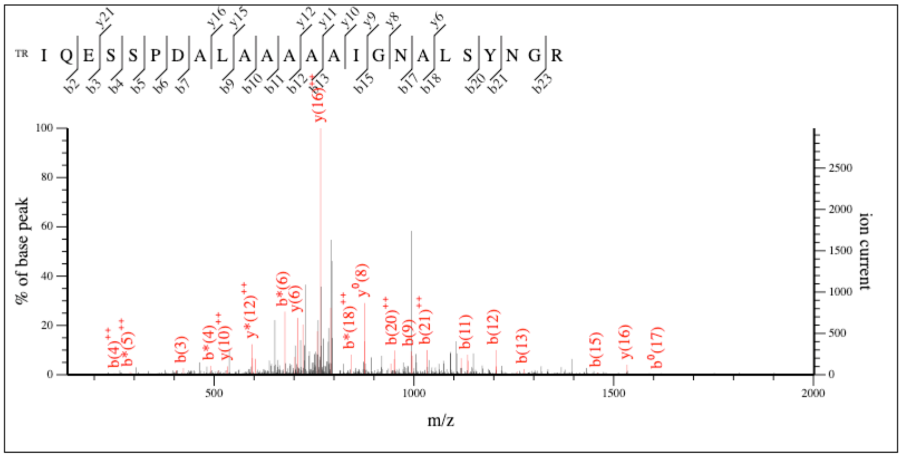


**Monoisotopic mass of neutral peptide Mr(calc):** 2410.2802
**Fixed modifications:** Carbamidomethyl (C) (apply to specified residues or termini only)
**Variable modifications:**
**N-term :** Trimethyl:2H(9) (Protein N-term)
**Ions Score:** 22 **Expect:** 0.024
**Matches :** 55/262 fragment ions using 131 most intense peaks (help)

| # | b | $b^{++}$ | $b^{*}$ | $b^{*++}$ | $b^{0}$ | $b^{0++}$ | Seq. | y | $y^{++}$ | $y^{*}$ | $y^{*++}$ | $y^{0}$ | $y^{0++}$ | # |
|---|---|---|---|---|---|---|---|---|---|---|---|---|---|---|
| 1 | 165.1948 | 83.1010 | | | | | I | | | | | | | 24 |
| 2 | ***293.2534*** | 147.1303 | 276.2268 | 138.6170 | | | Q | 2247.1000 | 1124.0536 | 2230.0735 | 1115.5404 | 2229.0894 | 1115.0484 | 23 |
| 3 | ***422.2960*** | 211.6516 | 405.2694 | 203.1383 | 404.2854 | 202.6463 | E | 2119.0414 | 1060.0244 | 2102.0149 | 1051.5111 | 2101.0309 | 1051.0191 | 22 |
| 4 | ***509.3280*** | ***255.1676*** | 492.3014 | 246.6544 | 491.3174 | 246.1623 | S | 1989.9988 | ***995.5031*** | 1972.9723 | 986.9898 | 1971.9883 | 986.4978 | 21 |
| 5 | ***596.3600*** | 298.6836 | 579.3335 | 290.1704 | 578.3494 | 289.6784 | S | 1902.9668 | ***951.9870*** | 1885.9403 | 943.4738 | 1884.9563 | 942.9818 | 20 |
| 6 | 693.4128 | 347.2100 | 676.3862 | 338.6967 | 675.4022 | 338.2047 | P | 1815.9348 | 908.4710 | 1798.9082 | 899.9578 | 1797.9242 | 899.4657 | 19 |
| 7 | 808.4397 | ***404.7235*** | 791.4132 | 396.2102 | 790.4291 | 395.7182 | D | 1718.8820 | 859.9446 | 1701.8555 | 851.4314 | 1700.8715 | 850.9394 | 18 |
| 8 | 879.4768 | 440.2421 | 862.4503 | 431.7288 | 861.4663 | 431.2368 | A | 1603.8551 | 802.4312 | 1586.8285 | 793.9179 | 1585.8445 | 793.4259 | 17 |
| 9 | ***992.5609*** | 496.7841 | 975.5343 | 488.2708 | 974.5503 | 487.7788 | L | ***1532.8180*** | ***766.9126*** | 1515.7914 | 758.3993 | 1514.8074 | 757.9073 | 16 |
| 10 | 1063.5980 | ***532.3026*** | 1046.5715 | 523.7894 | 1045.5874 | 523.2974 | A | 1419.7339 | 710.3706 | 1402.7074 | 701.8573 | 1401.7233 | 701.3653 | 15 |
| 11 | ***1134.6351*** | 567.8212 | 1117.6086 | 559.3079 | 1116.6246 | 558.8159 | A | 1348.6968 | 674.8520 | 1331.6702 | 666.3388 | 1330.6862 | 665.8467 | 14 |
| 12 | ***1205.6722*** | ***603.3398*** | 1188.6457 | 594.8265 | 1187.6617 | 594.3345 | A | 1277.6597 | 639.3335 | 1260.6331 | 630.8202 | 1259.6491 | 630.3282 | 13 |
| 13 | ***1276.7093*** | 638.8583 | 1259.6828 | 630.3450 | 1258.6988 | 629.8530 | A | ***1206.6226*** | ***603.8149*** | 1189.5960 | 595.3016 | 1188.6120 | 594.8096 | 12 |
| 14 | 1347.7465 | 674.3769 | 1330.7199 | 665.8636 | 1329.7359 | 665.3716 | A | ***1135.5854*** | 568.2964 | 1118.5589 | 559.7831 | 1117.5749 | 559.2911 | 11 |
| 15 | ***1460.8305*** | 730.9189 | 1443.8040 | 722.4056 | 1442.8200 | 721.9136 | I | 1064.5483 | ***532.7778*** | 1047.5218 | 524.2645 | 1046.5378 | 523.7725 | 10 |
| 16 | 1517.8520 | 759.4296 | 1500.8254 | 750.9164 | 1499.8414 | 750.4244 | G | ***951.4643*** | 476.2358 | 934.4377 | 467.7225 | 933.4537 | 467.2305 | 9 |
| 17 | 1631.8949 | 816.4511 | 1614.8684 | 807.9378 | 1613.8844 | 807.4458 | N | 894.4428 | 447.7250 | 877.4163 | 439.2118 | 876.4322 | 438.7198 | 8 |
| 18 | 1702.9320 | 851.9697 | 1685.9055 | 843.4564 | 1684.9215 | 842.9644 | A | 780.3999 | 390.7036 | 763.3733 | 382.1903 | 762.3893 | 381.6983 | 7 |
| 19 | 1816.0161 | 908.5117 | 1798.9895 | 899.9984 | 1798.0055 | 899.5064 | L | ***709.3628*** | 355.1850 | 692.3362 | 346.6717 | 691.3522 | 346.1797 | 6 |
| 20 | 1903.0481 | ***952.0277*** | 1886.0216 | 943.5144 | 1885.0376 | 943.0224 | S | ***596.2787*** | 298.6430 | 579.2522 | 290.1297 | 578.2681 | 289.6377 | 5 |
| 21 | 2066.1115 | ***1033.5594*** | 2049.0849 | 1025.0461 | 2048.1009 | 1024.5541 | Y | 509.2467 | ***255.1270*** | 492.2201 | 246.6137 | | | 4 |
| 22 | 2180.1544 | 1090.5808 | 2163.1278 | 1082.0676 | 2162.1438 | 1081.5755 | N | 346.1833 | 173.5953 | 329.1568 | 165.0820 | | | 3 |
| 23 | 2237.1758 | ***1119.0916*** | 2220.1493 | 1110.5783 | 2219.1653 | 1110.0863 | G | 232.1404 | 116.5738 | 215.1139 | 108.0606 | | | 2 |
| 24 | | | | | | | R | 175.1190 | 88.0631 | 158.0924 | 79.5498 | | | 1 |

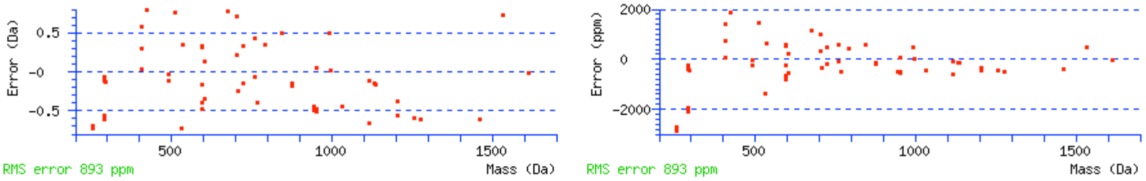


### 108. Hnm1 (1 peptide in 600mM fraction 3)

MS/MS Fragmentation of **SIRNDNASGGYMQPDQSSNASMHK**
Found in **P19807** in **Uniprot_yeastS288C**, Choline transport protein OS=Saccharomyces cerevisiae (strain ATCC 204508 / S288c) OX=559292 GN=HNM1 PE=1 SV=1

Match to Query 13535: 2660.231260 from(533.053528,5+) intensity(40910.3203) rtinseconds(1978.6729) index(4834)
Title: 20100530_methyl600mM_3.5394.5394.5
Data file D:\Panyue\Proteomic dataset\20100530_methyl600mM_3.RAW

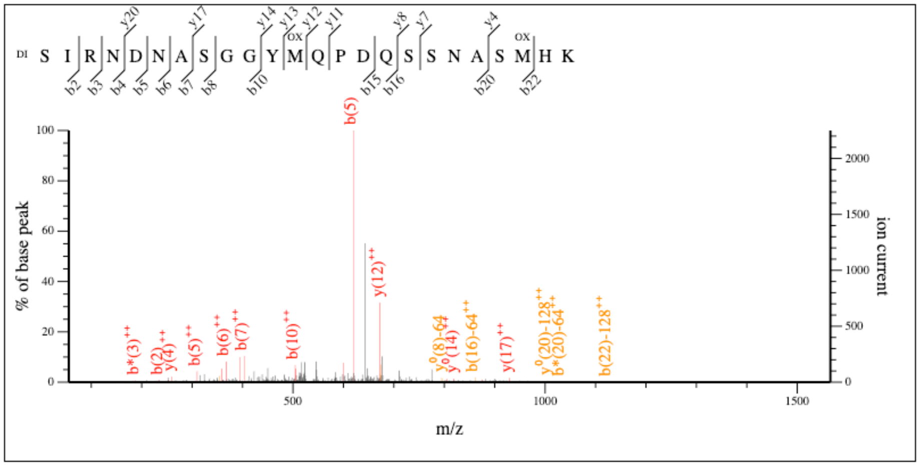

**Monoisotopic mass of neutral peptide Mr(calc):** 2660.1826
**Fixed modifications:** Carbamidomethyl (C) (apply to specified residues or termini only)
**Variable modifications:**
**N-term** : Dimethyl:2H(6) (Protein N-term)
**M12** : Oxidation (M), with neutral losses 63.9983(shown in table), 0.0000
**M22** : Oxidation (M), with neutral losses 63.9983(shown in table), 0.0000
**Ions Score:** 23 **Expect:** 0.026
**Matches :** 36/462 fragment ions using 56 most intense peaks (help)

| # | b | b++ | b* | b*++ | b0 | b0++ | Seq. | y | y++ | y* | y*++ | y0 | y0++ | # |
|---|---|---|---|---|---|---|---|---|---|---|---|---|---|---|
| 1 | 122.1083 | 61.5578 | | | 104.0977 | 52.5525 | S | | | | | | | 24 |
| 2 | 235.1923 | 118.0998 | | | 217.1818 | 109.0945 | I | 2412.0923 | 1206.5498 | 2395.0658 | 1198.0365 | 2394.0817 | 1197.5445 | 23 |
| 3 | 391.2934 | 196.1504 | 374.2669 | 187.6371 | 373.2829 | 187.1451 | R | 2299.0082 | 1150.0078 | 2281.9817 | 1141.4945 | 2280.9977 | 1141.0025 | 22 |
| 4 | 505.3364 | 253.1718 | 488.3098 | 244.6585 | 487.3258 | 244.1665 | N | 2142.9071 | 1071.9572 | 2125.8806 | 1063.4439 | 2124.8966 | 1062.9519 | 21 |
| 5 | 620.3633 | 310.6853 | 603.3368 | 302.1720 | 602.3527 | 301.6800 | D | 2028.8642 | 1014.9357 | 2011.8377 | 1006.4225 | 2010.8536 | 1005.9305 | 20 |
| 6 | 734.4062 | 367.7068 | 717.3797 | 359.1935 | 716.3957 | 358.7015 | N | 1913.8373 | 957.4223 | 1896.8107 | 948.9090 | 1895.8267 | 948.4170 | 19 |
| 7 | 805.4434 | 403.2253 | 788.4168 | 394.7120 | 787.4328 | 394.2200 | A | 1799.7943 | 900.4008 | 1782.7678 | 891.8875 | 1781.7838 | 891.3955 | 18 |
| 8 | 892.4754 | 446.7413 | 875.4488 | 438.2281 | 874.4648 | 437.7360 | S | 1728.7572 | 864.8822 | 1711.7307 | 856.3690 | 1710.7467 | 855.8770 | 17 |
| 9 | 949.4968 | 475.2521 | 932.4703 | 466.7388 | 931.4863 | 466.2468 | G | 1641.7252 | 821.3662 | 1624.6986 | 812.8530 | 1623.7146 | 812.3610 | 16 |
| 10 | 1006.5183 | 503.7628 | 989.4918 | 495.2495 | 988.5077 | 494.7575 | G | 1584.7037 | 792.8555 | 1567.6772 | 784.3422 | 1566.6932 | 783.8502 | 15 |
| 11 | 1169.5816 | 585.2945 | 1152.5551 | 576.7812 | 1151.5711 | 576.2892 | Y | 1527.6823 | 764.3448 | 1510.6557 | 755.8315 | 1509.6717 | 755.3395 | 14 |
| 12 | 1252.6188 | 626.8130 | 1235.5922 | 618.2997 | 1234.6082 | 617.8077 | M | 1364.6189 | 682.8131 | 1347.5924 | 674.2998 | 1346.6084 | 673.8078 | 13 |
| 13 | 1380.6773 | 690.8423 | 1363.6508 | 682.3290 | 1362.6668 | 681.8370 | Q | 1281.5818 | 641.2945 | 1264.5553 | 632.7813 | 1263.5713 | 632.2893 | 12 |
| 14 | 1477.7301 | 739.3687 | 1460.7035 | 730.8554 | 1459.7195 | 730.3634 | P | 1153.5232 | 577.2653 | 1136.4967 | 568.7520 | 1135.5127 | 568.2600 | 11 |
| 15 | 1592.7570 | 796.8822 | 1575.7305 | 788.3689 | 1574.7465 | 787.8769 | D | 1056.4705 | 528.7389 | 1039.4439 | 520.2256 | 1038.4599 | 519.7336 | 10 |
| 16 | 1720.8156 | 860.9114 | 1703.7891 | 852.3982 | 1702.8050 | 851.9062 | Q | 941.4435 | 471.2254 | 924.4170 | 462.7121 | 923.4330 | 462.2201 | 9 |
| 17 | 1807.8476 | 904.4275 | 1790.8211 | 895.9142 | 1789.8371 | 895.4222 | S | 813.3850 | 407.1961 | 796.3584 | 398.6828 | 795.3744 | 398.1908 | 8 |
| 18 | 1894.8797 | 947.9435 | 1877.8531 | 939.4302 | 1876.8691 | 938.9382 | S | 726.3529 | 363.6801 | 709.3264 | 355.1668 | 708.3424 | 354.6748 | 7 |
| 19 | 2008.9226 | 1004.9649 | 1991.8960 | 996.4517 | 1990.9120 | 995.9597 | N | 639.3209 | 320.1641 | 622.2944 | 311.6508 | 621.3103 | 311.1588 | 6 |
| 20 | 2079.9597 | 1040.4835 | 2062.9332 | 1031.9702 | 2061.9491 | 1031.4782 | A | 525.2780 | 263.1426 | 508.2514 | 254.6294 | 507.2674 | 254.1373 | 5 |
| 21 | 2166.9917 | 1083.9995 | 2149.9652 | 1075.4862 | 2148.9812 | 1074.9942 | S | 454.2409 | 227.6241 | 437.2143 | 219.1108 | 436.2303 | 218.6188 | 4 |
| 22 | 2250.0289 | 1125.5181 | 2233.0023 | 1117.0048 | 2232.0183 | 1116.5128 | M | 367.2088 | 184.1081 | 350.1823 | 175.5948 | | | 3 |
| 23 | 2387.0878 | 1194.0475 | 2370.0612 | 1185.5342 | 2369.0772 | 1185.0422 | H | 284.1717 | 142.5895 | 267.1452 | 134.0762 | | | 2 |
| 24 | | | | | | | K | 147.1128 | 74.0600 | 130.0863 | 65.5468 | | | 1 |

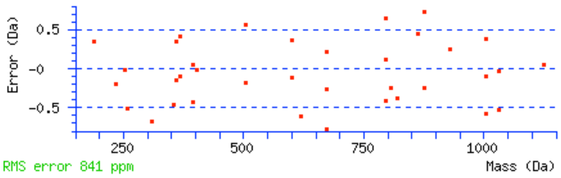


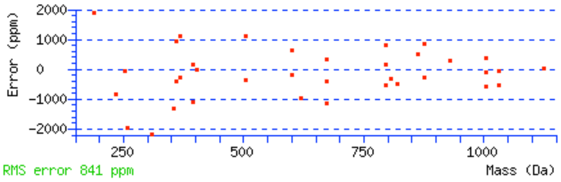


109. Eft1 (1 peptide in 600mM fraction 3)

MS/MS Fragmentation of **VAFTVDQMRSLMDK**
Found in **P32324** in **Uniprot_yeastS288C**, Elongation factor 2 OS=Saccharomyces cerevisiae (strain ATCC 204508 / S288c) OX=559292 GN=EFT1 PE=1 SV=1

Match to Query 7421: 1706.869238 from(854.441895,2+) intensity(41782.4805) rtinseconds(2518.6311) index(6151)
Title: 20100530_methyl600mM_3.6860.6860.2
Data file D:\Panyue\Proteomic dataset\20100530_methyl600mM_3.RAW

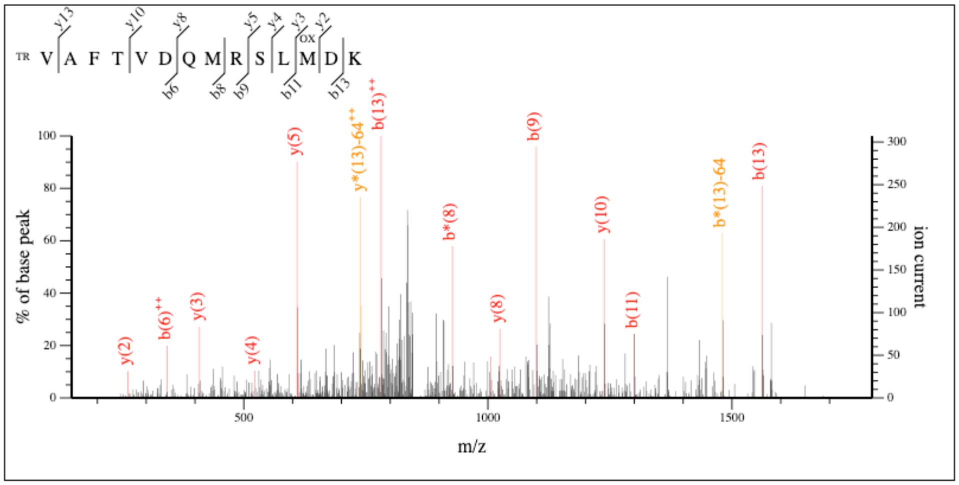


**Monoisotopic mass of neutral peptide Mr(calc):** 1706.9046
**Fixed modifications:** Carbamidomethyl (C) (apply to specified residues or termini only)
**Variable modifications:**
**N-term :** Trimethyl:2H(9) (Protein N-term)
**M12** : Oxidation (M), with neutral losses 0.0000(shown in table), 63.9983
**Ions Score:** 20 **Expect:** 0.14
**Matches :** 21/214 fragment ions using 30 most intense peaks (help)

| # | b | b++ | b* | b*++ | b0 | b0++ | Seq. | y | y++ | y* | y*++ | y0 | y0++ | # |
|---|---|---|---|---|---|---|---|---|---|---|---|---|---|---|
| 1 | 151.1791 | 76.0932 | | | | | V | | | | | | | 14 |
| 2 | 222.2162 | 111.6118 | | | | | A | 1557.7400 | 779.3736 | 1540.7134 | 770.8604 | 1539.7294 | 770.3683 | 13 |
| 3 | 369.2847 | 185.1460 | | | | | F | 1486.7029 | 743.8551 | 1469.6763 | 735.3418 | 1468.6923 | 734.8498 | 12 |
| 4 | 470.3323 | 235.6698 | | | 452.3218 | 226.6645 | T | 1339.6344 | 670.3209 | 1322.6079 | 661.8076 | 1321.6239 | 661.3156 | 11 |
| 5 | 569.4008 | 285.2040 | | | 551.3902 | 276.1987 | V | ***1238.5868*** | 619.7970 | 1221.5602 | 611.2837 | 1220.5762 | 610.7917 | 10 |
| 6 | 684.4277 | 342.7175 | | | 666.4171 | 333.7122 | D | 1139.5184 | 570.2628 | 1122.4918 | 561.7495 | 1121.5078 | 561.2575 | 9 |
| 7 | 812.4863 | 406.7468 | 795.4597 | 398.2335 | 794.4757 | 397.7415 | Q | ***1024.4914*** | 512.7493 | 1007.4649 | 504.2361 | 1006.4808 | 503.7441 | 8 |
| 8 | 943.5268 | 472.2670 | 926.5002 | 463.7537 | 925.5162 | 463.2617 | M | 896.4328 | 448.7201 | 879.4063 | 440.2068 | 878.4223 | 439.7148 | 7 |
| 9 | **1099.6279** | 550.3176 | 1082.6013 | 541.8043 | 1081.6173 | 541.3123 | R | 765.3924 | 383.1998 | 748.3658 | 374.6865 | 747.3818 | 374.1945 | 6 |
| 10 | 1186.6599 | 593.8336 | 1169.6333 | 585.3203 | 1168.6493 | 584.8283 | S | ***609.2912*** | 305.1493 | 592.2647 | 296.6360 | 591.2807 | 296.1440 | 5 |
| 11 | **1299.7440** | 650.3756 | 1282.7174 | 641.8623 | 1281.7334 | 641.3703 | L | ***522.2592*** | 261.6332 | 505.2327 | 253.1200 | 504.2486 | 252.6280 | 4 |
| 12 | 1446.7794 | 723.8933 | 1429.7528 | 715.3800 | 1428.7688 | 714.8880 | M | ***409.1751*** | 205.0912 | 392.1486 | 196.5779 | 391.1646 | 196.0859 | 3 |
| 13 | **1561.8063** | 781.4068 | 1544.7798 | 772.8935 | 1543.7957 | 772.4015 | D | ***262.1397*** | 131.5735 | 245.1132 | 123.0602 | 244.1292 | 122.5682 | 2 |
| 14 | | | | | | | K | 147.1128 | 74.0600 | 130.0863 | 65.5468 | | | 1 |

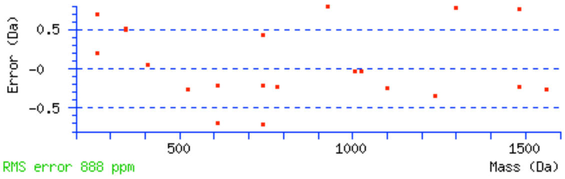


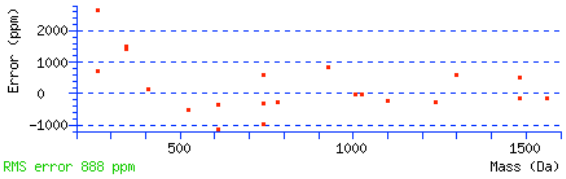


110. Ach1 (1 peptide in 1000mM fraction 1)

MS/MS Fragmentation of **TISNLLK**
Found in **P32316** in **Uniprot_yeastS288C**, Acetyl-CoA hydrolase OS=Saccharomyces cerevisiae (strain ATCC 204508 / S288c) OX=559292 GN=ACH1 PE=1 SV=2

Match to Query 2862: 821.549426 from(822.556702,1+) intensity(117999.4688) rtinseconds(1220.7012) index(2985)
Title: 20100530_methyl1000mM_1.3326.3326.1
Data file D:\Panyue\Proteomic dataset\20100530_methyl1000mM_1.RAW

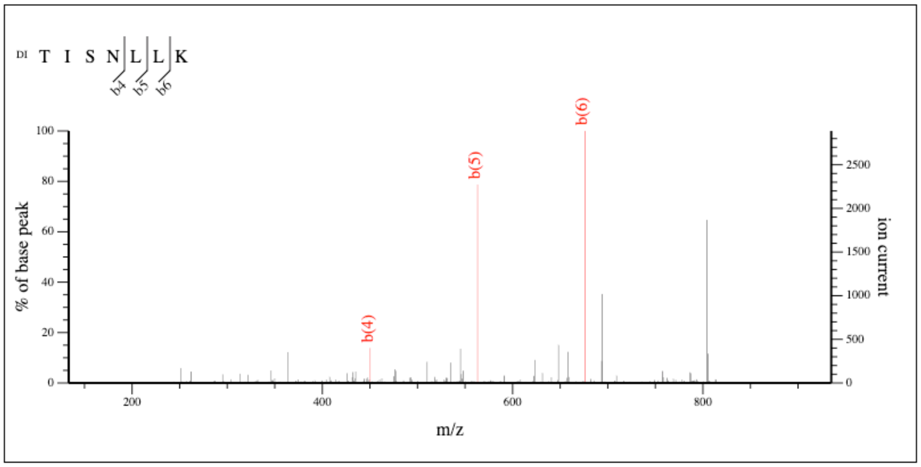


**Monoisotopic mass of neutral peptide Mr(calc):** 821.5493
**Fixed modifications:** Carbamidomethyl (C) (apply to specified residues or termini only)
**Variable modifications:**
**N-term :** Dimethyl:2H(6) (Protein N-term)
**Ions Score:** 20 **Expect:** 0.017
**Matches :** 3/29 fragment ions using 6 most intense peaks (help)

| # | b | b* | b⁰ | Seq. | y | y* | y⁰ | # |
|---|---|---|---|---|---|---|---|---|
| 1 | 136.1239 | | 118.1134 | T | | | | 7 |
| 2 | 249.2080 | | 231.1974 | I | 687.4400 | 670.4134 | 669.4294 | 6 |
| 3 | 336.2400 | | 318.2294 | S | 574.3559 | 557.3293 | 556.3453 | 5 |
| 4 | ***450.2829*** | 433.2564 | 432.2724 | N | 487.3239 | 470.2973 | | 4 |
| 5 | ***563.3670*** | 546.3404 | 545.3564 | L | 373.2809 | 356.2544 | | 3 |
| 6 | ***676.4511*** | 659.4245 | 658.4405 | L | 260.1969 | 243.1703 | | 2 |
| 7 | | | | K | 147.1128 | 130.0863 | | 1 |

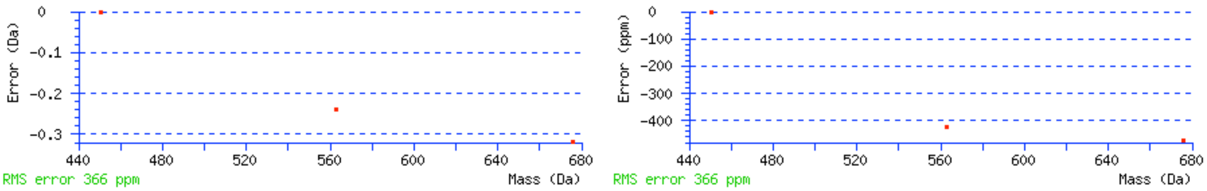


111. Mrpl17 (2 peptides in 1000mM fraction 2)

MS/MS Fragmentation of **KVNLMLK**
Found in **P36528** in **Uniprot_yeastS288C**, 54S ribosomal protein L17, mitochondrial OS=Saccharomyces cerevisiae (strain ATCC 204508 / S288c) OX=559292 GN=MRPL17 PE=1 SV=2

Match to Query 3065: 861.554722 from(431.784637,2+) intensity(168960.1562) rtinseconds(1082.8375) index(2638)
Title: 20100530_methyl1000mM_2.2941.2941.2
Data file D:\Panyue\Proteomic dataset\20100530_methyl1000mM_2.RAW

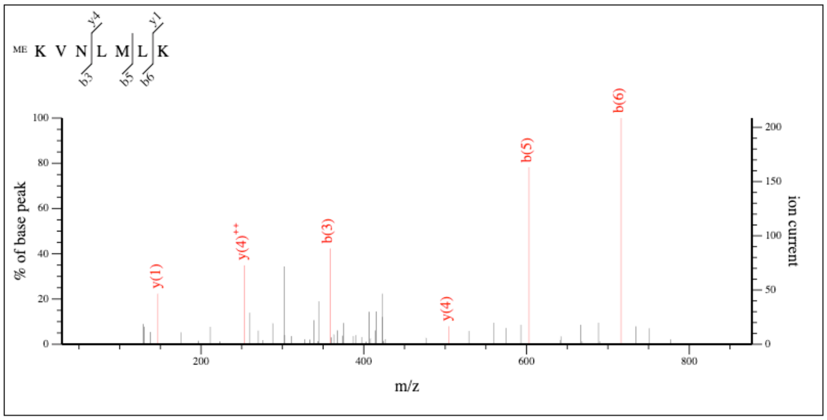


**Monoisotopic mass of neutral peptide Mr(calc):** 861.5549
**Fixed modifications:** Carbamidomethyl (C) (apply to specified residues or termini only)
**Variable modifications:**
**N-term :** Methyl:2H(3) (Protein N-term)
**Ions Score:** 23 **Expect:** 0.016
**Matches :** 8/48 fragment ions using 7 most intense peaks (help)

| # | b | b++ | b* | b*++ | Seq. | y | y++ | y* | y*++ | # |
|---|---|---|---|---|---|---|---|---|---|---|
| 1 | 146.1367 | 73.5720 | 129.1102 | 65.0587 | K |  |  |  |  | 7 |
| 2 | 245.2051 | 123.1062 | 228.1786 | 114.5929 | V | 717.4328 | **359.2200** | 700.4062 | 350.7067 | 6 |
| 3 | ***359.2481*** | 180.1277 | 342.2215 | 171.6144 | N | 618.3643 | 309.6858 | 601.3378 | 301.1725 | 5 |
| 4 | 472.3321 | 236.6697 | 455.3056 | 228.1564 | L | **504.3214** | **252.6643** | 487.2949 | 244.1511 | 4 |
| 5 | ***603.3726*** | 302.1899 | 586.3461 | 293.6767 | M | 391.2374 | 196.1223 | 374.2108 | 187.6090 | 3 |
| 6 | ***716.4567*** | 358.7320 | 699.4301 | 350.2187 | L | 260.1969 | 130.6021 | 243.1703 | 122.0888 | 2 |
| 7 |  |  |  |  | K | **147.1128** | 74.0600 | 130.0863 | 65.5468 | 1 |

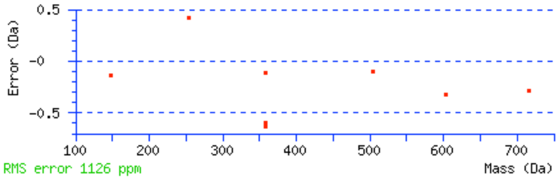


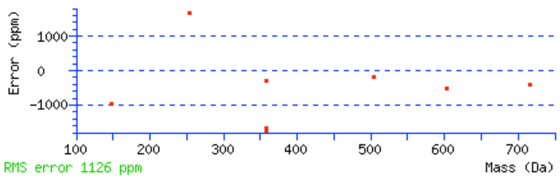

MS/MS Fragmentation of **KVNLMLK**
Found in **P36528** in **Uniprot_yeastS288C**, 54S ribosomal protein L17, mitochondrial OS=Saccharomyces cerevisiae (strain ATCC 204508 / S288c) OX=559292 GN=MRPL17 PE=1 SV=2

Match to Query 3064: 861.554296 from(431.784424,2+) intensity(186087.3281) rtinseconds(1079.6538) index(2629)
Title: 20100530_methyl1000mM_2.2931.2931.2
Data file D:\Panyue\Proteomic dataset\20100530_methyl1000mM_2.RAW

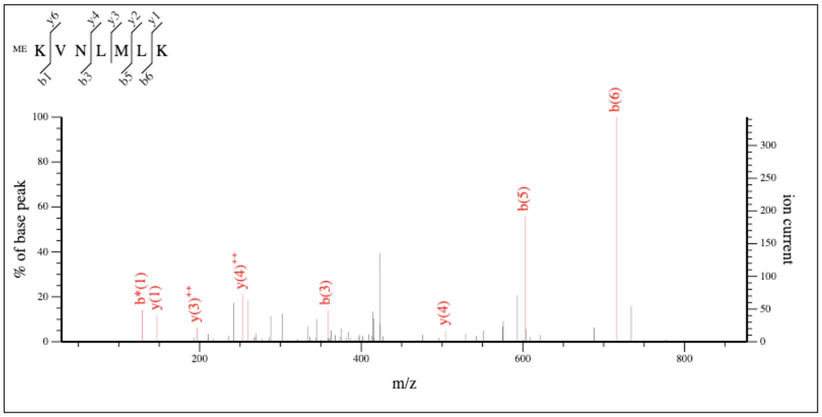


**Monoisotopic mass of neutral peptide Mr(calc):** 861.5549
**Fixed modifications:** Carbamidomethyl (C) (apply to specified residues or termini only)
**Variable modifications:**
**N-term :** Methyl:2H(3) (Protein N-term)
**Ions Score:** 25 **Expect:** 0.012
**Matches :** 12/48 fragment ions using 20 most intense peaks (help)

| # | b | b++ | b* | b*++ | Seq. | y | y++ | y* | y*++ | # |
|---|---|---|---|---|---|---|---|---|---|---|
| 1 | 146.1367 | 73.5720 | 129.1102 | 65.0587 | K | | | | | 7 |
| 2 | 245.2051 | 123.1062 | 228.1786 | 114.5929 | V | *717.4328* | *359.2200* | 700.4062 | 350.7067 | 6 |
| 3 | **359.2481** | 180.1277 | 342.2215 | 171.6144 | N | 618.3643 | 309.6858 | 601.3378 | 301.1725 | 5 |
| 4 | 472.3321 | 236.6697 | 455.3056 | 228.1564 | L | *504.3214* | *252.6643* | 487.2949 | 244.1511 | 4 |
| 5 | **603.3726** | 302.1899 | 586.3461 | 293.6767 | M | 391.2374 | *196.1223* | 374.2108 | 187.6090 | 3 |
| 6 | **716.4567** | 358.7320 | 699.4301 | 350.2187 | L | *260.1969* | 130.6021 | 243.1703 | 122.0888 | 2 |
| 7 | | | | | K | *147.1128* | 74.0600 | 130.0863 | 65.5468 | 1 |

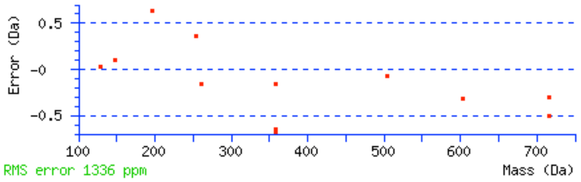


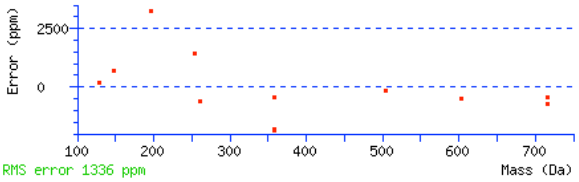


112. Kes1 (1 peptide in 1000mM fraction 2)

MS/MS Fragmentation of **SQYASSSSWTSFLK**
Found in **P35844** in **Uniprot_yeastS288C**, Oxysterol-binding protein homolog 4 OS=Saccharomyces cerevisiae (strain ATCC 204508 / S288c) OX=559292 GN=KES1 PE=1 SV=1

Match to Query 8853: 1594.845448 from(798.430000,2+) intensity(102156.6641) rtinseconds(1155.3997) index(2821)
Title: 20100530_methyl1000mM_2.3144.3144.
Data file D:\Panyue\Proteomic dataset\20100530_methyl1000mM_2.RAW

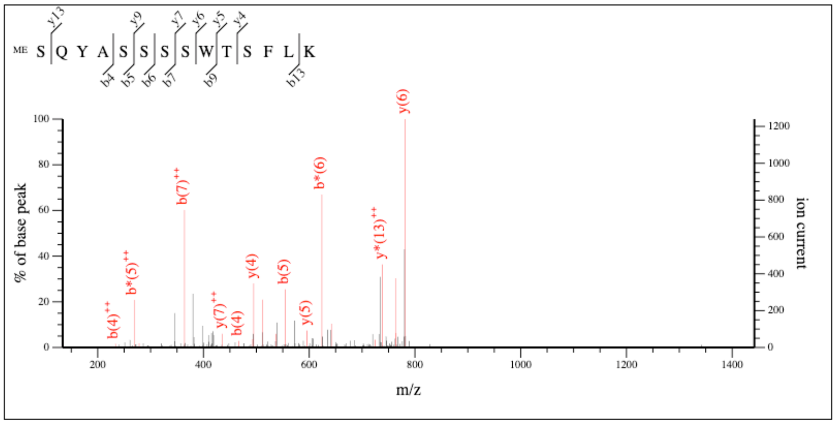


**Monoisotopic mass of neutral peptide Mr(calc):** 1594.7707
**Fixed modifications:** Carbamidomethyl (C) (apply to specified residues or termini only)
**Variable modifications:**
**N-term :** Methyl:2H(3) (Protein N-term)
**Ions Score:** 23 **Expect:** 0.074
**Matches :** 24/148 fragment ions using 45 most intense peaks (help)

| # | b | b++ | b* | b*++ | b0 | b0++ | Seq. | y | y++ | y* | y*++ | y0 | y0++ | # |
|---|---|---|---|---|---|---|---|---|---|---|---|---|---|---|
| 1 | 105.0738 | 53.0405 | | | 87.0632 | 44.0352 | S | | | | | | | 14 |
| 2 | ***233.1324*** | 117.0698 | 216.1058 | 108.5565 | 215.1218 | 108.0645 | Q | 1491.7114 | 746.3594 | 1474.6849 | 737.8461 | 1473.7009 | 737.3541 | 13 |
| 3 | 396.1957 | 198.6015 | 379.1691 | 190.0882 | 378.1851 | 189.5962 | Y | 1363.6529 | 682.3301 | 1346.6263 | 673.8168 | 1345.6423 | 673.3248 | 12 |
| 4 | ***467.2328*** | ***234.1200*** | 450.2063 | 225.6068 | 449.2222 | 225.1148 | A | 1200.5895 | 600.7984 | 1183.5630 | 592.2851 | 1182.5790 | 591.7931 | 11 |
| 5 | ***554.2648*** | ***277.6361*** | 537.2383 | 269.1228 | 536.2543 | 268.6308 | S | 1129.5524 | 565.2798 | 1112.5259 | 556.7666 | 1111.5418 | 556.2746 | 10 |
| 6 | ***641.2969*** | 321.1521 | 624.2703 | 312.6388 | 623.2863 | 312.1468 | S | 1042.5204 | 521.7638 | 1025.4938 | 513.2506 | 1024.5098 | 512.7585 | 9 |
| 7 | 728.3289 | ***364.6681*** | 711.3023 | 356.1548 | 710.3183 | 355.6628 | S | 955.4884 | 478.2478 | 938.4618 | 469.7345 | 937.4778 | 469.2425 | 8 |
| 8 | 815.3609 | 408.1841 | 798.3344 | 399.6708 | 797.3504 | 399.1788 | S | 868.4563 | 434.7318 | 851.4298 | 426.2185 | 850.4458 | 425.7265 | 7 |
| 9 | 1001.4402 | 501.2238 | 984.4137 | 492.7105 | 983.4297 | 492.2185 | W | ***781.4243*** | 391.2158 | 764.3978 | 382.7025 | 763.4137 | 382.2105 | 6 |
| 10 | 1102.4879 | 551.7476 | 1085.4614 | 543.2343 | 1084.4773 | 542.7423 | T | ***595.3450*** | 298.1761 | 578.3184 | 289.6629 | 577.3344 | 289.1709 | 5 |
| 11 | 1189.5199 | ***595.2636*** | 1172.4934 | 586.7503 | 1171.5094 | 586.2583 | S | ***494.2973*** | 247.6523 | 477.2708 | 239.1390 | 476.2867 | 238.6470 | 4 |
| 12 | 1336.5884 | 668.7978 | 1319.5618 | 660.2845 | 1318.5778 | 659.7925 | F | 407.2653 | 204.1363 | 390.2387 | 195.6230 | | | 3 |
| 13 | 1449.6724 | ***725.3398*** | 1432.6459 | 716.8266 | 1431.6618 | 716.3346 | L | 260.1969 | 130.6021 | 243.1703 | 122.0888 | | | 2 |
| 14 | | | | | | | K | 147.1128 | 74.0600 | 130.0863 | 65.5468 | | | 1 |

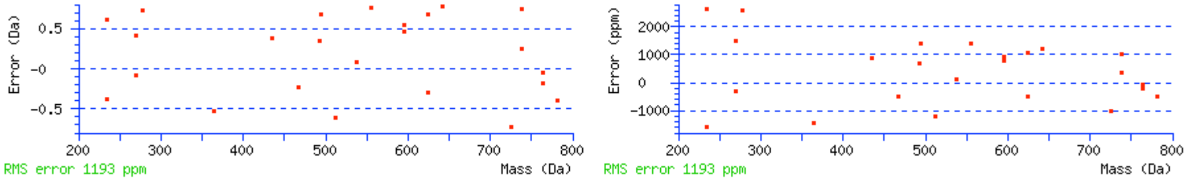


113. Hem13 (1 peptide in 1000mM fraction 2)

MS/MS Fragmentation of **PAPQDPR**
Found in **P11353** in **Uniprot_yeastS288C**, Oxygen-dependent coproporphyrinogen-III oxidase OS=Saccharomyces cerevisiae (strain ATCC 204508 / S288c) OX=559292 GN=HEM13 PE=1 SV=2

Match to Query 2388: 796.464221 from(797.471497,1+) intensity(125777.7812) rtinseconds(1319.6937) index(3240)
Title: 20100530_methyl1000mM_2.3611.3611.1
Data file D:\Panyue\Proteomic dataset\20100530_methyl1000mM_2.RAW

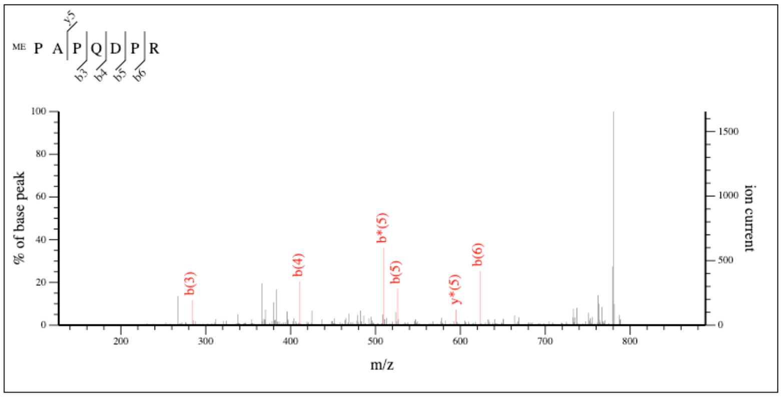


**Monoisotopic mass of neutral peptide Mr(calc):** 796.4271
**Fixed modifications:** Carbamidomethyl (C) (apply to specified residues or termini only)
**Variable modifications:**
**N-term :** Methyl:2H(3) (Protein N-term)
**Ions Score:** 25 **Expect:** 0.03
**Matches :** 6/27 fragment ions using 12 most intense peaks (help)

| # | b | b* | b⁰ | Seq. | y | y* | y⁰ | # |
|---|---|---|---|---|---|---|---|---|
| 1 | 115.0945 | | | P | | | | 7 |
| 2 | 186.1316 | | | A | 683.3471 | 666.3206 | 665.3365 | 6 |
| 3 | ***283.1844*** | | | P | 612.3100 | 595.2835 | 594.2994 | 5 |
| 4 | ***411.2430*** | 394.2164 | | Q | 515.2572 | 498.2307 | 497.2467 | 4 |
| 5 | ***526.2699*** | 509.2434 | 508.2594 | D | 387.1987 | 370.1721 | 369.1881 | 3 |
| 6 | ***623.3227*** | 606.2961 | 605.3121 | P | 272.1717 | 255.1452 | | 2 |
| 7 | | | | R | 175.1190 | 158.0924 | | 1 |

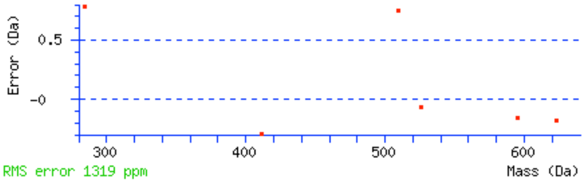


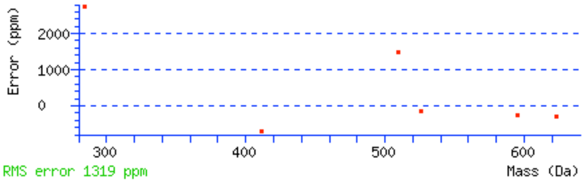


### 114. RSOLAG22IIIB_01053 (1 peptide in 1000mM fraction 3)

MS/MS Fragmentation of **SQATAPTERPGMFR**
Found in **A0A0K6G1L8** in **Uniprot_yeastS288C**, Carboxylic acid transporter protein homolog [Saccharomyces cerevisiae S288c] OS=Rhizoctonia solani OX=456999 GN=RSOLAG22IIIB_01053 PE=4 SV=1

Match to Query 8219: 1614.878758 from(808.446655,2+) intensity(232930.4844) rtinseconds(2879.7727) index(6908)
Title: 20100530_methyl1000mM_3.7699.7699.2
Data file D:\Panyue\Proteomic dataset\20100530_methyl1000mM_3.RAW

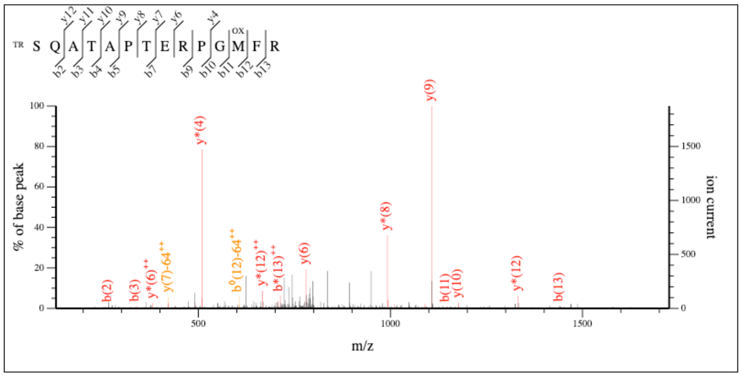

**Monoisotopic mass of neutral peptide Mr(calc):** 1614.8498
**Fixed modifications:** Carbamidomethyl (C) (apply to specified residues or termini only)
**Variable modifications:**
**N-term :** Trimethyl:2H(9) (Protein N-term)
**M12** : Oxidation (M), with neutral losses 0.0000(shown in table), 63.9983
**Ions Score:** 29 **Expect:** 0.0075
**Matches :** 32/212 fragment ions using 96 most intense peaks (help)

| # | b | $b^{++}$ | $b^{*}$ | $b^{*++}$ | $b^{0}$ | $b^{0++}$ | Seq. | y | $y^{++}$ | $y^{*}$ | $y^{*++}$ | $y^{0}$ | $y^{0++}$ | # |
|---|---|---|---|---|---|---|---|---|---|---|---|---|---|---|
| 1 | 139.1427 | 70.0750 | | | 121.1322 | 61.0697 | S | | | | | | | 14 |
| 2 | 267.2013 | 134.1043 | 250.1748 | 125.5910 | 249.1908 | 125.0990 | Q | 1477.7216 | 739.3645 | 1460.6951 | 730.8512 | 1459.7111 | 730.3592 | 13 |
| 3 | 338.2384 | 169.6229 | 321.2119 | 161.1096 | 320.2279 | 160.6176 | A | 1349.6630 | 675.3352 | 1332.6365 | 666.8219 | 1331.6525 | 666.3299 | 12 |
| 4 | 439.2861 | 220.1467 | 422.2596 | 211.6334 | 421.2756 | 211.1414 | T | 1278.6259 | 639.8166 | 1261.5994 | 631.3033 | 1260.6154 | 630.8113 | 11 |
| 5 | 510.3232 | 255.6653 | 493.2967 | 247.1520 | 492.3127 | 246.6600 | A | 1177.5783 | 589.2928 | 1160.5517 | 580.7795 | 1159.5677 | 580.2875 | 10 |
| 6 | 607.3760 | 304.1916 | 590.3494 | 295.6784 | 589.3654 | 295.1864 | P | 1106.5411 | 553.7742 | 1089.5146 | 545.2609 | 1088.5306 | 544.7689 | 9 |
| 7 | 708.4237 | 354.7155 | 691.3971 | 346.2022 | 690.4131 | 345.7102 | T | 1009.4884 | 505.2478 | 992.4618 | 496.7346 | 991.4778 | 496.2425 | 8 |
| 8 | 837.4663 | 419.2368 | 820.4397 | 410.7235 | 819.4557 | 410.2315 | E | 908.4407 | 454.7240 | 891.4141 | 446.2107 | 890.4301 | 445.7187 | 7 |
| 9 | 993.5674 | 497.2873 | 976.5408 | 488.7741 | 975.5568 | 488.2820 | R | 779.3981 | 390.2027 | 762.3716 | 381.6894 | | | 6 |
| 10 | 1090.6201 | 545.8137 | 1073.5936 | 537.3004 | 1072.6096 | 536.8084 | P | 623.2970 | 312.1521 | 606.2704 | 303.6389 | | | 5 |
| 11 | 1147.6416 | 574.3244 | 1130.6151 | 565.8112 | 1129.6310 | 565.3192 | G | 526.2442 | 263.6258 | 509.2177 | 255.1125 | | | 4 |
| 12 | 1294.6770 | 647.8421 | 1277.6505 | 639.3289 | 1276.6664 | 638.8369 | M | 469.2228 | 235.1150 | 452.1962 | 226.6017 | | | 3 |
| 13 | 1441.7454 | 721.3763 | 1424.7189 | 712.8631 | 1423.7349 | 712.3711 | F | 322.1874 | 161.5973 | 305.1608 | 153.0840 | | | 2 |
| 14 | | | | | | | R | 175.1190 | 88.0631 | 158.0924 | 79.5498 | | | 1 |

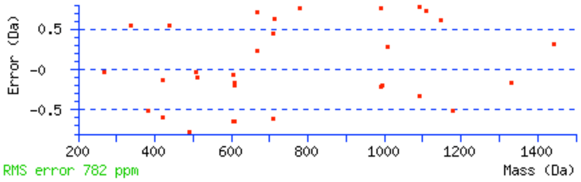


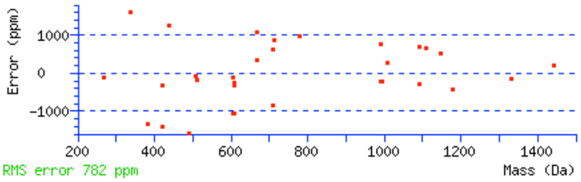
